# MORT, a locus for apoptosis in the human immunodeficiency virus-type 1 antisense gene: implications for AIDS, Cancer, and Covid-19

**DOI:** 10.1101/2020.07.01.180828

**Authors:** Linda B. Ludwig, Michael S. Albert

## Abstract

Apoptosis, or programmed cell death, is a fundamental requirement for life in multicellular organisms, including humans, and a mechanism to maintain homeostasis and prevent unwarranted cellular proliferations such as cancer. An antisense gene in HIV-1 (*Hap*) induces apoptosis in human cells. Apoptotic T cell death following HIV-1 infection leads to a compromised immune system and eventually AIDS (acquired immunodeficiency syndrome). A review of several studies that focused on long-term survivors of HIV-1 reveals that these survivors had deletion-mutations in *Hap*. A subset of these survivors changed course and experienced CD4+ T cell death and progression to AIDS. These individuals had virus that regained *Hap* gene sequence that had previously been deleted. Analysis of the changes in the genetic sequences with *in vivo* progression of the revertant HIV-1 virus allowed identification of a specific region in *Hap* we are calling MORT. MORT, in *Hap* RNA forms a primary microRNA-like structure. Potential human mRNAs targeted by MORT mi/siRNAs include gene/RNA sequences of X-linked inhibitor of apoptosis (XIAP), survivin, and apollon, along with many other human gene sites/RNAs. Thus MORT may be acting as an RNA antagonist to cellular IAPs thereby inducing apoptotic cell death. Surprisingly, additional potential MORT targets include viral sites in human SARS-CoV-2, including the protease, nsp5 RNA. Future uses for RNA therapy and a hypothesis for an HIV intrinsic mechanism utilizing MORT for viral anti-viral (or anti-microbial) and HIV anti-immune cell defense are proposed.

## Introduction

To die or not to die: that is the question. The acquired immunodeficiency syndrome (AIDS) caused by the human immunodeficiency virus type 1 (HIV-1) has long been known to be associated with the death of the CD4+ T cell (Kelleher and Zaunders, 2006)). While some controversy has been associated with the mechanism of the demise of this central cell pivotal in both B and T cell immunity, laboratory observations clearly reveal the depletion of thymocyte and CD4^+^ T cell populations as HIV-1 infection progresses, except in rare individuals (Barmania and Pepper, 2013; Liu, et al, 1996; Dean, et al, 1996; Samson, et al, 1996; Carrington, et al, 1999). If one puts aside the human host side of the pathogenesis equation, such as those individuals lacking a competent co-receptor (the CCR5 deletion) (Barmania and Pepper, 2013; Liu, et al, 1996; Dean, et al, 1996; Samson, et al, 1996; Carrington, et al, 1999) on the immune cells to facilitate HIV entry who thereby acquire some resistance to infection by the virus: what about the virus itself induces cell death? Is there a genetic component of HIV-1 that has reliably been shown to be associated with T cell death?

The human T cell intrinsically faces the decision for division, anergy, or apoptosis at multiple stages, both in developmental processes and in ramping up (and down) a response to infection (Kolb, et al, 2017; Kerr, et al, 1972; Clarke, et al, 1993; Schwartz and Osborne, 1993; Cohen, 1993). When it comes time to die-what signal or signals must occur within the cell? More particularly, is there a signal for cell death contributed by human immunodeficiency virus?

Here we propose that a specific region of the *Hap* antisense gene located within the HIV-1 long terminal repeat (LTR) is contributing to the programmed cell death or apoptosis observed with HIV infection (Ludwig, et al, 2006; Ludwig, 2008). *Hap* RNA initiates in the repeat (R) region of the HIV-1 provirus and is transcribed in the reverse direction (antisense) to the other HIV genes, overlapping the beginning portion of TAR region DNA, the TATA box, the SP1 sites and NFKb sites controlling HIV-1 sense transcription, as well as 200 nucleotides of the 3’ end of the nef gene (Figure 1B) (Ludwig, et al, 2006; Ludwig, 2008). The introduction of this antisense gene into human cells induced programmed cell death or apoptosis, as classically defined (Kerr, et al, 1972; Clarke, et al, 1993; Schwartz and Osborne, 1993; Cohen, 1993). In most experiments, performed over a decade between 1995 (the discovery of the HIV antisense initiator (HIVaINR) (U.S. patent 5,919,677) and 2005, it was a race to isolate the RNA or proteins(s) from human cells before apoptosis occurred.

**Figure 1.**
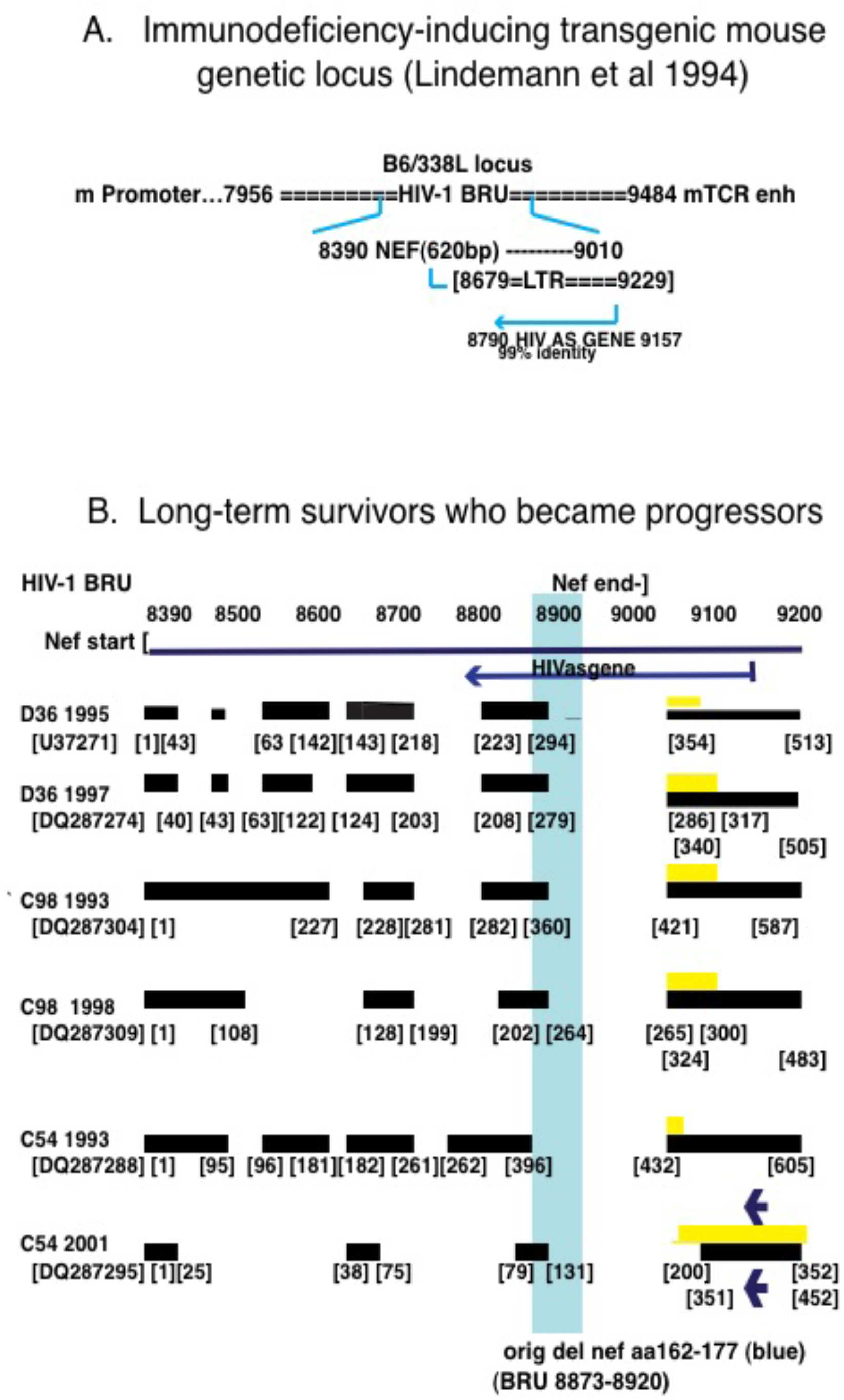
Indications that a specific HIV-1 genetic locus contributes to cell death. A. Lindemann, et al, 1994 showed that the HIV-1 BRU nef and long terminal repeat (LTR) genetic sequences linked to a mouse T cell receptor (TCR) beta chain promoter and mouse TCR beta chain core enhancer were sufficient to induce a phenotype of severe combined immunodeficiency and T cell loss in mice incorporating the transgene. Remarkably, their transgene construct rearranged in vivo, such that the B6/338L transgenic mouse locus had the mouse TCR enhancer <u>downstream</u> of the mouse TCR promoter linked to the HIV-1 BRU sequences. This situated the region of the HIV antisense gene (HIVasgene) within the LTR close to the enhancer elements (which can function in either direction). The HIV-1 BRU sequence (Genbank accession number K02013) (Wain-Hobson, et al, 1985) used in their construct is 99% identical to the corresponding region of the HIVas gene used in constructing the HAP-FLAG vector, previously described (Genbank accession number CQ767337) (Ludwig, et al, 2006; Ludwig, 2008). B. The HIV-1 nef and LTR sequences obtained from virus isolated from peripheral blood mononuclear cells from three long-term survivors who became progressors was aligned using BLAST and visual inspection (Blastn suite) on the NCBI website (National Center for Biotechnology Information), U.S. National Library of Medicine) with sequences from HIV-1 BRU (Genbank K02013) (Wain-Hobson, et al, 1985) and the HIV antisense gene sequence 27 (from U.S. patents 7700725, 7700726, and 8765461, and European patent 1359221 (Genbank CQ767337)) (Ludwig, et al, 2006; Ludwig, 2008). Black blocks represent intact sequence, gaps represent deletions, and grey blocks show altered/duplicated enhancer regions. The Sydney Blood Bank Cohort (SBBC) individuals who became progressors are shown, including the original blood donor who was infected in 1980 (D36) and blood recipients C98 and C54, who received blood transfusions in 1982 and 1984, respectively (Gorry, et al, 2007). The HIV-1 nef/LTR sequence data is from GenBank accession numbers: D36 1995 (U37271, sequence 1-513 (Deacon, et al, 1995)), D36 1997 (DQ287274, sequence 1-505 (Churchill, et al, 2006)), C98 1993 (DQ287304, sequence 1-587 (Churchill, et al, 2006)), C98 1998 (DQ287309, sequence 1-483 (Churchill, et al, 2006)), C54 1993 (DQ287288, sequence 1-605 (Churchill, et al, 2006)), and C54 2001 (DQ287295, sequence 1-452 (Churchill, et al, 2006). With C54 2001 between sequence 200-452 there is complete duplication of the HIV-1 enhancer/promoter region, including the HIV antisense initiator (two reverse arrows). The original common SBBC deletion of nef aa 162-177, corresponding to BRU sequence 8873-8920 is indicated throughout by the blue filled rectangle (Greenway, et al, 1998). The alignment is not to scale.

Because of the enhanced apoptotic cell death observed with introduction of a recombinant eukaryotic vector incorporating the *Hap* gene sequence (GenBank CQ767337 nt 14-367), we undertook a reanalysis of observations in two “in vivo” systems: a) a mouse model of AIDS (Lindemann, et al, 1994), with a nef/LTR transgene containing near-identical sequence to that of the *Hap* gene, and b) comparison of the genetic changes in the nef/LTR deleted viral species isolated in a group of HIV-1-infected individuals who changed from long term non-progressor (LTNP) status to progression to AIDS, so-called “slow progressors” (Gorry, et al, 2007; Deacon, et al, 1995; Churchill et al, 2004; Churchill, et al, 2006; Dyer, et al, 1999; Dyer, et al 1997; Greenway, et al 1998; Rhodes, et al, 1999; Learmont, et al 1992; Oelrichs, et al 1998; Zaunders, et al 2011; Learmont, et al, 1999).

The slow progressors within the Sydney Blood Bank Cohort (SBBC) remarkably demonstrated both the “loss of function” as well as “gain of function” required for T cell death, in a natural human system prior to the institution of anti-retroviral therapy (Gorry, et al, 2007; Deacon, et al, 1995; Churchill et al, 2004; Churchill, et al, 2006; Dyer, et al, 1999; Dyer, et al 1997; Greenway, et al 1998; Rhodes, et al, 1999; Learmont, et al 1992; Oelrichs, et al 1998; Zaunders, et al 2011; Learmont, et al, 1999). The original viral deletion within these slow progressors (SP) encompassed a portion of the HIV antisense gene that was critical in the formation of an RNA primary microRNA-like secondary/tertiary structure that we are calling MORT.

The HIV antisense gene sequence (GenBank CQ767337) from 270-300 shares 14 to 23 nucleotides of sequence homology with sites in or bordering 177 human genes, significant identity with seven human microRNAs (identified by BLAST on MirGeneDB), and a proposed mi/siRNA forms complementary base-pairing structures with multiple human genes important in the apoptosis pathway. X-linked inhibitor of apoptosis (XIAP), survivin, and Apollon are amongst some of the inhibitor of apoptosis proteins potentially targeted by MORT. This suggests a mechanism whereby the HIV-1 virus can induce apoptosis within the human host cell, with MORT as an antagonist to IAPs, and has potential implications for AIDS latency and applications for cancer. In yet another twist, the HIV antisense gene shares 7 sites with 11-14 nt sequence identity or complementarity with the novel coronavirus, SARS-CoV 2; implications of this will also be discussed.

## Results

### A transgenic mouse model of AIDS

Lindemann, et al, 1994 described a transgenic mouse model of AIDS in which the transgene incorporated only the intact HIV-1 nef and long terminal repeat (nef/LTR) genomic region (BRU, Genbank accession number K02013), and thereby also the HIV antisense gene (*Hap*), under the control of mouse T-cell receptor beta chain-specific promoter and enhancer elements (Figure 1 A). Dramatic changes in the mouse lymphocyte populations were induced in the transgenic mice along with a severe immunodeficiency (Lindemann, et al, 1994). Transgenic mice demonstrating the most significant and severe changes in lymphocyte populations (B6/338L) had a single aberrant copy of the transgene integrated but also rearranged such that the mouse T-cell receptor (TCR) beta chain enhancer was downstream, instead of upstream of the mouse TCR promoter and HIV-1 nef/3’long terminal repeat sequences (Figure 1A), (Lindemann, et al, 1994). This rearrangement situated the enhancer close to the region of the HIV antisense gene in the LTR and in position to help drive antisense transcription off the intrinsic HIV-1 antisense initiator (HIVaINR) in TAR region DNA (Figure S1C) (Ludwig, et al, 2006; Ludwig, 2008). The transgene construct contained HIV-1 BRU genome sequences from 7956 through 9484 and thus contained the intact HIV antisense gene in its entirety. Blast comparison (https://blast.ncbi.nim.nih.gov/Blast.cgi) of the BRU sequence in the mouse transgene with the HIV antisense gene (GenBank CQ767337.1) showed that the corresponding genetic regions were 99% identical (Figure 1A) (Ludwig, et al, 2006; Ludwig, 2008).

They were unable to demonstrate Nef protein production in this transgenic mouse line, thereby suggesting the possibility some other genetic component caused the phenotypic changes in these mice (Lindemann, et al, 1994). Mouse CD4+CD8+ and CD4+CD8-thymocytes were depleted early in ontogeny and offspring of line B6/338L were characterized by a profound immunodeficiency and high mortality rate (Lindemann, et al, 1994). A 200-300 fold decrease in total CD4+ thymocyte numbers in heterozygous B6/338L mice as compared to non-transgenic control littermates was described (Lindemann, et al, 1994). Thus CD4+ T-cell death was occurring as a consequence of the introduction of the transgene containing the nef/LTR and also HIV antisense gene sequences.

### The HIV antisense gene could induce apoptosis

Our experience showed that vectors producing the HIV antisense gene RNA from the intrinsic HIV antisense initiator (HIVaINR) sequences induced apoptosis within three days, but with an alternate cytomegalovirus (CMV) promoter driving antisense RNA transcription, one had even less time before apoptosis occurred (Figure 2A, see also Figure S1A) (Ludwig, et al, 2006; Ludwig, 2008). Characterization of the Hap RNA and protein(s) has previously been described (Ludwig, et al, 2006). Transfection of human cells with HIV-1 antisense gene sequences (CQ767337 bp 14-367) directionally cloned into a recombinant vector with a CMV promoter and a carboxyterminal FLAG (DYKDDDDK) epitope-encoding sequence, called HAP-FLAG, induced apoptosis within 24-36 hours, or even sooner in the absence of Tat (Figure 2), (Ludwig, et al, 2006). While the HeLa (Maddon, et al, 1986), HL2/3 (Felber and Pavlakis, 1988; Ciminale, et al, 1990), and Clone69T1-RevEnv (Yu, et al, 1996) cells transfected with control p15cxCAT (Durand, et al, 1988) and pCMVTag vectors demonstrated occasional mitotic figures, the same cells transfected with the HAP-FLAG vector containing the HIV antisense gene sequences demonstrated enhanced numbers of apoptotic cells (Figure 2A, HAP-FLAG, top row).

**Figure 2.**
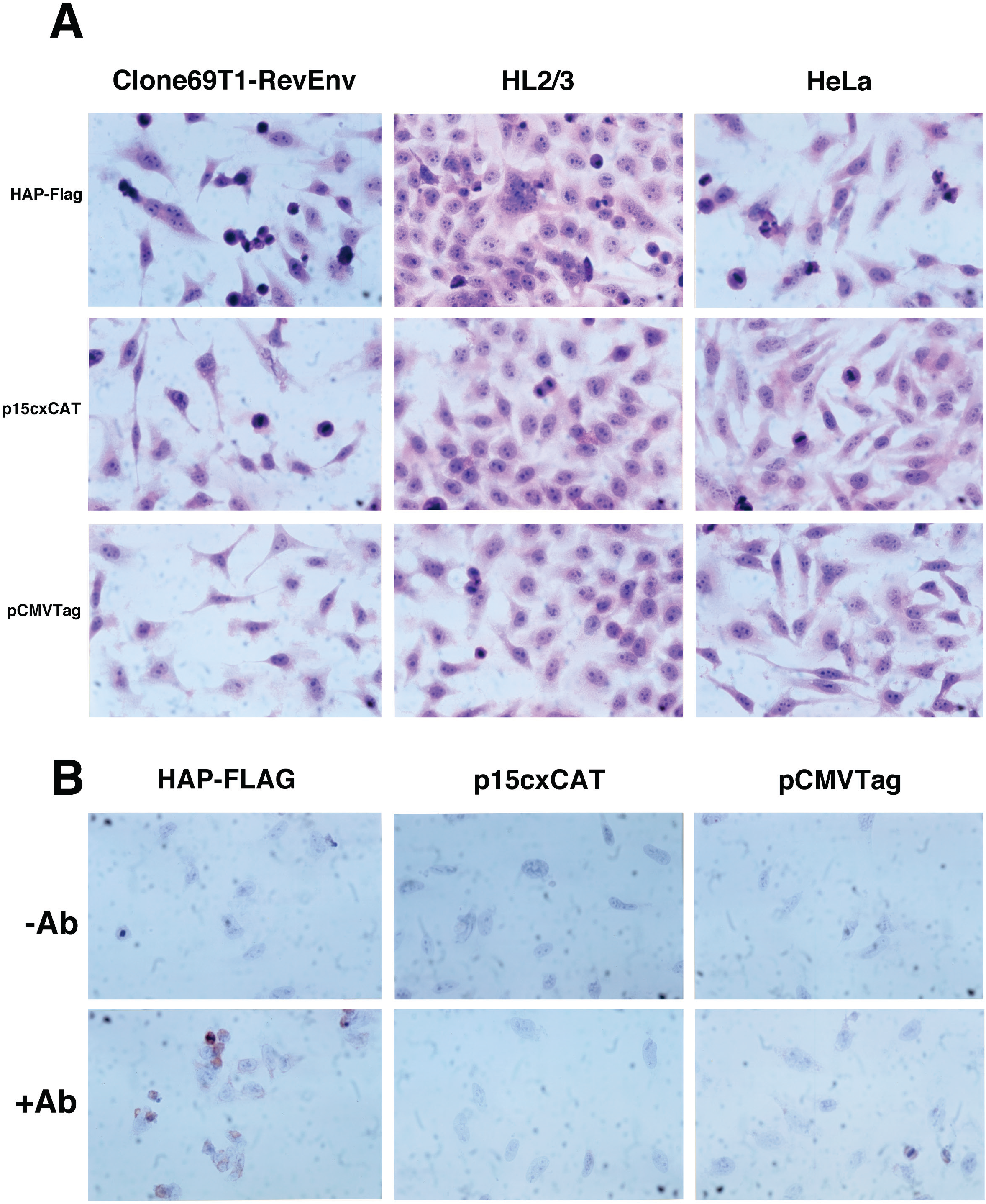
HIV antisense gene sequences incorporated into a recombinant vector produce RNA and protein(s) and induce apoptosis. (See also supplemental Figure S1.) A. Recombinant HIV-1 antisense gene sequences linked to a carboxy-terminal FLAG sequence (DYKDDDDK) (HAP-FLAG (Ludwig, et al, 2006)) were transfected into three separate cell lines and increased cell death (apoptosis) over cells transfected with control gene sequences (magnification 400X). Identical starting numbers (5 × 10^5^) of HL2/3 cells (Felber and Pavlakis, 1988; Ciminale, et al, 1990), HeLa cells (Maddon, et al, 1986) and Clone69T1RevEnv cells (induced to express Rev and Env by the withdrawal of tetracycline (Yu, et al, 1996)) were cultured in 6-well plates with cover slips, stabilized overnight, and then transfected with either the recombinant HIV-1 antisense gene-FLAG vector (HAP-FLAG) (Ludwig, et al, 2006), the interleukin 2 enhancer-CAT vector (Durand, et al, 1988) or the control luciferase-FLAG vector (Stratagene), as indicated. The cells were incubated 18 hours, then the coverslips with cells removed, fixed in 95% alcohol, placed on slides and stained with hematoxylin and eosin (H & E). A surgical pathologist, who was unaware of the experimental conditions, independently assessed slides and counted apoptotic cells per high power field (hpf). Apoptotic cells/hpf were as follows, SEM for 3 counts: For HeLa (HAP-FLAG: 4.3 +/−0.6; IL-2 –CAT: 1.7+/−0.6: luc-FLAG: 1.7+/−1.5); for Clone 69T1RevEnv (HAP-FLAG: 10.3 +/−4.5; IL-2-CAT 1.3 +/−0.6; luc-FLAG 1.3 +/− 1.5); for HL2/3 cells (HAP-FLAG 8.3 +/− 2.5; IL-2 CAT 3+/−2.6; luc-FLAG: 4.7 +/− 2.9). Mitotic figures can be observed (see middle slide, HL2/3 transfected with p15cxCAT control). B. Duplicate sets of Hela cells (as above) were transfected with HAP-FLAG, or control vectors p15cxCAT or pCMVTag, (as indicated) under identical experimental conditions. Following (early) removal of coverslips from 6-well plates, and fixing as above, they were either incubated with monoclonal antibody to FLAG (+ Ab) or no Ab (− Ab), followed by immunoperoxidase staining for HAP-FLAG detection (Ludwig, et al, 2006).

The HeLa cell line was also transfected with the three separate vectors (HAP-FLAG, p15cxCAT, and pCMVTag) under identical experimental conditions, and then analyzed for protein expression using a monoclonal anti-FLAG antibody (+Ab) or control (-Ab) (Figure 2B). In addition to demonstrating the perinuclear location of the HAP-FLAG protein (Figure 2B, HAP-FLAG, +Ab), in this experiment the well coverslips had to be pulled early (within hours) because under light microscopy the cells were already “balling up” and floating off the coverslips (lbl, observation).

Figure 2 shows that apoptosis was enhanced by the HIV antisense gene regardless of whether the human cell line already produced the HIV-1 Rev and Env proteins (Clone69T1-RevEnv) (Yu, et al, 1996), all the HIV-1 proteins except reverse transcriptase (HL2/3) (Felber and Pavlakis, 1988; Ciminale, et al, 1990) or was the control cell line, HeLa. This evidence strongly suggests a contributing agent for the apoptotic cell death (from the HIV-1 genome) is localized to the HIV antisense gene in the long terminal repeat of HIV (Ludwig, et al, 2006; Ludwig, 2008). These experiments also demonstrate that all three human HeLa cell lines, which are long-lived because they are immortalized cancer cell lines, can be induced to undergo apoptosis with the introduction of a vector expressing the HIV antisense gene.

### Discovery of MORT in the HIV antisense gene

A vast literature exists involving research on human long term survivors (LTS) infected with attenuated nef/LTR –deleted HIV-1 strains (Gorry, et al, 2007; Deacon, et al, 1995; Churchill et al, 2004; Churchill, et al, 2006; Kirchhoff, et al, 1995; Mariani, et al, 1996; Dyer, et al, 1999; Dyer, et al 1997; Salvi, et al 1998; Greenway, et al 1998; Rhodes, et al, 1999; Learmont, et al 1992; Oelrichs, et al 1998; Zaunders, et al 2011; Learmont, et al, 1999). Because the intact nef/LTR region of the HIV genome, which also contains the HIV antisense gene, appeared to be sufficient for inducing cytopathogenicity and an AIDS phenotype in mice (Lindemann, et al, 1994), we reasoned that the virus isolated from human long-term survivors might provide further clues as to the viral genetic regions important for cell death. The separate issues of <u>host</u> genetic changes, such as mutations/deletions in CCR5, a chemokine receptor on the host CD4 + T cell important in viral entry, or viral defects affecting viral entry or infectious capability are also important, but are not the subject of this paper.

The Sydney blood bank cohort (SBBC) included 8 individuals (C98, C54, C49, C64, C18, C135, C83 and C124) infected with an attenuated, nef/LTR deleted HIV-1 strain by blood products from a single HIV-1 infected donor (D36) (Gorry, et al, 2007; Deacon, et al, 1995; Churchill et al, 2004; Churchill, et al, 2006; Dyer, et al, 1999; Dyer, et al 1997; Greenway, et al 1998; Rhodes, et al, 1999; Learmont, et al 1992; Oelrichs, et al 1998; Zaunders, et al 2011; Learmont, et al, 1999). The original deletion in the HIV-1 nef/LTR causing attenuation of the virus was shown to be in a region of nef-long terminal repeat overlap (Figure 1B, blue rectangle)(Greenway, et al 1998). Early studies demonstrated the absence of antibody reactivity to a single peptide in an array of peptides corresponding to aa 162-177 of Nef protein (Figure 1B) (Greenway, et al 1998). They also reported failure to amplify wild-type HIV-1 sequences with PCR primers in the conserved deletion (Learmont, et al, 1999). The “accident of nature” represented by this human cohort allowed study of the natural course of viral infection and host immune responses, as well as the alterations in the genomic sequence of HIV-1 over time. Despite infection with the same D36 virus, the infected recipients had a varied clinical course (Gorry, et al, 2007; Deacon, et al, 1995; Churchill et al, 2004; Churchill, et al, 2006; Dyer, et al, 1999; Dyer, et al 1997; Greenway, et al 1998; Rhodes, et al, 1999; Learmont, et al 1992; Oelrichs, et al 1998; Zaunders, et al 2011; Learmont, et al, 1999). Some cohort members had stable CD4+T-cell counts and low plasma HIV RNA levels for up to 29 years without antiretroviral therapy (C49, C64, and C135) and were called elite non-progressors (Gorry, et al, 2007, Zaunders, et al 2011).

However, there were three slow progressors (SP) (D36, C54, and C98) that experienced a profoundly different clinical course, with a decline in CD4+T cells following many years of asymptomatic infections (Gorry, et al, 2007). Full-length proviral sequence analysis showed the nef-LTR mutations were the only unusual HIV-1 feature in these infected individuals (Oelrichs, et al 1998). The original HIV-1 deletion of the SP donor and recipients completely overlapped the HIV antisense gene sequence from nucleotides (nt) 237-284 (Figures 1B (blue rectangle), 3, 4 (red arrows)) (Ludwig, et al, 2006; Ludwig, 2008).

#### Slow progressor viruses regain a portion of HIV antisense gene sequence while losing nef sequence

Specific sequence comparisons of these SBBC SP HIV-1 genomic sequences deposited in GenBank, and cloned prior to any anti-retroviral therapy, was undertaken. Sequences were aligned with HIV-1 BRU (LAV-1) from the start of Nef at 8390 through the repeat (R) region sequence at 9193 (Figure 1B) (Gorry, et al, 2007; Deacon, et al, 1995; Churchill et al, 2004; Churchill, et al, 2006, Oelrichs, et al 1998, Zaunders, et al 2011). This included the HIV antisense initiator and start site at BRU 9156 (Figure 1B, as diagrammed) (Ludwig, et al, 2006; Ludwig, 2008). The original deletion of Nef aa 162-177 corresponding to HIV-1 BRU sequence 8873-8920 and HIV antisense gene sequence 284-237 (overlapping in the reverse direction) was shared by D36 with the SBBC blood product recipients (blue shaded rectangle, Figure 1B) (Greenway, et al 1998). The D36 viral genome changed over time to regain some of the deleted sequence (D36, Figures 1B, 3, 4) (Gorry, et al, 2007; Deacon, et al, 1995; Churchill et al, 2004; Churchill, et al, 2006, Oelrichs, et al 1998, Zaunders, et al 2011). This regaining of sequence in the original deletion (blue rectangle) can also be observed in the other SP C98 and C54 viruses (Figure 1B, Figure 3A). Comparison of C54 viral genomic alterations between 1993 and 2001 (9 and 17 years post transfusion) most clearly illustrates the gain of sequence in the region of the original SBBC deletion while more of the nef sequence is lost (C54, Figure 1B). The most significant decline in CD4+ T cells appeared 12-13 years post-transfusion, and co-incident with the gain of sequence in the original deletion region (Learmont, et al, 1999). Nef deletions increased in all three individuals who went from stable CD4^+^ T cell counts to SP status (Gorry, et al, 2007, Churchill et al, 2004, Churchill, et al, 2006), making it difficult to postulate that Nef was responsible for CD4+ T cell loss and disease progression in these individuals. It is interesting also to note that C54 HIV-1 viral species acquired a complete duplication of the entire enhancer and promoter region and thereby also duplicated the HIVaINR with the potential for two antisense RNA initiation sites (Figure 1B, C54 2001, reverse arrows: duplicated enhancer regions indicated). In our experiments, only a single intrinsic start site from the HIVaINR was observed (Figure S1C, Ludwig, et al, 2006).

**Figure 3.**
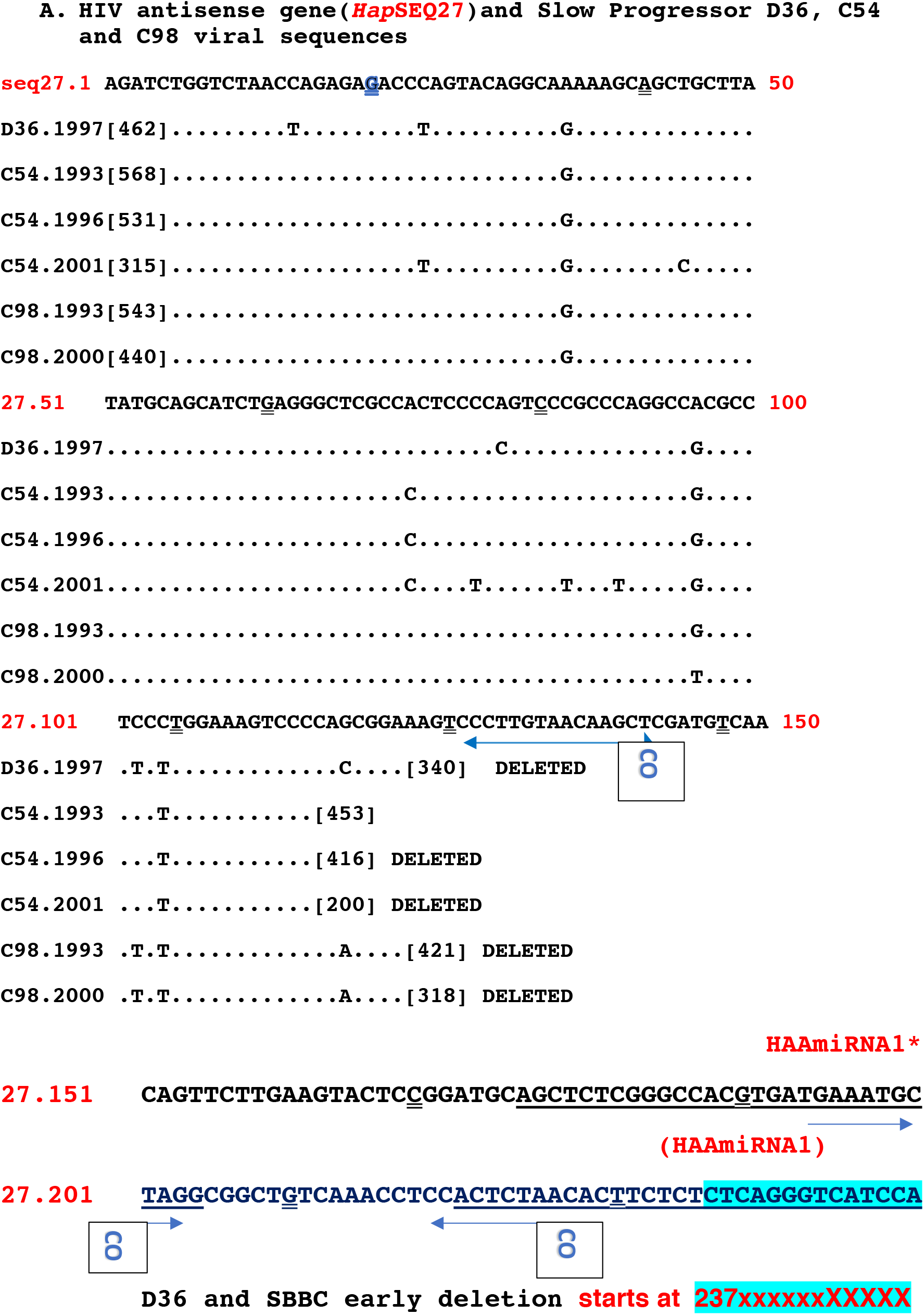
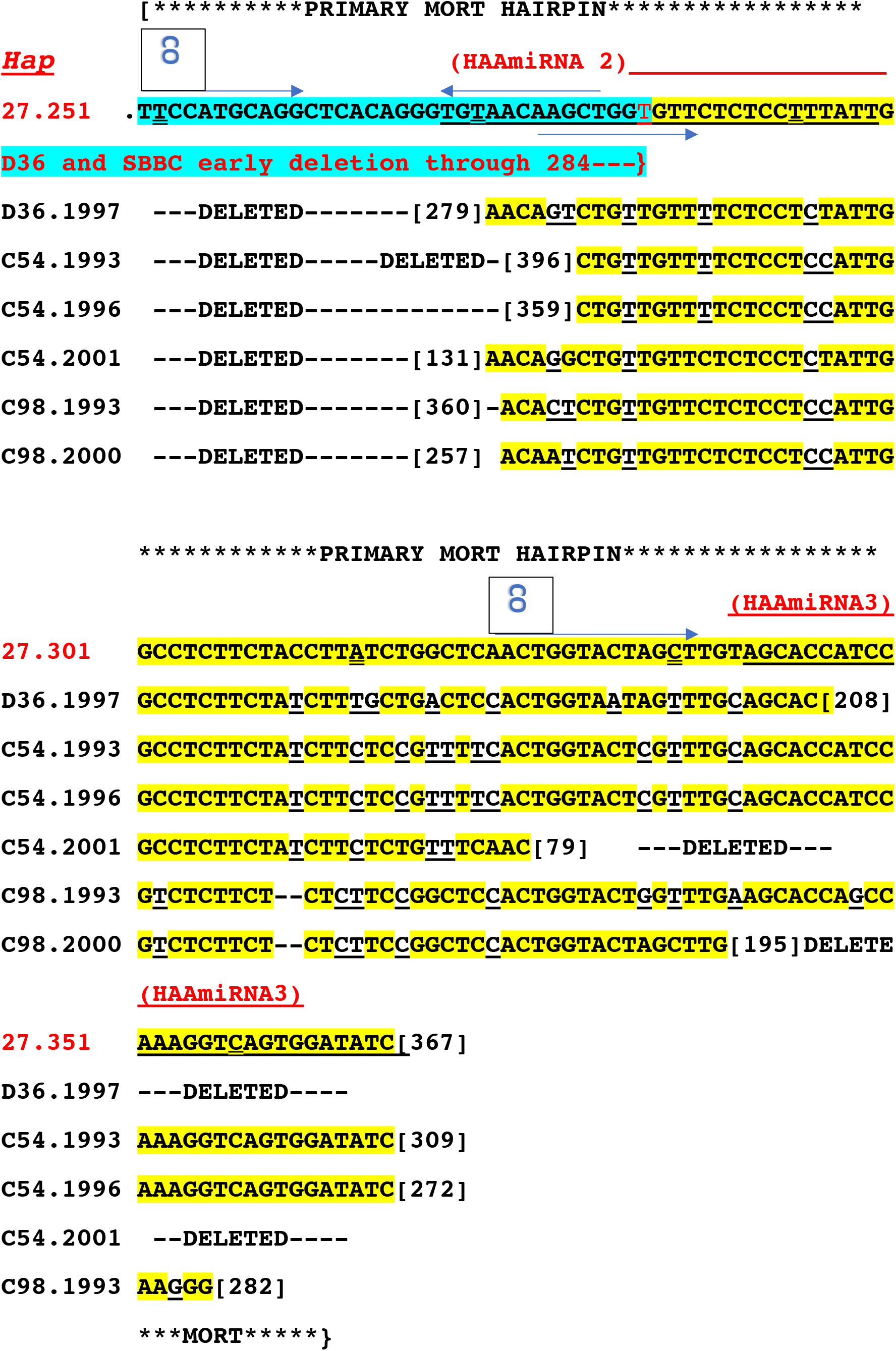
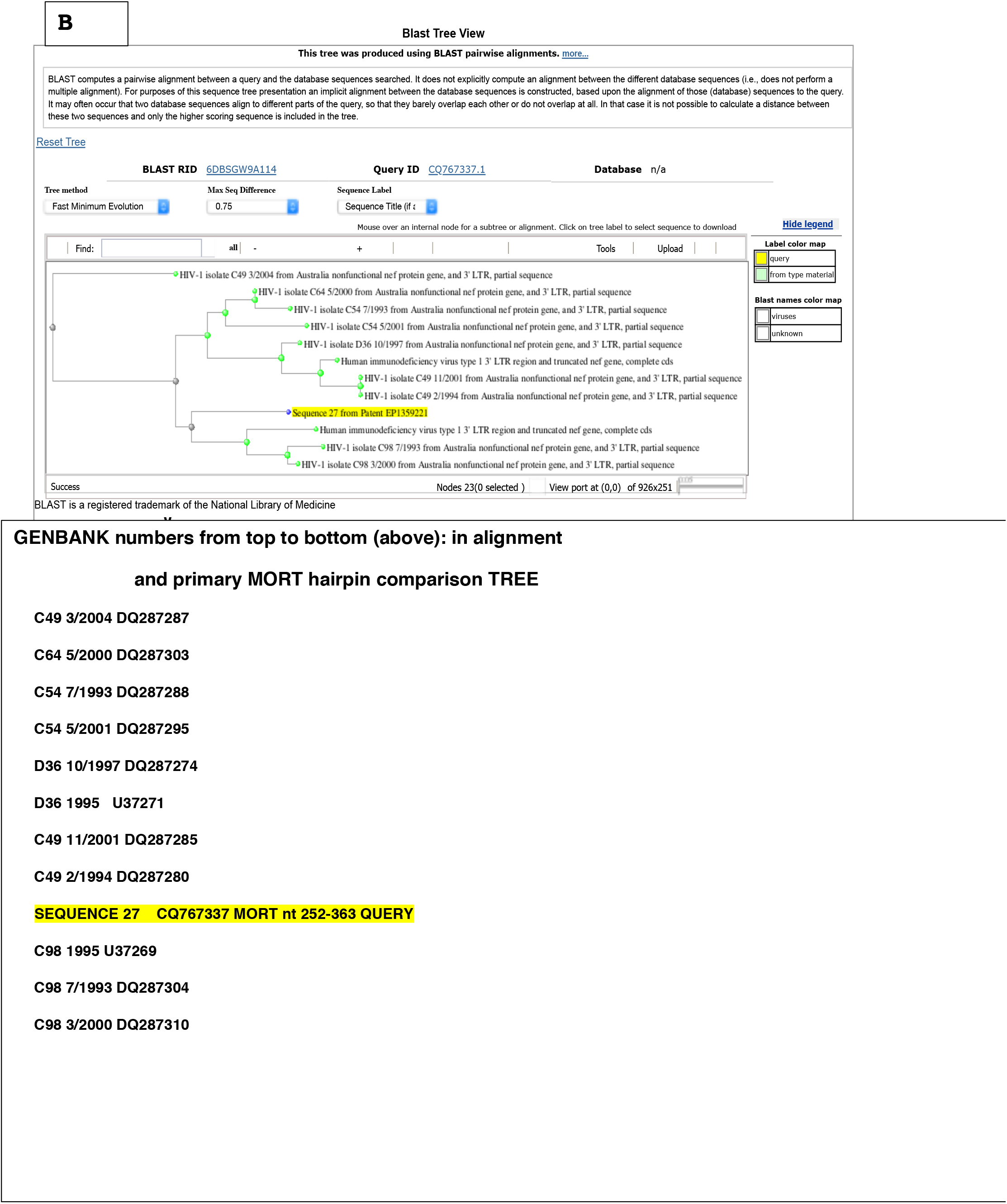
**A.** Comparison of the slow progressor HIV-1 sequences over time reveals sequence is regained in a specific locus of the HIV antisense gene seq 27 (CQ767337) nt 274-284 (a genetic deletion originally shared by all the SBBC donor and recipient viruses) while nef sequence is progressively lost. The D36 and shared SBBC early deletion is indicated in red type (blue shading) (X). The GenBank accession numbers are as indicated in Figure 1B, with the addition of C98.2000: GenBank accession DQ287310 and C54.1996: GenBank accession DQ287291 (Churchill, et al, 2006); GenBank specific nucleotide (nt) sequence for each individual SP is indicated in brackets. Sequences are shown from the beginning (HIVaINR start site) to the end of HIV antisense gene, with the exception of the duplicated enhancer regions (diagrammed in gray blocks in Figure 1B). Previously proposed HAAmiRNAs 1, 2, and 3 are underlined (Ludwig 2008). The region corresponding to the primary MORT RNA hairpin structure is indicated (***). Potential Dicer cuts (double-underlined) of dsRNA are indicated at 21 nt intervals (Ludwig 2008). Sites identical (forward blue arrow) or complementary to (backward blue arrow) SARS-CoV-2 RNA (CO) are indicated and were obtained by NCBI nBLAST comparison between CQ767337 and MT007544.1 (Wuhan seafood market pneumonia virus isolate Australia/VIC01/2020). Seven sites with identities between 11/11 to 14/15 nt were identified and are shown with arrows. **B.** MORT phylogeny of slow progressor (D36, C98, and C54) and elite non-progressor (C49, C64) SBBC sequences compared to MORT sequence 252-363 following BLASTn alignment (NCBI). GenBank accession numbers for each sequence is indicated below tree.

A study of separate HIV-1 viruses isolated from D36, either in 1995 (prior to progression to AIDS) or from 1999 (after disease progression) found that the 1999 isolate induced greater levels of CD4+ T cell cytotoxicity than the 1995 isolate in an ex vivo human lymphoid cell culture system (Jekle, et al, 2002). This illustrates the enhanced cell death that occurred coincident with evolution of the D36 viral genome. This is also consistent with our observations of apoptosis induced by the introduction of the intact HIV antisense gene into human cells (Figure 2 and data not shown).

#### Alignment of SBBC slow progressor viral sequences with the HIV antisense gene reveals sequence nt 274-284 (originally deleted) is regained, corresponding to sequence in a small noncoding antisense RNA (snc #208, Althaus, et al, 2012)

Others have analyzed the evolution of the nef/LTR sequences from SBBC viruses (Gorry, et al, 2007; Deacon, et al, 1995; Churchill et al, 2004; Churchill, et al, 2006, Oelrichs, et al 1998, Zaunders, et al, 2011). All of these studies viewed the deletion from the genomic RNA strand (sense) direction, instead of bi-directionally, despite the fact that HIV-1 must replicate using a dsDNA proviral form. Using the gene sequences from these studies (deposited in GenBank), we re-aligned the sequences to the HIV antisense gene located in the proviral LTR using BLAST and visual alignment (Figure 3A, *Hap* sequence 27, at top). Sequence 27 (CQ767337) is identical to that cloned into the recombinant HAP-FLAG vector used to produce RNA and proteins, as previously described in detail (Ludwig, et al, 2006). Figure 3A illustrates the striking alterations in the derived HIV antisense gene sequences of viruses isolated from the donor D36 and two SP recipients, C54 and C98, as compared to the original SBBC deletion. The region of the original SBBC deletion corresponds to HIV antisense gene sequence 237-284 (Figure 3A, in red).

D36, C54, and C98 gained sequence from HIV antisense nucleotides (nt) 274-284, which had previously been deleted. The regained sequence was part of the previously proposed <u>H</u>IV<u>a</u>INR <u>a</u>ntisense RNA <u>HAA</u>miRNA 2 site (underlined, Figure 3A) (Ludwig, 2008). **Particularly exciting was that a 22 nt small noncoding RNA has been cloned and sequenced from HIV-infected cells following selection for antisense small RNAs and corresponds to nt 274-295** (Althaus, et al, 2012). In contrast, the SBBC SP genetic regions overlapping the HIV-1 core promoter and enhancer elements (TATA box, Sp1 and NFkB sites) are predominantly conserved and are 93-96% identical to corresponding HIV-1 BRU sequences (see region corresponding to HIV antisense gene sequence 27.1 through 27.123 (Figure 3A). For simplicity, the enhancer duplication regions of the SBBC SP are not shown in Figure 3A, but are indicated as overlapping rectangles in Figure 1B.

The phylogeny of the SBBC progressor versus nonprogressor sequences as compared to HIV antisense gene MORT primary (miRNA) sequence 252-363 is shown in Figure 3B (NCBI Blast Tree View Widget). The *in vivo* progression of the C98 viruses (below Query *Hap* MORT sequences (in yellow)) can be compared to the elite non-progressor C49 (above Query sequence). C49 sequence did not appear to change in the MORT primary region between 1994 and 2001, and while closer to D36 1995 sequences is the furthest distance from *Hap* MORT (seq 27 from patent EP1359221) (Figure 3B)

Figure 4 illustrates M-fold analysis (Zuker, 2003; Markham and Zuker, 2008) of the HIV antisense gene RNA (CQ767337) used in our experiments. The original deletion of SBBC/D36 viral species is indicated by red line, later deletions for D36 viral species are outlined in yellow, and where D36 virus actually regained sequence is indicated (purple lines) (Figure 4). Over time, between the original infection in 1980 and genetic analysis of the virus in 1995 and 1997, the D36 viral genome underwent significant changes in the HIV antisense gene sequence region 274-345 (corresponding to BRU sequence 8882-8812).

**Figure 4.**
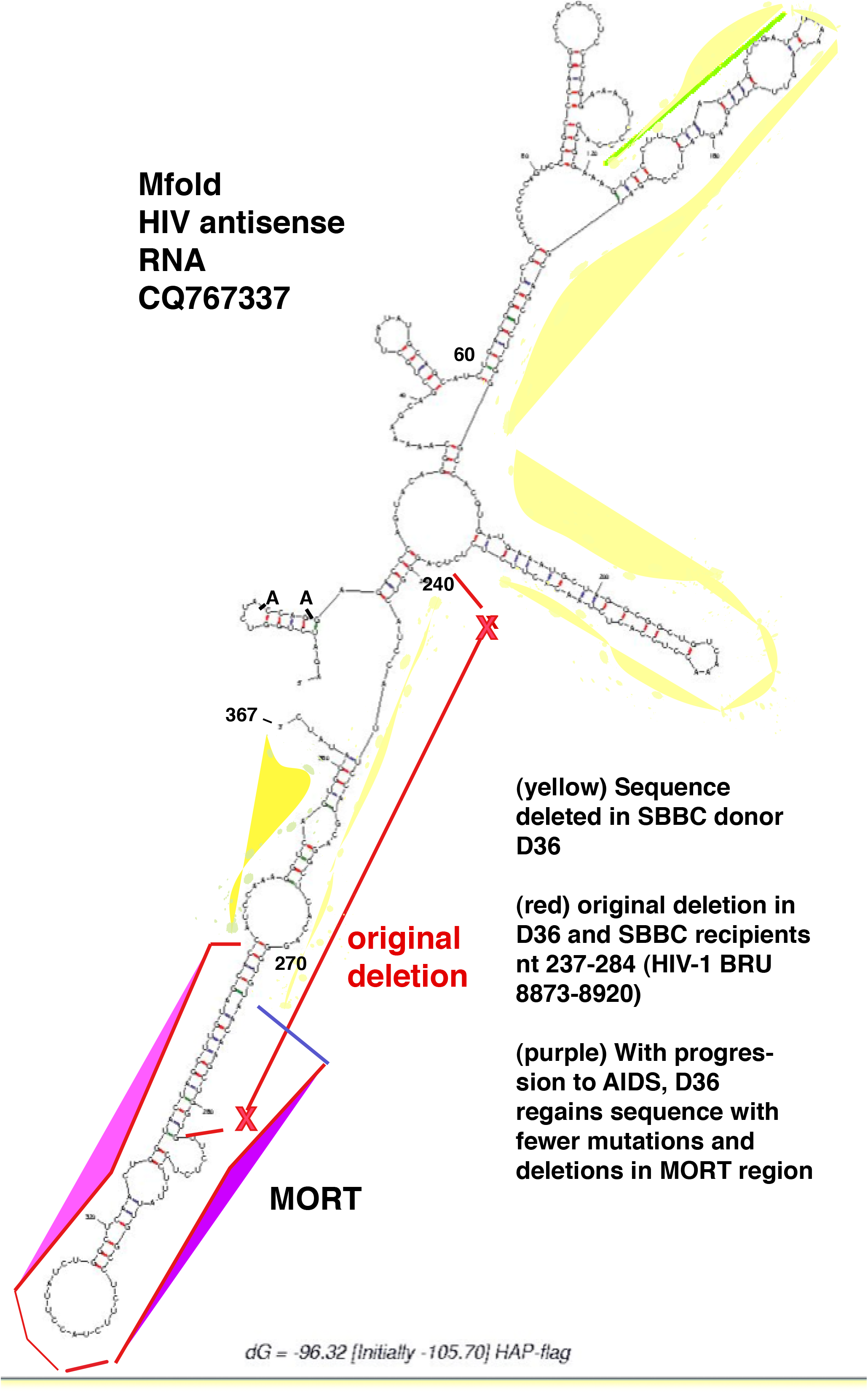
Mfold (Zuker, 2003; Markham and Zuker, 2008) analysis of the HIV antisense gene sequence 27 (CQ767337) reveals a 33 bp hairpin structure from nt 252-363 resembling a primary microRNA structure. The original D36 deletion is indicated in red, with the region of change in later sequence (D36.1997) in purple. We are calling this 33 bp hairpin structure MORT.

The revertant changes in D36, C54 and C98 viral genomes to include the intact HIV antisense gene region 274-284, while maintaining adjacent hairpin-forming sequence up to nt 345 occurs along with alterations of the clinical status in these individuals to progressors. The donor (D36) was infected in 1980, but by 1996-1998 six of seven CD4 lymphocyte counts were 500 or fewer cells per cubic millimeter with the lowest value of 282 in Sept 1998 (Learmont, et al, 1999). He was diagnosed with HIV dementia in 12/1998 and began antiretroviral therapy on 1/1999 (reviewed in Gorry, et al, 2007, Zaunders, et al, 2011). SP C54 and C98 experienced significant decreases in CD4+T cells averaging 58 and 73 cells per cubic millimeter per year (Learmont, et al, 1999) and C98 went on antiretroviral therapy in 11/1999 (Gorry, et al, 2007).

Thus a “regain of function” of CD4+T cell death was observed *in vivo* following the reemergence of a specific genetic locus in the HIV antisense gene, a locus of apoptosis. These genetic changes would clearly impact the formation of a RNA secondary/tertiary structure, a 33 bp hairpin we are calling MORT (Figure 4).

### MORT sequence is shared with human genes

NIH NCBI BLAST server and the GenBank database (http://www.ncbi.nim.nih.gov/GenBank/) was used to search the human genome database against a small segment of the HIV antisense gene, CQ767337 nt 270-300, that formed the 5p side of the 33 bp hairpin structure and included this locus of apoptosis (Figure 4). This search revealed shared homology with a large number of sites adjacent or in human genes (see also Table S1 and Figure S6). A hairpin implies often complementary sequence and thus we also used Blast to search the HIV antisense RNA from nucleotides (nt) 330-345 (3p), against the human gene database (data not shown). HIV *nef* sequences are complementary and in the opposite orientation from the HIV antisense gene nt 147 through 367 but regulatory elements from the antisense gene have never previously been described. Thus it was a surprise to observe more than 177 human genomic sites shared significant homology or identity, with 14/14 to 23/23 nt identical to MORT 270-300 (Table S1, Figure S6).

This suggests that once again the human immunodeficiency virus has borrowed from its human host, and also that the shared function of this bit of viral DNA/RNA may have more universal implications (and applications) than previously suspected.

### How MORT might function

Mfold (Zuker, 2003) structure of the HIV-1 antisense RNA that was utilized in our experiments (Ludwig, et al, 2006; Ludwig, 2008) demonstrates an intact 33 bp hairpin with a 15 nt terminal loop from CQ767337 antisense RNA sequence nt 252-363, consistent with a primary microRNA (miRNA)-like structure (labeled MORT, Figure 4). However, all the early SBBC SP viral RNA would be incapable of forming this structure because of the original deletion in the SBBC viral dsDNA genome corresponding to *Hap* gene nt 237-284 (Figures 3, 4). With biological progression or evolution of the virus *in vivo*, the D36 (and other SP) viruses regained sequence from MORT region 274-284, as well as maintaining contiguous sequence such that a revertant hairpin structure with 19 base-paired nucleotides was possible (Figures 3, 4). This reconstituted sequence (and structure) was in evolution at the same time as other surrounding sequence in the nef and nef/LTR region was lost (Figure 4, compare with Figure 1B, 3A). Corresponding to this evolution of the HIV-1 virus, the donor D36, for example, presented a phenotype of enhanced T cell loss and was diagnosed with AIDS dementia (reviewed in Gorry, et al, 2007, Zaunders, et al, 2011).

If one presumes that at least 16 nt complementarity are required between the 5p and 3p hairpin sequences to form a metazoan small interfering RNA (siRNA) or microRNA, it seems that the revertant viral species would support this requisite with a 5p arm (274-299 (26nt)) and 3p arm (321-343/345 (22nt)) (Fromm, et al, 2015; Elbashir, et al, 2001; Bartel, 2009).

As further confirmation, a unique small noncoding (nc) antisense RNA (corresponding to the MORT sequence from 274-295) has previously been identified by cloning and sequencing of small ncRNAs in HIV-1 infected human cells (Althaus, et al., 2012, Sequence No 208). This suggests potential parameters for both 5p and 3p intrinsic HIV si/miRNAs (Figure 5A, top, underlined).

**Figure 5.**
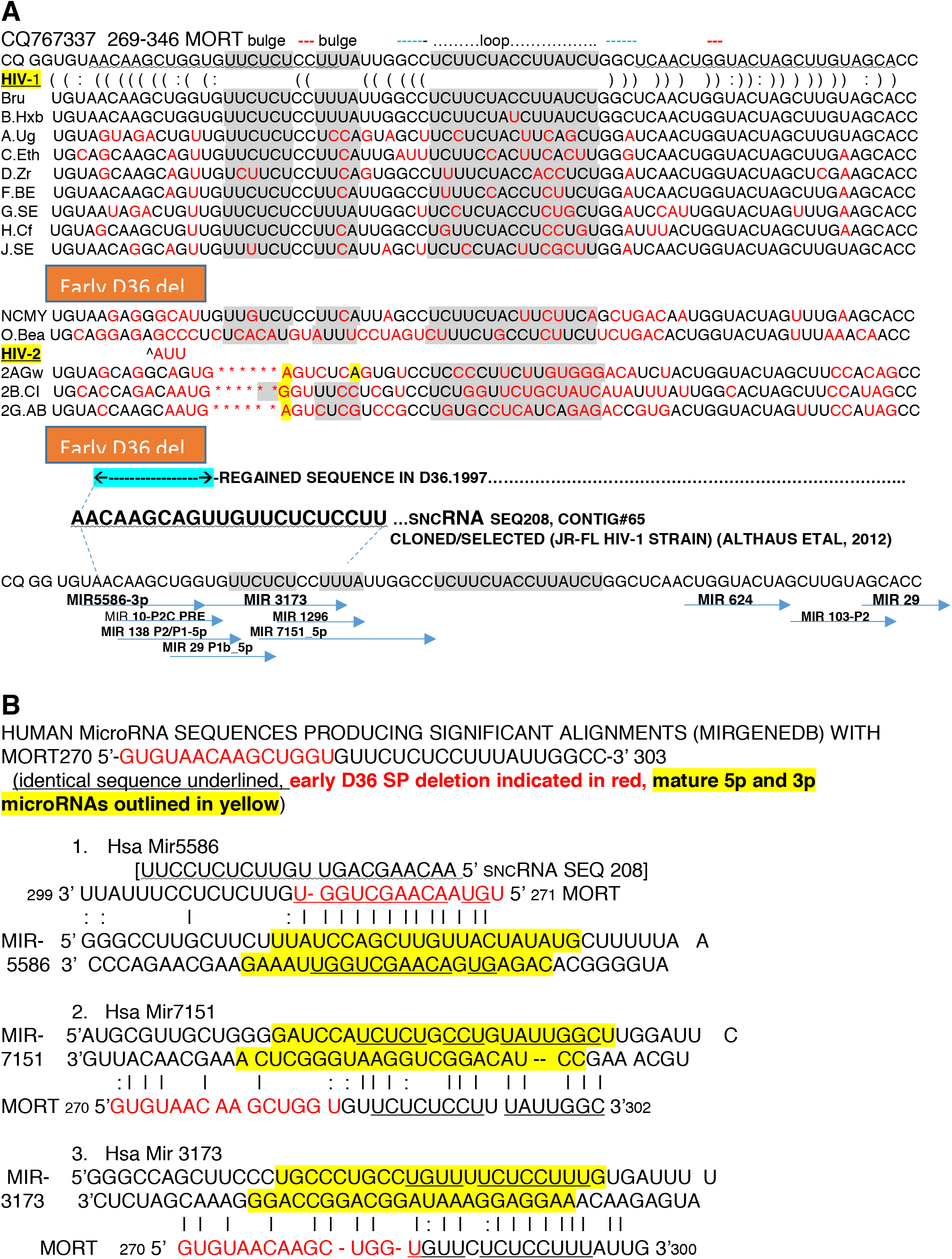
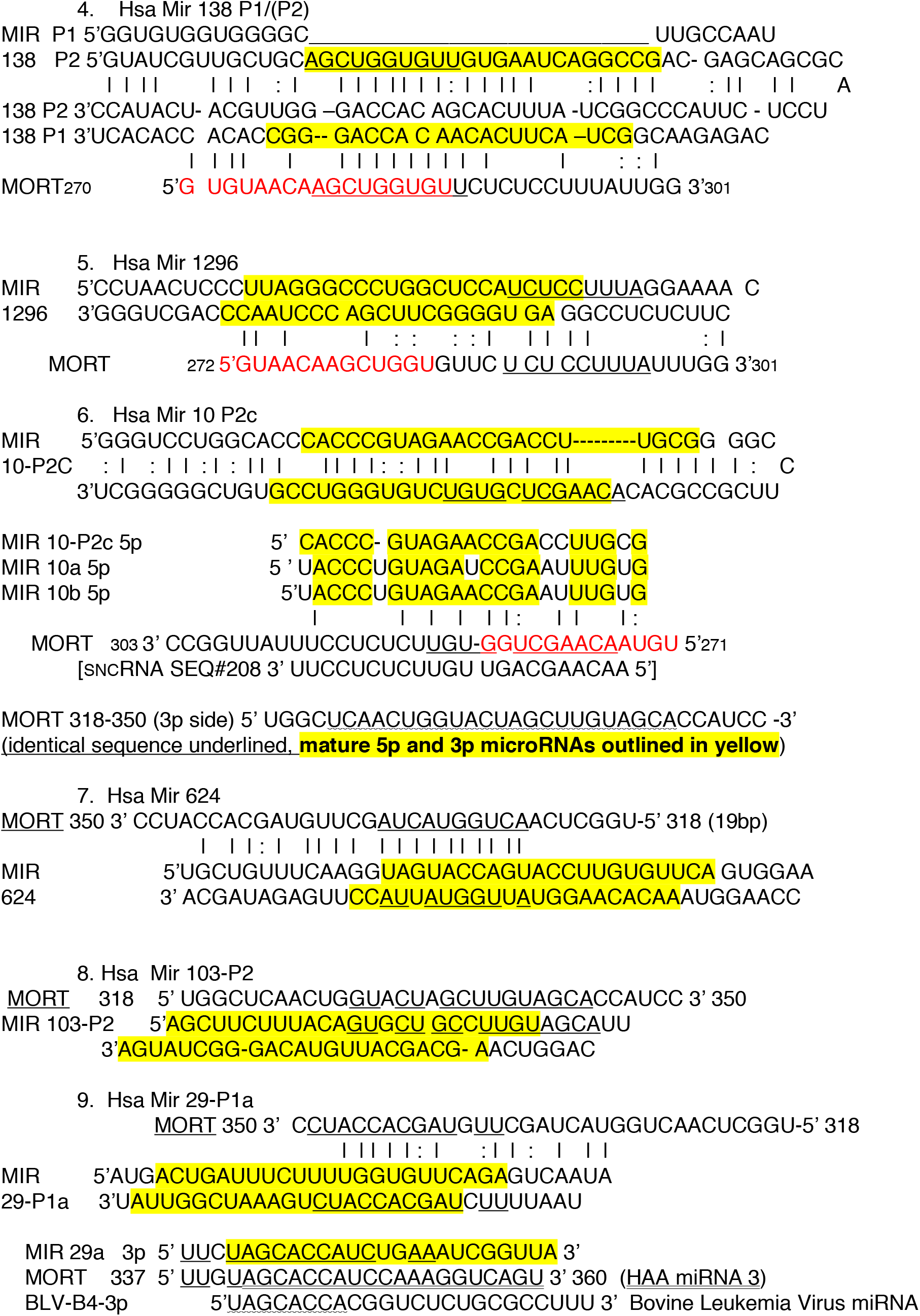
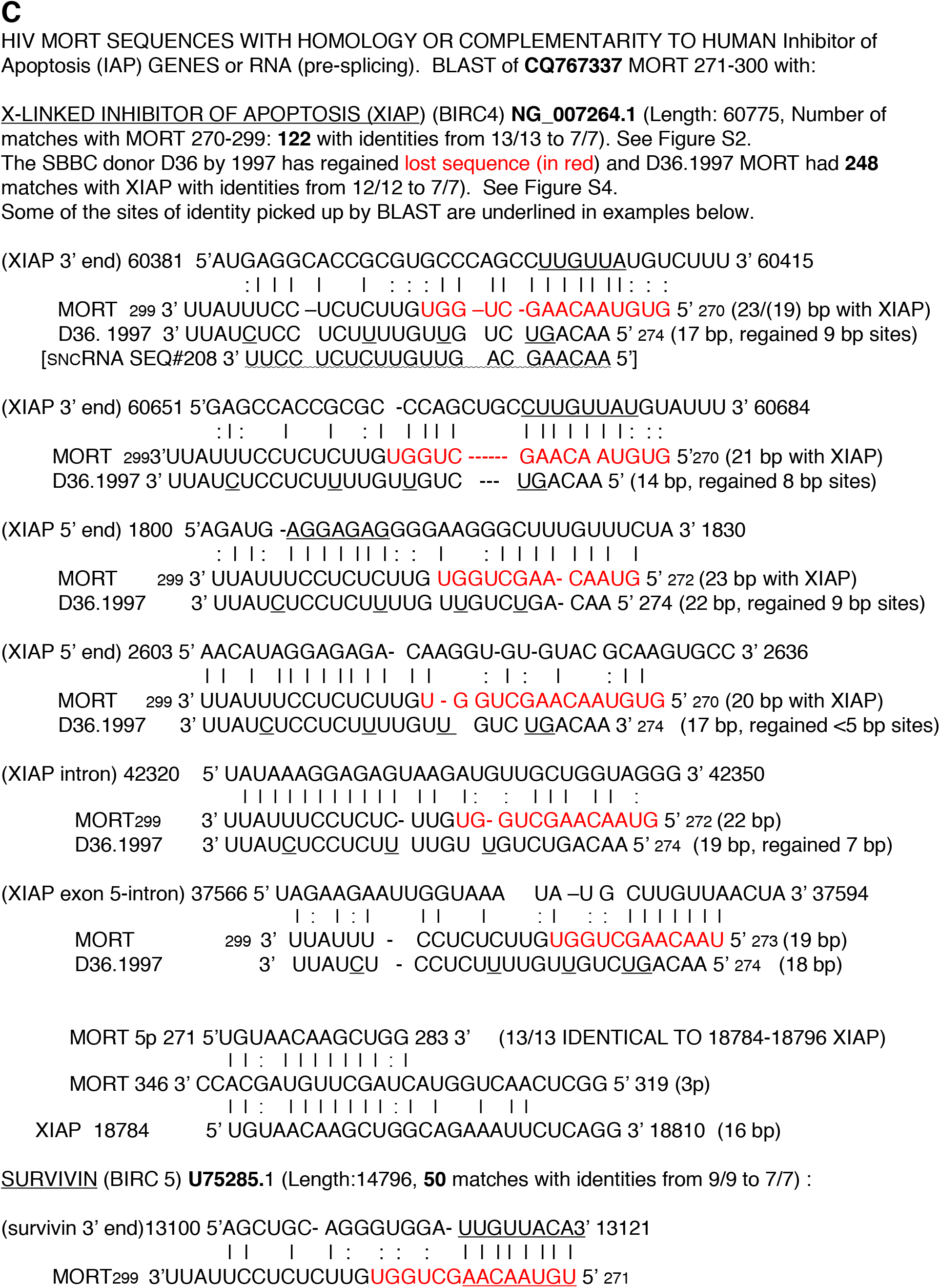
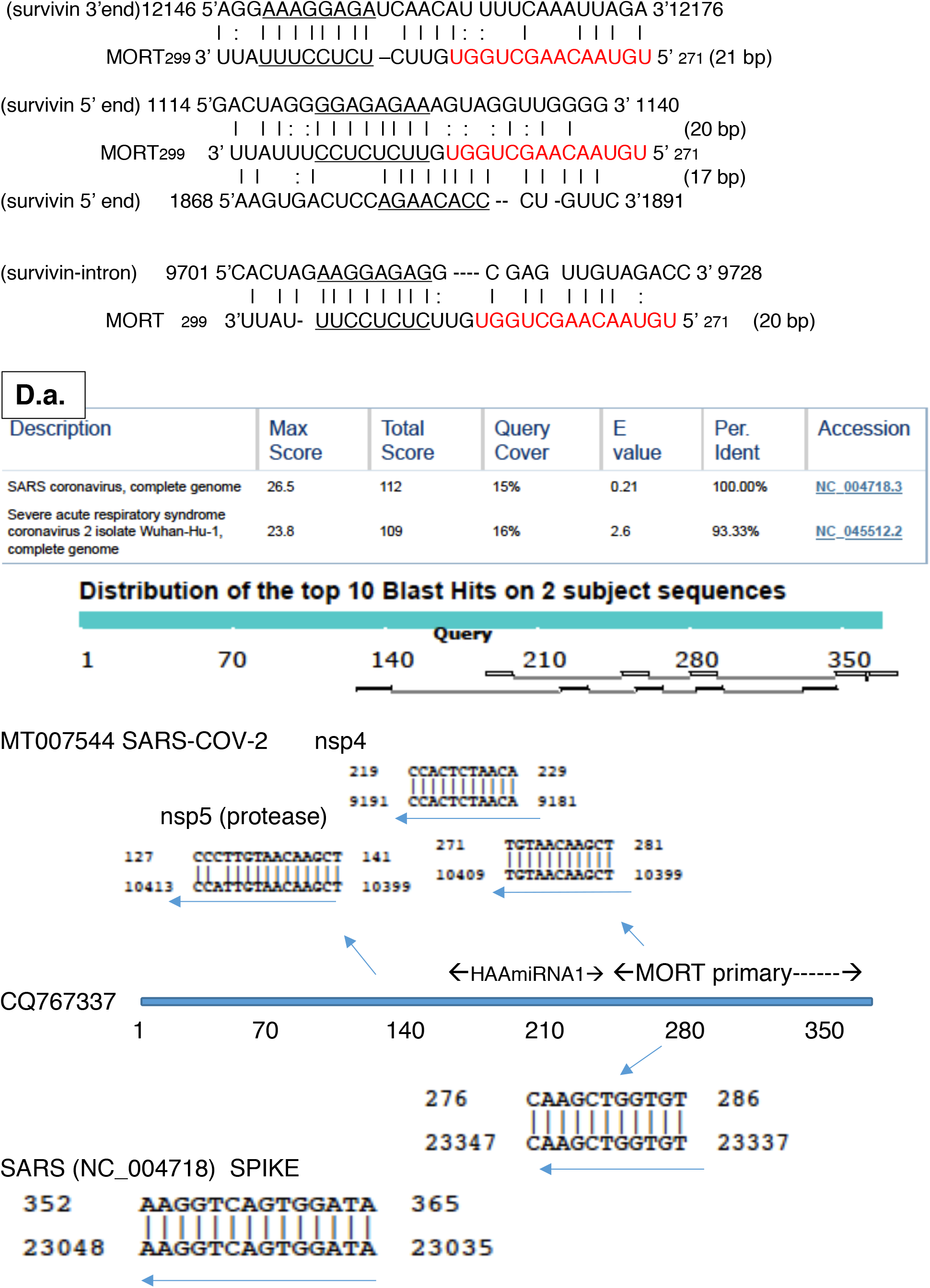
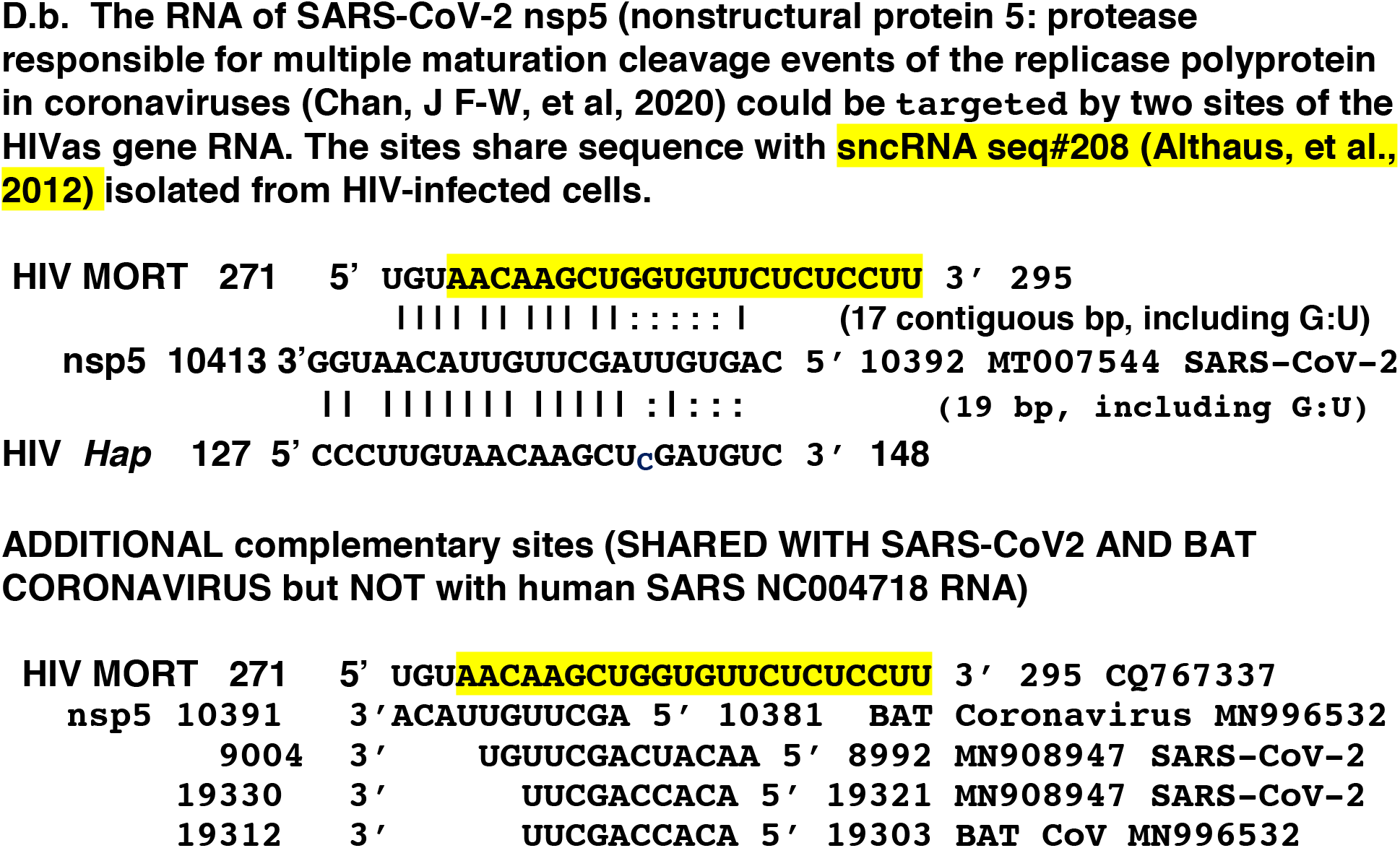
MORT includes characteristics of a microRNA precursor, with conservation across the M clades, sites of homology with human microRNAs, and diverse potential human and viral targets, including SARS-CoV-2. **A.** There is conservation across HIV-1 clades A-J in regions of MORT that would require base-pairing between the 5p (5 prime) and 3p (3 prime) sides of a precursor microRNA structure, but less conservation in less related species, especially notable in HIV-2, A, B, and G (labeled 2AGW, 2B.C1, and 2G.AB) (Kuiken, et al, 2001). A previously identified small noncoding antisense RNA found in HIV-infected cells (Althaus, et al, 2012) is indicated in turquoise below the CQ767337 sequence and suggests where Drosha cuts on the 5p side. An example of the intrinsic antisense RNA (identical to CQ767337) made “in vitro” is shown in supplemental Figure S1A, S1B, S1C. **B.** MirGeneDB (Fromm, et al, 2015) shows that there is significant identity of MORT on the 5p and 3p sides with human microRNAs. MORT 270-303 shares homology regions with seven human microRNAs; MORT 318-350 with three human microRNAs (identical sequence underlined). The presumed mature miRNAs suggested by sncRNA 208 (Althaus, et al, 2012) are underlined. See also supplemental Figure S3 for original search data. **C.** BLAST comparison using NCBI website (National Center for Biotechnology Information; U.S. National Library of Medicine) of MORT 271-300 (GenBank CQ767337) with inhibitor of apoptosis proteins survivin (GenBank U75285), and X-linked inhibitor of apoptosis (XIAP): GenBank accession NG_007264. D36.1997 MORT (GenBank DQ 287274) also has sites of identity with XIAP. See supplemental Figures S2, S4, S5, which includes additional BLAST against Apollon (BIRC 6; NC 000002.12, with 705 matches). **D** BLAST comparison using NCBI website (National Center for Biotechnology Information; U.S. National Library of Medicine) of the HIV antisense gene sequence (GenBank accession CQ767337) with SARS (GenBank accession NC004718) and SARS-CoV-2 (GenBank accession NC045512, MT007544) reveals interaction at disparate sites. Several of the *Hap* gene RNA complementary sites are directed to the same region in SARS-CoV-2 nsp5 RNA (**a** and **b**). The MORT region RNA complementary to SARS is instead directed to the SPIKE protein RNA.

#### Conservation of MORT across HIV-1 but not HIV-2 clades

There is conservation of MORT sequences forming the 33 bp hairpin, as is apparent when the precursor miRNA MORT hairpin is compared with the HIV-1 group M subtype or clade sequences (Figure 5A) (Kuiken, et al, HIV Sequence Compendium 2001). Sequences forming part of the complementary base-paired regions and necessary for the hairpin formation are more conserved between subtypes than the terminal hairpin loop sequences or bulges (Figure 5A). However, there are some significant strain differences, particularly in laboratory adapted types such as pNL4-3, which could alter stability of MORT hairpin formation (Figure S1B).

In contrast, with further distance on the lentiviral genome phylogenetic tree (Kuiken, et al, 2001), the MORT region of sequence becomes much less conserved. Human immunodeficiency virus type 2 (HIV-2) sequences are much more variable, to the extent that an identical MORT secondary/tertiary hairpin structure could not form. The HIV-2 subtype G sequence has 18 nt mutated or deleted in the region corresponding to MORT 5p sequence 271-299, and also represents a much less pathogenic virus for humans (see 2G.AB, Figure 5A).

This is compelling because, as discussed above, the original D36 deletion that was associated with an attenuated HIV and a non-progressor phenotype also had sequence missing from HIV antisense sequence 237-284, of which 14 nt correspond to this region. Revertant virus D36.1997 then regained 274-284, as well as retaining adjacent sequence to 345 (as diagrammed below Figure 5A).

#### MORT shares sequence with human microRNAs

Finally, blast comparison on the database for robustly supported miRNAs, MirGeneDB, (www.mirgenedb.org) yielded significant homologies (E value between 0.51 and 1.8: Figure S3) between the proposed precursor miRNA MORT and **seven** human microRNAs on the 5p side (Figure 5B, examples 1-6 and supplemental Figure S3), as well as their characterized homologues in three other species (Fromm, et al, 2015). The previously identified sncRNA SEQ#208 from HIV-infected cells (Althaus, et al, 2012) is shown above MORT sequence (example 1) with sequence corresponding to the original deletion of D36 shown in red. MORT 272-284 shares 12/13 identical nucleotides (underlined) with MIR-5586 3p mature microRNA. Precursor MORT shares 15/17 identical nucleotides with MIR-7151 5p microRNA (MORT 287-302), 12/13 identical nucleotides with MIR-3173 (MORT 284-296), 10/10 identical nucleotides with both MIR-138 P1 and P2 5p “seed” region microRNA (MORT 278-287), and 11/12 identical nucleotides with MIR-10-P2C (MORT 275-286) (Figure 5B, identical sequence underlined, examples 1-6). The original D36 (and shared SBBC) HIV-1 deleted sequence that was later regained is shown in red within the MORT sequence, and is suggestive of how this deletion might have impacted MORT activity (Figure 5B).

Furthermore, the 3p side of MORT, particularly between 325-353, demonstrates sequence homology with human microRNAs 624 3p, 103-P2 (5p side) and microRNA 29-P1a 3p (Figure 5B, underlined, examples 7-9) (Fromm, et al, 2015). The homology with human miRNA 29a 3p is of particular interest because another retrovirus, Bovine Leukemia Virus, has been shown to produce a microRNA, BLV-B4-3p, that shares partial sequence identity and shared common targets with host miRNA 29a (Kincaid, et al 2012). BLV-B4-3p, with 8 nt shared seed region, can be compared to MORT region 337-360 with 10 nt shared sequence to human miRNA29a 3p seed region (Figure 5B, example 9).

This would support the notion of this MORT locus for apoptosis actually being a site for miRNA generation. It also further implicates MORT in impacting the genes regulated by these human microRNAs, allowing an amplification of MORT gene effect. However, it should be pointed out that the entire HIV antisense gene RNA is capable of forming an exactly complementary duplex with the 3’ end of HIV sense mRNAs (such as nef mRNA) and genomic RNA. Thus Dicer could cut at 21/22 nt intervals for the entire length, producing siRNA, as noted previously (Ludwig, 2008).

#### MORT proposed as a regulatory small RNA or miRNA with target binding sites that include X-linked Inhibitor of Apoptosis (XIAP), survivin, and other inhibitor of apoptosis protein gene mRNA/DNA

MicroRNAs act as regulators of gene expression by direct base-pairing, often to specific target sites in messenger RNA 3’ untranslated regions (UTR), and perform in conjunction with the human Argonaute (AGO) proteins in a miRNA-induced silencing complex (miRISC) (Bartel, 2009; Broughton, et al, 2016; Berezikov, et al, 2007; Helwak, et al, 2013). Canonical animal mature microRNA is formed from longer hairpin-bearing, primary transcripts following sequential endonucleolytic cleavages by the RNase III enzymes Drosha, and then Dicer (Lee, et al, 2003; Du and Zamore, 2005). For an early review of pivotal work on miRNA and siRNA biogenesis see (Filipowicz, et al, 2005). The 33 bp MORT hairpin (Figure 4) has both the requisite terminal loop (15 nt) and flanking sequence for Drosha to identify and then cut about 22 bp away from the junction of the terminal loop and the adjacent stem (Zeng, et al, 2005). This would generate a precursor miRNA-like structure, as shown in Figure 5A, top, (underlined) that could then presumably be further recognized and cut by Dicer in conjunction with TRBP (Chendrimada, et al, 2005; Lee, et al, 2003; Du and Zamore, 2005; Filipowicz, et al, 2005; Zeng, et al, 2005). However, alternative pathways of biogenesis of small regulatory RNAs exist (Bogerd, et al 2010). MiRNAs called mirtrons can derive from short introns cut by Dicer alone (Berezikov, et al, 2007). More recently an alternative pathway bypassing Drosha and Dicer but associated with Argonaute proteins and derived from short introns (80-100 nucleotides) has also been described (Hansen, et al, 2016).

Human miRNA:mRNA duplexes selected on AGO have complex base-pairing interactions possible (Broughton, et al, 2016; Helwak, et al, 2013). More than 60% of the seed region (6-8 nt at the 5’ end of miRNA) interactions had mismatched or bulged nucleotides, and were also often accompanied by specific, non-seed base-pairing (Helwak, et al, 2013). Furthermore, functional miRNA-binding sites were found in all areas of the human target mRNA (Helwak, et al, 2013). With this in mind, we utilized the NIH NCBI BLAST site to investigate whether MORT region (5p) from 271-299 might have potential target sites in genes important in apoptosis pathways, without specifying specific regions (such as 3’ untranslated mRNA) of the genes. We paid special attention to the region nt 274-295, inasmuch as this has already been documented as an HIV-1 antisense small ncRNA (Althaus, et al, 2012), as well as including the sequence regained in revertant SP viruses (Figure 5C, D36.1997).

Remarkably, the HAP-FLAG MORT region 271-299 (CQ767337) demonstrated significant alignments with 122 matches on the human X-linked inhibitor of apoptosis (XIAP) gene reference sequence NG_007264.1. These regions matched either the coding or complementary strand and were identical from 13/13 to 7/7 nt (Figure 5C, supplemental Figure S2). MORT sequence on the 3p strand, nt 322-350 also yielded 98 significant alignments with identities of 9/9 to 7/7 bases suggesting either 5p or 3p regions of MORT could be functioning (Figure 5C and data not shown).

Significantly, investigation of the HIV-1 MORT region 274-300 from the SBBC donor D36 virus in 1997 (D36.1997) demonstrated <u>247</u> BLAST hits with from 12/12 to 7/7 identities matched to the human XIAP sequence (supplemental Figure S4).

One of the inhibitor of apoptosis (IAP) proteins shown to have a functional impact on thymocyte development, and therefore potentially relevant for T cell dysfunction observed in AIDS, was survivin (Ambrosini, et al, 1997; Okada, et al, 2004). BLAST analysis picked up 50 sites in the human apoptosis inhibitor survivin gene (BIRC 5) (GenBank U75285) with between 7/7 to 9/9 nt identical (+/+) or complementary (+/−) to MORT 271-300 (Figure 5C). Visual inspection of the survivin gene sequence around these sites yielded multiple regions in survivin that had sites with up to 21 base-pairs (including G:U) with MORT 271-299 (Figure 5C). If one restricts the viable miRNA region to 274-295 (as with documented SNCRNA 208) there are still up to 16/22 base-paired/MORT:survivin segments, and multiple regions (5’UTR, 3’ UTR and coding) involved between MORT:target, as was found by analysis of human miRNA:mRNA targets bound to AGO (Helwak, et al, 2013). Target sites in the survivin gene also included intronic regions, as well as 5’ and 3’ mRNA ends (Figure 5C).

Further investigation of MORT 271-299 sequences against other mammalian genes encoding inhibitor of apoptosis proteins also suggested potential target sites within Apollon, with BLAST analysis yielding 705 sites with 7/7-13/13 identity (Apollon, supplemental Figure S5). Of interest, many of these sites appear to be in intronic regions (Figure S5). Because other investigators may find other sites useful than those illustrated, we have included the raw data results of BLAST searches for survivin, XIAP, and apollon genes versus MORT271.299 (supplemental Figures S2, S4, S5).

#### HIV antisense gene and SARS-CoV-2 shared or complementary sequences

We considered the possibility that the novel coronavirus, SARS-CoV-2, and HIV-1 might share strategies and/or sequences, despite disparate sizes and replication mechanisms. We were surprised to find that the HIV antisense gene (CQ767337) and SARS-CoV-2 (NC045512, MT007544) shared seven matches of between 11/11 and 14/15 sequence identity (+/+ or +/−) on BLASTn analysis (NCBI). Four of these HIV-1 sites represented identical sequence (+/+) (Figure 3, forward arrows) and three were capable of complementary base-pairing (+/−) to SARS-CoV 2 sequence (Figure 3, backward arrow-CO). Curiously, two of the sites are capable of base-pairing with an identical region in the RNA of nsp5, the protease responsible for cutting much of the orf1ab polyprotein (Figure 5D), On further inspection HIV MORT region 271-295 has the capacity for 17 contiguous base-pairs with MT007544 SARS-CoV-2 nsp5 RNA seq.10392-10413 (Figure 5D.b.). This degree of complementarity could be associated with cleavage of a target mRNA if accompanied by the RNA-induced silencing complex, RISC. HIV *Hap* sequence 127-148 overlaps this same site in SARS-CoV-2 nsp5 RNA and also represents the only HIV antisense gene sequence that is not situated in one of the previously described HAAmiRNAs (Ludwig, 2008) or MORT region (Figure 5D) Of interest, while a similar portion of MORT 271-281 is complementary to Bat Coronavirus MN996532 nsp5 seq 10381-10391, no such site with complementary base-pairing is observed with human SARS NC004718 nsp5 RNA (Figure 5D.b.) The interaction between MORT and SARS appears to be at disparate sites from that with SARS-CoV-2, with several sites focused on the SARS (NC004718) SPIKE protein RNA (Figure 5D.a and b).

## Discussion

There is still no cure for or vaccine to prevent HIV-induced human AIDS (acquired immunodeficiency syndrome). Part of the problem appears to be an inability to induce protective immunity in the human host. It would be impossible to induce immunity if also inducing apoptosis of the respondent immune cells. A major consequence of HIV infection has long been known to be the apoptotic cell death of both directly infected CD4+ T cells and CD4+ cells in close proximity, called uninfected bystander cells (reviewed in Gougeon and Montagnier, 1993; Finkel, et al, 1995; Garg, et al, 2012; Garg and Joshi, 2016; Nardacci, et al, 2015; Kim, et al 2018). While the viral envelope glycoproteins gp120/gp41 (Env) are important for binding to the target human cell CD4 and chemokine receptor (CCR5/CXCR4) molecules, gp 41 is of particular importance in the fusion or partial fusion (hemi-fusion) process and presumed Env-mediated bystander cell death (Garg, et al, 2012).

Yet *in vivo* studies show that genetic mutations or deletions in the HIV envelope gene do not abrogate an AIDS-like phenotype (Hanna, et al, 1998 a,b). In a transgenic (Tg) mouse model where the HIV-1 genome was expressed under the control of regulatory sequences for CD4 (CD4C), the same subset of cells in the mouse thymus and periphery were found to be infected as in humans, with subsequent development of a severe AIDS-like disease (Hanna, et al, 1998 a,b). However, delivery of the intact HIV-1 envelope gene while gag, pol, vif, vpr, tat, vpu and nef genes were mutated did not induce an AIDS-like or MAIDS phenotype, and Tg mice bearing intact Env did not succumb early to a fatal disease (Hanna, et al, 1998). The opposite strategy, with only Env mutated in the HIV-1 proviral transgene, still led to a fatal disease very similar to that seen in the intact CD4C/HIV wild-type Tg mice (Hanna, et al, 1998). Thus, the Env gene did not contribute to the AIDS-like phenotype observed in the HIV transgenic mice *in vivo*, although the transgene was targeted to the right cells by the CD4C regulatory elements (Hanna, et al, 1998 a,b). However, a transgenic (Tg) mouse line exhibited the AIDS phenotype in which <u>only</u> the nef/LTR region of the HIV-1 provirus was incorporated in the transgene, along with mouse T cell receptor promoter/enhancer elements (Lindemann, et al, 1994). No nef protein was demonstrated as being produced in the Tg mice (Lindemann, et al, 1994).

We noted that this DNA transgene incorporated sequence that contained all of the HIV antisense gene, which is located within the HIV-1 LTR (Ludwig, et al, 2006; Ludwig, 2008). The incorporated viral transgene sequence was from the BRU isolate, (GenBank K02013.1), and was 99% identical to the HIV antisense gene (GenBank CQ767337) incorporated into vectors used in our studies (Figures 1–5) (Ludwig, et al, 2006; Ludwig, 2008). Experiments over ten years in our laboratory (from 1995-2005) continually ran the roadblock of cell death induced by the HIV antisense gene. This was observed whether the HIV antisense gene RNA was expressed off the intrinsic HIV-1 antisense initiator (HIVaINR), or from a recombinant vector with a CMV promoter (Figure 2, Figure S1 and as previously described (Ludwig, et al, 2006; Ludwig, 2008). Morphologic studies demonstrated classical features of apoptosis that were enhanced by the HIV antisense gene whether or not the cells also expressed envelope (gp120/gp41) and rev, or all the other HIV-1 proteins except reverse transcriptase (Figure 2, HAP-FLAG). The Hela cell line is a long-lived human cell line from a patient who had cancer, so the HIV antisense gene was also inducing apoptotic cell death in a cancer cell line not expressing any other HIV genes (Figure 2).

The most compelling evidence *in vivo* for a specific region of the HIV-1 DNA provirus as important in CD4+ T cell death and the AIDS phenotype, however, comes from retrospective analysis of studies of the virus in evolution within the human hosts of the Sydney Blood Bank Cohort (SBBC) (Gorry, et al, 2007; Deacon, et al, 1995; Churchill et al, 2004; Churchill, et al, 2006; Dyer, et al, 1999; Dyer, et al 1997; Greenway, et al 1998; Rhodes, et al, 1999; Learmont, et al 1992; Oelrichs, et al 1998; Zaunders, et al, 2011; Learmont, et al, 1999; Jekle, et al, 2002). The SBBC individuals were infected by blood products from a <u>single</u> donor with an attenuated, nef/LTR-deleted virus (D36 in Figures 1, 3). D36 virus replicated *in vivo* in different human hosts, with different outcomes observed. Some individuals, either because of innate host ability to resist infection or the viral genetic changes with replication, have continued for 27 plus years without immunodeficiency (reviewed in Gorry, et al, 2007; Zaunders, et al, 2011). Three individuals, including the donor, with 17 plus years of infection, became phenotypic “revertants” and progressed to AIDS (Gorry, et al, 2007).

These “revertant” viral genetic changes enabled identification of a “locus for apoptosis”. Reanalysis using the GenBank data of the viral sequences and alignment with the HIV antisense gene (GenBank CQ767337) showed a specific region of the *Hap* antisense RNA was involved in the evolution of the viral changes over time (abbreviated and summarized in Figure 3). The SBBC original deletion would abrogate formation of a primary microRNA-like structure (MORT) in the antisense gene RNA, whereas the revertant D36, C54, and C98 viral species regained sequence necessary for hairpin formation (Figures 1, 3A, 3B, and 4). In this process, these revertant viral species thereby regained sequence that would enable formation of an antisense sncRNA already identified in HIV-infected cells (Althaus, et al, 2012). We must stress that no laboratory experiment or set of experiments could possibly duplicate this “experiment of nature” with the study of HIV-1 virus replicating (and mutating) *in vivo* over 17-plus years within the human hosts, along with their progression to AIDS.

### A role for regulatory sncRNAs in HIV

Some have proposed HIV-1 regulatory RNAs or microRNAs, all originating from the sense HIV-1 RNA strand (Ouellet, et al, 2008; Omoto, et al, 2004; Klase, et al, 2007; Bennasser, et al, 2004, Harwig, et al, 2016, Ouellet, et al 2013), whilst others have found this unlikely (Whisnant, et al, 2013; Lin and Cullen, 2007; Pfeffer, et al, 2005, reviewed in Balasubramaniam, et al, 2018,). The argument that no retroviral miRNAs are made because cleavage by Drosha leads to degradation of the primary-miRNA precursor, which would also be the HIV genomic RNA, (Whisnant, et al 2013) assumes transcription occurs only in the sense direction. This argument would not apply to *Hap* antisense strand RNA encoding the HIVaINR antisense (HAA) miRNAs 1,2, and 3 previously proposed (Ludwig, 2008), nor to the MORT primary miRNA-like structure, proposed herein (Figure 4). Figure S1B shows the contribution of HIV strain differences to MORT 252-363 hairpin sequence and inferred stability, which may have a bearing on different laboratories seeing different results. For instance, visual inspection of readily available HIV laboratory isolates (of B subtype sequences) revealed that some of the strains commonly used, when compared to HIV antisense gene sequence (CQ767337), had alterations as follows: LAI was nearly identical, HXB2 had 4 mutations (m) in the primary MORT hairpin and 9 m in antisense gene, SF2: 6 m in MORT; 19 m in antisense gene, NY5: 9 m in MORT; 26 m in antisense gene, and NL4-3: <u>13 m in MORT and 30 mutations</u> in the HIV antisense gene (HIV sequence database compendium 1995 (https://www.hiv.lanl.gov)). Strain sequence differences in the primary MORT hairpin make it apparent that the capacity to base-pair in the stem and form a stable precursor miRNA, and therefore MORT miRNA would be affected, particularly for NL4-3 (see Figure S1B).

The *Hap* gene is unique and unlike the ASP (Miller, 1988; Michaels, et al, 1994; Landry, et al, 2007; Cassan, et al, 2016). *Hap* has protein(s) encoded within the LTR (<u>not the</u> <u>Env</u> region of the DNA), and antisense RNA begins off the HIVaINR in TAR region DNA and terminates within the U3 region (Ludwig, et al, 2006; Ludwig, 2008, Figure S1C)). We had previously proposed this represents an intrinsic RNA regulatory system for the virus (see U.S. patent 5,919,677, Figures 1 and 2, filed in 1997, issued in 1999, also Ludwig, 2008). Others have found the retrovirus, bovine leukemia virus (BLV), utilizes RNA polymerase III to transcribe subgenomic transcripts encoding its miRNAs, demonstrating an alternative mechanism that avoids genomic RNA alterations (Kincaid, et al, 2012).

We strongly believe that HIV-1 does make si/miRNAs but that this has been missed for a variety of reasons based on strain, timing, and perhaps also packaging (see hypothesis below). HIV-1 strain as well as changes that are incurred by growth of isolates *in vit*ro have been shown to have an impact on cell mortality. Thus, viral microRNAs that impact cell mortality should be looked for within hours, not days, when utilizing cells or cell lines (Peden, et al, 1991, Ludwig, et al, 2006; Ludwig, 2008). This notion of early apoptosis with HIV infection is also suggested by studies showing that the CD4+ T cell half-life is 24 hours *in vivo* (Simon and Ho, 2003). Despite the original observations for small, regulatory RNAs acting as antisense translational repressors for developmental transitions in worm larvae in the C. elegans system (Lee and Ambros, 2001), only a few studies have looked for antisense sncRNA in HIV-1 (Althaus, et al, 2012, Schopman, et al, 2012). Yet, with pre-selection for HIV-1 sequences in either strand orientation, the yield of viral vs human sncRNAs was greatly enhanced: 892 individual HIV-1sncRNAs with the selection strategy (Althaus, et al, 2012) as opposed to 125 viral sncRNA clones amongst 47,773 human clones (Yeung, et al, 2009). In addition, the JR-FL strain was utilized to select for small sense and antisense HIV-1 RNAs in HIV-infected cells, and HIV-1 small RNAs were found, including a small antisense 22 nt RNA (Seq. No. 208) mapping exactly to the 5p hairpin region corresponding to MORT 274-295 (Althaus, et al, 2012) (diagrammed in Figure 5A, Figure S1B). This represents the <u>identical</u> region regained (274-284) in revertant SP viruses D36, C54, and C98 (Figures 3, 4, 5A) as the individuals progressed to AIDS.

We believe these studies point the finger at MORT, a primary microRNA-like structure found within the *Hap* antisense RNA (Figure 4), and suggest a mechanism for the early apoptotic death observed in CD4+ T cells (and surrounding bystander cells). The potential for target sites for MORT mi/siRNA in the genes or RNAs of multiple human genes (Figure 5, Table 1, Figure S6), as well as inhibitor of apoptosis protein genes/mRNAs such as X-linked inhibitor of apoptosis (XIAP), survivin, and apollon (Figures S2, S4, S5) require further analysis. The homology to human microRNA segments also implicates the primary MORT hairpin as an RNA regulator (Figures 5B, S3). Studies to elucidate the potential for MORT as an RNA antagonist for the inhibitor of apoptosis protein(s) mRNAs might prove useful for future cancer therapeutics and/or the issue of HIV-1 latency.

**Table 1.**

| Accession | chromosome | description | query sequence | subject sequence | identities | other sites |
| --- | --- | --- | --- | --- | --- | --- |
| NC_018934.2 | X | DEAD box | 282-299 | 23172409-23172426 | 18 of 18 |  |
|  |  | myotubularin-related | 281-298 | 63593776-63593793 | 17 of 18 |  |
|  |  | FERM,PDZ domain | 279-296 | 12757175-12757192 | 17 of 18 |  |
| NC_000023.1 | X | DEAD box | 282-299 | 23123020-23123037 | 18 of 18 |  |
|  |  | mid1;Bcl-6 corepress | 275-290 | 39409735-39409720 | 16 of 16 |  |
| NC_018925.2 | 15 | Meis 2 isoform h242217 | 275-299 | 38105572-38105596 | 23 of 25 | NC_000015.1<br>0 37694082 |
|  |  | arrestin-domain | 274-289 | 98380710-98380695 | 16 of 16 |  |
|  |  | Na-nucleoside co-transporter | 275-292 | 45758855-45758872 | 17 of 18 |  |
|  |  | NT3 growth factor receptor isoform c, m | 279-296 | 88826214-88826197 | 17of18 |  |
| NC_018933.2 | 22 | breakpoint cluster region isoform protein | 279-295 | 23793663-23793679 | 17 of 17 | NC_000024.1<br>1 23439248 |
| NT_187632.1 | 22 genomic |  | 279-295 | 85206-85222 | 17 of 17 |  |
| NC_018913.2 | 2 | tomoregulin2precurso | 273-289 | 195000610-195000594 | 17 of 17 |  |
|  |  | prototransforming growth factor alpha iso | 278-296 | 70616196-70616214 | 18 of 19 |  |
|  |  | ALK tyrosine kinase receptor precursor | 279-296 | 29727509-29727526 | 17 of 18 |  |
|  |  | max dimerization protein 1 isoform 1 | 281-298 | 70086566-70086583 | 17 of 18 |  |
|  |  | sprouty-related EVH1 domain protein2 iso b | 279-296 | 66401278-66401261 | 17 of 18 |  |
|  |  | translin isoform | 281-298 | 123536502-123536485 | 17 of 18 |  |
|  |  | N chimserin isoform2N | 274-291 | 175833140-175833123 | 17 OF 18 |  |
| NC_018915.2 | 4 | putative methyl-transferaseNSUN7 | 280-300 | 40782049-40782029 | 20 OF 21 |  |
|  |  | serine-rich,coiled-coil | 283-299 | 92618998-92618982 | 17 of 17 |  |
|  |  | rap guanine nucleotide exchange factor 2 isox | 279-297 | 159162621-159162639 primary | 18 of 19 |  |

### MORT, a mechanism to kill cancer utilizing apoptosis pathways?

Other viruses have been shown to have genes that impact their host cell survival and apoptosis pathways (Clem, et al, 1991; Birnbaum, et al, 1994; Crook, et al, 1993; Clem and Miller, 1994). The baculovirus p35, Cp-IAP and Op-IAP genes encode proteins that function as viral inhibitors of apoptosis (IAP) (Clem, et al, 1991; Birnbaum, et al, 1994; Crook, et al, 1993; Clem and Miller, 1994). The shared sequence of the Baculovirus Inhibitor of apoptosis-Repeat (BIR) domains was utilized to search for and identify homolog IAPs in other species, including drosophila and humans (Duckett, et al, 1996).

The link between evasion of apoptosis and cancer has been appreciated for a long time (Hanahan and Weinberg, 2000, Hanahan and Weinberg, 2011). Cancer results as cells acquire genes that lead not only to aberrant cellular proliferation, but also resistance to apoptosis. In this regard, it is interesting that potential targets of MORT include X-linked inhibitor of apoptosis (X-IAP), Apollon, and the survivin gene/mRNA (Figure 5C). Survivin appears at the crossroads of at least three mammalian homeostatic networks: control of mitosis, the cellular stress response, and the regulation of apoptosis (Altieri, 2013). The inhibitor of apoptosis (IAP) family contain several members that appear to play significant roles in cancer, with antagonists of IAPs representing a modality for cancer therapy in humans (Finlay, et al, 2017). With specific targeting to cancer cells (instead of T cells), perhaps this MORT region of the HIV antisense gene might prove useful as a RNA tool against the IAP proteins that are up-regulated in cancer.

### Hypothesis: What if HIV-1 utilizes the exosome generation pathway to move Dicer, Ago, and primary/precursor/miRNAs, including MORT, in “RNA grenade” packages to be released against attacking immune cells or invading micro-organisms, including other viruses?

It has long been known that human cells release exosomes, which are less than 150 nm in diameter and consist of a variety of proteins and nucleic acids protected by a lipid bilayer (reviewed in Hessvik, N P, et al, 2018). Valadi, et al., 2007 showed that an array of microRNAs and mRNAs released by exosomes were then taken up and functioned within recipient cells. Cancer cells can release exosomes that contain Dicer, Ago 2, and TAR-RNA binding protein (TRBP) along with pre-microRNAs. These exosomes display cell-independent ability to process pre-miRNAs into miRNAs (Melo, S.A., et al., 2014). Thus, cancer cells already demonstrate that exosomal release and up-take into other cells can reprogram target cell transcriptome(s) (Melo, S.A., et al, 2014). This would be a remarkably efficient mechanism whereby an HIV-1-infected cell could mediate changes and genetic exchange with surrounding bystander cells. MORT pre-miRNA or mi/siRNAs, prepackaged with the RISC-loading complex proteins Dicer, TRBP, and Ago within the exosome, and then released and taken-up by recipient cells could facilitate the bystander CD4+ T cell/other immune cell death observed with HIV infection. The bulk of cells induced to die are the surrounding “bystander” cells (Finkel, et al, 1995) and may represent surrounding host cells specifically responding to HIV infection (CD4+ T cells, cytotoxic T cells, dendritic cells). We do not know the contribution of *Hap* gene proteins or other HIV proteins to an exosomal-mediated process, but MORT miRNA acting as an antagonist to IAP mRNAs via the RNA interference or silencing pathways within cells could provide an explanation for the rapid apoptotic cell death observed following introduction of a recombinant vector containing the 252-363 MORT hairpin (Figures 2, 4, 5, S1). If the RISC loading complex is associated early with MORT, packaged into late endosomes/multi-vesicular bodies (MVB), fused with the plasma membrane, and released as exosomes, these associated miRNAs made by HIV may prove more difficult to detect by standard methods.

This provides some insight into the inability to develop effective protective immunity to HIV: if the surrounding, bystander immune cells (CD4 + T cell, dendritic cell, etc.) that can recognize specific HIV epitopes in context of MHC are killed off before they can even proliferate and amplify the immune response?

Viruses are great imitators and scavengers. If a host cell genetic element can be implemented to further viral advantage, it often will be. Thus, it is not a surprise that HIV-1 has “borrowed” genetic elements (including portions of MORT) from its human host (Figure 5, Supplemental Table 1, Figure S6). However, it is more puzzling to observe the shared homologies between the HIV antisense gene *Hap* and SARS-CoV-2 (Figure 5D) Seven sites with between 11/11 to 13/15 nt +/+ or +/− were identified by nBLAST (Figures 3, 5D). Two of the sites in *Hap* with complementary sequence to SARS-CoV-2 are directed to the same region in SARS-CoV-2 nsp5 (protease) RNA (Figure 5Da,b). The protease nsp5 is responsible for cutting much of the SARS-CoV-2 orf1ab polyprotein gene product (Chan, J F-W, et al, 2020). Shared sequence between two viruses related to the novel coronavirus (human SARS, and Bat coronavirus), reveal an interesting pattern. Both Bat coronavirus and SARS-CoV-2, but not human SARS, could be targeted within their orf1ab RNA by the HIV-1 antisense gene MORT 271-287 region mi/siRNA. However an alternate sequence in SARS is targeted corresponding to the Spike protein gene/mRNA (Figure 5D). The extensive (17 bp) complementary region between HIV-1 MORT and SARS-CoV-2 nsp5 mRNA immediately suggests an RNA silencing method for use against the novel coronavirus-2019. Why not use what cancer exosomes have taught us to manufacture a RNA therapy: a MORT-mi/siRNA:Ago “slicer” against the nsp5 mRNA protease (protein dicer)? In addition, the “RNA therapy” exosomes could incorporate specific targeting molecules such as SPIKE antibody (Fab2) so as to only kill SARS-CoV-2-infected cells?

Here, we propose a strategy and/or functional mechanism for HIV-1 to have evolved a death-inducing “regulatory” RNA or MORT. Viruses are not only opportunistic, but they are “selfish”. When they take over cellular functions to replicate, they turn off cellular functions for intrinsic cellular use. They may also grab opportunities to prevent any outside intervention either from immune attack or other micro-organisms (such as viruses) that also want to use the cell or cell programs to replicate.

## Methods

### Cells, transfections and analysis of apoptosis

The following reagents and cell lines were obtained through the AIDS Research and Reference Reagent Program, Division of AIDS, NIAID, NIH: the HL2/3 cell line containing stably integrated copies of the HIV-1 molecular clone HXB2/3gpt from Drs. Barbara K Felber and George N. Pavlakis (Felber and Pavlakis, 1988; Ciminale, et al, 1990); HeLa, a human cervical epithelial carcinoma cell line from Dr. Richard Axel (Maddon, et al, 1986), Clone69T1RevEnv from Dr. Joseph Dougherty (Yu, et al, 1996) and pHIV-CAT from Drs. Gary Nabel and Neil Perkins (Nabel and Baltimore, 1987). pHIV-CAT was used to amplify by PCR the desired U3-R sequences for cloning of HAP-FLAG, and the HIV-1 sequences are derived from the original French isolate BRU, also called LAV strain (Wain-Hobson, et al, 1991). Clone 69T1RevEnv was a selected clone from the transfection of HeLa-derived HtTa-1 with T1RevEnv, and could be induced to express HIV-1 Rev and Env proteins by removal of tetracycline from the culture medium (Yu, et al, 1996). HeLa and Clone 69T1RevEnv were used as control cell lines for HL2/3 (Felber and Pavlakis, 1988; Ciminale, et al, 1990) in these experiments. HL2/3 was selected for high-level production of Gag, Env, Tat, Rev, and Nef proteins, but did not produce detectable reverse transcriptase or mature virions (Felber and Pavlakis, 1988; Ciminale, et al, 1990). All cell lines were maintained and propagated in cell cultures as recommended by the contributor.

The HIV-1 antisense gene sequences corresponding to SEQ ID 27 nt 14-367 of U.S. patents 7700725, 7700726, and 8765461, and European patent 1359221 (GenBank accession number CQ767337) were directionally cloned into the mammalian expression vector pCMVTag 4a with a carboxyterminal FLAG epitope tag (DYKDDDDK) (HAP-FLAG), as previously described (Ludwig, et al, 2006; Ludwig, 2008). The HIV antisense gene sequence and orientation were confirmed by sequencing. The pCMVTag4a (GenBank accession number AF073000) and pCMVTag 4 expression control plasmid containing the firefly luciferase gene fused with the FLAG tag (luc-FLAG) were obtained from Stratagene. The plasmid 15cxCAT contained the interleukin-2 (IL-2) enhancer and promoter driving the CAT gene and was a kind gift from Dr. Gerald R. Crabtree (Durand, et al, 1988).

Transfections of HeLa (Maddon, et al, 1986), Clone69T1RevEnv (Yu, et al, 1996), and HL2/3 (Felber and Pavlakis, 1988; Ciminale, et al, 1990) with HAP-FLAG and the control vectors, pCMVTag4 (luc-FLAG) and p15cxCAT (IL-2-CAT) were performed simultaneously with identical transfectam and solution reagent concentrations, as recommended by the manufacturer (Stratagene) and as previously described in detail (Ludwig, et al, 2006). The same number of each cell type (5 × 105 cells) were plated in four 6-well plates with coverslips (within the wells), grown overnight at 37 C, 5%CO2 and then underwent transfection four ways (transfectam solution alone, or with either HAP-FLAG, IL-2CAT or luc-FLAG vectors). Coverslips were removed from 6-well plates, transferred to 95% ethanol, and then mounted on superfrost slides using cytoseal 60. After drying ON, they either were submitted for H&E staining or immunoperoxidase staining, as previously described (Ludwig, et al, 2006).

#### Analysis of apoptosis

Slides were surveyed at low power, then three areas selected for closer analysis and counting (400x total magnification). Numbers represent data presented as mean +/− SEM of three counts per hpf. The pathologist was unaware of experimental conditions while performing this analysis. Histologically, apoptosis is characterized by condensation and hyperchromasia of the cell’s nucleus, followed by nuclear fragmentation and cell shrinkage. The nucleus and cytoplasm eventually break apart into fragments called “apoptotic bodies” which are eventually phagocytized by mononuclear cells (Kerr, et al, 1972).

| Accession | chromosome | description | query sequence | subject sequence | identities | other sites |
| --- | --- | --- | --- | --- | --- | --- |
| NC_018915.2 | 4 | APOBEC-3G like | 273-287 | 171115912-171115526 | 15 of 15 |  |
|  |  | TRIM L ligase | 280-294 | 190089502-190089488 | 15 of 15 | 190701563- |
|  |  | insulin-like growth factor binding protein7 | 272-289 | 61004533-61004550 | 17 of 18 |  |
|  |  | mineralocorticoid receptor isoform | 281-298 | 149481706-149481723 | 17 of 18 |  |
|  |  | janus kinase and micro-tubule interacting protein 1 isoform1 | 279-296 | 6077308-6077291 | 17 of 18 |  |
|  |  | polyadenylate-binding protein 4 like | 279-296 | 135591436-135591419 | 17 of 18 |  |
| NC_018916.2 | 5 | zinc finger protein | 277-297 | 123806586-123806606 | 20 of 21 |  |
|  |  | little elongation complex subunit | 284-300 | 5607063-5607079 | 17 of 17 |  |
|  |  | tenurin-2 | 281-297 | 166996269-166996253 | 17 of 17 |  |
|  |  | endoplasmic reticulum aminopeptidase 1 iso a | 277-292 | 95557173-95557158 | 16 of 16 |  |
|  |  | casein kinase 1 iso gamma 3 | 282-300 | 122677387-122677405 | 18 of 19 |  |
|  |  | disabled homolog 2 iso | 281-298 | 39648101-39648118 | 17 of 18 |  |
|  |  | transcription factor BTF3 iso | 282-299 | 72234367-72234384 | 17 of 18 |  |
|  |  | semaphorin-5a precurs | 278-299 | 9198489-9198468 | 20 of 22 |  |
|  |  | fibroblast growth factor 1 isoform 2 precursor | 278-295 | 141507573-141507556 | 17 of 18 |  |
|  |  | serine-threonine protein kinase 10 | 276-295 | 170923406-170923389 | 17 of 18 |  |
| NC_018919.2 | 7 | mitotic spindle assembly checkpoint proteinMAD1 isoform | 279-295 | 1876177-1876161 | 17 of 17 |  |
|  |  | forkhead box L1-like | 271-289 | 457086-457104 | 18 of 19 |  |
|  |  | homeobox proteinMOX | 278-295 | 16102717-16102734 | 17 of 18 |  |
|  |  | transcriptional activator | 280-297 | 42733743-42733760 | 17 of 18 |  |
|  |  | frizzled | 279-296 | 90978083-90978066 | 17 of 18 |  |
|  |  | reelin isoform b precurs | 281-298 | 103456111-103456094 | 17 of 18 |  |
| NC_018919.2 | 8 | dual specificity protein phosphatase 4 isoform | 277-293 | 29518700-29518684 | 17 of 17 |  |
|  |  | tumor necrosis factor receptor family 11B | 284-299 | 120022458-120022443 | 16 of 16 |  |
|  |  | zinc finger protein 754 | 280-298 | 81676064-81676044 | 18 of 19 |  |
|  |  | calbindin | 273-291 | 91414660-91414642 | 18 of 19 |  |
|  |  | XK related protein 6 | 277-294 | 10975434-10975451 | 17 of 18 |  |
|  |  | DNA repair scaffolding protein isoform 1 | 275-292 | 48490003-48489986 | 17 of 18 |  |
|  |  | hepatocyte nuclear factor 4 gamma | 281-298 | 76729513-76729496 | 17 of 18 |  |
| NC_000024.10 | Y, PRIMARY | deleted in azospermia multiple isoforms | 281-297,299 | multiple | 17 of 17, also 18 of 19 |  |
|  |  | argonaute | 274-296 | 36504106-36504128 | 21 of 23 |  |
|  |  | plexin A2 precursor | 277-294 | 210033085-210033102 | 17 of 18 |  |
|  |  | cytosolic carboxypeptidase 6 | 278-295 | 50476263-50476246 | 17 of 18 |  |
|  |  | leucine-rich repeat containing protein 7 | 281-298 | 70484877-70484860 | 17 of 18 |  |
|  |  | endophilin B1 isoform | 271-296 | 87361618-87361593 | 23 of 25 |  |
|  |  | small conductance Ca-activated K channel pro | 279-296 | 156114194-156114177 | 17 of 18 |  |
|  |  | retinoic acid receptor RXR gamma a iso a | 277-294 | 166832684-166832667 | 17 of 18 |  |
|  |  | transmembrane coiled-coiled protein 2 | 279-296 | 206667063-206667046 | 17 of 18 |  |
|  |  | dynein heavy chain14 | 282-298 | 226679495-226679478 | 17 of 18 |  |
| NC_018914.2 | 3, ALTERNATE | ERC protein 2 | 279-294 | 55687644-55687659 | 16 of 16 |  |
|  |  | tRNA splicing endonuclease subunit | 281-299 | 12495870-12495888 | 18 of 19 |  |
|  |  | contactin-3 precursor | 281-299 | 74722742-74722760 | 18 of 19 |  |
|  |  | A disintegrin and metallo proteinase | 276-294 | 64650914-64650896 | 18 of 19 |  |
|  |  | tyrosine protein kinase RYK isoform 1 | 281-299 | 133844607-133844589 | 18 of 19 |  |
|  |  | programmed cell death 6 interacting protein | 286-299 | 33919563-33919576 | 14 of 14 |  |
|  |  | IQmotif-containing J schwannamin-interacting protein 1 | 277-294 | 159290919-159290936 | 17 of 18 |  |
| NC_018917.2 | 6, ALTERNATE | transcription factor HV EP2 | 280-299 | 143486767-143486786 | 19 of 20 |  |
|  |  | F-box LRR repeat protein | 282-299 | 99971296-99971313 | 17 of 18 |  |
|  |  | glutamate receptor, ionotropic | 282-298 | 102810361-102810378 | 17 of 18 |  |
|  |  | mannosyl-oligosaccharide | 281-298 | 121384069-121384086 | 17 of 18 |  |
|  |  | vascular non-inflammatory molecule | 273-290 | 133331637-133331654 | 17 of 18 |  |

## Author Contributions

L.B.L conceived of the project, organized and evaluated the data, and wrote the paper. M.S.A. re-evaluated the pathology section and helped edit the paper. Both authors read and approve the paper.

## Acknowledgments

We are indebted to the researchers who shared HIV-1 genetic sequences with the NCBI and GenBank, as well as their papers with the research community. We also appreciate the NIH, NCBI and the ability to BLAST because of the Human Genome Project. We are further indebted to all the many individuals who contributed research materials, including cells and cell lines, which made the human research possible. We are grateful also to the National Institutes of Health for earlier funding from 1996-2004 (NIAID R29AI38114, R21AI46960, and R21 AI52006 (LBL)). Gratitude to Phil Dvoretsky, M.D., pathologist, who evaluated the original work done more than 20 years ago.

## References

1. Althaus, C. F., Vongrad, V., Niederost, B., Joos, B., Di Giallonardo, F., Rieder, P., Pavlovic, J., Trkola, A., Gunthard, H. F., Metzner, K.J., and Fischer, M. (2012) Tailored enrichment strategy detects low abundant small noncoding RNAs in HIV-1 infected cells. Retrovirology 9:27.

2. Altieri, D.C. (2013) Targeting survivin in cancer. Cancer Lett. (332 (2)), 225–228.

3. Ambrosini, G., Adida, C., and Altieri, D.C. (1997) A novel anti-apoptosis gene, survivin, expressed in cancer and lymphoma. Nat Med 3(8), 917–921.

4. Balasubramaniam, M., Pandhare, J, and Dash, C. (2018) Are microRNAs important players in HIV-1 infection? An update. Viruses 10, 110;doi:10.3390/v10030110.

5. Barmania, F., and Pepper, M.S. (2013). C-C chemokine receptor type five (CCR5): an emerging target for the control of HIV infection. Applied & Translational Genomics 2, 3–16.

6. Bartel, D.P. (2009) MicroRNAs:target recognition and regulatory functions. Cell 136:215–233.

7. Bennasser, Y., Le, S.Y., Yeung, M.L., and Jeang, K.T. (2004) HIV-1 encoded candidate micro-RNAs and their cellular targets. Retrovirology 1, 43.

8. Berezikov, E, Chung, W-J., Willis, J., Cuppen, E., and Lai, E. C. (2007) Mammalian Mirtron genes. Mol. Cell 28: 328–336.

9. Birnbaum MJ, Clem RJ, and Miller LK. (1994) An apoptosis-inhibiting gene from a nuclear polyhedrosis virus encoding a polypeptide with Cys/His sequence motifs. J Virol 68, 2521–2528.

10. Bogerd, H.P., Kamowski, H.W., Cai, X., Shin, J., Pohlers, M., and Cullen, B.R. (2010). A mammalian herpesvirus uses noncanonical expression and processing mechanisms to generate viral microRNAs. Mol Cell 37 (1):135–142.

11. Broughton, J. P., Lovci, M.T., Huang, J.L., Yeo, G.W., and Pasquinelli, A.E. (2016). Pairing beyond the seed supports microRNA targeting specificity. Mol. Cell 64:320–333.

12. Carrington, M., Dean, M., Martin, M.P., and O’Brien, S.J. (1999). Genetics of HIV-1 infection: chemokine receptor CCR5 polymorphism and its consequences. Human Molecular Genetics 8, 1939–1945.

13. Cassan E, Arigon-Chifolieau AM, Mesnard JM, Gross A, Cascuel O. (2016) Concomitant emergence of the antisense protein gene of HIV-1 and of the pandemic. Proc Natl Acad Sci U.S. A. 113 (41); 11537–11542.

14. Chan, J F-W, Kok, K-H, Zhu, Z., Chu, H., To, K K-W, Yuan, S., and Yuen, K-Y. (2020) Genomic characterization of the 2019 novel human pathogenic coronavirus isolated from a patient with atypical pneumonia after visiting Wuhan. Emerging Microbes & Infections (9) https://doi.org/10.1080/22221751.2020.1719902

15. Chendrimada, T.P., Gregory, R.I., Kumaraswamy, E., Norman, J., Cooch, N., Nishikura, K., Shiekhattar, R. (2005). TRBP recruits the Dicer complex to Ago 2 for microRNA processing and gene silencing. Nature 436, 740–744.

16. Churchill M., Sterjovski, J., Gray, L., Cowley, D., Chatfield, C., Learmont, J., Sullivan, J. S., Crowe, S.M., Mills, J., Brew, B.J., Wesselingh, S.L., McPhee, D.A., and Gorry, P.R. (2004). Longitudinal analysis of nef/long termnal repeat-deleted HIV-1 in blood and cerebrospinal fluid of a long-term survivor who developed HIV-associated dementia. J. Infect. Dis. 190 (12), 2181–2186.

17. Churchill, M. J., Rhodes, D. I., Learmont, J.C., Sullivan, J.S., Wesselingh, S. L., Cooke, J.R., Deacon, N.J., and Gorry, P.R. (2006). Longitudinal analysis of human immunodeficiency virus type 1 nef/long terminal repeat sequences in a cohort of long-term survivors infected from a single source. J. Virol. 80 (2), 1047–1052.

18. Ciminale, V., Felber, B.K., Campbell, M. and Pavlakis, G.N. (1990) A bioassay for HIV-1 based on Env-CD4 interaction. AIDS Res Hum Retroviruses 6, 1281–1287.

19. Clarke, AR., Purdie, CA, Harrison, DJ, Morris, RG, Bird, CC, Hooper, ML and Wyllie, AH. (1993). Thymocyte apoptosis induced by p53-dependent and independent pathways. Nature 362, 849–852.

20. Clem, RJ, Fechneimer M, and Miller, LK. (1991) Prevention of apoptosis by a baculovirus gene during infection of insect cells. Science 254, 1388–1390.

21. Clem, R.J. and Miller, L.K. (1994) Control of programmed cell death by the baculovirus genes p35 and iap. Mol. Cell Biol., 14, 5212–5222.

22. Cohen, J.J. (1993) Apoptosis. Immunology Today 14 (3), 126–130.

23. Crook, N.E., Clem, R.J. and Miller, I.K. (1993). An apoptosis-inhibiting baculovirus gene with a zinc finger-like motif. J. Virol 67, 2168–2174.

24. Deacon, N.J., Tsykin, A., Solomon, A., Smith, K., Ludford-Menting, M., Hooker, D.J., McPhee, D.A., Greenway, A.L., Ellett, A., Chatfield, C., Lawson, V.A., Crowe, S., Maerz, A., Sonza, S., Learmont, J., Sullivan, J.S., Cuningham, A., Dwyer, D., Dowton, D., Mills, J. (1995) Genomic structure of an attenuated quasi species of HIV-1 from a blood transfusion donor and recipients. Science 270, 988–991.

25. Dean, M., Carrington, M., Winkler, C., et al (1996). Genetic restriction of HIV-1 infection and progression to AIDS by a deletion allele of the CKR5 structural gene. Science, 273, 1856–62 {Erratum, Science 1996, 274:1069}.

26. Dyer, W.B., Ogg, G.S., Demoitie, M.A., Jin, X., Geczy, A.F., Rowland-Jones, S.L., McMichael, A.J., Nixon, D.F., Sullivan, J.S. (1999) Strong human immunodeficiency virus (HIV)-specific cytotoxic T-lymphocyte activity in Sydney Blood Bank Cohort patients infected with nef-defective HIV type 1. J. Virol. 73, 436–443.

27. Dyer, W.B., Geczy, A.F., Kent, S.J., McIntyre, L.B., Blasdall, S.A., Learmont, J.C., Sullivan, J.S. (1997) Lymphoproliferative immune function in the Sydney Blood Bank Cohort, infected with natural nef/long terminal repeat mutants, and in other long-term survivors of transfusion acquired HIV-1 infection. AIDS 11, 1565–1574.

28. Du, T., and Zamore, P.D. (2005) microPrimer: the biogenesis and function of microRNA. Development 132: 4645–4652.

29. Duckett, C.S., Nava, V.E., Gedrich, R.W., Clem, R.J., VanDongen, J.L., Gilfillan, M.C., Shiels, H., Hardwick, J.M., and Thompson, C.B. (1996). A conserved family of cellular genes related to the baculovirus *iap* gene and encoding apoptosis inhibitors. The EMBO J., 15 (11), 2685–2694.

30. Durand, D. B., Shaw, J., Bush, M.R., Replogle, R.E., Belagaje, R., and Crabtree, G.R. (1988). Characterization of antigen receptor response elements within the interleukin-2 enhancer. Mol.Cell. Biol. 8, 1715–1724.

31. Elbashir, S. M., Lendeckel, W, and Tuschl, T. (2001) RNA interference is mediated by 21- and 22-nucleotide RNAs. Genes & Development 15: 188–200.

32. Felber, B.K., and Pavlakis, G.N.(1988). A quantitative bioassay for HIV-1 based on trans-activation. Science 239, 184–187.

33. Filipowicz, W., Jaskiewicz, L., Kolb, F.A, and Pillai, R.S. (2005) Post-transcriptional gene silencing by siRNAs and miRNAs. Current Opinion in Structural Biology 15:331–341.

34. Finkel, T., Tudor-Williams, G., Banda, N., Cotton, M., Curiel,T., Monks, C., Baba,T, Ruprecht, R. and Kupfer, A (1995) Apoptosis occurs predominantly in bystander cells and not in productively infected cells of HIV-and SIV infected lymph nodes. Nat. Med 1, 129–134.

35. Finlay, D., Teriete, P., Vamos, M., Cosford, NDP., and Vuori, K. (2017) Inducing death in tumor cells:roles of the inhibitor of apoptosis proteins {version 1; referees;3 approved} F1000Research 2017, 6(F1000Faculty Rev):587

36. Fromm, B., Billipp, T., Peck, L.E., Johansen, M., Tarver, J.E., King, B.L., Newcomb, J.M., Sepere, L.F., Flatmark, K., Hovig, E., and Peterson, K.J. (2015). A uniform system for the annotation of human microRNA genes and the evolution of the human microRNAome. Annu. Rev. Genet. 49, 213–242.

37. Garg, H., Mohl, J. and Joshi, A (2012) HIV-1 induced bystander apoptosis. Viruses 4, 3020–3043.

38. Garg, H and Joshi, A. (2016) Host and viral factors in HIV-mediated bystander apoptosis. Viruses 9, 237

39. Gorry, P.R., McPhee, D.A., Verity, E., Dyer, W.B., Wesselingh, S. L., Learmont, J., Sullivan, J.S., Roche, M., Zaunders, J.J., Gabuzda, D., Crowe, S. M., Mills, J., Lewin, S. R., Brew, B. J., Cunningham, A.L., and Churchill, M.J (2007). Pathogenicity and immunogenicity of attenuated, nef-deleted HIV-1 strains in vivo. Retrovirology 4:66.

40. Gougeon, M. and Montagnier, L (1993) Apoptosis in AIDS. Science 260, 1269–1270.

41. Greenway, A.L., Mills, J., Rhodes, D., Deacon, N.J., McPhee, D.A. (1998) Serological detection of attenuated HIV-1 variants with nef gene deletions. AIDS 12, 555–561.

42. Hanahan, D. and Weinberg RA (2000) The hallmarks of cancer. Cell 100, 1:57–70.

43. Hanahan, D. and Weinberg RA (2011) Hallmarks of cancer: the next generation. Cell, 144 (5)646–674.

44. Hanna, Z., Kay, D.G., Rebai, N., Guimond, A., Jothy, S., and Jolicoeur, P. (1998) Nef harbors a major determinant of pathogenicity for an AIDS-like disease induced by HIV-1 in transgenic mice. Cell 95, 163–175.

45. Hanna, Z., Kay, D.G., Cool, M., Jothy, S., Rabai, N., and Jolicoeur, P. (1998). Transgenic mice expressing human immunodeficiency virus type 1 in immune cells develop a severe AIDS-like disease. J. Virol. 72, 121–132.

46. Hansen, T.B., Veno, M.T., Jensen, T.I., Schaefer, A., Damgaard, C. K., and Kjems, J. (2016). Argonaute-associated short introns are a novel class of gene regulators. Nat. Commun. 7:11538 doi: 10.1038/ncomms11538.

47. Harwig, A., Jongejan, A., van Kampen, A.H.C., Berkhout, B, and Das, A.T. (2016) Tat-dependent production of an HIV-1 TAR-encoded miRNA-like small RNA. NAR 44, 9; 4340–4353, doi:10.1093/nar/gkw167.

48. Helwak, A., Kudia, G., Dudnakova, T., and Tollervey, D. (2013). Mapping the human miRNA interactome by CLASH reveals frequent noncanonical binding. Cell 153 (3), 654–665.

49. Hessvik, N.P., and Liorente, A. (2018) Current knowledge on exosome biogenesis and release. Cell. Mol. Life Sci. 75, 193–208.

50. Jekle A, Schramm, B., Jayakumar, P., Trautner, V., Schols, D., De Clercz, E., Mills, J., Crowe, S.M., and Goldsmith, M.A. (2002) Coreceptor phenotype of natural human immunodeficiency virus with nef-deleted evolves in vivo, leading to increased virulence. J Virol. 76 (14), 6966–6973.

51. Kelleher, A.D., and Zaunders, J.J. (2006). Decimated or missing in action: CD4+ T cells as targets and effectors in the pathogenesis of primary HIV infection. Curr. HIV/AIDS Rep. 3 (1), 5–12.

52. Kerr, JFR, Wyllie, AH., and Currie, AR. (1972) Apoptosis: A basic biological phenomenon with wide-ranging implications in tissue kinetics. Br J Cancer 26, 239–257.

53. Kim, Y., Anderson, J.L., and Lewin, S.R. (2018) Getting the “kill” into “shock and kill”: strategies to eliminate latent HIV. Cell Host Microbe 23(1),14–26, doi:10.1016/j.chom.2017.12.004

54. Kincaid, R.P., Burke, J.M., and Sullivan, C.S. (2012) RNA virus microRNA that mimics a B-cell oncomiR. PNAS (109) 8: 3077–3082.

55. Kirchhoff, F., Greenough, T.C., Brettler, D.B., Sullivan, J.L., Desrosiers, R.C. (1995) Brief report: absence of intact nef sequences in a long-term survivor with nonprogressive HIV-1 infection. N. Engl. J. Med. 332, 228–232.

56. Klase, Z., Kale, P., Winograd, R., Gupta, M.V., Heydarian, M., Berro, R., McCaffrey, T., and Kashanchi, F. (2007) HIV-1 TAR element is processed by Dicer to yield a viral micro-RNA involved in chromatin remodeling of the viral LTR. BMC Mol. Biol. 8, 63.

57. Kolb, J.P., Oguin, T. H., Oberst, A., and Martinez, J. (2017) Programmed cell death and inflammation: winter is coming. Trends in Immunology (Cell Press Reviews) 38 (10), 705–718.

58. Kuiken, C., Foley, B., Hahn, B., Marx, P., McCutchan, F., Mellors, J., Wolinsky S., and Korber, B., editors. HIV Sequence Compendium 2001, published by Theoretical Biology and Biophysics Group, Los Alamos National Laboratory.

59. Landry S, Halin M, Lefort S, Audet B, Vaquero C, Mesnard JM, Barbeau B. (2007). Detection, characterization and regulation of antisense transcripts in HIV-1. Retrovirology 4:71.

60. Learmont J, Tindall B, Evans L, Cunningham A, Cunningham P, Wells J, Penny R, Kaldor J, and Cooper D.A. (1992) Long-term symptomless HIV-1 infection in recipients of blood products from a single donor. Lancet 340, 863–867.

61. Learmont JC, Geczy AF, Mills J, Ashton LJ, Raynes-Greenow C H, Garsia RJ, Dyer W B, McIntyre L, Oelrichs R B, Rhodes DI, Deacon NJ, and Sullivan JS. (1999). Immunologic and virologic status after 14 to 18 years of infection with an attenuated strain of HIV-1. N Engl J Med 340 (22), 1715–1722.

62. Lee, Y., Ahn, C., Han, J., Choi, H., Kim, J., Yim., J., Lee, J., Provost, P., Radmark, O., Kim, S., and Kim V.N. (2003). The nuclear RNase III Drosha initiates microRNA processing. Nature 425: 415–419.

63. Lin, J. and Cullen, B.R. (2007). Analysis of the interaction of primate retroviruses with the human RNA interference machinery. J.Virol. 81 (22): 12218–12226.

64. Lindemann, D., Wilhelm, R., Renard, P., Althage, A., Zinkernagel, R., Mous, J. (1994) Severe immunodeficiency associated with a human immunodeficiency virus 1 NEF/3’ –long terminal repeat transgene. J. Exp. Med. 179, 797–806.

65. Liu, R., Paxton, W.A., Choe, S., Ceradini, D., Martin, S.R., Horuk, R., et al. (1996) Homozygous defect in HIV-1 coreceptor accounts for resistance of some multiply-exposed individuals to HIV-1 infection. Cell 86, 367–377.

66. Ludwig, L.B., Hughes, B.J., and Schwartz, S.A. (1995) Biotinylated probes in the electrophoretic mobility shift assay to examine specific dsDNA, ssDNA or RNA-protein interactions. Nucleic Acids Res 23, 3792–3793.

67. Ludwig, L.B., Ambrus, J.L.,Jr., Krawczyk, K.A, Sharma, S., Brooks, S., Hsiao, C-B. and Schwartz, S.A. (2006) Human Immunodeficiency Virus-Type 1 LTR DNA contains an intrinsic gene producing antisense RNA and protein products. Retrovirology 3:80.

68. Ludwig, L.B. (2008). RNA silencing and HIV: A hypothesis for the etiology of the severe combined immunodeficiency induced by the virus. Retrovirology, 5:79.

69. Maddon, P.J., Dalgleish, A.G., McDougal, J.S., Clapham, P.R., Weiss, RA., and Axel, R. (1986). The T4 gene encodes the AIDS virus receptor and is expressed in the immune system and the brain. Cell 47, 333–348

70. Mariani, R., Kirchhoff, F., Greenough, T.C., Sullivan, J.L., Desrosiers, R.C., Skowronski, J. (1996) High frequency of defective nef alleles in a long-term survivor with nonprogressive human immunodeficiency virus type 1 infection. J. Virol. 70, 7752–7764.

71. Markham, N.R., and Zuker, M. (2008). UNAFold: Software for nucleic acid folding and Hybridization, in Data, Sequence Analysis, and Evolution, J. Keith, ed., Bioinformatics: Volume 2, chapter 1, p3–31, Humana Press Inc.

72. Melo, S.A., Sugimoto, H., O’Connell, J.T., Kato, N., Villanueva, A., Vidal, A., Qiu, L., Vitkin, E., Perelman, L.T., Melo, C.A., Lucci, A., Ivan, C., Calin G. A., and Kalluri, R. (2014) Cancer exosomes perform cell-independent microRNA biogenesis and promote tumorigenesis. Cancer Cell 26 (5), 707–721.

73. Michael NL, Vahe M T, d’Arcy L, Ehrenberg PK, Mosca JD, Rappaport J, Redfield RR. (1994) Negative-strand RNA transcripts are produced in human immunodeficiency virus type 1-infected cells and patients by a novel promoter downregulated by Tat. J. Virol 68:979–987.

74. Miller, R. H. (1998) Human immunodeficiency virus may encode a novel protein on the genomic DNA plus strand. Science 239, 1420–1422.

75. Nable, G and Baltimore, D. (1987) An inducible transcription factor activates expression of human immunodeficiency virus in T cells (published erratum appears in Nature (1990) 344 (6262), 178) Nature 326, 711–713.

76. Nardacci, R., Perfettini, J-L., Grieco, L., Thieffry, D., Kroemer G., and Piacentini, M. (2015) Syncytial apoptosis signaling network induced by the HIV-1 envelope glycoprotein complex: an overview. Cell Death and Disease 6,e1846;doi:10.1038/cddis.2015.204.

77. Oelrichs R, Tsykin A, Rhodes D, Solomon A, Ellett A, McPhee D, and Deacon N. (1998) Genomic sequence of HIV Type 1 from four members of the Sydney Blood Bank Cohort of long-term nonprogressors. AIDS Res and Human retroviruses 14 (9), 811–814.

78. Okada, H., Bakal, C., Shahinian, A., Elia, A., Wakeham, A., Suh, W-K., Duncan, G.S., Ciofani, M., Rottapel, R., Zuniga-Pflucker, J.C., and Mak T.W. (2004) Survivin loss in thymocytes triggers p53-mediated growth arrest and p53-independent cell death. J Exp Med (199)3: 399–410.

79. Omoto, S., Ito, M., Tsutsumi, Y., Ichikawa, Y., Okuyama, H., Brisibe, E.A., Saksena, N.K., and Fujii, Y.R. (2004) HIV-1 nef suppression by virally encoded microRNA. Retrovirology 1:44.

80. Ouellet, D.L., Vigneault-Edwards, J., Letourneau, K., Gobeil, L-A., Plante, I., Burnett, J.C., Rossi, J.J., and Provost, P. (2013) Regulation of host gene expression by HIV-1 TAR microRNAs. Retrovirology 10:86.

81. Ouellet, D.L., Plante, I., Landry, P., Barat, C.m Janelle, M.E., Flamand, L., Tremblay, M.J., and Provost, P. (2008) Identification of functional microRNAs released through asymmetrical processing of HIV-1 TAR element. Nucleic Acids Res. 36, 2353–2365.

82. Peden, K., Emerman, M., and Montagnier, L. (1991) Changes in growth properties on passage in tissue culture of viruses derived from infectious molecular clones of HIV-1Lai, HIV-1Mal, and HIV-1Eli. Virology 185 (2) 661–672.

83. Pfeffer, S., Sewer, A., Lagos-Quintana, M., Sheridan, R., Sander, C., Grasser, F.A., vanDyk, L.F., Ho, C.K., Shuman, S., Chien, M., Russo, J.J., Ju, J., Randall, G., Lindenbach, B.D., Rice, C.M., Simon, V., Ho, D.D., Zavolan, M., and Tuschl, T. (2005) Identification of microRNAs of the herpesvirus family. Nat Methods 2 (4): 269–276.

84. Rhodes, D., Solomon, A., Bolton, W., Wood, J., Sullivan, j., Learmont, J. and Deacon, N. (1999). Identification of a new recipient in the Sydney Blood Bank Cohort: a long-term HIV type 1-infected sero-indeterminate individual. AIDS Res. Hum. Retroviruses 15 (16), 1433–1439.

85. Salvi, R., Garbuglia, A.R., Di Caro, A., Pulciani, S., Montella, F., Benedetto, A. (1998) Grossly defective nef gene sequences in a human immunodeficiency virus type 1-seropositive long-term nonprogressor. J. Virol. 72, 3646–3657.

86. Samson, M., Libert, F., Doranz, B.J., Rucker, J., Liesnard, C., and Farber, C.M. (1996). Resistance to HIV-1 infection in Caucasian individuals bearing mutant alleles of the CCR5 chemokine receptor gene. Nature 382

87. Schopman, N.C., Willemsen, M., Liu, Y.P., Bradley, T., van Kampen, A., Baas, F., Berkhout, B., and Haasnoot, J. (2012) Deep sequencing of virus-infected cells reveals HIV-encoded small RNAs. Nucleic Acids Res. 40: 414–427.

88. Schwartz, L.M., and Osborne, B. A. (1993) Programmed cell death, apoptosis and killer genes. Immunology Today 14 (12) 582–590

89. Simon, V., Ho, D.D. (2003) HIV-1 dynamics in vivo: implications for therapy. Nat. Rev. Microbiol. 1, 181–190.

90. Valadi, H., Ekstrom, K., Bossios, A., Sjostrand, M., Lee, J. J., and Lotvall, J.O. (2007) Exosome-mediated transfer of mRNAs and microRNAs is a novel mechanism of genetic exchange between cells. Nature Cell Biology 9 (6), 654–659.

91. Wain-Hobson, S., Sonigo, P., Danos, O., Cole., S., and Alizon, M. (1985) Nucleotide sequence of the AIDS virus, LAV. Cell 40 (1), 9–17.

92. Wain-Hobson, S., Vartanian, J.P., Henry, M., Chenciner, N., Cheynier, R., Delassus, S., Martins, L.P., Sala, M., Nugeyre, M.T., Guetard, D., et. al. (1991) LAV revisited: origins of the early HIV-1 isolates from Institut Pasteur. Science 252 (5008) 961–965.

93. Whisnant, A.W., Bogerd, H.P., Flores, O., Ho, P., Powers, J.G., Sharova, N., Stevenson, M., Chen, C-H., and Cullen B.R. (2013) In-depth analysis of the interaction of HIV-1 with cellular microRNA biogenesis and effector mechanisms. mBio 4, 2:e00193–13.

94. Yeung, M.L., Bennasser, Y., Watashi, K., Le, S-Y., Houzet, L., and Jeang, K-T. (2009) Pyrosequencing of small non-coding RNAs in HIV-1 infected cells: evidence for the processing of a viral-cellular double-stranded RNA hybrid. Nucleic Acids Res. 37 (19); 6575–6586.

95. Yu, H., Rabson, A.B., Kaul, M., and Dougherty, J.P. (1996). Inducible human immunodeficiency virus type 1 packaging cell lines. J.Virol. 70, 4530–4537.

96. Zuker, M. (2003). Mfold web server for nucleic acid folding and hybridization prediction. Nucleic Acids Res. 31 (13), 3406–3415.

97. Zaunders J, Dyer WB, and Churchill M. (2011) The Sydney Blood Bank Cohort: implications for viral fitness as a cause of elite control.. Curr Opin HIV AIDS 6 (3), 151–156.

98. Zeng, Y., Yi, R., and Cullen, B. R. (2005) Recognition and cleavage of primary microRNA precursors by the nuclear processing enzyme Drosha. The EMBO J., 24: 138–148.

