## Supplemental Figure S1 for "MORT, a locus for apoptosis in the human immunodeficiency virus-type 1 antisense gene: implications for AIDS, Cancer, and Covid-19"

#### Supplemental Figure S1A

M 1 2 3 4 5 6 7 8 9 10 11 M 12

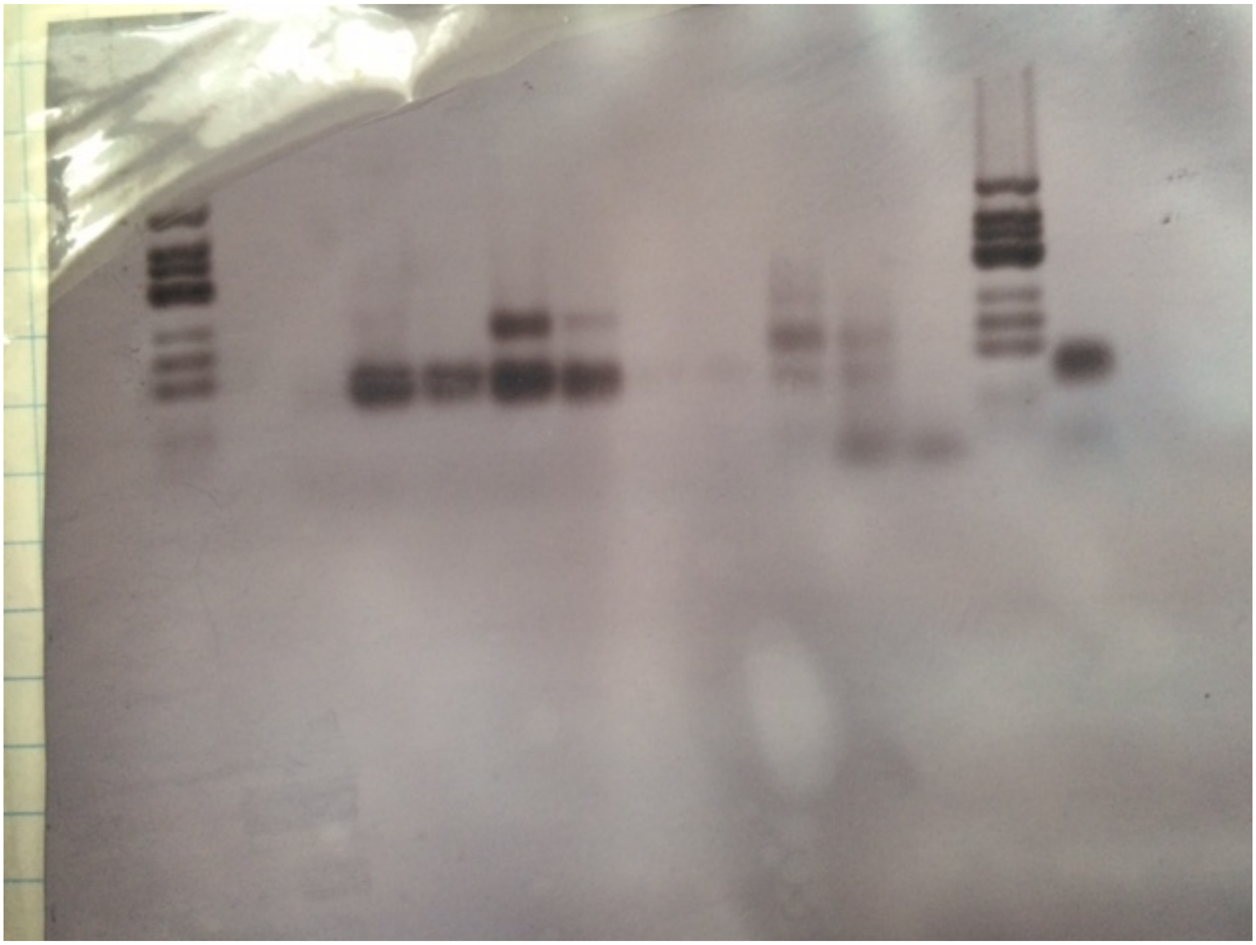

HL2/3----------------→ pr ←--IS + -→ M HL2/3

TRANSF: MOCK pHIV pHIV/wt^ HAP-FLAG

RNA ISOLATED 9/1999 10/1999

REVERSE TRANSCRIPTION (AVA1), THEN PCR
WITH AVA1 AND BIOTINYLATED-441 PRIMERS

### FIGURE S1B Strain differences in MORT hairpin

MORT 252-363
hairpin mutations

BRU – identical seq.
HXB2 o (4 mutations)
SF2 + (6 mutations)
RF # (8 mutations)
NL4-3 x 13mutations

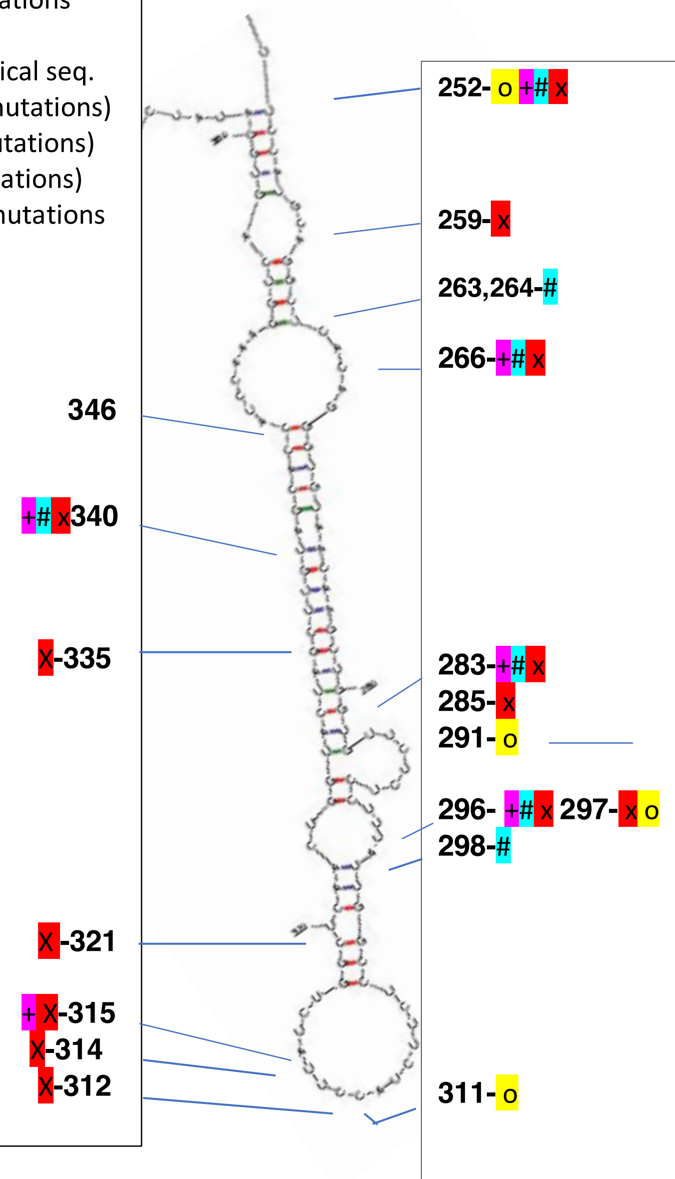

**Hap RNA sequence compared to NL4-3**

```
441
1 agaucugguc uaaccagaga gacccaguac aggcaaaaag cagcugcuua uaugcagcau   27 (Hap)
  agaucugguc uaaccagaga gacccaguac aggcaaaaag cagcugcuua uaugcagcau   pNL{3'ltr}
  agauguggcuc uaaccagaga gacccaguac aggcaaaaag cagcugcuua uauguagcau   5'ltr

61 cugagggcuc gccacucccc agucccgccc aggccacgcc ucccuggaaa gucccccagcg   27
   cugagggcuc gccacucccc agucccgccc aggccacgcc ucccuggaaa gucccccagcg   pNL{3'ltr}
   cugagggcuc gccacucccc agucccgccc aggccacacc ucccuggaaa gucccccagcg   5'ltr

121 gaaagucccu uguaacaagc ucgaugucaa caguucuuga aguacuccgg augcagcucu   27
    gaaagucccu uguagcaagc ucgaugucag caguucuuga aguacuccgg augcagcucu   pNL{3'ltr}
    gaaagucccu uguagaaagc ucgaugucag cagucuuugu aguacuccgg augcagcucu   5'ltr
Ava I
181 cgggccacgu gaugaaaugc uaggcggcug ucaaaccucc acucuaacac uucucucuca   27
    cgggccacgu gaugaaaugc uaggcggcug ucaaaccucc acucuaacac uucucucuca   pNL{3'ltr}
    cgggccaugu gacgaaaugc uaggaggcug ucaaacuucc acacuaauac uucucccucc   5'ltr

AvaII
241 gggucaucca uuccaugcag gcucacaggg uguaacaagc ugguguucuc uccuuuauug   27
    gggucaucca uuccaugcag gcucacaggg uguaacaagc ugguguucuc uccuuuauug   pNL{3'ltr}
    ggguccucca ucccaugcug gcucauaggg uguaacaagc uguucuucuc uccuuauuug   5'ltr
MORT hairpin

301 gccucuucua ccuuaucugg cucaacuggu acuagcuugu agcaccaucc aaaggucagu   27
    gccucuucua ccuuaucugg cucaacuggu acuagcuugu agcaccaucc aaaggucagu   pNL{3'ltr}
    gccucuucua cuugcucugg uucaacuggu acuaacuuga agcaccaucc aaaggucagu   5'ltr
MORT hairpin

361 ggauauc     27
    ggauaucu    pNL{3'ltr}
    ggauaucu    5'ltr
```

#### Figure S1C

**Primer extension of HIV antisense transcripts intrinsically made off HIVaINR in R region (LTR)**

Templates O, A, and B

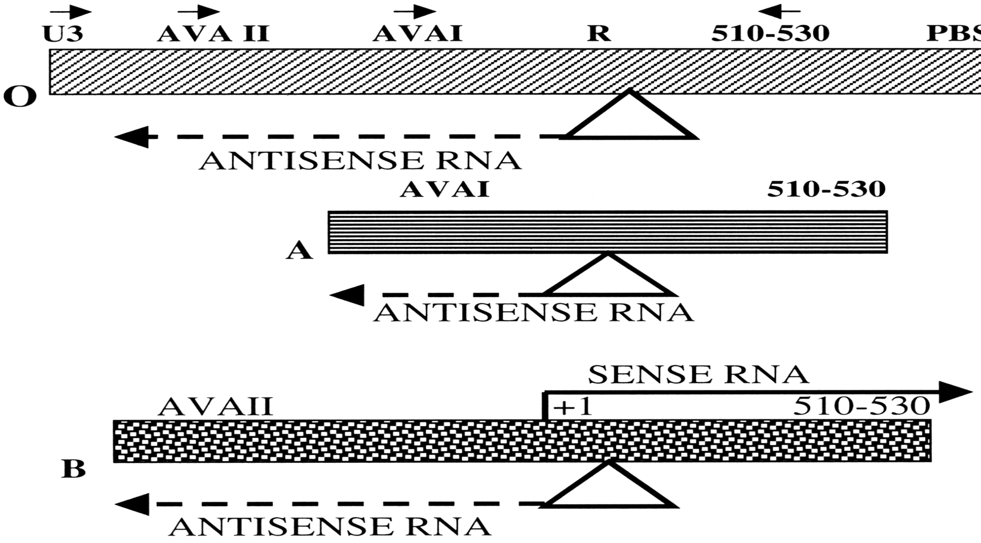


B.b B.a A.b A.a A.a (-) (-) M B.b B.a A.b A.a (-) (-) M A.a TEMPLATE (code above)
\+ + + + + + - + + + + + + NUCLEAR EXTRACT
\+ + (plus alpha-amanitin)

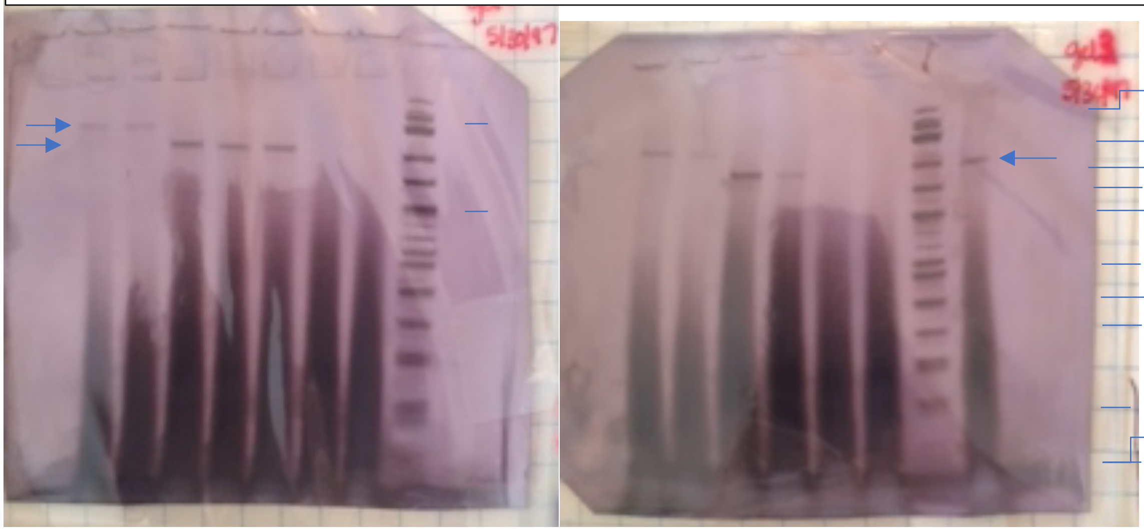

M
500.553
427,417,413
311
249
200
151
140, 118
100
82
66
48,42,40

Ava II...> <.Ava I ................> M Ava II.>Ava I.............> M Ava I B-PRIMERS used in Primer extension

Figure S1A (related to Figure 2)

RT-PCR shows that the HL2/3 cell line does not make the HIVaINR antisense RNA from its HXB3 LTR (lanes 1,2-mock transfected) whereas the pHIV-CAT transfected (lanes 3,4), and pHIV-CAT and pWT/^- transfected (lanes 5,6) HL2/3 cells clearly make HIVaINR antisense RNA from the transfected HIV-1 LTRs.

Lanes 1-6 cells were transfected and RNA isolated in a separate experiment from the transfection of HL2/3 with the HAP-FLAG recombinant vector on 10/1999; HAP-FLAG also makes the antisense RNA (CQ767337) from a CMV promoter (lane 12). Transfections were performed as previously described (Ludwig, et al, 2006). All RNAs were analyzed in the same RT-PCR experiment for HIVaINR antisense RNA using the Ava 1 primer (sense) 5'TTTCATCACGTGGCCCGAGAGC3' for the reverse transcription step, and Ava 1 and biotinylated-441 primers (5'CCAGAGAGACCCAGTACAGGCAAAA3') in the PCR step, as previously described (Ludwig, et al, 2006).

The colorimetric detection of the blot of the RT-PCR was as described in detail (Ludwig, et al, 1996, Ludwig, et al, 2006). Lane 7 received the primers alone control (pr) and lanes 9-11 received the internal standard control (IS+) for the RT-PCR experiment. Marker lanes received biotinylated phage X174-Hinf 1 fragments- the triplet of markers 200, 249, 311 show the HIV-1 antisense biotin-labeled RT-PCR products migrate at the expected size (¯183 bp) for both intrinsically made ( from HIV-1 LTR (lanes 3-6)) and pCMV-promoter (HAP-FLAG) (lane 12).

Figure S1B

Comparison of sequences demonstrates strain differences with mutations indicated between HIV strains in the MORT hairpin. Bru is identical to Hap, but NL4-3 has 13 mutations in the MORT region of the Hap gene, as shown. Many of these mutations would impair base-pairing in the MORT hairpin, as confirmed by Mfold analysis (not shown).

Figure S1C

Primer extension shows intrinsic antisense RNA transcripts initiating from HIVaINR

Transcription was studied using Drosophila extracts (nuclear extracts +/-, Promega) as a source of eukaryotic RNA polymerases and transcription factors, and variably-sized, U3R HIV-1 DNA templates (from (a) pHIV-CAT (BRU strain sequences) or (b) pNLgag (3'LTR NL43 strain sequences). Templates A and B are as diagrammed at top, and previously described (Ludwig, et al, 2006)). Samples receiving alpha-amanitin (Promega) are indicated (+).

RNA was then isolated (DNAse treated, and ethanol precipitated). Primer extension was then performed on purified RNA samples, in duplicate, using the indicated biotin-labeled primers (labeled Ava I or Ava II for antisense transcript detection; 510-530 for sense transcript detection). Complementary DNA products were analyzed following electrophoresis on precast, denaturing PAGE 6%TBE-urea gels (Novex), transferred to Biodyne B membranes (Pall), and colorimetric detection performed, as previously described (Ludwig, et al, 2006, Ludwig, et al 1996). The molecular weight marker was biotinylated ØX174/Hinf 1 fragments (M).

Primer extension demonstrated a single start site from the HIVaINR, as can be seen by a <u>single</u>, expected size, <u>biotin-labeled cDNA</u> band (arrow) when either biotinylated primers T7Ava I (217 nt ), or T7Ava II ( 278 nt ) were used in the primer extension, as indicated. The primers are 42 nt, and are indicated with sequence underlined in Figure S1B. The no template controls received no DNA (-) in the transcription reaction and had no specific bands, with (+) or without Drosophila nuclear extracts (-) as indicated above blots. While Drosophila nuclear extracts function in transcription reactions providing eukaryotic RNA polymerase II or III, a surprise finding was that alpha-amanitin (a potent inhibitor of RNA polymerase II) did not inhibit transcription in these experiments.
These experiments were performed in 1997; of interest, our (self-biotinylated) primers used in primer extension still allowed photography of blots in 2020 (Ludwig, et al, 1995).
