## Supplemental Figure S2 for "MORT, a locus for apoptosis in the human immunodeficiency virus-type 1 antisense gene: implications for AIDS, Cancer, and Covid-19"

**BLAST Results**

**Blast 2 sequences**

#### Job title: MORT271.299.VS.XIAP

| | |
|---|---|
| **RID** | 552BXBJB114 (Expires on 01-09 03:56 am) |
| **Query ID** | CQ767337.1 |
| **Description** | Sequence 27 from Patent EP1359221. |
| **Molecule type** | rna |
| **Query Length** | 28 |

| | |
|---|---|
| **Subject ID** | NG_007264.1 |
| **Description** | Homo sapiens X-linked inhibitor of apoptosis (XIAP), RefSeqGene (LRG_19) on chromosome X<br>See details |
| **Molecule type** | nucleic acid |
| **Subject Length** | 60775 |
| **Program** | BLASTN 2.7.1+ |

### Graphic Summary

**Distribution of the top 122 Blast Hits on 1 subject sequences**

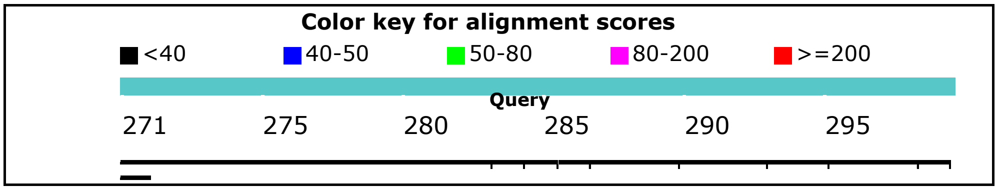

### Dot Matrix View

### Descriptions

Sequences producing significant alignments:

| Description | Max score | Total score | Query cover | E value | Ident | Accession |
|---|---|---|---|---|---|---|
| Homo sapiens X-linked inhibitor of apoptosis (XIAP), RefSeqGene (LRG_19) on chromosome X | 26.3 | 1866 | 100% | 0.014 | 100% | NG_007264.1 |

### Alignments

Homo sapiens X-linked inhibitor of apoptosis (XIAP), RefSeqGene (LRG_19) on chromosome X

Sequence ID: **NG_007264.1** Length: 60775 Number of Matches: 122

Range 1: 18784 to 18796

| Score | Expect | Identities | Gaps | Strand | Frame |
|---|---|---|---|---|---|
| 26.3 bits(13) | 0.014() | 13/13(100%) | 0/13(0%) | Plus/Plus | |

Features:

```
Query  271    TGTAACAAGCTGG  283
              |||||||||||||
Sbjct  18784  TGTAACAAGCTGG  18796
```

Range 2: 21827 to 21837

| Score | Expect | Identities | Gaps | Strand | Frame |
|---|---|---|---|---|---|
| 22.3 bits(11) | 0.22() | 11/11(100%) | 0/11(0%) | Plus/Minus | |

Features:

```
Query  287    TCTCTCCTTTA  297
              |||||||||||
Sbjct  21837  TCTCTCCTTTA  21827
```

Range 3: 42321 to 42331

| Score | Expect | Identities | Gaps | Strand | Frame |
|---|---|---|---|---|---|
| 22.3 bits(11) | 0.22() | 11/11(100%) | 0/11(0%) | Plus/Minus | |

Features:

```
Query  288    CTCTCCTTTAT  298
              |||||||||||
Sbjct  42331  CTCTCCTTTAT  42321
```

Range 4: 46200 to 46210

| Score | Expect | Identities | Gaps | Strand | Frame |
|---|---|---|---|---|---|
| 22.3 bits(11) | 0.22() | 11/11(100%) | 0/11(0%) | Plus/Plus | |

Features:

```
Query  287    TCTCTCCTTTA  297
              |||||||||||
Sbjct  46200  TCTCTCCTTTA  46210
```

Range 5: 15699 to 15707

| Score | Expect | Identities | Gaps | Strand | Frame |
|---|---|---|---|---|---|
| 18.3 bits(9) | 3.5() | 9/9(100%) | 0/9(0%) | Plus/Minus | |

Features:

```
Query  276    CAAGCTGGT  284
              |||||||||
Sbjct  15707  CAAGCTGGT  15699
```

Range 6: 36970 to 36978

| Score | Expect | Identities | Gaps | Strand | Frame |
|---|---|---|---|---|---|
| 18.3 bits(9) | 3.5() | 9/9(100%) | 0/9(0%) | Plus/Minus | |

Features:

```
Query  276    CAAGCTGGT  284
              |||||||||
Sbjct  36978  CAAGCTGGT  36970
```

Range 7: 41701 to 41709

| Score | Expect | Identities | Gaps | Strand | Frame |
|---|---|---|---|---|---|
| 18.3 bits(9) | 3.5() | 9/9(100%) | 0/9(0%) | Plus/Plus | |

Features:

```
Query  281    TGGTGTTCT  289
              |||||||||
Sbjct  41701  TGGTGTTCT  41709
```

Range 8: 50139 to 50147

| Score | Expect | Identities | Gaps | Strand | Frame |
|---|---|---|---|---|---|
| 18.3 bits(9) | 3.5() | 9/9(100%) | 0/9(0%) | Plus/Plus | |

Features:

```
Query  287    TCTCTCCTT  295
              |||||||||
Sbjct  50139  TCTCTCCTT  50147
```

Range 9: 53478 to 53486

| Score | Expect | Identities | Gaps | Strand | Frame |
|---|---|---|---|---|---|
| 18.3 bits(9) | 3.5() | 9/9(100%) | 0/9(0%) | Plus/Plus | |

Features:

```
Query  284    TGTTCTCTC  292
              |||||||||
Sbjct  53478  TGTTCTCTC  53486
```

Range 10: 58481 to 58489

| Score | Expect | Identities | Gaps | Strand | Frame |
|---|---|---|---|---|---|
| 18.3 bits(9) | 3.5() | 9/9(100%) | 0/9(0%) | Plus/Plus | |

Features:

```
Query  271    TGTAACAAG  279
              |||||||||
Sbjct  58481  TGTAACAAG  58489
```

Range 11: 106 to 113

| Score | Expect | Identities | Gaps | Strand | Frame |
|---|---|---|---|---|---|
| 16.4 bits(8) | 14() | 8/8(100%) | 0/8(0%) | Plus/Plus | |

Features:

```
Query  276  CAAGCTGG  283
            ||||||||
Sbjct  106  CAAGCTGG  113
```

Range 12: 424 to 431

| Score | Expect | Identities | Gaps | Strand | Frame |
|---|---|---|---|---|---|
| 16.4 bits(8) | 14() | 8/8(100%) | 0/8(0%) | Plus/Plus | |

Features:

```
Query  289  TCTCCTTT  296
            ||||||||
Sbjct  424  TCTCCTTT  431
```

Range 13: 929 to 936

| Score | Expect | Identities | Gaps | Strand | Frame |
|---|---|---|---|---|---|
| 16.4 bits(8) | 14() | 8/8(100%) | 0/8(0%) | Plus/Minus | |

Features:

```
Query  289  TCTCCTTT  296
            ||||||||
Sbjct  936  TCTCCTTT  929
```

Range 14: 1609 to 1616

| Score | Expect | Identities | Gaps | Strand | Frame |
|---|---|---|---|---|---|
| 16.4 bits(8) | 14() | 8/8(100%) | 0/8(0%) | Plus/Plus | |

Features:

```
Query  271   TGTAACAA  278
             ||||||||
Sbjct  1609  TGTAACAA  1616
```

Range 15: 2608 to 2615

| Score | Expect | Identities | Gaps | Strand | Frame |
|---|---|---|---|---|---|
| 16.4 bits(8) | 14() | 8/8(100%) | 0/8(0%) | Plus/Minus | |

Features:

```
Query  287   TCTCTCCT  294
             ||||||||
Sbjct  2615  TCTCTCCT  2608
```

Range 16: 5104 to 5111

| Score | Expect | Identities | Gaps | Strand | Frame |
|---|---|---|---|---|---|
| 16.4 bits(8) | 14() | 8/8(100%) | 0/8(0%) | Plus/Minus | |

Features:

```
Query  275   ACAAGCTG  282
             ||||||||
Sbjct  5111  ACAAGCTG  5104
```

Range 17: 5506 to 5513

| Score | Expect | Identities | Gaps | Strand | Frame |
|---|---|---|---|---|---|
| 16.4 bits(8) | 14() | 8/8(100%) | 0/8(0%) | Plus/Plus | |

Features:

```
Query  278   AGCTGGTG  285
             ||||||||
```

```
Sbjct  5506  AGCTGGTG  5513
```

Range 18: 6670 to 6677

| Score | Expect | Identities | Gaps | Strand | Frame |
|---|---|---|---|---|---|
| 16.4 bits(8) | 14() | 8/8(100%) | 0/8(0%) | Plus/Plus | |

Features:

```
Query  287   TCTCTCCT  294
             ||||||||
Sbjct  6670  TCTCTCCT  6677
```

Range 19: 9000 to 9007

| Score | Expect | Identities | Gaps | Strand | Frame |
|---|---|---|---|---|---|
| 16.4 bits(8) | 14() | 8/8(100%) | 0/8(0%) | Plus/Plus | |

Features:

```
Query  289   TCTCCTTT  296
             ||||||||
Sbjct  9000  TCTCCTTT  9007
```

Range 20: 13503 to 13510

| Score | Expect | Identities | Gaps | Strand | Frame |
|---|---|---|---|---|---|
| 16.4 bits(8) | 14() | 8/8(100%) | 0/8(0%) | Plus/Minus | |

Features:

```
Query  276    CAAGCTGG  283
              ||||||||
Sbjct  13510  CAAGCTGG  13503
```

Range 21: 13615 to 13622

| Score | Expect | Identities | Gaps | Strand | Frame |
|---|---|---|---|---|---|
| 16.4 bits(8) | 14() | 8/8(100%) | 0/8(0%) | Plus/Plus | |

Features:

```
Query  291    TCCTTTAT  298
              ||||||||
Sbjct  13615  TCCTTTAT  13622
```

Range 22: 19231 to 19238

| Score | Expect | Identities | Gaps | Strand | Frame |
|---|---|---|---|---|---|
| 16.4 bits(8) | 14() | 8/8(100%) | 0/8(0%) | Plus/Minus | |

Features:

```
Query  277    AAGCTGGT  284
              ||||||||
Sbjct  19238  AAGCTGGT  19231
```

Range 23: 22374 to 22381

| Score | Expect | Identities | Gaps | Strand | Frame |
|---|---|---|---|---|---|
| 16.4 bits(8) | 14() | 8/8(100%) | 0/8(0%) | Plus/Minus | |

Features:

```
Query  271    TGTAACAA  278
              ||||||||
Sbjct  22381  TGTAACAA  22374
```

Range 24: 24609 to 24616

| Score | Expect | Identities | Gaps | Strand | Frame |
|---|---|---|---|---|---|
| 16.4 bits(8) | 14() | 8/8(100%) | 0/8(0%) | Plus/Plus | |

Features:

```
Query  292    CCTTTATT  299
              ||||||||
Sbjct  24609  CCTTTATT  24616
```

Range 25: 25675 to 25682

| Score | Expect | Identities | Gaps | Strand | Frame |
|---|---|---|---|---|---|
| 16.4 bits(8) | 14() | 8/8(100%) | 0/8(0%) | Plus/Plus | |

Features:

```
Query  288    CTCTCCTT  295
              ||||||||
Sbjct  25675  CTCTCCTT  25682
```

Range 26: 28895 to 28902

| Score | Expect | Identities | Gaps | Strand | Frame |
|---|---|---|---|---|---|
| 16.4 bits(8) | 14() | 8/8(100%) | 0/8(0%) | Plus/Minus | |

Features:

```
Query  276    CAAGCTGG  283
              ||||||||
Sbjct  28902  CAAGCTGG  28895
```

Range 27: 30420 to 30427

| Score | Expect | Identities | Gaps | Strand | Frame |
|---|---|---|---|---|---|
| 16.4 bits(8) | 14() | 8/8(100%) | 0/8(0%) | Plus/Plus | |

Features:

```
Query  284    TGTTCTCT  291
              ||||||||
Sbjct  30420  TGTTCTCT  30427
```

Range 28: 37546 to 37557

| Score | Expect | Identities | Gaps | Strand | Frame |
|---|---|---|---|---|---|
| 16.4 bits(8) | 14() | 11/12(92%) | 0/12(0%) | Plus/Minus | |

Features:

```
Query  281    TGGTGTTCTCTC  292
              ||||||| ||||
Sbjct  37557  TGGTGTTTTCTC  37546
```

Range 29: 37585 to 37592

| Score | Expect | Identities | Gaps | Strand | Frame |
|---|---|---|---|---|---|
| 16.4 bits(8) | 14() | 8/8(100%) | 0/8(0%) | Plus/Minus | |

Features:

```
Query  273    TAACAAGC  280
              ||||||||
Sbjct  37592  TAACAAGC  37585
```

Range 30: 40556 to 40563

| Score | Expect | Identities | Gaps | Strand | Frame |
|---|---|---|---|---|---|

| | | | | |
|---|---|---|---|---|
| 16.4 bits(8) | 14() | 8/8(100%) | 0/8(0%) | Plus/Plus |

Features:

```
Query  289    TCTCCTTT  296
              ||||||||
Sbjct  40556  TCTCCTTT  40563
```

Range 31: 44724 to 44735

| Score | Expect | Identities | Gaps | Strand | Frame |
|---|---|---|---|---|---|
| 16.4 bits(8) | 14() | 11/12(92%) | 0/12(0%) | Plus/Plus | |

Features:

```
Query  283    GTGTTCTCTCCT  294
              |||| |||||||
Sbjct  44724  GTGTCCTCTCCT  44735
```

Range 32: 47102 to 47109

| Score | Expect | Identities | Gaps | Strand | Frame |
|---|---|---|---|---|---|
| 16.4 bits(8) | 14() | 8/8(100%) | 0/8(0%) | Plus/Minus | |

Features:

```
Query  288    CTCTCCTT  295
              ||||||||
Sbjct  47109  CTCTCCTT  47102
```

Range 33: 47481 to 47488

| Score | Expect | Identities | Gaps | Strand | Frame |
|---|---|---|---|---|---|
| 16.4 bits(8) | 14() | 8/8(100%) | 0/8(0%) | Plus/Plus | |

Features:

```
Query  292    CCTTTATT  299
              ||||||||
Sbjct  47481  CCTTTATT  47488
```

Range 34: 47694 to 47701

| Score | Expect | Identities | Gaps | Strand | Frame |
|---|---|---|---|---|---|
| 16.4 bits(8) | 14() | 8/8(100%) | 0/8(0%) | Plus/Plus | |

Features:

```
Query  292    CCTTTATT  299
              ||||||||
Sbjct  47694  CCTTTATT  47701
```

Range 35: 49101 to 49108

| Score | Expect | Identities | Gaps | Strand | Frame |
|---|---|---|---|---|---|
| 16.4 bits(8) | 14() | 8/8(100%) | 0/8(0%) | Plus/Plus | |

Features:

```
Query  289    TCTCCTTT  296
              ||||||||
Sbjct  49101  TCTCCTTT  49108
```

Range 36: 50018 to 50025

| Score | Expect | Identities | Gaps | Strand | Frame |
|---|---|---|---|---|---|
| 16.4 bits(8) | 14() | 8/8(100%) | 0/8(0%) | Plus/Plus | |

Features:

```
Query  290    CTCCTTTA  297
              ||||||||
Sbjct  50018  CTCCTTTA  50025
```

Range 37: 50150 to 50157

| Score | Expect | Identities | Gaps | Strand | Frame |
|---|---|---|---|---|---|
| 16.4 bits(8) | 14() | 8/8(100%) | 0/8(0%) | Plus/Minus | |

Features:

```
Query  274    AACAAGCT  281
              ||||||||
Sbjct  50157  AACAAGCT  50150
```

Range 38: 348 to 354

| Score | Expect | Identities | Gaps | Strand | Frame |
|---|---|---|---|---|---|
| 14.4 bits(7) | 55() | 7/7(100%) | 0/7(0%) | Plus/Plus | |

Features:

```
Query  291  TCCTTTA  297
            |||||||
Sbjct  348  TCCTTTA  354
```

Range 39: 1423 to 1429

| Score | Expect | Identities | Gaps | Strand | Frame |
|---|---|---|---|---|---|
| 14.4 bits(7) | 55() | 7/7(100%) | 0/7(0%) | Plus/Plus | |

Features:

```
Query  272   GTAACAA  278
             |||||||
Sbjct  1423  GTAACAA  1429
```

Range 40: 1805 to 1811

| Score | Expect | Identities | Gaps | Strand | Frame |
|---|---|---|---|---|---|
| 14.4 bits(7) | 55() | 7/7(100%) | 0/7(0%) | Plus/Minus | |

Features:

```
Query  288   CTCTCCT  294
             |||||||
Sbjct  1811  CTCTCCT  1805
```

Range 41: 1841 to 1847

| Score | Expect | Identities | Gaps | Strand | Frame |
|---|---|---|---|---|---|
| 14.4 bits(7) | 55() | 7/7(100%) | 0/7(0%) | Plus/Plus | |

Features:

```
Query  287   TCTCTCC  293
             |||||||
Sbjct  1841  TCTCTCC  1847
```

Range 42: 1943 to 1949

| Score | Expect | Identities | Gaps | Strand | Frame |
|---|---|---|---|---|---|
| 14.4 bits(7) | 55() | 7/7(100%) | 0/7(0%) | Plus/Minus | |

Features:

```
Query  281   TGGTGTT  287
             |||||||
Sbjct  1949  TGGTGTT  1943
```

Range 43: 2144 to 2150

| Score | Expect | Identities | Gaps | Strand | Frame |
|---|---|---|---|---|---|
| 14.4 bits(7) | 55() | 7/7(100%) | 0/7(0%) | Plus/Minus | |

Features:

```
Query  285   GTTCTCT  291
             |||||||
Sbjct  2150  GTTCTCT  2144
```

Range 44: 3559 to 3565

| Score | Expect | Identities | Gaps | Strand | Frame |
|---|---|---|---|---|---|
| 14.4 bits(7) | 55() | 7/7(100%) | 0/7(0%) | Plus/Minus | |

Features:

```
Query  287   TCTCTCC  293
             |||||||
Sbjct  3565  TCTCTCC  3559
```

Range 45: 3826 to 3832

| Score | Expect | Identities | Gaps | Strand | Frame |
|---|---|---|---|---|---|
| 14.4 bits(7) | 55() | 7/7(100%) | 0/7(0%) | Plus/Plus | |

Features:

```
Query  288   CTCTCCT  294
             |||||||
Sbjct  3826  CTCTCCT  3832
```

Range 46: 4316 to 4322

| Score | Expect | Identities | Gaps | Strand | Frame |
|---|---|---|---|---|---|
| 14.4 bits(7) | 55() | 7/7(100%) | 0/7(0%) | Plus/Minus | |

Features:

```
Query  283   GTGTTCT  289
             |||||||
Sbjct  4322  GTGTTCT  4316
```

Range 47: 5147 to 5153

| Score | Expect | Identities | Gaps | Strand | Frame |
|---|---|---|---|---|---|
| 14.4 bits(7) | 55() | 7/7(100%) | 0/7(0%) | Plus/Plus | |

Features:

```
Query  288   CTCTCCT  294
             |||||||
Sbjct  5147  CTCTCCT  5153
```

Range 48: 5899 to 5905

| Score | Expect | Identities | Gaps | Strand | Frame |
|---|---|---|---|---|---|
| 14.4 bits(7) | 55() | 7/7(100%) | 0/7(0%) | Plus/Minus | |

Features:

```
Query  271   TGTAACA  277
             |||||||
Sbjct  5905  TGTAACA  5899
```

Range 49: 6857 to 6863

| Score | Expect | Identities | Gaps | Strand | Frame |
|---|---|---|---|---|---|

| | | | | |
|---|---|---|---|---|
| 14.4 bits(7) | 55() | 7/7(100%) | 0/7(0%) | Plus/Plus |

Features:

```
Query  281   TGGTGTT  287
             |||||||
Sbjct  6857  TGGTGTT  6863
```

Range 50: 7540 to 7546

| Score | Expect | Identities | Gaps | Strand | Frame |
|---|---|---|---|---|---|
| 14.4 bits(7) | 55() | 7/7(100%) | 0/7(0%) | Plus/Minus | |

Features:

```
Query  278   AGCTGGT  284
             |||||||
Sbjct  7546  AGCTGGT  7540
```

Range 51: 8918 to 8924

| Score | Expect | Identities | Gaps | Strand | Frame |
|---|---|---|---|---|---|
| 14.4 bits(7) | 55() | 7/7(100%) | 0/7(0%) | Plus/Minus | |

Features:

```
Query  278   AGCTGGT  284
             |||||||
Sbjct  8924  AGCTGGT  8918
```

Range 52: 8997 to 9003

| Score | Expect | Identities | Gaps | Strand | Frame |
|---|---|---|---|---|---|
| 14.4 bits(7) | 55() | 7/7(100%) | 0/7(0%) | Plus/Plus | |

Features:

```
Query  284   TGTTCTC  290
             |||||||
Sbjct  8997  TGTTCTC  9003
```

Range 53: 9775 to 9781

| Score | Expect | Identities | Gaps | Strand | Frame |
|---|---|---|---|---|---|
| 14.4 bits(7) | 55() | 7/7(100%) | 0/7(0%) | Plus/Minus | |

Features:

```
Query  280   CTGGTGT  286
             |||||||
Sbjct  9781  CTGGTGT  9775
```

Range 54: 9973 to 9979

| Score | Expect | Identities | Gaps | Strand | Frame |
|---|---|---|---|---|---|
| 14.4 bits(7) | 55() | 7/7(100%) | 0/7(0%) | Plus/Minus | |

Features:

```
Query  291   TCCTTTA  297
             |||||||
Sbjct  9979  TCCTTTA  9973
```

Range 55: 13107 to 13113

| Score | Expect | Identities | Gaps | Strand | Frame |
|---|---|---|---|---|---|
| 14.4 bits(7) | 55() | 7/7(100%) | 0/7(0%) | Plus/Plus | |

Features:

```
Query  274    AACAAGC  280
              |||||||
Sbjct  13107  AACAAGC  13113
```

Range 56: 13781 to 13787

| Score | Expect | Identities | Gaps | Strand | Frame |
|---|---|---|---|---|---|
| 14.4 bits(7) | 55() | 7/7(100%) | 0/7(0%) | Plus/Minus | |

Features:

```
Query  273    TAACAAG  279
              |||||||
Sbjct  13787  TAACAAG  13781
```

Range 57: 14186 to 14192

| Score | Expect | Identities | Gaps | Strand | Frame |
|---|---|---|---|---|---|
| 14.4 bits(7) | 55() | 7/7(100%) | 0/7(0%) | Plus/Plus | |

Features:

```
Query  277    AAGCTGG  283
              |||||||
Sbjct  14186  AAGCTGG  14192
```

Range 58: 14330 to 14336

| Score | Expect | Identities | Gaps | Strand | Frame |
|---|---|---|---|---|---|
| 14.4 bits(7) | 55() | 7/7(100%) | 0/7(0%) | Plus/Minus | |

Features:

```
Query  293    CTTTATT  299
              |||||||
Sbjct  14336  CTTTATT  14330
```

Range 59: 15235 to 15241

| Score | Expect | Identities | Gaps | Strand | Frame |
|---|---|---|---|---|---|
| 14.4 bits(7) | 55() | 7/7(100%) | 0/7(0%) | Plus/Plus | |

Features:

```
Query  291    TCCTTTA  297
              |||||||
Sbjct  15235  TCCTTTA  15241
```

Range 60: 17290 to 17296

| Score | Expect | Identities | Gaps | Strand | Frame |
|---|---|---|---|---|---|
| 14.4 bits(7) | 55() | 7/7(100%) | 0/7(0%) | Plus/Minus | |

Features:

```
Query  284    TGTTCTC  290
              |||||||
Sbjct  17296  TGTTCTC  17290
```

Range 61: 22244 to 22250

| Score | Expect | Identities | Gaps | Strand | Frame |
|---|---|---|---|---|---|
| 14.4 bits(7) | 55() | 7/7(100%) | 0/7(0%) | Plus/Minus | |

Features:

```
Query  281    TGGTGTT  287
              |||||||
Sbjct  22250  TGGTGTT  22244
```

Range 62: 22808 to 22814

| Score | Expect | Identities | Gaps | Strand | Frame |
|---|---|---|---|---|---|
| 14.4 bits(7) | 55() | 7/7(100%) | 0/7(0%) | Plus/Minus | |

Features:

```
Query  291    TCCTTTA  297
              |||||||
Sbjct  22814  TCCTTTA  22808
```

Range 63: 23038 to 23044

| Score | Expect | Identities | Gaps | Strand | Frame |
|---|---|---|---|---|---|
| 14.4 bits(7) | 55() | 7/7(100%) | 0/7(0%) | Plus/Plus | |

Features:

```
Query  293    CTTTATT  299
              |||||||
Sbjct  23038  CTTTATT  23044
```

Range 64: 23283 to 23289

| Score | Expect | Identities | Gaps | Strand | Frame |
|---|---|---|---|---|---|
| 14.4 bits(7) | 55() | 7/7(100%) | 0/7(0%) | Plus/Plus | |

Features:

```
Query  284    TGTTCTC  290
              |||||||
Sbjct  23283  TGTTCTC  23289
```

Range 65: 23907 to 23913

| Score | Expect | Identities | Gaps | Strand | Frame |
|---|---|---|---|---|---|
| 14.4 bits(7) | 55() | 7/7(100%) | 0/7(0%) | Plus/Plus | |

Features:

```
Query  274    AACAAGC  280
              |||||||
Sbjct  23907  AACAAGC  23913
```

Range 66: 24016 to 24022

| Score | Expect | Identities | Gaps | Strand | Frame |
|---|---|---|---|---|---|
| 14.4 bits(7) | 55() | 7/7(100%) | 0/7(0%) | Plus/Minus | |

Features:

```
Query  284    TGTTCTC  290
              |||||||
Sbjct  24022  TGTTCTC  24016
```

Range 67: 25592 to 25598

| Score | Expect | Identities | Gaps | Strand | Frame |
|---|---|---|---|---|---|
| 14.4 bits(7) | 55() | 7/7(100%) | 0/7(0%) | Plus/Minus | |

Features:

```
Query  280    CTGGTGT  286
              |||||||
Sbjct  25598  CTGGTGT  25592
```

Range 68: 25898 to 25904

| Score | Expect | Identities | Gaps | Strand | Frame |
|---|---|---|---|---|---|

| Score | Expect | Identities | Gaps | Strand | Frame |
|---|---|---|---|---|---|
| 14.4 bits(7) | 55() | 7/7(100%) | 0/7(0%) | Plus/Plus | |

Features:

```
Query  292    CCTTTAT  298
              |||||||
Sbjct  25898  CCTTTAT  25904
```

Range 69: 26017 to 26023

| Score | Expect | Identities | Gaps | Strand | Frame |
|---|---|---|---|---|---|
| 14.4 bits(7) | 55() | 7/7(100%) | 0/7(0%) | Plus/Plus | |

Features:

```
Query  293    CTTTATT  299
              |||||||
Sbjct  26017  CTTTATT  26023
```

Range 70: 27007 to 27013

| Score | Expect | Identities | Gaps | Strand | Frame |
|---|---|---|---|---|---|
| 14.4 bits(7) | 55() | 7/7(100%) | 0/7(0%) | Plus/Plus | |

Features:

```
Query  291    TCCTTTA  297
              |||||||
Sbjct  27007  TCCTTTA  27013
```

Range 71: 28959 to 28965

| Score | Expect | Identities | Gaps | Strand | Frame |
|---|---|---|---|---|---|
| 14.4 bits(7) | 55() | 7/7(100%) | 0/7(0%) | Plus/Minus | |

Features:

```
Query  292    CCTTTAT  298
              |||||||
Sbjct  28965  CCTTTAT  28959
```

Range 72: 29336 to 29342

| Score | Expect | Identities | Gaps | Strand | Frame |
|---|---|---|---|---|---|
| 14.4 bits(7) | 55() | 7/7(100%) | 0/7(0%) | Plus/Minus | |

Features:

```
Query  272    GTAACAA  278
              |||||||
Sbjct  29342  GTAACAA  29336
```

Range 73: 29623 to 29629

| Score | Expect | Identities | Gaps | Strand | Frame |
|---|---|---|---|---|---|
| 14.4 bits(7) | 55() | 7/7(100%) | 0/7(0%) | Plus/Minus | |

Features:

```
Query  278    AGCTGGT  284
              |||||||
Sbjct  29629  AGCTGGT  29623
```

Range 74: 29685 to 29691

| Score | Expect | Identities | Gaps | Strand | Frame |
|---|---|---|---|---|---|
| 14.4 bits(7) | 55() | 7/7(100%) | 0/7(0%) | Plus/Minus | |

Features:

```
Query  272    GTAACAA  278
              |||||||
Sbjct  29691  GTAACAA  29685
```

Range 75: 29822 to 29828

| Score | Expect | Identities | Gaps | Strand | Frame |
|---|---|---|---|---|---|
| 14.4 bits(7) | 55() | 7/7(100%) | 0/7(0%) | Plus/Minus | |

Features:

```
Query  271    TGTAACA  277
              |||||||
Sbjct  29828  TGTAACA  29822
```

Range 76: 29910 to 29916

| Score | Expect | Identities | Gaps | Strand | Frame |
|---|---|---|---|---|---|
| 14.4 bits(7) | 55() | 7/7(100%) | 0/7(0%) | Plus/Minus | |

Features:

```
Query  283    GTGTTCT  289
              |||||||
Sbjct  29916  GTGTTCT  29910
```

Range 77: 30635 to 30641

| Score | Expect | Identities | Gaps | Strand | Frame |
|---|---|---|---|---|---|
| 14.4 bits(7) | 55() | 7/7(100%) | 0/7(0%) | Plus/Minus | |

Features:

```
Query  289    TCTCCTT  295
              |||||||
Sbjct  30641  TCTCCTT  30635
```

Range 78: 31119 to 31125

| Score | Expect | Identities | Gaps | Strand | Frame |
|---|---|---|---|---|---|
| 14.4 bits(7) | 55() | 7/7(100%) | 0/7(0%) | Plus/Minus | |

Features:

```
Query  283    GTGTTCT  289
              |||||||
Sbjct  31125  GTGTTCT  31119
```

Range 79: 31302 to 31308

| Score | Expect | Identities | Gaps | Strand | Frame |
|---|---|---|---|---|---|
| 14.4 bits(7) | 55() | 7/7(100%) | 0/7(0%) | Plus/Plus | |

Features:

```
Query  273    TAACAAG  279
              |||||||
Sbjct  31302  TAACAAG  31308
```

Range 80: 31312 to 31318

| Score | Expect | Identities | Gaps | Strand | Frame |
|---|---|---|---|---|---|
| 14.4 bits(7) | 55() | 7/7(100%) | 0/7(0%) | Plus/Minus | |

Features:

```
Query  276    CAAGCTG  282
              |||||||
Sbjct  31318  CAAGCTG  31312
```

Range 81: 31348 to 31354

| Score | Expect | Identities | Gaps | Strand | Frame |
|---|---|---|---|---|---|
| 14.4 bits(7) | 55() | 7/7(100%) | 0/7(0%) | Plus/Plus | |

Features:

```
Query  293    CTTTATT  299
              |||||||
Sbjct  31348  CTTTATT  31354
```

Range 82: 31756 to 31762

| Score | Expect | Identities | Gaps | Strand | Frame |
|---|---|---|---|---|---|
| 14.4 bits(7) | 55() | 7/7(100%) | 0/7(0%) | Plus/Minus | |

Features:

```
Query  273    TAACAAG  279
              |||||||
Sbjct  31762  TAACAAG  31756
```

Range 83: 31915 to 31921

| Score | Expect | Identities | Gaps | Strand | Frame |
|---|---|---|---|---|---|
| 14.4 bits(7) | 55() | 7/7(100%) | 0/7(0%) | Plus/Plus | |

Features:

```
Query  287    TCTCTCC  293
              |||||||
Sbjct  31915  TCTCTCC  31921
```

Range 84: 32846 to 32852

| Score | Expect | Identities | Gaps | Strand | Frame |
|---|---|---|---|---|---|
| 14.4 bits(7) | 55() | 7/7(100%) | 0/7(0%) | Plus/Minus | |

Features:

```
Query  293    CTTTATT  299
              |||||||
Sbjct  32852  CTTTATT  32846
```

Range 85: 33153 to 33159

| Score | Expect | Identities | Gaps | Strand | Frame |
|---|---|---|---|---|---|
| 14.4 bits(7) | 55() | 7/7(100%) | 0/7(0%) | Plus/Plus | |

Features:

```
Query  272    GTAACAA  278
              |||||||
Sbjct  33153  GTAACAA  33159
```

Range 86: 34720 to 34726

| Score | Expect | Identities | Gaps | Strand | Frame |
|---|---|---|---|---|---|
| 14.4 bits(7) | 55() | 7/7(100%) | 0/7(0%) | Plus/Minus | |

Features:

```
Query  289    TCTCCTT  295
              |||||||
Sbjct  34726  TCTCCTT  34720
```

Range 87: 37428 to 37434

| Score | Expect | Identities | Gaps | Strand | Frame |
|---|---|---|---|---|---|

| | | | | |
|---|---|---|---|---|
| 14.4 bits(7) | 55() | 7/7(100%) | 0/7(0%) | Plus/Minus |

Features:

```
Query  293    CTTTATT  299
              |||||||
Sbjct  37434  CTTTATT  37428
```

Range 88: 37526 to 37532

| Score | Expect | Identities | Gaps | Strand | Frame |
|---|---|---|---|---|---|
| 14.4 bits(7) | 55() | 7/7(100%) | 0/7(0%) | Plus/Minus | |

Features:

```
Query  271    TGTAACA  277
              |||||||
Sbjct  37532  TGTAACA  37526
```

Range 89: 38917 to 38923

| Score | Expect | Identities | Gaps | Strand | Frame |
|---|---|---|---|---|---|
| 14.4 bits(7) | 55() | 7/7(100%) | 0/7(0%) | Plus/Minus | |

Features:

```
Query  277    AAGCTGG  283
              |||||||
Sbjct  38923  AAGCTGG  38917
```

Range 90: 39552 to 39558

| Score | Expect | Identities | Gaps | Strand | Frame |
|---|---|---|---|---|---|
| 14.4 bits(7) | 55() | 7/7(100%) | 0/7(0%) | Plus/Minus | |

Features:

```
Query  290    CTCCTTT  296
              |||||||
Sbjct  39558  CTCCTTT  39552
```

Range 91: 39881 to 39887

| Score | Expect | Identities | Gaps | Strand | Frame |
|---|---|---|---|---|---|
| 14.4 bits(7) | 55() | 7/7(100%) | 0/7(0%) | Plus/Minus | |

Features:

```
Query  283    GTGTTCT  289
              |||||||
Sbjct  39887  GTGTTCT  39881
```

Range 92: 40165 to 40171

| Score | Expect | Identities | Gaps | Strand | Frame |
|---|---|---|---|---|---|
| 14.4 bits(7) | 55() | 7/7(100%) | 0/7(0%) | Plus/Minus | |

Features:

```
Query  274    AACAAGC  280
              |||||||
Sbjct  40171  AACAAGC  40165
```

Range 93: 40250 to 40256

| Score | Expect | Identities | Gaps | Strand | Frame |
|---|---|---|---|---|---|
| 14.4 bits(7) | 55() | 7/7(100%) | 0/7(0%) | Plus/Plus | |

Features:

```
Query  285    GTTCTCT  291
              |||||||
Sbjct  40250  GTTCTCT  40256
```

Range 94: 41054 to 41060

| Score | Expect | Identities | Gaps | Strand | Frame |
|---|---|---|---|---|---|
| 14.4 bits(7) | 55() | 7/7(100%) | 0/7(0%) | Plus/Minus | |

Features:

```
Query  286    TTCTCTC  292
              |||||||
Sbjct  41060  TTCTCTC  41054
```

Range 95: 41490 to 41496

| Score | Expect | Identities | Gaps | Strand | Frame |
|---|---|---|---|---|---|
| 14.4 bits(7) | 55() | 7/7(100%) | 0/7(0%) | Plus/Plus | |

Features:

```
Query  289    TCTCCTT  295
              |||||||
Sbjct  41490  TCTCCTT  41496
```

Range 96: 42171 to 42177

| Score | Expect | Identities | Gaps | Strand | Frame |
|---|---|---|---|---|---|
| 14.4 bits(7) | 55() | 7/7(100%) | 0/7(0%) | Plus/Plus | |

Features:

```
Query  281    TGGTGTT  287
              |||||||
Sbjct  42171  TGGTGTT  42177
```

Range 97: 44805 to 44811

| Score | Expect | Identities | Gaps | Strand | Frame |
|---|---|---|---|---|---|
| 14.4 bits(7) | 55() | 7/7(100%) | 0/7(0%) | Plus/Plus | |

Features:

```
Query  293    CTTTATT  299
              |||||||
Sbjct  44805  CTTTATT  44811
```

Range 98: 44934 to 44940

| Score | Expect | Identities | Gaps | Strand | Frame |
|---|---|---|---|---|---|
| 14.4 bits(7) | 55() | 7/7(100%) | 0/7(0%) | Plus/Plus | |

Features:

```
Query  272    GTAACAA  278
              |||||||
Sbjct  44934  GTAACAA  44940
```

Range 99: 45636 to 45642

| Score | Expect | Identities | Gaps | Strand | Frame |
|---|---|---|---|---|---|
| 14.4 bits(7) | 55() | 7/7(100%) | 0/7(0%) | Plus/Plus | |

Features:

```
Query  276    CAAGCTG  282
              |||||||
Sbjct  45636  CAAGCTG  45642
```

Range 100: 46304 to 46310

| Score | Expect | Identities | Gaps | Strand | Frame |
|---|---|---|---|---|---|
| 14.4 bits(7) | 55() | 7/7(100%) | 0/7(0%) | Plus/Plus | |

Features:

```
Query  285    GTTCTCT  291
              |||||||
Sbjct  46304  GTTCTCT  46310
```

Range 101: 46571 to 46577

| Score | Expect | Identities | Gaps | Strand | Frame |
|---|---|---|---|---|---|
| 14.4 bits(7) | 55() | 7/7(100%) | 0/7(0%) | Plus/Plus | |

Features:

```
Query  275    ACAAGCT  281
              |||||||
Sbjct  46571  ACAAGCT  46577
```

Range 102: 47134 to 47140

| Score | Expect | Identities | Gaps | Strand | Frame |
|---|---|---|---|---|---|
| 14.4 bits(7) | 55() | 7/7(100%) | 0/7(0%) | Plus/Plus | |

Features:

```
Query  288    CTCTCCT  294
              |||||||
Sbjct  47134  CTCTCCT  47140
```

Range 103: 47382 to 47388

| Score | Expect | Identities | Gaps | Strand | Frame |
|---|---|---|---|---|---|
| 14.4 bits(7) | 55() | 7/7(100%) | 0/7(0%) | Plus/Plus | |

Features:

```
Query  271    TGTAACA  277
              |||||||
Sbjct  47382  TGTAACA  47388
```

Range 104: 49098 to 49104

| Score | Expect | Identities | Gaps | Strand | Frame |
|---|---|---|---|---|---|
| 14.4 bits(7) | 55() | 7/7(100%) | 0/7(0%) | Plus/Plus | |

Features:

```
Query  284    TGTTCTC  290
              |||||||
Sbjct  49098  TGTTCTC  49104
```

Range 105: 49152 to 49158

| Score | Expect | Identities | Gaps | Strand | Frame |
|---|---|---|---|---|---|
| 14.4 bits(7) | 55() | 7/7(100%) | 0/7(0%) | Plus/Plus | |

Features:

```
Query  281    TGGTGTT  287
              |||||||
Sbjct  49152  TGGTGTT  49158
```

Range 106: 50393 to 50399

| Score | Expect | Identities | Gaps | Strand | Frame |
|---|---|---|---|---|---|

| 14.4 bits(7) | 55() | 7/7(100%) | 0/7(0%) | Plus/Plus |
|---|---|---|---|---|

Features:

```
Query  283    GTGTTCT  289
              |||||||
Sbjct  50393  GTGTTCT  50399
```

Range 107: 51186 to 51192

| Score | Expect | Identities | Gaps | Strand | Frame |
|---|---|---|---|---|---|
| 14.4 bits(7) | 55() | 7/7(100%) | 0/7(0%) | Plus/Plus | |

Features:

```
Query  290    CTCCTTT  296
              |||||||
Sbjct  51186  CTCCTTT  51192
```

Range 108: 51381 to 51387

| Score | Expect | Identities | Gaps | Strand | Frame |
|---|---|---|---|---|---|
| 14.4 bits(7) | 55() | 7/7(100%) | 0/7(0%) | Plus/Plus | |

Features:

```
Query  290    CTCCTTT  296
              |||||||
Sbjct  51381  CTCCTTT  51387
```

Range 109: 51557 to 51563

| Score | Expect | Identities | Gaps | Strand | Frame |
|---|---|---|---|---|---|
| 14.4 bits(7) | 55() | 7/7(100%) | 0/7(0%) | Plus/Plus | |

Features:

```
Query  293    CTTTATT  299
              |||||||
Sbjct  51557  CTTTATT  51563
```

Range 110: 52431 to 52437

| Score | Expect | Identities | Gaps | Strand | Frame |
|---|---|---|---|---|---|
| 14.4 bits(7) | 55() | 7/7(100%) | 0/7(0%) | Plus/Plus | |

Features:

```
Query  279    GCTGGTG  285
              |||||||
Sbjct  52431  GCTGGTG  52437
```

Range 111: 53764 to 53770

| Score | Expect | Identities | Gaps | Strand | Frame |
|---|---|---|---|---|---|
| 14.4 bits(7) | 55() | 7/7(100%) | 0/7(0%) | Plus/Plus | |

Features:

```
Query  284    TGTTCTC  290
              |||||||
Sbjct  53764  TGTTCTC  53770
```

Range 112: 54561 to 54567

| Score | Expect | Identities | Gaps | Strand | Frame |
|---|---|---|---|---|---|
| 14.4 bits(7) | 55() | 7/7(100%) | 0/7(0%) | Plus/Plus | |

Features:

```
Query  273    TAACAAG  279
              |||||||
Sbjct  54561  TAACAAG  54567
```

Range 113: 54904 to 54910

| Score | Expect | Identities | Gaps | Strand | Frame |
|---|---|---|---|---|---|
| 14.4 bits(7) | 55() | 7/7(100%) | 0/7(0%) | Plus/Plus | |

Features:

```
Query  293    CTTTATT  299
              |||||||
Sbjct  54904  CTTTATT  54910
```

Range 114: 55740 to 55746

| Score | Expect | Identities | Gaps | Strand | Frame |
|---|---|---|---|---|---|
| 14.4 bits(7) | 55() | 7/7(100%) | 0/7(0%) | Plus/Minus | |

Features:

```
Query  273    TAACAAG  279
              |||||||
Sbjct  55746  TAACAAG  55740
```

Range 115: 55816 to 55822

| Score | Expect | Identities | Gaps | Strand | Frame |
|---|---|---|---|---|---|
| 14.4 bits(7) | 55() | 7/7(100%) | 0/7(0%) | Plus/Minus | |

Features:

```
Query  289    TCTCCTT  295
              |||||||
Sbjct  55822  TCTCCTT  55816
```

Range 116: 55970 to 55976

| Score | Expect | Identities | Gaps | Strand | Frame |
|---|---|---|---|---|---|
| 14.4 bits(7) | 55() | 7/7(100%) | 0/7(0%) | Plus/Plus | |

Features:

```
Query  281    TGGTGTT  287
              |||||||
Sbjct  55970  TGGTGTT  55976
```

Range 117: 57444 to 57450

| Score | Expect | Identities | Gaps | Strand | Frame |
|---|---|---|---|---|---|
| 14.4 bits(7) | 55() | 7/7(100%) | 0/7(0%) | Plus/Minus | |

Features:

```
Query  293    CTTTATT  299
              |||||||
Sbjct  57450  CTTTATT  57444
```

Range 118: 59010 to 59016

| Score | Expect | Identities | Gaps | Strand | Frame |
|---|---|---|---|---|---|
| 14.4 bits(7) | 55() | 7/7(100%) | 0/7(0%) | Plus/Minus | |

Features:

```
Query  281    TGGTGTT  287
              |||||||
Sbjct  59016  TGGTGTT  59010
```

Range 119: 59219 to 59225

| Score | Expect | Identities | Gaps | Strand | Frame |
|---|---|---|---|---|---|
| 14.4 bits(7) | 55() | 7/7(100%) | 0/7(0%) | Plus/Minus | |

Features:

```
Query  293    CTTTATT  299
              |||||||
Sbjct  59225  CTTTATT  59219
```

Range 120: 60402 to 60408

| Score | Expect | Identities | Gaps | Strand | Frame |
|---|---|---|---|---|---|
| 14.4 bits(7) | 55() | 7/7(100%) | 0/7(0%) | Plus/Minus | |

Features:

```
Query  273    TAACAAG  279
              |||||||
Sbjct  60408  TAACAAG  60402
```

Range 121: 60438 to 60444

| Score | Expect | Identities | Gaps | Strand | Frame |
|---|---|---|---|---|---|
| 14.4 bits(7) | 55() | 7/7(100%) | 0/7(0%) | Plus/Plus | |

Features:

```
Query  286    TTCTCTC  292
              |||||||
Sbjct  60438  TTCTCTC  60444
```

Range 122: 60671 to 60677

| Score | Expect | Identities | Gaps | Strand | Frame |
|---|---|---|---|---|---|
| 14.4 bits(7) | 55() | 7/7(100%) | 0/7(0%) | Plus/Minus | |

Features:

```
Query  273    TAACAAG  279
              |||||||
Sbjct  60677  TAACAAG  60671
```

BLAST is a registered trademark of the National Library of Medicine

Support center Mailing list YouTube

- National Library Of Medicine
- National Institutes Of Health
- U.S. Department of Health & Human Services
- USA.gov

**NCBI**

*National Center for Biotechnology Information, U.S. National Library of Medicine 8600 Rockville Pike, Bethesda MD, 20894 USA*
Policies and Guidelines | Contact
