## Supplemental Figure S3 for "MORT, a locus for apoptosis in the human immunodeficiency virus-type 1 antisense gene: implications for AIDS, Cancer, and Covid-19"

**MirGeneDB**

Home | Browse | Search

Blast results for query sequence:

GUGUAACAAGCUGGUGUUCUCUCCUUUAUUGGCC

**BLASTN 2.2.27+**

**Reference**:
Zheng Zhang, Scott Schwartz, Lukas Wagner, and Webb Miller (2000),
"A greedy algorithm for aligning DNA sequences", J Comput Biol 2000;
7(1-2):203-14.

Database: ALL.fas
4,302 sequences; 151,598 total letters

**Query=**
Length=34

| Sequences producing significant alignments: | Score (Bits) | E Value |
|---|---|---|
| Hsa-Mir-5586_pre | 23.3 | 0.20 |
| Hsa-Mir-5586_5p | 23.3 | 0.20 |
| Dre-Mir-138-P2_pre | 19.6 | 2.6 |
| Dre-Mir-138-P2_5p | 19.6 | 2.6 |
| Dre-Mir-138-P1_pre | 19.6 | 2.6 |
| Dre-Mir-138-P1_5p | 19.6 | 2.6 |
| Gga-Mir-138-P2_pre | 19.6 | 2.6 |
| Gga-Mir-138-P2_5p | 19.6 | 2.6 |
| Gga-Mir-138-P1_pre | 19.6 | 2.6 |
| Gga-Mir-138-P1_5p | 19.6 | 2.6 |
| Mmu-Mir-138-P2_pre | 19.6 | 2.6 |
| Mmu-Mir-138-P2_5p | 19.6 | 2.6 |
| Mmu-Mir-138-P1_pre | 19.6 | 2.6 |
| Mmu-Mir-138-P1_5p | 19.6 | 2.6 |
| Hsa-Mir-7151_pre | 19.6 | 2.6 |
| Hsa-Mir-7151_5p | 19.6 | 2.6 |
| Hsa-Mir-5586_3p* | 19.6 | 2.6 |
| Hsa-Mir-3173_pre | 19.6 | 2.6 |
| Hsa-Mir-3173_5p | 19.6 | 2.6 |
| Hsa-Mir-138-P2_pre | 19.6 | 2.6 |
| Hsa-Mir-138-P2_5p | 19.6 | 2.6 |
| Hsa-Mir-138-P1_pre | 19.6 | 2.6 |
| Hsa-Mir-138-P1_5p | 19.6 | 2.6 |
| Dre-Mir-732_pre | 17.7 | 9.2 |
| Dre-Mir-732_3p | 17.7 | 9.2 |
| Dre-Mir-10-P2a2_pre | 17.7 | 9.2 |
| Dre-Mir-10-P2a1_pre | 17.7 | 9.2 |
| Dre-Mir-30-P1b_pre | 17.7 | 9.2 |
| Gga-Mir-10-P2a_pre | 17.7 | 9.2 |
| Gga-Mir-7-P3_pre | 17.7 | 9.2 |
| Mmu-Mir-3079_pre | 17.7 | 9.2 |
| Mmu-Mir-192-P1_pre | 17.7 | 9.2 |
| Mmu-Mir-129-P1_pre | 17.7 | 9.2 |
| Mmu-Mir-29-P1b_pre | 17.7 | 9.2 |
| Mmu-Mir-29-P1b_5p* | 17.7 | 9.2 |
| Mmu-Mir-10-P2c_pre | 17.7 | 9.2 |
| Mmu-Mir-10-P2a_pre | 17.7 | 9.2 |
| Hsa-Mir-1296_pre | 17.7 | 9.2 |
| Hsa-Mir-10-P2c_pre | 17.7 | 9.2 |
| Hsa-Mir-29-P1b_pre | 17.7 | 9.2 |
| Hsa-Mir-29-P1b_5p* | 17.7 | 9.2 |

> Hsa-Mir-5586_pre
Length=60

```
 Score = 23.3 bits (12),  Expect = 0.20
 Identities = 12/12 (100%), Gaps = 0/12 (0%)
 Strand=Plus/Minus
```

```
Query  3   GTAACAAGCTGG  14
           ||||||||||||
Sbjct  16  GTAACAAGCTGG  5


 Score = 19.6 bits (10),  Expect = 2.6
 Identities = 10/10 (100%), Gaps = 0/10 (0%)
 Strand=Plus/Plus

Query  6   ACAAGCTGGT  15
           ||||||||||
Sbjct  46  ACAAGCTGGT  55
```

> Hsa-Mir-5586_5p
Length=22

```
 Score = 23.3 bits (12),  Expect = 0.20
 Identities = 12/12 (100%), Gaps = 0/12 (0%)
 Strand=Plus/Minus

Query  3   GTAACAAGCTGG  14
           ||||||||||||
Sbjct  16  GTAACAAGCTGG  5
```

> Dre-Mir-138-P2_pre
Length=68

```
 Score = 19.6 bits (10),  Expect = 2.6
 Identities = 10/10 (100%), Gaps = 0/10 (0%)
 Strand=Plus/Plus

Query  9   AGCTGGTGTT  18
           ||||||||||
Sbjct  1   AGCTGGTGTT  10
```

> Dre-Mir-138-P2_5p
Length=23

```
 Score = 19.6 bits (10),  Expect = 2.6
 Identities = 10/10 (100%), Gaps = 0/10 (0%)
 Strand=Plus/Plus

Query  9   AGCTGGTGTT  18
           ||||||||||
Sbjct  1   AGCTGGTGTT  10
```

> Dre-Mir-138-P1_pre
Length=62

```
 Score = 19.6 bits (10),  Expect = 2.6
 Identities = 10/10 (100%), Gaps = 0/10 (0%)
 Strand=Plus/Plus

Query  9   AGCTGGTGTT  18
           ||||||||||
Sbjct  1   AGCTGGTGTT  10
```

> Dre-Mir-138-P1_5p
Length=23

```
 Score = 19.6 bits (10),  Expect = 2.6
 Identities = 10/10 (100%), Gaps = 0/10 (0%)
 Strand=Plus/Plus

Query  9   AGCTGGTGTT  18
           ||||||||||
Sbjct  1   AGCTGGTGTT  10
```

> Gga-Mir-138-P2_pre
Length=68

```
 Score = 19.6 bits (10),  Expect = 2.6
 Identities = 10/10 (100%), Gaps = 0/10 (0%)
 Strand=Plus/Plus

Query  9   AGCTGGTGTT  18
           ||||||||||
Sbjct  1   AGCTGGTGTT  10
```

> Gga-Mir-138-P2_5p

```
Length=23

 Score = 19.6 bits (10),  Expect = 2.6
 Identities = 10/10 (100%), Gaps = 0/10 (0%)
 Strand=Plus/Plus

Query  9   AGCTGGTGTT  18
           ||||||||||
Sbjct  1   AGCTGGTGTT  10
```

> Gga-Mir-138-P1_pre

```
Length=61

 Score = 19.6 bits (10),  Expect = 2.6
 Identities = 10/10 (100%), Gaps = 0/10 (0%)
 Strand=Plus/Plus

Query  9   AGCTGGTGTT  18
           ||||||||||
Sbjct  1   AGCTGGTGTT  10
```

> Gga-Mir-138-P1_5p

```
Length=23

 Score = 19.6 bits (10),  Expect = 2.6
 Identities = 10/10 (100%), Gaps = 0/10 (0%)
 Strand=Plus/Plus

Query  9   AGCTGGTGTT  18
           ||||||||||
Sbjct  1   AGCTGGTGTT  10
```

> Mmu-Mir-138-P2_pre

```
Length=69

 Score = 19.6 bits (10),  Expect = 2.6
 Identities = 10/10 (100%), Gaps = 0/10 (0%)
 Strand=Plus/Plus

Query  9   AGCTGGTGTT  18
           ||||||||||
Sbjct  1   AGCTGGTGTT  10
```

> Mmu-Mir-138-P2_5p

```
Length=23

 Score = 19.6 bits (10),  Expect = 2.6
 Identities = 10/10 (100%), Gaps = 0/10 (0%)
 Strand=Plus/Plus

Query  9   AGCTGGTGTT  18
           ||||||||||
Sbjct  1   AGCTGGTGTT  10
```

> Mmu-Mir-138-P1_pre

```
Length=61

 Score = 19.6 bits (10),  Expect = 2.6
 Identities = 10/10 (100%), Gaps = 0/10 (0%)
 Strand=Plus/Plus

Query  9   AGCTGGTGTT  18
           ||||||||||
Sbjct  1   AGCTGGTGTT  10
```

> Mmu-Mir-138-P1_5p

```
Length=23

 Score = 19.6 bits (10),  Expect = 2.6
 Identities = 10/10 (100%), Gaps = 0/10 (0%)
 Strand=Plus/Plus

Query  9   AGCTGGTGTT  18
           ||||||||||
Sbjct  1   AGCTGGTGTT  10
```

> Hsa-Mir-7151_pre

```
Length=60

 Score = 19.6 bits (10),  Expect = 2.6
```

```
 Identities = 15/17 (88%), Gaps = 1/17 (6%)
 Strand=Plus/Plus

Query  18  TCTCT-CCTTTATTGGC  33
           ||||| ||| |||||||
Sbjct  7   TCTCTGCCTGTATTGGC  23
```

> Hsa-Mir-7151_5p

```
Length=24

 Score = 19.6 bits (10),  Expect = 2.6
 Identities = 15/17 (88%), Gaps = 1/17 (6%)
 Strand=Plus/Plus

Query  18  TCTCT-CCTTTATTGGC  33
           ||||| ||| |||||||
Sbjct  7   TCTCTGCCTGTATTGGC  23
```

> Hsa-Mir-5586_3p*

```
Length=22

 Score = 19.6 bits (10),  Expect = 2.6
 Identities = 10/10 (100%), Gaps = 0/10 (0%)
 Strand=Plus/Plus

Query  6   ACAAGCTGGT  15
           ||||||||||
Sbjct  8   ACAAGCTGGT  17
```

> Hsa-Mir-3173_pre

```
Length=62

 Score = 19.6 bits (10),  Expect = 2.6
 Identities = 12/13 (92%), Gaps = 0/13 (0%)
 Strand=Plus/Plus

Query  15  TGTTCTCTCCTTT  27
           |||| ||||||||
Sbjct  10  TGTTTTCTCCTTT  22
```

> Hsa-Mir-3173_5p

```
Length=23

 Score = 19.6 bits (10),  Expect = 2.6
 Identities = 12/13 (92%), Gaps = 0/13 (0%)
 Strand=Plus/Plus

Query  15  TGTTCTCTCCTTT  27
           |||| ||||||||
Sbjct  10  TGTTTTCTCCTTT  22
```

> Hsa-Mir-138-P2_pre

```
Length=68

 Score = 19.6 bits (10),  Expect = 2.6
 Identities = 10/10 (100%), Gaps = 0/10 (0%)
 Strand=Plus/Plus

Query  9   AGCTGGTGTT  18
           ||||||||||
Sbjct  1   AGCTGGTGTT  10
```

> Hsa-Mir-138-P2_5p

```
Length=23

 Score = 19.6 bits (10),  Expect = 2.6
 Identities = 10/10 (100%), Gaps = 0/10 (0%)
 Strand=Plus/Plus

Query  9   AGCTGGTGTT  18
           ||||||||||
Sbjct  1   AGCTGGTGTT  10
```

> Hsa-Mir-138-P1_pre

```
Length=61

 Score = 19.6 bits (10),  Expect = 2.6
 Identities = 10/10 (100%), Gaps = 0/10 (0%)
 Strand=Plus/Plus
```

```
Query  9   AGCTGGTGTT  18
           ||||||||||
Sbjct  1   AGCTGGTGTT  10
```

> Hsa-Mir-138-P1_5p

```
Length=23

 Score = 19.6 bits (10),  Expect = 2.6
 Identities = 10/10 (100%), Gaps = 0/10 (0%)
 Strand=Plus/Plus

Query  9   AGCTGGTGTT  18
           ||||||||||
Sbjct  1   AGCTGGTGTT  10
```

> Dre-Mir-732_pre

```
Length=59

 Score = 17.7 bits (9),  Expect = 9.2
 Identities = 11/12 (92%), Gaps = 0/12 (0%)
 Strand=Plus/Minus

Query  16  GTTCTCTCCTTT  27
           ||||||| ||||
Sbjct  52  GTTCTCTGCTTT  41
```

> Dre-Mir-732_3p

```
Length=22

 Score = 17.7 bits (9),  Expect = 9.2
 Identities = 11/12 (92%), Gaps = 0/12 (0%)
 Strand=Plus/Minus

Query  16  GTTCTCTCCTTT  27
           ||||||| ||||
Sbjct  15  GTTCTCTGCTTT  4
```

> Dre-Mir-10-P2a2_pre

```
Length=57

 Score = 17.7 bits (9),  Expect = 9.2
 Identities = 11/12 (92%), Gaps = 0/12 (0%)
 Strand=Plus/Plus

Query  6   ACAAGCTGGTGT  17
           ||||||| ||||
Sbjct  35  ACAAGCTCGTGT  46
```

> Dre-Mir-10-P2a1_pre

```
Length=57

 Score = 17.7 bits (9),  Expect = 9.2
 Identities = 11/12 (92%), Gaps = 0/12 (0%)
 Strand=Plus/Minus

Query  6   ACAAGCTGGTGT  17
           ||||||| ||||
Sbjct  44  ACAAGCTTGTGT  33
```

> Dre-Mir-30-P1b_pre

```
Length=62

 Score = 17.7 bits (9),  Expect = 9.2
 Identities = 9/9 (100%), Gaps = 0/9 (0%)
 Strand=Plus/Plus

Query  8   AAGCTGGTG  16
           |||||||||
Sbjct  20  AAGCTGGTG  28
```

> Gga-Mir-10-P2a_pre

```
Length=57

 Score = 17.7 bits (9),  Expect = 9.2
 Identities = 11/12 (92%), Gaps = 0/12 (0%)
 Strand=Plus/Minus

Query  6   ACAAGCTGGTGT  17
           ||||||| ||||
Sbjct  44  ACAAGCTTGTGT  33
```

```
> Gga-Mir-7-P3_pre
Length=63

 Score = 17.7 bits (9),  Expect = 9.2
 Identities = 13/15 (87%), Gaps = 0/15 (0%)
 Strand=Plus/Minus

Query  15  TGTTCTCTCCTTTAT  29
           |||| || |||||||
Sbjct  46  TGTTGTCACCTTTAT  32


> Mmu-Mir-3079_pre
Length=63

 Score = 17.7 bits (9),  Expect = 9.2
 Identities = 11/12 (92%), Gaps = 0/12 (0%)
 Strand=Plus/Plus

Query  8   AAGCTGGTGTTC  19
           ||||||| ||||
Sbjct  17  AAGCTGGAGTTC  28


> Mmu-Mir-192-P1_pre
Length=65

 Score = 17.7 bits (9),  Expect = 9.2
 Identities = 9/9 (100%), Gaps = 0/9 (0%)
 Strand=Plus/Plus

Query  17  TTCTCTCCT  25
           |||||||||
Sbjct  31  TTCTCTCCT  39


> Mmu-Mir-129-P1_pre
Length=65

 Score = 17.7 bits (9),  Expect = 9.2
 Identities = 9/9 (100%), Gaps = 0/9 (0%)
 Strand=Plus/Plus

Query  15  TGTTCTCTC  23
           |||||||||
Sbjct  22  TGTTCTCTC  30


> Mmu-Mir-29-P1b_pre
Length=58

 Score = 17.7 bits (9),  Expect = 9.2
 Identities = 9/9 (100%), Gaps = 0/9 (0%)
 Strand=Plus/Plus

Query  11  CTGGTGTTC  19
           |||||||||
Sbjct  12  CTGGTGTTC  20


> Mmu-Mir-29-P1b_5p*
Length=23

 Score = 17.7 bits (9),  Expect = 9.2
 Identities = 9/9 (100%), Gaps = 0/9 (0%)
 Strand=Plus/Plus

Query  11  CTGGTGTTC  19
           |||||||||
Sbjct  12  CTGGTGTTC  20


> Mmu-Mir-10-P2c_pre
Length=60

 Score = 17.7 bits (9),  Expect = 9.2
 Identities = 11/12 (92%), Gaps = 0/12 (0%)
 Strand=Plus/Plus

Query  6   ACAAGCTGGTGT  17
           ||||||| ||||
Sbjct  38  ACAAGCTCGTGT  49


> Mmu-Mir-10-P2a_pre
```

```
Length=57

 Score = 17.7 bits (9),  Expect = 9.2
 Identities = 11/12 (92%), Gaps = 0/12 (0%)
 Strand=Plus/Minus

Query  6   ACAAGCTGGTGT  17
           ||||||| ||||
Sbjct  44  ACAAGCTTGTGT  33


 Score = 17.7 bits (9),  Expect = 9.2
 Identities = 11/12 (92%), Gaps = 0/12 (0%)
 Strand=Plus/Plus

Query  6   ACAAGCTGGTGT  17
           ||||||| ||||
Sbjct  35  ACAAGCTTGTGT  46


> Hsa-Mir-1296_pre
Length=65

 Score = 17.7 bits (9),  Expect = 9.2
 Identities = 9/9 (100%), Gaps = 0/9 (0%)
 Strand=Plus/Plus

Query  20  TCTCCTTTA  28
           |||||||||
Sbjct  18  TCTCCTTTA  26


> Hsa-Mir-10-P2c_pre
Length=60

 Score = 17.7 bits (9),  Expect = 9.2
 Identities = 11/12 (92%), Gaps = 0/12 (0%)
 Strand=Plus/Plus

Query  6   ACAAGCTGGTGT  17
           ||||||| ||||
Sbjct  38  ACAAGCTCGTGT  49


> Hsa-Mir-29-P1b_pre
Length=58

 Score = 17.7 bits (9),  Expect = 9.2
 Identities = 9/9 (100%), Gaps = 0/9 (0%)
 Strand=Plus/Plus

Query  11  CTGGTGTTC  19
           |||||||||
Sbjct  12  CTGGTGTTC  20


> Hsa-Mir-29-P1b_5p*
Length=23

 Score = 17.7 bits (9),  Expect = 9.2
 Identities = 9/9 (100%), Gaps = 0/9 (0%)
 Strand=Plus/Plus

Query  11  CTGGTGTTC  19
           |||||||||
Sbjct  12  CTGGTGTTC  20



Lambda      K        H
    1.33    0.621     1.12

Gapped
Lambda      K        H
    1.28    0.460    0.850

Effective search space used: 2009112


  Database: ALL.fas
    Posted date:  Nov 9, 2015  3:11 PM
  Number of letters in database: 151,598
  Number of sequences in database:  4,302



Matrix: blastn matrix 1 -2
```

```
Gap Penalties: Existence: 0, Extension: 2.5
```

---

For any errors, false omissions, or false inclusions please contact Kevin Peterson at.
