## Supplemental Figure S4 for "MORT, a locus for apoptosis in the human immunodeficiency virus-type 1 antisense gene: implications for AIDS, Cancer, and Covid-19"

**BLAST Results**

**Blast 2 sequences**

**Job title: D36MORT1997.VS.XIAP**

| | |
|---|---|
| **RID** | 5524BW23114 (Expires on 01-09 03:52 am) |
| **Query ID** | lcl\|Query_236255 |
| **Description** | None |
| **Molecule type** | nucleic acid |
| **Query Length** | 26 |

| | |
|---|---|
| **Subject ID** | NG_007264.1 |
| **Description** | Homo sapiens X-linked inhibitor of apoptosis (XIAP), RefSeqGene (LRG_19) on chromosome X<br>See details |
| **Molecule type** | nucleic acid |
| **Subject Length** | 60775 |
| **Program** | BLASTN 2.7.1+ |

### Graphic Summary

**Distribution of the top 247 Blast Hits on 1 subject sequences**

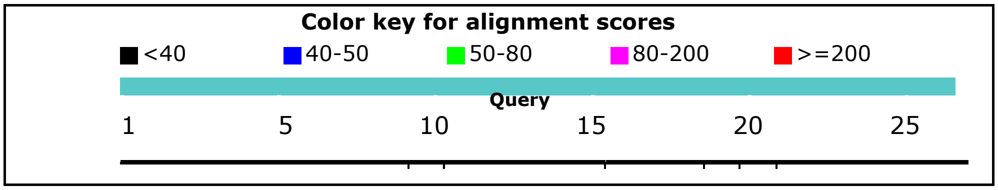

### Dot Matrix View

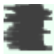

### Descriptions

Sequences producing significant alignments:

| Description | Max score | Total score | Query cover | E value | Ident | Accession |
|---|---|---|---|---|---|---|
| Homo sapiens X-linked inhibitor of apoptosis (XIAP), RefSeqGene (LRG_19) on chromosome X | 24.3 | 3674 | 100% | 0.048 | 100% | NG_007264.1 |

### Alignments

Homo sapiens X-linked inhibitor of apoptosis (XIAP), RefSeqGene (LRG_19) on chromosome X
Sequence ID: **NG_007264.1** Length: 60775 Number of Matches: 247
Range 1: 45674 to 45685

| Score | Expect | Identities | Gaps | Strand | Frame |
|---|---|---|---|---|---|
| 24.3 bits(12) | 0.048() | 12/12(100%) | 0/12(0%) | Plus/Minus | |

Features:

```
Query  4      AGTCTGTTGTTT  15
              ||||||||||||
Sbjct  45685  AGTCTGTTGTTT  45674
```

Range 2: 3492 to 3501

| Score | Expect | Identities | Gaps | Strand | Frame |
|---|---|---|---|---|---|
| 20.3 bits(10) | 0.74() | 10/10(100%) | 0/10(0%) | Plus/Minus | |

Features:

```
Query  9     GTTGTTTTCT  18
             ||||||||||
Sbjct  3501  GTTGTTTTCT  3492
```

Range 3: 30233 to 30242

| Score | Expect | Identities | Gaps | Strand | Frame |
|---|---|---|---|---|---|
| 20.3 bits(10) | 0.74() | 10/10(100%) | 0/10(0%) | Plus/Minus | |

Features:

```
Query  10     TTGTTTTCTC  19
              ||||||||||
Sbjct  30242  TTGTTTTCTC  30233
```

Range 4: 42879 to 42892

| Score | Expect | Identities | Gaps | Strand | Frame |
|---|---|---|---|---|---|
| 20.3 bits(10) | 0.74() | 13/14(93%) | 0/14(0%) | Plus/Plus | |

Features:

```
Query  13     TTTTCTCCTCTATT  26
              ||||||||| ||||
Sbjct  42879  TTTTCTCCTGTATT  42892
```

Range 5: 51519 to 51528

| Score | Expect | Identities | Gaps | Strand | Frame |
|---|---|---|---|---|---|
| 20.3 bits(10) | 0.74() | 10/10(100%) | 0/10(0%) | Plus/Plus | |

Features:

```
Query  10     TTGTTTTCTC  19
              ||||||||||
Sbjct  51519  TTGTTTTCTC  51528
```

Range 6: 4071 to 4079

| Score | Expect | Identities | Gaps | Strand | Frame |
|---|---|---|---|---|---|
| 18.3 bits(9) | 2.9() | 9/9(100%) | 0/9(0%) | Plus/Plus | |

Features:

```
Query  18    TCCTCTATT  26
             |||||||||
Sbjct  4071  TCCTCTATT  4079
```

Range 7: 5614 to 5622

| Score | Expect | Identities | Gaps | Strand | Frame |
|---|---|---|---|---|---|
| 18.3 bits(9) | 2.9() | 9/9(100%) | 0/9(0%) | Plus/Plus | |

Features:

```
Query  10    TTGTTTTCT  18
             |||||||||
Sbjct  5614  TTGTTTTCT  5622
```

Range 8: 17293 to 17301

| Score | Expect | Identities | Gaps | Strand | Frame |
|---|---|---|---|---|---|
| 18.3 bits(9) | 2.9() | 9/9(100%) | 0/9(0%) | Plus/Plus | |

Features:

```
Query  1      AACAGTCTG  9
              |||||||||
Sbjct  17293  AACAGTCTG  17301
```

Range 9: 26124 to 26132

| Score | Expect | Identities | Gaps | Strand | Frame |
|---|---|---|---|---|---|
| 18.3 bits(9) | 2.9() | 9/9(100%) | 0/9(0%) | Plus/Plus | |

Features:

```
Query  10     TTGTTTTCT  18
              |||||||||
Sbjct  26124  TTGTTTTCT  26132
```

Range 10: 30378 to 30386

| Score | Expect | Identities | Gaps | Strand | Frame |
|---|---|---|---|---|---|
| 18.3 bits(9) | 2.9() | 9/9(100%) | 0/9(0%) | Plus/Plus | |

Features:

```
Query  4      AGTCTGTTG  12
              |||||||||
Sbjct  30378  AGTCTGTTG  30386
```

Range 11: 37546 to 37554

| Score | Expect | Identities | Gaps | Strand | Frame |
|---|---|---|---|---|---|
| 18.3 bits(9) | 2.9() | 9/9(100%) | 0/9(0%) | Plus/Minus | |

Features:

```
Query  11     TGTTTTCTC  19
              |||||||||
Sbjct  37554  TGTTTTCTC  37546
```

Range 12: 49341 to 49349

| Score | Expect | Identities | Gaps | Strand | Frame |
|---|---|---|---|---|---|
| 18.3 bits(9) | 2.9() | 9/9(100%) | 0/9(0%) | Plus/Plus | |

Features:

```
Query  12     GTTTTCTCC  20
              |||||||||
Sbjct  49341  GTTTTCTCC  49349
```

Range 13: 51825 to 51833

| Score | Expect | Identities | Gaps | Strand | Frame |
|---|---|---|---|---|---|
| 18.3 bits(9) | 2.9() | 9/9(100%) | 0/9(0%) | Plus/Minus | |

Features:

```
Query  15     TTCTCCTCT  23
              |||||||||
Sbjct  51833  TTCTCCTCT  51825
```

Range 14: 53434 to 53442

| Score | Expect | Identities | Gaps | Strand | Frame |
|---|---|---|---|---|---|
| 18.3 bits(9) | 2.9() | 9/9(100%) | 0/9(0%) | Plus/Plus | |

Features:

```
Query  10     TTGTTTTCT  18
              |||||||||
Sbjct  53434  TTGTTTTCT  53442
```

Range 15: 396 to 403

| Score | Expect | Identities | Gaps | Strand | Frame |
|---|---|---|---|---|---|
| 16.4 bits(8) | 12() | 8/8(100%) | 0/8(0%) | Plus/Plus | |

Features:

```
Query  19   CCTCTATT  26
            ||||||||
Sbjct  396  CCTCTATT  403
```

Range 16: 422 to 429

| Score | Expect | Identities | Gaps | Strand | Frame |
|---|---|---|---|---|---|
| 16.4 bits(8) | 12() | 8/8(100%) | 0/8(0%) | Plus/Plus | |

Features:

```
Query  14   TTTCTCCT  21
            ||||||||
Sbjct  422  TTTCTCCT  429
```

Range 17: 583 to 590

| Score | Expect | Identities | Gaps | Strand | Frame |
|---|---|---|---|---|---|
| 16.4 bits(8) | 12() | 8/8(100%) | 0/8(0%) | Plus/Plus | |

Features:

```
Query  15   TTCTCCTC  22
            ||||||||
```

```
Sbjct  583  TTCTCCTC  590
```

Range 18: 1448 to 1455

| Score | Expect | Identities | Gaps | Strand | Frame |
| --- | --- | --- | --- | --- | --- |
| 16.4 bits(8) | 12() | 8/8(100%) | 0/8(0%) | Plus/Minus | |

Features:

```
Query  15    TTCTCCTC  22
             ||||||||
Sbjct  1455  TTCTCCTC  1448
```

Range 19: 1539 to 1546

| Score | Expect | Identities | Gaps | Strand | Frame |
| --- | --- | --- | --- | --- | --- |
| 16.4 bits(8) | 12() | 8/8(100%) | 0/8(0%) | Plus/Minus | |

Features:

```
Query  15    TTCTCCTC  22
             ||||||||
Sbjct  1546  TTCTCCTC  1539
```

Range 20: 4378 to 4385

| Score | Expect | Identities | Gaps | Strand | Frame |
| --- | --- | --- | --- | --- | --- |
| 16.4 bits(8) | 12() | 8/8(100%) | 0/8(0%) | Plus/Plus | |

Features:

```
Query  15    TTCTCCTC  22
             ||||||||
Sbjct  4378  TTCTCCTC  4385
```

Range 21: 6370 to 6377

| Score | Expect | Identities | Gaps | Strand | Frame |
| --- | --- | --- | --- | --- | --- |
| 16.4 bits(8) | 12() | 8/8(100%) | 0/8(0%) | Plus/Minus | |

Features:

```
Query  15    TTCTCCTC  22
             ||||||||
Sbjct  6377  TTCTCCTC  6370
```

Range 22: 11830 to 11837

| Score | Expect | Identities | Gaps | Strand | Frame |
| --- | --- | --- | --- | --- | --- |
| 16.4 bits(8) | 12() | 8/8(100%) | 0/8(0%) | Plus/Plus | |

Features:

```
Query  9      GTTGTTTT  16
              ||||||||
Sbjct  11830  GTTGTTTT  11837
```

Range 23: 13848 to 13855

| Score | Expect | Identities | Gaps | Strand | Frame |
| --- | --- | --- | --- | --- | --- |
| 16.4 bits(8) | 12() | 8/8(100%) | 0/8(0%) | Plus/Minus | |

Features:

```
Query  3      CAGTCTGT  10
              ||||||||
Sbjct  13855  CAGTCTGT  13848
```

Range 24: 17866 to 17873

| Score | Expect | Identities | Gaps | Strand | Frame |
|---|---|---|---|---|---|
| 16.4 bits(8) | 12() | 8/8(100%) | 0/8(0%) | Plus/Plus | |

Features:

```
Query  7      CTGTTGTT  14
              ||||||||
Sbjct  17866  CTGTTGTT  17873
```

Range 25: 20457 to 20464

| Score | Expect | Identities | Gaps | Strand | Frame |
|---|---|---|---|---|---|
| 16.4 bits(8) | 12() | 8/8(100%) | 0/8(0%) | Plus/Plus | |

Features:

```
Query  11     TGTTTTCT  18
              ||||||||
Sbjct  20457  TGTTTTCT  20464
```

Range 26: 23371 to 23378

| Score | Expect | Identities | Gaps | Strand | Frame |
|---|---|---|---|---|---|
| 16.4 bits(8) | 12() | 8/8(100%) | 0/8(0%) | Plus/Minus | |

Features:

```
Query  11     TGTTTTCT  18
              ||||||||
Sbjct  23378  TGTTTTCT  23371
```

Range 27: 24460 to 24467

| Score | Expect | Identities | Gaps | Strand | Frame |
|---|---|---|---|---|---|
| 16.4 bits(8) | 12() | 8/8(100%) | 0/8(0%) | Plus/Minus | |

Features:

```
Query  13     TTTTCTCC  20
              ||||||||
Sbjct  24467  TTTTCTCC  24460
```

Range 28: 25032 to 25039

| Score | Expect | Identities | Gaps | Strand | Frame |
|---|---|---|---|---|---|
| 16.4 bits(8) | 12() | 8/8(100%) | 0/8(0%) | Plus/Minus | |

Features:

```
Query  18     TCCTCTAT  25
              ||||||||
Sbjct  25039  TCCTCTAT  25032
```

Range 29: 27166 to 27173

| Score | Expect | Identities | Gaps | Strand | Frame |
|---|---|---|---|---|---|
| 16.4 bits(8) | 12() | 8/8(100%) | 0/8(0%) | Plus/Plus | |

Features:

```
Query  11     TGTTTTCT  18
              ||||||||
Sbjct  27166  TGTTTTCT  27173
```

Range 30: 30010 to 30017

| Score | Expect | Identities | Gaps | Strand | Frame |
|---|---|---|---|---|---|

| | | | | |
|---|---|---|---|---|
| 16.4 bits(8) | 12() | 8/8(100%) | 0/8(0%) | Plus/Minus |

Features:

```
Query  9      GTTGTTTT  16
              ||||||||
Sbjct  30017  GTTGTTTT  30010
```

Range 31: 41207 to 41214

| Score | Expect | Identities | Gaps | Strand | Frame |
|---|---|---|---|---|---|
| 16.4 bits(8) | 12() | 8/8(100%) | 0/8(0%) | Plus/Plus | |

Features:

```
Query  1      AACAGTCT  8
              ||||||||
Sbjct  41207  AACAGTCT  41214
```

Range 32: 41359 to 41366

| Score | Expect | Identities | Gaps | Strand | Frame |
|---|---|---|---|---|---|
| 16.4 bits(8) | 12() | 8/8(100%) | 0/8(0%) | Plus/Plus | |

Features:

```
Query  13     TTTTCTCC  20
              ||||||||
Sbjct  41359  TTTTCTCC  41366
```

Range 33: 42150 to 42157

| Score | Expect | Identities | Gaps | Strand | Frame |
|---|---|---|---|---|---|
| 16.4 bits(8) | 12() | 8/8(100%) | 0/8(0%) | Plus/Plus | |

Features:

```
Query  11     TGTTTTCT  18
              ||||||||
Sbjct  42150  TGTTTTCT  42157
```

Range 34: 47747 to 47754

| Score | Expect | Identities | Gaps | Strand | Frame |
|---|---|---|---|---|---|
| 16.4 bits(8) | 12() | 8/8(100%) | 0/8(0%) | Plus/Minus | |

Features:

```
Query  15     TTCTCCTC  22
              ||||||||
Sbjct  47754  TTCTCCTC  47747
```

Range 35: 49232 to 49239

| Score | Expect | Identities | Gaps | Strand | Frame |
|---|---|---|---|---|---|
| 16.4 bits(8) | 12() | 8/8(100%) | 0/8(0%) | Plus/Plus | |

Features:

```
Query  12     GTTTTCTC  19
              ||||||||
Sbjct  49232  GTTTTCTC  49239
```

Range 36: 50184 to 50191

| Score | Expect | Identities | Gaps | Strand | Frame |
|---|---|---|---|---|---|
| 16.4 bits(8) | 12() | 8/8(100%) | 0/8(0%) | Plus/Plus | |

Features:

```
Query  13     TTTTCTCC  20
              ||||||||
Sbjct  50184  TTTTCTCC  50191
```

Range 37: 50508 to 50515

| Score | Expect | Identities | Gaps | Strand | Frame |
|---|---|---|---|---|---|
| 16.4 bits(8) | 12() | 8/8(100%) | 0/8(0%) | Plus/Plus | |

Features:

```
Query  10     TTGTTTTC  17
              ||||||||
Sbjct  50508  TTGTTTTC  50515
```

Range 38: 51341 to 51348

| Score | Expect | Identities | Gaps | Strand | Frame |
|---|---|---|---|---|---|
| 16.4 bits(8) | 12() | 8/8(100%) | 0/8(0%) | Plus/Plus | |

Features:

```
Query  19     CCTCTATT  26
              ||||||||
Sbjct  51341  CCTCTATT  51348
```

Range 39: 54316 to 54323

| Score | Expect | Identities | Gaps | Strand | Frame |
|---|---|---|---|---|---|
| 16.4 bits(8) | 12() | 8/8(100%) | 0/8(0%) | Plus/Minus | |

Features:

```
Query  13     TTTTCTCC  20
              ||||||||
Sbjct  54323  TTTTCTCC  54316
```

Range 40: 55167 to 55174

| Score | Expect | Identities | Gaps | Strand | Frame |
|---|---|---|---|---|---|
| 16.4 bits(8) | 12() | 8/8(100%) | 0/8(0%) | Plus/Plus | |

Features:

```
Query  9      GTTGTTTT  16
              ||||||||
Sbjct  55167  GTTGTTTT  55174
```

Range 41: 57092 to 57099

| Score | Expect | Identities | Gaps | Strand | Frame |
|---|---|---|---|---|---|
| 16.4 bits(8) | 12() | 8/8(100%) | 0/8(0%) | Plus/Plus | |

Features:

```
Query  7      CTGTTGTT  14
              ||||||||
Sbjct  57092  CTGTTGTT  57099
```

Range 42: 57510 to 57517

| Score | Expect | Identities | Gaps | Strand | Frame |
|---|---|---|---|---|---|
| 16.4 bits(8) | 12() | 8/8(100%) | 0/8(0%) | Plus/Plus | |

Features:

```
Query  1      AACAGTCT  8
              ||||||||
Sbjct  57510  AACAGTCT  57517
```

Range 43: 11 to 17

| Score | Expect | Identities | Gaps | Strand | Frame |
|---|---|---|---|---|---|
| 14.4 bits(7) | 46() | 7/7(100%) | 0/7(0%) | Plus/Plus | |

Features:

```
Query  15  TTCTCCT  21
           |||||||
Sbjct  11  TTCTCCT  17
```

Range 44: 881 to 887

| Score | Expect | Identities | Gaps | Strand | Frame |
|---|---|---|---|---|---|
| 14.4 bits(7) | 46() | 7/7(100%) | 0/7(0%) | Plus/Minus | |

Features:

```
Query  10   TTGTTTT  16
            |||||||
Sbjct  887  TTGTTTT  881
```

Range 45: 1099 to 1105

| Score | Expect | Identities | Gaps | Strand | Frame |
|---|---|---|---|---|---|
| 14.4 bits(7) | 46() | 7/7(100%) | 0/7(0%) | Plus/Minus | |

Features:

```
Query  14    TTTCTCC  20
             |||||||
Sbjct  1105  TTTCTCC  1099
```

Range 46: 1185 to 1191

| Score | Expect | Identities | Gaps | Strand | Frame |
|---|---|---|---|---|---|
| 14.4 bits(7) | 46() | 7/7(100%) | 0/7(0%) | Plus/Minus | |

Features:

```
Query  15    TTCTCCT  21
             |||||||
Sbjct  1191  TTCTCCT  1185
```

Range 47: 1804 to 1810

| Score | Expect | Identities | Gaps | Strand | Frame |
|---|---|---|---|---|---|
| 14.4 bits(7) | 46() | 7/7(100%) | 0/7(0%) | Plus/Minus | |

Features:

```
Query  16    TCTCCTC  22
             |||||||
Sbjct  1810  TCTCCTC  1804
```

Range 48: 2950 to 2956

| Score | Expect | Identities | Gaps | Strand | Frame |
|---|---|---|---|---|---|
| 14.4 bits(7) | 46() | 7/7(100%) | 0/7(0%) | Plus/Plus | |

Features:

```
Query  15    TTCTCCT  21
             |||||||
Sbjct  2950  TTCTCCT  2956
```

Range 49: 3519 to 3525

| Score | Expect | Identities | Gaps | Strand | Frame |
|---|---|---|---|---|---|

| 14.4 bits(7) | 46() | 7/7(100%) | 0/7(0%) | Plus/Minus |
|---|---|---|---|---|

Features:

```
Query  10    TTGTTTT  16
             |||||||
Sbjct  3525  TTGTTTT  3519
```

Range 50: 3827 to 3833

| Score | Expect | Identities | Gaps | Strand | Frame |
|---|---|---|---|---|---|
| 14.4 bits(7) | 46() | 7/7(100%) | 0/7(0%) | Plus/Plus | |

Features:

```
Query  16    TCTCCTC  22
             |||||||
Sbjct  3827  TCTCCTC  3833
```

Range 51: 4033 to 4039

| Score | Expect | Identities | Gaps | Strand | Frame |
|---|---|---|---|---|---|
| 14.4 bits(7) | 46() | 7/7(100%) | 0/7(0%) | Plus/Minus | |

Features:

```
Query  4     AGTCTGT  10
             |||||||
Sbjct  4039  AGTCTGT  4033
```

Range 52: 4383 to 4389

| Score | Expect | Identities | Gaps | Strand | Frame |
|---|---|---|---|---|---|
| 14.4 bits(7) | 46() | 7/7(100%) | 0/7(0%) | Plus/Plus | |

Features:

```
Query  17    CTCCTCT  23
             |||||||
Sbjct  4383  CTCCTCT  4389
```

Range 53: 4898 to 4904

| Score | Expect | Identities | Gaps | Strand | Frame |
|---|---|---|---|---|---|
| 14.4 bits(7) | 46() | 7/7(100%) | 0/7(0%) | Plus/Plus | |

Features:

```
Query  14    TTTCTCC  20
             |||||||
Sbjct  4898  TTTCTCC  4904
```

Range 54: 5148 to 5154

| Score | Expect | Identities | Gaps | Strand | Frame |
|---|---|---|---|---|---|
| 14.4 bits(7) | 46() | 7/7(100%) | 0/7(0%) | Plus/Plus | |

Features:

```
Query  16    TCTCCTC  22
             |||||||
Sbjct  5148  TCTCCTC  5154
```

Range 55: 5560 to 5566

| Score | Expect | Identities | Gaps | Strand | Frame |
|---|---|---|---|---|---|
| 14.4 bits(7) | 46() | 7/7(100%) | 0/7(0%) | Plus/Minus | |

Features:

```
Query  10    TTGTTTT  16
             |||||||
Sbjct  5566  TTGTTTT  5560
```

Range 56: 6655 to 6661

| Score | Expect | Identities | Gaps | Strand | Frame |
|---|---|---|---|---|---|
| 14.4 bits(7) | 46() | 7/7(100%) | 0/7(0%) | Plus/Plus | |

Features:

```
Query  12    GTTTTCT  18
             |||||||
Sbjct  6655  GTTTTCT  6661
```

Range 57: 7467 to 7473

| Score | Expect | Identities | Gaps | Strand | Frame |
|---|---|---|---|---|---|
| 14.4 bits(7) | 46() | 7/7(100%) | 0/7(0%) | Plus/Minus | |

Features:

```
Query  3     CAGTCTG  9
             |||||||
Sbjct  7473  CAGTCTG  7467
```

Range 58: 7564 to 7570

| Score | Expect | Identities | Gaps | Strand | Frame |
|---|---|---|---|---|---|
| 14.4 bits(7) | 46() | 7/7(100%) | 0/7(0%) | Plus/Minus | |

Features:

```
Query  15    TTCTCCT  21
             |||||||
Sbjct  7570  TTCTCCT  7564
```

Range 59: 7638 to 7644

| Score | Expect | Identities | Gaps | Strand | Frame |
|---|---|---|---|---|---|
| 14.4 bits(7) | 46() | 7/7(100%) | 0/7(0%) | Plus/Minus | |

Features:

```
Query  7     CTGTTGT  13
             |||||||
Sbjct  7644  CTGTTGT  7638
```

Range 60: 7897 to 7903

| Score | Expect | Identities | Gaps | Strand | Frame |
|---|---|---|---|---|---|
| 14.4 bits(7) | 46() | 7/7(100%) | 0/7(0%) | Plus/Minus | |

Features:

```
Query  10    TTGTTTT  16
             |||||||
Sbjct  7903  TTGTTTT  7897
```

Range 61: 8160 to 8166

| Score | Expect | Identities | Gaps | Strand | Frame |
|---|---|---|---|---|---|
| 14.4 bits(7) | 46() | 7/7(100%) | 0/7(0%) | Plus/Minus | |

Features:

```
Query  15    TTCTCCT  21
             |||||||
Sbjct  8166  TTCTCCT  8160
```

Range 62: 8255 to 8261

| Score | Expect | Identities | Gaps | Strand | Frame |
|---|---|---|---|---|---|
| 14.4 bits(7) | 46() | 7/7(100%) | 0/7(0%) | Plus/Minus | |

Features:

```
Query  10    TTGTTTT  16
             |||||||
Sbjct  8261  TTGTTTT  8255
```

Range 63: 8317 to 8323

| Score | Expect | Identities | Gaps | Strand | Frame |
|---|---|---|---|---|---|
| 14.4 bits(7) | 46() | 7/7(100%) | 0/7(0%) | Plus/Plus | |

Features:

```
Query  10    TTGTTTT  16
             |||||||
Sbjct  8317  TTGTTTT  8323
```

Range 64: 8323 to 8329

| Score | Expect | Identities | Gaps | Strand | Frame |
|---|---|---|---|---|---|
| 14.4 bits(7) | 46() | 7/7(100%) | 0/7(0%) | Plus/Plus | |

Features:

```
Query  10    TTGTTTT  16
             |||||||
Sbjct  8323  TTGTTTT  8329
```

Range 65: 8504 to 8510

| Score | Expect | Identities | Gaps | Strand | Frame |
|---|---|---|---|---|---|
| 14.4 bits(7) | 46() | 7/7(100%) | 0/7(0%) | Plus/Plus | |

Features:

```
Query  14    TTTCTCC  20
             |||||||
Sbjct  8504  TTTCTCC  8510
```

Range 66: 8999 to 9005

| Score | Expect | Identities | Gaps | Strand | Frame |
|---|---|---|---|---|---|
| 14.4 bits(7) | 46() | 7/7(100%) | 0/7(0%) | Plus/Plus | |

Features:

```
Query  15    TTCTCCT  21
             |||||||
Sbjct  8999  TTCTCCT  9005
```

Range 67: 9238 to 9244

| Score | Expect | Identities | Gaps | Strand | Frame |
|---|---|---|---|---|---|
| 14.4 bits(7) | 46() | 7/7(100%) | 0/7(0%) | Plus/Minus | |

Features:

```
Query  10    TTGTTTT  16
             |||||||
Sbjct  9244  TTGTTTT  9238
```

Range 68: 9303 to 9309

| Score | Expect | Identities | Gaps | Strand | Frame |
|---|---|---|---|---|---|

| | | | | |
|---|---|---|---|---|
| 14.4 bits(7) | 46() | 7/7(100%) | 0/7(0%) | Plus/Plus |

Features:

```
Query  6     TCTGTTG  12
             |||||||
Sbjct  9303  TCTGTTG  9309
```

Range 69: 9375 to 9381

| Score | Expect | Identities | Gaps | Strand | Frame |
|---|---|---|---|---|---|
| 14.4 bits(7) | 46() | 7/7(100%) | 0/7(0%) | Plus/Plus | |

Features:

```
Query  15    TTCTCCT  21
             |||||||
Sbjct  9375  TTCTCCT  9381
```

Range 70: 9708 to 9714

| Score | Expect | Identities | Gaps | Strand | Frame |
|---|---|---|---|---|---|
| 14.4 bits(7) | 46() | 7/7(100%) | 0/7(0%) | Plus/Plus | |

Features:

```
Query  15    TTCTCCT  21
             |||||||
Sbjct  9708  TTCTCCT  9714
```

Range 71: 9825 to 9831

| Score | Expect | Identities | Gaps | Strand | Frame |
|---|---|---|---|---|---|
| 14.4 bits(7) | 46() | 7/7(100%) | 0/7(0%) | Plus/Plus | |

Features:

```
Query  5     GTCTGTT  11
             |||||||
Sbjct  9825  GTCTGTT  9831
```

Range 72: 9856 to 9862

| Score | Expect | Identities | Gaps | Strand | Frame |
|---|---|---|---|---|---|
| 14.4 bits(7) | 46() | 7/7(100%) | 0/7(0%) | Plus/Plus | |

Features:

```
Query  10    TTGTTTT  16
             |||||||
Sbjct  9856  TTGTTTT  9862
```

Range 73: 10433 to 10439

| Score | Expect | Identities | Gaps | Strand | Frame |
|---|---|---|---|---|---|
| 14.4 bits(7) | 46() | 7/7(100%) | 0/7(0%) | Plus/Minus | |

Features:

```
Query  15     TTCTCCT  21
              |||||||
Sbjct  10439  TTCTCCT  10433
```

Range 74: 10569 to 10575

| Score | Expect | Identities | Gaps | Strand | Frame |
|---|---|---|---|---|---|
| 14.4 bits(7) | 46() | 7/7(100%) | 0/7(0%) | Plus/Plus | |

Features:

```
Query  10     TTGTTTT  16
              |||||||
Sbjct  10569  TTGTTTT  10575
```

Range 75: 10849 to 10855

| Score | Expect | Identities | Gaps | Strand | Frame |
|---|---|---|---|---|---|
| 14.4 bits(7) | 46() | 7/7(100%) | 0/7(0%) | Plus/Plus | |

Features:

```
Query  15     TTCTCCT  21
              |||||||
Sbjct  10849  TTCTCCT  10855
```

Range 76: 11978 to 11984

| Score | Expect | Identities | Gaps | Strand | Frame |
|---|---|---|---|---|---|
| 14.4 bits(7) | 46() | 7/7(100%) | 0/7(0%) | Plus/Plus | |

Features:

```
Query  15     TTCTCCT  21
              |||||||
Sbjct  11978  TTCTCCT  11984
```

Range 77: 12282 to 12288

| Score | Expect | Identities | Gaps | Strand | Frame |
|---|---|---|---|---|---|
| 14.4 bits(7) | 46() | 7/7(100%) | 0/7(0%) | Plus/Plus | |

Features:

```
Query  15     TTCTCCT  21
              |||||||
Sbjct  12282  TTCTCCT  12288
```

Range 78: 12582 to 12588

| Score | Expect | Identities | Gaps | Strand | Frame |
|---|---|---|---|---|---|
| 14.4 bits(7) | 46() | 7/7(100%) | 0/7(0%) | Plus/Plus | |

Features:

```
Query  15     TTCTCCT  21
              |||||||
Sbjct  12582  TTCTCCT  12588
```

Range 79: 13097 to 13103

| Score | Expect | Identities | Gaps | Strand | Frame |
|---|---|---|---|---|---|
| 14.4 bits(7) | 46() | 7/7(100%) | 0/7(0%) | Plus/Minus | |

Features:

```
Query  10     TTGTTTT  16
              |||||||
Sbjct  13103  TTGTTTT  13097
```

Range 80: 13537 to 13543

| Score | Expect | Identities | Gaps | Strand | Frame |
|---|---|---|---|---|---|
| 14.4 bits(7) | 46() | 7/7(100%) | 0/7(0%) | Plus/Minus | |

Features:

```
Query  10     TTGTTTT  16
              |||||||
Sbjct  13543  TTGTTTT  13537
```

Range 81: 13674 to 13680

| Score | Expect | Identities | Gaps | Strand | Frame |
|---|---|---|---|---|---|
| 14.4 bits(7) | 46() | 7/7(100%) | 0/7(0%) | Plus/Plus | |

Features:

```
Query  10     TTGTTTT  16
              |||||||
Sbjct  13674  TTGTTTT  13680
```

Range 82: 14130 to 14136

| Score | Expect | Identities | Gaps | Strand | Frame |
|---|---|---|---|---|---|
| 14.4 bits(7) | 46() | 7/7(100%) | 0/7(0%) | Plus/Minus | |

Features:

```
Query  6      TCTGTTG  12
              |||||||
Sbjct  14136  TCTGTTG  14130
```

Range 83: 14547 to 14553

| Score | Expect | Identities | Gaps | Strand | Frame |
|---|---|---|---|---|---|
| 14.4 bits(7) | 46() | 7/7(100%) | 0/7(0%) | Plus/Plus | |

Features:

```
Query  1      AACAGTC  7
              |||||||
Sbjct  14547  AACAGTC  14553
```

Range 84: 14626 to 14632

| Score | Expect | Identities | Gaps | Strand | Frame |
|---|---|---|---|---|---|
| 14.4 bits(7) | 46() | 7/7(100%) | 0/7(0%) | Plus/Minus | |

Features:

```
Query  11     TGTTTTC  17
              |||||||
Sbjct  14632  TGTTTTC  14626
```

Range 85: 15354 to 15360

| Score | Expect | Identities | Gaps | Strand | Frame |
|---|---|---|---|---|---|
| 14.4 bits(7) | 46() | 7/7(100%) | 0/7(0%) | Plus/Minus | |

Features:

```
Query  13     TTTTCTC  19
              |||||||
Sbjct  15360  TTTTCTC  15354
```

Range 86: 15560 to 15566

| Score | Expect | Identities | Gaps | Strand | Frame |
|---|---|---|---|---|---|
| 14.4 bits(7) | 46() | 7/7(100%) | 0/7(0%) | Plus/Plus | |

Features:

```
Query  3      CAGTCTG  9
              |||||||
Sbjct  15560  CAGTCTG  15566
```

Range 87: 15606 to 15612

| Score | Expect | Identities | Gaps | Strand | Frame |
|---|---|---|---|---|---|

| Score | Expect | Identities | Gaps | Strand | Frame |
|---|---|---|---|---|---|
| 14.4 bits(7) | 46() | 7/7(100%) | 0/7(0%) | Plus/Plus | |

Features:

```
Query  9      GTTGTTT  15
              |||||||
Sbjct  15606  GTTGTTT  15612
```

Range 88: 15851 to 15857

| Score | Expect | Identities | Gaps | Strand | Frame |
|---|---|---|---|---|---|
| 14.4 bits(7) | 46() | 7/7(100%) | 0/7(0%) | Plus/Plus | |

Features:

```
Query  15     TTCTCCT  21
              |||||||
Sbjct  15851  TTCTCCT  15857
```

Range 89: 16464 to 16470

| Score | Expect | Identities | Gaps | Strand | Frame |
|---|---|---|---|---|---|
| 14.4 bits(7) | 46() | 7/7(100%) | 0/7(0%) | Plus/Plus | |

Features:

```
Query  10     TTGTTTT  16
              |||||||
Sbjct  16464  TTGTTTT  16470
```

Range 90: 16472 to 16478

| Score | Expect | Identities | Gaps | Strand | Frame |
|---|---|---|---|---|---|
| 14.4 bits(7) | 46() | 7/7(100%) | 0/7(0%) | Plus/Plus | |

Features:

```
Query  10     TTGTTTT  16
              |||||||
Sbjct  16472  TTGTTTT  16478
```

Range 91: 17397 to 17403

| Score | Expect | Identities | Gaps | Strand | Frame |
|---|---|---|---|---|---|
| 14.4 bits(7) | 46() | 7/7(100%) | 0/7(0%) | Plus/Minus | |

Features:

```
Query  15     TTCTCCT  21
              |||||||
Sbjct  17403  TTCTCCT  17397
```

Range 92: 19431 to 19437

| Score | Expect | Identities | Gaps | Strand | Frame |
|---|---|---|---|---|---|
| 14.4 bits(7) | 46() | 7/7(100%) | 0/7(0%) | Plus/Plus | |

Features:

```
Query  10     TTGTTTT  16
              |||||||
Sbjct  19431  TTGTTTT  19437
```

Range 93: 19531 to 19537

| Score | Expect | Identities | Gaps | Strand | Frame |
|---|---|---|---|---|---|
| 14.4 bits(7) | 46() | 7/7(100%) | 0/7(0%) | Plus/Plus | |

Features:

```
Query  6      TCTGTTG  12
              |||||||
Sbjct  19531  TCTGTTG  19537
```

Range 94: 19603 to 19609

| Score | Expect | Identities | Gaps | Strand | Frame |
|---|---|---|---|---|---|
| 14.4 bits(7) | 46() | 7/7(100%) | 0/7(0%) | Plus/Plus | |

Features:

```
Query  15     TTCTCCT  21
              |||||||
Sbjct  19603  TTCTCCT  19609
```

Range 95: 19931 to 19937

| Score | Expect | Identities | Gaps | Strand | Frame |
|---|---|---|---|---|---|
| 14.4 bits(7) | 46() | 7/7(100%) | 0/7(0%) | Plus/Plus | |

Features:

```
Query  15     TTCTCCT  21
              |||||||
Sbjct  19931  TTCTCCT  19937
```

Range 96: 20826 to 20832

| Score | Expect | Identities | Gaps | Strand | Frame |
|---|---|---|---|---|---|
| 14.4 bits(7) | 46() | 7/7(100%) | 0/7(0%) | Plus/Plus | |

Features:

```
Query  10     TTGTTTT  16
              |||||||
Sbjct  20826  TTGTTTT  20832
```

Range 97: 20851 to 20857

| Score | Expect | Identities | Gaps | Strand | Frame |
|---|---|---|---|---|---|
| 14.4 bits(7) | 46() | 7/7(100%) | 0/7(0%) | Plus/Plus | |

Features:

```
Query  7      CTGTTGT  13
              |||||||
Sbjct  20851  CTGTTGT  20857
```

Range 98: 20924 to 20930

| Score | Expect | Identities | Gaps | Strand | Frame |
|---|---|---|---|---|---|
| 14.4 bits(7) | 46() | 7/7(100%) | 0/7(0%) | Plus/Plus | |

Features:

```
Query  3      CAGTCTG  9
              |||||||
Sbjct  20924  CAGTCTG  20930
```

Range 99: 21081 to 21087

| Score | Expect | Identities | Gaps | Strand | Frame |
|---|---|---|---|---|---|
| 14.4 bits(7) | 46() | 7/7(100%) | 0/7(0%) | Plus/Minus | |

Features:

```
Query  10     TTGTTTT  16
              |||||||
Sbjct  21087  TTGTTTT  21081
```

Range 100: 21343 to 21349

| Score | Expect | Identities | Gaps | Strand | Frame |
|---|---|---|---|---|---|
| 14.4 bits(7) | 46() | 7/7(100%) | 0/7(0%) | Plus/Minus | |

Features:

```
Query  12     GTTTTCT  18
              |||||||
Sbjct  21349  GTTTTCT  21343
```

Range 101: 21486 to 21492

| Score | Expect | Identities | Gaps | Strand | Frame |
|---|---|---|---|---|---|
| 14.4 bits(7) | 46() | 7/7(100%) | 0/7(0%) | Plus/Minus | |

Features:

```
Query  6      TCTGTTG  12
              |||||||
Sbjct  21492  TCTGTTG  21486
```

Range 102: 22387 to 22393

| Score | Expect | Identities | Gaps | Strand | Frame |
|---|---|---|---|---|---|
| 14.4 bits(7) | 46() | 7/7(100%) | 0/7(0%) | Plus/Minus | |

Features:

```
Query  6      TCTGTTG  12
              |||||||
Sbjct  22393  TCTGTTG  22387
```

Range 103: 22396 to 22402

| Score | Expect | Identities | Gaps | Strand | Frame |
|---|---|---|---|---|---|
| 14.4 bits(7) | 46() | 7/7(100%) | 0/7(0%) | Plus/Minus | |

Features:

```
Query  8      TGTTGTT  14
              |||||||
Sbjct  22402  TGTTGTT  22396
```

Range 104: 22469 to 22475

| Score | Expect | Identities | Gaps | Strand | Frame |
|---|---|---|---|---|---|
| 14.4 bits(7) | 46() | 7/7(100%) | 0/7(0%) | Plus/Plus | |

Features:

```
Query  13     TTTTCTC  19
              |||||||
Sbjct  22469  TTTTCTC  22475
```

Range 105: 22958 to 22964

| Score | Expect | Identities | Gaps | Strand | Frame |
|---|---|---|---|---|---|
| 14.4 bits(7) | 46() | 7/7(100%) | 0/7(0%) | Plus/Plus | |

Features:

```
Query  10     TTGTTTT  16
              |||||||
Sbjct  22958  TTGTTTT  22964
```

Range 106: 23164 to 23170

| Score | Expect | Identities | Gaps | Strand | Frame |
|---|---|---|---|---|---|

| Score | Expect | Identities | Gaps | Strand | Frame |
|---|---|---|---|---|---|
| 14.4 bits(7) | 46() | 7/7(100%) | 0/7(0%) | Plus/Minus | |

Features:

```
Query  9      GTTGTTT  15
              |||||||
Sbjct  23170  GTTGTTT  23164
```

Range 107: 23725 to 23731

| Score | Expect | Identities | Gaps | Strand | Frame |
|---|---|---|---|---|---|
| 14.4 bits(7) | 46() | 7/7(100%) | 0/7(0%) | Plus/Minus | |

Features:

```
Query  1      AACAGTC  7
              |||||||
Sbjct  23731  AACAGTC  23725
```

Range 108: 23745 to 23751

| Score | Expect | Identities | Gaps | Strand | Frame |
|---|---|---|---|---|---|
| 14.4 bits(7) | 46() | 7/7(100%) | 0/7(0%) | Plus/Plus | |

Features:

```
Query  15     TTCTCCT  21
              |||||||
Sbjct  23745  TTCTCCT  23751
```

Range 109: 23803 to 23809

| Score | Expect | Identities | Gaps | Strand | Frame |
|---|---|---|---|---|---|
| 14.4 bits(7) | 46() | 7/7(100%) | 0/7(0%) | Plus/Plus | |

Features:

```
Query  5      GTCTGTT  11
              |||||||
Sbjct  23803  GTCTGTT  23809
```

Range 110: 24497 to 24503

| Score | Expect | Identities | Gaps | Strand | Frame |
|---|---|---|---|---|---|
| 14.4 bits(7) | 46() | 7/7(100%) | 0/7(0%) | Plus/Plus | |

Features:

```
Query  13     TTTTCTC  19
              |||||||
Sbjct  24497  TTTTCTC  24503
```

Range 111: 24922 to 24928

| Score | Expect | Identities | Gaps | Strand | Frame |
|---|---|---|---|---|---|
| 14.4 bits(7) | 46() | 7/7(100%) | 0/7(0%) | Plus/Minus | |

Features:

```
Query  12     GTTTTCT  18
              |||||||
Sbjct  24928  GTTTTCT  24922
```

Range 112: 25230 to 25236

| Score | Expect | Identities | Gaps | Strand | Frame |
|---|---|---|---|---|---|
| 14.4 bits(7) | 46() | 7/7(100%) | 0/7(0%) | Plus/Minus | |

Features:

```
Query  12     GTTTTCT  18
              |||||||
Sbjct  25236  GTTTTCT  25230
```

Range 113: 25446 to 25452

| Score | Expect | Identities | Gaps | Strand | Frame |
|---|---|---|---|---|---|
| 14.4 bits(7) | 46() | 7/7(100%) | 0/7(0%) | Plus/Minus | |

Features:

```
Query  15     TTCTCCT  21
              |||||||
Sbjct  25452  TTCTCCT  25446
```

Range 114: 25561 to 25567

| Score | Expect | Identities | Gaps | Strand | Frame |
|---|---|---|---|---|---|
| 14.4 bits(7) | 46() | 7/7(100%) | 0/7(0%) | Plus/Minus | |

Features:

```
Query  14     TTTCTCC  20
              |||||||
Sbjct  25567  TTTCTCC  25561
```

Range 115: 25672 to 25678

| Score | Expect | Identities | Gaps | Strand | Frame |
|---|---|---|---|---|---|
| 14.4 bits(7) | 46() | 7/7(100%) | 0/7(0%) | Plus/Plus | |

Features:

```
Query  17     CTCCTCT  23
              |||||||
Sbjct  25672  CTCCTCT  25678
```

Range 116: 25926 to 25932

| Score | Expect | Identities | Gaps | Strand | Frame |
|---|---|---|---|---|---|
| 14.4 bits(7) | 46() | 7/7(100%) | 0/7(0%) | Plus/Plus | |

Features:

```
Query  13     TTTTCTC  19
              |||||||
Sbjct  25926  TTTTCTC  25932
```

Range 117: 26400 to 26406

| Score | Expect | Identities | Gaps | Strand | Frame |
|---|---|---|---|---|---|
| 14.4 bits(7) | 46() | 7/7(100%) | 0/7(0%) | Plus/Minus | |

Features:

```
Query  9      GTTGTTT  15
              |||||||
Sbjct  26406  GTTGTTT  26400
```

Range 118: 26450 to 26456

| Score | Expect | Identities | Gaps | Strand | Frame |
|---|---|---|---|---|---|
| 14.4 bits(7) | 46() | 7/7(100%) | 0/7(0%) | Plus/Plus | |

Features:

```
Query  15     TTCTCCT  21
              |||||||
Sbjct  26450  TTCTCCT  26456
```

Range 119: 27429 to 27435

| Score | Expect | Identities | Gaps | Strand | Frame |
|---|---|---|---|---|---|
| 14.4 bits(7) | 46() | 7/7(100%) | 0/7(0%) | Plus/Plus | |

Features:

```
Query  3      CAGTCTG  9
              |||||||
Sbjct  27429  CAGTCTG  27435
```

Range 120: 27578 to 27584

| Score | Expect | Identities | Gaps | Strand | Frame |
|---|---|---|---|---|---|
| 14.4 bits(7) | 46() | 7/7(100%) | 0/7(0%) | Plus/Minus | |

Features:

```
Query  6      TCTGTTG  12
              |||||||
Sbjct  27584  TCTGTTG  27578
```

Range 121: 27595 to 27601

| Score | Expect | Identities | Gaps | Strand | Frame |
|---|---|---|---|---|---|
| 14.4 bits(7) | 46() | 7/7(100%) | 0/7(0%) | Plus/Plus | |

Features:

```
Query  10     TTGTTTT  16
              |||||||
Sbjct  27595  TTGTTTT  27601
```

Range 122: 27687 to 27693

| Score | Expect | Identities | Gaps | Strand | Frame |
|---|---|---|---|---|---|
| 14.4 bits(7) | 46() | 7/7(100%) | 0/7(0%) | Plus/Plus | |

Features:

```
Query  15     TTCTCCT  21
              |||||||
Sbjct  27687  TTCTCCT  27693
```

Range 123: 27791 to 27797

| Score | Expect | Identities | Gaps | Strand | Frame |
|---|---|---|---|---|---|
| 14.4 bits(7) | 46() | 7/7(100%) | 0/7(0%) | Plus/Plus | |

Features:

```
Query  14     TTTCTCC  20
              |||||||
Sbjct  27791  TTTCTCC  27797
```

Range 124: 28405 to 28411

| Score | Expect | Identities | Gaps | Strand | Frame |
|---|---|---|---|---|---|
| 14.4 bits(7) | 46() | 7/7(100%) | 0/7(0%) | Plus/Plus | |

Features:

```
Query  15     TTCTCCT  21
              |||||||
Sbjct  28405  TTCTCCT  28411
```

Range 125: 28492 to 28498

| Score | Expect | Identities | Gaps | Strand | Frame |
|---|---|---|---|---|---|

| 14.4 bits(7) | 46() | 7/7(100%) | 0/7(0%) | Plus/Plus |
|---|---|---|---|---|

Features:

```
Query  14     TTTCTCC  20
              |||||||
Sbjct  28492  TTTCTCC  28498
```

Range 126: 28748 to 28754

| Score | Expect | Identities | Gaps | Strand | Frame |
|---|---|---|---|---|---|
| 14.4 bits(7) | 46() | 7/7(100%) | 0/7(0%) | Plus/Plus | |

Features:

```
Query  11     TGTTTTC  17
              |||||||
Sbjct  28748  TGTTTTC  28754
```

Range 127: 29105 to 29111

| Score | Expect | Identities | Gaps | Strand | Frame |
|---|---|---|---|---|---|
| 14.4 bits(7) | 46() | 7/7(100%) | 0/7(0%) | Plus/Minus | |

Features:

```
Query  14     TTTCTCC  20
              |||||||
Sbjct  29111  TTTCTCC  29105
```

Range 128: 29480 to 29486

| Score | Expect | Identities | Gaps | Strand | Frame |
|---|---|---|---|---|---|
| 14.4 bits(7) | 46() | 7/7(100%) | 0/7(0%) | Plus/Plus | |

Features:

```
Query  7      CTGTTGT  13
              |||||||
Sbjct  29480  CTGTTGT  29486
```

Range 129: 31151 to 31157

| Score | Expect | Identities | Gaps | Strand | Frame |
|---|---|---|---|---|---|
| 14.4 bits(7) | 46() | 7/7(100%) | 0/7(0%) | Plus/Plus | |

Features:

```
Query  10     TTGTTTT  16
              |||||||
Sbjct  31151  TTGTTTT  31157
```

Range 130: 31571 to 31577

| Score | Expect | Identities | Gaps | Strand | Frame |
|---|---|---|---|---|---|
| 14.4 bits(7) | 46() | 7/7(100%) | 0/7(0%) | Plus/Plus | |

Features:

```
Query  10     TTGTTTT  16
              |||||||
Sbjct  31571  TTGTTTT  31577
```

Range 131: 31700 to 31706

| Score | Expect | Identities | Gaps | Strand | Frame |
|---|---|---|---|---|---|
| 14.4 bits(7) | 46() | 7/7(100%) | 0/7(0%) | Plus/Plus | |

Features:

```
Query  13     TTTTCTC  19
              |||||||
Sbjct  31700  TTTTCTC  31706
```

Range 132: 31732 to 31738

| Score | Expect | Identities | Gaps | Strand | Frame |
|---|---|---|---|---|---|
| 14.4 bits(7) | 46() | 7/7(100%) | 0/7(0%) | Plus/Plus | |

Features:

```
Query  10     TTGTTTT  16
              |||||||
Sbjct  31732  TTGTTTT  31738
```

Range 133: 31828 to 31834

| Score | Expect | Identities | Gaps | Strand | Frame |
|---|---|---|---|---|---|
| 14.4 bits(7) | 46() | 7/7(100%) | 0/7(0%) | Plus/Plus | |

Features:

```
Query  15     TTCTCCT  21
              |||||||
Sbjct  31828  TTCTCCT  31834
```

Range 134: 32430 to 32436

| Score | Expect | Identities | Gaps | Strand | Frame |
|---|---|---|---|---|---|
| 14.4 bits(7) | 46() | 7/7(100%) | 0/7(0%) | Plus/Plus | |

Features:

```
Query  10     TTGTTTT  16
              |||||||
Sbjct  32430  TTGTTTT  32436
```

Range 135: 32693 to 32699

| Score | Expect | Identities | Gaps | Strand | Frame |
|---|---|---|---|---|---|
| 14.4 bits(7) | 46() | 7/7(100%) | 0/7(0%) | Plus/Minus | |

Features:

```
Query  15     TTCTCCT  21
              |||||||
Sbjct  32699  TTCTCCT  32693
```

Range 136: 33044 to 33050

| Score | Expect | Identities | Gaps | Strand | Frame |
|---|---|---|---|---|---|
| 14.4 bits(7) | 46() | 7/7(100%) | 0/7(0%) | Plus/Minus | |

Features:

```
Query  15     TTCTCCT  21
              |||||||
Sbjct  33050  TTCTCCT  33044
```

Range 137: 33302 to 33308

| Score | Expect | Identities | Gaps | Strand | Frame |
|---|---|---|---|---|---|
| 14.4 bits(7) | 46() | 7/7(100%) | 0/7(0%) | Plus/Plus | |

Features:

```
Query  10     TTGTTTT  16
              |||||||
Sbjct  33302  TTGTTTT  33308
```

Range 138: 33411 to 33417

| Score | Expect | Identities | Gaps | Strand | Frame |
|---|---|---|---|---|---|
| 14.4 bits(7) | 46() | 7/7(100%) | 0/7(0%) | Plus/Plus | |

Features:

```
Query  11     TGTTTTC  17
              |||||||
Sbjct  33411  TGTTTTC  33417
```

Range 139: 33497 to 33503

| Score | Expect | Identities | Gaps | Strand | Frame |
|---|---|---|---|---|---|
| 14.4 bits(7) | 46() | 7/7(100%) | 0/7(0%) | Plus/Minus | |

Features:

```
Query  8      TGTTGTT  14
              |||||||
Sbjct  33503  TGTTGTT  33497
```

Range 140: 33928 to 33934

| Score | Expect | Identities | Gaps | Strand | Frame |
|---|---|---|---|---|---|
| 14.4 bits(7) | 46() | 7/7(100%) | 0/7(0%) | Plus/Plus | |

Features:

```
Query  6      TCTGTTG  12
              |||||||
Sbjct  33928  TCTGTTG  33934
```

Range 141: 33999 to 34005

| Score | Expect | Identities | Gaps | Strand | Frame |
|---|---|---|---|---|---|
| 14.4 bits(7) | 46() | 7/7(100%) | 0/7(0%) | Plus/Plus | |

Features:

```
Query  15     TTCTCCT  21
              |||||||
Sbjct  33999  TTCTCCT  34005
```

Range 142: 34030 to 34036

| Score | Expect | Identities | Gaps | Strand | Frame |
|---|---|---|---|---|---|
| 14.4 bits(7) | 46() | 7/7(100%) | 0/7(0%) | Plus/Minus | |

Features:

```
Query  7      CTGTTGT  13
              |||||||
Sbjct  34036  CTGTTGT  34030
```

Range 143: 34426 to 34432

| Score | Expect | Identities | Gaps | Strand | Frame |
|---|---|---|---|---|---|
| 14.4 bits(7) | 46() | 7/7(100%) | 0/7(0%) | Plus/Plus | |

Features:

```
Query  15     TTCTCCT  21
              |||||||
Sbjct  34426  TTCTCCT  34432
```

Range 144: 34670 to 34676

| Score | Expect | Identities | Gaps | Strand | Frame |
|---|---|---|---|---|---|

| Score | Expect | Identities | Gaps | Strand | Frame |
|---|---|---|---|---|---|
| 14.4 bits(7) | 46() | 7/7(100%) | 0/7(0%) | Plus/Plus | |

Features:

```
Query  8      TGTTGTT  14
              |||||||
Sbjct  34670  TGTTGTT  34676
```

Range 145: 34925 to 34931

| Score | Expect | Identities | Gaps | Strand | Frame |
|---|---|---|---|---|---|
| 14.4 bits(7) | 46() | 7/7(100%) | 0/7(0%) | Plus/Plus | |

Features:

```
Query  15     TTCTCCT  21
              |||||||
Sbjct  34925  TTCTCCT  34931
```

Range 146: 35650 to 35656

| Score | Expect | Identities | Gaps | Strand | Frame |
|---|---|---|---|---|---|
| 14.4 bits(7) | 46() | 7/7(100%) | 0/7(0%) | Plus/Minus | |

Features:

```
Query  15     TTCTCCT  21
              |||||||
Sbjct  35656  TTCTCCT  35650
```

Range 147: 35789 to 35795

| Score | Expect | Identities | Gaps | Strand | Frame |
|---|---|---|---|---|---|
| 14.4 bits(7) | 46() | 7/7(100%) | 0/7(0%) | Plus/Plus | |

Features:

```
Query  10     TTGTTTT  16
              |||||||
Sbjct  35789  TTGTTTT  35795
```

Range 148: 36203 to 36209

| Score | Expect | Identities | Gaps | Strand | Frame |
|---|---|---|---|---|---|
| 14.4 bits(7) | 46() | 7/7(100%) | 0/7(0%) | Plus/Plus | |

Features:

```
Query  10     TTGTTTT  16
              |||||||
Sbjct  36203  TTGTTTT  36209
```

Range 149: 36296 to 36302

| Score | Expect | Identities | Gaps | Strand | Frame |
|---|---|---|---|---|---|
| 14.4 bits(7) | 46() | 7/7(100%) | 0/7(0%) | Plus/Minus | |

Features:

```
Query  10     TTGTTTT  16
              |||||||
Sbjct  36302  TTGTTTT  36296
```

Range 150: 37076 to 37082

| Score | Expect | Identities | Gaps | Strand | Frame |
|---|---|---|---|---|---|
| 14.4 bits(7) | 46() | 7/7(100%) | 0/7(0%) | Plus/Minus | |

Features:

```
Query  15     TTCTCCT  21
              |||||||
Sbjct  37082  TTCTCCT  37076
```

Range 151: 37657 to 37663

| Score | Expect | Identities | Gaps | Strand | Frame |
|---|---|---|---|---|---|
| 14.4 bits(7) | 46() | 7/7(100%) | 0/7(0%) | Plus/Plus | |

Features:

```
Query  6      TCTGTTG  12
              |||||||
Sbjct  37657  TCTGTTG  37663
```

Range 152: 37729 to 37735

| Score | Expect | Identities | Gaps | Strand | Frame |
|---|---|---|---|---|---|
| 14.4 bits(7) | 46() | 7/7(100%) | 0/7(0%) | Plus/Plus | |

Features:

```
Query  15     TTCTCCT  21
              |||||||
Sbjct  37729  TTCTCCT  37735
```

Range 153: 38232 to 38238

| Score | Expect | Identities | Gaps | Strand | Frame |
|---|---|---|---|---|---|
| 14.4 bits(7) | 46() | 7/7(100%) | 0/7(0%) | Plus/Minus | |

Features:

```
Query  15     TTCTCCT  21
              |||||||
Sbjct  38238  TTCTCCT  38232
```

Range 154: 38332 to 38338

| Score | Expect | Identities | Gaps | Strand | Frame |
|---|---|---|---|---|---|
| 14.4 bits(7) | 46() | 7/7(100%) | 0/7(0%) | Plus/Minus | |

Features:

```
Query  10     TTGTTTT  16
              |||||||
Sbjct  38338  TTGTTTT  38332
```

Range 155: 38839 to 38845

| Score | Expect | Identities | Gaps | Strand | Frame |
|---|---|---|---|---|---|
| 14.4 bits(7) | 46() | 7/7(100%) | 0/7(0%) | Plus/Plus | |

Features:

```
Query  12     GTTTTCT  18
              |||||||
Sbjct  38839  GTTTTCT  38845
```

Range 156: 39477 to 39483

| Score | Expect | Identities | Gaps | Strand | Frame |
|---|---|---|---|---|---|
| 14.4 bits(7) | 46() | 7/7(100%) | 0/7(0%) | Plus/Minus | |

Features:

```
Query  15     TTCTCCT  21
              |||||||
Sbjct  39483  TTCTCCT  39477
```

Range 157: 39926 to 39932

| Score | Expect | Identities | Gaps | Strand | Frame |
|---|---|---|---|---|---|
| 14.4 bits(7) | 46() | 7/7(100%) | 0/7(0%) | Plus/Plus | |

Features:

```
Query  1      AACAGTC  7
              |||||||
Sbjct  39926  AACAGTC  39932
```

Range 158: 40390 to 40396

| Score | Expect | Identities | Gaps | Strand | Frame |
|---|---|---|---|---|---|
| 14.4 bits(7) | 46() | 7/7(100%) | 0/7(0%) | Plus/Plus | |

Features:

```
Query  13     TTTTCTC  19
              |||||||
Sbjct  40390  TTTTCTC  40396
```

Range 159: 40465 to 40471

| Score | Expect | Identities | Gaps | Strand | Frame |
|---|---|---|---|---|---|
| 14.4 bits(7) | 46() | 7/7(100%) | 0/7(0%) | Plus/Plus | |

Features:

```
Query  10     TTGTTTT  16
              |||||||
Sbjct  40465  TTGTTTT  40471
```

Range 160: 40547 to 40553

| Score | Expect | Identities | Gaps | Strand | Frame |
|---|---|---|---|---|---|
| 14.4 bits(7) | 46() | 7/7(100%) | 0/7(0%) | Plus/Plus | |

Features:

```
Query  5      GTCTGTT  11
              |||||||
Sbjct  40547  GTCTGTT  40553
```

Range 161: 41417 to 41423

| Score | Expect | Identities | Gaps | Strand | Frame |
|---|---|---|---|---|---|
| 14.4 bits(7) | 46() | 7/7(100%) | 0/7(0%) | Plus/Plus | |

Features:

```
Query  6      TCTGTTG  12
              |||||||
Sbjct  41417  TCTGTTG  41423
```

Range 162: 41426 to 41432

| Score | Expect | Identities | Gaps | Strand | Frame |
|---|---|---|---|---|---|
| 14.4 bits(7) | 46() | 7/7(100%) | 0/7(0%) | Plus/Plus | |

Features:

```
Query  3      CAGTCTG  9
              |||||||
Sbjct  41426  CAGTCTG  41432
```

Range 163: 41489 to 41495

| Score | Expect | Identities | Gaps | Strand | Frame |
|---|---|---|---|---|---|

| Score | Expect | Identities | Gaps | Strand | Frame |
|---|---|---|---|---|---|
| 14.4 bits(7) | 46() | 7/7(100%) | 0/7(0%) | Plus/Plus | |

Features:

```
Query  15     TTCTCCT  21
              |||||||
Sbjct  41489  TTCTCCT  41495
```

Range 164: 41840 to 41846

| Score | Expect | Identities | Gaps | Strand | Frame |
|---|---|---|---|---|---|
| 14.4 bits(7) | 46() | 7/7(100%) | 0/7(0%) | Plus/Plus | |

Features:

```
Query  15     TTCTCCT  21
              |||||||
Sbjct  41840  TTCTCCT  41846
```

Range 165: 41927 to 41933

| Score | Expect | Identities | Gaps | Strand | Frame |
|---|---|---|---|---|---|
| 14.4 bits(7) | 46() | 7/7(100%) | 0/7(0%) | Plus/Plus | |

Features:

```
Query  14     TTTCTCC  20
              |||||||
Sbjct  41927  TTTCTCC  41933
```

Range 166: 42254 to 42260

| Score | Expect | Identities | Gaps | Strand | Frame |
|---|---|---|---|---|---|
| 14.4 bits(7) | 46() | 7/7(100%) | 0/7(0%) | Plus/Plus | |

Features:

```
Query  10     TTGTTTT  16
              |||||||
Sbjct  42254  TTGTTTT  42260
```

Range 167: 42471 to 42477

| Score | Expect | Identities | Gaps | Strand | Frame |
|---|---|---|---|---|---|
| 14.4 bits(7) | 46() | 7/7(100%) | 0/7(0%) | Plus/Plus | |

Features:

```
Query  15     TTCTCCT  21
              |||||||
Sbjct  42471  TTCTCCT  42477
```

Range 168: 42748 to 42754

| Score | Expect | Identities | Gaps | Strand | Frame |
|---|---|---|---|---|---|
| 14.4 bits(7) | 46() | 7/7(100%) | 0/7(0%) | Plus/Minus | |

Features:

```
Query  10     TTGTTTT  16
              |||||||
Sbjct  42754  TTGTTTT  42748
```

Range 169: 42765 to 42771

| Score | Expect | Identities | Gaps | Strand | Frame |
|---|---|---|---|---|---|
| 14.4 bits(7) | 46() | 7/7(100%) | 0/7(0%) | Plus/Minus | |

Features:

```
Query  10     TTGTTTT  16
              |||||||
Sbjct  42771  TTGTTTT  42765
```

Range 170: 42781 to 42787

| Score | Expect | Identities | Gaps | Strand | Frame |
|---|---|---|---|---|---|
| 14.4 bits(7) | 46() | 7/7(100%) | 0/7(0%) | Plus/Minus | |

Features:

```
Query  15     TTCTCCT  21
              |||||||
Sbjct  42787  TTCTCCT  42781
```

Range 171: 42855 to 42861

| Score | Expect | Identities | Gaps | Strand | Frame |
|---|---|---|---|---|---|
| 14.4 bits(7) | 46() | 7/7(100%) | 0/7(0%) | Plus/Minus | |

Features:

```
Query  3      CAGTCTG  9
              |||||||
Sbjct  42861  CAGTCTG  42855
```

Range 172: 42934 to 42940

| Score | Expect | Identities | Gaps | Strand | Frame |
|---|---|---|---|---|---|
| 14.4 bits(7) | 46() | 7/7(100%) | 0/7(0%) | Plus/Plus | |

Features:

```
Query  8      TGTTGTT  14
              |||||||
Sbjct  42934  TGTTGTT  42940
```

Range 173: 43062 to 43068

| Score | Expect | Identities | Gaps | Strand | Frame |
|---|---|---|---|---|---|
| 14.4 bits(7) | 46() | 7/7(100%) | 0/7(0%) | Plus/Plus | |

Features:

```
Query  10     TTGTTTT  16
              |||||||
Sbjct  43062  TTGTTTT  43068
```

Range 174: 43126 to 43132

| Score | Expect | Identities | Gaps | Strand | Frame |
|---|---|---|---|---|---|
| 14.4 bits(7) | 46() | 7/7(100%) | 0/7(0%) | Plus/Minus | |

Features:

```
Query  6      TCTGTTG  12
              |||||||
Sbjct  43132  TCTGTTG  43126
```

Range 175: 43151 to 43157

| Score | Expect | Identities | Gaps | Strand | Frame |
|---|---|---|---|---|---|
| 14.4 bits(7) | 46() | 7/7(100%) | 0/7(0%) | Plus/Plus | |

Features:

```
Query  10     TTGTTTT  16
              |||||||
Sbjct  43151  TTGTTTT  43157
```

Range 176: 43161 to 43167

| Score | Expect | Identities | Gaps | Strand | Frame |
|---|---|---|---|---|---|
| 14.4 bits(7) | 46() | 7/7(100%) | 0/7(0%) | Plus/Plus | |

Features:

```
Query  10     TTGTTTT  16
              |||||||
Sbjct  43161  TTGTTTT  43167
```

Range 177: 43166 to 43172

| Score | Expect | Identities | Gaps | Strand | Frame |
|---|---|---|---|---|---|
| 14.4 bits(7) | 46() | 7/7(100%) | 0/7(0%) | Plus/Plus | |

Features:

```
Query  10     TTGTTTT  16
              |||||||
Sbjct  43166  TTGTTTT  43172
```

Range 178: 43174 to 43180

| Score | Expect | Identities | Gaps | Strand | Frame |
|---|---|---|---|---|---|
| 14.4 bits(7) | 46() | 7/7(100%) | 0/7(0%) | Plus/Plus | |

Features:

```
Query  10     TTGTTTT  16
              |||||||
Sbjct  43174  TTGTTTT  43180
```

Range 179: 43271 to 43277

| Score | Expect | Identities | Gaps | Strand | Frame |
|---|---|---|---|---|---|
| 14.4 bits(7) | 46() | 7/7(100%) | 0/7(0%) | Plus/Plus | |

Features:

```
Query  15     TTCTCCT  21
              |||||||
Sbjct  43271  TTCTCCT  43277
```

Range 180: 43595 to 43601

| Score | Expect | Identities | Gaps | Strand | Frame |
|---|---|---|---|---|---|
| 14.4 bits(7) | 46() | 7/7(100%) | 0/7(0%) | Plus/Plus | |

Features:

```
Query  17     CTCCTCT  23
              |||||||
Sbjct  43595  CTCCTCT  43601
```

Range 181: 44155 to 44161

| Score | Expect | Identities | Gaps | Strand | Frame |
|---|---|---|---|---|---|
| 14.4 bits(7) | 46() | 7/7(100%) | 0/7(0%) | Plus/Plus | |

Features:

```
Query  12     GTTTTCT  18
              |||||||
Sbjct  44155  GTTTTCT  44161
```

Range 182: 44601 to 44607

| Score | Expect | Identities | Gaps | Strand | Frame |
|---|---|---|---|---|---|

| Score | Expect | Identities | Gaps | Strand | Frame |
|---|---|---|---|---|---|
| 14.4 bits(7) | 46() | 7/7(100%) | 0/7(0%) | Plus/Plus | |

Features:

```
Query  9      GTTGTTT  15
              |||||||
Sbjct  44601  GTTGTTT  44607
```

Range 183: 44779 to 44785

| Score | Expect | Identities | Gaps | Strand | Frame |
|---|---|---|---|---|---|
| 14.4 bits(7) | 46() | 7/7(100%) | 0/7(0%) | Plus/Minus | |

Features:

```
Query  10     TTGTTTT  16
              |||||||
Sbjct  44785  TTGTTTT  44779
```

Range 184: 44936 to 44942

| Score | Expect | Identities | Gaps | Strand | Frame |
|---|---|---|---|---|---|
| 14.4 bits(7) | 46() | 7/7(100%) | 0/7(0%) | Plus/Minus | |

Features:

```
Query  8      TGTTGTT  14
              |||||||
Sbjct  44942  TGTTGTT  44936
```

Range 185: 45172 to 45178

| Score | Expect | Identities | Gaps | Strand | Frame |
|---|---|---|---|---|---|
| 14.4 bits(7) | 46() | 7/7(100%) | 0/7(0%) | Plus/Plus | |

Features:

```
Query  10     TTGTTTT  16
              |||||||
Sbjct  45172  TTGTTTT  45178
```

Range 186: 45850 to 45856

| Score | Expect | Identities | Gaps | Strand | Frame |
|---|---|---|---|---|---|
| 14.4 bits(7) | 46() | 7/7(100%) | 0/7(0%) | Plus/Plus | |

Features:

```
Query  18     TCCTCTA  24
              |||||||
Sbjct  45850  TCCTCTA  45856
```

Range 187: 46238 to 46244

| Score | Expect | Identities | Gaps | Strand | Frame |
|---|---|---|---|---|---|
| 14.4 bits(7) | 46() | 7/7(100%) | 0/7(0%) | Plus/Minus | |

Features:

```
Query  8      TGTTGTT  14
              |||||||
Sbjct  46244  TGTTGTT  46238
```

Range 188: 46347 to 46353

| Score | Expect | Identities | Gaps | Strand | Frame |
|---|---|---|---|---|---|
| 14.4 bits(7) | 46() | 7/7(100%) | 0/7(0%) | Plus/Plus | |

Features:

```
Query  15     TTCTCCT  21
              |||||||
Sbjct  46347  TTCTCCT  46353
```

Range 189: 46647 to 46653

| Score | Expect | Identities | Gaps | Strand | Frame |
|---|---|---|---|---|---|
| 14.4 bits(7) | 46() | 7/7(100%) | 0/7(0%) | Plus/Plus | |

Features:

```
Query  10     TTGTTTT  16
              |||||||
Sbjct  46647  TTGTTTT  46653
```

Range 190: 46741 to 46747

| Score | Expect | Identities | Gaps | Strand | Frame |
|---|---|---|---|---|---|
| 14.4 bits(7) | 46() | 7/7(100%) | 0/7(0%) | Plus/Plus | |

Features:

```
Query  15     TTCTCCT  21
              |||||||
Sbjct  46741  TTCTCCT  46747
```

Range 191: 47008 to 47014

| Score | Expect | Identities | Gaps | Strand | Frame |
|---|---|---|---|---|---|
| 14.4 bits(7) | 46() | 7/7(100%) | 0/7(0%) | Plus/Minus | |

Features:

```
Query  2      ACAGTCT  8
              |||||||
Sbjct  47014  ACAGTCT  47008
```

Range 192: 47032 to 47038

| Score | Expect | Identities | Gaps | Strand | Frame |
|---|---|---|---|---|---|
| 14.4 bits(7) | 46() | 7/7(100%) | 0/7(0%) | Plus/Plus | |

Features:

```
Query  13     TTTTCTC  19
              |||||||
Sbjct  47032  TTTTCTC  47038
```

Range 193: 47135 to 47141

| Score | Expect | Identities | Gaps | Strand | Frame |
|---|---|---|---|---|---|
| 14.4 bits(7) | 46() | 7/7(100%) | 0/7(0%) | Plus/Plus | |

Features:

```
Query  16     TCTCCTC  22
              |||||||
Sbjct  47135  TCTCCTC  47141
```

Range 194: 47313 to 47319

| Score | Expect | Identities | Gaps | Strand | Frame |
|---|---|---|---|---|---|
| 14.4 bits(7) | 46() | 7/7(100%) | 0/7(0%) | Plus/Minus | |

Features:

```
Query  13     TTTTCTC  19
              |||||||
Sbjct  47319  TTTTCTC  47313
```

Range 195: 48097 to 48103

| Score | Expect | Identities | Gaps | Strand | Frame |
|---|---|---|---|---|---|
| 14.4 bits(7) | 46() | 7/7(100%) | 0/7(0%) | Plus/Plus | |

Features:

```
Query  15     TTCTCCT  21
              |||||||
Sbjct  48097  TTCTCCT  48103
```

Range 196: 48184 to 48190

| Score | Expect | Identities | Gaps | Strand | Frame |
|---|---|---|---|---|---|
| 14.4 bits(7) | 46() | 7/7(100%) | 0/7(0%) | Plus/Plus | |

Features:

```
Query  14     TTTCTCC  20
              |||||||
Sbjct  48184  TTTCTCC  48190
```

Range 197: 48432 to 48438

| Score | Expect | Identities | Gaps | Strand | Frame |
|---|---|---|---|---|---|
| 14.4 bits(7) | 46() | 7/7(100%) | 0/7(0%) | Plus/Plus | |

Features:

```
Query  10     TTGTTTT  16
              |||||||
Sbjct  48432  TTGTTTT  48438
```

Range 198: 48437 to 48443

| Score | Expect | Identities | Gaps | Strand | Frame |
|---|---|---|---|---|---|
| 14.4 bits(7) | 46() | 7/7(100%) | 0/7(0%) | Plus/Plus | |

Features:

```
Query  10     TTGTTTT  16
              |||||||
Sbjct  48437  TTGTTTT  48443
```

Range 199: 48444 to 48450

| Score | Expect | Identities | Gaps | Strand | Frame |
|---|---|---|---|---|---|
| 14.4 bits(7) | 46() | 7/7(100%) | 0/7(0%) | Plus/Plus | |

Features:

```
Query  10     TTGTTTT  16
              |||||||
Sbjct  48444  TTGTTTT  48450
```

Range 200: 48791 to 48797

| Score | Expect | Identities | Gaps | Strand | Frame |
|---|---|---|---|---|---|
| 14.4 bits(7) | 46() | 7/7(100%) | 0/7(0%) | Plus/Minus | |

Features:

```
Query  9      GTTGTTT  15
              |||||||
Sbjct  48797  GTTGTTT  48791
```

Range 201: 48882 to 48888

| Score | Expect | Identities | Gaps | Strand | Frame |
|---|---|---|---|---|---|

| | | | | |
|---|---|---|---|---|
| 14.4 bits(7) | 46() | 7/7(100%) | 0/7(0%) | Plus/Plus |

Features:

```
Query  8      TGTTGTT  14
              |||||||
Sbjct  48882  TGTTGTT  48888
```

Range 202: 49047 to 49053

| Score | Expect | Identities | Gaps | Strand | Frame |
|---|---|---|---|---|---|
| 14.4 bits(7) | 46() | 7/7(100%) | 0/7(0%) | Plus/Plus | |

Features:

```
Query  10     TTGTTTT  16
              |||||||
Sbjct  49047  TTGTTTT  49053
```

Range 203: 49100 to 49106

| Score | Expect | Identities | Gaps | Strand | Frame |
|---|---|---|---|---|---|
| 14.4 bits(7) | 46() | 7/7(100%) | 0/7(0%) | Plus/Plus | |

Features:

```
Query  15     TTCTCCT  21
              |||||||
Sbjct  49100  TTCTCCT  49106
```

Range 204: 49407 to 49413

| Score | Expect | Identities | Gaps | Strand | Frame |
|---|---|---|---|---|---|
| 14.4 bits(7) | 46() | 7/7(100%) | 0/7(0%) | Plus/Plus | |

Features:

```
Query  10     TTGTTTT  16
              |||||||
Sbjct  49407  TTGTTTT  49413
```

Range 205: 49412 to 49418

| Score | Expect | Identities | Gaps | Strand | Frame |
|---|---|---|---|---|---|
| 14.4 bits(7) | 46() | 7/7(100%) | 0/7(0%) | Plus/Plus | |

Features:

```
Query  10     TTGTTTT  16
              |||||||
Sbjct  49412  TTGTTTT  49418
```

Range 206: 49422 to 49428

| Score | Expect | Identities | Gaps | Strand | Frame |
|---|---|---|---|---|---|
| 14.4 bits(7) | 46() | 7/7(100%) | 0/7(0%) | Plus/Plus | |

Features:

```
Query  10     TTGTTTT  16
              |||||||
Sbjct  49422  TTGTTTT  49428
```

Range 207: 49427 to 49433

| Score | Expect | Identities | Gaps | Strand | Frame |
|---|---|---|---|---|---|
| 14.4 bits(7) | 46() | 7/7(100%) | 0/7(0%) | Plus/Plus | |

Features:

```
Query  10     TTGTTTT  16
              |||||||
Sbjct  49427  TTGTTTT  49433
```

Range 208: 49432 to 49438

| Score | Expect | Identities | Gaps | Strand | Frame |
|---|---|---|---|---|---|
| 14.4 bits(7) | 46() | 7/7(100%) | 0/7(0%) | Plus/Plus | |

Features:

```
Query  10     TTGTTTT  16
              |||||||
Sbjct  49432  TTGTTTT  49438
```

Range 209: 49508 to 49514

| Score | Expect | Identities | Gaps | Strand | Frame |
|---|---|---|---|---|---|
| 14.4 bits(7) | 46() | 7/7(100%) | 0/7(0%) | Plus/Plus | |

Features:

```
Query  19     CCTCTAT  25
              |||||||
Sbjct  49508  CCTCTAT  49514
```

Range 210: 49909 to 49915

| Score | Expect | Identities | Gaps | Strand | Frame |
|---|---|---|---|---|---|
| 14.4 bits(7) | 46() | 7/7(100%) | 0/7(0%) | Plus/Plus | |

Features:

```
Query  18     TCCTCTA  24
              |||||||
Sbjct  49909  TCCTCTA  49915
```

Range 211: 50083 to 50089

| Score | Expect | Identities | Gaps | Strand | Frame |
|---|---|---|---|---|---|
| 14.4 bits(7) | 46() | 7/7(100%) | 0/7(0%) | Plus/Plus | |

Features:

```
Query  13     TTTTCTC  19
              |||||||
Sbjct  50083  TTTTCTC  50089
```

Range 212: 50218 to 50224

| Score | Expect | Identities | Gaps | Strand | Frame |
|---|---|---|---|---|---|
| 14.4 bits(7) | 46() | 7/7(100%) | 0/7(0%) | Plus/Plus | |

Features:

```
Query  15     TTCTCCT  21
              |||||||
Sbjct  50218  TTCTCCT  50224
```

Range 213: 50522 to 50528

| Score | Expect | Identities | Gaps | Strand | Frame |
|---|---|---|---|---|---|
| 14.4 bits(7) | 46() | 7/7(100%) | 0/7(0%) | Plus/Plus | |

Features:

```
Query  11     TGTTTTC  17
              |||||||
Sbjct  50522  TGTTTTC  50528
```

Range 214: 50928 to 50934

| Score | Expect | Identities | Gaps | Strand | Frame |
|---|---|---|---|---|---|
| 14.4 bits(7) | 46() | 7/7(100%) | 0/7(0%) | Plus/Plus | |

Features:

```
Query  7      CTGTTGT  13
              |||||||
Sbjct  50928  CTGTTGT  50934
```

Range 215: 51084 to 51090

| Score | Expect | Identities | Gaps | Strand | Frame |
|---|---|---|---|---|---|
| 14.4 bits(7) | 46() | 7/7(100%) | 0/7(0%) | Plus/Plus | |

Features:

```
Query  15     TTCTCCT  21
              |||||||
Sbjct  51084  TTCTCCT  51090
```

Range 216: 51146 to 51152

| Score | Expect | Identities | Gaps | Strand | Frame |
|---|---|---|---|---|---|
| 14.4 bits(7) | 46() | 7/7(100%) | 0/7(0%) | Plus/Minus | |

Features:

```
Query  11     TGTTTTC  17
              |||||||
Sbjct  51152  TGTTTTC  51146
```

Range 217: 51398 to 51404

| Score | Expect | Identities | Gaps | Strand | Frame |
|---|---|---|---|---|---|
| 14.4 bits(7) | 46() | 7/7(100%) | 0/7(0%) | Plus/Plus | |

Features:

```
Query  10     TTGTTTT  16
              |||||||
Sbjct  51398  TTGTTTT  51404
```

Range 218: 51725 to 51731

| Score | Expect | Identities | Gaps | Strand | Frame |
|---|---|---|---|---|---|
| 14.4 bits(7) | 46() | 7/7(100%) | 0/7(0%) | Plus/Plus | |

Features:

```
Query  11     TGTTTTC  17
              |||||||
Sbjct  51725  TGTTTTC  51731
```

Range 219: 51741 to 51747

| Score | Expect | Identities | Gaps | Strand | Frame |
|---|---|---|---|---|---|
| 14.4 bits(7) | 46() | 7/7(100%) | 0/7(0%) | Plus/Minus | |

Features:

```
Query  9      GTTGTTT  15
              |||||||
Sbjct  51747  GTTGTTT  51741
```

Range 220: 51775 to 51781

| Score | Expect | Identities | Gaps | Strand | Frame |
|---|---|---|---|---|---|

| | | | | |
|---|---|---|---|---|
| 14.4 bits(7) | 46() | 7/7(100%) | 0/7(0%) | Plus/Plus |

Features:

```
Query  13     TTTTCTC  19
              |||||||
Sbjct  51775  TTTTCTC  51781
```

Range 221: 52009 to 52015

| Score | Expect | Identities | Gaps | Strand | Frame |
|---|---|---|---|---|---|
| 14.4 bits(7) | 46() | 7/7(100%) | 0/7(0%) | Plus/Plus | |

Features:

```
Query  8      TGTTGTT  14
              |||||||
Sbjct  52009  TGTTGTT  52015
```

Range 222: 52397 to 52403

| Score | Expect | Identities | Gaps | Strand | Frame |
|---|---|---|---|---|---|
| 14.4 bits(7) | 46() | 7/7(100%) | 0/7(0%) | Plus/Plus | |

Features:

```
Query  10     TTGTTTT  16
              |||||||
Sbjct  52397  TTGTTTT  52403
```

Range 223: 52450 to 52456

| Score | Expect | Identities | Gaps | Strand | Frame |
|---|---|---|---|---|---|
| 14.4 bits(7) | 46() | 7/7(100%) | 0/7(0%) | Plus/Plus | |

Features:

```
Query  10     TTGTTTT  16
              |||||||
Sbjct  52450  TTGTTTT  52456
```

Range 224: 53420 to 53426

| Score | Expect | Identities | Gaps | Strand | Frame |
|---|---|---|---|---|---|
| 14.4 bits(7) | 46() | 7/7(100%) | 0/7(0%) | Plus/Plus | |

Features:

```
Query  18     TCCTCTA  24
              |||||||
Sbjct  53420  TCCTCTA  53426
```

Range 225: 53542 to 53548

| Score | Expect | Identities | Gaps | Strand | Frame |
|---|---|---|---|---|---|
| 14.4 bits(7) | 46() | 7/7(100%) | 0/7(0%) | Plus/Plus | |

Features:

```
Query  9      GTTGTTT  15
              |||||||
Sbjct  53542  GTTGTTT  53548
```

Range 226: 53766 to 53772

| Score | Expect | Identities | Gaps | Strand | Frame |
|---|---|---|---|---|---|
| 14.4 bits(7) | 46() | 7/7(100%) | 0/7(0%) | Plus/Plus | |

Features:

```
Query  15     TTCTCCT  21
              |||||||
Sbjct  53766  TTCTCCT  53772
```

Range 227: 53777 to 53783

| Score | Expect | Identities | Gaps | Strand | Frame |
|---|---|---|---|---|---|
| 14.4 bits(7) | 46() | 7/7(100%) | 0/7(0%) | Plus/Plus | |

Features:

```
Query  6      TCTGTTG  12
              |||||||
Sbjct  53777  TCTGTTG  53783
```

Range 228: 54303 to 54309

| Score | Expect | Identities | Gaps | Strand | Frame |
|---|---|---|---|---|---|
| 14.4 bits(7) | 46() | 7/7(100%) | 0/7(0%) | Plus/Plus | |

Features:

```
Query  12     GTTTTCT  18
              |||||||
Sbjct  54303  GTTTTCT  54309
```

Range 229: 54344 to 54350

| Score | Expect | Identities | Gaps | Strand | Frame |
|---|---|---|---|---|---|
| 14.4 bits(7) | 46() | 7/7(100%) | 0/7(0%) | Plus/Plus | |

Features:

```
Query  11     TGTTTTC  17
              |||||||
Sbjct  54344  TGTTTTC  54350
```

Range 230: 54420 to 54426

| Score | Expect | Identities | Gaps | Strand | Frame |
|---|---|---|---|---|---|
| 14.4 bits(7) | 46() | 7/7(100%) | 0/7(0%) | Plus/Plus | |

Features:

```
Query  10     TTGTTTT  16
              |||||||
Sbjct  54420  TTGTTTT  54426
```

Range 231: 55140 to 55146

| Score | Expect | Identities | Gaps | Strand | Frame |
|---|---|---|---|---|---|
| 14.4 bits(7) | 46() | 7/7(100%) | 0/7(0%) | Plus/Plus | |

Features:

```
Query  8      TGTTGTT  14
              |||||||
Sbjct  55140  TGTTGTT  55146
```

Range 232: 55277 to 55283

| Score | Expect | Identities | Gaps | Strand | Frame |
|---|---|---|---|---|---|
| 14.4 bits(7) | 46() | 7/7(100%) | 0/7(0%) | Plus/Plus | |

Features:

```
Query  10     TTGTTTT  16
              |||||||
Sbjct  55277  TTGTTTT  55283
```

Range 233: 55577 to 55583

| Score | Expect | Identities | Gaps | Strand | Frame |
|---|---|---|---|---|---|
| 14.4 bits(7) | 46() | 7/7(100%) | 0/7(0%) | Plus/Minus | |

Features:

```
Query  15     TTCTCCT  21
              |||||||
Sbjct  55583  TTCTCCT  55577
```

Range 234: 55649 to 55655

| Score | Expect | Identities | Gaps | Strand | Frame |
|---|---|---|---|---|---|
| 14.4 bits(7) | 46() | 7/7(100%) | 0/7(0%) | Plus/Minus | |

Features:

```
Query  6      TCTGTTG  12
              |||||||
Sbjct  55655  TCTGTTG  55649
```

Range 235: 56099 to 56105

| Score | Expect | Identities | Gaps | Strand | Frame |
|---|---|---|---|---|---|
| 14.4 bits(7) | 46() | 7/7(100%) | 0/7(0%) | Plus/Minus | |

Features:

```
Query  14     TTTCTCC  20
              |||||||
Sbjct  56105  TTTCTCC  56099
```

Range 236: 56186 to 56192

| Score | Expect | Identities | Gaps | Strand | Frame |
|---|---|---|---|---|---|
| 14.4 bits(7) | 46() | 7/7(100%) | 0/7(0%) | Plus/Minus | |

Features:

```
Query  15     TTCTCCT  21
              |||||||
Sbjct  56192  TTCTCCT  56186
```

Range 237: 56450 to 56456

| Score | Expect | Identities | Gaps | Strand | Frame |
|---|---|---|---|---|---|
| 14.4 bits(7) | 46() | 7/7(100%) | 0/7(0%) | Plus/Plus | |

Features:

```
Query  3      CAGTCTG  9
              |||||||
Sbjct  56450  CAGTCTG  56456
```

Range 238: 56856 to 56862

| Score | Expect | Identities | Gaps | Strand | Frame |
|---|---|---|---|---|---|
| 14.4 bits(7) | 46() | 7/7(100%) | 0/7(0%) | Plus/Plus | |

Features:

```
Query  11     TGTTTTC  17
              |||||||
Sbjct  56856  TGTTTTC  56862
```

Range 239: 57008 to 57014

| Score | Expect | Identities | Gaps | Strand | Frame |
|---|---|---|---|---|---|

| | | | | |
|---|---|---|---|---|
| 14.4 bits(7) | 46() | 7/7(100%) | 0/7(0%) | Plus/Minus |

Features:

```
Query  11     TGTTTTC  17
              |||||||
Sbjct  57014  TGTTTTC  57008
```

Range 240: 57017 to 57023

| Score | Expect | Identities | Gaps | Strand | Frame |
|---|---|---|---|---|---|
| 14.4 bits(7) | 46() | 7/7(100%) | 0/7(0%) | Plus/Minus | |

Features:

```
Query  13     TTTTCTC  19
              |||||||
Sbjct  57023  TTTTCTC  57017
```

Range 241: 58179 to 58185

| Score | Expect | Identities | Gaps | Strand | Frame |
|---|---|---|---|---|---|
| 14.4 bits(7) | 46() | 7/7(100%) | 0/7(0%) | Plus/Minus | |

Features:

```
Query  15     TTCTCCT  21
              |||||||
Sbjct  58185  TTCTCCT  58179
```

Range 242: 58274 to 58280

| Score | Expect | Identities | Gaps | Strand | Frame |
|---|---|---|---|---|---|
| 14.4 bits(7) | 46() | 7/7(100%) | 0/7(0%) | Plus/Minus | |

Features:

```
Query  10     TTGTTTT  16
              |||||||
Sbjct  58280  TTGTTTT  58274
```

Range 243: 58279 to 58285

| Score | Expect | Identities | Gaps | Strand | Frame |
|---|---|---|---|---|---|
| 14.4 bits(7) | 46() | 7/7(100%) | 0/7(0%) | Plus/Minus | |

Features:

```
Query  10     TTGTTTT  16
              |||||||
Sbjct  58285  TTGTTTT  58279
```

Range 244: 58284 to 58290

| Score | Expect | Identities | Gaps | Strand | Frame |
|---|---|---|---|---|---|
| 14.4 bits(7) | 46() | 7/7(100%) | 0/7(0%) | Plus/Minus | |

Features:

```
Query  10     TTGTTTT  16
              |||||||
Sbjct  58290  TTGTTTT  58284
```

Range 245: 59963 to 59969

| Score | Expect | Identities | Gaps | Strand | Frame |
|---|---|---|---|---|---|
| 14.4 bits(7) | 46() | 7/7(100%) | 0/7(0%) | Plus/Plus | |

Features:

```
Query  7      CTGTTGT  13
              |||||||
Sbjct  59963  CTGTTGT  59969
```

Range 246: 60473 to 60479

| Score | Expect | Identities | Gaps | Strand | Frame |
|---|---|---|---|---|---|
| 14.4 bits(7) | 46() | 7/7(100%) | 0/7(0%) | Plus/Plus | |

Features:

```
Query  15     TTCTCCT  21
              |||||||
Sbjct  60473  TTCTCCT  60479
```

Range 247: 60562 to 60568

| Score | Expect | Identities | Gaps | Strand | Frame |
|---|---|---|---|---|---|
| 14.4 bits(7) | 46() | 7/7(100%) | 0/7(0%) | Plus/Plus | |

Features:

```
Query  14     TTTCTCC  20
              |||||||
Sbjct  60562  TTTCTCC  60568
```

BLAST is a registered trademark of the National Library of Medicine

Support center Mailing list YouTube

- National Library Of Medicine
- National Institutes Of Health
- U.S. Department of Health & Human Services
- USA.gov

**NCBI**

*National Center for Biotechnology Information, U.S. National Library of Medicine 8600 Rockville Pike, Bethesda MD, 20894 USA*

Policies and Guidelines | Contact
