## Supplemental Figure S5 for "MORT, a locus for apoptosis in the human immunodeficiency virus-type 1 antisense gene: implications for AIDS, Cancer, and Covid-19"

BLAST ® » blastn suite-2sequences » RID-5TWKGCNS114

**BLAST Results**

**Blast 2 sequences**

**Job title: MORT271.299.VS.BIRC6**

| | |
|---|---|
| **RID** | 5TWKGCNS114 (Expires on 01-17 01:26 am) |
| **Query ID** | CQ767337.1 |
| **Description** | Sequence 27 from Patent EP1359221. |
| **Molecule type** | rna |
| **Query Length** | 28 |

| | |
|---|---|
| **Subject ID** | NC_000002.12 |
| **Description** | Homo sapiens chromosome 2, GRCh38.p7 Primary Assembly<br>See details |
| **Molecule type** | dna |
| **Subject Length** | 261992 |
| **Program** | BLASTN 2.7.1+ |

### Graphic Summary

**Distribution of the top 101 Blast Hits on 1 subject sequences**

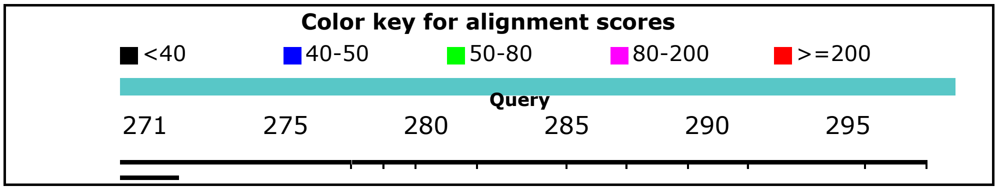

### Dot Matrix View

### Descriptions

Sequences producing significant alignments:

| Description | Max score | Total score | Query cover | E value | Ident | Accession |
|---|---|---|---|---|---|---|
| Homo sapiens chromosome 2, GRCh38.p7 Primary Assembly | 20.3 | 10534 | 100% | 3.6 | 100% | NC_000002.12 |

### Alignments

Homo sapiens chromosome 2, GRCh38.p7 Primary Assembly
Sequence ID: **NC_000002.12** Length: 242193529 Number of Matches: 705
Range 1: 32363510 to 32363519

| Score | Expect | Identities | Gaps | Strand | Frame |
|---|---|---|---|---|---|
| 20.3 bits(10) | 3.6() | 10/10(100%) | 0/10(0%) | Plus/Plus | |

Features:

```
Query  282       GGTGTTCTCT  291
                 ||||||||||
Sbjct  32363510  GGTGTTCTCT  32363519
```

Range 2: 32378213 to 32378222

| Score | Expect | Identities | Gaps | Strand | Frame |
|---|---|---|---|---|---|
| 20.3 bits(10) | 3.6() | 10/10(100%) | 0/10(0%) | Plus/Plus | |

Features:

```
Query  286       TTCTCTCCTT  295
                 ||||||||||
Sbjct  32378213  TTCTCTCCTT  32378222
```

Range 3: 32405828 to 32405837

| Score | Expect | Identities | Gaps | Strand | Frame |
|---|---|---|---|---|---|
| 20.3 bits(10) | 3.6() | 10/10(100%) | 0/10(0%) | Plus/Plus | |

Features:

```
Query  276       CAAGCTGGTG  285
                 ||||||||||
Sbjct  32405828  CAAGCTGGTG  32405837
```

Range 4: 32487352 to 32487361

| Score | Expect | Identities | Gaps | Strand | Frame |
|---|---|---|---|---|---|
| 20.3 bits(10) | 3.6() | 10/10(100%) | 0/10(0%) | Plus/Plus | |

Features:

```
Query  290       CTCCTTTATT  299
                 ||||||||||
Sbjct  32487352  CTCCTTTATT  32487361
```

Range 5: 32541805 to 32541814

| Score | Expect | Identities | Gaps | Strand | Frame |
|---|---|---|---|---|---|
| 20.3 bits(10) | 3.6() | 10/10(100%) | 0/10(0%) | Plus/Plus | |

Features:

```
Query  290       CTCCTTTATT  299
                 ||||||||||
Sbjct  32541805  CTCCTTTATT  32541814
```

Range 6: 32568137 to 32568146

| Score | Expect | Identities | Gaps | Strand | Frame |
|---|---|---|---|---|---|
| 20.3 bits(10) | 3.6() | 10/10(100%) | 0/10(0%) | Plus/Minus | |

Features:

```
Query  286       TTCTCTCCTT  295
                 ||||||||||
Sbjct  32568146  TTCTCTCCTT  32568137
```

Range 7: 32380365 to 32380373

| Score | Expect | Identities | Gaps | Strand | Frame |
|---|---|---|---|---|---|
| 18.3 bits(9) | 14() | 9/9(100%) | 0/9(0%) | Plus/Plus | |

Features:

```
Query  288       CTCTCCTTT  296
                 |||||||||
Sbjct  32380365  CTCTCCTTT  32380373
```

Range 8: 32433081 to 32433089

| Score | Expect | Identities | Gaps | Strand | Frame |
|---|---|---|---|---|---|
| 18.3 bits(9) | 14() | 9/9(100%) | 0/9(0%) | Plus/Minus | |

Features:

```
Query  291       TCCTTTATT  299
                 |||||||||
Sbjct  32433089  TCCTTTATT  32433081
```

Range 9: 32434299 to 32434311

| Score | Expect | Identities | Gaps | Strand | Frame |
|---|---|---|---|---|---|
| 18.3 bits(9) | 14() | 12/13(92%) | 0/13(0%) | Plus/Plus | |

Features:

```
Query  283       GTGTTCTCTCCTT  295
                 ||||| |||||||
Sbjct  32434299  GTGTTTTCTCCTT  32434311
```

Range 10: 32436772 to 32436780

| Score | Expect | Identities | Gaps | Strand | Frame |
|---|---|---|---|---|---|
| 18.3 bits(9) | 14() | 9/9(100%) | 0/9(0%) | Plus/Plus | |

Features:

```
Query  276       CAAGCTGGT  284
                 |||||||||
Sbjct  32436772  CAAGCTGGT  32436780
```

Range 11: 32441248 to 32441256

| Score | Expect | Identities | Gaps | Strand | Frame |
|---|---|---|---|---|---|
| 18.3 bits(9) | 14() | 9/9(100%) | 0/9(0%) | Plus/Plus | |

Features:

```
Query  291       TCCTTTATT  299
                 |||||||||
Sbjct  32441248  TCCTTTATT  32441256
```

Range 12: 32446440 to 32446448

| Score | Expect | Identities | Gaps | Strand | Frame |
|---|---|---|---|---|---|
| 18.3 bits(9) | 14() | 9/9(100%) | 0/9(0%) | Plus/Plus | |

Features:

```
Query  291       TCCTTTATT  299
                 |||||||||
Sbjct  32446440  TCCTTTATT  32446448
```

Range 13: 32457126 to 32457134

| Score | Expect | Identities | Gaps | Strand | Frame |
|---|---|---|---|---|---|
| 18.3 bits(9) | 14() | 9/9(100%) | 0/9(0%) | Plus/Minus | |

Features:

```
Query  279       GCTGGTGTT  287
                 |||||||||
Sbjct  32457134  GCTGGTGTT  32457126
```

Range 14: 32458475 to 32458483

| Score | Expect | Identities | Gaps | Strand | Frame |
|---|---|---|---|---|---|
| 18.3 bits(9) | 14() | 9/9(100%) | 0/9(0%) | Plus/Plus | |

Features:

```
Query  287       TCTCTCCTT  295
                 |||||||||
Sbjct  32458475  TCTCTCCTT  32458483
```

Range 15: 32461414 to 32461422

| Score | Expect | Identities | Gaps | Strand | Frame |
|---|---|---|---|---|---|
| 18.3 bits(9) | 14() | 9/9(100%) | 0/9(0%) | Plus/Plus | |

Features:

```
Query  276       CAAGCTGGT  284
                 |||||||||
Sbjct  32461414  CAAGCTGGT  32461422
```

Range 16: 32462510 to 32462518

| Score | Expect | Identities | Gaps | Strand | Frame |
|---|---|---|---|---|---|
| 18.3 bits(9) | 14() | 9/9(100%) | 0/9(0%) | Plus/Plus | |

Features:

```
Query  281       TGGTGTTCT  289
                 |||||||||
Sbjct  32462510  TGGTGTTCT  32462518
```

Range 17: 32471730 to 32471738

| Score | Expect | Identities | Gaps | Strand | Frame |
|---|---|---|---|---|---|
| 18.3 bits(9) | 14() | 9/9(100%) | 0/9(0%) | Plus/Minus | |

Features:

```
Query  291       TCCTTTATT  299
                 |||||||||
```

```
Sbjct  32471738  TCCTTTATT  32471730
```

Range 18: 32472648 to 32472656

| Score | Expect | Identities | Gaps | Strand | Frame |
|---|---|---|---|---|---|
| 18.3 bits(9) | 14() | 9/9(100%) | 0/9(0%) | Plus/Minus | |

Features:

```
Query  288       CTCTCCTTT  296
                 |||||||||
Sbjct  32472656  CTCTCCTTT  32472648
```

Range 19: 32473219 to 32473227

| Score | Expect | Identities | Gaps | Strand | Frame |
|---|---|---|---|---|---|
| 18.3 bits(9) | 14() | 9/9(100%) | 0/9(0%) | Plus/Minus | |

Features:

```
Query  274       AACAAGCTG  282
                 |||||||||
Sbjct  32473227  AACAAGCTG  32473219
```

Range 20: 32475205 to 32475213

| Score | Expect | Identities | Gaps | Strand | Frame |
|---|---|---|---|---|---|
| 18.3 bits(9) | 14() | 9/9(100%) | 0/9(0%) | Plus/Plus | |

Features:

```
Query  283       GTGTTCTCT  291
                 |||||||||
Sbjct  32475205  GTGTTCTCT  32475213
```

Range 21: 32482129 to 32482137

| Score | Expect | Identities | Gaps | Strand | Frame |
|---|---|---|---|---|---|
| 18.3 bits(9) | 14() | 9/9(100%) | 0/9(0%) | Plus/Minus | |

Features:

```
Query  289       TCTCCTTTA  297
                 |||||||||
Sbjct  32482137  TCTCCTTTA  32482129
```

Range 22: 32485090 to 32485098

| Score | Expect | Identities | Gaps | Strand | Frame |
|---|---|---|---|---|---|
| 18.3 bits(9) | 14() | 9/9(100%) | 0/9(0%) | Plus/Minus | |

Features:

```
Query  289       TCTCCTTTA  297
                 |||||||||
Sbjct  32485098  TCTCCTTTA  32485090
```

Range 23: 32488142 to 32488150

| Score | Expect | Identities | Gaps | Strand | Frame |
|---|---|---|---|---|---|
| 18.3 bits(9) | 14() | 9/9(100%) | 0/9(0%) | Plus/Minus | |

Features:

```
Query  276       CAAGCTGGT  284
                 |||||||||
Sbjct  32488150  CAAGCTGGT  32488142
```

Range 24: 32488972 to 32488980

| Score | Expect | Identities | Gaps | Strand | Frame |
|---|---|---|---|---|---|
| 18.3 bits(9) | 14() | 9/9(100%) | 0/9(0%) | Plus/Plus | |

Features:

```
Query  290       CTCCTTTAT  298
                 |||||||||
Sbjct  32488972  CTCCTTTAT  32488980
```

Range 25: 32500813 to 32500821

| Score | Expect | Identities | Gaps | Strand | Frame |
|---|---|---|---|---|---|
| 18.3 bits(9) | 14() | 9/9(100%) | 0/9(0%) | Plus/Plus | |

Features:

```
Query  276       CAAGCTGGT  284
                 |||||||||
Sbjct  32500813  CAAGCTGGT  32500821
```

Range 26: 32502193 to 32502201

| Score | Expect | Identities | Gaps | Strand | Frame |
|---|---|---|---|---|---|
| 18.3 bits(9) | 14() | 9/9(100%) | 0/9(0%) | Plus/Plus | |

Features:

```
Query  282       GGTGTTCTC  290
                 |||||||||
Sbjct  32502193  GGTGTTCTC  32502201
```

Range 27: 32510595 to 32510603

| Score | Expect | Identities | Gaps | Strand | Frame |
|---|---|---|---|---|---|
| 18.3 bits(9) | 14() | 9/9(100%) | 0/9(0%) | Plus/Minus | |

Features:

```
Query  276       CAAGCTGGT  284
                 |||||||||
Sbjct  32510603  CAAGCTGGT  32510595
```

Range 28: 32513575 to 32513583

| Score | Expect | Identities | Gaps | Strand | Frame |
|---|---|---|---|---|---|
| 18.3 bits(9) | 14() | 9/9(100%) | 0/9(0%) | Plus/Minus | |

Features:

```
Query  276       CAAGCTGGT  284
                 |||||||||
Sbjct  32513583  CAAGCTGGT  32513575
```

Range 29: 32526486 to 32526494

| Score | Expect | Identities | Gaps | Strand | Frame |
|---|---|---|---|---|---|
| 18.3 bits(9) | 14() | 9/9(100%) | 0/9(0%) | Plus/Plus | |

Features:

```
Query  274       AACAAGCTG  282
                 |||||||||
Sbjct  32526486  AACAAGCTG  32526494
```

Range 30: 32556397 to 32556405

| Score | Expect | Identities | Gaps | Strand | Frame |
|---|---|---|---|---|---|

| | | | | |
|---|---|---|---|---|
| 18.3 bits(9) | 14() | 9/9(100%) | 0/9(0%) | Plus/Plus |

Features:

```
Query  286       TTCTCTCCT  294
                 |||||||||
Sbjct  32556397  TTCTCTCCT  32556405
```

Range 31: 32559647 to 32559655

| Score | Expect | Identities | Gaps | Strand | Frame |
|---|---|---|---|---|---|
| 18.3 bits(9) | 14() | 9/9(100%) | 0/9(0%) | Plus/Minus | |

Features:

```
Query  288       CTCTCCTTT  296
                 |||||||||
Sbjct  32559655  CTCTCCTTT  32559647
```

Range 32: 32570178 to 32570186

| Score | Expect | Identities | Gaps | Strand | Frame |
|---|---|---|---|---|---|
| 18.3 bits(9) | 14() | 9/9(100%) | 0/9(0%) | Plus/Plus | |

Features:

```
Query  291       TCCTTTATT  299
                 |||||||||
Sbjct  32570178  TCCTTTATT  32570186
```

Range 33: 32582067 to 32582075

| Score | Expect | Identities | Gaps | Strand | Frame |
|---|---|---|---|---|---|
| 18.3 bits(9) | 14() | 9/9(100%) | 0/9(0%) | Plus/Plus | |

Features:

```
Query  276       CAAGCTGGT  284
                 |||||||||
Sbjct  32582067  CAAGCTGGT  32582075
```

Range 34: 32590373 to 32590381

| Score | Expect | Identities | Gaps | Strand | Frame |
|---|---|---|---|---|---|
| 18.3 bits(9) | 14() | 9/9(100%) | 0/9(0%) | Plus/Plus | |

Features:

```
Query  288       CTCTCCTTT  296
                 |||||||||
Sbjct  32590373  CTCTCCTTT  32590381
```

Range 35: 32592243 to 32592251

| Score | Expect | Identities | Gaps | Strand | Frame |
|---|---|---|---|---|---|
| 18.3 bits(9) | 14() | 9/9(100%) | 0/9(0%) | Plus/Plus | |

Features:

```
Query  283       GTGTTCTCT  291
                 |||||||||
Sbjct  32592243  GTGTTCTCT  32592251
```

Range 36: 32597648 to 32597656

| Score | Expect | Identities | Gaps | Strand | Frame |
|---|---|---|---|---|---|
| 18.3 bits(9) | 14() | 9/9(100%) | 0/9(0%) | Plus/Plus | |

Features:

```
Query  285       GTTCTCTCC  293
                 |||||||||
Sbjct  32597648  GTTCTCTCC  32597656
```

Range 37: 32603435 to 32603443

| Score | Expect | Identities | Gaps | Strand | Frame |
|---|---|---|---|---|---|
| 18.3 bits(9) | 14() | 9/9(100%) | 0/9(0%) | Plus/Minus | |

Features:

```
Query  288       CTCTCCTTT  296
                 |||||||||
Sbjct  32603443  CTCTCCTTT  32603435
```

Range 38: 32603516 to 32603524

| Score | Expect | Identities | Gaps | Strand | Frame |
|---|---|---|---|---|---|
| 18.3 bits(9) | 14() | 9/9(100%) | 0/9(0%) | Plus/Minus | |

Features:

```
Query  276       CAAGCTGGT  284
                 |||||||||
Sbjct  32603524  CAAGCTGGT  32603516
```

Range 39: 32607440 to 32607448

| Score | Expect | Identities | Gaps | Strand | Frame |
|---|---|---|---|---|---|
| 18.3 bits(9) | 14() | 9/9(100%) | 0/9(0%) | Plus/Plus | |

Features:

```
Query  291       TCCTTTATT  299
                 |||||||||
Sbjct  32607440  TCCTTTATT  32607448
```

Range 40: 32608410 to 32608418

| Score | Expect | Identities | Gaps | Strand | Frame |
|---|---|---|---|---|---|
| 18.3 bits(9) | 14() | 9/9(100%) | 0/9(0%) | Plus/Plus | |

Features:

```
Query  288       CTCTCCTTT  296
                 |||||||||
Sbjct  32608410  CTCTCCTTT  32608418
```

Range 41: 32610751 to 32610759

| Score | Expect | Identities | Gaps | Strand | Frame |
|---|---|---|---|---|---|
| 18.3 bits(9) | 14() | 9/9(100%) | 0/9(0%) | Plus/Plus | |

Features:

```
Query  286       TTCTCTCCT  294
                 |||||||||
Sbjct  32610751  TTCTCTCCT  32610759
```

Range 42: 32614047 to 32614055

| Score | Expect | Identities | Gaps | Strand | Frame |
|---|---|---|---|---|---|
| 18.3 bits(9) | 14() | 9/9(100%) | 0/9(0%) | Plus/Minus | |

Features:

```
Query  289       TCTCCTTTA  297
                 |||||||||
Sbjct  32614055  TCTCCTTTA  32614047
```

Range 43: 32614428 to 32614436

| Score | Expect | Identities | Gaps | Strand | Frame |
|---|---|---|---|---|---|
| 18.3 bits(9) | 14() | 9/9(100%) | 0/9(0%) | Plus/Plus | |

Features:

```
Query  291       TCCTTTATT  299
                 |||||||||
Sbjct  32614428  TCCTTTATT  32614436
```

Range 44: 32616905 to 32616913

| Score | Expect | Identities | Gaps | Strand | Frame |
|---|---|---|---|---|---|
| 18.3 bits(9) | 14() | 9/9(100%) | 0/9(0%) | Plus/Minus | |

Features:

```
Query  276       CAAGCTGGT  284
                 |||||||||
Sbjct  32616913  CAAGCTGGT  32616905
```

Range 45: 32359018 to 32359025

| Score | Expect | Identities | Gaps | Strand | Frame |
|---|---|---|---|---|---|
| 16.4 bits(8) | 56() | 8/8(100%) | 0/8(0%) | Plus/Minus | |

Features:

```
Query  281       TGGTGTTC  288
                 ||||||||
Sbjct  32359025  TGGTGTTC  32359018
```

Range 46: 32359087 to 32359094

| Score | Expect | Identities | Gaps | Strand | Frame |
|---|---|---|---|---|---|
| 16.4 bits(8) | 56() | 8/8(100%) | 0/8(0%) | Plus/Plus | |

Features:

```
Query  287       TCTCTCCT  294
                 ||||||||
Sbjct  32359087  TCTCTCCT  32359094
```

Range 47: 32362222 to 32362229

| Score | Expect | Identities | Gaps | Strand | Frame |
|---|---|---|---|---|---|
| 16.4 bits(8) | 56() | 8/8(100%) | 0/8(0%) | Plus/Plus | |

Features:

```
Query  289       TCTCCTTT  296
                 ||||||||
Sbjct  32362222  TCTCCTTT  32362229
```

Range 48: 32363967 to 32363974

| Score | Expect | Identities | Gaps | Strand | Frame |
|---|---|---|---|---|---|
| 16.4 bits(8) | 56() | 8/8(100%) | 0/8(0%) | Plus/Plus | |

Features:

```
Query  271       TGTAACAA  278
                 ||||||||
Sbjct  32363967  TGTAACAA  32363974
```

Range 49: 32365686 to 32365693

| Score | Expect | Identities | Gaps | Strand | Frame |
|---|---|---|---|---|---|

| | | | | |
|---|---|---|---|---|
| 16.4 bits(8) | 56() | 8/8(100%) | 0/8(0%) | Plus/Plus |

Features:

```
Query  272       GTAACAAG  279
                 ||||||||
Sbjct  32365686  GTAACAAG  32365693
```

Range 50: 32368034 to 32368041

| Score | Expect | Identities | Gaps | Strand | Frame |
|---|---|---|---|---|---|
| 16.4 bits(8) | 56() | 8/8(100%) | 0/8(0%) | Plus/Minus | |

Features:

```
Query  287       TCTCTCCT  294
                 ||||||||
Sbjct  32368041  TCTCTCCT  32368034
```

Range 51: 32376222 to 32376229

| Score | Expect | Identities | Gaps | Strand | Frame |
|---|---|---|---|---|---|
| 16.4 bits(8) | 56() | 8/8(100%) | 0/8(0%) | Plus/Minus | |

Features:

```
Query  287       TCTCTCCT  294
                 ||||||||
Sbjct  32376229  TCTCTCCT  32376222
```

Range 52: 32378046 to 32378053

| Score | Expect | Identities | Gaps | Strand | Frame |
|---|---|---|---|---|---|
| 16.4 bits(8) | 56() | 8/8(100%) | 0/8(0%) | Plus/Minus | |

Features:

```
Query  274       AACAAGCT  281
                 ||||||||
Sbjct  32378053  AACAAGCT  32378046
```

Range 53: 32378273 to 32378280

| Score | Expect | Identities | Gaps | Strand | Frame |
|---|---|---|---|---|---|
| 16.4 bits(8) | 56() | 8/8(100%) | 0/8(0%) | Plus/Plus | |

Features:

```
Query  284       TGTTCTCT  291
                 ||||||||
Sbjct  32378273  TGTTCTCT  32378280
```

Range 54: 32379868 to 32379875

| Score | Expect | Identities | Gaps | Strand | Frame |
|---|---|---|---|---|---|
| 16.4 bits(8) | 56() | 8/8(100%) | 0/8(0%) | Plus/Minus | |

Features:

```
Query  275       ACAAGCTG  282
                 ||||||||
Sbjct  32379875  ACAAGCTG  32379868
```

Range 55: 32382054 to 32382061

| Score | Expect | Identities | Gaps | Strand | Frame |
|---|---|---|---|---|---|
| 16.4 bits(8) | 56() | 8/8(100%) | 0/8(0%) | Plus/Plus | |

Features:

```
Query  280       CTGGTGTT  287
                 ||||||||
Sbjct  32382054  CTGGTGTT  32382061
```

Range 56: 32382080 to 32382087

| Score | Expect | Identities | Gaps | Strand | Frame |
|---|---|---|---|---|---|
| 16.4 bits(8) | 56() | 8/8(100%) | 0/8(0%) | Plus/Minus | |

Features:

```
Query  289       TCTCCTTT  296
                 ||||||||
Sbjct  32382087  TCTCCTTT  32382080
```

Range 57: 32383828 to 32383835

| Score | Expect | Identities | Gaps | Strand | Frame |
|---|---|---|---|---|---|
| 16.4 bits(8) | 56() | 8/8(100%) | 0/8(0%) | Plus/Plus | |

Features:

```
Query  278       AGCTGGTG  285
                 ||||||||
Sbjct  32383828  AGCTGGTG  32383835
```

Range 58: 32385126 to 32385133

| Score | Expect | Identities | Gaps | Strand | Frame |
|---|---|---|---|---|---|
| 16.4 bits(8) | 56() | 8/8(100%) | 0/8(0%) | Plus/Plus | |

Features:

```
Query  288       CTCTCCTT  295
                 ||||||||
Sbjct  32385126  CTCTCCTT  32385133
```

Range 59: 32385530 to 32385537

| Score | Expect | Identities | Gaps | Strand | Frame |
|---|---|---|---|---|---|
| 16.4 bits(8) | 56() | 8/8(100%) | 0/8(0%) | Plus/Minus | |

Features:

```
Query  289       TCTCCTTT  296
                 ||||||||
Sbjct  32385537  TCTCCTTT  32385530
```

Range 60: 32390071 to 32390078

| Score | Expect | Identities | Gaps | Strand | Frame |
|---|---|---|---|---|---|
| 16.4 bits(8) | 56() | 8/8(100%) | 0/8(0%) | Plus/Plus | |

Features:

```
Query  279       GCTGGTGT  286
                 ||||||||
Sbjct  32390071  GCTGGTGT  32390078
```

Range 61: 32390860 to 32390867

| Score | Expect | Identities | Gaps | Strand | Frame |
|---|---|---|---|---|---|
| 16.4 bits(8) | 56() | 8/8(100%) | 0/8(0%) | Plus/Minus | |

Features:

```
Query  271       TGTAACAA  278
                 ||||||||
Sbjct  32390867  TGTAACAA  32390860
```

Range 62: 32392130 to 32392137

| Score | Expect | Identities | Gaps | Strand | Frame |
|---|---|---|---|---|---|
| 16.4 bits(8) | 56() | 8/8(100%) | 0/8(0%) | Plus/Plus | |

Features:

```
Query  276       CAAGCTGG  283
                 ||||||||
Sbjct  32392130  CAAGCTGG  32392137
```

Range 63: 32392839 to 32392846

| Score | Expect | Identities | Gaps | Strand | Frame |
|---|---|---|---|---|---|
| 16.4 bits(8) | 56() | 8/8(100%) | 0/8(0%) | Plus/Plus | |

Features:

```
Query  280       CTGGTGTT  287
                 ||||||||
Sbjct  32392839  CTGGTGTT  32392846
```

Range 64: 32399652 to 32399659

| Score | Expect | Identities | Gaps | Strand | Frame |
|---|---|---|---|---|---|
| 16.4 bits(8) | 56() | 8/8(100%) | 0/8(0%) | Plus/Minus | |

Features:

```
Query  291       TCCTTTAT  298
                 ||||||||
Sbjct  32399659  TCCTTTAT  32399652
```

Range 65: 32405667 to 32405674

| Score | Expect | Identities | Gaps | Strand | Frame |
|---|---|---|---|---|---|
| 16.4 bits(8) | 56() | 8/8(100%) | 0/8(0%) | Plus/Minus | |

Features:

```
Query  290       CTCCTTTA  297
                 ||||||||
Sbjct  32405674  CTCCTTTA  32405667
```

Range 66: 32408642 to 32408649

| Score | Expect | Identities | Gaps | Strand | Frame |
|---|---|---|---|---|---|
| 16.4 bits(8) | 56() | 8/8(100%) | 0/8(0%) | Plus/Plus | |

Features:

```
Query  292       CCTTTATT  299
                 ||||||||
Sbjct  32408642  CCTTTATT  32408649
```

Range 67: 32412522 to 32412529

| Score | Expect | Identities | Gaps | Strand | Frame |
|---|---|---|---|---|---|
| 16.4 bits(8) | 56() | 8/8(100%) | 0/8(0%) | Plus/Minus | |

Features:

```
Query  292       CCTTTATT  299
                 ||||||||
Sbjct  32412529  CCTTTATT  32412522
```

Range 68: 32415717 to 32415724

| Score | Expect | Identities | Gaps | Strand | Frame |
|---|---|---|---|---|---|

| | | | | |
|---|---|---|---|---|
| 16.4 bits(8) | 56() | 8/8(100%) | 0/8(0%) | Plus/Plus |

Features:

```
Query  290       CTCCTTTA  297
                 ||||||||
Sbjct  32415717  CTCCTTTA  32415724
```

Range 69: 32416755 to 32416762

| Score | Expect | Identities | Gaps | Strand | Frame |
|---|---|---|---|---|---|
| 16.4 bits(8) | 56() | 8/8(100%) | 0/8(0%) | Plus/Plus | |

Features:

```
Query  289       TCTCCTTT  296
                 ||||||||
Sbjct  32416755  TCTCCTTT  32416762
```

Range 70: 32420961 to 32420968

| Score | Expect | Identities | Gaps | Strand | Frame |
|---|---|---|---|---|---|
| 16.4 bits(8) | 56() | 8/8(100%) | 0/8(0%) | Plus/Minus | |

Features:

```
Query  277       AAGCTGGT  284
                 ||||||||
Sbjct  32420968  AAGCTGGT  32420961
```

Range 71: 32422018 to 32422025

| Score | Expect | Identities | Gaps | Strand | Frame |
|---|---|---|---|---|---|
| 16.4 bits(8) | 56() | 8/8(100%) | 0/8(0%) | Plus/Minus | |

Features:

```
Query  271       TGTAACAA  278
                 ||||||||
Sbjct  32422025  TGTAACAA  32422018
```

Range 72: 32426837 to 32426844

| Score | Expect | Identities | Gaps | Strand | Frame |
|---|---|---|---|---|---|
| 16.4 bits(8) | 56() | 8/8(100%) | 0/8(0%) | Plus/Plus | |

Features:

```
Query  285       GTTCTCTC  292
                 ||||||||
Sbjct  32426837  GTTCTCTC  32426844
```

Range 73: 32428325 to 32428332

| Score | Expect | Identities | Gaps | Strand | Frame |
|---|---|---|---|---|---|
| 16.4 bits(8) | 56() | 8/8(100%) | 0/8(0%) | Plus/Plus | |

Features:

```
Query  280       CTGGTGTT  287
                 ||||||||
Sbjct  32428325  CTGGTGTT  32428332
```

Range 74: 32433958 to 32433965

| Score | Expect | Identities | Gaps | Strand | Frame |
|---|---|---|---|---|---|
| 16.4 bits(8) | 56() | 8/8(100%) | 0/8(0%) | Plus/Minus | |

Features:

```
Query  284       TGTTCTCT  291
                 ||||||||
Sbjct  32433965  TGTTCTCT  32433958
```

Range 75: 32435279 to 32435290

| Score | Expect | Identities | Gaps | Strand | Frame |
|---|---|---|---|---|---|
| 16.4 bits(8) | 56() | 11/12(92%) | 0/12(0%) | Plus/Minus | |

Features:

```
Query  286       TTCTCTCCTTTA  297
                 ||||||| ||||
Sbjct  32435290  TTCTCTCTTTTA  32435279
```

Range 76: 32435841 to 32435848

| Score | Expect | Identities | Gaps | Strand | Frame |
|---|---|---|---|---|---|
| 16.4 bits(8) | 56() | 8/8(100%) | 0/8(0%) | Plus/Minus | |

Features:

```
Query  292       CCTTTATT  299
                 ||||||||
Sbjct  32435848  CCTTTATT  32435841
```

Range 77: 32436161 to 32436168

| Score | Expect | Identities | Gaps | Strand | Frame |
|---|---|---|---|---|---|
| 16.4 bits(8) | 56() | 8/8(100%) | 0/8(0%) | Plus/Plus | |

Features:

```
Query  273       TAACAAGC  280
                 ||||||||
Sbjct  32436161  TAACAAGC  32436168
```

Range 78: 32439625 to 32439632

| Score | Expect | Identities | Gaps | Strand | Frame |
|---|---|---|---|---|---|
| 16.4 bits(8) | 56() | 8/8(100%) | 0/8(0%) | Plus/Minus | |

Features:

```
Query  280       CTGGTGTT  287
                 ||||||||
Sbjct  32439632  CTGGTGTT  32439625
```

Range 79: 32439634 to 32439641

| Score | Expect | Identities | Gaps | Strand | Frame |
|---|---|---|---|---|---|
| 16.4 bits(8) | 56() | 8/8(100%) | 0/8(0%) | Plus/Plus | |

Features:

```
Query  272       GTAACAAG  279
                 ||||||||
Sbjct  32439634  GTAACAAG  32439641
```

Range 80: 32442796 to 32442803

| Score | Expect | Identities | Gaps | Strand | Frame |
|---|---|---|---|---|---|
| 16.4 bits(8) | 56() | 8/8(100%) | 0/8(0%) | Plus/Plus | |

Features:

```
Query  271       TGTAACAA  278
                 ||||||||
Sbjct  32442796  TGTAACAA  32442803
```

Range 81: 32445587 to 32445594

| Score | Expect | Identities | Gaps | Strand | Frame |
|---|---|---|---|---|---|
| 16.4 bits(8) | 56() | 8/8(100%) | 0/8(0%) | Plus/Minus | |

Features:

```
Query  275       ACAAGCTG  282
                 ||||||||
Sbjct  32445594  ACAAGCTG  32445587
```

Range 82: 32446548 to 32446555

| Score | Expect | Identities | Gaps | Strand | Frame |
|---|---|---|---|---|---|
| 16.4 bits(8) | 56() | 8/8(100%) | 0/8(0%) | Plus/Minus | |

Features:

```
Query  286       TTCTCTCC  293
                 ||||||||
Sbjct  32446555  TTCTCTCC  32446548
```

Range 83: 32450020 to 32450027

| Score | Expect | Identities | Gaps | Strand | Frame |
|---|---|---|---|---|---|
| 16.4 bits(8) | 56() | 8/8(100%) | 0/8(0%) | Plus/Minus | |

Features:

```
Query  287       TCTCTCCT  294
                 ||||||||
Sbjct  32450027  TCTCTCCT  32450020
```

Range 84: 32451672 to 32451679

| Score | Expect | Identities | Gaps | Strand | Frame |
|---|---|---|---|---|---|
| 16.4 bits(8) | 56() | 8/8(100%) | 0/8(0%) | Plus/Plus | |

Features:

```
Query  275       ACAAGCTG  282
                 ||||||||
Sbjct  32451672  ACAAGCTG  32451679
```

Range 85: 32453521 to 32453528

| Score | Expect | Identities | Gaps | Strand | Frame |
|---|---|---|---|---|---|
| 16.4 bits(8) | 56() | 8/8(100%) | 0/8(0%) | Plus/Minus | |

Features:

```
Query  272       GTAACAAG  279
                 ||||||||
Sbjct  32453528  GTAACAAG  32453521
```

Range 86: 32454903 to 32454910

| Score | Expect | Identities | Gaps | Strand | Frame |
|---|---|---|---|---|---|
| 16.4 bits(8) | 56() | 8/8(100%) | 0/8(0%) | Plus/Plus | |

Features:

```
Query  291       TCCTTTAT  298
                 ||||||||
Sbjct  32454903  TCCTTTAT  32454910
```

Range 87: 32462433 to 32462440

| Score | Expect | Identities | Gaps | Strand | Frame |
|---|---|---|---|---|---|

| | | | | |
|---|---|---|---|---|
| 16.4 bits(8) | 56() | 8/8(100%) | 0/8(0%) | Plus/Plus |

Features:

```
Query  292       CCTTTATT  299
                 ||||||||
Sbjct  32462433  CCTTTATT  32462440
```

Range 88: 32463151 to 32463158

| Score | Expect | Identities | Gaps | Strand | Frame |
|---|---|---|---|---|---|
| 16.4 bits(8) | 56() | 8/8(100%) | 0/8(0%) | Plus/Plus | |

Features:

```
Query  280       CTGGTGTT  287
                 ||||||||
Sbjct  32463151  CTGGTGTT  32463158
```

Range 89: 32463899 to 32463906

| Score | Expect | Identities | Gaps | Strand | Frame |
|---|---|---|---|---|---|
| 16.4 bits(8) | 56() | 8/8(100%) | 0/8(0%) | Plus/Plus | |

Features:

```
Query  289       TCTCCTTT  296
                 ||||||||
Sbjct  32463899  TCTCCTTT  32463906
```

Range 90: 32466732 to 32466739

| Score | Expect | Identities | Gaps | Strand | Frame |
|---|---|---|---|---|---|
| 16.4 bits(8) | 56() | 8/8(100%) | 0/8(0%) | Plus/Minus | |

Features:

```
Query  274       AACAAGCT  281
                 ||||||||
Sbjct  32466739  AACAAGCT  32466732
```

Range 91: 32471051 to 32471058

| Score | Expect | Identities | Gaps | Strand | Frame |
|---|---|---|---|---|---|
| 16.4 bits(8) | 56() | 8/8(100%) | 0/8(0%) | Plus/Plus | |

Features:

```
Query  279       GCTGGTGT  286
                 ||||||||
Sbjct  32471051  GCTGGTGT  32471058
```

Range 92: 32473696 to 32473703

| Score | Expect | Identities | Gaps | Strand | Frame |
|---|---|---|---|---|---|
| 16.4 bits(8) | 56() | 8/8(100%) | 0/8(0%) | Plus/Plus | |

Features:

```
Query  289       TCTCCTTT  296
                 ||||||||
Sbjct  32473696  TCTCCTTT  32473703
```

Range 93: 32475324 to 32475331

| Score | Expect | Identities | Gaps | Strand | Frame |
|---|---|---|---|---|---|
| 16.4 bits(8) | 56() | 8/8(100%) | 0/8(0%) | Plus/Minus | |

Features:

```
Query  283       GTGTTCTC  290
                 ||||||||
Sbjct  32475331  GTGTTCTC  32475324
```

Range 94: 32476577 to 32476584

| Score | Expect | Identities | Gaps | Strand | Frame |
|---|---|---|---|---|---|
| 16.4 bits(8) | 56() | 8/8(100%) | 0/8(0%) | Plus/Minus | |

Features:

```
Query  291       TCCTTTAT  298
                 ||||||||
Sbjct  32476584  TCCTTTAT  32476577
```

Range 95: 32477531 to 32477538

| Score | Expect | Identities | Gaps | Strand | Frame |
|---|---|---|---|---|---|
| 16.4 bits(8) | 56() | 8/8(100%) | 0/8(0%) | Plus/Plus | |

Features:

```
Query  286       TTCTCTCC  293
                 ||||||||
Sbjct  32477531  TTCTCTCC  32477538
```

Range 96: 32478303 to 32478310

| Score | Expect | Identities | Gaps | Strand | Frame |
|---|---|---|---|---|---|
| 16.4 bits(8) | 56() | 8/8(100%) | 0/8(0%) | Plus/Minus | |

Features:

```
Query  279       GCTGGTGT  286
                 ||||||||
Sbjct  32478310  GCTGGTGT  32478303
```

Range 97: 32479532 to 32479539

| Score | Expect | Identities | Gaps | Strand | Frame |
|---|---|---|---|---|---|
| 16.4 bits(8) | 56() | 8/8(100%) | 0/8(0%) | Plus/Plus | |

Features:

```
Query  278       AGCTGGTG  285
                 ||||||||
Sbjct  32479532  AGCTGGTG  32479539
```

Range 98: 32480022 to 32480029

| Score | Expect | Identities | Gaps | Strand | Frame |
|---|---|---|---|---|---|
| 16.4 bits(8) | 56() | 8/8(100%) | 0/8(0%) | Plus/Plus | |

Features:

```
Query  291       TCCTTTAT  298
                 ||||||||
Sbjct  32480022  TCCTTTAT  32480029
```

Range 99: 32484986 to 32484993

| Score | Expect | Identities | Gaps | Strand | Frame |
|---|---|---|---|---|---|
| 16.4 bits(8) | 56() | 8/8(100%) | 0/8(0%) | Plus/Plus | |

Features:

```
Query  284       TGTTCTCT  291
                 ||||||||
Sbjct  32484986  TGTTCTCT  32484993
```

Range 100: 32485062 to 32485069

| Score | Expect | Identities | Gaps | Strand | Frame |
|---|---|---|---|---|---|
| 16.4 bits(8) | 56() | 8/8(100%) | 0/8(0%) | Plus/Minus | |

Features:

```
Query  290       CTCCTTTA  297
                 ||||||||
Sbjct  32485069  CTCCTTTA  32485062
```

Range 101: 32490500 to 32490507

| Score | Expect | Identities | Gaps | Strand | Frame |
|---|---|---|---|---|---|
| 16.4 bits(8) | 56() | 8/8(100%) | 0/8(0%) | Plus/Minus | |

Features:

```
Query  279       GCTGGTGT  286
                 ||||||||
Sbjct  32490507  GCTGGTGT  32490500
```

Range 102: 32492020 to 32492027

| Score | Expect | Identities | Gaps | Strand | Frame |
|---|---|---|---|---|---|
| 16.4 bits(8) | 56() | 8/8(100%) | 0/8(0%) | Plus/Plus | |

Features:

```
Query  291       TCCTTTAT  298
                 ||||||||
Sbjct  32492020  TCCTTTAT  32492027
```

Range 103: 32495412 to 32495419

| Score | Expect | Identities | Gaps | Strand | Frame |
|---|---|---|---|---|---|
| 16.4 bits(8) | 56() | 8/8(100%) | 0/8(0%) | Plus/Minus | |

Features:

```
Query  292       CCTTTATT  299
                 ||||||||
Sbjct  32495419  CCTTTATT  32495412
```

Range 104: 32495762 to 32495769

| Score | Expect | Identities | Gaps | Strand | Frame |
|---|---|---|---|---|---|
| 16.4 bits(8) | 56() | 8/8(100%) | 0/8(0%) | Plus/Minus | |

Features:

```
Query  273       TAACAAGC  280
                 ||||||||
Sbjct  32495769  TAACAAGC  32495762
```

Range 105: 32498400 to 32498407

| Score | Expect | Identities | Gaps | Strand | Frame |
|---|---|---|---|---|---|
| 16.4 bits(8) | 56() | 8/8(100%) | 0/8(0%) | Plus/Plus | |

Features:

```
Query  292       CCTTTATT  299
                 ||||||||
Sbjct  32498400  CCTTTATT  32498407
```

Range 106: 32503043 to 32503050

| Score | Expect | Identities | Gaps | Strand | Frame |
|---|---|---|---|---|---|

| 16.4 bits(8) | 56() | 8/8(100%) | 0/8(0%) | Plus/Plus |
|---|---|---|---|---|

Features:

```
Query  281       TGGTGTTC  288
                 ||||||||
Sbjct  32503043  TGGTGTTC  32503050
```

Range 107: 32505238 to 32505245

| Score | Expect | Identities | Gaps | Strand | Frame |
|---|---|---|---|---|---|
| 16.4 bits(8) | 56() | 8/8(100%) | 0/8(0%) | Plus/Plus | |

Features:

```
Query  274       AACAAGCT  281
                 ||||||||
Sbjct  32505238  AACAAGCT  32505245
```

Range 108: 32508081 to 32508088

| Score | Expect | Identities | Gaps | Strand | Frame |
|---|---|---|---|---|---|
| 16.4 bits(8) | 56() | 8/8(100%) | 0/8(0%) | Plus/Minus | |

Features:

```
Query  275       ACAAGCTG  282
                 ||||||||
Sbjct  32508088  ACAAGCTG  32508081
```

Range 109: 32510114 to 32510121

| Score | Expect | Identities | Gaps | Strand | Frame |
|---|---|---|---|---|---|
| 16.4 bits(8) | 56() | 8/8(100%) | 0/8(0%) | Plus/Plus | |

Features:

```
Query  287       TCTCTCCT  294
                 ||||||||
Sbjct  32510114  TCTCTCCT  32510121
```

Range 110: 32514520 to 32514527

| Score | Expect | Identities | Gaps | Strand | Frame |
|---|---|---|---|---|---|
| 16.4 bits(8) | 56() | 8/8(100%) | 0/8(0%) | Plus/Plus | |

Features:

```
Query  284       TGTTCTCT  291
                 ||||||||
Sbjct  32514520  TGTTCTCT  32514527
```

Range 111: 32515907 to 32515914

| Score | Expect | Identities | Gaps | Strand | Frame |
|---|---|---|---|---|---|
| 16.4 bits(8) | 56() | 8/8(100%) | 0/8(0%) | Plus/Plus | |

Features:

```
Query  289       TCTCCTTT  296
                 ||||||||
Sbjct  32515907  TCTCCTTT  32515914
```

Range 112: 32516344 to 32516351

| Score | Expect | Identities | Gaps | Strand | Frame |
|---|---|---|---|---|---|
| 16.4 bits(8) | 56() | 8/8(100%) | 0/8(0%) | Plus/Plus | |

Features:

```
Query  287       TCTCTCCT  294
                 ||||||||
Sbjct  32516344  TCTCTCCT  32516351
```

Range 113: 32518036 to 32518043

| Score | Expect | Identities | Gaps | Strand | Frame |
|---|---|---|---|---|---|
| 16.4 bits(8) | 56() | 8/8(100%) | 0/8(0%) | Plus/Plus | |

Features:

```
Query  289       TCTCCTTT  296
                 ||||||||
Sbjct  32518036  TCTCCTTT  32518043
```

Range 114: 32523884 to 32523891

| Score | Expect | Identities | Gaps | Strand | Frame |
|---|---|---|---|---|---|
| 16.4 bits(8) | 56() | 8/8(100%) | 0/8(0%) | Plus/Plus | |

Features:

```
Query  287       TCTCTCCT  294
                 ||||||||
Sbjct  32523884  TCTCTCCT  32523891
```

Range 115: 32524765 to 32524772

| Score | Expect | Identities | Gaps | Strand | Frame |
|---|---|---|---|---|---|
| 16.4 bits(8) | 56() | 8/8(100%) | 0/8(0%) | Plus/Plus | |

Features:

```
Query  291       TCCTTTAT  298
                 ||||||||
Sbjct  32524765  TCCTTTAT  32524772
```

Range 116: 32526986 to 32526993

| Score | Expect | Identities | Gaps | Strand | Frame |
|---|---|---|---|---|---|
| 16.4 bits(8) | 56() | 8/8(100%) | 0/8(0%) | Plus/Plus | |

Features:

```
Query  280       CTGGTGTT  287
                 ||||||||
Sbjct  32526986  CTGGTGTT  32526993
```

Range 117: 32535359 to 32535366

| Score | Expect | Identities | Gaps | Strand | Frame |
|---|---|---|---|---|---|
| 16.4 bits(8) | 56() | 8/8(100%) | 0/8(0%) | Plus/Minus | |

Features:

```
Query  271       TGTAACAA  278
                 ||||||||
Sbjct  32535366  TGTAACAA  32535359
```

Range 118: 32535509 to 32535516

| Score | Expect | Identities | Gaps | Strand | Frame |
|---|---|---|---|---|---|
| 16.4 bits(8) | 56() | 8/8(100%) | 0/8(0%) | Plus/Plus | |

Features:

```
Query  283       GTGTTCTC  290
                 ||||||||
Sbjct  32535509  GTGTTCTC  32535516
```

Range 119: 32538446 to 32538453

| Score | Expect | Identities | Gaps | Strand | Frame |
|---|---|---|---|---|---|
| 16.4 bits(8) | 56() | 8/8(100%) | 0/8(0%) | Plus/Minus | |

Features:

```
Query  292       CCTTTATT  299
                 ||||||||
Sbjct  32538453  CCTTTATT  32538446
```

Range 120: 32541143 to 32541150

| Score | Expect | Identities | Gaps | Strand | Frame |
|---|---|---|---|---|---|
| 16.4 bits(8) | 56() | 8/8(100%) | 0/8(0%) | Plus/Plus | |

Features:

```
Query  271       TGTAACAA  278
                 ||||||||
Sbjct  32541143  TGTAACAA  32541150
```

Range 121: 32542033 to 32542040

| Score | Expect | Identities | Gaps | Strand | Frame |
|---|---|---|---|---|---|
| 16.4 bits(8) | 56() | 8/8(100%) | 0/8(0%) | Plus/Plus | |

Features:

```
Query  271       TGTAACAA  278
                 ||||||||
Sbjct  32542033  TGTAACAA  32542040
```

Range 122: 32542578 to 32542585

| Score | Expect | Identities | Gaps | Strand | Frame |
|---|---|---|---|---|---|
| 16.4 bits(8) | 56() | 8/8(100%) | 0/8(0%) | Plus/Plus | |

Features:

```
Query  289       TCTCCTTT  296
                 ||||||||
Sbjct  32542578  TCTCCTTT  32542585
```

Range 123: 32542796 to 32542803

| Score | Expect | Identities | Gaps | Strand | Frame |
|---|---|---|---|---|---|
| 16.4 bits(8) | 56() | 8/8(100%) | 0/8(0%) | Plus/Minus | |

Features:

```
Query  274       AACAAGCT  281
                 ||||||||
Sbjct  32542803  AACAAGCT  32542796
```

Range 124: 32543103 to 32543110

| Score | Expect | Identities | Gaps | Strand | Frame |
|---|---|---|---|---|---|
| 16.4 bits(8) | 56() | 8/8(100%) | 0/8(0%) | Plus/Minus | |

Features:

```
Query  274       AACAAGCT  281
                 ||||||||
Sbjct  32543110  AACAAGCT  32543103
```

Range 125: 32544210 to 32544217

| Score | Expect | Identities | Gaps | Strand | Frame |
|---|---|---|---|---|---|

| | | | | |
|---|---|---|---|---|
| 16.4 bits(8) | 56() | 8/8(100%) | 0/8(0%) | Plus/Minus |

Features:

```
Query  292       CCTTTATT  299
                 ||||||||
Sbjct  32544217  CCTTTATT  32544210
```

Range 126: 32544300 to 32544307

| Score | Expect | Identities | Gaps | Strand | Frame |
|---|---|---|---|---|---|
| 16.4 bits(8) | 56() | 8/8(100%) | 0/8(0%) | Plus/Minus | |

Features:

```
Query  284       TGTTCTCT  291
                 ||||||||
Sbjct  32544307  TGTTCTCT  32544300
```

Range 127: 32544735 to 32544742

| Score | Expect | Identities | Gaps | Strand | Frame |
|---|---|---|---|---|---|
| 16.4 bits(8) | 56() | 8/8(100%) | 0/8(0%) | Plus/Plus | |

Features:

```
Query  291       TCCTTTAT  298
                 ||||||||
Sbjct  32544735  TCCTTTAT  32544742
```

Range 128: 32545705 to 32545712

| Score | Expect | Identities | Gaps | Strand | Frame |
|---|---|---|---|---|---|
| 16.4 bits(8) | 56() | 8/8(100%) | 0/8(0%) | Plus/Minus | |

Features:

```
Query  274       AACAAGCT  281
                 ||||||||
Sbjct  32545712  AACAAGCT  32545705
```

Range 129: 32547355 to 32547362

| Score | Expect | Identities | Gaps | Strand | Frame |
|---|---|---|---|---|---|
| 16.4 bits(8) | 56() | 8/8(100%) | 0/8(0%) | Plus/Plus | |

Features:

```
Query  286       TTCTCTCC  293
                 ||||||||
Sbjct  32547355  TTCTCTCC  32547362
```

Range 130: 32550134 to 32550141

| Score | Expect | Identities | Gaps | Strand | Frame |
|---|---|---|---|---|---|
| 16.4 bits(8) | 56() | 8/8(100%) | 0/8(0%) | Plus/Plus | |

Features:

```
Query  292       CCTTTATT  299
                 ||||||||
Sbjct  32550134  CCTTTATT  32550141
```

Range 131: 32558758 to 32558765

| Score | Expect | Identities | Gaps | Strand | Frame |
|---|---|---|---|---|---|
| 16.4 bits(8) | 56() | 8/8(100%) | 0/8(0%) | Plus/Plus | |

Features:

```
Query  274       AACAAGCT  281
                 ||||||||
Sbjct  32558758  AACAAGCT  32558765
```

Range 132: 32559325 to 32559332

| Score | Expect | Identities | Gaps | Strand | Frame |
|---|---|---|---|---|---|
| 16.4 bits(8) | 56() | 8/8(100%) | 0/8(0%) | Plus/Minus | |

Features:

```
Query  290       CTCCTTTA  297
                 ||||||||
Sbjct  32559332  CTCCTTTA  32559325
```

Range 133: 32559438 to 32559445

| Score | Expect | Identities | Gaps | Strand | Frame |
|---|---|---|---|---|---|
| 16.4 bits(8) | 56() | 8/8(100%) | 0/8(0%) | Plus/Minus | |

Features:

```
Query  290       CTCCTTTA  297
                 ||||||||
Sbjct  32559445  CTCCTTTA  32559438
```

Range 134: 32565692 to 32565699

| Score | Expect | Identities | Gaps | Strand | Frame |
|---|---|---|---|---|---|
| 16.4 bits(8) | 56() | 8/8(100%) | 0/8(0%) | Plus/Minus | |

Features:

```
Query  291       TCCTTTAT  298
                 ||||||||
Sbjct  32565699  TCCTTTAT  32565692
```

Range 135: 32568667 to 32568678

| Score | Expect | Identities | Gaps | Strand | Frame |
|---|---|---|---|---|---|
| 16.4 bits(8) | 56() | 11/12(92%) | 0/12(0%) | Plus/Minus | |

Features:

```
Query  276       CAAGCTGGTGTT  287
                 |||| |||||||
Sbjct  32568678  CAAGATGGTGTT  32568667
```

Range 136: 32570073 to 32570080

| Score | Expect | Identities | Gaps | Strand | Frame |
|---|---|---|---|---|---|
| 16.4 bits(8) | 56() | 8/8(100%) | 0/8(0%) | Plus/Plus | |

Features:

```
Query  292       CCTTTATT  299
                 ||||||||
Sbjct  32570073  CCTTTATT  32570080
```

Range 137: 32570331 to 32570338

| Score | Expect | Identities | Gaps | Strand | Frame |
|---|---|---|---|---|---|
| 16.4 bits(8) | 56() | 8/8(100%) | 0/8(0%) | Plus/Minus | |

Features:

```
Query  275       ACAAGCTG  282
                 ||||||||
Sbjct  32570338  ACAAGCTG  32570331
```

Range 138: 32571437 to 32571444

| Score | Expect | Identities | Gaps | Strand | Frame |
|---|---|---|---|---|---|
| 16.4 bits(8) | 56() | 8/8(100%) | 0/8(0%) | Plus/Minus | |

Features:

```
Query  273       TAACAAGC  280
                 ||||||||
Sbjct  32571444  TAACAAGC  32571437
```

Range 139: 32571691 to 32571698

| Score | Expect | Identities | Gaps | Strand | Frame |
|---|---|---|---|---|---|
| 16.4 bits(8) | 56() | 8/8(100%) | 0/8(0%) | Plus/Minus | |

Features:

```
Query  277       AAGCTGGT  284
                 ||||||||
Sbjct  32571698  AAGCTGGT  32571691
```

Range 140: 32572448 to 32572455

| Score | Expect | Identities | Gaps | Strand | Frame |
|---|---|---|---|---|---|
| 16.4 bits(8) | 56() | 8/8(100%) | 0/8(0%) | Plus/Plus | |

Features:

```
Query  272       GTAACAAG  279
                 ||||||||
Sbjct  32572448  GTAACAAG  32572455
```

Range 141: 32577875 to 32577882

| Score | Expect | Identities | Gaps | Strand | Frame |
|---|---|---|---|---|---|
| 16.4 bits(8) | 56() | 8/8(100%) | 0/8(0%) | Plus/Minus | |

Features:

```
Query  284       TGTTCTCT  291
                 ||||||||
Sbjct  32577882  TGTTCTCT  32577875
```

Range 142: 32580269 to 32580276

| Score | Expect | Identities | Gaps | Strand | Frame |
|---|---|---|---|---|---|
| 16.4 bits(8) | 56() | 8/8(100%) | 0/8(0%) | Plus/Plus | |

Features:

```
Query  271       TGTAACAA  278
                 ||||||||
Sbjct  32580269  TGTAACAA  32580276
```

Range 143: 32588743 to 32588750

| Score | Expect | Identities | Gaps | Strand | Frame |
|---|---|---|---|---|---|
| 16.4 bits(8) | 56() | 8/8(100%) | 0/8(0%) | Plus/Minus | |

Features:

```
Query  292       CCTTTATT  299
                 ||||||||
Sbjct  32588750  CCTTTATT  32588743
```

Range 144: 32592119 to 32592126

| Score | Expect | Identities | Gaps | Strand | Frame |
|---|---|---|---|---|---|

| Score | Expect | Identities | Gaps | Strand | Frame |
|---|---|---|---|---|---|
| 16.4 bits(8) | 56() | 8/8(100%) | 0/8(0%) | Plus/Plus | |

Features:

```
Query  288       CTCTCCTT  295
                 ||||||||
Sbjct  32592119  CTCTCCTT  32592126
```

Range 145: 32594190 to 32594197

| Score | Expect | Identities | Gaps | Strand | Frame |
|---|---|---|---|---|---|
| 16.4 bits(8) | 56() | 8/8(100%) | 0/8(0%) | Plus/Plus | |

Features:

```
Query  284       TGTTCTCT  291
                 ||||||||
Sbjct  32594190  TGTTCTCT  32594197
```

Range 146: 32596496 to 32596503

| Score | Expect | Identities | Gaps | Strand | Frame |
|---|---|---|---|---|---|
| 16.4 bits(8) | 56() | 8/8(100%) | 0/8(0%) | Plus/Plus | |

Features:

```
Query  292       CCTTTATT  299
                 ||||||||
Sbjct  32596496  CCTTTATT  32596503
```

Range 147: 32596968 to 32596975

| Score | Expect | Identities | Gaps | Strand | Frame |
|---|---|---|---|---|---|
| 16.4 bits(8) | 56() | 8/8(100%) | 0/8(0%) | Plus/Plus | |

Features:

```
Query  285       GTTCTCTC  292
                 ||||||||
Sbjct  32596968  GTTCTCTC  32596975
```

Range 148: 32597740 to 32597747

| Score | Expect | Identities | Gaps | Strand | Frame |
|---|---|---|---|---|---|
| 16.4 bits(8) | 56() | 8/8(100%) | 0/8(0%) | Plus/Plus | |

Features:

```
Query  289       TCTCCTTT  296
                 ||||||||
Sbjct  32597740  TCTCCTTT  32597747
```

Range 149: 32598172 to 32598179

| Score | Expect | Identities | Gaps | Strand | Frame |
|---|---|---|---|---|---|
| 16.4 bits(8) | 56() | 8/8(100%) | 0/8(0%) | Plus/Plus | |

Features:

```
Query  276       CAAGCTGG  283
                 ||||||||
Sbjct  32598172  CAAGCTGG  32598179
```

Range 150: 32598326 to 32598333

| Score | Expect | Identities | Gaps | Strand | Frame |
|---|---|---|---|---|---|
| 16.4 bits(8) | 56() | 8/8(100%) | 0/8(0%) | Plus/Plus | |

Features:

```
Query  277       AAGCTGGT  284
                 ||||||||
Sbjct  32598326  AAGCTGGT  32598333
```

Range 151: 32605374 to 32605381

| Score | Expect | Identities | Gaps | Strand | Frame |
|---|---|---|---|---|---|
| 16.4 bits(8) | 56() | 8/8(100%) | 0/8(0%) | Plus/Minus | |

Features:

```
Query  289       TCTCCTTT  296
                 ||||||||
Sbjct  32605381  TCTCCTTT  32605374
```

Range 152: 32610550 to 32610557

| Score | Expect | Identities | Gaps | Strand | Frame |
|---|---|---|---|---|---|
| 16.4 bits(8) | 56() | 8/8(100%) | 0/8(0%) | Plus/Plus | |

Features:

```
Query  282       GGTGTTCT  289
                 ||||||||
Sbjct  32610550  GGTGTTCT  32610557
```

Range 153: 32612498 to 32612505

| Score | Expect | Identities | Gaps | Strand | Frame |
|---|---|---|---|---|---|
| 16.4 bits(8) | 56() | 8/8(100%) | 0/8(0%) | Plus/Minus | |

Features:

```
Query  273       TAACAAGC  280
                 ||||||||
Sbjct  32612505  TAACAAGC  32612498
```

Range 154: 32618767 to 32618774

| Score | Expect | Identities | Gaps | Strand | Frame |
|---|---|---|---|---|---|
| 16.4 bits(8) | 56() | 8/8(100%) | 0/8(0%) | Plus/Minus | |

Features:

```
Query  287       TCTCTCCT  294
                 ||||||||
Sbjct  32618774  TCTCTCCT  32618767
```

Range 155: 32357326 to 32357332

| Score | Expect | Identities | Gaps | Strand | Frame |
|---|---|---|---|---|---|
| 14.4 bits(7) | 223() | 7/7(100%) | 0/7(0%) | Plus/Plus | |

Features:

```
Query  279       GCTGGTG  285
                 |||||||
Sbjct  32357326  GCTGGTG  32357332
```

Range 156: 32357892 to 32357898

| Score | Expect | Identities | Gaps | Strand | Frame |
|---|---|---|---|---|---|
| 14.4 bits(7) | 223() | 7/7(100%) | 0/7(0%) | Plus/Plus | |

Features:

```
Query  279       GCTGGTG  285
                 |||||||
Sbjct  32357892  GCTGGTG  32357898
```

Range 157: 32357974 to 32357980

| Score | Expect | Identities | Gaps | Strand | Frame |
|---|---|---|---|---|---|
| 14.4 bits(7) | 223() | 7/7(100%) | 0/7(0%) | Plus/Plus | |

Features:

```
Query  284       TGTTCTC  290
                 |||||||
Sbjct  32357974  TGTTCTC  32357980
```

Range 158: 32358325 to 32358331

| Score | Expect | Identities | Gaps | Strand | Frame |
|---|---|---|---|---|---|
| 14.4 bits(7) | 223() | 7/7(100%) | 0/7(0%) | Plus/Minus | |

Features:

```
Query  288       CTCTCCT  294
                 |||||||
Sbjct  32358331  CTCTCCT  32358325
```

Range 159: 32360039 to 32360045

| Score | Expect | Identities | Gaps | Strand | Frame |
|---|---|---|---|---|---|
| 14.4 bits(7) | 223() | 7/7(100%) | 0/7(0%) | Plus/Minus | |

Features:

```
Query  286       TTCTCTC  292
                 |||||||
Sbjct  32360045  TTCTCTC  32360039
```

Range 160: 32360492 to 32360498

| Score | Expect | Identities | Gaps | Strand | Frame |
|---|---|---|---|---|---|
| 14.4 bits(7) | 223() | 7/7(100%) | 0/7(0%) | Plus/Plus | |

Features:

```
Query  289       TCTCCTT  295
                 |||||||
Sbjct  32360492  TCTCCTT  32360498
```

Range 161: 32361564 to 32361570

| Score | Expect | Identities | Gaps | Strand | Frame |
|---|---|---|---|---|---|
| 14.4 bits(7) | 223() | 7/7(100%) | 0/7(0%) | Plus/Plus | |

Features:

```
Query  281       TGGTGTT  287
                 |||||||
Sbjct  32361564  TGGTGTT  32361570
```

Range 162: 32362168 to 32362174

| Score | Expect | Identities | Gaps | Strand | Frame |
|---|---|---|---|---|---|
| 14.4 bits(7) | 223() | 7/7(100%) | 0/7(0%) | Plus/Minus | |

Features:

```
Query  279       GCTGGTG  285
                 |||||||
Sbjct  32362174  GCTGGTG  32362168
```

Range 163: 32362735 to 32362741

| Score | Expect | Identities | Gaps | Strand | Frame |
|---|---|---|---|---|---|

| Score | Expect | Identities | Gaps | Strand | Frame |
|---|---|---|---|---|---|
| 14.4 bits(7) | 223() | 7/7(100%) | 0/7(0%) | Plus/Minus | |

Features:

```
Query  290       CTCCTTT  296
                 |||||||
Sbjct  32362741  CTCCTTT  32362735
```

Range 164: 32363631 to 32363637

| Score | Expect | Identities | Gaps | Strand | Frame |
|---|---|---|---|---|---|
| 14.4 bits(7) | 223() | 7/7(100%) | 0/7(0%) | Plus/Minus | |

Features:

```
Query  287       TCTCTCC  293
                 |||||||
Sbjct  32363637  TCTCTCC  32363631
```

Range 165: 32364548 to 32364554

| Score | Expect | Identities | Gaps | Strand | Frame |
|---|---|---|---|---|---|
| 14.4 bits(7) | 223() | 7/7(100%) | 0/7(0%) | Plus/Plus | |

Features:

```
Query  283       GTGTTCT  289
                 |||||||
Sbjct  32364548  GTGTTCT  32364554
```

Range 166: 32366279 to 32366285

| Score | Expect | Identities | Gaps | Strand | Frame |
|---|---|---|---|---|---|
| 14.4 bits(7) | 223() | 7/7(100%) | 0/7(0%) | Plus/Minus | |

Features:

```
Query  288       CTCTCCT  294
                 |||||||
Sbjct  32366285  CTCTCCT  32366279
```

Range 167: 32366393 to 32366399

| Score | Expect | Identities | Gaps | Strand | Frame |
|---|---|---|---|---|---|
| 14.4 bits(7) | 223() | 7/7(100%) | 0/7(0%) | Plus/Plus | |

Features:

```
Query  293       CTTTATT  299
                 |||||||
Sbjct  32366393  CTTTATT  32366399
```

Range 168: 32366399 to 32366405

| Score | Expect | Identities | Gaps | Strand | Frame |
|---|---|---|---|---|---|
| 14.4 bits(7) | 223() | 7/7(100%) | 0/7(0%) | Plus/Plus | |

Features:

```
Query  291       TCCTTTA  297
                 |||||||
Sbjct  32366399  TCCTTTA  32366405
```

Range 169: 32367453 to 32367459

| Score | Expect | Identities | Gaps | Strand | Frame |
|---|---|---|---|---|---|
| 14.4 bits(7) | 223() | 7/7(100%) | 0/7(0%) | Plus/Minus | |

Features:

```
Query  281       TGGTGTT  287
                 |||||||
Sbjct  32367459  TGGTGTT  32367453
```

Range 170: 32368215 to 32368221

| Score | Expect | Identities | Gaps | Strand | Frame |
|---|---|---|---|---|---|
| 14.4 bits(7) | 223() | 7/7(100%) | 0/7(0%) | Plus/Plus | |

Features:

```
Query  278       AGCTGGT  284
                 |||||||
Sbjct  32368215  AGCTGGT  32368221
```

Range 171: 32368872 to 32368878

| Score | Expect | Identities | Gaps | Strand | Frame |
|---|---|---|---|---|---|
| 14.4 bits(7) | 223() | 7/7(100%) | 0/7(0%) | Plus/Minus | |

Features:

```
Query  282       GGTGTTC  288
                 |||||||
Sbjct  32368878  GGTGTTC  32368872
```

Range 172: 32369074 to 32369080

| Score | Expect | Identities | Gaps | Strand | Frame |
|---|---|---|---|---|---|
| 14.4 bits(7) | 223() | 7/7(100%) | 0/7(0%) | Plus/Plus | |

Features:

```
Query  281       TGGTGTT  287
                 |||||||
Sbjct  32369074  TGGTGTT  32369080
```

Range 173: 32369752 to 32369758

| Score | Expect | Identities | Gaps | Strand | Frame |
|---|---|---|---|---|---|
| 14.4 bits(7) | 223() | 7/7(100%) | 0/7(0%) | Plus/Plus | |

Features:

```
Query  285       GTTCTCT  291
                 |||||||
Sbjct  32369752  GTTCTCT  32369758
```

Range 174: 32370113 to 32370119

| Score | Expect | Identities | Gaps | Strand | Frame |
|---|---|---|---|---|---|
| 14.4 bits(7) | 223() | 7/7(100%) | 0/7(0%) | Plus/Minus | |

Features:

```
Query  291       TCCTTTA  297
                 |||||||
Sbjct  32370119  TCCTTTA  32370113
```

Range 175: 32370434 to 32370440

| Score | Expect | Identities | Gaps | Strand | Frame |
|---|---|---|---|---|---|
| 14.4 bits(7) | 223() | 7/7(100%) | 0/7(0%) | Plus/Plus | |

Features:

```
Query  285       GTTCTCT  291
                 |||||||
Sbjct  32370434  GTTCTCT  32370440
```

Range 176: 32370779 to 32370785

| Score | Expect | Identities | Gaps | Strand | Frame |
|---|---|---|---|---|---|
| 14.4 bits(7) | 223() | 7/7(100%) | 0/7(0%) | Plus/Plus | |

Features:

```
Query  293       CTTTATT  299
                 |||||||
Sbjct  32370779  CTTTATT  32370785
```

Range 177: 32370915 to 32370921

| Score | Expect | Identities | Gaps | Strand | Frame |
|---|---|---|---|---|---|
| 14.4 bits(7) | 223() | 7/7(100%) | 0/7(0%) | Plus/Plus | |

Features:

```
Query  271       TGTAACA  277
                 |||||||
Sbjct  32370915  TGTAACA  32370921
```

Range 178: 32372114 to 32372120

| Score | Expect | Identities | Gaps | Strand | Frame |
|---|---|---|---|---|---|
| 14.4 bits(7) | 223() | 7/7(100%) | 0/7(0%) | Plus/Plus | |

Features:

```
Query  291       TCCTTTA  297
                 |||||||
Sbjct  32372114  TCCTTTA  32372120
```

Range 179: 32372335 to 32372341

| Score | Expect | Identities | Gaps | Strand | Frame |
|---|---|---|---|---|---|
| 14.4 bits(7) | 223() | 7/7(100%) | 0/7(0%) | Plus/Plus | |

Features:

```
Query  284       TGTTCTC  290
                 |||||||
Sbjct  32372335  TGTTCTC  32372341
```

Range 180: 32372370 to 32372376

| Score | Expect | Identities | Gaps | Strand | Frame |
|---|---|---|---|---|---|
| 14.4 bits(7) | 223() | 7/7(100%) | 0/7(0%) | Plus/Minus | |

Features:

```
Query  272       GTAACAA  278
                 |||||||
Sbjct  32372376  GTAACAA  32372370
```

Range 181: 32373052 to 32373058

| Score | Expect | Identities | Gaps | Strand | Frame |
|---|---|---|---|---|---|
| 14.4 bits(7) | 223() | 7/7(100%) | 0/7(0%) | Plus/Plus | |

Features:

```
Query  291       TCCTTTA  297
                 |||||||
Sbjct  32373052  TCCTTTA  32373058
```

Range 182: 32373557 to 32373563

| Score | Expect | Identities | Gaps | Strand | Frame |
|---|---|---|---|---|---|

| Score | Expect | Identities | Gaps | Strand | Frame |
|---|---|---|---|---|---|
| 14.4 bits(7) | 223() | 7/7(100%) | 0/7(0%) | Plus/Minus | |

Features:

```
Query  281       TGGTGTT  287
                 |||||||
Sbjct  32373563  TGGTGTT  32373557
```

Range 183: 32373632 to 32373638

| Score | Expect | Identities | Gaps | Strand | Frame |
|---|---|---|---|---|---|
| 14.4 bits(7) | 223() | 7/7(100%) | 0/7(0%) | Plus/Plus | |

Features:

```
Query  279       GCTGGTG  285
                 |||||||
Sbjct  32373632  GCTGGTG  32373638
```

Range 184: 32373771 to 32373777

| Score | Expect | Identities | Gaps | Strand | Frame |
|---|---|---|---|---|---|
| 14.4 bits(7) | 223() | 7/7(100%) | 0/7(0%) | Plus/Minus | |

Features:

```
Query  289       TCTCCTT  295
                 |||||||
Sbjct  32373777  TCTCCTT  32373771
```

Range 185: 32373785 to 32373791

| Score | Expect | Identities | Gaps | Strand | Frame |
|---|---|---|---|---|---|
| 14.4 bits(7) | 223() | 7/7(100%) | 0/7(0%) | Plus/Minus | |

Features:

```
Query  280       CTGGTGT  286
                 |||||||
Sbjct  32373791  CTGGTGT  32373785
```

Range 186: 32374094 to 32374100

| Score | Expect | Identities | Gaps | Strand | Frame |
|---|---|---|---|---|---|
| 14.4 bits(7) | 223() | 7/7(100%) | 0/7(0%) | Plus/Plus | |

Features:

```
Query  292       CCTTTAT  298
                 |||||||
Sbjct  32374094  CCTTTAT  32374100
```

Range 187: 32375735 to 32375741

| Score | Expect | Identities | Gaps | Strand | Frame |
|---|---|---|---|---|---|
| 14.4 bits(7) | 223() | 7/7(100%) | 0/7(0%) | Plus/Plus | |

Features:

```
Query  286       TTCTCTC  292
                 |||||||
Sbjct  32375735  TTCTCTC  32375741
```

Range 188: 32376752 to 32376758

| Score | Expect | Identities | Gaps | Strand | Frame |
|---|---|---|---|---|---|
| 14.4 bits(7) | 223() | 7/7(100%) | 0/7(0%) | Plus/Minus | |

Features:

```
Query  293       CTTTATT  299
                 |||||||
Sbjct  32376758  CTTTATT  32376752
```

Range 189: 32377124 to 32377130

| Score | Expect | Identities | Gaps | Strand | Frame |
|---|---|---|---|---|---|
| 14.4 bits(7) | 223() | 7/7(100%) | 0/7(0%) | Plus/Minus | |

Features:

```
Query  271       TGTAACA  277
                 |||||||
Sbjct  32377130  TGTAACA  32377124
```

Range 190: 32377746 to 32377752

| Score | Expect | Identities | Gaps | Strand | Frame |
|---|---|---|---|---|---|
| 14.4 bits(7) | 223() | 7/7(100%) | 0/7(0%) | Plus/Minus | |

Features:

```
Query  276       CAAGCTG  282
                 |||||||
Sbjct  32377752  CAAGCTG  32377746
```

Range 191: 32377849 to 32377855

| Score | Expect | Identities | Gaps | Strand | Frame |
|---|---|---|---|---|---|
| 14.4 bits(7) | 223() | 7/7(100%) | 0/7(0%) | Plus/Plus | |

Features:

```
Query  291       TCCTTTA  297
                 |||||||
Sbjct  32377849  TCCTTTA  32377855
```

Range 192: 32377962 to 32377968

| Score | Expect | Identities | Gaps | Strand | Frame |
|---|---|---|---|---|---|
| 14.4 bits(7) | 223() | 7/7(100%) | 0/7(0%) | Plus/Plus | |

Features:

```
Query  281       TGGTGTT  287
                 |||||||
Sbjct  32377962  TGGTGTT  32377968
```

Range 193: 32378188 to 32378194

| Score | Expect | Identities | Gaps | Strand | Frame |
|---|---|---|---|---|---|
| 14.4 bits(7) | 223() | 7/7(100%) | 0/7(0%) | Plus/Minus | |

Features:

```
Query  288       CTCTCCT  294
                 |||||||
Sbjct  32378194  CTCTCCT  32378188
```

Range 194: 32378253 to 32378259

| Score | Expect | Identities | Gaps | Strand | Frame |
|---|---|---|---|---|---|
| 14.4 bits(7) | 223() | 7/7(100%) | 0/7(0%) | Plus/Plus | |

Features:

```
Query  288       CTCTCCT  294
                 |||||||
Sbjct  32378253  CTCTCCT  32378259
```

Range 195: 32378362 to 32378368

| Score | Expect | Identities | Gaps | Strand | Frame |
|---|---|---|---|---|---|
| 14.4 bits(7) | 223() | 7/7(100%) | 0/7(0%) | Plus/Plus | |

Features:

```
Query  276       CAAGCTG  282
                 |||||||
Sbjct  32378362  CAAGCTG  32378368
```

Range 196: 32378979 to 32378985

| Score | Expect | Identities | Gaps | Strand | Frame |
|---|---|---|---|---|---|
| 14.4 bits(7) | 223() | 7/7(100%) | 0/7(0%) | Plus/Plus | |

Features:

```
Query  283       GTGTTCT  289
                 |||||||
Sbjct  32378979  GTGTTCT  32378985
```

Range 197: 32378988 to 32378994

| Score | Expect | Identities | Gaps | Strand | Frame |
|---|---|---|---|---|---|
| 14.4 bits(7) | 223() | 7/7(100%) | 0/7(0%) | Plus/Minus | |

Features:

```
Query  293       CTTTATT  299
                 |||||||
Sbjct  32378994  CTTTATT  32378988
```

Range 198: 32379021 to 32379027

| Score | Expect | Identities | Gaps | Strand | Frame |
|---|---|---|---|---|---|
| 14.4 bits(7) | 223() | 7/7(100%) | 0/7(0%) | Plus/Minus | |

Features:

```
Query  273       TAACAAG  279
                 |||||||
Sbjct  32379027  TAACAAG  32379021
```

Range 199: 32379053 to 32379059

| Score | Expect | Identities | Gaps | Strand | Frame |
|---|---|---|---|---|---|
| 14.4 bits(7) | 223() | 7/7(100%) | 0/7(0%) | Plus/Minus | |

Features:

```
Query  293       CTTTATT  299
                 |||||||
Sbjct  32379059  CTTTATT  32379053
```

Range 200: 32379314 to 32379320

| Score | Expect | Identities | Gaps | Strand | Frame |
|---|---|---|---|---|---|
| 14.4 bits(7) | 223() | 7/7(100%) | 0/7(0%) | Plus/Minus | |

Features:

```
Query  278       AGCTGGT  284
                 |||||||
Sbjct  32379320  AGCTGGT  32379314
```

Range 201: 32379591 to 32379597

| Score | Expect | Identities | Gaps | Strand | Frame |
|---|---|---|---|---|---|

| | | | | |
|---|---|---|---|---|
| 14.4 bits(7) | 223() | 7/7(100%) | 0/7(0%) | Plus/Plus |

Features:

```
Query  291       TCCTTTA  297
                 |||||||
Sbjct  32379591  TCCTTTA  32379597
```

Range 202: 32379690 to 32379696

| Score | Expect | Identities | Gaps | Strand | Frame |
|---|---|---|---|---|---|
| 14.4 bits(7) | 223() | 7/7(100%) | 0/7(0%) | Plus/Plus | |

Features:

```
Query  293       CTTTATT  299
                 |||||||
Sbjct  32379690  CTTTATT  32379696
```

Range 203: 32380889 to 32380895

| Score | Expect | Identities | Gaps | Strand | Frame |
|---|---|---|---|---|---|
| 14.4 bits(7) | 223() | 7/7(100%) | 0/7(0%) | Plus/Minus | |

Features:

```
Query  273       TAACAAG  279
                 |||||||
Sbjct  32380895  TAACAAG  32380889
```

Range 204: 32383534 to 32383540

| Score | Expect | Identities | Gaps | Strand | Frame |
|---|---|---|---|---|---|
| 14.4 bits(7) | 223() | 7/7(100%) | 0/7(0%) | Plus/Plus | |

Features:

```
Query  293       CTTTATT  299
                 |||||||
Sbjct  32383534  CTTTATT  32383540
```

Range 205: 32383854 to 32383860

| Score | Expect | Identities | Gaps | Strand | Frame |
|---|---|---|---|---|---|
| 14.4 bits(7) | 223() | 7/7(100%) | 0/7(0%) | Plus/Minus | |

Features:

```
Query  272       GTAACAA  278
                 |||||||
Sbjct  32383860  GTAACAA  32383854
```

Range 206: 32383930 to 32383936

| Score | Expect | Identities | Gaps | Strand | Frame |
|---|---|---|---|---|---|
| 14.4 bits(7) | 223() | 7/7(100%) | 0/7(0%) | Plus/Plus | |

Features:

```
Query  289       TCTCCTT  295
                 |||||||
Sbjct  32383930  TCTCCTT  32383936
```

Range 207: 32384334 to 32384340

| Score | Expect | Identities | Gaps | Strand | Frame |
|---|---|---|---|---|---|
| 14.4 bits(7) | 223() | 7/7(100%) | 0/7(0%) | Plus/Minus | |

Features:

```
Query  277       AAGCTGG  283
                 |||||||
Sbjct  32384340  AAGCTGG  32384334
```

Range 208: 32384809 to 32384815

| Score | Expect | Identities | Gaps | Strand | Frame |
|---|---|---|---|---|---|
| 14.4 bits(7) | 223() | 7/7(100%) | 0/7(0%) | Plus/Minus | |

Features:

```
Query  287       TCTCTCC  293
                 |||||||
Sbjct  32384815  TCTCTCC  32384809
```

Range 209: 32385402 to 32385408

| Score | Expect | Identities | Gaps | Strand | Frame |
|---|---|---|---|---|---|
| 14.4 bits(7) | 223() | 7/7(100%) | 0/7(0%) | Plus/Plus | |

Features:

```
Query  279       GCTGGTG  285
                 |||||||
Sbjct  32385402  GCTGGTG  32385408
```

Range 210: 32385876 to 32385882

| Score | Expect | Identities | Gaps | Strand | Frame |
|---|---|---|---|---|---|
| 14.4 bits(7) | 223() | 7/7(100%) | 0/7(0%) | Plus/Plus | |

Features:

```
Query  281       TGGTGTT  287
                 |||||||
Sbjct  32385876  TGGTGTT  32385882
```

Range 211: 32385879 to 32385885

| Score | Expect | Identities | Gaps | Strand | Frame |
|---|---|---|---|---|---|
| 14.4 bits(7) | 223() | 7/7(100%) | 0/7(0%) | Plus/Minus | |

Features:

```
Query  271       TGTAACA  277
                 |||||||
Sbjct  32385885  TGTAACA  32385879
```

Range 212: 32386293 to 32386299

| Score | Expect | Identities | Gaps | Strand | Frame |
|---|---|---|---|---|---|
| 14.4 bits(7) | 223() | 7/7(100%) | 0/7(0%) | Plus/Minus | |

Features:

```
Query  293       CTTTATT  299
                 |||||||
Sbjct  32386299  CTTTATT  32386293
```

Range 213: 32387010 to 32387016

| Score | Expect | Identities | Gaps | Strand | Frame |
|---|---|---|---|---|---|
| 14.4 bits(7) | 223() | 7/7(100%) | 0/7(0%) | Plus/Plus | |

Features:

```
Query  293       CTTTATT  299
                 |||||||
Sbjct  32387010  CTTTATT  32387016
```

Range 214: 32387581 to 32387587

| Score | Expect | Identities | Gaps | Strand | Frame |
|---|---|---|---|---|---|
| 14.4 bits(7) | 223() | 7/7(100%) | 0/7(0%) | Plus/Plus | |

Features:

```
Query  277       AAGCTGG  283
                 |||||||
Sbjct  32387581  AAGCTGG  32387587
```

Range 215: 32388661 to 32388667

| Score | Expect | Identities | Gaps | Strand | Frame |
|---|---|---|---|---|---|
| 14.4 bits(7) | 223() | 7/7(100%) | 0/7(0%) | Plus/Plus | |

Features:

```
Query  283       GTGTTCT  289
                 |||||||
Sbjct  32388661  GTGTTCT  32388667
```

Range 216: 32388666 to 32388672

| Score | Expect | Identities | Gaps | Strand | Frame |
|---|---|---|---|---|---|
| 14.4 bits(7) | 223() | 7/7(100%) | 0/7(0%) | Plus/Plus | |

Features:

```
Query  293       CTTTATT  299
                 |||||||
Sbjct  32388666  CTTTATT  32388672
```

Range 217: 32391162 to 32391168

| Score | Expect | Identities | Gaps | Strand | Frame |
|---|---|---|---|---|---|
| 14.4 bits(7) | 223() | 7/7(100%) | 0/7(0%) | Plus/Plus | |

Features:

```
Query  276       CAAGCTG  282
                 |||||||
Sbjct  32391162  CAAGCTG  32391168
```

Range 218: 32391270 to 32391276

| Score | Expect | Identities | Gaps | Strand | Frame |
|---|---|---|---|---|---|
| 14.4 bits(7) | 223() | 7/7(100%) | 0/7(0%) | Plus/Plus | |

Features:

```
Query  281       TGGTGTT  287
                 |||||||
Sbjct  32391270  TGGTGTT  32391276
```

Range 219: 32391716 to 32391722

| Score | Expect | Identities | Gaps | Strand | Frame |
|---|---|---|---|---|---|
| 14.4 bits(7) | 223() | 7/7(100%) | 0/7(0%) | Plus/Plus | |

Features:

```
Query  291       TCCTTTA  297
                 |||||||
Sbjct  32391716  TCCTTTA  32391722
```

Range 220: 32391844 to 32391850

| Score | Expect | Identities | Gaps | Strand | Frame |
|---|---|---|---|---|---|

| | | | | |
|---|---|---|---|---|
| 14.4 bits(7) | 223() | 7/7(100%) | 0/7(0%) | Plus/Plus |

Features:

```
Query  289       TCTCCTT  295
                 |||||||
Sbjct  32391844  TCTCCTT  32391850
```

Range 221: 32392280 to 32392286

| Score | Expect | Identities | Gaps | Strand | Frame |
|---|---|---|---|---|---|
| 14.4 bits(7) | 223() | 7/7(100%) | 0/7(0%) | Plus/Minus | |

Features:

```
Query  274       AACAAGC  280
                 |||||||
Sbjct  32392286  AACAAGC  32392280
```

Range 222: 32393272 to 32393278

| Score | Expect | Identities | Gaps | Strand | Frame |
|---|---|---|---|---|---|
| 14.4 bits(7) | 223() | 7/7(100%) | 0/7(0%) | Plus/Minus | |

Features:

```
Query  288       CTCTCCT  294
                 |||||||
Sbjct  32393278  CTCTCCT  32393272
```

Range 223: 32393489 to 32393495

| Score | Expect | Identities | Gaps | Strand | Frame |
|---|---|---|---|---|---|
| 14.4 bits(7) | 223() | 7/7(100%) | 0/7(0%) | Plus/Minus | |

Features:

```
Query  293       CTTTATT  299
                 |||||||
Sbjct  32393495  CTTTATT  32393489
```

Range 224: 32393509 to 32393515

| Score | Expect | Identities | Gaps | Strand | Frame |
|---|---|---|---|---|---|
| 14.4 bits(7) | 223() | 7/7(100%) | 0/7(0%) | Plus/Plus | |

Features:

```
Query  293       CTTTATT  299
                 |||||||
Sbjct  32393509  CTTTATT  32393515
```

Range 225: 32394001 to 32394007

| Score | Expect | Identities | Gaps | Strand | Frame |
|---|---|---|---|---|---|
| 14.4 bits(7) | 223() | 7/7(100%) | 0/7(0%) | Plus/Minus | |

Features:

```
Query  271       TGTAACA  277
                 |||||||
Sbjct  32394007  TGTAACA  32394001
```

Range 226: 32394552 to 32394558

| Score | Expect | Identities | Gaps | Strand | Frame |
|---|---|---|---|---|---|
| 14.4 bits(7) | 223() | 7/7(100%) | 0/7(0%) | Plus/Plus | |

Features:

```
Query  279       GCTGGTG  285
                 |||||||
Sbjct  32394552  GCTGGTG  32394558
```

Range 227: 32394650 to 32394656

| Score | Expect | Identities | Gaps | Strand | Frame |
|---|---|---|---|---|---|
| 14.4 bits(7) | 223() | 7/7(100%) | 0/7(0%) | Plus/Minus | |

Features:

```
Query  289       TCTCCTT  295
                 |||||||
Sbjct  32394656  TCTCCTT  32394650
```

Range 228: 32395932 to 32395938

| Score | Expect | Identities | Gaps | Strand | Frame |
|---|---|---|---|---|---|
| 14.4 bits(7) | 223() | 7/7(100%) | 0/7(0%) | Plus/Minus | |

Features:

```
Query  287       TCTCTCC  293
                 |||||||
Sbjct  32395938  TCTCTCC  32395932
```

Range 229: 32397095 to 32397101

| Score | Expect | Identities | Gaps | Strand | Frame |
|---|---|---|---|---|---|
| 14.4 bits(7) | 223() | 7/7(100%) | 0/7(0%) | Plus/Plus | |

Features:

```
Query  286       TTCTCTC  292
                 |||||||
Sbjct  32397095  TTCTCTC  32397101
```

Range 230: 32397100 to 32397106

| Score | Expect | Identities | Gaps | Strand | Frame |
|---|---|---|---|---|---|
| 14.4 bits(7) | 223() | 7/7(100%) | 0/7(0%) | Plus/Plus | |

Features:

```
Query  287       TCTCTCC  293
                 |||||||
Sbjct  32397100  TCTCTCC  32397106
```

Range 231: 32397929 to 32397935

| Score | Expect | Identities | Gaps | Strand | Frame |
|---|---|---|---|---|---|
| 14.4 bits(7) | 223() | 7/7(100%) | 0/7(0%) | Plus/Minus | |

Features:

```
Query  279       GCTGGTG  285
                 |||||||
Sbjct  32397935  GCTGGTG  32397929
```

Range 232: 32398713 to 32398719

| Score | Expect | Identities | Gaps | Strand | Frame |
|---|---|---|---|---|---|
| 14.4 bits(7) | 223() | 7/7(100%) | 0/7(0%) | Plus/Plus | |

Features:

```
Query  293       CTTTATT  299
                 |||||||
Sbjct  32398713  CTTTATT  32398719
```

Range 233: 32399873 to 32399879

| Score | Expect | Identities | Gaps | Strand | Frame |
|---|---|---|---|---|---|
| 14.4 bits(7) | 223() | 7/7(100%) | 0/7(0%) | Plus/Minus | |

Features:

```
Query  282       GGTGTTC  288
                 |||||||
Sbjct  32399879  GGTGTTC  32399873
```

Range 234: 32400114 to 32400120

| Score | Expect | Identities | Gaps | Strand | Frame |
|---|---|---|---|---|---|
| 14.4 bits(7) | 223() | 7/7(100%) | 0/7(0%) | Plus/Minus | |

Features:

```
Query  273       TAACAAG  279
                 |||||||
Sbjct  32400120  TAACAAG  32400114
```

Range 235: 32400282 to 32400288

| Score | Expect | Identities | Gaps | Strand | Frame |
|---|---|---|---|---|---|
| 14.4 bits(7) | 223() | 7/7(100%) | 0/7(0%) | Plus/Plus | |

Features:

```
Query  283       GTGTTCT  289
                 |||||||
Sbjct  32400282  GTGTTCT  32400288
```

Range 236: 32400933 to 32400939

| Score | Expect | Identities | Gaps | Strand | Frame |
|---|---|---|---|---|---|
| 14.4 bits(7) | 223() | 7/7(100%) | 0/7(0%) | Plus/Plus | |

Features:

```
Query  272       GTAACAA  278
                 |||||||
Sbjct  32400933  GTAACAA  32400939
```

Range 237: 32401121 to 32401127

| Score | Expect | Identities | Gaps | Strand | Frame |
|---|---|---|---|---|---|
| 14.4 bits(7) | 223() | 7/7(100%) | 0/7(0%) | Plus/Plus | |

Features:

```
Query  293       CTTTATT  299
                 |||||||
Sbjct  32401121  CTTTATT  32401127
```

Range 238: 32401404 to 32401410

| Score | Expect | Identities | Gaps | Strand | Frame |
|---|---|---|---|---|---|
| 14.4 bits(7) | 223() | 7/7(100%) | 0/7(0%) | Plus/Plus | |

Features:

```
Query  272       GTAACAA  278
                 |||||||
Sbjct  32401404  GTAACAA  32401410
```

Range 239: 32401552 to 32401558

| Score | Expect | Identities | Gaps | Strand | Frame |
|---|---|---|---|---|---|

| Score | Expect | Identities | Gaps | Strand | Frame |
|---|---|---|---|---|---|
| 14.4 bits(7) | 223() | 7/7(100%) | 0/7(0%) | Plus/Minus | |

Features:

```
Query  289       TCTCCTT  295
                 |||||||
Sbjct  32401558  TCTCCTT  32401552
```

Range 240: 32401614 to 32401620

| Score | Expect | Identities | Gaps | Strand | Frame |
|---|---|---|---|---|---|
| 14.4 bits(7) | 223() | 7/7(100%) | 0/7(0%) | Plus/Minus | |

Features:

```
Query  289       TCTCCTT  295
                 |||||||
Sbjct  32401620  TCTCCTT  32401614
```

Range 241: 32401657 to 32401663

| Score | Expect | Identities | Gaps | Strand | Frame |
|---|---|---|---|---|---|
| 14.4 bits(7) | 223() | 7/7(100%) | 0/7(0%) | Plus/Minus | |

Features:

```
Query  271       TGTAACA  277
                 |||||||
Sbjct  32401663  TGTAACA  32401657
```

Range 242: 32402170 to 32402176

| Score | Expect | Identities | Gaps | Strand | Frame |
|---|---|---|---|---|---|
| 14.4 bits(7) | 223() | 7/7(100%) | 0/7(0%) | Plus/Plus | |

Features:

```
Query  289       TCTCCTT  295
                 |||||||
Sbjct  32402170  TCTCCTT  32402176
```

Range 243: 32403639 to 32403645

| Score | Expect | Identities | Gaps | Strand | Frame |
|---|---|---|---|---|---|
| 14.4 bits(7) | 223() | 7/7(100%) | 0/7(0%) | Plus/Minus | |

Features:

```
Query  271       TGTAACA  277
                 |||||||
Sbjct  32403645  TGTAACA  32403639
```

Range 244: 32404143 to 32404149

| Score | Expect | Identities | Gaps | Strand | Frame |
|---|---|---|---|---|---|
| 14.4 bits(7) | 223() | 7/7(100%) | 0/7(0%) | Plus/Plus | |

Features:

```
Query  284       TGTTCTC  290
                 |||||||
Sbjct  32404143  TGTTCTC  32404149
```

Range 245: 32405059 to 32405065

| Score | Expect | Identities | Gaps | Strand | Frame |
|---|---|---|---|---|---|
| 14.4 bits(7) | 223() | 7/7(100%) | 0/7(0%) | Plus/Plus | |

Features:

```
Query  291       TCCTTTA  297
                 |||||||
Sbjct  32405059  TCCTTTA  32405065
```

Range 246: 32406882 to 32406888

| Score | Expect | Identities | Gaps | Strand | Frame |
|---|---|---|---|---|---|
| 14.4 bits(7) | 223() | 7/7(100%) | 0/7(0%) | Plus/Plus | |

Features:

```
Query  293       CTTTATT  299
                 |||||||
Sbjct  32406882  CTTTATT  32406888
```

Range 247: 32406899 to 32406905

| Score | Expect | Identities | Gaps | Strand | Frame |
|---|---|---|---|---|---|
| 14.4 bits(7) | 223() | 7/7(100%) | 0/7(0%) | Plus/Minus | |

Features:

```
Query  293       CTTTATT  299
                 |||||||
Sbjct  32406905  CTTTATT  32406899
```

Range 248: 32407429 to 32407435

| Score | Expect | Identities | Gaps | Strand | Frame |
|---|---|---|---|---|---|
| 14.4 bits(7) | 223() | 7/7(100%) | 0/7(0%) | Plus/Minus | |

Features:

```
Query  286       TTCTCTC  292
                 |||||||
Sbjct  32407435  TTCTCTC  32407429
```

Range 249: 32408446 to 32408452

| Score | Expect | Identities | Gaps | Strand | Frame |
|---|---|---|---|---|---|
| 14.4 bits(7) | 223() | 7/7(100%) | 0/7(0%) | Plus/Minus | |

Features:

```
Query  273       TAACAAG  279
                 |||||||
Sbjct  32408452  TAACAAG  32408446
```

Range 250: 32409790 to 32409796

| Score | Expect | Identities | Gaps | Strand | Frame |
|---|---|---|---|---|---|
| 14.4 bits(7) | 223() | 7/7(100%) | 0/7(0%) | Plus/Plus | |

Features:

```
Query  279       GCTGGTG  285
                 |||||||
Sbjct  32409790  GCTGGTG  32409796
```

Range 251: 32409796 to 32409802

| Score | Expect | Identities | Gaps | Strand | Frame |
|---|---|---|---|---|---|
| 14.4 bits(7) | 223() | 7/7(100%) | 0/7(0%) | Plus/Minus | |

Features:

```
Query  274       AACAAGC  280
                 |||||||
Sbjct  32409802  AACAAGC  32409796
```

Range 252: 32410018 to 32410024

| Score | Expect | Identities | Gaps | Strand | Frame |
|---|---|---|---|---|---|
| 14.4 bits(7) | 223() | 7/7(100%) | 0/7(0%) | Plus/Minus | |

Features:

```
Query  293       CTTTATT  299
                 |||||||
Sbjct  32410024  CTTTATT  32410018
```

Range 253: 32411168 to 32411174

| Score | Expect | Identities | Gaps | Strand | Frame |
|---|---|---|---|---|---|
| 14.4 bits(7) | 223() | 7/7(100%) | 0/7(0%) | Plus/Plus | |

Features:

```
Query  278       AGCTGGT  284
                 |||||||
Sbjct  32411168  AGCTGGT  32411174
```

Range 254: 32411691 to 32411697

| Score | Expect | Identities | Gaps | Strand | Frame |
|---|---|---|---|---|---|
| 14.4 bits(7) | 223() | 7/7(100%) | 0/7(0%) | Plus/Plus | |

Features:

```
Query  284       TGTTCTC  290
                 |||||||
Sbjct  32411691  TGTTCTC  32411697
```

Range 255: 32412575 to 32412581

| Score | Expect | Identities | Gaps | Strand | Frame |
|---|---|---|---|---|---|
| 14.4 bits(7) | 223() | 7/7(100%) | 0/7(0%) | Plus/Minus | |

Features:

```
Query  293       CTTTATT  299
                 |||||||
Sbjct  32412581  CTTTATT  32412575
```

Range 256: 32412732 to 32412738

| Score | Expect | Identities | Gaps | Strand | Frame |
|---|---|---|---|---|---|
| 14.4 bits(7) | 223() | 7/7(100%) | 0/7(0%) | Plus/Minus | |

Features:

```
Query  286       TTCTCTC  292
                 |||||||
Sbjct  32412738  TTCTCTC  32412732
```

Range 257: 32412849 to 32412855

| Score | Expect | Identities | Gaps | Strand | Frame |
|---|---|---|---|---|---|
| 14.4 bits(7) | 223() | 7/7(100%) | 0/7(0%) | Plus/Plus | |

Features:

```
Query  286       TTCTCTC  292
                 |||||||
Sbjct  32412849  TTCTCTC  32412855
```

Range 258: 32414539 to 32414545

| Score | Expect | Identities | Gaps | Strand | Frame |
|---|---|---|---|---|---|

| Score | Expect | Identities | Gaps | Strand | Frame |
|---|---|---|---|---|---|
| 14.4 bits(7) | 223() | 7/7(100%) | 0/7(0%) | Plus/Minus | |

Features:

```
Query  275       ACAAGCT  281
                 |||||||
Sbjct  32414545  ACAAGCT  32414539
```

Range 259: 32415699 to 32415705

| Score | Expect | Identities | Gaps | Strand | Frame |
|---|---|---|---|---|---|
| 14.4 bits(7) | 223() | 7/7(100%) | 0/7(0%) | Plus/Minus | |

Features:

```
Query  279       GCTGGTG  285
                 |||||||
Sbjct  32415705  GCTGGTG  32415699
```

Range 260: 32415961 to 32415967

| Score | Expect | Identities | Gaps | Strand | Frame |
|---|---|---|---|---|---|
| 14.4 bits(7) | 223() | 7/7(100%) | 0/7(0%) | Plus/Minus | |

Features:

```
Query  286       TTCTCTC  292
                 |||||||
Sbjct  32415967  TTCTCTC  32415961
```

Range 261: 32416081 to 32416087

| Score | Expect | Identities | Gaps | Strand | Frame |
|---|---|---|---|---|---|
| 14.4 bits(7) | 223() | 7/7(100%) | 0/7(0%) | Plus/Minus | |

Features:

```
Query  290       CTCCTTT  296
                 |||||||
Sbjct  32416087  CTCCTTT  32416081
```

Range 262: 32416830 to 32416836

| Score | Expect | Identities | Gaps | Strand | Frame |
|---|---|---|---|---|---|
| 14.4 bits(7) | 223() | 7/7(100%) | 0/7(0%) | Plus/Plus | |

Features:

```
Query  290       CTCCTTT  296
                 |||||||
Sbjct  32416830  CTCCTTT  32416836
```

Range 263: 32416879 to 32416885

| Score | Expect | Identities | Gaps | Strand | Frame |
|---|---|---|---|---|---|
| 14.4 bits(7) | 223() | 7/7(100%) | 0/7(0%) | Plus/Plus | |

Features:

```
Query  276       CAAGCTG  282
                 |||||||
Sbjct  32416879  CAAGCTG  32416885
```

Range 264: 32419637 to 32419643

| Score | Expect | Identities | Gaps | Strand | Frame |
|---|---|---|---|---|---|
| 14.4 bits(7) | 223() | 7/7(100%) | 0/7(0%) | Plus/Minus | |

Features:

```
Query  293       CTTTATT  299
                 |||||||
Sbjct  32419643  CTTTATT  32419637
```

Range 265: 32419871 to 32419877

| Score | Expect | Identities | Gaps | Strand | Frame |
|---|---|---|---|---|---|
| 14.4 bits(7) | 223() | 7/7(100%) | 0/7(0%) | Plus/Plus | |

Features:

```
Query  291       TCCTTTA  297
                 |||||||
Sbjct  32419871  TCCTTTA  32419877
```

Range 266: 32422343 to 32422349

| Score | Expect | Identities | Gaps | Strand | Frame |
|---|---|---|---|---|---|
| 14.4 bits(7) | 223() | 7/7(100%) | 0/7(0%) | Plus/Minus | |

Features:

```
Query  272       GTAACAA  278
                 |||||||
Sbjct  32422349  GTAACAA  32422343
```

Range 267: 32422579 to 32422585

| Score | Expect | Identities | Gaps | Strand | Frame |
|---|---|---|---|---|---|
| 14.4 bits(7) | 223() | 7/7(100%) | 0/7(0%) | Plus/Minus | |

Features:

```
Query  272       GTAACAA  278
                 |||||||
Sbjct  32422585  GTAACAA  32422579
```

Range 268: 32422671 to 32422677

| Score | Expect | Identities | Gaps | Strand | Frame |
|---|---|---|---|---|---|
| 14.4 bits(7) | 223() | 7/7(100%) | 0/7(0%) | Plus/Minus | |

Features:

```
Query  274       AACAAGC  280
                 |||||||
Sbjct  32422677  AACAAGC  32422671
```

Range 269: 32422683 to 32422689

| Score | Expect | Identities | Gaps | Strand | Frame |
|---|---|---|---|---|---|
| 14.4 bits(7) | 223() | 7/7(100%) | 0/7(0%) | Plus/Plus | |

Features:

```
Query  292       CCTTTAT  298
                 |||||||
Sbjct  32422683  CCTTTAT  32422689
```

Range 270: 32424547 to 32424553

| Score | Expect | Identities | Gaps | Strand | Frame |
|---|---|---|---|---|---|
| 14.4 bits(7) | 223() | 7/7(100%) | 0/7(0%) | Plus/Minus | |

Features:

```
Query  274       AACAAGC  280
                 |||||||
Sbjct  32424553  AACAAGC  32424547
```

Range 271: 32425555 to 32425561

| Score | Expect | Identities | Gaps | Strand | Frame |
|---|---|---|---|---|---|
| 14.4 bits(7) | 223() | 7/7(100%) | 0/7(0%) | Plus/Plus | |

Features:

```
Query  286       TTCTCTC  292
                 |||||||
Sbjct  32425555  TTCTCTC  32425561
```

Range 272: 32425577 to 32425583

| Score | Expect | Identities | Gaps | Strand | Frame |
|---|---|---|---|---|---|
| 14.4 bits(7) | 223() | 7/7(100%) | 0/7(0%) | Plus/Minus | |

Features:

```
Query  274       AACAAGC  280
                 |||||||
Sbjct  32425583  AACAAGC  32425577
```

Range 273: 32425636 to 32425642

| Score | Expect | Identities | Gaps | Strand | Frame |
|---|---|---|---|---|---|
| 14.4 bits(7) | 223() | 7/7(100%) | 0/7(0%) | Plus/Minus | |

Features:

```
Query  285       GTTCTCT  291
                 |||||||
Sbjct  32425642  GTTCTCT  32425636
```

Range 274: 32425881 to 32425887

| Score | Expect | Identities | Gaps | Strand | Frame |
|---|---|---|---|---|---|
| 14.4 bits(7) | 223() | 7/7(100%) | 0/7(0%) | Plus/Minus | |

Features:

```
Query  279       GCTGGTG  285
                 |||||||
Sbjct  32425887  GCTGGTG  32425881
```

Range 275: 32425952 to 32425958

| Score | Expect | Identities | Gaps | Strand | Frame |
|---|---|---|---|---|---|
| 14.4 bits(7) | 223() | 7/7(100%) | 0/7(0%) | Plus/Plus | |

Features:

```
Query  278       AGCTGGT  284
                 |||||||
Sbjct  32425952  AGCTGGT  32425958
```

Range 276: 32426232 to 32426238

| Score | Expect | Identities | Gaps | Strand | Frame |
|---|---|---|---|---|---|
| 14.4 bits(7) | 223() | 7/7(100%) | 0/7(0%) | Plus/Plus | |

Features:

```
Query  293       CTTTATT  299
                 |||||||
Sbjct  32426232  CTTTATT  32426238
```

Range 277: 32426269 to 32426275

| Score | Expect | Identities | Gaps | Strand | Frame |
|---|---|---|---|---|---|

| Score | Expect | Identities | Gaps | Strand | Frame |
|---|---|---|---|---|---|
| 14.4 bits(7) | 223() | 7/7(100%) | 0/7(0%) | Plus/Plus | |

Features:

```
Query  287       TCTCTCC  293
                 |||||||
Sbjct  32426269  TCTCTCC  32426275
```

Range 278: 32426636 to 32426642

| Score | Expect | Identities | Gaps | Strand | Frame |
|---|---|---|---|---|---|
| 14.4 bits(7) | 223() | 7/7(100%) | 0/7(0%) | Plus/Minus | |

Features:

```
Query  271       TGTAACA  277
                 |||||||
Sbjct  32426642  TGTAACA  32426636
```

Range 279: 32426735 to 32426741

| Score | Expect | Identities | Gaps | Strand | Frame |
|---|---|---|---|---|---|
| 14.4 bits(7) | 223() | 7/7(100%) | 0/7(0%) | Plus/Plus | |

Features:

```
Query  293       CTTTATT  299
                 |||||||
Sbjct  32426735  CTTTATT  32426741
```

Range 280: 32426802 to 32426808

| Score | Expect | Identities | Gaps | Strand | Frame |
|---|---|---|---|---|---|
| 14.4 bits(7) | 223() | 7/7(100%) | 0/7(0%) | Plus/Minus | |

Features:

```
Query  272       GTAACAA  278
                 |||||||
Sbjct  32426808  GTAACAA  32426802
```

Range 281: 32427389 to 32427395

| Score | Expect | Identities | Gaps | Strand | Frame |
|---|---|---|---|---|---|
| 14.4 bits(7) | 223() | 7/7(100%) | 0/7(0%) | Plus/Plus | |

Features:

```
Query  289       TCTCCTT  295
                 |||||||
Sbjct  32427389  TCTCCTT  32427395
```

Range 282: 32428292 to 32428298

| Score | Expect | Identities | Gaps | Strand | Frame |
|---|---|---|---|---|---|
| 14.4 bits(7) | 223() | 7/7(100%) | 0/7(0%) | Plus/Plus | |

Features:

```
Query  286       TTCTCTC  292
                 |||||||
Sbjct  32428292  TTCTCTC  32428298
```

Range 283: 32428381 to 32428387

| Score | Expect | Identities | Gaps | Strand | Frame |
|---|---|---|---|---|---|
| 14.4 bits(7) | 223() | 7/7(100%) | 0/7(0%) | Plus/Plus | |

Features:

```
Query  289       TCTCCTT  295
                 |||||||
Sbjct  32428381  TCTCCTT  32428387
```

Range 284: 32429228 to 32429234

| Score | Expect | Identities | Gaps | Strand | Frame |
|---|---|---|---|---|---|
| 14.4 bits(7) | 223() | 7/7(100%) | 0/7(0%) | Plus/Minus | |

Features:

```
Query  291       TCCTTTA  297
                 |||||||
Sbjct  32429234  TCCTTTA  32429228
```

Range 285: 32429262 to 32429268

| Score | Expect | Identities | Gaps | Strand | Frame |
|---|---|---|---|---|---|
| 14.4 bits(7) | 223() | 7/7(100%) | 0/7(0%) | Plus/Plus | |

Features:

```
Query  292       CCTTTAT  298
                 |||||||
Sbjct  32429262  CCTTTAT  32429268
```

Range 286: 32430726 to 32430732

| Score | Expect | Identities | Gaps | Strand | Frame |
|---|---|---|---|---|---|
| 14.4 bits(7) | 223() | 7/7(100%) | 0/7(0%) | Plus/Plus | |

Features:

```
Query  293       CTTTATT  299
                 |||||||
Sbjct  32430726  CTTTATT  32430732
```

Range 287: 32431006 to 32431012

| Score | Expect | Identities | Gaps | Strand | Frame |
|---|---|---|---|---|---|
| 14.4 bits(7) | 223() | 7/7(100%) | 0/7(0%) | Plus/Minus | |

Features:

```
Query  290       CTCCTTT  296
                 |||||||
Sbjct  32431012  CTCCTTT  32431006
```

Range 288: 32431036 to 32431042

| Score | Expect | Identities | Gaps | Strand | Frame |
|---|---|---|---|---|---|
| 14.4 bits(7) | 223() | 7/7(100%) | 0/7(0%) | Plus/Minus | |

Features:

```
Query  281       TGGTGTT  287
                 |||||||
Sbjct  32431042  TGGTGTT  32431036
```

Range 289: 32431737 to 32431743

| Score | Expect | Identities | Gaps | Strand | Frame |
|---|---|---|---|---|---|
| 14.4 bits(7) | 223() | 7/7(100%) | 0/7(0%) | Plus/Minus | |

Features:

```
Query  274       AACAAGC  280
                 |||||||
Sbjct  32431743  AACAAGC  32431737
```

Range 290: 32431940 to 32431946

| Score | Expect | Identities | Gaps | Strand | Frame |
|---|---|---|---|---|---|
| 14.4 bits(7) | 223() | 7/7(100%) | 0/7(0%) | Plus/Minus | |

Features:

```
Query  286       TTCTCTC  292
                 |||||||
Sbjct  32431946  TTCTCTC  32431940
```

Range 291: 32432464 to 32432470

| Score | Expect | Identities | Gaps | Strand | Frame |
|---|---|---|---|---|---|
| 14.4 bits(7) | 223() | 7/7(100%) | 0/7(0%) | Plus/Plus | |

Features:

```
Query  292       CCTTTAT  298
                 |||||||
Sbjct  32432464  CCTTTAT  32432470
```

Range 292: 32432548 to 32432554

| Score | Expect | Identities | Gaps | Strand | Frame |
|---|---|---|---|---|---|
| 14.4 bits(7) | 223() | 7/7(100%) | 0/7(0%) | Plus/Plus | |

Features:

```
Query  282       GGTGTTC  288
                 |||||||
Sbjct  32432548  GGTGTTC  32432554
```

Range 293: 32433502 to 32433508

| Score | Expect | Identities | Gaps | Strand | Frame |
|---|---|---|---|---|---|
| 14.4 bits(7) | 223() | 7/7(100%) | 0/7(0%) | Plus/Plus | |

Features:

```
Query  286       TTCTCTC  292
                 |||||||
Sbjct  32433502  TTCTCTC  32433508
```

Range 294: 32433612 to 32433618

| Score | Expect | Identities | Gaps | Strand | Frame |
|---|---|---|---|---|---|
| 14.4 bits(7) | 223() | 7/7(100%) | 0/7(0%) | Plus/Plus | |

Features:

```
Query  293       CTTTATT  299
                 |||||||
Sbjct  32433612  CTTTATT  32433618
```

Range 295: 32433686 to 32433692

| Score | Expect | Identities | Gaps | Strand | Frame |
|---|---|---|---|---|---|
| 14.4 bits(7) | 223() | 7/7(100%) | 0/7(0%) | Plus/Minus | |

Features:

```
Query  275       ACAAGCT  281
                 |||||||
Sbjct  32433692  ACAAGCT  32433686
```

Range 296: 32433774 to 32433780

| Score | Expect | Identities | Gaps | Strand | Frame |
|---|---|---|---|---|---|

| Score | Expect | Identities | Gaps | Strand | Frame |
|---|---|---|---|---|---|
| 14.4 bits(7) | 223() | 7/7(100%) | 0/7(0%) | Plus/Minus | |

Features:

```
Query  293       CTTTATT  299
                 |||||||
Sbjct  32433780  CTTTATT  32433774
```

Range 297: 32435622 to 32435628

| Score | Expect | Identities | Gaps | Strand | Frame |
|---|---|---|---|---|---|
| 14.4 bits(7) | 223() | 7/7(100%) | 0/7(0%) | Plus/Minus | |

Features:

```
Query  290       CTCCTTT  296
                 |||||||
Sbjct  32435628  CTCCTTT  32435622
```

Range 298: 32435691 to 32435697

| Score | Expect | Identities | Gaps | Strand | Frame |
|---|---|---|---|---|---|
| 14.4 bits(7) | 223() | 7/7(100%) | 0/7(0%) | Plus/Minus | |

Features:

```
Query  283       GTGTTCT  289
                 |||||||
Sbjct  32435697  GTGTTCT  32435691
```

Range 299: 32436175 to 32436181

| Score | Expect | Identities | Gaps | Strand | Frame |
|---|---|---|---|---|---|
| 14.4 bits(7) | 223() | 7/7(100%) | 0/7(0%) | Plus/Minus | |

Features:

```
Query  273       TAACAAG  279
                 |||||||
Sbjct  32436181  TAACAAG  32436175
```

Range 300: 32436957 to 32436963

| Score | Expect | Identities | Gaps | Strand | Frame |
|---|---|---|---|---|---|
| 14.4 bits(7) | 223() | 7/7(100%) | 0/7(0%) | Plus/Minus | |

Features:

```
Query  276       CAAGCTG  282
                 |||||||
Sbjct  32436963  CAAGCTG  32436957
```

Range 301: 32437947 to 32437953

| Score | Expect | Identities | Gaps | Strand | Frame |
|---|---|---|---|---|---|
| 14.4 bits(7) | 223() | 7/7(100%) | 0/7(0%) | Plus/Plus | |

Features:

```
Query  281       TGGTGTT  287
                 |||||||
Sbjct  32437947  TGGTGTT  32437953
```

Range 302: 32438335 to 32438341

| Score | Expect | Identities | Gaps | Strand | Frame |
|---|---|---|---|---|---|
| 14.4 bits(7) | 223() | 7/7(100%) | 0/7(0%) | Plus/Plus | |

Features:

```
Query  271       TGTAACA  277
                 |||||||
Sbjct  32438335  TGTAACA  32438341
```

Range 303: 32438553 to 32438559

| Score | Expect | Identities | Gaps | Strand | Frame |
|---|---|---|---|---|---|
| 14.4 bits(7) | 223() | 7/7(100%) | 0/7(0%) | Plus/Plus | |

Features:

```
Query  283       GTGTTCT  289
                 |||||||
Sbjct  32438553  GTGTTCT  32438559
```

Range 304: 32439863 to 32439869

| Score | Expect | Identities | Gaps | Strand | Frame |
|---|---|---|---|---|---|
| 14.4 bits(7) | 223() | 7/7(100%) | 0/7(0%) | Plus/Plus | |

Features:

```
Query  293       CTTTATT  299
                 |||||||
Sbjct  32439863  CTTTATT  32439869
```

Range 305: 32440252 to 32440258

| Score | Expect | Identities | Gaps | Strand | Frame |
|---|---|---|---|---|---|
| 14.4 bits(7) | 223() | 7/7(100%) | 0/7(0%) | Plus/Plus | |

Features:

```
Query  292       CCTTTAT  298
                 |||||||
Sbjct  32440252  CCTTTAT  32440258
```

Range 306: 32440974 to 32440980

| Score | Expect | Identities | Gaps | Strand | Frame |
|---|---|---|---|---|---|
| 14.4 bits(7) | 223() | 7/7(100%) | 0/7(0%) | Plus/Plus | |

Features:

```
Query  284       TGTTCTC  290
                 |||||||
Sbjct  32440974  TGTTCTC  32440980
```

Range 307: 32441192 to 32441198

| Score | Expect | Identities | Gaps | Strand | Frame |
|---|---|---|---|---|---|
| 14.4 bits(7) | 223() | 7/7(100%) | 0/7(0%) | Plus/Plus | |

Features:

```
Query  275       ACAAGCT  281
                 |||||||
Sbjct  32441192  ACAAGCT  32441198
```

Range 308: 32441481 to 32441487

| Score | Expect | Identities | Gaps | Strand | Frame |
|---|---|---|---|---|---|
| 14.4 bits(7) | 223() | 7/7(100%) | 0/7(0%) | Plus/Minus | |

Features:

```
Query  273       TAACAAG  279
                 |||||||
Sbjct  32441487  TAACAAG  32441481
```

Range 309: 32441516 to 32441522

| Score | Expect | Identities | Gaps | Strand | Frame |
|---|---|---|---|---|---|
| 14.4 bits(7) | 223() | 7/7(100%) | 0/7(0%) | Plus/Minus | |

Features:

```
Query  281       TGGTGTT  287
                 |||||||
Sbjct  32441522  TGGTGTT  32441516
```

Range 310: 32441657 to 32441663

| Score | Expect | Identities | Gaps | Strand | Frame |
|---|---|---|---|---|---|
| 14.4 bits(7) | 223() | 7/7(100%) | 0/7(0%) | Plus/Minus | |

Features:

```
Query  274       AACAAGC  280
                 |||||||
Sbjct  32441663  AACAAGC  32441657
```

Range 311: 32441864 to 32441870

| Score | Expect | Identities | Gaps | Strand | Frame |
|---|---|---|---|---|---|
| 14.4 bits(7) | 223() | 7/7(100%) | 0/7(0%) | Plus/Minus | |

Features:

```
Query  271       TGTAACA  277
                 |||||||
Sbjct  32441870  TGTAACA  32441864
```

Range 312: 32442082 to 32442088

| Score | Expect | Identities | Gaps | Strand | Frame |
|---|---|---|---|---|---|
| 14.4 bits(7) | 223() | 7/7(100%) | 0/7(0%) | Plus/Minus | |

Features:

```
Query  271       TGTAACA  277
                 |||||||
Sbjct  32442088  TGTAACA  32442082
```

Range 313: 32442994 to 32443000

| Score | Expect | Identities | Gaps | Strand | Frame |
|---|---|---|---|---|---|
| 14.4 bits(7) | 223() | 7/7(100%) | 0/7(0%) | Plus/Minus | |

Features:

```
Query  293       CTTTATT  299
                 |||||||
Sbjct  32443000  CTTTATT  32442994
```

Range 314: 32443420 to 32443426

| Score | Expect | Identities | Gaps | Strand | Frame |
|---|---|---|---|---|---|
| 14.4 bits(7) | 223() | 7/7(100%) | 0/7(0%) | Plus/Minus | |

Features:

```
Query  280       CTGGTGT  286
                 |||||||
Sbjct  32443426  CTGGTGT  32443420
```

Range 315: 32443878 to 32443884

| Score | Expect | Identities | Gaps | Strand | Frame |
|---|---|---|---|---|---|

| | | | | |
|---|---|---|---|---|
| 14.4 bits(7) | 223() | 7/7(100%) | 0/7(0%) | Plus/Plus |

Features:

```
Query  271       TGTAACA  277
                 |||||||
Sbjct  32443878  TGTAACA  32443884
```

Range 316: 32444434 to 32444440

| Score | Expect | Identities | Gaps | Strand | Frame |
|---|---|---|---|---|---|
| 14.4 bits(7) | 223() | 7/7(100%) | 0/7(0%) | Plus/Plus | |

Features:

```
Query  285       GTTCTCT  291
                 |||||||
Sbjct  32444434  GTTCTCT  32444440
```

Range 317: 32444503 to 32444509

| Score | Expect | Identities | Gaps | Strand | Frame |
|---|---|---|---|---|---|
| 14.4 bits(7) | 223() | 7/7(100%) | 0/7(0%) | Plus/Plus | |

Features:

```
Query  281       TGGTGTT  287
                 |||||||
Sbjct  32444503  TGGTGTT  32444509
```

Range 318: 32444766 to 32444772

| Score | Expect | Identities | Gaps | Strand | Frame |
|---|---|---|---|---|---|
| 14.4 bits(7) | 223() | 7/7(100%) | 0/7(0%) | Plus/Minus | |

Features:

```
Query  274       AACAAGC  280
                 |||||||
Sbjct  32444772  AACAAGC  32444766
```

Range 319: 32445477 to 32445483

| Score | Expect | Identities | Gaps | Strand | Frame |
|---|---|---|---|---|---|
| 14.4 bits(7) | 223() | 7/7(100%) | 0/7(0%) | Plus/Minus | |

Features:

```
Query  290       CTCCTTT  296
                 |||||||
Sbjct  32445483  CTCCTTT  32445477
```

Range 320: 32445639 to 32445645

| Score | Expect | Identities | Gaps | Strand | Frame |
|---|---|---|---|---|---|
| 14.4 bits(7) | 223() | 7/7(100%) | 0/7(0%) | Plus/Minus | |

Features:

```
Query  281       TGGTGTT  287
                 |||||||
Sbjct  32445645  TGGTGTT  32445639
```

Range 321: 32445728 to 32445734

| Score | Expect | Identities | Gaps | Strand | Frame |
|---|---|---|---|---|---|
| 14.4 bits(7) | 223() | 7/7(100%) | 0/7(0%) | Plus/Minus | |

Features:

```
Query  285       GTTCTCT  291
                 |||||||
Sbjct  32445734  GTTCTCT  32445728
```

Range 322: 32446612 to 32446618

| Score | Expect | Identities | Gaps | Strand | Frame |
|---|---|---|---|---|---|
| 14.4 bits(7) | 223() | 7/7(100%) | 0/7(0%) | Plus/Minus | |

Features:

```
Query  277       AAGCTGG  283
                 |||||||
Sbjct  32446618  AAGCTGG  32446612
```

Range 323: 32448608 to 32448614

| Score | Expect | Identities | Gaps | Strand | Frame |
|---|---|---|---|---|---|
| 14.4 bits(7) | 223() | 7/7(100%) | 0/7(0%) | Plus/Minus | |

Features:

```
Query  277       AAGCTGG  283
                 |||||||
Sbjct  32448614  AAGCTGG  32448608
```

Range 324: 32448946 to 32448952

| Score | Expect | Identities | Gaps | Strand | Frame |
|---|---|---|---|---|---|
| 14.4 bits(7) | 223() | 7/7(100%) | 0/7(0%) | Plus/Minus | |

Features:

```
Query  291       TCCTTTA  297
                 |||||||
Sbjct  32448952  TCCTTTA  32448946
```

Range 325: 32449635 to 32449641

| Score | Expect | Identities | Gaps | Strand | Frame |
|---|---|---|---|---|---|
| 14.4 bits(7) | 223() | 7/7(100%) | 0/7(0%) | Plus/Minus | |

Features:

```
Query  286       TTCTCTC  292
                 |||||||
Sbjct  32449641  TTCTCTC  32449635
```

Range 326: 32450909 to 32450915

| Score | Expect | Identities | Gaps | Strand | Frame |
|---|---|---|---|---|---|
| 14.4 bits(7) | 223() | 7/7(100%) | 0/7(0%) | Plus/Plus | |

Features:

```
Query  271       TGTAACA  277
                 |||||||
Sbjct  32450909  TGTAACA  32450915
```

Range 327: 32451176 to 32451182

| Score | Expect | Identities | Gaps | Strand | Frame |
|---|---|---|---|---|---|
| 14.4 bits(7) | 223() | 7/7(100%) | 0/7(0%) | Plus/Plus | |

Features:

```
Query  289       TCTCCTT  295
                 |||||||
Sbjct  32451176  TCTCCTT  32451182
```

Range 328: 32451332 to 32451338

| Score | Expect | Identities | Gaps | Strand | Frame |
|---|---|---|---|---|---|
| 14.4 bits(7) | 223() | 7/7(100%) | 0/7(0%) | Plus/Plus | |

Features:

```
Query  283       GTGTTCT  289
                 |||||||
Sbjct  32451332  GTGTTCT  32451338
```

Range 329: 32451466 to 32451472

| Score | Expect | Identities | Gaps | Strand | Frame |
|---|---|---|---|---|---|
| 14.4 bits(7) | 223() | 7/7(100%) | 0/7(0%) | Plus/Plus | |

Features:

```
Query  281       TGGTGTT  287
                 |||||||
Sbjct  32451466  TGGTGTT  32451472
```

Range 330: 32451642 to 32451648

| Score | Expect | Identities | Gaps | Strand | Frame |
|---|---|---|---|---|---|
| 14.4 bits(7) | 223() | 7/7(100%) | 0/7(0%) | Plus/Minus | |

Features:

```
Query  274       AACAAGC  280
                 |||||||
Sbjct  32451648  AACAAGC  32451642
```

Range 331: 32451688 to 32451694

| Score | Expect | Identities | Gaps | Strand | Frame |
|---|---|---|---|---|---|
| 14.4 bits(7) | 223() | 7/7(100%) | 0/7(0%) | Plus/Minus | |

Features:

```
Query  274       AACAAGC  280
                 |||||||
Sbjct  32451694  AACAAGC  32451688
```

Range 332: 32452021 to 32452027

| Score | Expect | Identities | Gaps | Strand | Frame |
|---|---|---|---|---|---|
| 14.4 bits(7) | 223() | 7/7(100%) | 0/7(0%) | Plus/Plus | |

Features:

```
Query  271       TGTAACA  277
                 |||||||
Sbjct  32452021  TGTAACA  32452027
```

Range 333: 32452720 to 32452726

| Score | Expect | Identities | Gaps | Strand | Frame |
|---|---|---|---|---|---|
| 14.4 bits(7) | 223() | 7/7(100%) | 0/7(0%) | Plus/Plus | |

Features:

```
Query  291       TCCTTTA  297
                 |||||||
Sbjct  32452720  TCCTTTA  32452726
```

Range 334: 32453119 to 32453125

| Score | Expect | Identities | Gaps | Strand | Frame |
|---|---|---|---|---|---|

| | | | | |
|---|---|---|---|---|
| 14.4 bits(7) | 223() | 7/7(100%) | 0/7(0%) | Plus/Plus |

Features:

```
Query  276       CAAGCTG  282
                 |||||||
Sbjct  32453119  CAAGCTG  32453125
```

Range 335: 32453152 to 32453158

| Score | Expect | Identities | Gaps | Strand | Frame |
|---|---|---|---|---|---|
| 14.4 bits(7) | 223() | 7/7(100%) | 0/7(0%) | Plus/Plus | |

Features:

```
Query  283       GTGTTCT  289
                 |||||||
Sbjct  32453152  GTGTTCT  32453158
```

Range 336: 32453333 to 32453339

| Score | Expect | Identities | Gaps | Strand | Frame |
|---|---|---|---|---|---|
| 14.4 bits(7) | 223() | 7/7(100%) | 0/7(0%) | Plus/Minus | |

Features:

```
Query  273       TAACAAG  279
                 |||||||
Sbjct  32453339  TAACAAG  32453333
```

Range 337: 32453544 to 32453550

| Score | Expect | Identities | Gaps | Strand | Frame |
|---|---|---|---|---|---|
| 14.4 bits(7) | 223() | 7/7(100%) | 0/7(0%) | Plus/Minus | |

Features:

```
Query  271       TGTAACA  277
                 |||||||
Sbjct  32453550  TGTAACA  32453544
```

Range 338: 32453990 to 32453996

| Score | Expect | Identities | Gaps | Strand | Frame |
|---|---|---|---|---|---|
| 14.4 bits(7) | 223() | 7/7(100%) | 0/7(0%) | Plus/Minus | |

Features:

```
Query  293       CTTTATT  299
                 |||||||
Sbjct  32453996  CTTTATT  32453990
```

Range 339: 32454736 to 32454742

| Score | Expect | Identities | Gaps | Strand | Frame |
|---|---|---|---|---|---|
| 14.4 bits(7) | 223() | 7/7(100%) | 0/7(0%) | Plus/Plus | |

Features:

```
Query  278       AGCTGGT  284
                 |||||||
Sbjct  32454736  AGCTGGT  32454742
```

Range 340: 32454782 to 32454788

| Score | Expect | Identities | Gaps | Strand | Frame |
|---|---|---|---|---|---|
| 14.4 bits(7) | 223() | 7/7(100%) | 0/7(0%) | Plus/Plus | |

Features:

```
Query  293       CTTTATT  299
                 |||||||
Sbjct  32454782  CTTTATT  32454788
```

Range 341: 32455510 to 32455516

| Score | Expect | Identities | Gaps | Strand | Frame |
|---|---|---|---|---|---|
| 14.4 bits(7) | 223() | 7/7(100%) | 0/7(0%) | Plus/Minus | |

Features:

```
Query  277       AAGCTGG  283
                 |||||||
Sbjct  32455516  AAGCTGG  32455510
```

Range 342: 32455840 to 32455846

| Score | Expect | Identities | Gaps | Strand | Frame |
|---|---|---|---|---|---|
| 14.4 bits(7) | 223() | 7/7(100%) | 0/7(0%) | Plus/Plus | |

Features:

```
Query  293       CTTTATT  299
                 |||||||
Sbjct  32455840  CTTTATT  32455846
```

Range 343: 32456744 to 32456750

| Score | Expect | Identities | Gaps | Strand | Frame |
|---|---|---|---|---|---|
| 14.4 bits(7) | 223() | 7/7(100%) | 0/7(0%) | Plus/Plus | |

Features:

```
Query  283       GTGTTCT  289
                 |||||||
Sbjct  32456744  GTGTTCT  32456750
```

Range 344: 32456764 to 32456770

| Score | Expect | Identities | Gaps | Strand | Frame |
|---|---|---|---|---|---|
| 14.4 bits(7) | 223() | 7/7(100%) | 0/7(0%) | Plus/Plus | |

Features:

```
Query  293       CTTTATT  299
                 |||||||
Sbjct  32456764  CTTTATT  32456770
```

Range 345: 32457409 to 32457415

| Score | Expect | Identities | Gaps | Strand | Frame |
|---|---|---|---|---|---|
| 14.4 bits(7) | 223() | 7/7(100%) | 0/7(0%) | Plus/Plus | |

Features:

```
Query  290       CTCCTTT  296
                 |||||||
Sbjct  32457409  CTCCTTT  32457415
```

Range 346: 32458900 to 32458906

| Score | Expect | Identities | Gaps | Strand | Frame |
|---|---|---|---|---|---|
| 14.4 bits(7) | 223() | 7/7(100%) | 0/7(0%) | Plus/Plus | |

Features:

```
Query  278       AGCTGGT  284
                 |||||||
Sbjct  32458900  AGCTGGT  32458906
```

Range 347: 32459004 to 32459010

| Score | Expect | Identities | Gaps | Strand | Frame |
|---|---|---|---|---|---|
| 14.4 bits(7) | 223() | 7/7(100%) | 0/7(0%) | Plus/Plus | |

Features:

```
Query  272       GTAACAA  278
                 |||||||
Sbjct  32459004  GTAACAA  32459010
```

Range 348: 32459275 to 32459281

| Score | Expect | Identities | Gaps | Strand | Frame |
|---|---|---|---|---|---|
| 14.4 bits(7) | 223() | 7/7(100%) | 0/7(0%) | Plus/Plus | |

Features:

```
Query  290       CTCCTTT  296
                 |||||||
Sbjct  32459275  CTCCTTT  32459281
```

Range 349: 32459445 to 32459451

| Score | Expect | Identities | Gaps | Strand | Frame |
|---|---|---|---|---|---|
| 14.4 bits(7) | 223() | 7/7(100%) | 0/7(0%) | Plus/Plus | |

Features:

```
Query  285       GTTCTCT  291
                 |||||||
Sbjct  32459445  GTTCTCT  32459451
```

Range 350: 32459596 to 32459602

| Score | Expect | Identities | Gaps | Strand | Frame |
|---|---|---|---|---|---|
| 14.4 bits(7) | 223() | 7/7(100%) | 0/7(0%) | Plus/Plus | |

Features:

```
Query  292       CCTTTAT  298
                 |||||||
Sbjct  32459596  CCTTTAT  32459602
```

Range 351: 32461066 to 32461072

| Score | Expect | Identities | Gaps | Strand | Frame |
|---|---|---|---|---|---|
| 14.4 bits(7) | 223() | 7/7(100%) | 0/7(0%) | Plus/Plus | |

Features:

```
Query  284       TGTTCTC  290
                 |||||||
Sbjct  32461066  TGTTCTC  32461072
```

Range 352: 32461073 to 32461079

| Score | Expect | Identities | Gaps | Strand | Frame |
|---|---|---|---|---|---|
| 14.4 bits(7) | 223() | 7/7(100%) | 0/7(0%) | Plus/Plus | |

Features:

```
Query  288       CTCTCCT  294
                 |||||||
Sbjct  32461073  CTCTCCT  32461079
```

Range 353: 32461078 to 32461084

| Score | Expect | Identities | Gaps | Strand | Frame |
|---|---|---|---|---|---|

| | | | | |
|---|---|---|---|---|
| 14.4 bits(7) | 223() | 7/7(100%) | 0/7(0%) | Plus/Plus |

Features:

```
Query  288       CTCTCCT  294
                 |||||||
Sbjct  32461078  CTCTCCT  32461084
```

Range 354: 32461083 to 32461089

| Score | Expect | Identities | Gaps | Strand | Frame |
|---|---|---|---|---|---|
| 14.4 bits(7) | 223() | 7/7(100%) | 0/7(0%) | Plus/Plus | |

Features:

```
Query  288       CTCTCCT  294
                 |||||||
Sbjct  32461083  CTCTCCT  32461089
```

Range 355: 32461088 to 32461094

| Score | Expect | Identities | Gaps | Strand | Frame |
|---|---|---|---|---|---|
| 14.4 bits(7) | 223() | 7/7(100%) | 0/7(0%) | Plus/Plus | |

Features:

```
Query  288       CTCTCCT  294
                 |||||||
Sbjct  32461088  CTCTCCT  32461094
```

Range 356: 32461093 to 32461099

| Score | Expect | Identities | Gaps | Strand | Frame |
|---|---|---|---|---|---|
| 14.4 bits(7) | 223() | 7/7(100%) | 0/7(0%) | Plus/Plus | |

Features:

```
Query  288       CTCTCCT  294
                 |||||||
Sbjct  32461093  CTCTCCT  32461099
```

Range 357: 32461098 to 32461104

| Score | Expect | Identities | Gaps | Strand | Frame |
|---|---|---|---|---|---|
| 14.4 bits(7) | 223() | 7/7(100%) | 0/7(0%) | Plus/Plus | |

Features:

```
Query  288       CTCTCCT  294
                 |||||||
Sbjct  32461098  CTCTCCT  32461104
```

Range 358: 32461103 to 32461109

| Score | Expect | Identities | Gaps | Strand | Frame |
|---|---|---|---|---|---|
| 14.4 bits(7) | 223() | 7/7(100%) | 0/7(0%) | Plus/Plus | |

Features:

```
Query  288       CTCTCCT  294
                 |||||||
Sbjct  32461103  CTCTCCT  32461109
```

Range 359: 32461108 to 32461114

| Score | Expect | Identities | Gaps | Strand | Frame |
|---|---|---|---|---|---|
| 14.4 bits(7) | 223() | 7/7(100%) | 0/7(0%) | Plus/Plus | |

Features:

```
Query  288       CTCTCCT  294
                 |||||||
Sbjct  32461108  CTCTCCT  32461114
```

Range 360: 32461113 to 32461119

| Score | Expect | Identities | Gaps | Strand | Frame |
|---|---|---|---|---|---|
| 14.4 bits(7) | 223() | 7/7(100%) | 0/7(0%) | Plus/Plus | |

Features:

```
Query  288       CTCTCCT  294
                 |||||||
Sbjct  32461113  CTCTCCT  32461119
```

Range 361: 32461118 to 32461124

| Score | Expect | Identities | Gaps | Strand | Frame |
|---|---|---|---|---|---|
| 14.4 bits(7) | 223() | 7/7(100%) | 0/7(0%) | Plus/Plus | |

Features:

```
Query  288       CTCTCCT  294
                 |||||||
Sbjct  32461118  CTCTCCT  32461124
```

Range 362: 32461123 to 32461129

| Score | Expect | Identities | Gaps | Strand | Frame |
|---|---|---|---|---|---|
| 14.4 bits(7) | 223() | 7/7(100%) | 0/7(0%) | Plus/Plus | |

Features:

```
Query  288       CTCTCCT  294
                 |||||||
Sbjct  32461123  CTCTCCT  32461129
```

Range 363: 32461585 to 32461591

| Score | Expect | Identities | Gaps | Strand | Frame |
|---|---|---|---|---|---|
| 14.4 bits(7) | 223() | 7/7(100%) | 0/7(0%) | Plus/Plus | |

Features:

```
Query  281       TGGTGTT  287
                 |||||||
Sbjct  32461585  TGGTGTT  32461591
```

Range 364: 32461758 to 32461764

| Score | Expect | Identities | Gaps | Strand | Frame |
|---|---|---|---|---|---|
| 14.4 bits(7) | 223() | 7/7(100%) | 0/7(0%) | Plus/Plus | |

Features:

```
Query  292       CCTTTAT  298
                 |||||||
Sbjct  32461758  CCTTTAT  32461764
```

Range 365: 32462311 to 32462317

| Score | Expect | Identities | Gaps | Strand | Frame |
|---|---|---|---|---|---|
| 14.4 bits(7) | 223() | 7/7(100%) | 0/7(0%) | Plus/Minus | |

Features:

```
Query  277       AAGCTGG  283
                 |||||||
Sbjct  32462317  AAGCTGG  32462311
```

Range 366: 32462499 to 32462505

| Score | Expect | Identities | Gaps | Strand | Frame |
|---|---|---|---|---|---|
| 14.4 bits(7) | 223() | 7/7(100%) | 0/7(0%) | Plus/Plus | |

Features:

```
Query  287       TCTCTCC  293
                 |||||||
Sbjct  32462499  TCTCTCC  32462505
```

Range 367: 32463020 to 32463026

| Score | Expect | Identities | Gaps | Strand | Frame |
|---|---|---|---|---|---|
| 14.4 bits(7) | 223() | 7/7(100%) | 0/7(0%) | Plus/Plus | |

Features:

```
Query  293       CTTTATT  299
                 |||||||
Sbjct  32463020  CTTTATT  32463026
```

Range 368: 32464111 to 32464117

| Score | Expect | Identities | Gaps | Strand | Frame |
|---|---|---|---|---|---|
| 14.4 bits(7) | 223() | 7/7(100%) | 0/7(0%) | Plus/Plus | |

Features:

```
Query  280       CTGGTGT  286
                 |||||||
Sbjct  32464111  CTGGTGT  32464117
```

Range 369: 32464515 to 32464521

| Score | Expect | Identities | Gaps | Strand | Frame |
|---|---|---|---|---|---|
| 14.4 bits(7) | 223() | 7/7(100%) | 0/7(0%) | Plus/Plus | |

Features:

```
Query  277       AAGCTGG  283
                 |||||||
Sbjct  32464515  AAGCTGG  32464521
```

Range 370: 32465935 to 32465941

| Score | Expect | Identities | Gaps | Strand | Frame |
|---|---|---|---|---|---|
| 14.4 bits(7) | 223() | 7/7(100%) | 0/7(0%) | Plus/Plus | |

Features:

```
Query  272       GTAACAA  278
                 |||||||
Sbjct  32465935  GTAACAA  32465941
```

Range 371: 32466108 to 32466114

| Score | Expect | Identities | Gaps | Strand | Frame |
|---|---|---|---|---|---|
| 14.4 bits(7) | 223() | 7/7(100%) | 0/7(0%) | Plus/Minus | |

Features:

```
Query  273       TAACAAG  279
                 |||||||
Sbjct  32466114  TAACAAG  32466108
```

Range 372: 32466218 to 32466224

| Score | Expect | Identities | Gaps | Strand | Frame |
|---|---|---|---|---|---|

| | | | | |
|---|---|---|---|---|
| 14.4 bits(7) | 223() | 7/7(100%) | 0/7(0%) | Plus/Plus |

Features:

```
Query  278       AGCTGGT  284
                 |||||||
Sbjct  32466218  AGCTGGT  32466224
```

Range 373: 32466663 to 32466669

| Score | Expect | Identities | Gaps | Strand | Frame |
|---|---|---|---|---|---|
| 14.4 bits(7) | 223() | 7/7(100%) | 0/7(0%) | Plus/Plus | |

Features:

```
Query  291       TCCTTTA  297
                 |||||||
Sbjct  32466663  TCCTTTA  32466669
```

Range 374: 32466764 to 32466770

| Score | Expect | Identities | Gaps | Strand | Frame |
|---|---|---|---|---|---|
| 14.4 bits(7) | 223() | 7/7(100%) | 0/7(0%) | Plus/Plus | |

Features:

```
Query  271       TGTAACA  277
                 |||||||
Sbjct  32466764  TGTAACA  32466770
```

Range 375: 32467514 to 32467520

| Score | Expect | Identities | Gaps | Strand | Frame |
|---|---|---|---|---|---|
| 14.4 bits(7) | 223() | 7/7(100%) | 0/7(0%) | Plus/Plus | |

Features:

```
Query  286       TTCTCTC  292
                 |||||||
Sbjct  32467514  TTCTCTC  32467520
```

Range 376: 32469232 to 32469238

| Score | Expect | Identities | Gaps | Strand | Frame |
|---|---|---|---|---|---|
| 14.4 bits(7) | 223() | 7/7(100%) | 0/7(0%) | Plus/Plus | |

Features:

```
Query  271       TGTAACA  277
                 |||||||
Sbjct  32469232  TGTAACA  32469238
```

Range 377: 32469760 to 32469766

| Score | Expect | Identities | Gaps | Strand | Frame |
|---|---|---|---|---|---|
| 14.4 bits(7) | 223() | 7/7(100%) | 0/7(0%) | Plus/Minus | |

Features:

```
Query  279       GCTGGTG  285
                 |||||||
Sbjct  32469766  GCTGGTG  32469760
```

Range 378: 32469919 to 32469925

| Score | Expect | Identities | Gaps | Strand | Frame |
|---|---|---|---|---|---|
| 14.4 bits(7) | 223() | 7/7(100%) | 0/7(0%) | Plus/Plus | |

Features:

```
Query  286       TTCTCTC  292
                 |||||||
Sbjct  32469919  TTCTCTC  32469925
```

Range 379: 32469946 to 32469952

| Score | Expect | Identities | Gaps | Strand | Frame |
|---|---|---|---|---|---|
| 14.4 bits(7) | 223() | 7/7(100%) | 0/7(0%) | Plus/Minus | |

Features:

```
Query  273       TAACAAG  279
                 |||||||
Sbjct  32469952  TAACAAG  32469946
```

Range 380: 32470318 to 32470324

| Score | Expect | Identities | Gaps | Strand | Frame |
|---|---|---|---|---|---|
| 14.4 bits(7) | 223() | 7/7(100%) | 0/7(0%) | Plus/Plus | |

Features:

```
Query  283       GTGTTCT  289
                 |||||||
Sbjct  32470318  GTGTTCT  32470324
```

Range 381: 32471005 to 32471011

| Score | Expect | Identities | Gaps | Strand | Frame |
|---|---|---|---|---|---|
| 14.4 bits(7) | 223() | 7/7(100%) | 0/7(0%) | Plus/Plus | |

Features:

```
Query  287       TCTCTCC  293
                 |||||||
Sbjct  32471005  TCTCTCC  32471011
```

Range 382: 32471235 to 32471241

| Score | Expect | Identities | Gaps | Strand | Frame |
|---|---|---|---|---|---|
| 14.4 bits(7) | 223() | 7/7(100%) | 0/7(0%) | Plus/Minus | |

Features:

```
Query  278       AGCTGGT  284
                 |||||||
Sbjct  32471241  AGCTGGT  32471235
```

Range 383: 32471323 to 32471329

| Score | Expect | Identities | Gaps | Strand | Frame |
|---|---|---|---|---|---|
| 14.4 bits(7) | 223() | 7/7(100%) | 0/7(0%) | Plus/Plus | |

Features:

```
Query  275       ACAAGCT  281
                 |||||||
Sbjct  32471323  ACAAGCT  32471329
```

Range 384: 32471328 to 32471334

| Score | Expect | Identities | Gaps | Strand | Frame |
|---|---|---|---|---|---|
| 14.4 bits(7) | 223() | 7/7(100%) | 0/7(0%) | Plus/Plus | |

Features:

```
Query  290       CTCCTTT  296
                 |||||||
Sbjct  32471328  CTCCTTT  32471334
```

Range 385: 32471397 to 32471403

| Score | Expect | Identities | Gaps | Strand | Frame |
|---|---|---|---|---|---|
| 14.4 bits(7) | 223() | 7/7(100%) | 0/7(0%) | Plus/Minus | |

Features:

```
Query  274       AACAAGC  280
                 |||||||
Sbjct  32471403  AACAAGC  32471397
```

Range 386: 32471612 to 32471618

| Score | Expect | Identities | Gaps | Strand | Frame |
|---|---|---|---|---|---|
| 14.4 bits(7) | 223() | 7/7(100%) | 0/7(0%) | Plus/Minus | |

Features:

```
Query  291       TCCTTTA  297
                 |||||||
Sbjct  32471618  TCCTTTA  32471612
```

Range 387: 32471658 to 32471664

| Score | Expect | Identities | Gaps | Strand | Frame |
|---|---|---|---|---|---|
| 14.4 bits(7) | 223() | 7/7(100%) | 0/7(0%) | Plus/Plus | |

Features:

```
Query  273       TAACAAG  279
                 |||||||
Sbjct  32471658  TAACAAG  32471664
```

Range 388: 32471678 to 32471684

| Score | Expect | Identities | Gaps | Strand | Frame |
|---|---|---|---|---|---|
| 14.4 bits(7) | 223() | 7/7(100%) | 0/7(0%) | Plus/Plus | |

Features:

```
Query  293       CTTTATT  299
                 |||||||
Sbjct  32471678  CTTTATT  32471684
```

Range 389: 32472535 to 32472541

| Score | Expect | Identities | Gaps | Strand | Frame |
|---|---|---|---|---|---|
| 14.4 bits(7) | 223() | 7/7(100%) | 0/7(0%) | Plus/Minus | |

Features:

```
Query  286       TTCTCTC  292
                 |||||||
Sbjct  32472541  TTCTCTC  32472535
```

Range 390: 32473439 to 32473445

| Score | Expect | Identities | Gaps | Strand | Frame |
|---|---|---|---|---|---|
| 14.4 bits(7) | 223() | 7/7(100%) | 0/7(0%) | Plus/Plus | |

Features:

```
Query  291       TCCTTTA  297
                 |||||||
Sbjct  32473439  TCCTTTA  32473445
```

Range 391: 32474544 to 32474550

| Score | Expect | Identities | Gaps | Strand | Frame |
|---|---|---|---|---|---|

| | | | | |
|---|---|---|---|---|
| 14.4 bits(7) | 223() | 7/7(100%) | 0/7(0%) | Plus/Plus |

Features:

```
Query  286       TTCTCTC  292
                 |||||||
Sbjct  32474544  TTCTCTC  32474550
```

Range 392: 32475919 to 32475925

| Score | Expect | Identities | Gaps | Strand | Frame |
|---|---|---|---|---|---|
| 14.4 bits(7) | 223() | 7/7(100%) | 0/7(0%) | Plus/Plus | |

Features:

```
Query  286       TTCTCTC  292
                 |||||||
Sbjct  32475919  TTCTCTC  32475925
```

Range 393: 32476028 to 32476034

| Score | Expect | Identities | Gaps | Strand | Frame |
|---|---|---|---|---|---|
| 14.4 bits(7) | 223() | 7/7(100%) | 0/7(0%) | Plus/Minus | |

Features:

```
Query  287       TCTCTCC  293
                 |||||||
Sbjct  32476034  TCTCTCC  32476028
```

Range 394: 32476280 to 32476286

| Score | Expect | Identities | Gaps | Strand | Frame |
|---|---|---|---|---|---|
| 14.4 bits(7) | 223() | 7/7(100%) | 0/7(0%) | Plus/Plus | |

Features:

```
Query  272       GTAACAA  278
                 |||||||
Sbjct  32476280  GTAACAA  32476286
```

Range 395: 32476313 to 32476319

| Score | Expect | Identities | Gaps | Strand | Frame |
|---|---|---|---|---|---|
| 14.4 bits(7) | 223() | 7/7(100%) | 0/7(0%) | Plus/Plus | |

Features:

```
Query  274       AACAAGC  280
                 |||||||
Sbjct  32476313  AACAAGC  32476319
```

Range 396: 32476881 to 32476887

| Score | Expect | Identities | Gaps | Strand | Frame |
|---|---|---|---|---|---|
| 14.4 bits(7) | 223() | 7/7(100%) | 0/7(0%) | Plus/Plus | |

Features:

```
Query  281       TGGTGTT  287
                 |||||||
Sbjct  32476881  TGGTGTT  32476887
```

Range 397: 32477154 to 32477160

| Score | Expect | Identities | Gaps | Strand | Frame |
|---|---|---|---|---|---|
| 14.4 bits(7) | 223() | 7/7(100%) | 0/7(0%) | Plus/Minus | |

Features:

```
Query  271       TGTAACA  277
                 |||||||
Sbjct  32477160  TGTAACA  32477154
```

Range 398: 32478931 to 32478937

| Score | Expect | Identities | Gaps | Strand | Frame |
|---|---|---|---|---|---|
| 14.4 bits(7) | 223() | 7/7(100%) | 0/7(0%) | Plus/Minus | |

Features:

```
Query  292       CCTTTAT  298
                 |||||||
Sbjct  32478937  CCTTTAT  32478931
```

Range 399: 32479079 to 32479085

| Score | Expect | Identities | Gaps | Strand | Frame |
|---|---|---|---|---|---|
| 14.4 bits(7) | 223() | 7/7(100%) | 0/7(0%) | Plus/Plus | |

Features:

```
Query  293       CTTTATT  299
                 |||||||
Sbjct  32479079  CTTTATT  32479085
```

Range 400: 32479642 to 32479648

| Score | Expect | Identities | Gaps | Strand | Frame |
|---|---|---|---|---|---|
| 14.4 bits(7) | 223() | 7/7(100%) | 0/7(0%) | Plus/Plus | |

Features:

```
Query  293       CTTTATT  299
                 |||||||
Sbjct  32479642  CTTTATT  32479648
```

Range 401: 32479673 to 32479679

| Score | Expect | Identities | Gaps | Strand | Frame |
|---|---|---|---|---|---|
| 14.4 bits(7) | 223() | 7/7(100%) | 0/7(0%) | Plus/Minus | |

Features:

```
Query  293       CTTTATT  299
                 |||||||
Sbjct  32479679  CTTTATT  32479673
```

Range 402: 32479693 to 32479699

| Score | Expect | Identities | Gaps | Strand | Frame |
|---|---|---|---|---|---|
| 14.4 bits(7) | 223() | 7/7(100%) | 0/7(0%) | Plus/Minus | |

Features:

```
Query  293       CTTTATT  299
                 |||||||
Sbjct  32479699  CTTTATT  32479693
```

Range 403: 32480855 to 32480861

| Score | Expect | Identities | Gaps | Strand | Frame |
|---|---|---|---|---|---|
| 14.4 bits(7) | 223() | 7/7(100%) | 0/7(0%) | Plus/Plus | |

Features:

```
Query  284       TGTTCTC  290
                 |||||||
Sbjct  32480855  TGTTCTC  32480861
```

Range 404: 32481218 to 32481224

| Score | Expect | Identities | Gaps | Strand | Frame |
|---|---|---|---|---|---|
| 14.4 bits(7) | 223() | 7/7(100%) | 0/7(0%) | Plus/Plus | |

Features:

```
Query  271       TGTAACA  277
                 |||||||
Sbjct  32481218  TGTAACA  32481224
```

Range 405: 32481402 to 32481408

| Score | Expect | Identities | Gaps | Strand | Frame |
|---|---|---|---|---|---|
| 14.4 bits(7) | 223() | 7/7(100%) | 0/7(0%) | Plus/Plus | |

Features:

```
Query  291       TCCTTTA  297
                 |||||||
Sbjct  32481402  TCCTTTA  32481408
```

Range 406: 32481479 to 32481485

| Score | Expect | Identities | Gaps | Strand | Frame |
|---|---|---|---|---|---|
| 14.4 bits(7) | 223() | 7/7(100%) | 0/7(0%) | Plus/Minus | |

Features:

```
Query  290       CTCCTTT  296
                 |||||||
Sbjct  32481485  CTCCTTT  32481479
```

Range 407: 32482526 to 32482532

| Score | Expect | Identities | Gaps | Strand | Frame |
|---|---|---|---|---|---|
| 14.4 bits(7) | 223() | 7/7(100%) | 0/7(0%) | Plus/Minus | |

Features:

```
Query  279       GCTGGTG  285
                 |||||||
Sbjct  32482532  GCTGGTG  32482526
```

Range 408: 32482768 to 32482774

| Score | Expect | Identities | Gaps | Strand | Frame |
|---|---|---|---|---|---|
| 14.4 bits(7) | 223() | 7/7(100%) | 0/7(0%) | Plus/Plus | |

Features:

```
Query  285       GTTCTCT  291
                 |||||||
Sbjct  32482768  GTTCTCT  32482774
```

Range 409: 32483215 to 32483221

| Score | Expect | Identities | Gaps | Strand | Frame |
|---|---|---|---|---|---|
| 14.4 bits(7) | 223() | 7/7(100%) | 0/7(0%) | Plus/Plus | |

Features:

```
Query  293       CTTTATT  299
                 |||||||
Sbjct  32483215  CTTTATT  32483221
```

Range 410: 32483954 to 32483960

| Score | Expect | Identities | Gaps | Strand | Frame |
|---|---|---|---|---|---|

| Score | Expect | Identities | Gaps | Strand | Frame |
|---|---|---|---|---|---|
| 14.4 bits(7) | 223() | 7/7(100%) | 0/7(0%) | Plus/Plus | |

Features:

```
Query  284       TGTTCTC  290
                 |||||||
Sbjct  32483954  TGTTCTC  32483960
```

Range 411: 32484090 to 32484096

| Score | Expect | Identities | Gaps | Strand | Frame |
|---|---|---|---|---|---|
| 14.4 bits(7) | 223() | 7/7(100%) | 0/7(0%) | Plus/Plus | |

Features:

```
Query  284       TGTTCTC  290
                 |||||||
Sbjct  32484090  TGTTCTC  32484096
```

Range 412: 32485019 to 32485025

| Score | Expect | Identities | Gaps | Strand | Frame |
|---|---|---|---|---|---|
| 14.4 bits(7) | 223() | 7/7(100%) | 0/7(0%) | Plus/Plus | |

Features:

```
Query  291       TCCTTTA  297
                 |||||||
Sbjct  32485019  TCCTTTA  32485025
```

Range 413: 32485348 to 32485354

| Score | Expect | Identities | Gaps | Strand | Frame |
|---|---|---|---|---|---|
| 14.4 bits(7) | 223() | 7/7(100%) | 0/7(0%) | Plus/Plus | |

Features:

```
Query  293       CTTTATT  299
                 |||||||
Sbjct  32485348  CTTTATT  32485354
```

Range 414: 32485407 to 32485413

| Score | Expect | Identities | Gaps | Strand | Frame |
|---|---|---|---|---|---|
| 14.4 bits(7) | 223() | 7/7(100%) | 0/7(0%) | Plus/Minus | |

Features:

```
Query  272       GTAACAA  278
                 |||||||
Sbjct  32485413  GTAACAA  32485407
```

Range 415: 32485465 to 32485471

| Score | Expect | Identities | Gaps | Strand | Frame |
|---|---|---|---|---|---|
| 14.4 bits(7) | 223() | 7/7(100%) | 0/7(0%) | Plus/Plus | |

Features:

```
Query  290       CTCCTTT  296
                 |||||||
Sbjct  32485465  CTCCTTT  32485471
```

Range 416: 32485531 to 32485537

| Score | Expect | Identities | Gaps | Strand | Frame |
|---|---|---|---|---|---|
| 14.4 bits(7) | 223() | 7/7(100%) | 0/7(0%) | Plus/Minus | |

Features:

```
Query  283       GTGTTCT  289
                 |||||||
Sbjct  32485537  GTGTTCT  32485531
```

Range 417: 32486259 to 32486265

| Score | Expect | Identities | Gaps | Strand | Frame |
|---|---|---|---|---|---|
| 14.4 bits(7) | 223() | 7/7(100%) | 0/7(0%) | Plus/Minus | |

Features:

```
Query  291       TCCTTTA  297
                 |||||||
Sbjct  32486265  TCCTTTA  32486259
```

Range 418: 32486382 to 32486388

| Score | Expect | Identities | Gaps | Strand | Frame |
|---|---|---|---|---|---|
| 14.4 bits(7) | 223() | 7/7(100%) | 0/7(0%) | Plus/Minus | |

Features:

```
Query  271       TGTAACA  277
                 |||||||
Sbjct  32486388  TGTAACA  32486382
```

Range 419: 32486810 to 32486816

| Score | Expect | Identities | Gaps | Strand | Frame |
|---|---|---|---|---|---|
| 14.4 bits(7) | 223() | 7/7(100%) | 0/7(0%) | Plus/Plus | |

Features:

```
Query  283       GTGTTCT  289
                 |||||||
Sbjct  32486810  GTGTTCT  32486816
```

Range 420: 32487444 to 32487450

| Score | Expect | Identities | Gaps | Strand | Frame |
|---|---|---|---|---|---|
| 14.4 bits(7) | 223() | 7/7(100%) | 0/7(0%) | Plus/Plus | |

Features:

```
Query  286       TTCTCTC  292
                 |||||||
Sbjct  32487444  TTCTCTC  32487450
```

Range 421: 32487714 to 32487720

| Score | Expect | Identities | Gaps | Strand | Frame |
|---|---|---|---|---|---|
| 14.4 bits(7) | 223() | 7/7(100%) | 0/7(0%) | Plus/Minus | |

Features:

```
Query  277       AAGCTGG  283
                 |||||||
Sbjct  32487720  AAGCTGG  32487714
```

Range 422: 32487904 to 32487910

| Score | Expect | Identities | Gaps | Strand | Frame |
|---|---|---|---|---|---|
| 14.4 bits(7) | 223() | 7/7(100%) | 0/7(0%) | Plus/Plus | |

Features:

```
Query  293       CTTTATT  299
                 |||||||
Sbjct  32487904  CTTTATT  32487910
```

Range 423: 32488178 to 32488184

| Score | Expect | Identities | Gaps | Strand | Frame |
|---|---|---|---|---|---|
| 14.4 bits(7) | 223() | 7/7(100%) | 0/7(0%) | Plus/Minus | |

Features:

```
Query  293       CTTTATT  299
                 |||||||
Sbjct  32488184  CTTTATT  32488178
```

Range 424: 32489264 to 32489270

| Score | Expect | Identities | Gaps | Strand | Frame |
|---|---|---|---|---|---|
| 14.4 bits(7) | 223() | 7/7(100%) | 0/7(0%) | Plus/Plus | |

Features:

```
Query  286       TTCTCTC  292
                 |||||||
Sbjct  32489264  TTCTCTC  32489270
```

Range 425: 32489382 to 32489388

| Score | Expect | Identities | Gaps | Strand | Frame |
|---|---|---|---|---|---|
| 14.4 bits(7) | 223() | 7/7(100%) | 0/7(0%) | Plus/Minus | |

Features:

```
Query  279       GCTGGTG  285
                 |||||||
Sbjct  32489388  GCTGGTG  32489382
```

Range 426: 32490197 to 32490203

| Score | Expect | Identities | Gaps | Strand | Frame |
|---|---|---|---|---|---|
| 14.4 bits(7) | 223() | 7/7(100%) | 0/7(0%) | Plus/Plus | |

Features:

```
Query  283       GTGTTCT  289
                 |||||||
Sbjct  32490197  GTGTTCT  32490203
```

Range 427: 32490602 to 32490608

| Score | Expect | Identities | Gaps | Strand | Frame |
|---|---|---|---|---|---|
| 14.4 bits(7) | 223() | 7/7(100%) | 0/7(0%) | Plus/Minus | |

Features:

```
Query  274       AACAAGC  280
                 |||||||
Sbjct  32490608  AACAAGC  32490602
```

Range 428: 32491423 to 32491429

| Score | Expect | Identities | Gaps | Strand | Frame |
|---|---|---|---|---|---|
| 14.4 bits(7) | 223() | 7/7(100%) | 0/7(0%) | Plus/Plus | |

Features:

```
Query  278       AGCTGGT  284
                 |||||||
Sbjct  32491423  AGCTGGT  32491429
```

Range 429: 32491452 to 32491458

| Score | Expect | Identities | Gaps | Strand | Frame |
|---|---|---|---|---|---|

| 14.4 bits(7) | 223() | 7/7(100%) | 0/7(0%) | Plus/Minus |
|---|---|---|---|---|

Features:

```
Query  288       CTCTCCT  294
                 |||||||
Sbjct  32491458  CTCTCCT  32491452
```

Range 430: 32492041 to 32492047

| Score | Expect | Identities | Gaps | Strand | Frame |
|---|---|---|---|---|---|
| 14.4 bits(7) | 223() | 7/7(100%) | 0/7(0%) | Plus/Minus | |

Features:

```
Query  273       TAACAAG  279
                 |||||||
Sbjct  32492047  TAACAAG  32492041
```

Range 431: 32492277 to 32492283

| Score | Expect | Identities | Gaps | Strand | Frame |
|---|---|---|---|---|---|
| 14.4 bits(7) | 223() | 7/7(100%) | 0/7(0%) | Plus/Minus | |

Features:

```
Query  274       AACAAGC  280
                 |||||||
Sbjct  32492283  AACAAGC  32492277
```

Range 432: 32492415 to 32492421

| Score | Expect | Identities | Gaps | Strand | Frame |
|---|---|---|---|---|---|
| 14.4 bits(7) | 223() | 7/7(100%) | 0/7(0%) | Plus/Minus | |

Features:

```
Query  286       TTCTCTC  292
                 |||||||
Sbjct  32492421  TTCTCTC  32492415
```

Range 433: 32492685 to 32492691

| Score | Expect | Identities | Gaps | Strand | Frame |
|---|---|---|---|---|---|
| 14.4 bits(7) | 223() | 7/7(100%) | 0/7(0%) | Plus/Minus | |

Features:

```
Query  290       CTCCTTT  296
                 |||||||
Sbjct  32492691  CTCCTTT  32492685
```

Range 434: 32492813 to 32492819

| Score | Expect | Identities | Gaps | Strand | Frame |
|---|---|---|---|---|---|
| 14.4 bits(7) | 223() | 7/7(100%) | 0/7(0%) | Plus/Plus | |

Features:

```
Query  293       CTTTATT  299
                 |||||||
Sbjct  32492813  CTTTATT  32492819
```

Range 435: 32493557 to 32493563

| Score | Expect | Identities | Gaps | Strand | Frame |
|---|---|---|---|---|---|
| 14.4 bits(7) | 223() | 7/7(100%) | 0/7(0%) | Plus/Plus | |

Features:

```
Query  275       ACAAGCT  281
                 |||||||
Sbjct  32493557  ACAAGCT  32493563
```

Range 436: 32495056 to 32495062

| Score | Expect | Identities | Gaps | Strand | Frame |
|---|---|---|---|---|---|
| 14.4 bits(7) | 223() | 7/7(100%) | 0/7(0%) | Plus/Minus | |

Features:

```
Query  293       CTTTATT  299
                 |||||||
Sbjct  32495062  CTTTATT  32495056
```

Range 437: 32495205 to 32495211

| Score | Expect | Identities | Gaps | Strand | Frame |
|---|---|---|---|---|---|
| 14.4 bits(7) | 223() | 7/7(100%) | 0/7(0%) | Plus/Minus | |

Features:

```
Query  291       TCCTTTA  297
                 |||||||
Sbjct  32495211  TCCTTTA  32495205
```

Range 438: 32495574 to 32495580

| Score | Expect | Identities | Gaps | Strand | Frame |
|---|---|---|---|---|---|
| 14.4 bits(7) | 223() | 7/7(100%) | 0/7(0%) | Plus/Plus | |

Features:

```
Query  293       CTTTATT  299
                 |||||||
Sbjct  32495574  CTTTATT  32495580
```

Range 439: 32495657 to 32495663

| Score | Expect | Identities | Gaps | Strand | Frame |
|---|---|---|---|---|---|
| 14.4 bits(7) | 223() | 7/7(100%) | 0/7(0%) | Plus/Minus | |

Features:

```
Query  272       GTAACAA  278
                 |||||||
Sbjct  32495663  GTAACAA  32495657
```

Range 440: 32496650 to 32496656

| Score | Expect | Identities | Gaps | Strand | Frame |
|---|---|---|---|---|---|
| 14.4 bits(7) | 223() | 7/7(100%) | 0/7(0%) | Plus/Plus | |

Features:

```
Query  293       CTTTATT  299
                 |||||||
Sbjct  32496650  CTTTATT  32496656
```

Range 441: 32496792 to 32496798

| Score | Expect | Identities | Gaps | Strand | Frame |
|---|---|---|---|---|---|
| 14.4 bits(7) | 223() | 7/7(100%) | 0/7(0%) | Plus/Minus | |

Features:

```
Query  274       AACAAGC  280
                 |||||||
Sbjct  32496798  AACAAGC  32496792
```

Range 442: 32496836 to 32496842

| Score | Expect | Identities | Gaps | Strand | Frame |
|---|---|---|---|---|---|
| 14.4 bits(7) | 223() | 7/7(100%) | 0/7(0%) | Plus/Plus | |

Features:

```
Query  290       CTCCTTT  296
                 |||||||
Sbjct  32496836  CTCCTTT  32496842
```

Range 443: 32497180 to 32497186

| Score | Expect | Identities | Gaps | Strand | Frame |
|---|---|---|---|---|---|
| 14.4 bits(7) | 223() | 7/7(100%) | 0/7(0%) | Plus/Plus | |

Features:

```
Query  281       TGGTGTT  287
                 |||||||
Sbjct  32497180  TGGTGTT  32497186
```

Range 444: 32497601 to 32497607

| Score | Expect | Identities | Gaps | Strand | Frame |
|---|---|---|---|---|---|
| 14.4 bits(7) | 223() | 7/7(100%) | 0/7(0%) | Plus/Plus | |

Features:

```
Query  291       TCCTTTA  297
                 |||||||
Sbjct  32497601  TCCTTTA  32497607
```

Range 445: 32498896 to 32498902

| Score | Expect | Identities | Gaps | Strand | Frame |
|---|---|---|---|---|---|
| 14.4 bits(7) | 223() | 7/7(100%) | 0/7(0%) | Plus/Plus | |

Features:

```
Query  272       GTAACAA  278
                 |||||||
Sbjct  32498896  GTAACAA  32498902
```

Range 446: 32499851 to 32499857

| Score | Expect | Identities | Gaps | Strand | Frame |
|---|---|---|---|---|---|
| 14.4 bits(7) | 223() | 7/7(100%) | 0/7(0%) | Plus/Minus | |

Features:

```
Query  273       TAACAAG  279
                 |||||||
Sbjct  32499857  TAACAAG  32499851
```

Range 447: 32501046 to 32501052

| Score | Expect | Identities | Gaps | Strand | Frame |
|---|---|---|---|---|---|
| 14.4 bits(7) | 223() | 7/7(100%) | 0/7(0%) | Plus/Plus | |

Features:

```
Query  271       TGTAACA  277
                 |||||||
Sbjct  32501046  TGTAACA  32501052
```

Range 448: 32501333 to 32501339

| Score | Expect | Identities | Gaps | Strand | Frame |
|---|---|---|---|---|---|

| 14.4 bits(7) | 223() | 7/7(100%) | 0/7(0%) | Plus/Plus |
|---|---|---|---|---|

Features:

```
Query  284       TGTTCTC  290
                 |||||||
Sbjct  32501333  TGTTCTC  32501339
```

Range 449: 32502046 to 32502052

| Score | Expect | Identities | Gaps | Strand | Frame |
|---|---|---|---|---|---|
| 14.4 bits(7) | 223() | 7/7(100%) | 0/7(0%) | Plus/Minus | |

Features:

```
Query  293       CTTTATT  299
                 |||||||
Sbjct  32502052  CTTTATT  32502046
```

Range 450: 32502942 to 32502948

| Score | Expect | Identities | Gaps | Strand | Frame |
|---|---|---|---|---|---|
| 14.4 bits(7) | 223() | 7/7(100%) | 0/7(0%) | Plus/Minus | |

Features:

```
Query  271       TGTAACA  277
                 |||||||
Sbjct  32502948  TGTAACA  32502942
```

Range 451: 32503149 to 32503155

| Score | Expect | Identities | Gaps | Strand | Frame |
|---|---|---|---|---|---|
| 14.4 bits(7) | 223() | 7/7(100%) | 0/7(0%) | Plus/Minus | |

Features:

```
Query  293       CTTTATT  299
                 |||||||
Sbjct  32503155  CTTTATT  32503149
```

Range 452: 32503287 to 32503293

| Score | Expect | Identities | Gaps | Strand | Frame |
|---|---|---|---|---|---|
| 14.4 bits(7) | 223() | 7/7(100%) | 0/7(0%) | Plus/Plus | |

Features:

```
Query  278       AGCTGGT  284
                 |||||||
Sbjct  32503287  AGCTGGT  32503293
```

Range 453: 32504321 to 32504327

| Score | Expect | Identities | Gaps | Strand | Frame |
|---|---|---|---|---|---|
| 14.4 bits(7) | 223() | 7/7(100%) | 0/7(0%) | Plus/Minus | |

Features:

```
Query  293       CTTTATT  299
                 |||||||
Sbjct  32504327  CTTTATT  32504321
```

Range 454: 32504950 to 32504956

| Score | Expect | Identities | Gaps | Strand | Frame |
|---|---|---|---|---|---|
| 14.4 bits(7) | 223() | 7/7(100%) | 0/7(0%) | Plus/Plus | |

Features:

```
Query  293       CTTTATT  299
                 |||||||
Sbjct  32504950  CTTTATT  32504956
```

Range 455: 32505036 to 32505042

| Score | Expect | Identities | Gaps | Strand | Frame |
|---|---|---|---|---|---|
| 14.4 bits(7) | 223() | 7/7(100%) | 0/7(0%) | Plus/Minus | |

Features:

```
Query  278       AGCTGGT  284
                 |||||||
Sbjct  32505042  AGCTGGT  32505036
```

Range 456: 32505166 to 32505172

| Score | Expect | Identities | Gaps | Strand | Frame |
|---|---|---|---|---|---|
| 14.4 bits(7) | 223() | 7/7(100%) | 0/7(0%) | Plus/Minus | |

Features:

```
Query  289       TCTCCTT  295
                 |||||||
Sbjct  32505172  TCTCCTT  32505166
```

Range 457: 32505235 to 32505241

| Score | Expect | Identities | Gaps | Strand | Frame |
|---|---|---|---|---|---|
| 14.4 bits(7) | 223() | 7/7(100%) | 0/7(0%) | Plus/Minus | |

Features:

```
Query  284       TGTTCTC  290
                 |||||||
Sbjct  32505241  TGTTCTC  32505235
```

Range 458: 32505459 to 32505465

| Score | Expect | Identities | Gaps | Strand | Frame |
|---|---|---|---|---|---|
| 14.4 bits(7) | 223() | 7/7(100%) | 0/7(0%) | Plus/Plus | |

Features:

```
Query  286       TTCTCTC  292
                 |||||||
Sbjct  32505459  TTCTCTC  32505465
```

Range 459: 32506415 to 32506421

| Score | Expect | Identities | Gaps | Strand | Frame |
|---|---|---|---|---|---|
| 14.4 bits(7) | 223() | 7/7(100%) | 0/7(0%) | Plus/Minus | |

Features:

```
Query  292       CCTTTAT  298
                 |||||||
Sbjct  32506421  CCTTTAT  32506415
```

Range 460: 32506552 to 32506558

| Score | Expect | Identities | Gaps | Strand | Frame |
|---|---|---|---|---|---|
| 14.4 bits(7) | 223() | 7/7(100%) | 0/7(0%) | Plus/Minus | |

Features:

```
Query  290       CTCCTTT  296
                 |||||||
Sbjct  32506558  CTCCTTT  32506552
```

Range 461: 32506667 to 32506673

| Score | Expect | Identities | Gaps | Strand | Frame |
|---|---|---|---|---|---|
| 14.4 bits(7) | 223() | 7/7(100%) | 0/7(0%) | Plus/Plus | |

Features:

```
Query  282       GGTGTTC  288
                 |||||||
Sbjct  32506667  GGTGTTC  32506673
```

Range 462: 32506937 to 32506943

| Score | Expect | Identities | Gaps | Strand | Frame |
|---|---|---|---|---|---|
| 14.4 bits(7) | 223() | 7/7(100%) | 0/7(0%) | Plus/Plus | |

Features:

```
Query  277       AAGCTGG  283
                 |||||||
Sbjct  32506937  AAGCTGG  32506943
```

Range 463: 32507142 to 32507148

| Score | Expect | Identities | Gaps | Strand | Frame |
|---|---|---|---|---|---|
| 14.4 bits(7) | 223() | 7/7(100%) | 0/7(0%) | Plus/Minus | |

Features:

```
Query  290       CTCCTTT  296
                 |||||||
Sbjct  32507148  CTCCTTT  32507142
```

Range 464: 32508691 to 32508697

| Score | Expect | Identities | Gaps | Strand | Frame |
|---|---|---|---|---|---|
| 14.4 bits(7) | 223() | 7/7(100%) | 0/7(0%) | Plus/Plus | |

Features:

```
Query  273       TAACAAG  279
                 |||||||
Sbjct  32508691  TAACAAG  32508697
```

Range 465: 32510536 to 32510542

| Score | Expect | Identities | Gaps | Strand | Frame |
|---|---|---|---|---|---|
| 14.4 bits(7) | 223() | 7/7(100%) | 0/7(0%) | Plus/Plus | |

Features:

```
Query  291       TCCTTTA  297
                 |||||||
Sbjct  32510536  TCCTTTA  32510542
```

Range 466: 32512157 to 32512163

| Score | Expect | Identities | Gaps | Strand | Frame |
|---|---|---|---|---|---|
| 14.4 bits(7) | 223() | 7/7(100%) | 0/7(0%) | Plus/Plus | |

Features:

```
Query  293       CTTTATT  299
                 |||||||
Sbjct  32512157  CTTTATT  32512163
```

Range 467: 32512426 to 32512432

| Score | Expect | Identities | Gaps | Strand | Frame |
|---|---|---|---|---|---|

| Score | Expect | Identities | Gaps | Strand | Frame |
|---|---|---|---|---|---|
| 14.4 bits(7) | 223() | 7/7(100%) | 0/7(0%) | Plus/Minus | |

Features:

```
Query  273       TAACAAG  279
                 |||||||
Sbjct  32512432  TAACAAG  32512426
```

Range 468: 32512942 to 32512948

| Score | Expect | Identities | Gaps | Strand | Frame |
|---|---|---|---|---|---|
| 14.4 bits(7) | 223() | 7/7(100%) | 0/7(0%) | Plus/Minus | |

Features:

```
Query  273       TAACAAG  279
                 |||||||
Sbjct  32512948  TAACAAG  32512942
```

Range 469: 32513054 to 32513060

| Score | Expect | Identities | Gaps | Strand | Frame |
|---|---|---|---|---|---|
| 14.4 bits(7) | 223() | 7/7(100%) | 0/7(0%) | Plus/Plus | |

Features:

```
Query  289       TCTCCTT  295
                 |||||||
Sbjct  32513054  TCTCCTT  32513060
```

Range 470: 32513628 to 32513634

| Score | Expect | Identities | Gaps | Strand | Frame |
|---|---|---|---|---|---|
| 14.4 bits(7) | 223() | 7/7(100%) | 0/7(0%) | Plus/Plus | |

Features:

```
Query  277       AAGCTGG  283
                 |||||||
Sbjct  32513628  AAGCTGG  32513634
```

Range 471: 32514495 to 32514501

| Score | Expect | Identities | Gaps | Strand | Frame |
|---|---|---|---|---|---|
| 14.4 bits(7) | 223() | 7/7(100%) | 0/7(0%) | Plus/Plus | |

Features:

```
Query  271       TGTAACA  277
                 |||||||
Sbjct  32514495  TGTAACA  32514501
```

Range 472: 32514881 to 32514887

| Score | Expect | Identities | Gaps | Strand | Frame |
|---|---|---|---|---|---|
| 14.4 bits(7) | 223() | 7/7(100%) | 0/7(0%) | Plus/Minus | |

Features:

```
Query  293       CTTTATT  299
                 |||||||
Sbjct  32514887  CTTTATT  32514881
```

Range 473: 32515829 to 32515835

| Score | Expect | Identities | Gaps | Strand | Frame |
|---|---|---|---|---|---|
| 14.4 bits(7) | 223() | 7/7(100%) | 0/7(0%) | Plus/Minus | |

Features:

```
Query  292       CCTTTAT  298
                 |||||||
Sbjct  32515835  CCTTTAT  32515829
```

Range 474: 32516645 to 32516651

| Score | Expect | Identities | Gaps | Strand | Frame |
|---|---|---|---|---|---|
| 14.4 bits(7) | 223() | 7/7(100%) | 0/7(0%) | Plus/Plus | |

Features:

```
Query  277       AAGCTGG  283
                 |||||||
Sbjct  32516645  AAGCTGG  32516651
```

Range 475: 32516876 to 32516882

| Score | Expect | Identities | Gaps | Strand | Frame |
|---|---|---|---|---|---|
| 14.4 bits(7) | 223() | 7/7(100%) | 0/7(0%) | Plus/Plus | |

Features:

```
Query  271       TGTAACA  277
                 |||||||
Sbjct  32516876  TGTAACA  32516882
```

Range 476: 32516934 to 32516940

| Score | Expect | Identities | Gaps | Strand | Frame |
|---|---|---|---|---|---|
| 14.4 bits(7) | 223() | 7/7(100%) | 0/7(0%) | Plus/Plus | |

Features:

```
Query  289       TCTCCTT  295
                 |||||||
Sbjct  32516934  TCTCCTT  32516940
```

Range 477: 32516938 to 32516944

| Score | Expect | Identities | Gaps | Strand | Frame |
|---|---|---|---|---|---|
| 14.4 bits(7) | 223() | 7/7(100%) | 0/7(0%) | Plus/Minus | |

Features:

```
Query  273       TAACAAG  279
                 |||||||
Sbjct  32516944  TAACAAG  32516938
```

Range 478: 32517543 to 32517549

| Score | Expect | Identities | Gaps | Strand | Frame |
|---|---|---|---|---|---|
| 14.4 bits(7) | 223() | 7/7(100%) | 0/7(0%) | Plus/Plus | |

Features:

```
Query  286       TTCTCTC  292
                 |||||||
Sbjct  32517543  TTCTCTC  32517549
```

Range 479: 32517733 to 32517739

| Score | Expect | Identities | Gaps | Strand | Frame |
|---|---|---|---|---|---|
| 14.4 bits(7) | 223() | 7/7(100%) | 0/7(0%) | Plus/Minus | |

Features:

```
Query  277       AAGCTGG  283
                 |||||||
Sbjct  32517739  AAGCTGG  32517733
```

Range 480: 32517872 to 32517878

| Score | Expect | Identities | Gaps | Strand | Frame |
|---|---|---|---|---|---|
| 14.4 bits(7) | 223() | 7/7(100%) | 0/7(0%) | Plus/Minus | |

Features:

```
Query  273       TAACAAG  279
                 |||||||
Sbjct  32517878  TAACAAG  32517872
```

Range 481: 32518516 to 32518522

| Score | Expect | Identities | Gaps | Strand | Frame |
|---|---|---|---|---|---|
| 14.4 bits(7) | 223() | 7/7(100%) | 0/7(0%) | Plus/Plus | |

Features:

```
Query  292       CCTTTAT  298
                 |||||||
Sbjct  32518516  CCTTTAT  32518522
```

Range 482: 32519020 to 32519026

| Score | Expect | Identities | Gaps | Strand | Frame |
|---|---|---|---|---|---|
| 14.4 bits(7) | 223() | 7/7(100%) | 0/7(0%) | Plus/Plus | |

Features:

```
Query  293       CTTTATT  299
                 |||||||
Sbjct  32519020  CTTTATT  32519026
```

Range 483: 32519209 to 32519215

| Score | Expect | Identities | Gaps | Strand | Frame |
|---|---|---|---|---|---|
| 14.4 bits(7) | 223() | 7/7(100%) | 0/7(0%) | Plus/Plus | |

Features:

```
Query  284       TGTTCTC  290
                 |||||||
Sbjct  32519209  TGTTCTC  32519215
```

Range 484: 32520210 to 32520216

| Score | Expect | Identities | Gaps | Strand | Frame |
|---|---|---|---|---|---|
| 14.4 bits(7) | 223() | 7/7(100%) | 0/7(0%) | Plus/Minus | |

Features:

```
Query  272       GTAACAA  278
                 |||||||
Sbjct  32520216  GTAACAA  32520210
```

Range 485: 32520422 to 32520428

| Score | Expect | Identities | Gaps | Strand | Frame |
|---|---|---|---|---|---|
| 14.4 bits(7) | 223() | 7/7(100%) | 0/7(0%) | Plus/Minus | |

Features:

```
Query  271       TGTAACA  277
                 |||||||
Sbjct  32520428  TGTAACA  32520422
```

Range 486: 32520542 to 32520548

| Score | Expect | Identities | Gaps | Strand | Frame |
|---|---|---|---|---|---|

| | | | | |
|---|---|---|---|---|
| 14.4 bits(7) | 223() | 7/7(100%) | 0/7(0%) | Plus/Minus |

Features:

```
Query  274       AACAAGC  280
                 |||||||
Sbjct  32520548  AACAAGC  32520542
```

Range 487: 32521450 to 32521456

| Score | Expect | Identities | Gaps | Strand | Frame |
|---|---|---|---|---|---|
| 14.4 bits(7) | 223() | 7/7(100%) | 0/7(0%) | Plus/Plus | |

Features:

```
Query  286       TTCTCTC  292
                 |||||||
Sbjct  32521450  TTCTCTC  32521456
```

Range 488: 32521804 to 32521810

| Score | Expect | Identities | Gaps | Strand | Frame |
|---|---|---|---|---|---|
| 14.4 bits(7) | 223() | 7/7(100%) | 0/7(0%) | Plus/Plus | |

Features:

```
Query  292       CCTTTAT  298
                 |||||||
Sbjct  32521804  CCTTTAT  32521810
```

Range 489: 32522145 to 32522151

| Score | Expect | Identities | Gaps | Strand | Frame |
|---|---|---|---|---|---|
| 14.4 bits(7) | 223() | 7/7(100%) | 0/7(0%) | Plus/Plus | |

Features:

```
Query  289       TCTCCTT  295
                 |||||||
Sbjct  32522145  TCTCCTT  32522151
```

Range 490: 32522239 to 32522245

| Score | Expect | Identities | Gaps | Strand | Frame |
|---|---|---|---|---|---|
| 14.4 bits(7) | 223() | 7/7(100%) | 0/7(0%) | Plus/Plus | |

Features:

```
Query  284       TGTTCTC  290
                 |||||||
Sbjct  32522239  TGTTCTC  32522245
```

Range 491: 32522250 to 32522256

| Score | Expect | Identities | Gaps | Strand | Frame |
|---|---|---|---|---|---|
| 14.4 bits(7) | 223() | 7/7(100%) | 0/7(0%) | Plus/Plus | |

Features:

```
Query  293       CTTTATT  299
                 |||||||
Sbjct  32522250  CTTTATT  32522256
```

Range 492: 32524357 to 32524363

| Score | Expect | Identities | Gaps | Strand | Frame |
|---|---|---|---|---|---|
| 14.4 bits(7) | 223() | 7/7(100%) | 0/7(0%) | Plus/Minus | |

Features:

```
Query  271       TGTAACA  277
                 |||||||
Sbjct  32524363  TGTAACA  32524357
```

Range 493: 32524577 to 32524583

| Score | Expect | Identities | Gaps | Strand | Frame |
|---|---|---|---|---|---|
| 14.4 bits(7) | 223() | 7/7(100%) | 0/7(0%) | Plus/Minus | |

Features:

```
Query  293       CTTTATT  299
                 |||||||
Sbjct  32524583  CTTTATT  32524577
```

Range 494: 32525791 to 32525797

| Score | Expect | Identities | Gaps | Strand | Frame |
|---|---|---|---|---|---|
| 14.4 bits(7) | 223() | 7/7(100%) | 0/7(0%) | Plus/Minus | |

Features:

```
Query  290       CTCCTTT  296
                 |||||||
Sbjct  32525797  CTCCTTT  32525791
```

Range 495: 32526290 to 32526296

| Score | Expect | Identities | Gaps | Strand | Frame |
|---|---|---|---|---|---|
| 14.4 bits(7) | 223() | 7/7(100%) | 0/7(0%) | Plus/Plus | |

Features:

```
Query  272       GTAACAA  278
                 |||||||
Sbjct  32526290  GTAACAA  32526296
```

Range 496: 32526395 to 32526401

| Score | Expect | Identities | Gaps | Strand | Frame |
|---|---|---|---|---|---|
| 14.4 bits(7) | 223() | 7/7(100%) | 0/7(0%) | Plus/Plus | |

Features:

```
Query  281       TGGTGTT  287
                 |||||||
Sbjct  32526395  TGGTGTT  32526401
```

Range 497: 32526436 to 32526442

| Score | Expect | Identities | Gaps | Strand | Frame |
|---|---|---|---|---|---|
| 14.4 bits(7) | 223() | 7/7(100%) | 0/7(0%) | Plus/Plus | |

Features:

```
Query  272       GTAACAA  278
                 |||||||
Sbjct  32526436  GTAACAA  32526442
```

Range 498: 32526545 to 32526551

| Score | Expect | Identities | Gaps | Strand | Frame |
|---|---|---|---|---|---|
| 14.4 bits(7) | 223() | 7/7(100%) | 0/7(0%) | Plus/Plus | |

Features:

```
Query  281       TGGTGTT  287
                 |||||||
Sbjct  32526545  TGGTGTT  32526551
```

Range 499: 32526789 to 32526795

| Score | Expect | Identities | Gaps | Strand | Frame |
|---|---|---|---|---|---|
| 14.4 bits(7) | 223() | 7/7(100%) | 0/7(0%) | Plus/Minus | |

Features:

```
Query  290       CTCCTTT  296
                 |||||||
Sbjct  32526795  CTCCTTT  32526789
```

Range 500: 32527507 to 32527513

| Score | Expect | Identities | Gaps | Strand | Frame |
|---|---|---|---|---|---|
| 14.4 bits(7) | 223() | 7/7(100%) | 0/7(0%) | Plus/Minus | |

Features:

```
Query  290       CTCCTTT  296
                 |||||||
Sbjct  32527513  CTCCTTT  32527507
```

Range 501: 32527734 to 32527740

| Score | Expect | Identities | Gaps | Strand | Frame |
|---|---|---|---|---|---|
| 14.4 bits(7) | 223() | 7/7(100%) | 0/7(0%) | Plus/Plus | |

Features:

```
Query  277       AAGCTGG  283
                 |||||||
Sbjct  32527734  AAGCTGG  32527740
```

Range 502: 32527869 to 32527875

| Score | Expect | Identities | Gaps | Strand | Frame |
|---|---|---|---|---|---|
| 14.4 bits(7) | 223() | 7/7(100%) | 0/7(0%) | Plus/Minus | |

Features:

```
Query  281       TGGTGTT  287
                 |||||||
Sbjct  32527875  TGGTGTT  32527869
```

Range 503: 32528782 to 32528788

| Score | Expect | Identities | Gaps | Strand | Frame |
|---|---|---|---|---|---|
| 14.4 bits(7) | 223() | 7/7(100%) | 0/7(0%) | Plus/Plus | |

Features:

```
Query  293       CTTTATT  299
                 |||||||
Sbjct  32528782  CTTTATT  32528788
```

Range 504: 32529227 to 32529233

| Score | Expect | Identities | Gaps | Strand | Frame |
|---|---|---|---|---|---|
| 14.4 bits(7) | 223() | 7/7(100%) | 0/7(0%) | Plus/Minus | |

Features:

```
Query  293       CTTTATT  299
                 |||||||
Sbjct  32529233  CTTTATT  32529227
```

Range 505: 32529312 to 32529318

| Score | Expect | Identities | Gaps | Strand | Frame |
|---|---|---|---|---|---|

| Score | Expect | Identities | Gaps | Strand | Frame |
|---|---|---|---|---|---|
| 14.4 bits(7) | 223() | 7/7(100%) | 0/7(0%) | Plus/Plus | |

Features:

```
Query  271       TGTAACA  277
                 |||||||
Sbjct  32529312  TGTAACA  32529318
```

Range 506: 32529682 to 32529688

| Score | Expect | Identities | Gaps | Strand | Frame |
|---|---|---|---|---|---|
| 14.4 bits(7) | 223() | 7/7(100%) | 0/7(0%) | Plus/Minus | |

Features:

```
Query  277       AAGCTGG  283
                 |||||||
Sbjct  32529688  AAGCTGG  32529682
```

Range 507: 32529693 to 32529699

| Score | Expect | Identities | Gaps | Strand | Frame |
|---|---|---|---|---|---|
| 14.4 bits(7) | 223() | 7/7(100%) | 0/7(0%) | Plus/Plus | |

Features:

```
Query  288       CTCTCCT  294
                 |||||||
Sbjct  32529693  CTCTCCT  32529699
```

Range 508: 32530339 to 32530345

| Score | Expect | Identities | Gaps | Strand | Frame |
|---|---|---|---|---|---|
| 14.4 bits(7) | 223() | 7/7(100%) | 0/7(0%) | Plus/Plus | |

Features:

```
Query  277       AAGCTGG  283
                 |||||||
Sbjct  32530339  AAGCTGG  32530345
```

Range 509: 32530395 to 32530401

| Score | Expect | Identities | Gaps | Strand | Frame |
|---|---|---|---|---|---|
| 14.4 bits(7) | 223() | 7/7(100%) | 0/7(0%) | Plus/Minus | |

Features:

```
Query  286       TTCTCTC  292
                 |||||||
Sbjct  32530401  TTCTCTC  32530395
```

Range 510: 32530796 to 32530802

| Score | Expect | Identities | Gaps | Strand | Frame |
|---|---|---|---|---|---|
| 14.4 bits(7) | 223() | 7/7(100%) | 0/7(0%) | Plus/Plus | |

Features:

```
Query  286       TTCTCTC  292
                 |||||||
Sbjct  32530796  TTCTCTC  32530802
```

Range 511: 32531019 to 32531025

| Score | Expect | Identities | Gaps | Strand | Frame |
|---|---|---|---|---|---|
| 14.4 bits(7) | 223() | 7/7(100%) | 0/7(0%) | Plus/Minus | |

Features:

```
Query  291       TCCTTTA  297
                 |||||||
Sbjct  32531025  TCCTTTA  32531019
```

Range 512: 32531151 to 32531157

| Score | Expect | Identities | Gaps | Strand | Frame |
|---|---|---|---|---|---|
| 14.4 bits(7) | 223() | 7/7(100%) | 0/7(0%) | Plus/Plus | |

Features:

```
Query  273       TAACAAG  279
                 |||||||
Sbjct  32531151  TAACAAG  32531157
```

Range 513: 32531807 to 32531813

| Score | Expect | Identities | Gaps | Strand | Frame |
|---|---|---|---|---|---|
| 14.4 bits(7) | 223() | 7/7(100%) | 0/7(0%) | Plus/Plus | |

Features:

```
Query  271       TGTAACA  277
                 |||||||
Sbjct  32531807  TGTAACA  32531813
```

Range 514: 32532485 to 32532491

| Score | Expect | Identities | Gaps | Strand | Frame |
|---|---|---|---|---|---|
| 14.4 bits(7) | 223() | 7/7(100%) | 0/7(0%) | Plus/Plus | |

Features:

```
Query  289       TCTCCTT  295
                 |||||||
Sbjct  32532485  TCTCCTT  32532491
```

Range 515: 32532498 to 32532504

| Score | Expect | Identities | Gaps | Strand | Frame |
|---|---|---|---|---|---|
| 14.4 bits(7) | 223() | 7/7(100%) | 0/7(0%) | Plus/Minus | |

Features:

```
Query  280       CTGGTGT  286
                 |||||||
Sbjct  32532504  CTGGTGT  32532498
```

Range 516: 32533431 to 32533437

| Score | Expect | Identities | Gaps | Strand | Frame |
|---|---|---|---|---|---|
| 14.4 bits(7) | 223() | 7/7(100%) | 0/7(0%) | Plus/Minus | |

Features:

```
Query  286       TTCTCTC  292
                 |||||||
Sbjct  32533437  TTCTCTC  32533431
```

Range 517: 32533450 to 32533456

| Score | Expect | Identities | Gaps | Strand | Frame |
|---|---|---|---|---|---|
| 14.4 bits(7) | 223() | 7/7(100%) | 0/7(0%) | Plus/Plus | |

Features:

```
Query  274       AACAAGC  280
                 |||||||
Sbjct  32533450  AACAAGC  32533456
```

Range 518: 32533755 to 32533761

| Score | Expect | Identities | Gaps | Strand | Frame |
|---|---|---|---|---|---|
| 14.4 bits(7) | 223() | 7/7(100%) | 0/7(0%) | Plus/Plus | |

Features:

```
Query  277       AAGCTGG  283
                 |||||||
Sbjct  32533755  AAGCTGG  32533761
```

Range 519: 32534636 to 32534642

| Score | Expect | Identities | Gaps | Strand | Frame |
|---|---|---|---|---|---|
| 14.4 bits(7) | 223() | 7/7(100%) | 0/7(0%) | Plus/Minus | |

Features:

```
Query  289       TCTCCTT  295
                 |||||||
Sbjct  32534642  TCTCCTT  32534636
```

Range 520: 32535123 to 32535129

| Score | Expect | Identities | Gaps | Strand | Frame |
|---|---|---|---|---|---|
| 14.4 bits(7) | 223() | 7/7(100%) | 0/7(0%) | Plus/Plus | |

Features:

```
Query  273       TAACAAG  279
                 |||||||
Sbjct  32535123  TAACAAG  32535129
```

Range 521: 32535188 to 32535194

| Score | Expect | Identities | Gaps | Strand | Frame |
|---|---|---|---|---|---|
| 14.4 bits(7) | 223() | 7/7(100%) | 0/7(0%) | Plus/Minus | |

Features:

```
Query  286       TTCTCTC  292
                 |||||||
Sbjct  32535194  TTCTCTC  32535188
```

Range 522: 32535202 to 32535208

| Score | Expect | Identities | Gaps | Strand | Frame |
|---|---|---|---|---|---|
| 14.4 bits(7) | 223() | 7/7(100%) | 0/7(0%) | Plus/Minus | |

Features:

```
Query  278       AGCTGGT  284
                 |||||||
Sbjct  32535208  AGCTGGT  32535202
```

Range 523: 32535299 to 32535305

| Score | Expect | Identities | Gaps | Strand | Frame |
|---|---|---|---|---|---|
| 14.4 bits(7) | 223() | 7/7(100%) | 0/7(0%) | Plus/Plus | |

Features:

```
Query  293       CTTTATT  299
                 |||||||
Sbjct  32535299  CTTTATT  32535305
```

Range 524: 32535541 to 32535547

| Score | Expect | Identities | Gaps | Strand | Frame |
|---|---|---|---|---|---|

| Score | Expect | Identities | Gaps | Strand | Frame |
|---|---|---|---|---|---|
| 14.4 bits(7) | 223() | 7/7(100%) | 0/7(0%) | Plus/Minus | |

Features:

```
Query  284       TGTTCTC  290
                 |||||||
Sbjct  32535547  TGTTCTC  32535541
```

Range 525: 32535600 to 32535606

| Score | Expect | Identities | Gaps | Strand | Frame |
|---|---|---|---|---|---|
| 14.4 bits(7) | 223() | 7/7(100%) | 0/7(0%) | Plus/Minus | |

Features:

```
Query  277       AAGCTGG  283
                 |||||||
Sbjct  32535606  AAGCTGG  32535600
```

Range 526: 32536354 to 32536360

| Score | Expect | Identities | Gaps | Strand | Frame |
|---|---|---|---|---|---|
| 14.4 bits(7) | 223() | 7/7(100%) | 0/7(0%) | Plus/Plus | |

Features:

```
Query  281       TGGTGTT  287
                 |||||||
Sbjct  32536354  TGGTGTT  32536360
```

Range 527: 32536747 to 32536753

| Score | Expect | Identities | Gaps | Strand | Frame |
|---|---|---|---|---|---|
| 14.4 bits(7) | 223() | 7/7(100%) | 0/7(0%) | Plus/Minus | |

Features:

```
Query  277       AAGCTGG  283
                 |||||||
Sbjct  32536753  AAGCTGG  32536747
```

Range 528: 32536955 to 32536961

| Score | Expect | Identities | Gaps | Strand | Frame |
|---|---|---|---|---|---|
| 14.4 bits(7) | 223() | 7/7(100%) | 0/7(0%) | Plus/Plus | |

Features:

```
Query  289       TCTCCTT  295
                 |||||||
Sbjct  32536955  TCTCCTT  32536961
```

Range 529: 32537071 to 32537077

| Score | Expect | Identities | Gaps | Strand | Frame |
|---|---|---|---|---|---|
| 14.4 bits(7) | 223() | 7/7(100%) | 0/7(0%) | Plus/Plus | |

Features:

```
Query  280       CTGGTGT  286
                 |||||||
Sbjct  32537071  CTGGTGT  32537077
```

Range 530: 32537146 to 32537152

| Score | Expect | Identities | Gaps | Strand | Frame |
|---|---|---|---|---|---|
| 14.4 bits(7) | 223() | 7/7(100%) | 0/7(0%) | Plus/Minus | |

Features:

```
Query  289       TCTCCTT  295
                 |||||||
Sbjct  32537152  TCTCCTT  32537146
```

Range 531: 32537192 to 32537198

| Score | Expect | Identities | Gaps | Strand | Frame |
|---|---|---|---|---|---|
| 14.4 bits(7) | 223() | 7/7(100%) | 0/7(0%) | Plus/Plus | |

Features:

```
Query  271       TGTAACA  277
                 |||||||
Sbjct  32537192  TGTAACA  32537198
```

Range 532: 32537239 to 32537245

| Score | Expect | Identities | Gaps | Strand | Frame |
|---|---|---|---|---|---|
| 14.4 bits(7) | 223() | 7/7(100%) | 0/7(0%) | Plus/Plus | |

Features:

```
Query  273       TAACAAG  279
                 |||||||
Sbjct  32537239  TAACAAG  32537245
```

Range 533: 32537318 to 32537324

| Score | Expect | Identities | Gaps | Strand | Frame |
|---|---|---|---|---|---|
| 14.4 bits(7) | 223() | 7/7(100%) | 0/7(0%) | Plus/Minus | |

Features:

```
Query  281       TGGTGTT  287
                 |||||||
Sbjct  32537324  TGGTGTT  32537318
```

Range 534: 32537486 to 32537492

| Score | Expect | Identities | Gaps | Strand | Frame |
|---|---|---|---|---|---|
| 14.4 bits(7) | 223() | 7/7(100%) | 0/7(0%) | Plus/Plus | |

Features:

```
Query  274       AACAAGC  280
                 |||||||
Sbjct  32537486  AACAAGC  32537492
```

Range 535: 32538215 to 32538221

| Score | Expect | Identities | Gaps | Strand | Frame |
|---|---|---|---|---|---|
| 14.4 bits(7) | 223() | 7/7(100%) | 0/7(0%) | Plus/Minus | |

Features:

```
Query  290       CTCCTTT  296
                 |||||||
Sbjct  32538221  CTCCTTT  32538215
```

Range 536: 32538463 to 32538469

| Score | Expect | Identities | Gaps | Strand | Frame |
|---|---|---|---|---|---|
| 14.4 bits(7) | 223() | 7/7(100%) | 0/7(0%) | Plus/Minus | |

Features:

```
Query  293       CTTTATT  299
                 |||||||
Sbjct  32538469  CTTTATT  32538463
```

Range 537: 32539112 to 32539118

| Score | Expect | Identities | Gaps | Strand | Frame |
|---|---|---|---|---|---|
| 14.4 bits(7) | 223() | 7/7(100%) | 0/7(0%) | Plus/Minus | |

Features:

```
Query  293       CTTTATT  299
                 |||||||
Sbjct  32539118  CTTTATT  32539112
```

Range 538: 32541000 to 32541006

| Score | Expect | Identities | Gaps | Strand | Frame |
|---|---|---|---|---|---|
| 14.4 bits(7) | 223() | 7/7(100%) | 0/7(0%) | Plus/Minus | |

Features:

```
Query  273       TAACAAG  279
                 |||||||
Sbjct  32541006  TAACAAG  32541000
```

Range 539: 32541102 to 32541108

| Score | Expect | Identities | Gaps | Strand | Frame |
|---|---|---|---|---|---|
| 14.4 bits(7) | 223() | 7/7(100%) | 0/7(0%) | Plus/Plus | |

Features:

```
Query  291       TCCTTTA  297
                 |||||||
Sbjct  32541102  TCCTTTA  32541108
```

Range 540: 32541237 to 32541243

| Score | Expect | Identities | Gaps | Strand | Frame |
|---|---|---|---|---|---|
| 14.4 bits(7) | 223() | 7/7(100%) | 0/7(0%) | Plus/Minus | |

Features:

```
Query  292       CCTTTAT  298
                 |||||||
Sbjct  32541243  CCTTTAT  32541237
```

Range 541: 32541346 to 32541352

| Score | Expect | Identities | Gaps | Strand | Frame |
|---|---|---|---|---|---|
| 14.4 bits(7) | 223() | 7/7(100%) | 0/7(0%) | Plus/Plus | |

Features:

```
Query  277       AAGCTGG  283
                 |||||||
Sbjct  32541346  AAGCTGG  32541352
```

Range 542: 32542050 to 32542056

| Score | Expect | Identities | Gaps | Strand | Frame |
|---|---|---|---|---|---|
| 14.4 bits(7) | 223() | 7/7(100%) | 0/7(0%) | Plus/Plus | |

Features:

```
Query  293       CTTTATT  299
                 |||||||
Sbjct  32542050  CTTTATT  32542056
```

Range 543: 32542254 to 32542260

| Score | Expect | Identities | Gaps | Strand | Frame |
|---|---|---|---|---|---|

| Score | Expect | Identities | Gaps | Strand | Frame |
|---|---|---|---|---|---|
| 14.4 bits(7) | 223() | 7/7(100%) | 0/7(0%) | Plus/Minus | |

Features:

```
Query  291       TCCTTTA  297
                 |||||||
Sbjct  32542260  TCCTTTA  32542254
```

Range 544: 32542468 to 32542474

| Score | Expect | Identities | Gaps | Strand | Frame |
|---|---|---|---|---|---|
| 14.4 bits(7) | 223() | 7/7(100%) | 0/7(0%) | Plus/Minus | |

Features:

```
Query  293       CTTTATT  299
                 |||||||
Sbjct  32542474  CTTTATT  32542468
```

Range 545: 32542712 to 32542718

| Score | Expect | Identities | Gaps | Strand | Frame |
|---|---|---|---|---|---|
| 14.4 bits(7) | 223() | 7/7(100%) | 0/7(0%) | Plus/Minus | |

Features:

```
Query  292       CCTTTAT  298
                 |||||||
Sbjct  32542718  CCTTTAT  32542712
```

Range 546: 32543126 to 32543132

| Score | Expect | Identities | Gaps | Strand | Frame |
|---|---|---|---|---|---|
| 14.4 bits(7) | 223() | 7/7(100%) | 0/7(0%) | Plus/Plus | |

Features:

```
Query  293       CTTTATT  299
                 |||||||
Sbjct  32543126  CTTTATT  32543132
```

Range 547: 32543255 to 32543261

| Score | Expect | Identities | Gaps | Strand | Frame |
|---|---|---|---|---|---|
| 14.4 bits(7) | 223() | 7/7(100%) | 0/7(0%) | Plus/Plus | |

Features:

```
Query  271       TGTAACA  277
                 |||||||
Sbjct  32543255  TGTAACA  32543261
```

Range 548: 32543478 to 32543484

| Score | Expect | Identities | Gaps | Strand | Frame |
|---|---|---|---|---|---|
| 14.4 bits(7) | 223() | 7/7(100%) | 0/7(0%) | Plus/Minus | |

Features:

```
Query  286       TTCTCTC  292
                 |||||||
Sbjct  32543484  TTCTCTC  32543478
```

Range 549: 32543832 to 32543838

| Score | Expect | Identities | Gaps | Strand | Frame |
|---|---|---|---|---|---|
| 14.4 bits(7) | 223() | 7/7(100%) | 0/7(0%) | Plus/Plus | |

Features:

```
Query  284       TGTTCTC  290
                 |||||||
Sbjct  32543832  TGTTCTC  32543838
```

Range 550: 32545210 to 32545216

| Score | Expect | Identities | Gaps | Strand | Frame |
|---|---|---|---|---|---|
| 14.4 bits(7) | 223() | 7/7(100%) | 0/7(0%) | Plus/Minus | |

Features:

```
Query  283       GTGTTCT  289
                 |||||||
Sbjct  32545216  GTGTTCT  32545210
```

Range 551: 32545220 to 32545226

| Score | Expect | Identities | Gaps | Strand | Frame |
|---|---|---|---|---|---|
| 14.4 bits(7) | 223() | 7/7(100%) | 0/7(0%) | Plus/Minus | |

Features:

```
Query  283       GTGTTCT  289
                 |||||||
Sbjct  32545226  GTGTTCT  32545220
```

Range 552: 32545921 to 32545927

| Score | Expect | Identities | Gaps | Strand | Frame |
|---|---|---|---|---|---|
| 14.4 bits(7) | 223() | 7/7(100%) | 0/7(0%) | Plus/Minus | |

Features:

```
Query  293       CTTTATT  299
                 |||||||
Sbjct  32545927  CTTTATT  32545921
```

Range 553: 32545959 to 32545965

| Score | Expect | Identities | Gaps | Strand | Frame |
|---|---|---|---|---|---|
| 14.4 bits(7) | 223() | 7/7(100%) | 0/7(0%) | Plus/Plus | |

Features:

```
Query  282       GGTGTTC  288
                 |||||||
Sbjct  32545959  GGTGTTC  32545965
```

Range 554: 32547959 to 32547965

| Score | Expect | Identities | Gaps | Strand | Frame |
|---|---|---|---|---|---|
| 14.4 bits(7) | 223() | 7/7(100%) | 0/7(0%) | Plus/Plus | |

Features:

```
Query  274       AACAAGC  280
                 |||||||
Sbjct  32547959  AACAAGC  32547965
```

Range 555: 32548988 to 32548994

| Score | Expect | Identities | Gaps | Strand | Frame |
|---|---|---|---|---|---|
| 14.4 bits(7) | 223() | 7/7(100%) | 0/7(0%) | Plus/Minus | |

Features:

```
Query  272       GTAACAA  278
                 |||||||
Sbjct  32548994  GTAACAA  32548988
```

Range 556: 32549658 to 32549664

| Score | Expect | Identities | Gaps | Strand | Frame |
|---|---|---|---|---|---|
| 14.4 bits(7) | 223() | 7/7(100%) | 0/7(0%) | Plus/Minus | |

Features:

```
Query  283       GTGTTCT  289
                 |||||||
Sbjct  32549664  GTGTTCT  32549658
```

Range 557: 32550038 to 32550044

| Score | Expect | Identities | Gaps | Strand | Frame |
|---|---|---|---|---|---|
| 14.4 bits(7) | 223() | 7/7(100%) | 0/7(0%) | Plus/Plus | |

Features:

```
Query  283       GTGTTCT  289
                 |||||||
Sbjct  32550038  GTGTTCT  32550044
```

Range 558: 32550099 to 32550105

| Score | Expect | Identities | Gaps | Strand | Frame |
|---|---|---|---|---|---|
| 14.4 bits(7) | 223() | 7/7(100%) | 0/7(0%) | Plus/Plus | |

Features:

```
Query  274       AACAAGC  280
                 |||||||
Sbjct  32550099  AACAAGC  32550105
```

Range 559: 32550383 to 32550389

| Score | Expect | Identities | Gaps | Strand | Frame |
|---|---|---|---|---|---|
| 14.4 bits(7) | 223() | 7/7(100%) | 0/7(0%) | Plus/Plus | |

Features:

```
Query  291       TCCTTTA  297
                 |||||||
Sbjct  32550383  TCCTTTA  32550389
```

Range 560: 32551046 to 32551052

| Score | Expect | Identities | Gaps | Strand | Frame |
|---|---|---|---|---|---|
| 14.4 bits(7) | 223() | 7/7(100%) | 0/7(0%) | Plus/Plus | |

Features:

```
Query  272       GTAACAA  278
                 |||||||
Sbjct  32551046  GTAACAA  32551052
```

Range 561: 32552343 to 32552349

| Score | Expect | Identities | Gaps | Strand | Frame |
|---|---|---|---|---|---|
| 14.4 bits(7) | 223() | 7/7(100%) | 0/7(0%) | Plus/Plus | |

Features:

```
Query  278       AGCTGGT  284
                 |||||||
Sbjct  32552343  AGCTGGT  32552349
```

Range 562: 32553863 to 32553869

| Score | Expect | Identities | Gaps | Strand | Frame |
|---|---|---|---|---|---|

| | | | | |
|---|---|---|---|---|
| 14.4 bits(7) | 223() | 7/7(100%) | 0/7(0%) | Plus/Minus |

Features:

```
Query  293       CTTTATT  299
                 |||||||
Sbjct  32553869  CTTTATT  32553863
```

Range 563: 32553880 to 32553886

| Score | Expect | Identities | Gaps | Strand | Frame |
|---|---|---|---|---|---|
| 14.4 bits(7) | 223() | 7/7(100%) | 0/7(0%) | Plus/Plus | |

Features:

```
Query  283       GTGTTCT  289
                 |||||||
Sbjct  32553880  GTGTTCT  32553886
```

Range 564: 32555063 to 32555069

| Score | Expect | Identities | Gaps | Strand | Frame |
|---|---|---|---|---|---|
| 14.4 bits(7) | 223() | 7/7(100%) | 0/7(0%) | Plus/Plus | |

Features:

```
Query  293       CTTTATT  299
                 |||||||
Sbjct  32555063  CTTTATT  32555069
```

Range 565: 32555217 to 32555223

| Score | Expect | Identities | Gaps | Strand | Frame |
|---|---|---|---|---|---|
| 14.4 bits(7) | 223() | 7/7(100%) | 0/7(0%) | Plus/Minus | |

Features:

```
Query  293       CTTTATT  299
                 |||||||
Sbjct  32555223  CTTTATT  32555217
```

Range 566: 32555750 to 32555756

| Score | Expect | Identities | Gaps | Strand | Frame |
|---|---|---|---|---|---|
| 14.4 bits(7) | 223() | 7/7(100%) | 0/7(0%) | Plus/Minus | |

Features:

```
Query  279       GCTGGTG  285
                 |||||||
Sbjct  32555756  GCTGGTG  32555750
```

Range 567: 32555951 to 32555957

| Score | Expect | Identities | Gaps | Strand | Frame |
|---|---|---|---|---|---|
| 14.4 bits(7) | 223() | 7/7(100%) | 0/7(0%) | Plus/Plus | |

Features:

```
Query  293       CTTTATT  299
                 |||||||
Sbjct  32555951  CTTTATT  32555957
```

Range 568: 32557090 to 32557096

| Score | Expect | Identities | Gaps | Strand | Frame |
|---|---|---|---|---|---|
| 14.4 bits(7) | 223() | 7/7(100%) | 0/7(0%) | Plus/Plus | |

Features:

```
Query  293       CTTTATT  299
                 |||||||
Sbjct  32557090  CTTTATT  32557096
```

Range 569: 32557631 to 32557637

| Score | Expect | Identities | Gaps | Strand | Frame |
|---|---|---|---|---|---|
| 14.4 bits(7) | 223() | 7/7(100%) | 0/7(0%) | Plus/Minus | |

Features:

```
Query  291       TCCTTTA  297
                 |||||||
Sbjct  32557637  TCCTTTA  32557631
```

Range 570: 32557654 to 32557660

| Score | Expect | Identities | Gaps | Strand | Frame |
|---|---|---|---|---|---|
| 14.4 bits(7) | 223() | 7/7(100%) | 0/7(0%) | Plus/Minus | |

Features:

```
Query  273       TAACAAG  279
                 |||||||
Sbjct  32557660  TAACAAG  32557654
```

Range 571: 32558372 to 32558378

| Score | Expect | Identities | Gaps | Strand | Frame |
|---|---|---|---|---|---|
| 14.4 bits(7) | 223() | 7/7(100%) | 0/7(0%) | Plus/Plus | |

Features:

```
Query  290       CTCCTTT  296
                 |||||||
Sbjct  32558372  CTCCTTT  32558378
```

Range 572: 32559287 to 32559293

| Score | Expect | Identities | Gaps | Strand | Frame |
|---|---|---|---|---|---|
| 14.4 bits(7) | 223() | 7/7(100%) | 0/7(0%) | Plus/Plus | |

Features:

```
Query  271       TGTAACA  277
                 |||||||
Sbjct  32559287  TGTAACA  32559293
```

Range 573: 32559336 to 32559342

| Score | Expect | Identities | Gaps | Strand | Frame |
|---|---|---|---|---|---|
| 14.4 bits(7) | 223() | 7/7(100%) | 0/7(0%) | Plus/Plus | |

Features:

```
Query  276       CAAGCTG  282
                 |||||||
Sbjct  32559336  CAAGCTG  32559342
```

Range 574: 32559355 to 32559361

| Score | Expect | Identities | Gaps | Strand | Frame |
|---|---|---|---|---|---|
| 14.4 bits(7) | 223() | 7/7(100%) | 0/7(0%) | Plus/Plus | |

Features:

```
Query  281       TGGTGTT  287
                 |||||||
Sbjct  32559355  TGGTGTT  32559361
```

Range 575: 32560443 to 32560449

| Score | Expect | Identities | Gaps | Strand | Frame |
|---|---|---|---|---|---|
| 14.4 bits(7) | 223() | 7/7(100%) | 0/7(0%) | Plus/Plus | |

Features:

```
Query  288       CTCTCCT  294
                 |||||||
Sbjct  32560443  CTCTCCT  32560449
```

Range 576: 32563170 to 32563176

| Score | Expect | Identities | Gaps | Strand | Frame |
|---|---|---|---|---|---|
| 14.4 bits(7) | 223() | 7/7(100%) | 0/7(0%) | Plus/Minus | |

Features:

```
Query  271       TGTAACA  277
                 |||||||
Sbjct  32563176  TGTAACA  32563170
```

Range 577: 32563593 to 32563599

| Score | Expect | Identities | Gaps | Strand | Frame |
|---|---|---|---|---|---|
| 14.4 bits(7) | 223() | 7/7(100%) | 0/7(0%) | Plus/Minus | |

Features:

```
Query  290       CTCCTTT  296
                 |||||||
Sbjct  32563599  CTCCTTT  32563593
```

Range 578: 32564446 to 32564452

| Score | Expect | Identities | Gaps | Strand | Frame |
|---|---|---|---|---|---|
| 14.4 bits(7) | 223() | 7/7(100%) | 0/7(0%) | Plus/Plus | |

Features:

```
Query  281       TGGTGTT  287
                 |||||||
Sbjct  32564446  TGGTGTT  32564452
```

Range 579: 32564874 to 32564880

| Score | Expect | Identities | Gaps | Strand | Frame |
|---|---|---|---|---|---|
| 14.4 bits(7) | 223() | 7/7(100%) | 0/7(0%) | Plus/Plus | |

Features:

```
Query  283       GTGTTCT  289
                 |||||||
Sbjct  32564874  GTGTTCT  32564880
```

Range 580: 32564936 to 32564942

| Score | Expect | Identities | Gaps | Strand | Frame |
|---|---|---|---|---|---|
| 14.4 bits(7) | 223() | 7/7(100%) | 0/7(0%) | Plus/Minus | |

Features:

```
Query  290       CTCCTTT  296
                 |||||||
Sbjct  32564942  CTCCTTT  32564936
```

Range 581: 32565068 to 32565074

| Score | Expect | Identities | Gaps | Strand | Frame |
|---|---|---|---|---|---|

| Score | Expect | Identities | Gaps | Strand | Frame |
|---|---|---|---|---|---|
| 14.4 bits(7) | 223() | 7/7(100%) | 0/7(0%) | Plus/Plus | |

Features:

```
Query  283       GTGTTCT  289
                 |||||||
Sbjct  32565068  GTGTTCT  32565074
```

Range 582: 32565414 to 32565420

| Score | Expect | Identities | Gaps | Strand | Frame |
|---|---|---|---|---|---|
| 14.4 bits(7) | 223() | 7/7(100%) | 0/7(0%) | Plus/Plus | |

Features:

```
Query  284       TGTTCTC  290
                 |||||||
Sbjct  32565414  TGTTCTC  32565420
```

Range 583: 32565619 to 32565625

| Score | Expect | Identities | Gaps | Strand | Frame |
|---|---|---|---|---|---|
| 14.4 bits(7) | 223() | 7/7(100%) | 0/7(0%) | Plus/Plus | |

Features:

```
Query  273       TAACAAG  279
                 |||||||
Sbjct  32565619  TAACAAG  32565625
```

Range 584: 32569794 to 32569800

| Score | Expect | Identities | Gaps | Strand | Frame |
|---|---|---|---|---|---|
| 14.4 bits(7) | 223() | 7/7(100%) | 0/7(0%) | Plus/Plus | |

Features:

```
Query  293       CTTTATT  299
                 |||||||
Sbjct  32569794  CTTTATT  32569800
```

Range 585: 32571179 to 32571185

| Score | Expect | Identities | Gaps | Strand | Frame |
|---|---|---|---|---|---|
| 14.4 bits(7) | 223() | 7/7(100%) | 0/7(0%) | Plus/Plus | |

Features:

```
Query  293       CTTTATT  299
                 |||||||
Sbjct  32571179  CTTTATT  32571185
```

Range 586: 32571273 to 32571279

| Score | Expect | Identities | Gaps | Strand | Frame |
|---|---|---|---|---|---|
| 14.4 bits(7) | 223() | 7/7(100%) | 0/7(0%) | Plus/Minus | |

Features:

```
Query  286       TTCTCTC  292
                 |||||||
Sbjct  32571279  TTCTCTC  32571273
```

Range 587: 32571633 to 32571639

| Score | Expect | Identities | Gaps | Strand | Frame |
|---|---|---|---|---|---|
| 14.4 bits(7) | 223() | 7/7(100%) | 0/7(0%) | Plus/Plus | |

Features:

```
Query  286       TTCTCTC  292
                 |||||||
Sbjct  32571633  TTCTCTC  32571639
```

Range 588: 32571830 to 32571836

| Score | Expect | Identities | Gaps | Strand | Frame |
|---|---|---|---|---|---|
| 14.4 bits(7) | 223() | 7/7(100%) | 0/7(0%) | Plus/Minus | |

Features:

```
Query  272       GTAACAA  278
                 |||||||
Sbjct  32571836  GTAACAA  32571830
```

Range 589: 32571871 to 32571877

| Score | Expect | Identities | Gaps | Strand | Frame |
|---|---|---|---|---|---|
| 14.4 bits(7) | 223() | 7/7(100%) | 0/7(0%) | Plus/Plus | |

Features:

```
Query  286       TTCTCTC  292
                 |||||||
Sbjct  32571871  TTCTCTC  32571877
```

Range 590: 32572061 to 32572067

| Score | Expect | Identities | Gaps | Strand | Frame |
|---|---|---|---|---|---|
| 14.4 bits(7) | 223() | 7/7(100%) | 0/7(0%) | Plus/Plus | |

Features:

```
Query  293       CTTTATT  299
                 |||||||
Sbjct  32572061  CTTTATT  32572067
```

Range 591: 32572127 to 32572133

| Score | Expect | Identities | Gaps | Strand | Frame |
|---|---|---|---|---|---|
| 14.4 bits(7) | 223() | 7/7(100%) | 0/7(0%) | Plus/Plus | |

Features:

```
Query  290       CTCCTTT  296
                 |||||||
Sbjct  32572127  CTCCTTT  32572133
```

Range 592: 32572392 to 32572398

| Score | Expect | Identities | Gaps | Strand | Frame |
|---|---|---|---|---|---|
| 14.4 bits(7) | 223() | 7/7(100%) | 0/7(0%) | Plus/Minus | |

Features:

```
Query  277       AAGCTGG  283
                 |||||||
Sbjct  32572398  AAGCTGG  32572392
```

Range 593: 32572955 to 32572961

| Score | Expect | Identities | Gaps | Strand | Frame |
|---|---|---|---|---|---|
| 14.4 bits(7) | 223() | 7/7(100%) | 0/7(0%) | Plus/Minus | |

Features:

```
Query  271       TGTAACA  277
                 |||||||
Sbjct  32572961  TGTAACA  32572955
```

Range 594: 32573908 to 32573914

| Score | Expect | Identities | Gaps | Strand | Frame |
|---|---|---|---|---|---|
| 14.4 bits(7) | 223() | 7/7(100%) | 0/7(0%) | Plus/Plus | |

Features:

```
Query  280       CTGGTGT  286
                 |||||||
Sbjct  32573908  CTGGTGT  32573914
```

Range 595: 32573964 to 32573970

| Score | Expect | Identities | Gaps | Strand | Frame |
|---|---|---|---|---|---|
| 14.4 bits(7) | 223() | 7/7(100%) | 0/7(0%) | Plus/Plus | |

Features:

```
Query  284       TGTTCTC  290
                 |||||||
Sbjct  32573964  TGTTCTC  32573970
```

Range 596: 32575145 to 32575151

| Score | Expect | Identities | Gaps | Strand | Frame |
|---|---|---|---|---|---|
| 14.4 bits(7) | 223() | 7/7(100%) | 0/7(0%) | Plus/Plus | |

Features:

```
Query  289       TCTCCTT  295
                 |||||||
Sbjct  32575145  TCTCCTT  32575151
```

Range 597: 32575262 to 32575268

| Score | Expect | Identities | Gaps | Strand | Frame |
|---|---|---|---|---|---|
| 14.4 bits(7) | 223() | 7/7(100%) | 0/7(0%) | Plus/Minus | |

Features:

```
Query  285       GTTCTCT  291
                 |||||||
Sbjct  32575268  GTTCTCT  32575262
```

Range 598: 32576513 to 32576519

| Score | Expect | Identities | Gaps | Strand | Frame |
|---|---|---|---|---|---|
| 14.4 bits(7) | 223() | 7/7(100%) | 0/7(0%) | Plus/Plus | |

Features:

```
Query  290       CTCCTTT  296
                 |||||||
Sbjct  32576513  CTCCTTT  32576519
```

Range 599: 32576666 to 32576672

| Score | Expect | Identities | Gaps | Strand | Frame |
|---|---|---|---|---|---|
| 14.4 bits(7) | 223() | 7/7(100%) | 0/7(0%) | Plus/Minus | |

Features:

```
Query  273       TAACAAG  279
                 |||||||
Sbjct  32576672  TAACAAG  32576666
```

Range 600: 32576747 to 32576753

| Score | Expect | Identities | Gaps | Strand | Frame |
|---|---|---|---|---|---|

| 14.4 bits(7) | 223() | 7/7(100%) | 0/7(0%) | Plus/Plus |
|---|---|---|---|---|

Features:

```
Query  293       CTTTATT  299
                 |||||||
Sbjct  32576747  CTTTATT  32576753
```

Range 601: 32577480 to 32577486

| Score | Expect | Identities | Gaps | Strand | Frame |
|---|---|---|---|---|---|
| 14.4 bits(7) | 223() | 7/7(100%) | 0/7(0%) | Plus/Minus | |

Features:

```
Query  271       TGTAACA  277
                 |||||||
Sbjct  32577486  TGTAACA  32577480
```

Range 602: 32577742 to 32577748

| Score | Expect | Identities | Gaps | Strand | Frame |
|---|---|---|---|---|---|
| 14.4 bits(7) | 223() | 7/7(100%) | 0/7(0%) | Plus/Minus | |

Features:

```
Query  280       CTGGTGT  286
                 |||||||
Sbjct  32577748  CTGGTGT  32577742
```

Range 603: 32577811 to 32577817

| Score | Expect | Identities | Gaps | Strand | Frame |
|---|---|---|---|---|---|
| 14.4 bits(7) | 223() | 7/7(100%) | 0/7(0%) | Plus/Plus | |

Features:

```
Query  293       CTTTATT  299
                 |||||||
Sbjct  32577811  CTTTATT  32577817
```

Range 604: 32578428 to 32578434

| Score | Expect | Identities | Gaps | Strand | Frame |
|---|---|---|---|---|---|
| 14.4 bits(7) | 223() | 7/7(100%) | 0/7(0%) | Plus/Plus | |

Features:

```
Query  283       GTGTTCT  289
                 |||||||
Sbjct  32578428  GTGTTCT  32578434
```

Range 605: 32578902 to 32578908

| Score | Expect | Identities | Gaps | Strand | Frame |
|---|---|---|---|---|---|
| 14.4 bits(7) | 223() | 7/7(100%) | 0/7(0%) | Plus/Minus | |

Features:

```
Query  293       CTTTATT  299
                 |||||||
Sbjct  32578908  CTTTATT  32578902
```

Range 606: 32579211 to 32579217

| Score | Expect | Identities | Gaps | Strand | Frame |
|---|---|---|---|---|---|
| 14.4 bits(7) | 223() | 7/7(100%) | 0/7(0%) | Plus/Minus | |

Features:

```
Query  286       TTCTCTC  292
                 |||||||
Sbjct  32579217  TTCTCTC  32579211
```

Range 607: 32579705 to 32579711

| Score | Expect | Identities | Gaps | Strand | Frame |
|---|---|---|---|---|---|
| 14.4 bits(7) | 223() | 7/7(100%) | 0/7(0%) | Plus/Minus | |

Features:

```
Query  293       CTTTATT  299
                 |||||||
Sbjct  32579711  CTTTATT  32579705
```

Range 608: 32579725 to 32579731

| Score | Expect | Identities | Gaps | Strand | Frame |
|---|---|---|---|---|---|
| 14.4 bits(7) | 223() | 7/7(100%) | 0/7(0%) | Plus/Plus | |

Features:

```
Query  277       AAGCTGG  283
                 |||||||
Sbjct  32579725  AAGCTGG  32579731
```

Range 609: 32579766 to 32579772

| Score | Expect | Identities | Gaps | Strand | Frame |
|---|---|---|---|---|---|
| 14.4 bits(7) | 223() | 7/7(100%) | 0/7(0%) | Plus/Plus | |

Features:

```
Query  271       TGTAACA  277
                 |||||||
Sbjct  32579766  TGTAACA  32579772
```

Range 610: 32580342 to 32580348

| Score | Expect | Identities | Gaps | Strand | Frame |
|---|---|---|---|---|---|
| 14.4 bits(7) | 223() | 7/7(100%) | 0/7(0%) | Plus/Minus | |

Features:

```
Query  291       TCCTTTA  297
                 |||||||
Sbjct  32580348  TCCTTTA  32580342
```

Range 611: 32580933 to 32580939

| Score | Expect | Identities | Gaps | Strand | Frame |
|---|---|---|---|---|---|
| 14.4 bits(7) | 223() | 7/7(100%) | 0/7(0%) | Plus/Plus | |

Features:

```
Query  289       TCTCCTT  295
                 |||||||
Sbjct  32580933  TCTCCTT  32580939
```

Range 612: 32580992 to 32580998

| Score | Expect | Identities | Gaps | Strand | Frame |
|---|---|---|---|---|---|
| 14.4 bits(7) | 223() | 7/7(100%) | 0/7(0%) | Plus/Plus | |

Features:

```
Query  284       TGTTCTC  290
                 |||||||
Sbjct  32580992  TGTTCTC  32580998
```

Range 613: 32581301 to 32581307

| Score | Expect | Identities | Gaps | Strand | Frame |
|---|---|---|---|---|---|
| 14.4 bits(7) | 223() | 7/7(100%) | 0/7(0%) | Plus/Minus | |

Features:

```
Query  285       GTTCTCT  291
                 |||||||
Sbjct  32581307  GTTCTCT  32581301
```

Range 614: 32581779 to 32581785

| Score | Expect | Identities | Gaps | Strand | Frame |
|---|---|---|---|---|---|
| 14.4 bits(7) | 223() | 7/7(100%) | 0/7(0%) | Plus/Minus | |

Features:

```
Query  290       CTCCTTT  296
                 |||||||
Sbjct  32581785  CTCCTTT  32581779
```

Range 615: 32582221 to 32582227

| Score | Expect | Identities | Gaps | Strand | Frame |
|---|---|---|---|---|---|
| 14.4 bits(7) | 223() | 7/7(100%) | 0/7(0%) | Plus/Plus | |

Features:

```
Query  277       AAGCTGG  283
                 |||||||
Sbjct  32582221  AAGCTGG  32582227
```

Range 616: 32582377 to 32582383

| Score | Expect | Identities | Gaps | Strand | Frame |
|---|---|---|---|---|---|
| 14.4 bits(7) | 223() | 7/7(100%) | 0/7(0%) | Plus/Plus | |

Features:

```
Query  290       CTCCTTT  296
                 |||||||
Sbjct  32582377  CTCCTTT  32582383
```

Range 617: 32582850 to 32582856

| Score | Expect | Identities | Gaps | Strand | Frame |
|---|---|---|---|---|---|
| 14.4 bits(7) | 223() | 7/7(100%) | 0/7(0%) | Plus/Plus | |

Features:

```
Query  285       GTTCTCT  291
                 |||||||
Sbjct  32582850  GTTCTCT  32582856
```

Range 618: 32583398 to 32583404

| Score | Expect | Identities | Gaps | Strand | Frame |
|---|---|---|---|---|---|
| 14.4 bits(7) | 223() | 7/7(100%) | 0/7(0%) | Plus/Plus | |

Features:

```
Query  281       TGGTGTT  287
                 |||||||
Sbjct  32583398  TGGTGTT  32583404
```

Range 619: 32584724 to 32584730

| Score | Expect | Identities | Gaps | Strand | Frame |
|---|---|---|---|---|---|

| Score | Expect | Identities | Gaps | Strand | Frame |
|---|---|---|---|---|---|
| 14.4 bits(7) | 223() | 7/7(100%) | 0/7(0%) | Plus/Plus | |

Features:

```
Query  293       CTTTATT  299
                 |||||||
Sbjct  32584724  CTTTATT  32584730
```

Range 620: 32585963 to 32585969

| Score | Expect | Identities | Gaps | Strand | Frame |
|---|---|---|---|---|---|
| 14.4 bits(7) | 223() | 7/7(100%) | 0/7(0%) | Plus/Minus | |

Features:

```
Query  271       TGTAACA  277
                 |||||||
Sbjct  32585969  TGTAACA  32585963
```

Range 621: 32585990 to 32585996

| Score | Expect | Identities | Gaps | Strand | Frame |
|---|---|---|---|---|---|
| 14.4 bits(7) | 223() | 7/7(100%) | 0/7(0%) | Plus/Minus | |

Features:

```
Query  293       CTTTATT  299
                 |||||||
Sbjct  32585996  CTTTATT  32585990
```

Range 622: 32586336 to 32586342

| Score | Expect | Identities | Gaps | Strand | Frame |
|---|---|---|---|---|---|
| 14.4 bits(7) | 223() | 7/7(100%) | 0/7(0%) | Plus/Plus | |

Features:

```
Query  293       CTTTATT  299
                 |||||||
Sbjct  32586336  CTTTATT  32586342
```

Range 623: 32586907 to 32586913

| Score | Expect | Identities | Gaps | Strand | Frame |
|---|---|---|---|---|---|
| 14.4 bits(7) | 223() | 7/7(100%) | 0/7(0%) | Plus/Plus | |

Features:

```
Query  284       TGTTCTC  290
                 |||||||
Sbjct  32586907  TGTTCTC  32586913
```

Range 624: 32586915 to 32586921

| Score | Expect | Identities | Gaps | Strand | Frame |
|---|---|---|---|---|---|
| 14.4 bits(7) | 223() | 7/7(100%) | 0/7(0%) | Plus/Plus | |

Features:

```
Query  285       GTTCTCT  291
                 |||||||
Sbjct  32586915  GTTCTCT  32586921
```

Range 625: 32587064 to 32587070

| Score | Expect | Identities | Gaps | Strand | Frame |
|---|---|---|---|---|---|
| 14.4 bits(7) | 223() | 7/7(100%) | 0/7(0%) | Plus/Plus | |

Features:

```
Query  278       AGCTGGT  284
                 |||||||
Sbjct  32587064  AGCTGGT  32587070
```

Range 626: 32588585 to 32588591

| Score | Expect | Identities | Gaps | Strand | Frame |
|---|---|---|---|---|---|
| 14.4 bits(7) | 223() | 7/7(100%) | 0/7(0%) | Plus/Plus | |

Features:

```
Query  290       CTCCTTT  296
                 |||||||
Sbjct  32588585  CTCCTTT  32588591
```

Range 627: 32588608 to 32588614

| Score | Expect | Identities | Gaps | Strand | Frame |
|---|---|---|---|---|---|
| 14.4 bits(7) | 223() | 7/7(100%) | 0/7(0%) | Plus/Minus | |

Features:

```
Query  291       TCCTTTA  297
                 |||||||
Sbjct  32588614  TCCTTTA  32588608
```

Range 628: 32588937 to 32588943

| Score | Expect | Identities | Gaps | Strand | Frame |
|---|---|---|---|---|---|
| 14.4 bits(7) | 223() | 7/7(100%) | 0/7(0%) | Plus/Plus | |

Features:

```
Query  278       AGCTGGT  284
                 |||||||
Sbjct  32588937  AGCTGGT  32588943
```

Range 629: 32589142 to 32589148

| Score | Expect | Identities | Gaps | Strand | Frame |
|---|---|---|---|---|---|
| 14.4 bits(7) | 223() | 7/7(100%) | 0/7(0%) | Plus/Minus | |

Features:

```
Query  286       TTCTCTC  292
                 |||||||
Sbjct  32589148  TTCTCTC  32589142
```

Range 630: 32589219 to 32589225

| Score | Expect | Identities | Gaps | Strand | Frame |
|---|---|---|---|---|---|
| 14.4 bits(7) | 223() | 7/7(100%) | 0/7(0%) | Plus/Plus | |

Features:

```
Query  283       GTGTTCT  289
                 |||||||
Sbjct  32589219  GTGTTCT  32589225
```

Range 631: 32589458 to 32589464

| Score | Expect | Identities | Gaps | Strand | Frame |
|---|---|---|---|---|---|
| 14.4 bits(7) | 223() | 7/7(100%) | 0/7(0%) | Plus/Plus | |

Features:

```
Query  277       AAGCTGG  283
                 |||||||
Sbjct  32589458  AAGCTGG  32589464
```

Range 632: 32590551 to 32590557

| Score | Expect | Identities | Gaps | Strand | Frame |
|---|---|---|---|---|---|
| 14.4 bits(7) | 223() | 7/7(100%) | 0/7(0%) | Plus/Plus | |

Features:

```
Query  293       CTTTATT  299
                 |||||||
Sbjct  32590551  CTTTATT  32590557
```

Range 633: 32591112 to 32591118

| Score | Expect | Identities | Gaps | Strand | Frame |
|---|---|---|---|---|---|
| 14.4 bits(7) | 223() | 7/7(100%) | 0/7(0%) | Plus/Plus | |

Features:

```
Query  283       GTGTTCT  289
                 |||||||
Sbjct  32591112  GTGTTCT  32591118
```

Range 634: 32591526 to 32591532

| Score | Expect | Identities | Gaps | Strand | Frame |
|---|---|---|---|---|---|
| 14.4 bits(7) | 223() | 7/7(100%) | 0/7(0%) | Plus/Minus | |

Features:

```
Query  293       CTTTATT  299
                 |||||||
Sbjct  32591532  CTTTATT  32591526
```

Range 635: 32591668 to 32591674

| Score | Expect | Identities | Gaps | Strand | Frame |
|---|---|---|---|---|---|
| 14.4 bits(7) | 223() | 7/7(100%) | 0/7(0%) | Plus/Plus | |

Features:

```
Query  293       CTTTATT  299
                 |||||||
Sbjct  32591668  CTTTATT  32591674
```

Range 636: 32591990 to 32591996

| Score | Expect | Identities | Gaps | Strand | Frame |
|---|---|---|---|---|---|
| 14.4 bits(7) | 223() | 7/7(100%) | 0/7(0%) | Plus/Plus | |

Features:

```
Query  293       CTTTATT  299
                 |||||||
Sbjct  32591990  CTTTATT  32591996
```

Range 637: 32592226 to 32592232

| Score | Expect | Identities | Gaps | Strand | Frame |
|---|---|---|---|---|---|
| 14.4 bits(7) | 223() | 7/7(100%) | 0/7(0%) | Plus/Minus | |

Features:

```
Query  291       TCCTTTA  297
                 |||||||
Sbjct  32592232  TCCTTTA  32592226
```

Range 638: 32592830 to 32592836

| Score | Expect | Identities | Gaps | Strand | Frame |
|---|---|---|---|---|---|

| 14.4 bits(7) | 223() | 7/7(100%) | 0/7(0%) | Plus/Plus |
|---|---|---|---|---|

Features:

```
Query  282       GGTGTTC  288
                 |||||||
Sbjct  32592830  GGTGTTC  32592836
```

Range 639: 32592937 to 32592943

| Score | Expect | Identities | Gaps | Strand | Frame |
|---|---|---|---|---|---|
| 14.4 bits(7) | 223() | 7/7(100%) | 0/7(0%) | Plus/Plus | |

Features:

```
Query  285       GTTCTCT  291
                 |||||||
Sbjct  32592937  GTTCTCT  32592943
```

Range 640: 32593580 to 32593586

| Score | Expect | Identities | Gaps | Strand | Frame |
|---|---|---|---|---|---|
| 14.4 bits(7) | 223() | 7/7(100%) | 0/7(0%) | Plus/Plus | |

Features:

```
Query  291       TCCTTTA  297
                 |||||||
Sbjct  32593580  TCCTTTA  32593586
```

Range 641: 32594287 to 32594293

| Score | Expect | Identities | Gaps | Strand | Frame |
|---|---|---|---|---|---|
| 14.4 bits(7) | 223() | 7/7(100%) | 0/7(0%) | Plus/Minus | |

Features:

```
Query  284       TGTTCTC  290
                 |||||||
Sbjct  32594293  TGTTCTC  32594287
```

Range 642: 32594343 to 32594349

| Score | Expect | Identities | Gaps | Strand | Frame |
|---|---|---|---|---|---|
| 14.4 bits(7) | 223() | 7/7(100%) | 0/7(0%) | Plus/Plus | |

Features:

```
Query  292       CCTTTAT  298
                 |||||||
Sbjct  32594343  CCTTTAT  32594349
```

Range 643: 32594498 to 32594504

| Score | Expect | Identities | Gaps | Strand | Frame |
|---|---|---|---|---|---|
| 14.4 bits(7) | 223() | 7/7(100%) | 0/7(0%) | Plus/Minus | |

Features:

```
Query  276       CAAGCTG  282
                 |||||||
Sbjct  32594504  CAAGCTG  32594498
```

Range 644: 32594828 to 32594834

| Score | Expect | Identities | Gaps | Strand | Frame |
|---|---|---|---|---|---|
| 14.4 bits(7) | 223() | 7/7(100%) | 0/7(0%) | Plus/Plus | |

Features:

```
Query  293       CTTTATT  299
                 |||||||
Sbjct  32594828  CTTTATT  32594834
```

Range 645: 32595246 to 32595252

| Score | Expect | Identities | Gaps | Strand | Frame |
|---|---|---|---|---|---|
| 14.4 bits(7) | 223() | 7/7(100%) | 0/7(0%) | Plus/Plus | |

Features:

```
Query  271       TGTAACA  277
                 |||||||
Sbjct  32595246  TGTAACA  32595252
```

Range 646: 32595616 to 32595622

| Score | Expect | Identities | Gaps | Strand | Frame |
|---|---|---|---|---|---|
| 14.4 bits(7) | 223() | 7/7(100%) | 0/7(0%) | Plus/Plus | |

Features:

```
Query  292       CCTTTAT  298
                 |||||||
Sbjct  32595616  CCTTTAT  32595622
```

Range 647: 32595939 to 32595945

| Score | Expect | Identities | Gaps | Strand | Frame |
|---|---|---|---|---|---|
| 14.4 bits(7) | 223() | 7/7(100%) | 0/7(0%) | Plus/Minus | |

Features:

```
Query  276       CAAGCTG  282
                 |||||||
Sbjct  32595945  CAAGCTG  32595939
```

Range 648: 32596071 to 32596077

| Score | Expect | Identities | Gaps | Strand | Frame |
|---|---|---|---|---|---|
| 14.4 bits(7) | 223() | 7/7(100%) | 0/7(0%) | Plus/Plus | |

Features:

```
Query  293       CTTTATT  299
                 |||||||
Sbjct  32596071  CTTTATT  32596077
```

Range 649: 32596765 to 32596771

| Score | Expect | Identities | Gaps | Strand | Frame |
|---|---|---|---|---|---|
| 14.4 bits(7) | 223() | 7/7(100%) | 0/7(0%) | Plus/Plus | |

Features:

```
Query  283       GTGTTCT  289
                 |||||||
Sbjct  32596765  GTGTTCT  32596771
```

Range 650: 32597984 to 32597990

| Score | Expect | Identities | Gaps | Strand | Frame |
|---|---|---|---|---|---|
| 14.4 bits(7) | 223() | 7/7(100%) | 0/7(0%) | Plus/Minus | |

Features:

```
Query  293       CTTTATT  299
                 |||||||
Sbjct  32597990  CTTTATT  32597984
```

Range 651: 32598003 to 32598009

| Score | Expect | Identities | Gaps | Strand | Frame |
|---|---|---|---|---|---|
| 14.4 bits(7) | 223() | 7/7(100%) | 0/7(0%) | Plus/Plus | |

Features:

```
Query  289       TCTCCTT  295
                 |||||||
Sbjct  32598003  TCTCCTT  32598009
```

Range 652: 32598112 to 32598118

| Score | Expect | Identities | Gaps | Strand | Frame |
|---|---|---|---|---|---|
| 14.4 bits(7) | 223() | 7/7(100%) | 0/7(0%) | Plus/Minus | |

Features:

```
Query  290       CTCCTTT  296
                 |||||||
Sbjct  32598118  CTCCTTT  32598112
```

Range 653: 32598372 to 32598378

| Score | Expect | Identities | Gaps | Strand | Frame |
|---|---|---|---|---|---|
| 14.4 bits(7) | 223() | 7/7(100%) | 0/7(0%) | Plus/Minus | |

Features:

```
Query  271       TGTAACA  277
                 |||||||
Sbjct  32598378  TGTAACA  32598372
```

Range 654: 32598406 to 32598412

| Score | Expect | Identities | Gaps | Strand | Frame |
|---|---|---|---|---|---|
| 14.4 bits(7) | 223() | 7/7(100%) | 0/7(0%) | Plus/Plus | |

Features:

```
Query  286       TTCTCTC  292
                 |||||||
Sbjct  32598406  TTCTCTC  32598412
```

Range 655: 32599882 to 32599888

| Score | Expect | Identities | Gaps | Strand | Frame |
|---|---|---|---|---|---|
| 14.4 bits(7) | 223() | 7/7(100%) | 0/7(0%) | Plus/Plus | |

Features:

```
Query  292       CCTTTAT  298
                 |||||||
Sbjct  32599882  CCTTTAT  32599888
```

Range 656: 32601104 to 32601110

| Score | Expect | Identities | Gaps | Strand | Frame |
|---|---|---|---|---|---|
| 14.4 bits(7) | 223() | 7/7(100%) | 0/7(0%) | Plus/Plus | |

Features:

```
Query  275       ACAAGCT  281
                 |||||||
Sbjct  32601104  ACAAGCT  32601110
```

Range 657: 32601262 to 32601268

| Score | Expect | Identities | Gaps | Strand | Frame |
|---|---|---|---|---|---|

| Score | Expect | Identities | Gaps | Strand | Frame |
|---|---|---|---|---|---|
| 14.4 bits(7) | 223() | 7/7(100%) | 0/7(0%) | Plus/Minus | |

Features:

```
Query  289       TCTCCTT  295
                 |||||||
Sbjct  32601268  TCTCCTT  32601262
```

Range 658: 32601699 to 32601705

| Score | Expect | Identities | Gaps | Strand | Frame |
|---|---|---|---|---|---|
| 14.4 bits(7) | 223() | 7/7(100%) | 0/7(0%) | Plus/Minus | |

Features:

```
Query  283       GTGTTCT  289
                 |||||||
Sbjct  32601705  GTGTTCT  32601699
```

Range 659: 32601770 to 32601776

| Score | Expect | Identities | Gaps | Strand | Frame |
|---|---|---|---|---|---|
| 14.4 bits(7) | 223() | 7/7(100%) | 0/7(0%) | Plus/Plus | |

Features:

```
Query  285       GTTCTCT  291
                 |||||||
Sbjct  32601770  GTTCTCT  32601776
```

Range 660: 32601901 to 32601907

| Score | Expect | Identities | Gaps | Strand | Frame |
|---|---|---|---|---|---|
| 14.4 bits(7) | 223() | 7/7(100%) | 0/7(0%) | Plus/Minus | |

Features:

```
Query  290       CTCCTTT  296
                 |||||||
Sbjct  32601907  CTCCTTT  32601901
```

Range 661: 32602353 to 32602359

| Score | Expect | Identities | Gaps | Strand | Frame |
|---|---|---|---|---|---|
| 14.4 bits(7) | 223() | 7/7(100%) | 0/7(0%) | Plus/Plus | |

Features:

```
Query  284       TGTTCTC  290
                 |||||||
Sbjct  32602353  TGTTCTC  32602359
```

Range 662: 32602421 to 32602427

| Score | Expect | Identities | Gaps | Strand | Frame |
|---|---|---|---|---|---|
| 14.4 bits(7) | 223() | 7/7(100%) | 0/7(0%) | Plus/Plus | |

Features:

```
Query  277       AAGCTGG  283
                 |||||||
Sbjct  32602421  AAGCTGG  32602427
```

Range 663: 32603118 to 32603124

| Score | Expect | Identities | Gaps | Strand | Frame |
|---|---|---|---|---|---|
| 14.4 bits(7) | 223() | 7/7(100%) | 0/7(0%) | Plus/Minus | |

Features:

```
Query  293       CTTTATT  299
                 |||||||
Sbjct  32603124  CTTTATT  32603118
```

Range 664: 32604134 to 32604140

| Score | Expect | Identities | Gaps | Strand | Frame |
|---|---|---|---|---|---|
| 14.4 bits(7) | 223() | 7/7(100%) | 0/7(0%) | Plus/Plus | |

Features:

```
Query  286       TTCTCTC  292
                 |||||||
Sbjct  32604134  TTCTCTC  32604140
```

Range 665: 32604149 to 32604155

| Score | Expect | Identities | Gaps | Strand | Frame |
|---|---|---|---|---|---|
| 14.4 bits(7) | 223() | 7/7(100%) | 0/7(0%) | Plus/Plus | |

Features:

```
Query  271       TGTAACA  277
                 |||||||
Sbjct  32604149  TGTAACA  32604155
```

Range 666: 32604477 to 32604483

| Score | Expect | Identities | Gaps | Strand | Frame |
|---|---|---|---|---|---|
| 14.4 bits(7) | 223() | 7/7(100%) | 0/7(0%) | Plus/Minus | |

Features:

```
Query  293       CTTTATT  299
                 |||||||
Sbjct  32604483  CTTTATT  32604477
```

Range 667: 32604638 to 32604644

| Score | Expect | Identities | Gaps | Strand | Frame |
|---|---|---|---|---|---|
| 14.4 bits(7) | 223() | 7/7(100%) | 0/7(0%) | Plus/Plus | |

Features:

```
Query  286       TTCTCTC  292
                 |||||||
Sbjct  32604638  TTCTCTC  32604644
```

Range 668: 32604981 to 32604987

| Score | Expect | Identities | Gaps | Strand | Frame |
|---|---|---|---|---|---|
| 14.4 bits(7) | 223() | 7/7(100%) | 0/7(0%) | Plus/Plus | |

Features:

```
Query  276       CAAGCTG  282
                 |||||||
Sbjct  32604981  CAAGCTG  32604987
```

Range 669: 32605266 to 32605272

| Score | Expect | Identities | Gaps | Strand | Frame |
|---|---|---|---|---|---|
| 14.4 bits(7) | 223() | 7/7(100%) | 0/7(0%) | Plus/Plus | |

Features:

```
Query  291       TCCTTTA  297
                 |||||||
Sbjct  32605266  TCCTTTA  32605272
```

Range 670: 32606007 to 32606013

| Score | Expect | Identities | Gaps | Strand | Frame |
|---|---|---|---|---|---|
| 14.4 bits(7) | 223() | 7/7(100%) | 0/7(0%) | Plus/Plus | |

Features:

```
Query  285       GTTCTCT  291
                 |||||||
Sbjct  32606007  GTTCTCT  32606013
```

Range 671: 32606148 to 32606154

| Score | Expect | Identities | Gaps | Strand | Frame |
|---|---|---|---|---|---|
| 14.4 bits(7) | 223() | 7/7(100%) | 0/7(0%) | Plus/Plus | |

Features:

```
Query  293       CTTTATT  299
                 |||||||
Sbjct  32606148  CTTTATT  32606154
```

Range 672: 32606217 to 32606223

| Score | Expect | Identities | Gaps | Strand | Frame |
|---|---|---|---|---|---|
| 14.4 bits(7) | 223() | 7/7(100%) | 0/7(0%) | Plus/Minus | |

Features:

```
Query  271       TGTAACA  277
                 |||||||
Sbjct  32606223  TGTAACA  32606217
```

Range 673: 32606498 to 32606504

| Score | Expect | Identities | Gaps | Strand | Frame |
|---|---|---|---|---|---|
| 14.4 bits(7) | 223() | 7/7(100%) | 0/7(0%) | Plus/Minus | |

Features:

```
Query  289       TCTCCTT  295
                 |||||||
Sbjct  32606504  TCTCCTT  32606498
```

Range 674: 32607288 to 32607294

| Score | Expect | Identities | Gaps | Strand | Frame |
|---|---|---|---|---|---|
| 14.4 bits(7) | 223() | 7/7(100%) | 0/7(0%) | Plus/Plus | |

Features:

```
Query  292       CCTTTAT  298
                 |||||||
Sbjct  32607288  CCTTTAT  32607294
```

Range 675: 32608037 to 32608043

| Score | Expect | Identities | Gaps | Strand | Frame |
|---|---|---|---|---|---|
| 14.4 bits(7) | 223() | 7/7(100%) | 0/7(0%) | Plus/Plus | |

Features:

```
Query  292       CCTTTAT  298
                 |||||||
Sbjct  32608037  CCTTTAT  32608043
```

Range 676: 32608394 to 32608400

| Score | Expect | Identities | Gaps | Strand | Frame |
|---|---|---|---|---|---|

| | | | | |
|---|---|---|---|---|
| 14.4 bits(7) | 223() | 7/7(100%) | 0/7(0%) | Plus/Plus |

Features:

```
Query  288       CTCTCCT  294
                 |||||||
Sbjct  32608394  CTCTCCT  32608400
```

Range 677: 32608720 to 32608726

| Score | Expect | Identities | Gaps | Strand | Frame |
|---|---|---|---|---|---|
| 14.4 bits(7) | 223() | 7/7(100%) | 0/7(0%) | Plus/Plus | |

Features:

```
Query  293       CTTTATT  299
                 |||||||
Sbjct  32608720  CTTTATT  32608726
```

Range 678: 32608865 to 32608871

| Score | Expect | Identities | Gaps | Strand | Frame |
|---|---|---|---|---|---|
| 14.4 bits(7) | 223() | 7/7(100%) | 0/7(0%) | Plus/Minus | |

Features:

```
Query  289       TCTCCTT  295
                 |||||||
Sbjct  32608871  TCTCCTT  32608865
```

Range 679: 32609336 to 32609342

| Score | Expect | Identities | Gaps | Strand | Frame |
|---|---|---|---|---|---|
| 14.4 bits(7) | 223() | 7/7(100%) | 0/7(0%) | Plus/Plus | |

Features:

```
Query  287       TCTCTCC  293
                 |||||||
Sbjct  32609336  TCTCTCC  32609342
```

Range 680: 32610058 to 32610064

| Score | Expect | Identities | Gaps | Strand | Frame |
|---|---|---|---|---|---|
| 14.4 bits(7) | 223() | 7/7(100%) | 0/7(0%) | Plus/Minus | |

Features:

```
Query  272       GTAACAA  278
                 |||||||
Sbjct  32610064  GTAACAA  32610058
```

Range 681: 32610154 to 32610160

| Score | Expect | Identities | Gaps | Strand | Frame |
|---|---|---|---|---|---|
| 14.4 bits(7) | 223() | 7/7(100%) | 0/7(0%) | Plus/Plus | |

Features:

```
Query  281       TGGTGTT  287
                 |||||||
Sbjct  32610154  TGGTGTT  32610160
```

Range 682: 32610195 to 32610201

| Score | Expect | Identities | Gaps | Strand | Frame |
|---|---|---|---|---|---|
| 14.4 bits(7) | 223() | 7/7(100%) | 0/7(0%) | Plus/Plus | |

Features:

```
Query  292       CCTTTAT  298
                 |||||||
Sbjct  32610195  CCTTTAT  32610201
```

Range 683: 32611419 to 32611425

| Score | Expect | Identities | Gaps | Strand | Frame |
|---|---|---|---|---|---|
| 14.4 bits(7) | 223() | 7/7(100%) | 0/7(0%) | Plus/Minus | |

Features:

```
Query  274       AACAAGC  280
                 |||||||
Sbjct  32611425  AACAAGC  32611419
```

Range 684: 32611422 to 32611428

| Score | Expect | Identities | Gaps | Strand | Frame |
|---|---|---|---|---|---|
| 14.4 bits(7) | 223() | 7/7(100%) | 0/7(0%) | Plus/Plus | |

Features:

```
Query  284       TGTTCTC  290
                 |||||||
Sbjct  32611422  TGTTCTC  32611428
```

Range 685: 32611719 to 32611725

| Score | Expect | Identities | Gaps | Strand | Frame |
|---|---|---|---|---|---|
| 14.4 bits(7) | 223() | 7/7(100%) | 0/7(0%) | Plus/Minus | |

Features:

```
Query  272       GTAACAA  278
                 |||||||
Sbjct  32611725  GTAACAA  32611719
```

Range 686: 32612397 to 32612403

| Score | Expect | Identities | Gaps | Strand | Frame |
|---|---|---|---|---|---|
| 14.4 bits(7) | 223() | 7/7(100%) | 0/7(0%) | Plus/Plus | |

Features:

```
Query  271       TGTAACA  277
                 |||||||
Sbjct  32612397  TGTAACA  32612403
```

Range 687: 32612827 to 32612833

| Score | Expect | Identities | Gaps | Strand | Frame |
|---|---|---|---|---|---|
| 14.4 bits(7) | 223() | 7/7(100%) | 0/7(0%) | Plus/Minus | |

Features:

```
Query  286       TTCTCTC  292
                 |||||||
Sbjct  32612833  TTCTCTC  32612827
```

Range 688: 32613009 to 32613015

| Score | Expect | Identities | Gaps | Strand | Frame |
|---|---|---|---|---|---|
| 14.4 bits(7) | 223() | 7/7(100%) | 0/7(0%) | Plus/Plus | |

Features:

```
Query  287       TCTCTCC  293
                 |||||||
Sbjct  32613009  TCTCTCC  32613015
```

Range 689: 32614041 to 32614047

| Score | Expect | Identities | Gaps | Strand | Frame |
|---|---|---|---|---|---|
| 14.4 bits(7) | 223() | 7/7(100%) | 0/7(0%) | Plus/Plus | |

Features:

```
Query  289       TCTCCTT  295
                 |||||||
Sbjct  32614041  TCTCCTT  32614047
```

Range 690: 32614271 to 32614277

| Score | Expect | Identities | Gaps | Strand | Frame |
|---|---|---|---|---|---|
| 14.4 bits(7) | 223() | 7/7(100%) | 0/7(0%) | Plus/Minus | |

Features:

```
Query  277       AAGCTGG  283
                 |||||||
Sbjct  32614277  AAGCTGG  32614271
```

Range 691: 32614389 to 32614395

| Score | Expect | Identities | Gaps | Strand | Frame |
|---|---|---|---|---|---|
| 14.4 bits(7) | 223() | 7/7(100%) | 0/7(0%) | Plus/Minus | |

Features:

```
Query  277       AAGCTGG  283
                 |||||||
Sbjct  32614395  AAGCTGG  32614389
```

Range 692: 32615193 to 32615199

| Score | Expect | Identities | Gaps | Strand | Frame |
|---|---|---|---|---|---|
| 14.4 bits(7) | 223() | 7/7(100%) | 0/7(0%) | Plus/Plus | |

Features:

```
Query  286       TTCTCTC  292
                 |||||||
Sbjct  32615193  TTCTCTC  32615199
```

Range 693: 32615310 to 32615316

| Score | Expect | Identities | Gaps | Strand | Frame |
|---|---|---|---|---|---|
| 14.4 bits(7) | 223() | 7/7(100%) | 0/7(0%) | Plus/Plus | |

Features:

```
Query  289       TCTCCTT  295
                 |||||||
Sbjct  32615310  TCTCCTT  32615316
```

Range 694: 32615522 to 32615528

| Score | Expect | Identities | Gaps | Strand | Frame |
|---|---|---|---|---|---|
| 14.4 bits(7) | 223() | 7/7(100%) | 0/7(0%) | Plus/Plus | |

Features:

```
Query  292       CCTTTAT  298
                 |||||||
Sbjct  32615522  CCTTTAT  32615528
```

Range 695: 32615975 to 32615981

| Score | Expect | Identities | Gaps | Strand | Frame |
|---|---|---|---|---|---|

| | | | | |
|---|---|---|---|---|
| 14.4 bits(7) | 223() | 7/7(100%) | 0/7(0%) | Plus/Plus |

Features:

```
Query  286       TTCTCTC  292
                 |||||||
Sbjct  32615975  TTCTCTC  32615981
```

Range 696: 32616554 to 32616560

| Score | Expect | Identities | Gaps | Strand | Frame |
|---|---|---|---|---|---|
| 14.4 bits(7) | 223() | 7/7(100%) | 0/7(0%) | Plus/Plus | |

Features:

```
Query  284       TGTTCTC  290
                 |||||||
Sbjct  32616554  TGTTCTC  32616560
```

Range 697: 32616744 to 32616750

| Score | Expect | Identities | Gaps | Strand | Frame |
|---|---|---|---|---|---|
| 14.4 bits(7) | 223() | 7/7(100%) | 0/7(0%) | Plus/Plus | |

Features:

```
Query  286       TTCTCTC  292
                 |||||||
Sbjct  32616744  TTCTCTC  32616750
```

Range 698: 32617254 to 32617260

| Score | Expect | Identities | Gaps | Strand | Frame |
|---|---|---|---|---|---|
| 14.4 bits(7) | 223() | 7/7(100%) | 0/7(0%) | Plus/Plus | |

Features:

```
Query  278       AGCTGGT  284
                 |||||||
Sbjct  32617254  AGCTGGT  32617260
```

Range 699: 32617432 to 32617438

| Score | Expect | Identities | Gaps | Strand | Frame |
|---|---|---|---|---|---|
| 14.4 bits(7) | 223() | 7/7(100%) | 0/7(0%) | Plus/Plus | |

Features:

```
Query  291       TCCTTTA  297
                 |||||||
Sbjct  32617432  TCCTTTA  32617438
```

Range 700: 32617465 to 32617471

| Score | Expect | Identities | Gaps | Strand | Frame |
|---|---|---|---|---|---|
| 14.4 bits(7) | 223() | 7/7(100%) | 0/7(0%) | Plus/Minus | |

Features:

```
Query  276       CAAGCTG  282
                 |||||||
Sbjct  32617471  CAAGCTG  32617465
```

Range 701: 32617581 to 32617587

| Score | Expect | Identities | Gaps | Strand | Frame |
|---|---|---|---|---|---|
| 14.4 bits(7) | 223() | 7/7(100%) | 0/7(0%) | Plus/Plus | |

Features:

```
Query  271       TGTAACA  277
                 |||||||
Sbjct  32617581  TGTAACA  32617587
```

Range 702: 32618027 to 32618033

| Score | Expect | Identities | Gaps | Strand | Frame |
|---|---|---|---|---|---|
| 14.4 bits(7) | 223() | 7/7(100%) | 0/7(0%) | Plus/Minus | |

Features:

```
Query  289       TCTCCTT  295
                 |||||||
Sbjct  32618033  TCTCCTT  32618027
```

Range 703: 32618695 to 32618701

| Score | Expect | Identities | Gaps | Strand | Frame |
|---|---|---|---|---|---|
| 14.4 bits(7) | 223() | 7/7(100%) | 0/7(0%) | Plus/Plus | |

Features:

```
Query  290       CTCCTTT  296
                 |||||||
Sbjct  32618695  CTCCTTT  32618701
```

Range 704: 32618794 to 32618800

| Score | Expect | Identities | Gaps | Strand | Frame |
|---|---|---|---|---|---|
| 14.4 bits(7) | 223() | 7/7(100%) | 0/7(0%) | Plus/Minus | |

Features:

```
Query  271       TGTAACA  277
                 |||||||
Sbjct  32618800  TGTAACA  32618794
```

Range 705: 32618861 to 32618867

| Score | Expect | Identities | Gaps | Strand | Frame |
|---|---|---|---|---|---|
| 14.4 bits(7) | 223() | 7/7(100%) | 0/7(0%) | Plus/Minus | |

Features:

```
Query  293       CTTTATT  299
                 |||||||
Sbjct  32618867  CTTTATT  32618861
```

BLAST is a registered trademark of the National Library of Medicine

Support center Mailing list YouTube

- National Library Of Medicine
- National Institutes Of Health
- U.S. Department of Health & Human Services

- USA.gov

**NCBI**

*National Center for Biotechnology Information, U.S. National Library of Medicine 8600 Rockville Pike, Bethesda MD, 20894 USA*

Policies and Guidelines | Contact
