## Supplemental Figure S6 for "MORT, a locus for apoptosis in the human immunodeficiency virus-type 1 antisense gene: implications for AIDS, Cancer, and Covid-19"

### BLAST ®

### Basic Local Alignment Search Tool

NCBI/ BLAST/ blastn suite/ **Formatting Results - B285YCN201R**
Formatting options
Download
Blast report description

##### CQ767337:Sequence 27 from Patent EP1359221

**RID** B285YCN201R (Expires on 02-04 04:33 am)

**Query ID** gi|44909347|emb|CQ767337.1|
**Description** Sequence 27 from Patent EP1359221.
**Molecule type** rna
**Query Length** 367

**Database Name** Genome (all assemblies top-level)
**Description** Homo sapiens all assemblies [GCF_000001405.28 GCF_000306695.2] chromosomes plus unplaced and unlocalized scaffolds in Annotation Release 107
**Program** BLASTN 2.3.1+

#### Graphic Summary

##### Distribution of 177 Blast Hits on the Query Sequence

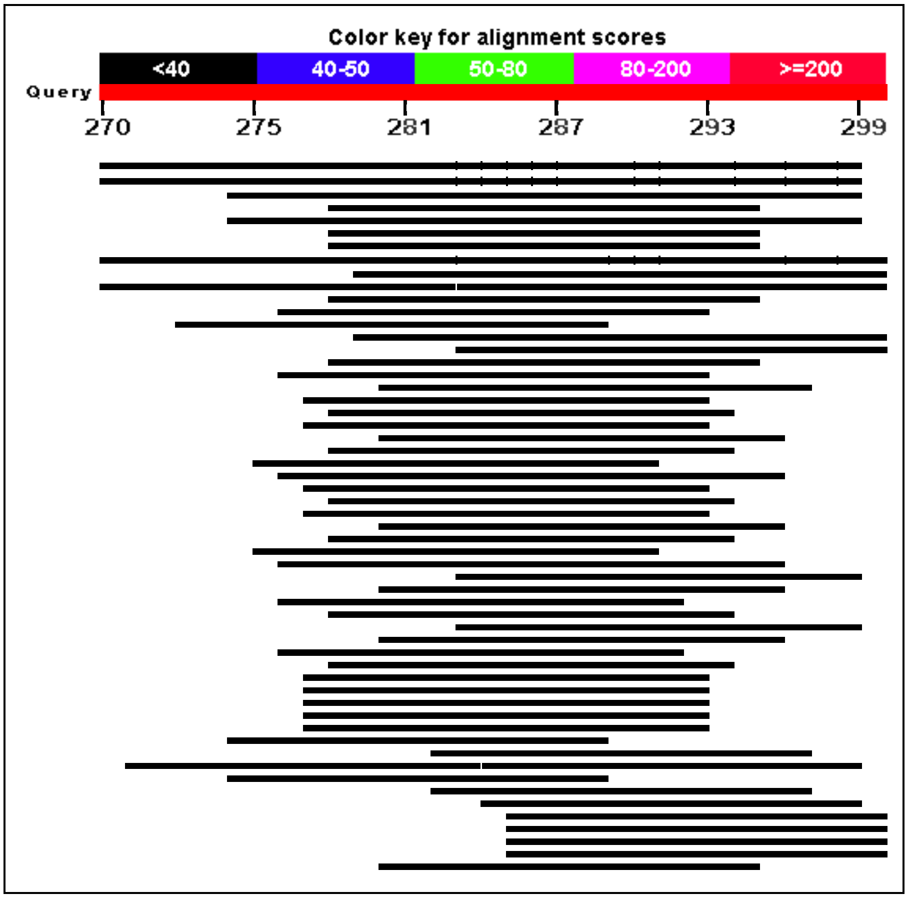

#### Descriptions

Sequences producing significant alignments:

| Description | Max score | Total score | Query cover | E value | Ident | Accession |
|---|---|---|---|---|---|---|
| Homo sapiens chromosome X, alternate assembly CHM1_1.1 | 36.2 | 1127 | 96% | 1.0 | 100% | NC_018934.2 |
| Homo sapiens chromosome X, GRCh38.p2 Primary Assembly | 36.2 | 1127 | 96% | 1.0 | 100% | NC_000023.11 |
| Homo sapiens chromosome 15, alternate assembly CHM1_1.1 | 34.2 | 752 | 87% | 4.1 | 92% | NC_018926.2 |
| Homo sapiens chromosome 22, alternate assembly CHM1_1.1 | 34.2 | 288 | 77% | 4.1 | 100% | NC_018933.2 |
| Homo sapiens chromosome 15, GRCh38.p2 Primary Assembly | 34.2 | 780 | 87% | 4.1 | 92% | NC_000015.10 |
| Homo sapiens chromosome 22, GRCh38.p2 Primary Assembly | 34.2 | 316 | 77% | 4.1 | 100% | NC_000022.11 |
| Homo sapiens chromosome 22 genomic scaffold, GRCh38.p2 alternate locus group ALT_REF_LOCI_1 HSCHR22_1_CTG6 | 34.2 | 34.2 | 54% | 4.1 | 100% | NT_187632.1 |
| Homo sapiens chromosome 2, alternate assembly CHM1_1.1 | 34.2 | 2091 | 100% | 4.1 | 100% | NC_018913.2 |
| Homo sapiens chromosome 4, alternate assembly CHM1_1.1 | 34.2 | 1792 | 96% | 4.1 | 95% | NC_018915.2 |
| Homo sapiens chromosome 5, alternate assembly CHM1_1.1 | 34.2 | 2147 | 90% | 4.1 | 100% | NC_018916.2 |
| Homo sapiens chromosome 7, alternate assembly CHM1_1.1 | 34.2 | 1462 | 96% | 4.1 | 100% | NC_018918.2 |
| Homo sapiens chromosome 8, alternate assembly CHM1_1.1 | 34.2 | 1607 | 96% | 4.1 | 100% | NC_018919.2 |
| Homo sapiens chromosome 2, GRCh38.p2 Primary Assembly | 34.2 | 2119 | 100% | 4.1 | 100% | NC_000002.12 |
| Homo sapiens chromosome 4, GRCh38.p2 Primary Assembly | 34.2 | 1819 | 96% | 4.1 | 95% | NC_000004.12 |
| Homo sapiens chromosome 5, GRCh38.p2 Primary Assembly | 34.2 | 2177 | 90% | 4.1 | 100% | NC_000005.10 |
| Homo sapiens chromosome 7, GRCh38.p2 Primary Assembly | 34.2 | 1460 | 96% | 4.1 | 100% | NC_000007.14 |
| Homo sapiens chromosome 8, GRCh38.p2 Primary Assembly | 34.2 | 1607 | 96% | 4.1 | 100% | NC_000008.11 |
| Homo sapiens chromosome Y, GRCh38.p2 Primary Assembly | 34.2 | 554 | 90% | 4.1 | 100% | NC_000024.10 |
| Homo sapiens chromosome 1, alternate assembly CHM1_1.1 | 32.2 | 2053 | 100% | 16 | 100% | NC_018912.2 |
| Homo sapiens chromosome 3, alternate assembly CHM1_1.1 | 32.2 | 1877 | 100% | 16 | 100% | NC_018914.2 |
| Homo sapiens chromosome 6, alternate assembly CHM1_1.1 | 32.2 | 1702 | 96% | 16 | 100% | NC_018917.2 |
| Homo sapiens chromosome 12, alternate assembly CHM1_1.1 | 32.2 | 1375 | 87% | 16 | 100% | NC_018923.2 |
| Homo sapiens chromosome 13, alternate assembly CHM1_1.1 | 32.2 | 514 | 90% | 16 | 100% | NC_018924.2 |

| Description | Max score | Total score | Query cover | E value | Ident | Accession |
|---|---|---|---|---|---|---|
| Homo sapiens chromosome 19, alternate assembly CHM1_1.1 | 32.2 | 435 | 87% | 16 | 100% | NC_018930.2 |
| Homo sapiens chromosome 20, alternate assembly CHM1_1.1 | 32.2 | 605 | 87% | 16 | 95% | NC_018931.2 |
| Homo sapiens chromosome 1, GRCh38.p2 Primary Assembly | 32.2 | 2085 | 100% | 16 | 100% | NC_000001.11 |
| Homo sapiens chromosome 3, GRCh38.p2 Primary Assembly | 32.2 | 1905 | 100% | 16 | 100% | NC_000003.12 |
| Homo sapiens chromosome 6, GRCh38.p2 Primary Assembly | 32.2 | 1702 | 96% | 16 | 100% | NC_000006.12 |
| Homo sapiens chromosome 12, GRCh38.p2 Primary Assembly | 32.2 | 1375 | 87% | 16 | 100% | NC_000012.12 |
| Homo sapiens chromosome 13, GRCh38.p2 Primary Assembly | 32.2 | 542 | 90% | 16 | 100% | NC_000013.11 |
| Homo sapiens chromosome 19, GRCh38.p2 Primary Assembly | 32.2 | 435 | 87% | 16 | 100% | NC_000019.10 |
| Homo sapiens chromosome 20, GRCh38.p2 Primary Assembly | 32.2 | 605 | 87% | 16 | 95% | NC_000020.11 |
| Homo sapiens chromosome 9, alternate assembly CHM1_1.1 | 32.2 | 1034 | 90% | 16 | 100% | NC_018920.2 |
| Homo sapiens chromosome 10, alternate assembly CHM1_1.1 | 32.2 | 1010 | 90% | 16 | 100% | NC_018921.2 |
| Homo sapiens chromosome 14, alternate assembly CHM1_1.1 | 32.2 | 687 | 87% | 16 | 100% | NC_018925.2 |
| Homo sapiens chromosome 16, alternate assembly CHM1_1.1 | 32.2 | 834 | 100% | 16 | 100% | NC_018927.2 |
| Homo sapiens chromosome 9, GRCh38.p2 Primary Assembly | 32.2 | 1091 | 90% | 16 | 100% | NC_000009.12 |
| Homo sapiens chromosome 10, GRCh38.p2 Primary Assembly | 32.2 | 980 | 90% | 16 | 100% | NC_000010.11 |
| Homo sapiens chromosome 14, GRCh38.p2 Primary Assembly | 32.2 | 657 | 87% | 16 | 100% | NC_000014.9 |
| Homo sapiens chromosome 16, GRCh38.p2 Primary Assembly | 32.2 | 834 | 100% | 16 | 100% | NC_000016.10 |
| Homo sapiens chromosome 6 genomic scaffold, GRCh38.p2 alternate locus group ALT_REF_LOCI_7 HSCHR6_MHC_SSTO_CTG1 | 32.2 | 32.2 | 51% | 16 | 100% | NT_167249.2 |
| Homo sapiens chromosome 6 genomic scaffold, GRCh38.p2 alternate locus group ALT_REF_LOCI_6 HSCHR6_MHC_QBL_CTG1 | 32.2 | 32.2 | 51% | 16 | 100% | NT_167248.2 |
| Homo sapiens chromosome 6 genomic scaffold, GRCh38.p2 alternate locus group ALT_REF_LOCI_4 HSCHR6_MHC_MANN_CTG1 | 32.2 | 32.2 | 51% | 16 | 100% | NT_167246.2 |
| Homo sapiens chromosome 6 genomic scaffold, GRCh38.p2 alternate locus group ALT_REF_LOCI_3 HSCHR6_MHC_DBB_CTG1 | 32.2 | 32.2 | 51% | 16 | 100% | NT_167245.2 |

| Description | Max score | Total score | Query cover | E value | Ident | Accession |
|---|---|---|---|---|---|---|
| Homo sapiens chromosome 6 genomic scaffold, GRCh38.p2 alternate locus group ALT_REF_LOCI_2 HSCHR6_MHC_COX_CTG1 | 32.2 | 32.2 | 51% | 16 | 100% | NT_113891.3 |
| Homo sapiens chromosome 11, alternate assembly CHM1_1.1 | 30.2 | 1214 | 96% | 65 | 100% | NC_018922.2 |
| Homo sapiens chromosome 17, alternate assembly CHM1_1.1 | 30.2 | 568 | 100% | 65 | 100% | NC_018928.2 |
| Homo sapiens chromosome 18, alternate assembly CHM1_1.1 | 30.2 | 831 | 90% | 65 | 100% | NC_018929.2 |
| Homo sapiens chromosome 11, GRCh38.p2 Primary Assembly | 30.2 | 1214 | 96% | 65 | 100% | NC_000011.10 |
| Homo sapiens chromosome 17, GRCh38.p2 Primary Assembly | 30.2 | 570 | 96% | 65 | 100% | NC_000017.11 |
| Homo sapiens chromosome 18, GRCh38.p2 Primary Assembly | 30.2 | 831 | 90% | 65 | 100% | NC_000018.10 |
| Homo sapiens chromosome 5 genomic scaffold, GRCh38.p2 alternate locus group ALT_REF_LOCI_1 HSCHR5_2_CTG1_1 | 30.2 | 58.5 | 48% | 65 | 100% | NW_003315917.2 |
| Homo sapiens chromosome 21, alternate assembly CHM1_1.1 | 30.2 | 397 | 90% | 65 | 100% | NC_018932.2 |
| Homo sapiens chromosome 21, GRCh38.p2 Primary Assembly | 30.2 | 397 | 90% | 65 | 100% | NC_000021.9 |
| Homo sapiens chromosome 5 genomic scaffold, GRCh38.p2 alternate locus group ALT_REF_LOCI_2 HSCHR5_1_CTG1_1 | 30.2 | 58.5 | 48% | 65 | 100% | NT_187651.1 |
| Homo sapiens chromosome 3 genomic scaffold, GRCh38.p2 alternate locus group ALT_REF_LOCI_1 HSCHR3_2_CTG3 | 30.2 | 30.2 | 48% | 65 | 100% | NT_187534.1 |
| Homo sapiens chromosome 19 genomic scaffold, GRCh38.p2 alternate locus group ALT_REF_LOCI_1 HSCHR19_1_CTG2 | 30.2 | 30.2 | 74% | 65 | 91% | NW_003315962.1 |
| Homo sapiens chromosome 13 genomic patch of type FIX, GRCh38.p2 PATCHES HG2291_PATCH | 28.2 | 28.2 | 45% | 256 | 100% | NW_011332699.1 |
| Homo sapiens chromosome 8 genomic scaffold, GRCh38.p2 alternate locus group ALT_REF_LOCI_2 HSCHR8_6_CTG1 | 28.2 | 28.2 | 45% | 256 | 100% | NT_187655.1 |
| Homo sapiens chromosome 4 genomic scaffold, GRCh38.p2 alternate locus group ALT_REF_LOCI_1 HSCHR4_1_CTG9 | 28.2 | 28.2 | 45% | 256 | 100% | NT_167250.2 |
| Homo sapiens chromosome 17 genomic scaffold, GRCh38.p2 alternate locus group ALT_REF_LOCI_1 HSCHR17_7_CTG4 | 28.2 | 28.2 | 45% | 256 | 100% | NT_187614.1 |

| Description | Max score | Total score | Query cover | E value | Ident | Accession |
|---|---|---|---|---|---|---|
| Homo sapiens chromosome 12 genomic scaffold, GRCh38.p2 alternate locus group ALT_REF_LOCI_1 HSCHR12_2_CTG2_1 | 28.2 | 28.2 | 45% | 256 | 100% | NW_003315941.1 |
| Homo sapiens chromosome 22 genomic patch of type NOVEL, GRCh38.p2 PATCHES HSCHR22_4_CTG1 | 28.2 | 28.2 | 45% | 256 | 100% | NW_009646207.1 |
| Homo sapiens chromosome 22 genomic patch of type NOVEL, GRCh38.p2 PATCHES HSCHR22_5_CTG1 | 28.2 | 28.2 | 45% | 256 | 100% | NW_009646208.1 |
| Homo sapiens chromosome 4 genomic scaffold, GRCh38.p2 alternate locus group ALT_REF_LOCI_3 HSCHR4_7_CTG12 | 28.2 | 28.2 | 45% | 256 | 100% | NT_187679.1 |
| Homo sapiens chromosome 22 genomic scaffold, GRCh38.p2 alternate locus group ALT_REF_LOCI_3 HSCHR22_3_CTG1 | 28.2 | 28.2 | 45% | 256 | 100% | NT_187682.1 |
| Homo sapiens chromosome 4 genomic scaffold, GRCh38.p2 alternate locus group ALT_REF_LOCI_2 HSCHR4_6_CTG12 | 28.2 | 28.2 | 45% | 256 | 100% | NT_187650.1 |
| Homo sapiens chromosome 8 genomic scaffold, GRCh38.p2 alternate locus group ALT_REF_LOCI_2 HSCHR8_5_CTG1 | 28.2 | 28.2 | 45% | 256 | 100% | NT_187654.1 |
| Homo sapiens chromosome 4 genomic scaffold, GRCh38.p2 alternate locus group ALT_REF_LOCI_1 HSCHR4_1_CTG4 | 28.2 | 28.2 | 45% | 256 | 100% | NT_187540.1 |
| Homo sapiens chromosome 4 genomic scaffold, GRCh38.p2 alternate locus group ALT_REF_LOCI_1 HSCHR4_3_CTG12 | 28.2 | 28.2 | 45% | 256 | 100% | NT_187543.1 |
| Homo sapiens chromosome 8 genomic scaffold, GRCh38.p2 alternate locus group ALT_REF_LOCI_1 HSCHR8_1_CTG1 | 28.2 | 28.2 | 45% | 256 | 100% | NT_187565.1 |
| Homo sapiens chromosome 8 genomic scaffold, GRCh38.p2 alternate locus group ALT_REF_LOCI_1 HSCHR8_8_CTG1 | 28.2 | 28.2 | 45% | 256 | 100% | NT_187576.1 |
| Homo sapiens chromosome 9 genomic scaffold, GRCh38.p2 alternate locus group ALT_REF_LOCI_1 HSCHR9_1_CTG5 | 28.2 | 28.2 | 45% | 256 | 100% | NT_187578.1 |
| Homo sapiens chromosome 19 genomic scaffold, GRCh38.p2 alternate locus group ALT_REF_LOCI_1 HSCHR19_3_CTG3_1 | 28.2 | 28.2 | 45% | 256 | 100% | NT_187620.1 |

| Description | Max score | Total score | Query cover | E value | Ident | Accession |
|---|---|---|---|---|---|---|
| Homo sapiens chromosome 22 genomic scaffold, GRCh38.p2 alternate locus group ALT_REF_LOCI_2 HSCHR22_2_CTG1 | 28.2 | 28.2 | 45% | 256 | 100% | NW_004504305.1 |
| Homo sapiens chromosome 22 genomic scaffold, GRCh38.p2 alternate locus group ALT_REF_LOCI_1 HSCHR22_1_CTG1 | 28.2 | 28.2 | 45% | 256 | 100% | NW_003315971.2 |
| Homo sapiens unplaced genomic scaffold, GRCh38.p2 Primary Assembly HSCHRUN_RANDOM_CTG16 | 28.2 | 56.5 | 45% | 256 | 100% | NT_167218.1 |

#### Alignments

Homo sapiens chromosome X, alternate assembly CHM1_1.1
Sequence ID: **ref|NC_018934.2|** Length: 155181468 Number of Matches: 39
Range 1: 23172409 to 23172426

| Score | Expect | Identities | Gaps | Strand | Frame |
|---|---|---|---|---|---|
| 36.2 bits(18) | 1.0() | 18/18(100%) | 0/18(0%) | Plus/Plus | |

Features:
**121012 bp at 5' side: DEAD box protein 53211631 bp at 3' side: patched domain-containing protein 1**

```
Query  282       GGTGTTCTCTCCTTTATT  299
                 ||||||||||||||||||
Sbjct  23172409  GGTGTTCTCTCCTTTATT  23172426
```

Range 2: 39300524 to 39300539

| Score | Expect | Identities | Gaps | Strand | Frame |
|---|---|---|---|---|---|
| 32.2 bits(16) | 16() | 16/16(100%) | 0/16(0%) | Plus/Minus | |

Features:
**604692 bp at 5' side: mid1-interacting protein 1642635 bp at 3' side: BCL-6 corepressor isoform b**

```
Query  275       ACAAGCTGGTGTTCTC  290
                 ||||||||||||||||
Sbjct  39300539  ACAAGCTGGTGTTCTC  39300524
```

Range 3: 132708151 to 132708166

| Score | Expect | Identities | Gaps | Strand | Frame |
|---|---|---|---|---|---|
| 32.2 bits(16) | 16() | 16/16(100%) | 0/16(0%) | Plus/Minus | |

Features:
**glypican-3 isoform 4 precursorglypican-3 isoform 3 precursor**

```
Query  279        GCTGGTGTTCTCTCCT  294
                  ||||||||||||||||
Sbjct  132708166  GCTGGTGTTCTCTCCT  132708151
```

Range 4: 18494987 to 18495001

| Score | Expect | Identities | Gaps | Strand | Frame |
|---|---|---|---|---|---|
| 30.2 bits(15) | 65() | 15/15(100%) | 0/15(0%) | Plus/Plus | |

Features:
**112416 bp at 5' side: sex comb on midleg-like protein 260866 bp at 3' side: cyclin-dependent kinase-like 5**

```
Query  284       TGTTCTCTCCTTTAT  298
                 |||||||||||||||
Sbjct  18494987  TGTTCTCTCCTTTAT  18495001
```

Range 5: 20356814 to 20356828

| Score | Expect | Identities | Gaps | Strand | Frame |
|---|---|---|---|---|---|
| 30.2 bits(15) | 65() | 15/15(100%) | 0/15(0%) | Plus/Plus | |

Features:
**41653 bp at 5' side: ribosomal protein S6 kinase alpha-31066615 bp at 3' side: connector enhancer of kinase suppressor of ras 2 isoform 1**

```
Query  282       GGTGTTCTCTCCTTT  296
                 |||||||||||||||
Sbjct  20356814  GGTGTTCTCTCCTTT  20356828
```

Range 6: 37868053 to 37868067

| Score | Expect | Identities | Gaps | Strand | Frame |
|---|---|---|---|---|---|
| 30.2 bits(15) | 65() | 15/15(100%) | 0/15(0%) | Plus/Minus | |

Features:
**130150 bp at 5' side: dynein light chain Tctex-type 312908 bp at 3' side: huntingtin-interacting protein M**

```
Query  282       GGTGTTCTCTCCTTT  296
                 |||||||||||||||
Sbjct  37868067  GGTGTTCTCTCCTTT  37868053
```

Range 7: 51953865 to 51953879

| Score | Expect | Identities | Gaps | Strand | Frame |
|---|---|---|---|---|---|
| 30.2 bits(15) | 65() | 15/15(100%) | 0/15(0%) | Plus/Minus | |

Features:
**234916 bp at 5' side: melanoma-associated antigen D1 isoform b54095 bp at 3' side: melanoma-associated antigen D4 isoform X1**

```
Query  282       GGTGTTCTCTCCTTT  296
                 |||||||||||||||
Sbjct  51953879  GGTGTTCTCTCCTTT  51953865
```

Range 8: 101166652 to 101166666

| Score | Expect | Identities | Gaps | Strand | Frame |
|---|---|---|---|---|---|
| 30.2 bits(15) | 65() | 15/15(100%) | 0/15(0%) | Plus/Minus | |

Features:
**130981 bp at 5' side: zinc finger matrin-type protein 1 isoform 4109049 bp at 3' side: transcription elongation factor A protein-like 2**

```
Query  280        CTGGTGTTCTCTCCT  294
                  |||||||||||||||
Sbjct  101166666  CTGGTGTTCTCTCCT  101166652
```

Range 9: 2207149 to 2207162

| Score | Expect | Identities | Gaps | Strand | Frame |
|---|---|---|---|---|---|
| 28.2 bits(14) | 256() | 14/14(100%) | 0/14(0%) | Plus/Plus | |

Features:
**dehydrogenase/reductase SDR family member on chromosome X...**

```
Query  284      TGTTCTCTCCTTTA  297
                ||||||||||||||
Sbjct  2207149  TGTTCTCTCCTTTA  2207162
```

Range 10: 13099897 to 13099910

| Score | Expect | Identities | Gaps | Strand | Frame |
|---|---|---|---|---|---|
| 28.2 bits(14) | 256() | 14/14(100%) | 0/14(0%) | Plus/Plus | |

Features:
**uncharacterized protein LOC105373133 isoform X2uncharacterized protein LOC105373133 isoform X1**

```
Query  274       AACAAGCTGGTGTT  287
                 ||||||||||||||
Sbjct  13099897  AACAAGCTGGTGTT  13099910
```

Range 11: 22527176 to 22527189

| Score | Expect | Identities | Gaps | Strand | Frame |
|---|---|---|---|---|---|
| 28.2 bits(14) | 256() | 14/14(100%) | 0/14(0%) | Plus/Plus | |

Features:
**204102 bp at 5' side: E3 ubiquitin-protein ligase ZNF645522313 bp at 3' side: DEAD box protein 53**

```
Query  283       GTGTTCTCTCCTTT  296
                 ||||||||||||||
Sbjct  22527176  GTGTTCTCTCCTTT  22527189
```

Range 12: 63593776 to 63593793

| Score | Expect | Identities | Gaps | Strand | Frame |
|---|---|---|---|---|---|
| 28.2 bits(14) | 256() | 17/18(94%) | 0/18(0%) | Plus/Plus | |

Features:
**85841 bp at 5' side: myotubularin-related protein 8436492 bp at 3' side: zinc finger C4H2 domain-containing protein isoform 3**

```
Query  281       TGGTGTTCTCTCCTTTAT  298
                 ||||||| ||||||||||
Sbjct  63593776  TGGTGTTATCTCCTTTAT  63593793
```

Range 13: 66796434 to 66796447

| Score | Expect | Identities | Gaps | Strand | Frame |
|---|---|---|---|---|---|
| 28.2 bits(14) | 256() | 14/14(100%) | 0/14(0%) | Plus/Plus | |

Features:
**androgen receptor isoform 1androgen receptor isoform 2**

```
Query  271       TGTAACAAGCTGGT  284
                 ||||||||||||||
Sbjct  66796434  TGTAACAAGCTGGT  66796447
```

Range 14: 72591600 to 72591613

| Score | Expect | Identities | Gaps | Strand | Frame |
|---|---|---|---|---|---|
| 28.2 bits(14) | 256() | 14/14(100%) | 0/14(0%) | Plus/Plus | |

Features:
**24606 bp at 5' side: homeobox protein CDX-484349 bp at 3' side: cysteine-rich hydrophobic domain-containing protein 1 iso...**

```
Query  286       TTCTCTCCTTTATT  299
                 ||||||||||||||
Sbjct  72591600  TTCTCTCCTTTATT  72591613
```

Range 15: 73874303 to 73874316

| Score | Expect | Identities | Gaps | Strand | Frame |
|---|---|---|---|---|---|
| 28.2 bits(14) | 256() | 14/14(100%) | 0/14(0%) | Plus/Plus | |

Features:
**15954 bp at 5' side: protein KIAA2022291798 bp at 3' side: ATP-binding cassette sub-family B member 7, mitochondrial...**

```
Query  286       TTCTCTCCTTTATT  299
                 ||||||||||||||
Sbjct  73874303  TTCTCTCCTTTATT  73874316
```

Range 16: 77801842 to 77801855

| Score | Expect | Identities | Gaps | Strand | Frame |
|---|---|---|---|---|---|
| 28.2 bits(14) | 256() | 14/14(100%) | 0/14(0%) | Plus/Plus | |

Features:
**379516 bp at 5' side: cysteinyl leukotriene receptor 13650 bp at 3' side: zinc finger CCHC domain-containing protein 5**

```
Query  283       GTGTTCTCTCCTTT  296
                 ||||||||||||||
Sbjct  77801842  GTGTTCTCTCCTTT  77801855
```

Range 17: 107521405 to 107521418

| Score | Expect | Identities | Gaps | Strand | Frame |
|---|---|---|---|---|---|
| 28.2 bits(14) | 256() | 14/14(100%) | 0/14(0%) | Plus/Plus | |

Features:
**collagen alpha-6(IV) chain isoform A precursorcollagen alpha-6(IV) chain isoform 5 precursor**

```
Query  279        GCTGGTGTTCTCTC  292
                  ||||||||||||||
Sbjct  107521405  GCTGGTGTTCTCTC  107521418
```

Range 18: 108451033 to 108451046

| Score | Expect | Identities | Gaps | Strand | Frame |
|---|---|---|---|---|---|
| 28.2 bits(14) | 256() | 14/14(100%) | 0/14(0%) | Plus/Plus | |

Features:
**560571 bp at 5' side: insulin receptor substrate 479357 bp at 3' side: retinal guanylyl cyclase 2**

```
Query  270        GTGTAACAAGCTGG  283
                  ||||||||||||||
Sbjct  108451033  GTGTAACAAGCTGG  108451046
```

Range 19: 124750414 to 124750427

| Score | Expect | Identities | Gaps | Strand | Frame |
|---|---|---|---|---|---|
| 28.2 bits(14) | 256() | 14/14(100%) | 0/14(0%) | Plus/Plus | |

Features:
**382445 bp at 5' side: uncharacterized protein LOC100129520458975 bp at 3' side: DDB1- and CUL4-associated factor 12-like protein 2**

```
Query  276        CAAGCTGGTGTTCT  289
                  ||||||||||||||
Sbjct  124750414  CAAGCTGGTGTTCT  124750427
```

Range 20: 140463432 to 140463445

| Score | Expect | Identities | Gaps | Strand | Frame |
|---|---|---|---|---|---|
| 28.2 bits(14) | 256() | 14/14(100%) | 0/14(0%) | Plus/Plus | |

Features:
**281100 bp at 5' side: protein LDOC1119419 bp at 3' side: sperm protein associated with the nucleus on the X chromo...**

```
Query  286        TTCTCTCCTTTATT  299
                  ||||||||||||||
Sbjct  140463432  TTCTCTCCTTTATT  140463445
```

Range 21: 140926710 to 140926723

| Score | Expect | Identities | Gaps | Strand | Frame |
|---|---|---|---|---|---|
| 28.2 bits(14) | 256() | 14/14(100%) | 0/14(0%) | Plus/Plus | |

Features:

**18747 bp at 5' side: melanoma-associated antigen C1275072 bp at 3' side: melanoma-associated antigen C2**

```
Query  282        GGTGTTCTCTCCTT  295
                  ||||||||||||||
Sbjct  140926710  GGTGTTCTCTCCTT  140926723
```

Range 22: 146475581 to 146475594

| Score | Expect | Identities | Gaps | Strand | Frame |
|---|---|---|---|---|---|
| 28.2 bits(14) | 256() | 14/14(100%) | 0/14(0%) | Plus/Plus | |

Features:
**211363 bp at 5' side: uncharacterized protein LOC105373347 isoform X1428958 bp at 3' side: fragile X mental retardation protein 1 isoform ISO1**

```
Query  286        TTCTCTCCTTTATT  299
                  ||||||||||||||
Sbjct  146475581  TTCTCTCCTTTATT  146475594
```

Range 23: 5882256 to 5882269

| Score | Expect | Identities | Gaps | Strand | Frame |
|---|---|---|---|---|---|
| 28.2 bits(14) | 256() | 14/14(100%) | 0/14(0%) | Plus/Minus | |

Features:
**neuroligin-4, X-linkedneuroligin-4, X-linked**

```
Query  286      TTCTCTCCTTTATT  299
                ||||||||||||||
Sbjct  5882269  TTCTCTCCTTTATT  5882256
```

Range 24: 12501864 to 12501877

| Score | Expect | Identities | Gaps | Strand | Frame |
|---|---|---|---|---|---|
| 28.2 bits(14) | 256() | 14/14(100%) | 0/14(0%) | Plus/Minus | |

Features:
**FERM and PDZ domain-containing protein 4**

```
Query  286       TTCTCTCCTTTATT  299
                 ||||||||||||||
Sbjct  12501877  TTCTCTCCTTTATT  12501864
```

Range 25: 12757175 to 12757192

| Score | Expect | Identities | Gaps | Strand | Frame |
|---|---|---|---|---|---|
| 28.2 bits(14) | 256() | 17/18(94%) | 0/18(0%) | Plus/Minus | |

Features:
**FERM and PDZ domain-containing protein 4**

```
Query  279       GCTGGTGTTCTCTCCTTT  296
                 ||||||||| ||||||||
Sbjct  12757192  GCTGGTGTTGTCTCCTTT  12757175
```

Range 26: 14978358 to 14978371

| Score | Expect | Identities | Gaps | Strand | Frame |
|---|---|---|---|---|---|
| 28.2 bits(14) | 256() | 14/14(100%) | 0/14(0%) | Plus/Minus | |

Features:
**10004 bp at 5' side: motile sperm domain-containing protein 2 isoform 2315211 bp at 3' side: ankyrin repeat and SOCS box protein 9 isoform 1**

```
Query  286       TTCTCTCCTTTATT  299
                 ||||||||||||||
Sbjct  14978371  TTCTCTCCTTTATT  14978358
```

Range 27: 18187789 to 18187802

| Score | Expect | Identities | Gaps | Strand | Frame |
|---|---|---|---|---|---|
| 28.2 bits(14) | 256() | 14/14(100%) | 0/14(0%) | Plus/Minus | |

Features:
**337271 bp at 5' side: retinoic acid-induced protein 2 isoform 225703 bp at 3' side: BEN domain-containing protein 2 isoform 1**

```
Query  286       TTCTCTCCTTTATT  299
                 ||||||||||||||
Sbjct  18187802  TTCTCTCCTTTATT  18187789
```

Range 28: 18328519 to 18328532

| Score | Expect | Identities | Gaps | Strand | Frame |
|---|---|---|---|---|---|
| 28.2 bits(14) | 256() | 14/14(100%) | 0/14(0%) | Plus/Minus | |

Features:
**sex comb on midleg-like protein 2**

```
Query  278       AGCTGGTGTTCTCT  291
                 ||||||||||||||
Sbjct  18328532  AGCTGGTGTTCTCT  18328519
```

Range 29: 23000218 to 23000231

| Score | Expect | Identities | Gaps | Strand | Frame |
|---|---|---|---|---|---|
| 28.2 bits(14) | 256() | 14/14(100%) | 0/14(0%) | Plus/Minus | |

Features:
**677144 bp at 5' side: E3 ubiquitin-protein ligase ZNF64549271 bp at 3' side: DEAD box protein 53**

```
Query  272       GTAACAAGCTGGTG  285
                 ||||||||||||||
Sbjct  23000231  GTAACAAGCTGGTG  23000218
```

Range 30: 29805717 to 29805730

| Score | Expect | Identities | Gaps | Strand | Frame |
|---|---|---|---|---|---|
| 28.2 bits(14) | 256() | 14/14(100%) | 0/14(0%) | Plus/Minus | |

Features:
**interleukin-1 receptor accessory protein-like 1 precursor**

```
Query  281       TGGTGTTCTCTCCT  294
                 ||||||||||||||
Sbjct  29805730  TGGTGTTCTCTCCT  29805717
```

Range 31: 39248320 to 39248333

| Score | Expect | Identities | Gaps | Strand | Frame |
|---|---|---|---|---|---|
| 28.2 bits(14) | 256() | 14/14(100%) | 0/14(0%) | Plus/Minus | |

Features:
**552488 bp at 5' side: mid1-interacting protein 1694841 bp at 3' side: BCL-6 corepressor isoform b**

```
Query  279       GCTGGTGTTCTCTC  292
                 ||||||||||||||
Sbjct  39248333  GCTGGTGTTCTCTC  39248320
```

Range 32: 45071728 to 45071741

| Score | Expect | Identities | Gaps | Strand | Frame |
|---|---|---|---|---|---|
| 28.2 bits(14) | 256() | 14/14(100%) | 0/14(0%) | Plus/Minus | |

Features:
**deleted in autism-related protein 1 isoform 1 precursordeleted in autism-related protein 1 isoform 2 precursor**

```
Query  286       TTCTCTCCTTTATT  299
                 ||||||||||||||
Sbjct  45071741  TTCTCTCCTTTATT  45071728
```

Range 33: 67364784 to 67364797

| Score | Expect | Identities | Gaps | Strand | Frame |
|---|---|---|---|---|---|
| 28.2 bits(14) | 256() | 14/14(100%) | 0/14(0%) | Plus/Minus | |

Features:
**oligophrenin-1**

```
Query  274       AACAAGCTGGTGTT  287
                 ||||||||||||||
Sbjct  67364797  AACAAGCTGGTGTT  67364784
```

Range 34: 74500271 to 74500284

| Score | Expect | Identities | Gaps | Strand | Frame |
|---|---|---|---|---|---|
| 28.2 bits(14) | 256() | 14/14(100%) | 0/14(0%) | Plus/Minus | |

Features:
**palmitoyltransferase ZDHHC15 isoform 2palmitoyltransferase ZDHHC15 isoform 1**

```
Query  286       TTCTCTCCTTTATT  299
                 ||||||||||||||
Sbjct  74500284  TTCTCTCCTTTATT  74500271
```

Range 35: 75001452 to 75001465

| Score | Expect | Identities | Gaps | Strand | Frame |
|---|---|---|---|---|---|
| 28.2 bits(14) | 256() | 14/14(100%) | 0/14(0%) | Plus/Minus | |

Features:
**103720 bp at 5' side: melanoma-associated antigen E2260008 bp at 3' side: uncharacterized protein LOC105373256**

```
Query  279       GCTGGTGTTCTCTC  292
                 ||||||||||||||
Sbjct  75001465  GCTGGTGTTCTCTC  75001452
```

Range 36: 79260674 to 79260687

| Score | Expect | Identities | Gaps | Strand | Frame |
|---|---|---|---|---|---|
| 28.2 bits(14) | 256() | 14/14(100%) | 0/14(0%) | Plus/Minus | |

Features:
**80846 bp at 5' side: T-box transcription factor TBX22 isoform 2330773 bp at 3' side: protein FAM46D**

```
Query  271       TGTAACAAGCTGGT  284
                 ||||||||||||||
Sbjct  79260687  TGTAACAAGCTGGT  79260674
```

Range 37: 93622655 to 93622668

| Score | Expect | Identities | Gaps | Strand | Frame |
|---|---|---|---|---|---|
| 28.2 bits(14) | 256() | 14/14(100%) | 0/14(0%) | Plus/Minus | |

Features:
**764810 bp at 5' side: protein FAM133A2210099 bp at 3' side: protein diaphanous homolog 2 isoform 156**

```
Query  286       TTCTCTCCTTTATT  299
                 ||||||||||||||
Sbjct  93622668  TTCTCTCCTTTATT  93622655
```

Range 38: 93923237 to 93923250

| Score | Expect | Identities | Gaps | Strand | Frame |
|---|---|---|---|---|---|
| 28.2 bits(14) | 256() | 14/14(100%) | 0/14(0%) | Plus/Minus | |

Features:
**1065392 bp at 5' side: protein FAM133A1909517 bp at 3' side: protein diaphanous homolog 2 isoform 156**

```
Query  273       TAACAAGCTGGTGT  286
                 ||||||||||||||
Sbjct  93923250  TAACAAGCTGGTGT  93923237
```

Range 39: 120885534 to 120885547

| Score | Expect | Identities | Gaps | Strand | Frame |
|---|---|---|---|---|---|
| 28.2 bits(14) | 256() | 14/14(100%) | 0/14(0%) | Plus/Minus | |

Features:
**791312 bp at 5' side: glutamate dehydrogenase 2, mitochondrial precursor1343726 bp at 3' side: glutamate receptor 3 isoform 2 precursor**

```
Query  273        TAACAAGCTGGTGT  286
                  ||||||||||||||
Sbjct  120885547  TAACAAGCTGGTGT  120885534
```

Homo sapiens chromosome X, GRCh38.p2 Primary Assembly
Sequence ID: **ref|NC_000023.11|** Length: 156040895 Number of Matches: 39
Range 1: 23123020 to 23123037

| Score | Expect | Identities | Gaps | Strand | Frame |
|---|---|---|---|---|---|
| 36.2 bits(18) | 1.0() | 18/18(100%) | 0/18(0%) | Plus/Plus | |

Features:
**121067 bp at 5' side: DEAD box protein 53211839 bp at 3' side: patched domain-containing protein 1 isoform X1**

```
Query  282       GGTGTTCTCTCCTTTATT  299
                 ||||||||||||||||||
Sbjct  23123020  GGTGTTCTCTCCTTTATT  23123037
```

Range 2: 39409720 to 39409735

| Score | Expect | Identities | Gaps | Strand | Frame |
|---|---|---|---|---|---|
| 32.2 bits(16) | 16() | 16/16(100%) | 0/16(0%) | Plus/Minus | |

Features:
**604222 bp at 5' side: mid1-interacting protein 1642374 bp at 3' side: BCL-6 corepressor isoform X2**

```
Query  275       ACAAGCTGGTGTTCTC  290
                 ||||||||||||||||
Sbjct  39409735  ACAAGCTGGTGTTCTC  39409720
```

Range 3: 133662424 to 133662439

| Score | Expect | Identities | Gaps | Strand | Frame |
|---|---|---|---|---|---|
| 32.2 bits(16) | 16() | 16/16(100%) | 0/16(0%) | Plus/Minus | |

Features:
**glypican-3 isoform 4 precursorglypican-3 isoform 3 precursor**

```
Query  279        GCTGGTGTTCTCTCCT  294
                  ||||||||||||||||
Sbjct  133662439  GCTGGTGTTCTCTCCT  133662424
```

Range 4: 18446543 to 18446557

| Score | Expect | Identities | Gaps | Strand | Frame |
|---|---|---|---|---|---|
| 30.2 bits(15) | 65() | 15/15(100%) | 0/15(0%) | Plus/Plus | |

Features:
**112472 bp at 5' side: sex comb on midleg-like protein 2 isoform X260540 bp at 3' side: cyclin-dependent kinase-like 5**

```
Query  284       TGTTCTCTCCTTTAT  298
                 |||||||||||||||
Sbjct  18446543  TGTTCTCTCCTTTAT  18446557
```

Range 5: 20308385 to 20308399

| Score | Expect | Identities | Gaps | Strand | Frame |
|---|---|---|---|---|---|
| 30.2 bits(15) | 65() | 15/15(100%) | 0/15(0%) | Plus/Plus | |

Features:
**73586 bp at 5' side: ribosomal protein S6 kinase alpha-3 isoform X71066499 bp at 3' side: connector enhancer of kinase suppressor of ras 2 isoform 1**

```
Query  282       GGTGTTCTCTCCTTT  296
                 |||||||||||||||
Sbjct  20308385  GGTGTTCTCTCCTTT  20308399
```

Range 6: 37977918 to 37977932

| Score | Expect | Identities | Gaps | Strand | Frame |
|---|---|---|---|---|---|
| 30.2 bits(15) | 65() | 15/15(100%) | 0/15(0%) | Plus/Minus | |

Features:
**130408 bp at 5' side: dynein light chain Tctex-type 312908 bp at 3' side: huntingtin-interacting protein M**

```
Query  282       GGTGTTCTCTCCTTT  296
                 |||||||||||||||
Sbjct  37977932  GGTGTTCTCTCCTTT  37977918
```

Range 7: 52136791 to 52136805

| Score | Expect | Identities | Gaps | Strand | Frame |
|---|---|---|---|---|---|
| 30.2 bits(15) | 65() | 15/15(100%) | 0/15(0%) | Plus/Minus | |

Features:
**68619 bp at 5' side: melanoma-associated antigen D4 isoform X549118 bp at 3' side: melanoma-associated antigen D4 isoform 3**

```
Query  282       GGTGTTCTCTCCTTT  296
                 |||||||||||||||
Sbjct  52136805  GGTGTTCTCTCCTTT  52136791
```

Range 8: 102017766 to 102017780

| Score | Expect | Identities | Gaps | Strand | Frame |
|---|---|---|---|---|---|
| 30.2 bits(15) | 65() | 15/15(100%) | 0/15(0%) | Plus/Minus | |

Features:
**85758 bp at 5' side: zinc finger matrin-type protein 1 isoform X6109051 bp at 3' side: transcription elongation factor A protein-like 2**

```
Query  280        CTGGTGTTCTCTCCT  294
                  |||||||||||||||
Sbjct  102017780  CTGGTGTTCTCTCCT  102017766
```

Range 9: 2258700 to 2258713

| Score | Expect | Identities | Gaps | Strand | Frame |
|---|---|---|---|---|---|
| 28.2 bits(14) | 256() | 14/14(100%) | 0/14(0%) | Plus/Plus | |

Features:
**dehydrogenase/reductase SDR family member on chromosome X...**

```
Query  284      TGTTCTCTCCTTTA  297
                ||||||||||||||
Sbjct  2258700  TGTTCTCTCCTTTA  2258713
```

Range 10: 13051220 to 13051233

| Score | Expect | Identities | Gaps | Strand | Frame |
|---|---|---|---|---|---|
| 28.2 bits(14) | 256() | 14/14(100%) | 0/14(0%) | Plus/Plus | |

Features:
**uncharacterized protein LOC105373133 isoform X2uncharacterized protein LOC105373133 isoform X1**

```
Query  274       AACAAGCTGGTGTT  287
                 ||||||||||||||
Sbjct  13051220  AACAAGCTGGTGTT  13051233
```

Range 11: 22478394 to 22478407

| Score | Expect | Identities | Gaps | Strand | Frame |
|---|---|---|---|---|---|
| 28.2 bits(14) | 256() | 14/14(100%) | 0/14(0%) | Plus/Plus | |

Features:
**204125 bp at 5' side: E3 ubiquitin-protein ligase ZNF645521651 bp at 3' side: DEAD box protein 53**

```
Query  283       GTGTTCTCTCCTTT  296
                 ||||||||||||||
Sbjct  22478394  GTGTTCTCTCCTTT  22478407
```

Range 12: 64481206 to 64481223

| Score | Expect | Identities | Gaps | Strand | Frame |
|---|---|---|---|---|---|
| 28.2 bits(14) | 256() | 17/18(94%) | 0/18(0%) | Plus/Plus | |

Features:
**85843 bp at 5' side: myotubularin-related protein 8436541 bp at 3' side: zinc finger C4H2 domain-containing protein isoform 3**

```
Query  281       TGGTGTTCTCTCCTTTAT  298
                 ||||||| ||||||||||
Sbjct  64481206  TGGTGTTATCTCCTTTAT  64481223
```

Range 13: 67684094 to 67684107

| Score | Expect | Identities | Gaps | Strand | Frame |
|---|---|---|---|---|---|
| 28.2 bits(14) | 256() | 14/14(100%) | 0/14(0%) | Plus/Plus | |

Features:
**androgen receptor isoform 1androgen receptor isoform 2**

```
Query  271       TGTAACAAGCTGGT  284
                 ||||||||||||||
Sbjct  67684094  TGTAACAAGCTGGT  67684107
```

Range 14: 73479191 to 73479204

| Score | Expect | Identities | Gaps | Strand | Frame |
|---|---|---|---|---|---|
| 28.2 bits(14) | 256() | 14/14(100%) | 0/14(0%) | Plus/Plus | |

Features:
**24606 bp at 5' side: homeobox protein CDX-484081 bp at 3' side: cysteine-rich hydrophobic domain-containing protein 1 iso...**

```
Query  286       TTCTCTCCTTTATT  299
                 ||||||||||||||
Sbjct  73479191  TTCTCTCCTTTATT  73479204
```

Range 15: 74761604 to 74761617

| Score | Expect | Identities | Gaps | Strand | Frame |
|---|---|---|---|---|---|
| 28.2 bits(14) | 256() | 14/14(100%) | 0/14(0%) | Plus/Plus | |

Features:
**15954 bp at 5' side: protein KIAA2022 isoform X1291753 bp at 3' side: ATP-binding cassette sub-family B member 7, mitochondrial...**

```
Query  286       TTCTCTCCTTTATT  299
                 ||||||||||||||
Sbjct  74761604  TTCTCTCCTTTATT  74761617
```

Range 16: 78653330 to 78653343

| Score | Expect | Identities | Gaps | Strand | Frame |
|---|---|---|---|---|---|

| | | | | |
|---|---|---|---|---|
| 28.2 bits(14) | 256() | 14/14(100%) | 0/14(0%) | Plus/Plus |

Features:
**379584 bp at 5' side: cysteinyl leukotriene receptor 13650 bp at 3' side: zinc finger CCHC domain-containing protein 5**

```
Query  283       GTGTTCTCTCCTTT  296
                 ||||||||||||||
Sbjct  78653330  GTGTTCTCTCCTTT  78653343
```

Range 17: 108367366 to 108367379

| Score | Expect | Identities | Gaps | Strand | Frame |
|---|---|---|---|---|---|
| 28.2 bits(14) | 256() | 14/14(100%) | 0/14(0%) | Plus/Plus | |

Features:
**collagen alpha-6(IV) chain isoform X1collagen alpha-6(IV) chain isoform 3 precursor**

```
Query  279        GCTGGTGTTCTCTC  292
                  ||||||||||||||
Sbjct  108367366  GCTGGTGTTCTCTC  108367379
```

Range 18: 109296533 to 109296546

| Score | Expect | Identities | Gaps | Strand | Frame |
|---|---|---|---|---|---|
| 28.2 bits(14) | 256() | 14/14(100%) | 0/14(0%) | Plus/Plus | |

Features:
**560189 bp at 5' side: insulin receptor substrate 479353 bp at 3' side: retinal guanylyl cyclase 2**

```
Query  270        GTGTAACAAGCTGG  283
                  ||||||||||||||
Sbjct  109296533  GTGTAACAAGCTGG  109296546
```

Range 19: 125705608 to 125705621

| Score | Expect | Identities | Gaps | Strand | Frame |
|---|---|---|---|---|---|
| 28.2 bits(14) | 256() | 14/14(100%) | 0/14(0%) | Plus/Plus | |

Features:
**382507 bp at 5' side: uncharacterized protein LOC100129520458912 bp at 3' side: DDB1- and CUL4-associated factor 12-like protein 2**

```
Query  276        CAAGCTGGTGTTCT  289
                  ||||||||||||||
Sbjct  125705608  CAAGCTGGTGTTCT  125705621
```

Range 20: 141464546 to 141464559

| Score | Expect | Identities | Gaps | Strand | Frame |
|---|---|---|---|---|---|
| 28.2 bits(14) | 256() | 14/14(100%) | 0/14(0%) | Plus/Plus | |

Features:
**222089 bp at 5' side: sperm protein associated with the nucleus on the X chromo...119179 bp at 3' side: sperm protein associated with the nucleus on the X chromo...**

```
Query  286        TTCTCTCCTTTATT  299
                  ||||||||||||||
Sbjct  141464546  TTCTCTCCTTTATT  141464559
```

Range 21: 141927580 to 141927593

| Score | Expect | Identities | Gaps | Strand | Frame |
|---|---|---|---|---|---|
| 28.2 bits(14) | 256() | 14/14(100%) | 0/14(0%) | Plus/Plus | |

Features:
**18747 bp at 5' side: melanoma-associated antigen C1 isoform X1275273 bp at 3' side: melanoma-associated antigen C2**

```
Query  282        GGTGTTCTCTCCTT  295
                  ||||||||||||||
Sbjct  141927580  GGTGTTCTCTCCTT  141927593
```

Range 22: 147482770 to 147482783

| Score | Expect | Identities | Gaps | Strand | Frame |
|---|---|---|---|---|---|
| 28.2 bits(14) | 256() | 14/14(100%) | 0/14(0%) | Plus/Plus | |

Features:
**211483 bp at 5' side: uncharacterized protein LOC105373347 isoform X2429397 bp at 3' side: fragile X mental retardation protein 1 isoform ISO12**

```
Query  286        TTCTCTCCTTTATT  299
                  ||||||||||||||
Sbjct  147482770  TTCTCTCCTTTATT  147482783
```

Range 23: 5932724 to 5932737

| Score | Expect | Identities | Gaps | Strand | Frame |
|---|---|---|---|---|---|
| 28.2 bits(14) | 256() | 14/14(100%) | 0/14(0%) | Plus/Minus | |

Features:
**neuroligin-4, X-linkedneuroligin-4, X-linked isoform X1**

```
Query  286      TTCTCTCCTTTATT  299
                ||||||||||||||
Sbjct  5932737  TTCTCTCCTTTATT  5932724
```

Range 24: 12453206 to 12453219

| Score | Expect | Identities | Gaps | Strand | Frame |
|---|---|---|---|---|---|
| 28.2 bits(14) | 256() | 14/14(100%) | 0/14(0%) | Plus/Minus | |

Features:
**FERM and PDZ domain-containing protein 4FERM and PDZ domain-containing protein 4 isoform X2**

```
Query  286       TTCTCTCCTTTATT  299
                 ||||||||||||||
Sbjct  12453219  TTCTCTCCTTTATT  12453206
```

Range 25: 12708672 to 12708689

| Score | Expect | Identities | Gaps | Strand | Frame |
|---|---|---|---|---|---|
| 28.2 bits(14) | 256() | 17/18(94%) | 0/18(0%) | Plus/Minus | |

Features:
**FERM and PDZ domain-containing protein 4FERM and PDZ domain-containing protein 4 isoform X2**

```
Query  279       GCTGGTGTTCTCTCCTTT  296
                 ||||||||| ||||||||
Sbjct  12708689  GCTGGTGTTGTCTCCTTT  12708672
```

Range 26: 14929813 to 14929826

| Score | Expect | Identities | Gaps | Strand | Frame |
|---|---|---|---|---|---|
| 28.2 bits(14) | 256() | 14/14(100%) | 0/14(0%) | Plus/Minus | |

Features:
**10004 bp at 5' side: motile sperm domain-containing protein 2 isoform 2314680 bp at 3' side: ankyrin repeat and SOCS box protein 9 isoform X3**

```
Query  286       TTCTCTCCTTTATT  299
                 ||||||||||||||
Sbjct  14929826  TTCTCTCCTTTATT  14929813
```

Range 27: 18139299 to 18139312

| Score | Expect | Identities | Gaps | Strand | Frame |
|---|---|---|---|---|---|
| 28.2 bits(14) | 256() | 14/14(100%) | 0/14(0%) | Plus/Minus | |

Features:
**337289 bp at 5' side: retinoic acid-induced protein 2 isoform X125697 bp at 3' side: BEN domain-containing protein 2 isoform 1**

```
Query  286       TTCTCTCCTTTATT  299
                 ||||||||||||||
Sbjct  18139312  TTCTCTCCTTTATT  18139299
```

Range 28: 18280011 to 18280024

| Score | Expect | Identities | Gaps | Strand | Frame |
|---|---|---|---|---|---|
| 28.2 bits(14) | 256() | 14/14(100%) | 0/14(0%) | Plus/Minus | |

Features:
**sex comb on midleg-like protein 2sex comb on midleg-like protein 2 isoform X1**

```
Query  278       AGCTGGTGTTCTCT  291
                 ||||||||||||||
Sbjct  18280024  AGCTGGTGTTCTCT  18280011
```

Range 29: 22950855 to 22950868

| Score | Expect | Identities | Gaps | Strand | Frame |
|---|---|---|---|---|---|
| 28.2 bits(14) | 256() | 14/14(100%) | 0/14(0%) | Plus/Minus | |

Features:
**676586 bp at 5' side: E3 ubiquitin-protein ligase ZNF64549190 bp at 3' side: DEAD box protein 53**

```
Query  272       GTAACAAGCTGGTG  285
                 ||||||||||||||
Sbjct  22950868  GTAACAAGCTGGTG  22950855
```

Range 30: 29756914 to 29756927

| Score | Expect | Identities | Gaps | Strand | Frame |
|---|---|---|---|---|---|
| 28.2 bits(14) | 256() | 14/14(100%) | 0/14(0%) | Plus/Minus | |

Features:
**interleukin-1 receptor accessory protein-like 1 precursorinterleukin-1 receptor accessory protein-like 1 isoform X1**

```
Query  281       TGGTGTTCTCTCCT  294
                 ||||||||||||||
Sbjct  29756927  TGGTGTTCTCTCCT  29756914
```

Range 31: 39357519 to 39357532

| Score | Expect | Identities | Gaps | Strand | Frame |
|---|---|---|---|---|---|
| 28.2 bits(14) | 256() | 14/14(100%) | 0/14(0%) | Plus/Minus | |

Features:
**552021 bp at 5' side: mid1-interacting protein 1694577 bp at 3' side: BCL-6 corepressor isoform X2**

```
Query  279       GCTGGTGTTCTCTC  292
                 ||||||||||||||
Sbjct  39357532  GCTGGTGTTCTCTC  39357519
```

Range 32: 45179825 to 45179838

| Score | Expect | Identities | Gaps | Strand | Frame |
|---|---|---|---|---|---|
| 28.2 bits(14) | 256() | 14/14(100%) | 0/14(0%) | Plus/Minus | |

Features:
**deleted in autism-related protein 1 isoform X2deleted in autism-related protein 1 isoform X1**

```
Query  286       TTCTCTCCTTTATT  299
                 ||||||||||||||
Sbjct  45179838  TTCTCTCCTTTATT  45179825
```

Range 33: 68252489 to 68252502

| Score | Expect | Identities | Gaps | Strand | Frame |
|---|---|---|---|---|---|
| 28.2 bits(14) | 256() | 14/14(100%) | 0/14(0%) | Plus/Minus | |

Features:
**oligophrenin-1 isoform X2oligophrenin-1 isoform X3**

```
Query  274       AACAAGCTGGTGTT  287
                 ||||||||||||||
Sbjct  68252502  AACAAGCTGGTGTT  68252489
```

Range 34: 75387607 to 75387620

| Score | Expect | Identities | Gaps | Strand | Frame |
|---|---|---|---|---|---|
| 28.2 bits(14) | 256() | 14/14(100%) | 0/14(0%) | Plus/Minus | |

Features:
**palmitoyltransferase ZDHHC15 isoform 1palmitoyltransferase ZDHHC15 isoform X1**

```
Query  286       TTCTCTCCTTTATT  299
                 ||||||||||||||
Sbjct  75387620  TTCTCTCCTTTATT  75387607
```

Range 35: 75888781 to 75888794

| Score | Expect | Identities | Gaps | Strand | Frame |
|---|---|---|---|---|---|
| 28.2 bits(14) | 256() | 14/14(100%) | 0/14(0%) | Plus/Minus | |

Features:
**103730 bp at 5' side: melanoma-associated antigen E2260358 bp at 3' side: uncharacterized protein LOC105373256**

```
Query  279       GCTGGTGTTCTCTC  292
                 ||||||||||||||
Sbjct  75888794  GCTGGTGTTCTCTC  75888781
```

Range 36: 80111957 to 80111970

| Score | Expect | Identities | Gaps | Strand | Frame |
|---|---|---|---|---|---|
| 28.2 bits(14) | 256() | 14/14(100%) | 0/14(0%) | Plus/Minus | |

Features:
**80846 bp at 5' side: T-box transcription factor TBX22 isoform X2330570 bp at 3' side: protein FAM46D**

```
Query  271       TGTAACAAGCTGGT  284
                 ||||||||||||||
Sbjct  80111970  TGTAACAAGCTGGT  80111957
```

Range 37: 94474966 to 94474979

| Score | Expect | Identities | Gaps | Strand | Frame |
|---|---|---|---|---|---|
| 28.2 bits(14) | 256() | 14/14(100%) | 0/14(0%) | Plus/Minus | |

Features:
**764800 bp at 5' side: protein FAM133A2210080 bp at 3' side: protein diaphanous homolog 2 isoform 156**

```
Query  286       TTCTCTCCTTTATT  299
                 ||||||||||||||
Sbjct  94474979  TTCTCTCCTTTATT  94474966
```

Range 38: 94775560 to 94775573

| Score | Expect | Identities | Gaps | Strand | Frame |
|---|---|---|---|---|---|
| 28.2 bits(14) | 256() | 14/14(100%) | 0/14(0%) | Plus/Minus | |

Features:
**1065394 bp at 5' side: protein FAM133A1909486 bp at 3' side: protein diaphanous homolog 2 isoform 156**

```
Query  273       TAACAAGCTGGTGT  286
                 ||||||||||||||
Sbjct  94775573  TAACAAGCTGGTGT  94775560
```

Range 39: 121840625 to 121840638

| Score | Expect | Identities | Gaps | Strand | Frame |
|---|---|---|---|---|---|
| 28.2 bits(14) | 256() | 14/14(100%) | 0/14(0%) | Plus/Minus | |

Features:
**791264 bp at 5' side: glutamate dehydrogenase 2, mitochondrial precursor1343898 bp at 3' side: glutamate receptor 3 isoform 3 precursor**

```
Query  273        TAACAAGCTGGTGT  286
                  ||||||||||||||
Sbjct  121840638  TAACAAGCTGGTGT  121840625
```

Homo sapiens chromosome 15, alternate assembly CHM1_1.1
Sequence ID: **ref|NC_018926.2|** Length: 102381530 Number of Matches: 26
Range 1: 38105572 to 38105596

| Score | Expect | Identities | Gaps | Strand | Frame |
|---|---|---|---|---|---|
| 34.2 bits(17) | 4.1() | 23/25(92%) | 0/25(0%) | Plus/Plus | |

Features:
**596914 bp at 5' side: homeobox protein Meis2 isoform h242217 bp at 3' side: transmembrane and coiled-coil domain-containing protein 5A**

```
Query  275       ACAAGCTGGTGTTCTCTCCTTTATT  299
                 |||||||||| | ||||||||||||
Sbjct  38105572  ACAAGCTGGTATCCTCTCCTTTATT  38105596
```

Range 2: 98380695 to 98380710

| Score | Expect | Identities | Gaps | Strand | Frame |
|---|---|---|---|---|---|
| 32.2 bits(16) | 16() | 16/16(100%) | 0/16(0%) | Plus/Minus | |

Features:
**24287 bp at 5' side: arrestin domain-containing protein 4443798 bp at 3' side: protein FAM169B**

```
Query  274       AACAAGCTGGTGTTCT  289
                 ||||||||||||||||
Sbjct  98380710  AACAAGCTGGTGTTCT  98380695
```

Range 3: 51134987 to 51135001

| Score | Expect | Identities | Gaps | Strand | Frame |
|---|---|---|---|---|---|
| 30.2 bits(15) | 65() | 15/15(100%) | 0/15(0%) | Plus/Plus | |

Features:
**signal peptide peptidase-like 2A precursor**

```
Query  275       ACAAGCTGGTGTTCT  289
                 |||||||||||||||
Sbjct  51134987  ACAAGCTGGTGTTCT  51135001
```

Range 4: 94209596 to 94209610

| Score | Expect | Identities | Gaps | Strand | Frame |
|---|---|---|---|---|---|
| 30.2 bits(15) | 65() | 15/15(100%) | 0/15(0%) | Plus/Plus | |

Features:
**618378 bp at 5' side: putative uncharacterized protein UNQ9370/PRO34162473303 bp at 3' side: multiple C2 and transmembrane domain-containing protein 2...**

```
Query  284       TGTTCTCTCCTTTAT  298
                 |||||||||||||||
Sbjct  94209596  TGTTCTCTCCTTTAT  94209610
```

Range 5: 63168238 to 63168252

| Score | Expect | Identities | Gaps | Strand | Frame |
|---|---|---|---|---|---|
| 30.2 bits(15) | 65() | 15/15(100%) | 0/15(0%) | Plus/Minus | |

Features:
**talin-2**

```
Query  282       GGTGTTCTCTCCTTT  296
                 |||||||||||||||
Sbjct  63168252  GGTGTTCTCTCCTTT  63168238
```

Range 6: 101670671 to 101670685

| Score | Expect | Identities | Gaps | Strand | Frame |
|---|---|---|---|---|---|
| 30.2 bits(15) | 65() | 15/15(100%) | 0/15(0%) | Plus/Minus | |

Features:
**U2 small nuclear ribonucleoprotein A'**

```
Query  286        TTCTCTCCTTTATTG  300
                  |||||||||||||||
Sbjct  101670685  TTCTCTCCTTTATTG  101670671
```

Range 7: 45758855 to 45758872

| Score | Expect | Identities | Gaps | Strand | Frame |
|---|---|---|---|---|---|
| 28.2 bits(14) | 256() | 17/18(94%) | 0/18(0%) | Plus/Plus | |

Features:
**72679 bp at 5' side: sodium/nucleoside cotransporter 213923 bp at 3' side: glycine amidinotransferase, mitochondrial precursor**

```
Query  275       ACAAGCTGGTGTTCTCTC  292
                 |||||| |||||||||||
Sbjct  45758855  ACAAGCAGGTGTTCTCTC  45758872
```

Range 8: 46530380 to 46530393

| Score | Expect | Identities | Gaps | Strand | Frame |
|---|---|---|---|---|---|
| 28.2 bits(14) | 256() | 14/14(100%) | 0/14(0%) | Plus/Plus | |

Features:
**429034 bp at 5' side: sulfide:quinone oxidoreductase, mitochondrial1639815 bp at 3' side: semaphorin-6D isoform 1 precursor**

```
Query  286       TTCTCTCCTTTATT  299
                 ||||||||||||||
Sbjct  46530380  TTCTCTCCTTTATT  46530393
```

Range 9: 52492480 to 52492493

| Score | Expect | Identities | Gaps | Strand | Frame |
|---|---|---|---|---|---|
| 28.2 bits(14) | 256() | 14/14(100%) | 0/14(0%) | Plus/Plus | |

Features:
**17290 bp at 5' side: mitogen-activated protein kinase 627860 bp at 3' side: bcl-2-like protein 10**

```
Query  284       TGTTCTCTCCTTTA  297
                 ||||||||||||||
Sbjct  52492480  TGTTCTCTCCTTTA  52492493
```

Range 10: 63109068 to 63109081

| Score | Expect | Identities | Gaps | Strand | Frame |
|---|---|---|---|---|---|
| 28.2 bits(14) | 256() | 14/14(100%) | 0/14(0%) | Plus/Plus | |

Features:
**talin-2**

```
Query  279       GCTGGTGTTCTCTC  292
                 ||||||||||||||
Sbjct  63109068  GCTGGTGTTCTCTC  63109081
```

Range 11: 67318106 to 67318119

| Score | Expect | Identities | Gaps | Strand | Frame |
|---|---|---|---|---|---|
| 28.2 bits(14) | 256() | 14/14(100%) | 0/14(0%) | Plus/Plus | |

Features:
**126157 bp at 5' side: mothers against decapentaplegic homolog 6158439 bp at 3' side: mothers against decapentaplegic homolog 3 isoform 1**

```
Query  287       TCTCTCCTTTATTG  300
                 ||||||||||||||
Sbjct  67318106  TCTCTCCTTTATTG  67318119
```

Range 12: 68748575 to 68748588

| Score | Expect | Identities | Gaps | Strand | Frame |
|---|---|---|---|---|---|
| 28.2 bits(14) | 256() | 14/14(100%) | 0/14(0%) | Plus/Plus | |

Features:
**integrin alpha-11 precursor**

```
Query  286       TTCTCTCCTTTATT  299
                 ||||||||||||||
Sbjct  68748575  TTCTCTCCTTTATT  68748588
```

Range 13: 69030123 to 69030136

| Score | Expect | Identities | Gaps | Strand | Frame |
|---|---|---|---|---|---|
| 28.2 bits(14) | 256() | 14/14(100%) | 0/14(0%) | Plus/Plus | |

Features:
**coronin-2B isoform 1**

```
Query  283       GTGTTCTCTCCTTT  296
                 ||||||||||||||
Sbjct  69030123  GTGTTCTCTCCTTT  69030136
```

Range 14: 71245933 to 71245946

| Score | Expect | Identities | Gaps | Strand | Frame |
|---|---|---|---|---|---|
| 28.2 bits(14) | 256() | 14/14(100%) | 0/14(0%) | Plus/Plus | |

Features:
**la-related protein 6 isoform 1**

```
Query  287       TCTCTCCTTTATTG  300
                 ||||||||||||||
Sbjct  71245933  TCTCTCCTTTATTG  71245946
```

Range 15: 80865360 to 80865373

| Score | Expect | Identities | Gaps | Strand | Frame |
|---|---|---|---|---|---|
| 28.2 bits(14) | 256() | 14/14(100%) | 0/14(0%) | Plus/Plus | |

Features:
**aryl hydrocarbon receptor nuclear translocator 2**

```
Query  286       TTCTCTCCTTTATT  299
                 ||||||||||||||
Sbjct  80865360  TTCTCTCCTTTATT  80865373
```

Range 16: 85505224 to 85505237

| Score | Expect | Identities | Gaps | Strand | Frame |
|---|---|---|---|---|---|
| 28.2 bits(14) | 256() | 14/14(100%) | 0/14(0%) | Plus/Plus | |

Features:
**high affinity cAMP-specific and IBMX-insensitive 3',5'-cy...high affinity cAMP-specific and IBMX-insensitive 3',5'-cy...**

```
Query  286       TTCTCTCCTTTATT  299
                 ||||||||||||||
```

```
Sbjct  85505224  TTCTCTCCTTTATT  85505237
```

Range 17: 90131694 to 90131707

| Score | Expect | Identities | Gaps | Strand | Frame |
|---|---|---|---|---|---|
| 28.2 bits(14) | 256() | 14/14(100%) | 0/14(0%) | Plus/Plus | |

Features:
**4250 bp at 5' side: WD repeat-containing protein 93 isoform 12496 bp at 3' side: mesoderm posterior protein 1**

```
Query  275       ACAAGCTGGTGTTC  288
                 ||||||||||||||
Sbjct  90131694  ACAAGCTGGTGTTC  90131707
```

Range 18: 91760624 to 91760637

| Score | Expect | Identities | Gaps | Strand | Frame |
|---|---|---|---|---|---|
| 28.2 bits(14) | 256() | 14/14(100%) | 0/14(0%) | Plus/Plus | |

Features:
**103398 bp at 5' side: putative protein TPRXL477741 bp at 3' side: solute carrier organic anion transporter family member 3A...**

```
Query  282       GGTGTTCTCTCCTT  295
                 ||||||||||||||
Sbjct  91760624  GGTGTTCTCTCCTT  91760637
```

Range 19: 44530103 to 44530116

| Score | Expect | Identities | Gaps | Strand | Frame |
|---|---|---|---|---|---|
| 28.2 bits(14) | 256() | 14/14(100%) | 0/14(0%) | Plus/Minus | |

Features:
**FERM domain-containing protein 5 isoform 2**

```
Query  286       TTCTCTCCTTTATT  299
                 ||||||||||||||
Sbjct  44530116  TTCTCTCCTTTATT  44530103
```

Range 20: 50944568 to 50944581

| Score | Expect | Identities | Gaps | Strand | Frame |
|---|---|---|---|---|---|
| 28.2 bits(14) | 256() | 14/14(100%) | 0/14(0%) | Plus/Minus | |

Features:
**inactive ubiquitin carboxyl-terminal hydrolase 50**

```
Query  286       TTCTCTCCTTTATT  299
                 ||||||||||||||
Sbjct  50944581  TTCTCTCCTTTATT  50944568
```

Range 21: 51572533 to 51572546

| Score | Expect | Identities | Gaps | Strand | Frame |
|---|---|---|---|---|---|
| 28.2 bits(14) | 256() | 14/14(100%) | 0/14(0%) | Plus/Minus | |

Features:
**66293 bp at 5' side: serine/arginine repetitive matrix protein 3-like isoform X148550 bp at 3' side: aromatase**

```
Query  286       TTCTCTCCTTTATT  299
                 ||||||||||||||
Sbjct  51572546  TTCTCTCCTTTATT  51572533
```

Range 22: 60726154 to 60726167

| Score | Expect | Identities | Gaps | Strand | Frame |
|---|---|---|---|---|---|
| 28.2 bits(14) | 256() | 14/14(100%) | 0/14(0%) | Plus/Minus | |

Features:

**309672 bp at 5' side: forkhead box protein B132096 bp at 3' side: annexin A2 isoform 2**

```
Query  280       CTGGTGTTCTCTCC  293
                 ||||||||||||||
Sbjct  60726167  CTGGTGTTCTCTCC  60726154
```

Range 23: 62322449 to 62322462

| Score | Expect | Identities | Gaps | Strand | Frame |
|---|---|---|---|---|---|
| 28.2 bits(14) | 256() | 14/14(100%) | 0/14(0%) | Plus/Minus | |

Features:
**vacuolar protein sorting-associated protein 13C isoform 1Avacuolar protein sorting-associated protein 13C isoform 2A**

```
Query  281       TGGTGTTCTCTCCT  294
                 ||||||||||||||
Sbjct  62322462  TGGTGTTCTCTCCT  62322449
```

Range 24: 88826197 to 88826214

| Score | Expect | Identities | Gaps | Strand | Frame |
|---|---|---|---|---|---|
| 28.2 bits(14) | 256() | 17/18(94%) | 0/18(0%) | Plus/Minus | |

Features:
**184959 bp at 5' side: NT-3 growth factor receptor isoform c precursor18423 bp at 3' side: 39S ribosomal protein L46, mitochondrial**

```
Query  279       GCTGGTGTTCTCTCCTTT  296
                 |||||||| |||||||||
Sbjct  88826214  GCTGGTGTCCTCTCCTTT  88826197
```

Range 25: 92303463 to 92303476

| Score | Expect | Identities | Gaps | Strand | Frame |
|---|---|---|---|---|---|
| 28.2 bits(14) | 256() | 14/14(100%) | 0/14(0%) | Plus/Minus | |

Features:
**solute carrier organic anion transporter family member 3A...solute carrier organic anion transporter family member 3A...**

```
Query  274       AACAAGCTGGTGTT  287
                 ||||||||||||||
Sbjct  92303476  AACAAGCTGGTGTT  92303463
```

Range 26: 101187435 to 101187448

| Score | Expect | Identities | Gaps | Strand | Frame |
|---|---|---|---|---|---|
| 28.2 bits(14) | 256() | 14/14(100%) | 0/14(0%) | Plus/Minus | |

Features:
**157787 bp at 5' side: ankyrin repeat and SOCS box protein 7 isoform 255148 bp at 3' side: uncharacterized protein LOC105369201 isoform X2**

```
Query  283        GTGTTCTCTCCTTT  296
                  ||||||||||||||
Sbjct  101187448  GTGTTCTCTCCTTT  101187435
```

Homo sapiens chromosome 22, alternate assembly CHM1_1.1

Sequence ID: **ref|NC_018933.2|** Length: 51262586 Number of Matches: 10

Range 1: 23793663 to 23793679

| Score | Expect | Identities | Gaps | Strand | Frame |
|---|---|---|---|---|---|
| 34.2 bits(17) | 4.1() | 17/17(100%) | 0/17(0%) | Plus/Plus | |

Features:
**123751 bp at 5' side: breakpoint cluster region protein isoform 2133935 bp at 3' side: immunoglobulin lambda-like polypeptide 1 isoform a precursor**

```
Query  279       GCTGGTGTTCTCTCCTT  295
                 |||||||||||||||||
Sbjct  23793663  GCTGGTGTTCTCTCCTT  23793679
```

Range 2: 27857565 to 27857578

| Score | Expect | Identities | Gaps | Strand | Frame |
|---|---|---|---|---|---|
| 28.2 bits(14) | 256() | 14/14(100%) | 0/14(0%) | Plus/Plus | |

Features:
**871522 bp at 5' side: beta-crystallin A4248593 bp at 3' side: probable tumor suppressor protein MN1**

```
Query  280       CTGGTGTTCTCTCC  293
                 ||||||||||||||
Sbjct  27857565  CTGGTGTTCTCTCC  27857578
```

Range 3: 31896996 to 31897009

| Score | Expect | Identities | Gaps | Strand | Frame |
|---|---|---|---|---|---|
| 28.2 bits(14) | 256() | 14/14(100%) | 0/14(0%) | Plus/Plus | |

Features:
**protein SFI1 homolog isoform aprotein SFI1 homolog isoform b**

```
Query  285       GTTCTCTCCTTTAT  298
                 ||||||||||||||
Sbjct  31896996  GTTCTCTCCTTTAT  31897009
```

Range 4: 36923947 to 36923960

| Score | Expect | Identities | Gaps | Strand | Frame |
|---|---|---|---|---|---|
| 28.2 bits(14) | 256() | 14/14(100%) | 0/14(0%) | Plus/Plus | |

Features:
**voltage-dependent calcium channel gamma-2 subunit**

```
Query  282       GGTGTTCTCTCCTT  295
                 ||||||||||||||
Sbjct  36923947  GGTGTTCTCTCCTT  36923960
```

Range 5: 18601797 to 18601810

| Score | Expect | Identities | Gaps | Strand | Frame |
|---|---|---|---|---|---|
| 28.2 bits(14) | 256() | 14/14(100%) | 0/14(0%) | Plus/Minus | |

Features:
**tubulin alpha-8 chain isoform 1**

```
Query  287       TCTCTCCTTTATTG  300
                 ||||||||||||||
Sbjct  18601810  TCTCTCCTTTATTG  18601797
```

Range 6: 35991216 to 35991229

| Score | Expect | Identities | Gaps | Strand | Frame |
|---|---|---|---|---|---|
| 28.2 bits(14) | 256() | 14/14(100%) | 0/14(0%) | Plus/Minus | |

Features:
**19076 bp at 5' side: myoglobin20120 bp at 3' side: apolipoprotein L6**

```
Query  285       GTTCTCTCCTTTAT  298
                 ||||||||||||||
Sbjct  35991229  GTTCTCTCCTTTAT  35991216
```

Range 7: 42472887 to 42472900

| Score | Expect | Identities | Gaps | Strand | Frame |
|---|---|---|---|---|---|
| 28.2 bits(14) | 256() | 14/14(100%) | 0/14(0%) | Plus/Minus | |

Features:
**25929 bp at 5' side: NADH dehydrogenase [ubiquinone] 1 alpha subcomplex subunit 69417 bp at 3' side: cytochrome P450**

**2D6 isoform 2**

```
Query  278       AGCTGGTGTTCTCT  291
                 ||||||||||||||
Sbjct  42472900  AGCTGGTGTTCTCT  42472887
```

Range 8: 45416645 to 45416658

| Score | Expect | Identities | Gaps | Strand | Frame |
|---|---|---|---|---|---|
| 28.2 bits(14) | 256() | 14/14(100%) | 0/14(0%) | Plus/Minus | |

Features:
**145695 bp at 5' side: PHD finger protein 21B isoform 4106138 bp at 3' side: nuclear pore complex protein Nup50 isoform b**

```
Query  280       CTGGTGTTCTCTCC  293
                 ||||||||||||||
Sbjct  45416658  CTGGTGTTCTCTCC  45416645
```

Range 9: 47963135 to 47963148

| Score | Expect | Identities | Gaps | Strand | Frame |
|---|---|---|---|---|---|
| 28.2 bits(14) | 256() | 14/14(100%) | 0/14(0%) | Plus/Minus | |

Features:
**435188 bp at 5' side: TBC1 domain family member 22A isoform d880979 bp at 3' side: protein FAM19A5 isoform 1**

```
Query  280       CTGGTGTTCTCTCC  293
                 ||||||||||||||
Sbjct  47963148  CTGGTGTTCTCTCC  47963135
```

Range 10: 49569555 to 49569568

| Score | Expect | Identities | Gaps | Strand | Frame |
|---|---|---|---|---|---|
| 28.2 bits(14) | 256() | 14/14(100%) | 0/14(0%) | Plus/Minus | |

Features:
**465501 bp at 5' side: protein FAM19A5 isoform 2 precursor403457 bp at 3' side: uncharacterized protein C22orf34**

```
Query  277       AAGCTGGTGTTCTC  290
                 ||||||||||||||
Sbjct  49569568  AAGCTGGTGTTCTC  49569555
```

Homo sapiens chromosome 15, GRCh38.p2 Primary Assembly
Sequence ID: **ref|NC_000015.10|** Length: 101991189 Number of Matches: 27
Range 1: 37694082 to 37694106

| Score | Expect | Identities | Gaps | Strand | Frame |
|---|---|---|---|---|---|
| 34.2 bits(17) | 4.1() | 23/25(92%) | 0/25(0%) | Plus/Plus | |

Features:
**599505 bp at 5' side: homeobox protein Meis2 isoform X7242218 bp at 3' side: transmembrane and coiled-coil domain-containing protein 5...**

```
Query  275       ACAAGCTGGTGTTCTCTCCTTTATT  299
                 |||||||||| | ||||||||||||
Sbjct  37694082  ACAAGCTGGTATCCTCTCCTTTATT  37694106
```

Range 2: 97995480 to 97995495

| Score | Expect | Identities | Gaps | Strand | Frame |
|---|---|---|---|---|---|
| 32.2 bits(16) | 16() | 16/16(100%) | 0/16(0%) | Plus/Minus | |

Features:
**24293 bp at 5' side: arrestin domain-containing protein 4444136 bp at 3' side: protein FAM169B**

```
Query  274       AACAAGCTGGTGTTCT  289
                 ||||||||||||||||
Sbjct  97995495  AACAAGCTGGTGTTCT  97995480
```

Range 3: 50724794 to 50724808

| Score | Expect | Identities | Gaps | Strand | Frame |
|---|---|---|---|---|---|
| 30.2 bits(15) | 65() | 15/15(100%) | 0/15(0%) | Plus/Plus | |

Features:
**signal peptide peptidase-like 2A isoform X3signal peptide peptidase-like 2A isoform X1**

```
Query  275       ACAAGCTGGTGTTCT  289
                 |||||||||||||||
Sbjct  50724794  ACAAGCTGGTGTTCT  50724808
```

Range 4: 93825364 to 93825378

| Score | Expect | Identities | Gaps | Strand | Frame |
|---|---|---|---|---|---|
| 30.2 bits(15) | 65() | 15/15(100%) | 0/15(0%) | Plus/Plus | |

Features:
**618625 bp at 5' side: putative uncharacterized protein UNQ9370/PRO34162472888 bp at 3' side: multiple C2 and transmembrane domain-containing protein 2...**

```
Query  284       TGTTCTCTCCTTTAT  298
                 |||||||||||||||
Sbjct  93825364  TGTTCTCTCCTTTAT  93825378
```

Range 5: 62756795 to 62756809

| Score | Expect | Identities | Gaps | Strand | Frame |
|---|---|---|---|---|---|
| 30.2 bits(15) | 65() | 15/15(100%) | 0/15(0%) | Plus/Minus | |

Features:
**talin-2 isoform X2talin-2**

```
Query  282       GGTGTTCTCTCCTTT  296
                 |||||||||||||||
Sbjct  62756809  GGTGTTCTCTCCTTT  62756795
```

Range 6: 101289480 to 101289494

| Score | Expect | Identities | Gaps | Strand | Frame |
|---|---|---|---|---|---|
| 30.2 bits(15) | 65() | 15/15(100%) | 0/15(0%) | Plus/Minus | |

Features:
**U2 small nuclear ribonucleoprotein A'**

```
Query  286        TTCTCTCCTTTATTG  300
                  |||||||||||||||
Sbjct  101289494  TTCTCTCCTTTATTG  101289480
```

Range 7: 45348157 to 45348174

| Score | Expect | Identities | Gaps | Strand | Frame |
|---|---|---|---|---|---|
| 28.2 bits(14) | 256() | 17/18(94%) | 0/18(0%) | Plus/Plus | |

Features:
**72644 bp at 5' side: sodium/nucleoside cotransporter 2 isoform X513935 bp at 3' side: glycine amidinotransferase, mitochondrial isoform X1**

```
Query  275       ACAAGCTGGTGTTCTCTC  292
                 |||||| |||||||||||
Sbjct  45348157  ACAAGCAGGTGTTCTCTC  45348174
```

Range 8: 46120068 to 46120081

| Score | Expect | Identities | Gaps | Strand | Frame |
|---|---|---|---|---|---|
| 28.2 bits(14) | 256() | 14/14(100%) | 0/14(0%) | Plus/Plus | |

Features:
**429038 bp at 5' side: sulfide:quinone oxidoreductase, mitochondrial1639718 bp at 3' side: semaphorin-6D isoform X8**

```
Query  286       TTCTCTCCTTTATT  299
                 ||||||||||||||
Sbjct  46120068  TTCTCTCCTTTATT  46120081
```

Range 9: 52082289 to 52082302

| Score | Expect | Identities | Gaps | Strand | Frame |
|---|---|---|---|---|---|
| 28.2 bits(14) | 256() | 14/14(100%) | 0/14(0%) | Plus/Plus | |

Features:
**17289 bp at 5' side: mitogen-activated protein kinase 627505 bp at 3' side: bcl-2-like protein 10 isoform X1**

```
Query  284       TGTTCTCTCCTTTA  297
                 ||||||||||||||
Sbjct  52082289  TGTTCTCTCCTTTA  52082302
```

Range 10: 62697648 to 62697661

| Score | Expect | Identities | Gaps | Strand | Frame |
|---|---|---|---|---|---|
| 28.2 bits(14) | 256() | 14/14(100%) | 0/14(0%) | Plus/Plus | |

Features:
**talin-2 isoform X2talin-2**

```
Query  279       GCTGGTGTTCTCTC  292
                 ||||||||||||||
Sbjct  62697648  GCTGGTGTTCTCTC  62697661
```

Range 11: 66907535 to 66907548

| Score | Expect | Identities | Gaps | Strand | Frame |
|---|---|---|---|---|---|
| 28.2 bits(14) | 256() | 14/14(100%) | 0/14(0%) | Plus/Plus | |

Features:
**126000 bp at 5' side: mothers against decapentaplegic homolog 6 isoform X1158607 bp at 3' side: mothers against decapentaplegic homolog 3 isoform 1**

```
Query  287       TCTCTCCTTTATTG  300
                 ||||||||||||||
Sbjct  66907535  TCTCTCCTTTATTG  66907548
```

Range 12: 68338276 to 68338289

| Score | Expect | Identities | Gaps | Strand | Frame |
|---|---|---|---|---|---|
| 28.2 bits(14) | 256() | 14/14(100%) | 0/14(0%) | Plus/Plus | |

Features:
**integrin alpha-11 isoform X1integrin alpha-11 precursor**

```
Query  286       TTCTCTCCTTTATT  299
                 ||||||||||||||
Sbjct  68338276  TTCTCTCCTTTATT  68338289
```

Range 13: 68619836 to 68619849

| Score | Expect | Identities | Gaps | Strand | Frame |
|---|---|---|---|---|---|
| 28.2 bits(14) | 256() | 14/14(100%) | 0/14(0%) | Plus/Plus | |

Features:
**coronin-2B isoform X2coronin-2B isoform 1**

```
Query  283       GTGTTCTCTCCTTT  296
                 ||||||||||||||
Sbjct  68619836  GTGTTCTCTCCTTT  68619849
```

Range 14: 70835340 to 70835353

| Score | Expect | Identities | Gaps | Strand | Frame |
|---|---|---|---|---|---|

| | | | | |
|---|---|---|---|---|
| 28.2 bits(14) | 256() | 14/14(100%) | 0/14(0%) | Plus/Plus |

Features:
**la-related protein 6 isoform 1**

```
Query  287       TCTCTCCTTTATTG  300
                 ||||||||||||||
Sbjct  70835340  TCTCTCCTTTATTG  70835353
```

Range 15: 80455000 to 80455013

| Score | Expect | Identities | Gaps | Strand | Frame |
|---|---|---|---|---|---|
| 28.2 bits(14) | 256() | 14/14(100%) | 0/14(0%) | Plus/Plus | |

Features:
**aryl hydrocarbon receptor nuclear translocator 2**

```
Query  286       TTCTCTCCTTTATT  299
                 ||||||||||||||
Sbjct  80455000  TTCTCTCCTTTATT  80455013
```

Range 16: 85120452 to 85120465

| Score | Expect | Identities | Gaps | Strand | Frame |
|---|---|---|---|---|---|
| 28.2 bits(14) | 256() | 14/14(100%) | 0/14(0%) | Plus/Plus | |

Features:
**high affinity cAMP-specific and IBMX-insensitive 3',5'-cy...high affinity cAMP-specific and IBMX-insensitive 3',5'-cy...**

```
Query  286       TTCTCTCCTTTATT  299
                 ||||||||||||||
Sbjct  85120452  TTCTCTCCTTTATT  85120465
```

Range 17: 89747635 to 89747648

| Score | Expect | Identities | Gaps | Strand | Frame |
|---|---|---|---|---|---|
| 28.2 bits(14) | 256() | 14/14(100%) | 0/14(0%) | Plus/Plus | |

Features:
**4063 bp at 5' side: WD repeat-containing protein 93 isoform X42496 bp at 3' side: mesoderm posterior protein 1**

```
Query  275       ACAAGCTGGTGTTC  288
                 ||||||||||||||
Sbjct  89747635  ACAAGCTGGTGTTC  89747648
```

Range 18: 91375756 to 91375769

| Score | Expect | Identities | Gaps | Strand | Frame |
|---|---|---|---|---|---|
| 28.2 bits(14) | 256() | 14/14(100%) | 0/14(0%) | Plus/Plus | |

Features:
**103389 bp at 5' side: putative protein TPRXL478140 bp at 3' side: solute carrier organic anion transporter family member 3A...**

```
Query  282       GGTGTTCTCTCCTT  295
                 ||||||||||||||
Sbjct  91375756  GGTGTTCTCTCCTT  91375769
```

Range 19: 44119623 to 44119636

| Score | Expect | Identities | Gaps | Strand | Frame |
|---|---|---|---|---|---|
| 28.2 bits(14) | 256() | 14/14(100%) | 0/14(0%) | Plus/Minus | |

Features:
**FERM domain-containing protein 5 isoform X2FERM domain-containing protein 5 isoform X1**

```
Query  286       TTCTCTCCTTTATT  299
                 ||||||||||||||
Sbjct  44119636  TTCTCTCCTTTATT  44119623
```

Range 20: 50534368 to 50534381

| Score | Expect | Identities | Gaps | Strand | Frame |
|---|---|---|---|---|---|
| 28.2 bits(14) | 256() | 14/14(100%) | 0/14(0%) | Plus/Minus | |

Features:
**inactive ubiquitin carboxyl-terminal hydrolase 50 isoform X2inactive ubiquitin carboxyl-terminal hydrolase 50 isoform X4**

```
Query  286       TTCTCTCCTTTATT  299
                 ||||||||||||||
Sbjct  50534381  TTCTCTCCTTTATT  50534368
```

Range 21: 51162241 to 51162254

| Score | Expect | Identities | Gaps | Strand | Frame |
|---|---|---|---|---|---|
| 28.2 bits(14) | 256() | 14/14(100%) | 0/14(0%) | Plus/Minus | |

Features:
**66326 bp at 5' side: uncharacterized protein LOC105370815 isoform X248554 bp at 3' side: aromatase isoform X1**

```
Query  286       TTCTCTCCTTTATT  299
                 ||||||||||||||
Sbjct  51162254  TTCTCTCCTTTATT  51162241
```

Range 22: 60315590 to 60315603

| Score | Expect | Identities | Gaps | Strand | Frame |
|---|---|---|---|---|---|
| 28.2 bits(14) | 256() | 14/14(100%) | 0/14(0%) | Plus/Minus | |

Features:
**309649 bp at 5' side: forkhead box protein B132027 bp at 3' side: annexin A2 isoform 1**

```
Query  280       CTGGTGTTCTCTCC  293
                 ||||||||||||||
Sbjct  60315603  CTGGTGTTCTCTCC  60315590
```

Range 23: 61910726 to 61910739

| Score | Expect | Identities | Gaps | Strand | Frame |
|---|---|---|---|---|---|
| 28.2 bits(14) | 256() | 14/14(100%) | 0/14(0%) | Plus/Minus | |

Features:
**vacuolar protein sorting-associated protein 13C isoform 1Avacuolar protein sorting-associated protein 13C isoform 2A**

```
Query  281       TGGTGTTCTCTCCT  294
                 ||||||||||||||
Sbjct  61910739  TGGTGTTCTCTCCT  61910726
```

Range 24: 88441173 to 88441190

| Score | Expect | Identities | Gaps | Strand | Frame |
|---|---|---|---|---|---|
| 28.2 bits(14) | 256() | 17/18(94%) | 0/18(0%) | Plus/Minus | |

Features:
**185020 bp at 5' side: NT-3 growth factor receptor isoform X1218423 bp at 3' side: 39S ribosomal protein L46, mitochondrial**

```
Query  279       GCTGGTGTTCTCTCCTTT  296
                 |||||||| |||||||||
Sbjct  88441190  GCTGGTGTCCTCTCCTTT  88441173
```

Range 25: 91919005 to 91919018

| Score | Expect | Identities | Gaps | Strand | Frame |
|---|---|---|---|---|---|
| 28.2 bits(14) | 256() | 14/14(100%) | 0/14(0%) | Plus/Minus | |

Features:
**solute carrier organic anion transporter family member 3A...solute carrier organic anion transporter family member 3A...**

```
Query  274       AACAAGCTGGTGTT  287
                 ||||||||||||||
```

```
Sbjct  91919018  AACAAGCTGGTGTT  91919005
```

Range 26: 92757919 to 92757936

| Score | Expect | Identities | Gaps | Strand | Frame |
|---|---|---|---|---|---|
| 28.2 bits(14) | 256() | 17/18(94%) | 0/18(0%) | Plus/Minus | |

Features:
**102260 bp at 5' side: membrane protein FAM174B precursor143302 bp at 3' side: chromodomain-helicase-DNA-binding protein 2 isoform 1**

```
Query  281       TGGTGTTCTCTCCTTTAT  298
                 |||||||||| |||||||
Sbjct  92757936  TGGTGTTCTCCCCTTTAT  92757919
```

Range 27: 100806303 to 100806316

| Score | Expect | Identities | Gaps | Strand | Frame |
|---|---|---|---|---|---|
| 28.2 bits(14) | 256() | 14/14(100%) | 0/14(0%) | Plus/Minus | |

Features:
**176253 bp at 5' side: ankyrin repeat and SOCS box protein 7 isoform 155100 bp at 3' side: uncharacterized protein LOC105369201 isoform X1**

```
Query  283        GTGTTCTCTCCTTT  296
                  ||||||||||||||
Sbjct  100806316  GTGTTCTCTCCTTT  100806303
```

Homo sapiens chromosome 22, GRCh38.p2 Primary Assembly

Sequence ID: **ref|NC_000022.11|** Length: 50818468 Number of Matches: 11

Range 1: 23439248 to 23439264

| Score | Expect | Identities | Gaps | Strand | Frame |
|---|---|---|---|---|---|
| 34.2 bits(17) | 4.1() | 17/17(100%) | 0/17(0%) | Plus/Plus | |

Features:
**123726 bp at 5' side: breakpoint cluster region protein isoform 1134002 bp at 3' side: immunoglobulin lambda-like polypeptide 1 isoform a precursor**

```
Query  279       GCTGGTGTTCTCTCCTT  295
                 |||||||||||||||||
Sbjct  23439248  GCTGGTGTTCTCTCCTT  23439264
```

Range 2: 10695252 to 10695265

| Score | Expect | Identities | Gaps | Strand | Frame |
|---|---|---|---|---|---|
| 28.2 bits(14) | 256() | 14/14(100%) | 0/14(0%) | Plus/Plus | |

Features:
**9288 bp at 5' side: Ig kappa chain V-I region Walker-like242630 bp at 3' side: protein FRG1-like**

```
Query  286       TTCTCTCCTTTATT  299
                 ||||||||||||||
Sbjct  10695252  TTCTCTCCTTTATT  10695265
```

Range 3: 27502880 to 27502893

| Score | Expect | Identities | Gaps | Strand | Frame |
|---|---|---|---|---|---|
| 28.2 bits(14) | 256() | 14/14(100%) | 0/14(0%) | Plus/Plus | |

Features:
**872393 bp at 5' side: beta-crystallin A4248022 bp at 3' side: probable tumor suppressor protein MN1**

```
Query  280       CTGGTGTTCTCTCC  293
                 ||||||||||||||
Sbjct  27502880  CTGGTGTTCTCTCC  27502893
```

Range 4: 31541238 to 31541251

| Score | Expect | Identities | Gaps | Strand | Frame |
|---|---|---|---|---|---|
| 28.2 bits(14) | 256() | 14/14(100%) | 0/14(0%) | Plus/Plus | |

Features:
**protein SFI1 homolog isoform X1protein SFI1 homolog isoform X6**

```
Query  285       GTTCTCTCCTTTAT  298
                 ||||||||||||||
Sbjct  31541238  GTTCTCTCCTTTAT  31541251
```

Range 5: 36568968 to 36568981

| Score | Expect | Identities | Gaps | Strand | Frame |
|---|---|---|---|---|---|
| 28.2 bits(14) | 256() | 14/14(100%) | 0/14(0%) | Plus/Plus | |

Features:
**voltage-dependent calcium channel gamma-2 subunit**

```
Query  282       GGTGTTCTCTCCTT  295
                 ||||||||||||||
Sbjct  36568968  GGTGTTCTCTCCTT  36568981
```

Range 6: 18119313 to 18119326

| Score | Expect | Identities | Gaps | Strand | Frame |
|---|---|---|---|---|---|
| 28.2 bits(14) | 256() | 14/14(100%) | 0/14(0%) | Plus/Minus | |

Features:
**tubulin alpha-8 chain isoform 1**

```
Query  287       TCTCTCCTTTATTG  300
                 ||||||||||||||
Sbjct  18119326  TCTCTCCTTTATTG  18119313
```

Range 7: 35636291 to 35636304

| Score | Expect | Identities | Gaps | Strand | Frame |
|---|---|---|---|---|---|
| 28.2 bits(14) | 256() | 14/14(100%) | 0/14(0%) | Plus/Minus | |

Features:
**19034 bp at 5' side: myoglobin20122 bp at 3' side: apolipoprotein L6**

```
Query  285       GTTCTCTCCTTTAT  298
                 ||||||||||||||
Sbjct  35636304  GTTCTCTCCTTTAT  35636291
```

Range 8: 42117123 to 42117136

| Score | Expect | Identities | Gaps | Strand | Frame |
|---|---|---|---|---|---|
| 28.2 bits(14) | 256() | 14/14(100%) | 0/14(0%) | Plus/Minus | |

Features:
**26301 bp at 5' side: NADH dehydrogenase [ubiquinone] 1 alpha subcomplex subunit 68761 bp at 3' side: cytochrome P450 2D6 isoform X1**

```
Query  278       AGCTGGTGTTCTCT  291
                 ||||||||||||||
Sbjct  42117136  AGCTGGTGTTCTCT  42117123
```

Range 9: 45062041 to 45062054

| Score | Expect | Identities | Gaps | Strand | Frame |
|---|---|---|---|---|---|
| 28.2 bits(14) | 256() | 14/14(100%) | 0/14(0%) | Plus/Minus | |

Features:
**52492 bp at 5' side: PHD finger protein 21B isoform X2106124 bp at 3' side: nuclear pore complex protein Nup50 isoform b**

```
Query  280       CTGGTGTTCTCTCC  293
                 ||||||||||||||
```

```
Sbjct  45062054  CTGGTGTTCTCTCC  45062041
```

Range 10: 47608617 to 47608630

| Score | Expect | Identities | Gaps | Strand | Frame |
|---|---|---|---|---|---|
| 28.2 bits(14) | 256() | 14/14(100%) | 0/14(0%) | Plus/Minus | |

Features:
**434991 bp at 5' side: TBC1 domain family member 22A isoform d880963 bp at 3' side: protein FAM19A5 isoform 1**

```
Query  280       CTGGTGTTCTCTCC  293
                 ||||||||||||||
Sbjct  47608630  CTGGTGTTCTCTCC  47608617
```

Range 11: 49214864 to 49214877

| Score | Expect | Identities | Gaps | Strand | Frame |
|---|---|---|---|---|---|
| 28.2 bits(14) | 256() | 14/14(100%) | 0/14(0%) | Plus/Minus | |

Features:
**465017 bp at 5' side: protein FAM19A5 isoform 2 precursor403038 bp at 3' side: uncharacterized protein C22orf34 isoform X2**

```
Query  277       AAGCTGGTGTTCTC  290
                 ||||||||||||||
Sbjct  49214877  AAGCTGGTGTTCTC  49214864
```

Homo sapiens chromosome 22 genomic scaffold, GRCh38.p2 alternate locus group ALT_REF_LOCI_1 HSCHR22_1_CTG6
Sequence ID: **ref|NT_187632.1|** Length: 186262 Number of Matches: 1
Range 1: 85206 to 85222

| Score | Expect | Identities | Gaps | Strand | Frame |
|---|---|---|---|---|---|
| 34.2 bits(17) | 4.1() | 17/17(100%) | 0/17(0%) | Plus/Plus | |

Features:

```
Query  279    GCTGGTGTTCTCTCCTT  295
              |||||||||||||||||
Sbjct  85206  GCTGGTGTTCTCTCCTT  85222
```

Homo sapiens chromosome 2, alternate assembly CHM1_1.1
Sequence ID: **ref|NC_018913.2|** Length: 243205335 Number of Matches: 73
Range 1: 195000594 to 195000610

| Score | Expect | Identities | Gaps | Strand | Frame |
|---|---|---|---|---|---|
| 34.2 bits(17) | 4.1() | 17/17(100%) | 0/17(0%) | Plus/Minus | |

Features:
**1935208 bp at 5' side: tomoregulin-2 precursor1550419 bp at 3' side: zinc transporter ZIP10 precursor**

```
Query  273        TAACAAGCTGGTGTTCT  289
                  |||||||||||||||||
Sbjct  195000610  TAACAAGCTGGTGTTCT  195000594
```

Range 2: 5481779 to 5481794

| Score | Expect | Identities | Gaps | Strand | Frame |
|---|---|---|---|---|---|
| 32.2 bits(16) | 16() | 16/16(100%) | 0/16(0%) | Plus/Minus | |

Features:
**1395595 bp at 5' side: uncharacterized protein LOC105373397280354 bp at 3' side: transcription factor SOX-11**

```
Query  275      ACAAGCTGGTGTTCTC  290
                ||||||||||||||||
Sbjct  5481794  ACAAGCTGGTGTTCTC  5481779
```

Range 3: 20095210 to 20095225

| Score | Expect | Identities | Gaps | Strand | Frame |
|---|---|---|---|---|---|
| 32.2 bits(16) | 16() | 16/16(100%) | 0/16(0%) | Plus/Minus | |

Features:
**WD repeat-containing protein 35 isoform 2WD repeat-containing protein 35 isoform 1**

```
Query  279       GCTGGTGTTCTCTCCT  294
                 ||||||||||||||||
Sbjct  20095225  GCTGGTGTTCTCTCCT  20095210
```

Range 4: 17137711 to 17137725

| Score | Expect | Identities | Gaps | Strand | Frame |
|---|---|---|---|---|---|
| 30.2 bits(15) | 65() | 15/15(100%) | 0/15(0%) | Plus/Plus | |

Features:
**438871 bp at 5' side: protein FAM49A483742 bp at 3' side: RAD51-associated protein 2**

```
Query  284       TGTTCTCTCCTTTAT  298
                 |||||||||||||||
Sbjct  17137711  TGTTCTCTCCTTTAT  17137725
```

Range 5: 48819019 to 48819033

| Score | Expect | Identities | Gaps | Strand | Frame |
|---|---|---|---|---|---|
| 30.2 bits(15) | 65() | 15/15(100%) | 0/15(0%) | Plus/Plus | |

Features:
**STON1-GTF2A1L protein isoform 2STON1-GTF2A1L protein isoform 1**

```
Query  284       TGTTCTCTCCTTTAT  298
                 |||||||||||||||
Sbjct  48819019  TGTTCTCTCCTTTAT  48819033
```

Range 6: 70616196 to 70616214

| Score | Expect | Identities | Gaps | Strand | Frame |
|---|---|---|---|---|---|
| 30.2 bits(15) | 65() | 18/19(95%) | 0/19(0%) | Plus/Plus | |

Features:
**protransforming growth factor alpha isoform 1 preproproteinprotransforming growth factor alpha isoform 2 preproprotein**

```
Query  278       AGCTGGTGTTCTCTCCTTT  296
                 ||||| |||||||||||||
Sbjct  70616196  AGCTGCTGTTCTCTCCTTT  70616214
```

Range 7: 222884214 to 222884228

| Score | Expect | Identities | Gaps | Strand | Frame |
|---|---|---|---|---|---|
| 30.2 bits(15) | 65() | 15/15(100%) | 0/15(0%) | Plus/Plus | |

Features:
**441462 bp at 5' side: ephrin type-A receptor 4 isoform b188154 bp at 3' side: paired box protein Pax-3 isoform PAX3e**

```
Query  280        CTGGTGTTCTCTCCT  294
                  |||||||||||||||
Sbjct  222884214  CTGGTGTTCTCTCCT  222884228
```

Range 8: 85420693 to 85420707

| Score | Expect | Identities | Gaps | Strand | Frame |
|---|---|---|---|---|---|
| 30.2 bits(15) | 65() | 15/15(100%) | 0/15(0%) | Plus/Minus | |

Features:
**transcription factor 7-like 1**

```
Query  277       AAGCTGGTGTTCTCT  291
                 |||||||||||||||
```

```
Sbjct  85420707  AAGCTGGTGTTCTCT  85420693
```

Range 9: 128535254 to 128535268

| Score | Expect | Identities | Gaps | Strand | Frame |
|---|---|---|---|---|---|
| 30.2 bits(15) | 65() | 15/15(100%) | 0/15(0%) | Plus/Minus | |

Features:
**2198 bp at 5' side: pre-mRNA 3' end processing protein WDR33 isoform 275402 bp at 3' side: DNA-directed RNA polymerase II subunit RPB4**

```
Query  286        TTCTCTCCTTTATTG  300
                  |||||||||||||||
Sbjct  128535268  TTCTCTCCTTTATTG  128535254
```

Range 10: 141632452 to 141632466

| Score | Expect | Identities | Gaps | Strand | Frame |
|---|---|---|---|---|---|
| 30.2 bits(15) | 65() | 15/15(100%) | 0/15(0%) | Plus/Minus | |

Features:
**low-density lipoprotein receptor-related protein 1B precu...**

```
Query  286        TTCTCTCCTTTATTG  300
                  |||||||||||||||
Sbjct  141632466  TTCTCTCCTTTATTG  141632452
```

Range 11: 151035460 to 151035474

| Score | Expect | Identities | Gaps | Strand | Frame |
|---|---|---|---|---|---|
| 30.2 bits(15) | 65() | 15/15(100%) | 0/15(0%) | Plus/Minus | |

Features:
**585933 bp at 5' side: methylmalonic aciduria and homocystinuria type D protein,...296812 bp at 3' side: rho-related GTP-binding protein RhoE precursor**

```
Query  286        TTCTCTCCTTTATTG  300
                  |||||||||||||||
Sbjct  151035474  TTCTCTCCTTTATTG  151035460
```

Range 12: 5423952 to 5423965

| Score | Expect | Identities | Gaps | Strand | Frame |
|---|---|---|---|---|---|
| 28.2 bits(14) | 256() | 14/14(100%) | 0/14(0%) | Plus/Plus | |

Features:
**1337768 bp at 5' side: uncharacterized protein LOC105373397338183 bp at 3' side: transcription factor SOX-11**

```
Query  286      TTCTCTCCTTTATT  299
                ||||||||||||||
Sbjct  5423952  TTCTCTCCTTTATT  5423965
```

Range 13: 22543097 to 22543110

| Score | Expect | Identities | Gaps | Strand | Frame |
|---|---|---|---|---|---|
| 28.2 bits(14) | 256() | 14/14(100%) | 0/14(0%) | Plus/Plus | |

Features:
**1247601 bp at 5' side: tudor domain-containing protein 151171327 bp at 3' side: kelch-like protein 29**

```
Query  283       GTGTTCTCTCCTTT  296
                 ||||||||||||||
Sbjct  22543097  GTGTTCTCTCCTTT  22543110
```

Range 14: 29727509 to 29727526

| Score | Expect | Identities | Gaps | Strand | Frame |
|---|---|---|---|---|---|
| 28.2 bits(14) | 256() | 17/18(94%) | 0/18(0%) | Plus/Plus | |

Features:
**ALK tyrosine kinase receptor precursor**

```
Query  279       GCTGGTGTTCTCTCCTTT  296
                 ||||||||||||| ||||
Sbjct  29727509  GCTGGTGTTCTCTGCTTT  29727526
```

Range 15: 40622470 to 40622483

| Score | Expect | Identities | Gaps | Strand | Frame |
|---|---|---|---|---|---|
| 28.2 bits(14) | 256() | 14/14(100%) | 0/14(0%) | Plus/Plus | |

Features:
**34695 bp at 5' side: sodium/calcium exchanger 1 isoform C precursor1472856 bp at 3' side: uncharacterized protein C2orf91**

```
Query  282       GGTGTTCTCTCCTT  295
                 ||||||||||||||
Sbjct  40622470  GGTGTTCTCTCCTT  40622483
```

Range 16: 70086566 to 70086583

| Score | Expect | Identities | Gaps | Strand | Frame |
|---|---|---|---|---|---|
| 28.2 bits(14) | 256() | 17/18(94%) | 0/18(0%) | Plus/Plus | |

Features:
**max dimerization protein 1 isoform 1max dimerization protein 1 isoform 3**

```
Query  281       TGGTGTTCTCTCCTTTAT  298
                 ||||||| ||||||||||
Sbjct  70086566  TGGTGTTATCTCCTTTAT  70086583
```

Range 17: 77480461 to 77480474

| Score | Expect | Identities | Gaps | Strand | Frame |
|---|---|---|---|---|---|
| 28.2 bits(14) | 256() | 14/14(100%) | 0/14(0%) | Plus/Plus | |

Features:
**leucine-rich repeat transmembrane neuronal protein 4 isof...leucine-rich repeat transmembrane neuronal protein 4 isof...**

```
Query  280       CTGGTGTTCTCTCC  293
                 ||||||||||||||
Sbjct  77480461  CTGGTGTTCTCTCC  77480474
```

Range 18: 79724556 to 79724569

| Score | Expect | Identities | Gaps | Strand | Frame |
|---|---|---|---|---|---|
| 28.2 bits(14) | 256() | 14/14(100%) | 0/14(0%) | Plus/Plus | |

Features:
**catenin alpha-2 isoform 4**

```
Query  283       GTGTTCTCTCCTTT  296
                 ||||||||||||||
Sbjct  79724556  GTGTTCTCTCCTTT  79724569
```

Range 19: 109983937 to 109983950

| Score | Expect | Identities | Gaps | Strand | Frame |
|---|---|---|---|---|---|
| 28.2 bits(14) | 256() | 14/14(100%) | 0/14(0%) | Plus/Plus | |

Features:
**SH3 domain-containing RING finger protein 3 precursor**

```
Query  286        TTCTCTCCTTTATT  299
                  ||||||||||||||
Sbjct  109983937  TTCTCTCCTTTATT  109983950
```

Range 20: 122762670 to 122762683

| Score | Expect | Identities | Gaps | Strand | Frame |
|---|---|---|---|---|---|
| 28.2 bits(14) | 256() | 14/14(100%) | 0/14(0%) | Plus/Plus | |

Features:
**236187 bp at 5' side: translin isoform 12024147 bp at 3' side: contactin-associated protein-like 5 precursor**

```
Query  279        GCTGGTGTTCTCTC  292
                  ||||||||||||||
Sbjct  122762670  GCTGGTGTTCTCTC  122762683
```

Range 21: 124639475 to 124639488

| Score | Expect | Identities | Gaps | Strand | Frame |
|---|---|---|---|---|---|
| 28.2 bits(14) | 256() | 14/14(100%) | 0/14(0%) | Plus/Plus | |

Features:
**2112992 bp at 5' side: translin isoform 1147342 bp at 3' side: contactin-associated protein-like 5 precursor**

```
Query  286        TTCTCTCCTTTATT  299
                  ||||||||||||||
Sbjct  124639475  TTCTCTCCTTTATT  124639488
```

Range 22: 126320303 to 126320316

| Score | Expect | Identities | Gaps | Strand | Frame |
|---|---|---|---|---|---|
| 28.2 bits(14) | 256() | 14/14(100%) | 0/14(0%) | Plus/Plus | |

Features:
**644865 bp at 5' side: contactin-associated protein-like 5 precursor977676 bp at 3' side: uncharacterized protein LOC105373602 isoform X1**

```
Query  284        TGTTCTCTCCTTTA  297
                  ||||||||||||||
Sbjct  126320303  TGTTCTCTCCTTTA  126320316
```

Range 23: 127035690 to 127035703

| Score | Expect | Identities | Gaps | Strand | Frame |
|---|---|---|---|---|---|
| 28.2 bits(14) | 256() | 14/14(100%) | 0/14(0%) | Plus/Plus | |

Features:
**1360252 bp at 5' side: contactin-associated protein-like 5 precursor262289 bp at 3' side: uncharacterized protein LOC105373602 isoform X1**

```
Query  282        GGTGTTCTCTCCTT  295
                  ||||||||||||||
Sbjct  127035690  GGTGTTCTCTCCTT  127035703
```

Range 24: 131893153 to 131893166

| Score | Expect | Identities | Gaps | Strand | Frame |
|---|---|---|---|---|---|
| 28.2 bits(14) | 256() | 14/14(100%) | 0/14(0%) | Plus/Plus | |

Features:
**pleckstrin homology domain-containing family B member 2 i...pleckstrin homology domain-containing family B member 2 i...**

```
Query  286        TTCTCTCCTTTATT  299
                  ||||||||||||||
Sbjct  131893153  TTCTCTCCTTTATT  131893166
```

Range 25: 138559588 to 138559601

| Score | Expect | Identities | Gaps | Strand | Frame |
|---|---|---|---|---|---|
| 28.2 bits(14) | 256() | 14/14(100%) | 0/14(0%) | Plus/Plus | |

Features:
**120870 bp at 5' side: thrombospondin type-1 domain-containing protein 7B166947 bp at 3' side: histamine N-methyltransferase isoform 3**

```
Query  283        GTGTTCTCTCCTTT  296
                  ||||||||||||||
Sbjct  138559588  GTGTTCTCTCCTTT  138559601
```

Range 26: 157092601 to 157092614

| Score | Expect | Identities | Gaps | Strand | Frame |
|---|---|---|---|---|---|
| 28.2 bits(14) | 256() | 14/14(100%) | 0/14(0%) | Plus/Plus | |

Features:
**1375100 bp at 5' side: G protein-activated inward rectifier potassium channel 1 ...95723 bp at 3' side: nuclear receptor subfamily 4 group A member 2**

```
Query  278        AGCTGGTGTTCTCT  291
                  ||||||||||||||
Sbjct  157092601  AGCTGGTGTTCTCT  157092614
```

Range 27: 159110703 to 159110716

| Score | Expect | Identities | Gaps | Strand | Frame |
|---|---|---|---|---|---|
| 28.2 bits(14) | 256() | 14/14(100%) | 0/14(0%) | Plus/Plus | |

Features:
**coiled-coil domain-containing protein 148 isoform 1coiled-coil domain-containing protein 148 isoform 3**

```
Query  286        TTCTCTCCTTTATT  299
                  ||||||||||||||
Sbjct  159110703  TTCTCTCCTTTATT  159110716
```

Range 28: 161332911 to 161332924

| Score | Expect | Identities | Gaps | Strand | Frame |
|---|---|---|---|---|---|
| 28.2 bits(14) | 256() | 14/14(100%) | 0/14(0%) | Plus/Plus | |

Features:
**102682 bp at 5' side: RNA-binding motif, single-stranded-interacting protein 1 ...709387 bp at 3' side: TRAF family member-associated NF-kappa-B activator isoform a**

```
Query  276        CAAGCTGGTGTTCT  289
                  ||||||||||||||
Sbjct  161332911  CAAGCTGGTGTTCT  161332924
```

Range 29: 161601840 to 161601853

| Score | Expect | Identities | Gaps | Strand | Frame |
|---|---|---|---|---|---|
| 28.2 bits(14) | 256() | 14/14(100%) | 0/14(0%) | Plus/Plus | |

Features:
**371611 bp at 5' side: RNA-binding motif, single-stranded-interacting protein 1 ...440458 bp at 3' side: TRAF family member-associated NF-kappa-B activator isoform a**

```
Query  286        TTCTCTCCTTTATT  299
                  ||||||||||||||
Sbjct  161601840  TTCTCTCCTTTATT  161601853
```

Range 30: 165798547 to 165798560

| Score | Expect | Identities | Gaps | Strand | Frame |
|---|---|---|---|---|---|
| 28.2 bits(14) | 256() | 14/14(100%) | 0/14(0%) | Plus/Plus | |

Features:
**putative sodium-coupled neutral amino acid transporter 11...putative sodium-coupled neutral amino acid transporter 11...**

```
Query  286        TTCTCTCCTTTATT  299
                  ||||||||||||||
Sbjct  165798547  TTCTCTCCTTTATT  165798560
```

Range 31: 182980680 to 182980693

| Score | Expect | Identities | Gaps | Strand | Frame |
|---|---|---|---|---|---|
| 28.2 bits(14) | 256() | 14/14(100%) | 0/14(0%) | Plus/Plus | |

Features:
**protein phosphatase 1 regulatory subunit 1C isoform 1protein phosphatase 1 regulatory subunit 1C isoform 2**

```
Query  276        CAAGCTGGTGTTCT  289
                  ||||||||||||||
Sbjct  182980680  CAAGCTGGTGTTCT  182980693
```

Range 32: 192223414 to 192223427

| Score | Expect | Identities | Gaps | Strand | Frame |
|---|---|---|---|---|---|
| 28.2 bits(14) | 256() | 14/14(100%) | 0/14(0%) | Plus/Plus | |

Features:
**unconventional myosin-Ib isoform 1unconventional myosin-Ib isoform 1**

```
Query  280        CTGGTGTTCTCTCC  293
                  ||||||||||||||
Sbjct  192223414  CTGGTGTTCTCTCC  192223427
```

Range 33: 201183365 to 201183378

| Score | Expect | Identities | Gaps | Strand | Frame |
|---|---|---|---|---|---|
| 28.2 bits(14) | 256() | 14/14(100%) | 0/14(0%) | Plus/Plus | |

Features:
**SPATS2-like protein isoform d**

```
Query  286        TTCTCTCCTTTATT  299
                  ||||||||||||||
Sbjct  201183365  TTCTCTCCTTTATT  201183378
```

Range 34: 212595034 to 212595047

| Score | Expect | Identities | Gaps | Strand | Frame |
|---|---|---|---|---|---|
| 28.2 bits(14) | 256() | 14/14(100%) | 0/14(0%) | Plus/Plus | |

Features:
**receptor tyrosine-protein kinase erbB-4 isoform JM-a/CVT-...receptor tyrosine-protein kinase erbB-4 isoform JM-a/CVT-...**

```
Query  277        AAGCTGGTGTTCTC  290
                  ||||||||||||||
Sbjct  212595034  AAGCTGGTGTTCTC  212595047
```

Range 35: 217915507 to 217915520

| Score | Expect | Identities | Gaps | Strand | Frame |
|---|---|---|---|---|---|
| 28.2 bits(14) | 256() | 14/14(100%) | 0/14(0%) | Plus/Plus | |

Features:
**184696 bp at 5' side: spermatid nuclear transition protein 1760061 bp at 3' side: tensin-1**

```
Query  286        TTCTCTCCTTTATT  299
                  ||||||||||||||
Sbjct  217915507  TTCTCTCCTTTATT  217915520
```

Range 36: 222268365 to 222268378

| Score | Expect | Identities | Gaps | Strand | Frame |
|---|---|---|---|---|---|
| 28.2 bits(14) | 256() | 14/14(100%) | 0/14(0%) | Plus/Plus | |

Features:
**1756095 bp at 5' side: anion exchange protein 3 isoform 127910 bp at 3' side: ephrin type-A receptor 4 isoform a precursor**

```
Query  287        TCTCTCCTTTATTG  300
                  ||||||||||||||
Sbjct  222268365  TCTCTCCTTTATTG  222268378
```

Range 37: 222536304 to 222536317

| Score | Expect | Identities | Gaps | Strand | Frame |
|---|---|---|---|---|---|
| 28.2 bits(14) | 256() | 14/14(100%) | 0/14(0%) | Plus/Plus | |

Features:
**93552 bp at 5' side: ephrin type-A receptor 4 isoform b536065 bp at 3' side: paired box protein Pax-3 isoform PAX3e**

```
Query  286        TTCTCTCCTTTATT  299
                  ||||||||||||||
Sbjct  222536304  TTCTCTCCTTTATT  222536317
```

Range 38: 225748178 to 225748191

| Score | Expect | Identities | Gaps | Strand | Frame |
|---|---|---|---|---|---|
| 28.2 bits(14) | 256() | 14/14(100%) | 0/14(0%) | Plus/Plus | |

Features:
**dedicator of cytokinesis protein 10 DOCK10.1dedicator of cytokinesis protein 10 DOCK10.2**

```
Query  286        TTCTCTCCTTTATT  299
                  ||||||||||||||
Sbjct  225748178  TTCTCTCCTTTATT  225748191
```

Range 39: 229832192 to 229832205

| Score | Expect | Identities | Gaps | Strand | Frame |
|---|---|---|---|---|---|
| 28.2 bits(14) | 256() | 14/14(100%) | 0/14(0%) | Plus/Plus | |

Features:
**779174 bp at 5' side: A-kinase anchor protein SPHKAP isoform 264303 bp at 3' side: PTB-containing, cubilin and LRP1-interacting protein isof...**

```
Query  286        TTCTCTCCTTTATT  299
                  ||||||||||||||
Sbjct  229832192  TTCTCTCCTTTATT  229832205
```

Range 40: 17616107 to 17616120

| Score | Expect | Identities | Gaps | Strand | Frame |
|---|---|---|---|---|---|
| 28.2 bits(14) | 256() | 14/14(100%) | 0/14(0%) | Plus/Minus | |

Features:
**917267 bp at 5' side: protein FAM49A5347 bp at 3' side: RAD51-associated protein 2**

```
Query  286       TTCTCTCCTTTATT  299
                 ||||||||||||||
Sbjct  17616120  TTCTCTCCTTTATT  17616107
```

Range 41: 22112452 to 22112465

| Score | Expect | Identities | Gaps | Strand | Frame |
|---|---|---|---|---|---|
| 28.2 bits(14) | 256() | 14/14(100%) | 0/14(0%) | Plus/Minus | |

Features:
**816956 bp at 5' side: tudor domain-containing protein 151601972 bp at 3' side: kelch-like protein 29**

```
Query  286       TTCTCTCCTTTATT  299
                 ||||||||||||||
Sbjct  22112465  TTCTCTCCTTTATT  22112452
```

Range 42: 35412203 to 35412216

| Score | Expect | Identities | Gaps | Strand | Frame |
|---|---|---|---|---|---|
| 28.2 bits(14) | 256() | 14/14(100%) | 0/14(0%) | Plus/Minus | |

Features:

**1670561 bp at 5' side: protein FAM98A isoform 21100579 bp at 3' side: cysteine-rich motor neuron 1 protein precursor**

```
Query  286       TTCTCTCCTTTATT  299
                 ||||||||||||||
Sbjct  35412216  TTCTCTCCTTTATT  35412203
```

Range 43: 58895339 to 58895352

| Score | Expect | Identities | Gaps | Strand | Frame |
|---|---|---|---|---|---|
| 28.2 bits(14) | 256() | 14/14(100%) | 0/14(0%) | Plus/Minus | |

Features:
**496662 bp at 5' side: E3 ubiquitin-protein ligase FANCL isoform 11714368 bp at 3' side: B-cell lymphoma/leukemia 11A isoform 3**

```
Query  283       GTGTTCTCTCCTTT  296
                 ||||||||||||||
Sbjct  58895352  GTGTTCTCTCCTTT  58895339
```

Range 44: 66401261 to 66401278

| Score | Expect | Identities | Gaps | Strand | Frame |
|---|---|---|---|---|---|
| 28.2 bits(14) | 256() | 17/18(94%) | 0/18(0%) | Plus/Minus | |

Features:
**876159 bp at 5' side: sprouty-related, EVH1 domain-containing protein 2 isoform b192901 bp at 3' side: homeobox protein Meis1**

```
Query  279       GCTGGTGTTCTCTCCTTT  296
                 ||||||||| ||||||||
Sbjct  66401278  GCTGGTGTTTTCTCCTTT  66401261
```

Range 45: 75125778 to 75125791

| Score | Expect | Identities | Gaps | Strand | Frame |
|---|---|---|---|---|---|
| 28.2 bits(14) | 256() | 14/14(100%) | 0/14(0%) | Plus/Minus | |

Features:
**DNA polymerase epsilon subunit 4**

```
Query  270       GTGTAACAAGCTGG  283
                 ||||||||||||||
Sbjct  75125791  GTGTAACAAGCTGG  75125778
```

Range 46: 80761805 to 80761818

| Score | Expect | Identities | Gaps | Strand | Frame |
|---|---|---|---|---|---|
| 28.2 bits(14) | 256() | 14/14(100%) | 0/14(0%) | Plus/Minus | |

Features:
**catenin alpha-2 isoform 4catenin alpha-2 isoform 3**

```
Query  286       TTCTCTCCTTTATT  299
                 ||||||||||||||
Sbjct  80761818  TTCTCTCCTTTATT  80761805
```

Range 47: 89152897 to 89152910

| Score | Expect | Identities | Gaps | Strand | Frame |
|---|---|---|---|---|---|
| 28.2 bits(14) | 256() | 14/14(100%) | 0/14(0%) | Plus/Minus | |

Features:
**8105 bp at 5' side: IGKV7-37855 bp at 3' side: IGKV2-4**

```
Query  286       TTCTCTCCTTTATT  299
                 ||||||||||||||
Sbjct  89152910  TTCTCTCCTTTATT  89152897
```

Range 48: 91934329 to 91934342

| Score | Expect | Identities | Gaps | Strand | Frame |
|---|---|---|---|---|---|
| 28.2 bits(14) | 256() | 14/14(100%) | 0/14(0%) | Plus/Minus | |

Features:
**44697 bp at 5' side: histone-lysine N-methyltransferase 2C-like3607291 bp at 3' side: tektin-4 isoform 1**

```
Query  286       TTCTCTCCTTTATT  299
                 ||||||||||||||
Sbjct  91934342  TTCTCTCCTTTATT  91934329
```

Range 49: 123536485 to 123536502

| Score | Expect | Identities | Gaps | Strand | Frame |
|---|---|---|---|---|---|
| 28.2 bits(14) | 256() | 17/18(94%) | 0/18(0%) | Plus/Minus | |

Features:
**1010002 bp at 5' side: translin isoform 11250328 bp at 3' side: contactin-associated protein-like 5 precursor**

```
Query  281        TGGTGTTCTCTCCTTTAT  298
                  ||||| ||||||||||||
Sbjct  123536502  TGGTGATCTCTCCTTTAT  123536485
```

Range 50: 127096356 to 127096369

| Score | Expect | Identities | Gaps | Strand | Frame |
|---|---|---|---|---|---|
| 28.2 bits(14) | 256() | 14/14(100%) | 0/14(0%) | Plus/Minus | |

Features:
**1420918 bp at 5' side: contactin-associated protein-like 5 precursor201623 bp at 3' side: uncharacterized protein LOC105373602 isoform X1**

```
Query  276        CAAGCTGGTGTTCT  289
                  ||||||||||||||
Sbjct  127096369  CAAGCTGGTGTTCT  127096356
```

Range 51: 131629969 to 131629982

| Score | Expect | Identities | Gaps | Strand | Frame |
|---|---|---|---|---|---|
| 28.2 bits(14) | 256() | 14/14(100%) | 0/14(0%) | Plus/Minus | |

Features:
**103732 bp at 5' side: APC membrane recruitment protein 362556 bp at 3' side: rho guanine nucleotide exchange factor 4 isoform a**

```
Query  283        GTGTTCTCTCCTTT  296
                  ||||||||||||||
Sbjct  131629982  GTGTTCTCTCCTTT  131629969
```

Range 52: 133647770 to 133647783

| Score | Expect | Identities | Gaps | Strand | Frame |
|---|---|---|---|---|---|
| 28.2 bits(14) | 256() | 14/14(100%) | 0/14(0%) | Plus/Minus | |

Features:
**nck-associated protein 5 isoform 1nck-associated protein 5 isoform 2**

```
Query  285        GTTCTCTCCTTTAT  298
                  ||||||||||||||
Sbjct  133647783  GTTCTCTCCTTTAT  133647770
```

Range 53: 135076472 to 135076485

| Score | Expect | Identities | Gaps | Strand | Frame |
|---|---|---|---|---|---|
| 28.2 bits(14) | 256() | 14/14(100%) | 0/14(0%) | Plus/Minus | |

Features:
**alpha-1,6-mannosylglycoprotein 6-beta-N-acetylglucosaminy...**

```
Query  286        TTCTCTCCTTTATT  299
```

```
                    ||||||||||||||
Sbjct  135076485  TTCTCTCCTTTATT  135076472
```

Range 54: 140207053 to 140207066

| Score | Expect | Identities | Gaps | Strand | Frame |
|---|---|---|---|---|---|
| 28.2 bits(14) | 256() | 14/14(100%) | 0/14(0%) | Plus/Minus | |

Features:
**664622 bp at 5' side: neurexophilin-2 precursor788998 bp at 3' side: low-density lipoprotein receptor-related protein 1B precu...**

```
Query  286        TTCTCTCCTTTATT  299
                  ||||||||||||||
Sbjct  140207066  TTCTCTCCTTTATT  140207053
```

Range 55: 143399181 to 143399194

| Score | Expect | Identities | Gaps | Strand | Frame |
|---|---|---|---|---|---|
| 28.2 bits(14) | 256() | 14/14(100%) | 0/14(0%) | Plus/Minus | |

Features:
**505745 bp at 5' side: low-density lipoprotein receptor-related protein 1B precu...248748 bp at 3' side: kynureninase isoform a**

```
Query  286        TTCTCTCCTTTATT  299
                  ||||||||||||||
Sbjct  143399194  TTCTCTCCTTTATT  143399181
```

Range 56: 146377051 to 146377064

| Score | Expect | Identities | Gaps | Strand | Frame |
|---|---|---|---|---|---|
| 28.2 bits(14) | 256() | 14/14(100%) | 0/14(0%) | Plus/Minus | |

Features:
**1097044 bp at 5' side: zinc finger E-box-binding homeobox 2 isoform 12231287 bp at 3' side: activin receptor type-2A isoform 1 precursor**

```
Query  283        GTGTTCTCTCCTTT  296
                  ||||||||||||||
Sbjct  146377064  GTGTTCTCTCCTTT  146377051
```

Range 57: 151265155 to 151265168

| Score | Expect | Identities | Gaps | Strand | Frame |
|---|---|---|---|---|---|
| 28.2 bits(14) | 256() | 14/14(100%) | 0/14(0%) | Plus/Minus | |

Features:
**815628 bp at 5' side: methylmalonic aciduria and homocystinuria type D protein,...67118 bp at 3' side: rho-related GTP-binding protein RhoE precursor**

```
Query  286        TTCTCTCCTTTATT  299
                  ||||||||||||||
Sbjct  151265168  TTCTCTCCTTTATT  151265155
```

Range 58: 151691972 to 151691985

| Score | Expect | Identities | Gaps | Strand | Frame |
|---|---|---|---|---|---|
| 28.2 bits(14) | 256() | 14/14(100%) | 0/14(0%) | Plus/Minus | |

Features:
**342256 bp at 5' side: rho-related GTP-binding protein RhoE precursor421275 bp at 3' side: RNA-binding protein 43**

```
Query  287        TCTCTCCTTTATTG  300
                  ||||||||||||||
Sbjct  151691985  TCTCTCCTTTATTG  151691972
```

Range 59: 152486363 to 152486376

| Score | Expect | Identities | Gaps | Strand | Frame |
|---|---|---|---|---|---|

| | | | | |
|---|---|---|---|---|
| 28.2 bits(14) | 256() | 14/14(100%) | 0/14(0%) | Plus/Minus |

Features:
**nebulin isoform 3nebulin isoform 2**

```
Query  283        GTGTTCTCTCCTTT  296
                  ||||||||||||||
Sbjct  152486376  GTGTTCTCTCCTTT  152486363
```

Range 60: 157051598 to 157051611

| Score | Expect | Identities | Gaps | Strand | Frame |
|---|---|---|---|---|---|
| 28.2 bits(14) | 256() | 14/14(100%) | 0/14(0%) | Plus/Minus | |

Features:
**1334097 bp at 5' side: G protein-activated inward rectifier potassium channel 1 ...136726 bp at 3' side: nuclear receptor subfamily 4 group A member 2**

```
Query  286        TTCTCTCCTTTATT  299
                  ||||||||||||||
Sbjct  157051611  TTCTCTCCTTTATT  157051598
```

Range 61: 157909032 to 157909045

| Score | Expect | Identities | Gaps | Strand | Frame |
|---|---|---|---|---|---|
| 28.2 bits(14) | 256() | 14/14(100%) | 0/14(0%) | Plus/Minus | |

Features:
**463573 bp at 5' side: glycerol-3-phosphate dehydrogenase, mitochondrial precursor211567 bp at 3' side: polypeptide N-acetylgalactosaminyltransferase 5**

```
Query  279        GCTGGTGTTCTCTC  292
                  ||||||||||||||
Sbjct  157909045  GCTGGTGTTCTCTC  157909032
```

Range 62: 175833123 to 175833140

| Score | Expect | Identities | Gaps | Strand | Frame |
|---|---|---|---|---|---|
| 28.2 bits(14) | 256() | 17/18(94%) | 0/18(0%) | Plus/Minus | |

Features:
**N-chimaerin isoform 2N-chimaerin isoform 1**

```
Query  274        AACAAGCTGGTGTTCTCT  291
                  ||||||||| ||||||||
Sbjct  175833140  AACAAGCTGCTGTTCTCT  175833123
```

Range 63: 178549583 to 178549596

| Score | Expect | Identities | Gaps | Strand | Frame |
|---|---|---|---|---|---|
| 28.2 bits(14) | 256() | 14/14(100%) | 0/14(0%) | Plus/Minus | |

Features:
**dual 3',5'-cyclic-AMP and -GMP phosphodiesterase 11A isof...dual 3',5'-cyclic-AMP and -GMP phosphodiesterase 11A isof...**

```
Query  286        TTCTCTCCTTTATT  299
                  ||||||||||||||
Sbjct  178549596  TTCTCTCCTTTATT  178549583
```

Range 64: 181773559 to 181773572

| Score | Expect | Identities | Gaps | Strand | Frame |
|---|---|---|---|---|---|
| 28.2 bits(14) | 256() | 14/14(100%) | 0/14(0%) | Plus/Minus | |

Features:
**909528 bp at 5' side: pre-mRNA-splicing factor CWC22 homolog79166 bp at 3' side: ubiquitin-conjugating enzyme E2 E3**

```
Query  286        TTCTCTCCTTTATT  299
                  ||||||||||||||
Sbjct  181773572  TTCTCTCCTTTATT  181773559
```

Range 65: 199481083 to 199481096

| Score | Expect | Identities | Gaps | Strand | Frame |
|---|---|---|---|---|---|
| 28.2 bits(14) | 256() | 14/14(100%) | 0/14(0%) | Plus/Minus | |

Features:
**463410 bp at 5' side: inactive phospholipase C-like protein 1662050 bp at 3' side: DNA-binding protein SATB2**

```
Query  286        TTCTCTCCTTTATT  299
                  ||||||||||||||
Sbjct  199481096  TTCTCTCCTTTATT  199481083
```

Range 66: 200184717 to 200184730

| Score | Expect | Identities | Gaps | Strand | Frame |
|---|---|---|---|---|---|
| 28.2 bits(14) | 256() | 14/14(100%) | 0/14(0%) | Plus/Minus | |

Features:
**DNA-binding protein SATB2DNA-binding protein SATB2**

```
Query  285        GTTCTCTCCTTTAT  298
                  ||||||||||||||
Sbjct  200184730  GTTCTCTCCTTTAT  200184717
```

Range 67: 212256501 to 212256514

| Score | Expect | Identities | Gaps | Strand | Frame |
|---|---|---|---|---|---|
| 28.2 bits(14) | 256() | 14/14(100%) | 0/14(0%) | Plus/Minus | |

Features:
**receptor tyrosine-protein kinase erbB-4 isoform JM-a/CVT-...receptor tyrosine-protein kinase erbB-4 isoform JM-a/CVT-...**

```
Query  284        TGTTCTCTCCTTTA  297
                  ||||||||||||||
Sbjct  212256514  TGTTCTCTCCTTTA  212256501
```

Range 68: 214833102 to 214833115

| Score | Expect | Identities | Gaps | Strand | Frame |
|---|---|---|---|---|---|
| 28.2 bits(14) | 256() | 14/14(100%) | 0/14(0%) | Plus/Minus | |

Features:
**sperm-associated antigen 16 protein isoform 1**

```
Query  286        TTCTCTCCTTTATT  299
                  ||||||||||||||
Sbjct  214833115  TTCTCTCCTTTATT  214833102
```

Range 69: 215362928 to 215362941

| Score | Expect | Identities | Gaps | Strand | Frame |
|---|---|---|---|---|---|
| 28.2 bits(14) | 256() | 14/14(100%) | 0/14(0%) | Plus/Minus | |

Features:
**von Willebrand factor C domain-containing protein 2-like ...**

```
Query  283        GTGTTCTCTCCTTT  296
                  ||||||||||||||
Sbjct  215362941  GTGTTCTCTCCTTT  215362928
```

Range 70: 225770959 to 225770972

| Score | Expect | Identities | Gaps | Strand | Frame |
|---|---|---|---|---|---|
| 28.2 bits(14) | 256() | 14/14(100%) | 0/14(0%) | Plus/Minus | |

Features:
**dedicator of cytokinesis protein 10 DOCK10.1dedicator of cytokinesis protein 10 DOCK10.2**

```
Query  284        TGTTCTCTCCTTTA  297
                  ||||||||||||||
```

```
Sbjct  225770972  TGTTCTCTCCTTTA  225770959
```

Range 71: 229419410 to 229419423

| Score | Expect | Identities | Gaps | Strand | Frame |
|---|---|---|---|---|---|
| 28.2 bits(14) | 256() | 14/14(100%) | 0/14(0%) | Plus/Minus | |

Features:
**366392 bp at 5' side: A-kinase anchor protein SPHKAP isoform 2477085 bp at 3' side: PTB-containing, cubilin and LRP1-interacting protein isof...**

```
Query  283        GTGTTCTCTCCTTT  296
                  ||||||||||||||
Sbjct  229419423  GTGTTCTCTCCTTT  229419410
```

Range 72: 231160529 to 231160542

| Score | Expect | Identities | Gaps | Strand | Frame |
|---|---|---|---|---|---|
| 28.2 bits(14) | 256() | 14/14(100%) | 0/14(0%) | Plus/Minus | |

Features:
**nuclear body protein SP140 isoform 1nuclear body protein SP140 isoform 3**

```
Query  284        TGTTCTCTCCTTTA  297
                  ||||||||||||||
Sbjct  231160542  TGTTCTCTCCTTTA  231160529
```

Range 73: 232340517 to 232340530

| Score | Expect | Identities | Gaps | Strand | Frame |
|---|---|---|---|---|---|
| 28.2 bits(14) | 256() | 14/14(100%) | 0/14(0%) | Plus/Minus | |

Features:
**5637 bp at 5' side: nucleolin55038 bp at 3' side: neuromedin-U receptor 1**

```
Query  287        TCTCTCCTTTATTG  300
                  ||||||||||||||
Sbjct  232340530  TCTCTCCTTTATTG  232340517
```

Homo sapiens chromosome 4, alternate assembly CHM1_1.1
Sequence ID: **ref|NC_018915.2|** Length: 191040880 Number of Matches: 62
Range 1: 40782029 to 40782049

| Score | Expect | Identities | Gaps | Strand | Frame |
|---|---|---|---|---|---|
| 34.2 bits(17) | 4.1() | 20/21(95%) | 0/21(0%) | Plus/Minus | |

Features:
**putative methyltransferase NSUN7**

```
Query  280       CTGGTGTTCTCTCCTTTATTG  300
                 ||||| |||||||||||||||
Sbjct  40782049  CTGGTCTTCTCTCCTTTATTG  40782029
```

Range 2: 92618982 to 92618998

| Score | Expect | Identities | Gaps | Strand | Frame |
|---|---|---|---|---|---|
| 34.2 bits(17) | 4.1() | 17/17(100%) | 0/17(0%) | Plus/Minus | |

Features:
**122260 bp at 5' side: serine-rich coiled-coil domain-containing protein 1 isofo...583282 bp at 3' side: glutamate receptor ionotropic, delta-2 isoform 1 precursor**

```
Query  283       GTGTTCTCTCCTTTATT  299
                 |||||||||||||||||
Sbjct  92618998  GTGTTCTCTCCTTTATT  92618982
```

Range 3: 92427430 to 92427445

| Score | Expect | Identities | Gaps | Strand | Frame |
|---|---|---|---|---|---|
| 32.2 bits(16) | 16() | 16/16(100%) | 0/16(0%) | Plus/Plus | |

Features:
**serine-rich coiled-coil domain-containing protein 1 isofo...**

```
Query  284       TGTTCTCTCCTTTATT  299
                 ||||||||||||||||
Sbjct  92427430  TGTTCTCTCCTTTATT  92427445
```

Range 4: 181747246 to 181747261

| Score | Expect | Identities | Gaps | Strand | Frame |
|---|---|---|---|---|---|
| 32.2 bits(16) | 16() | 16/16(100%) | 0/16(0%) | Plus/Plus | |

Features:
**3174085 bp at 5' side: uncharacterized protein LOC285500 isoform X21474448 bp at 3' side: teneurin-3**

```
Query  279        GCTGGTGTTCTCTCCT  294
                  ||||||||||||||||
Sbjct  181747246  GCTGGTGTTCTCTCCT  181747261
```

Range 5: 54587425 to 54587439

| Score | Expect | Identities | Gaps | Strand | Frame |
|---|---|---|---|---|---|
| 30.2 bits(15) | 65() | 15/15(100%) | 0/15(0%) | Plus/Plus | |

Features:
**109141 bp at 5' side: COMM domain-containing protein 5-like324522 bp at 3' side: cysteine-rich hydrophobic domain-containing protein 2**

```
Query  271       TGTAACAAGCTGGTG  285
                 |||||||||||||||
Sbjct  54587425  TGTAACAAGCTGGTG  54587439
```

Range 6: 136512706 to 136512720

| Score | Expect | Identities | Gaps | Strand | Frame |
|---|---|---|---|---|---|
| 30.2 bits(15) | 65() | 15/15(100%) | 0/15(0%) | Plus/Plus | |

Features:
**1413905 bp at 5' side: polyadenylate-binding protein 4-like1905967 bp at 3' side: protocadherin-18 isoform 2 precursor**

```
Query  286        TTCTCTCCTTTATTG  300
                  |||||||||||||||
Sbjct  136512706  TTCTCTCCTTTATTG  136512720
```

Range 7: 151648426 to 151648440

| Score | Expect | Identities | Gaps | Strand | Frame |
|---|---|---|---|---|---|
| 30.2 bits(15) | 65() | 15/15(100%) | 0/15(0%) | Plus/Plus | |

Features:
**lipopolysaccharide-responsive and beige-like anchor prote...lipopolysaccharide-responsive and beige-like anchor prote...**

```
Query  284        TGTTCTCTCCTTTAT  298
                  |||||||||||||||
Sbjct  151648426  TGTTCTCTCCTTTAT  151648440
```

Range 8: 160060274 to 160060292

| Score | Expect | Identities | Gaps | Strand | Frame |
|---|---|---|---|---|---|
| 30.2 bits(15) | 65() | 18/19(95%) | 0/19(0%) | Plus/Plus | |

Features:
**127543 bp at 5' side: uncharacterized protein C4orf45105519 bp at 3' side: rap guanine nucleotide exchange factor 2**

```
Query  279        GCTGGTGTTCTCTCCTTTA  297
                  |||||||||||||| ||||
```

```
Sbjct  160060274  GCTGGTGTTCTCTCATTTA  160060292
```

Range 9: 160835956 to 160835970

| Score | Expect | Identities | Gaps | Strand | Frame |
|---|---|---|---|---|---|
| 30.2 bits(15) | 65() | 15/15(100%) | 0/15(0%) | Plus/Plus | |

Features:
**580172 bp at 5' side: rap guanine nucleotide exchange factor 21447391 bp at 3' side: follistatin-related protein 5 isoform c precursor**

```
Query  283        GTGTTCTCTCCTTTA  297
                  |||||||||||||||
Sbjct  160835956  GTGTTCTCTCCTTTA  160835970
```

Range 10: 171115512 to 171115526

| Score | Expect | Identities | Gaps | Strand | Frame |
|---|---|---|---|---|---|
| 30.2 bits(15) | 65() | 15/15(100%) | 0/15(0%) | Plus/Plus | |

Features:
**111069 bp at 5' side: DNA dC->dU-editing enzyme APOBEC-3G-like1596630 bp at 3' side: polypeptide N-acetylgalactosaminyltransferase-like 6**

```
Query  273        TAACAAGCTGGTGTT  287
                  |||||||||||||||
Sbjct  171115512  TAACAAGCTGGTGTT  171115526
```

Range 11: 60160772 to 60160786

| Score | Expect | Identities | Gaps | Strand | Frame |
|---|---|---|---|---|---|
| 30.2 bits(15) | 65() | 15/15(100%) | 0/15(0%) | Plus/Minus | |

Features:
**2148439 bp at 5' side: insulin-like growth factor-binding protein 7 isoform 2 pr...2238095 bp at 3' side: latrophilin-3 precursor**

```
Query  285       GTTCTCTCCTTTATT  299
                 |||||||||||||||
Sbjct  60160786  GTTCTCTCCTTTATT  60160772
```

Range 12: 77447831 to 77447845

| Score | Expect | Identities | Gaps | Strand | Frame |
|---|---|---|---|---|---|
| 30.2 bits(15) | 65() | 15/15(100%) | 0/15(0%) | Plus/Minus | |

Features:
**protein Shroom3**

```
Query  282       GGTGTTCTCTCCTTT  296
                 |||||||||||||||
Sbjct  77447845  GGTGTTCTCTCCTTT  77447831
```

Range 13: 131447391 to 131447405

| Score | Expect | Identities | Gaps | Strand | Frame |
|---|---|---|---|---|---|
| 30.2 bits(15) | 65() | 15/15(100%) | 0/15(0%) | Plus/Minus | |

Features:
**1437586 bp at 5' side: UPF0462 protein C4orf331179574 bp at 3' side: uncharacterized protein LOC105379408**

```
Query  284        TGTTCTCTCCTTTAT  298
                  |||||||||||||||
Sbjct  131447405  TGTTCTCTCCTTTAT  131447391
```

Range 14: 174736877 to 174736891

| Score | Expect | Identities | Gaps | Strand | Frame |
|---|---|---|---|---|---|
| 30.2 bits(15) | 65() | 15/15(100%) | 0/15(0%) | Plus/Minus | |

Features:
**309623 bp at 5' side: heart- and neural crest derivatives-expressed protein 2398125 bp at 3' side: F-box only protein 8**

```
Query  273        TAACAAGCTGGTGTT  287
                  |||||||||||||||
Sbjct  174736891  TAACAAGCTGGTGTT  174736877
```

Range 15: 190089488 to 190089502

| Score | Expect | Identities | Gaps | Strand | Frame |
|---|---|---|---|---|---|
| 30.2 bits(15) | 65() | 15/15(100%) | 0/15(0%) | Plus/Minus | |

Features:
**1044507 bp at 5' side: probable E3 ubiquitin-protein ligase TRIML1756994 bp at 3' side: protein FRG1**

```
Query  280        CTGGTGTTCTCTCCT  294
                  |||||||||||||||
Sbjct  190089502  CTGGTGTTCTCTCCT  190089488
```

Range 16: 3195713 to 3195726

| Score | Expect | Identities | Gaps | Strand | Frame |
|---|---|---|---|---|---|
| 28.2 bits(14) | 256() | 14/14(100%) | 0/14(0%) | Plus/Plus | |

Features:
**huntingtin**

```
Query  279      GCTGGTGTTCTCTC  292
                ||||||||||||||
Sbjct  3195713  GCTGGTGTTCTCTC  3195726
```

Range 17: 3222378 to 3222391

| Score | Expect | Identities | Gaps | Strand | Frame |
|---|---|---|---|---|---|
| 28.2 bits(14) | 256() | 14/14(100%) | 0/14(0%) | Plus/Plus | |

Features:
**huntingtin**

```
Query  276      CAAGCTGGTGTTCT  289
                ||||||||||||||
Sbjct  3222378  CAAGCTGGTGTTCT  3222391
```

Range 18: 8398538 to 8398551

| Score | Expect | Identities | Gaps | Strand | Frame |
|---|---|---|---|---|---|
| 28.2 bits(14) | 256() | 14/14(100%) | 0/14(0%) | Plus/Plus | |

Features:
**peroxisomal acyl-coenzyme A oxidase 3 isoform aperoxisomal acyl-coenzyme A oxidase 3 isoform b**

```
Query  286      TTCTCTCCTTTATT  299
                ||||||||||||||
Sbjct  8398538  TTCTCTCCTTTATT  8398551
```

Range 19: 10767689 to 10767702

| Score | Expect | Identities | Gaps | Strand | Frame |
|---|---|---|---|---|---|
| 28.2 bits(14) | 256() | 14/14(100%) | 0/14(0%) | Plus/Plus | |

Features:
**100003 bp at 5' side: cytokine-dependent hematopoietic cell linker631071 bp at 3' side: heparan sulfate glucosamine 3-O-sulfotransferase 1 precursor**

```
Query  285       GTTCTCTCCTTTAT  298
                 ||||||||||||||
Sbjct  10767689  GTTCTCTCCTTTAT  10767702
```

Range 20: 23298385 to 23298398

| Score | Expect | Identities | Gaps | Strand | Frame |
|---|---|---|---|---|---|
| 28.2 bits(14) | 256() | 14/14(100%) | 0/14(0%) | Plus/Plus | |

Features:
**479903 bp at 5' side: cytosolic beta-glucosidase isoform c496975 bp at 3' side: peroxisome proliferator-activated receptor gamma coactiva...**

```
Query  281       TGGTGTTCTCTCCT  294
                 ||||||||||||||
Sbjct  23298385  TGGTGTTCTCTCCT  23298398
```

Range 21: 28462457 to 28462470

| Score | Expect | Identities | Gaps | Strand | Frame |
|---|---|---|---|---|---|
| 28.2 bits(14) | 256() | 14/14(100%) | 0/14(0%) | Plus/Plus | |

Features:
**1438281 bp at 5' side: stromal interaction molecule 2 isoform 3 precursor2261987 bp at 3' side: protocadherin-7 isoform d precursor**

```
Query  276       CAAGCTGGTGTTCT  289
                 ||||||||||||||
Sbjct  28462457  CAAGCTGGTGTTCT  28462470
```

Range 22: 56181418 to 56181431

| Score | Expect | Identities | Gaps | Strand | Frame |
|---|---|---|---|---|---|
| 28.2 bits(14) | 256() | 14/14(100%) | 0/14(0%) | Plus/Plus | |

Features:
**154705 bp at 5' side: vascular endothelial growth factor receptor 2 precursor66227 bp at 3' side: polyprenol reductase**

```
Query  283       GTGTTCTCTCCTTT  296
                 ||||||||||||||
Sbjct  56181418  GTGTTCTCTCCTTT  56181431
```

Range 23: 61004533 to 61004550

| Score | Expect | Identities | Gaps | Strand | Frame |
|---|---|---|---|---|---|
| 28.2 bits(14) | 256() | 17/18(94%) | 0/18(0%) | Plus/Plus | |

Features:
**2992200 bp at 5' side: insulin-like growth factor-binding protein 7 isoform 2 pr...1394331 bp at 3' side: latrophilin-3 precursor**

```
Query  272       GTAACAAGCTGGTGTTCT  289
                 ||||||| ||||||||||
Sbjct  61004533  GTAACAATCTGGTGTTCT  61004550
```

Range 24: 67227764 to 67227777

| Score | Expect | Identities | Gaps | Strand | Frame |
|---|---|---|---|---|---|
| 28.2 bits(14) | 256() | 14/14(100%) | 0/14(0%) | Plus/Plus | |

Features:
**654795 bp at 5' side: ephrin type-A receptor 5 isoform e1147734 bp at 3' side: centromere protein C**

```
Query  271       TGTAACAAGCTGGT  284
                 ||||||||||||||
Sbjct  67227764  TGTAACAAGCTGGT  67227777
```

Range 25: 69877246 to 69877259

| Score | Expect | Identities | Gaps | Strand | Frame |
|---|---|---|---|---|---|
| 28.2 bits(14) | 256() | 14/14(100%) | 0/14(0%) | Plus/Plus | |

Features:
**23177 bp at 5' side: UDP-glucuronosyltransferase 2A3 precursor121303 bp at 3' side: UDP-glucuronosyltransferase 2B7 precursor**

```
Query  286       TTCTCTCCTTTATT  299
                 ||||||||||||||
Sbjct  69877246  TTCTCTCCTTTATT  69877259
```

Range 26: 79076135 to 79076148

| Score | Expect | Identities | Gaps | Strand | Frame |
|---|---|---|---|---|---|
| 28.2 bits(14) | 256() | 14/14(100%) | 0/14(0%) | Plus/Plus | |

Features:
**extracellular matrix protein FRAS1 isoform 1 precursorextracellular matrix protein FRAS1 isoform 2 precursor**

```
Query  283       GTGTTCTCTCCTTT  296
                 ||||||||||||||
Sbjct  79076135  GTGTTCTCTCCTTT  79076148
```

Range 27: 86370492 to 86370505

| Score | Expect | Identities | Gaps | Strand | Frame |
|---|---|---|---|---|---|
| 28.2 bits(14) | 256() | 14/14(100%) | 0/14(0%) | Plus/Plus | |

Features:
**611531 bp at 5' side: WD repeat and FYVE domain-containing protein 398642 bp at 3' side: rho GTPase-activating protein 24 isoform 1**

```
Query  286       TTCTCTCCTTTATT  299
                 ||||||||||||||
Sbjct  86370492  TTCTCTCCTTTATT  86370505
```

Range 28: 94135991 to 94136004

| Score | Expect | Identities | Gaps | Strand | Frame |
|---|---|---|---|---|---|
| 28.2 bits(14) | 256() | 14/14(100%) | 0/14(0%) | Plus/Plus | |

Features:
**glutamate receptor ionotropic, delta-2 isoform 1 precursorglutamate receptor ionotropic, delta-2 isoform 2 precursor**

```
Query  284       TGTTCTCTCCTTTA  297
                 ||||||||||||||
Sbjct  94135991  TGTTCTCTCCTTTA  94136004
```

Range 29: 97381547 to 97381560

| Score | Expect | Identities | Gaps | Strand | Frame |
|---|---|---|---|---|---|
| 28.2 bits(14) | 256() | 14/14(100%) | 0/14(0%) | Plus/Plus | |

Features:
**642408 bp at 5' side: pyruvate dehydrogenase E1 component subunit alpha, testis...1075425 bp at 3' side: sperm-tail PG-rich repeat-containing protein 2**

```
Query  278       AGCTGGTGTTCTCT  291
                 ||||||||||||||
Sbjct  97381547  AGCTGGTGTTCTCT  97381560
```

Range 30: 108776226 to 108776239

| Score | Expect | Identities | Gaps | Strand | Frame |
|---|---|---|---|---|---|
| 28.2 bits(14) | 256() | 14/14(100%) | 0/14(0%) | Plus/Plus | |

Features:
**158393 bp at 5' side: bifunctional 3'-phosphoadenosine 5'-phosphosulfate syntha...16934 bp at 3' side: phosphatidylcholine:ceramide cholinephosphotransferase 2**

```
Query  286        TTCTCTCCTTTATT  299
                  ||||||||||||||
Sbjct  108776226  TTCTCTCCTTTATT  108776239
```

Range 31: 110542903 to 110542916

| Score | Expect | Identities | Gaps | Strand | Frame |
|---|---|---|---|---|---|
| 28.2 bits(14) | 256() | 14/14(100%) | 0/14(0%) | Plus/Plus | |

Features:
**calcium uniporter regulatory subunit MCUb, mitochondrial**

```
Query  286        TTCTCTCCTTTATT  299
                  ||||||||||||||
Sbjct  110542903  TTCTCTCCTTTATT  110542916
```

Range 32: 113435455 to 113435468

| Score | Expect | Identities | Gaps | Strand | Frame |
|---|---|---|---|---|---|
| 28.2 bits(14) | 256() | 14/14(100%) | 0/14(0%) | Plus/Plus | |

Features:
**22002 bp at 5' side: neurogenin-22074 bp at 3' side: protein ZGRF1**

```
Query  286        TTCTCTCCTTTATT  299
                  ||||||||||||||
Sbjct  113435455  TTCTCTCCTTTATT  113435468
```

Range 33: 113630256 to 113630269

| Score | Expect | Identities | Gaps | Strand | Frame |
|---|---|---|---|---|---|
| 28.2 bits(14) | 256() | 14/14(100%) | 0/14(0%) | Plus/Plus | |

Features:
**75280 bp at 5' side: la-related protein 7 isoform 1171867 bp at 3' side: ankyrin-2 isoform 3**

```
Query  282        GGTGTTCTCTCCTT  295
                  ||||||||||||||
Sbjct  113630256  GGTGTTCTCTCCTT  113630269
```

Range 34: 116474577 to 116474590

| Score | Expect | Identities | Gaps | Strand | Frame |
|---|---|---|---|---|---|
| 28.2 bits(14) | 256() | 14/14(100%) | 0/14(0%) | Plus/Plus | |

Features:
**500010 bp at 5' side: bifunctional heparan sulfate N-deacetylase/N-sulfotransfe...1507305 bp at 3' side: translocating chain-associated membrane protein 1-like 1**

```
Query  286        TTCTCTCCTTTATT  299
                  ||||||||||||||
Sbjct  116474577  TTCTCTCCTTTATT  116474590
```

Range 35: 143989895 to 143989908

| Score | Expect | Identities | Gaps | Strand | Frame |
|---|---|---|---|---|---|
| 28.2 bits(14) | 256() | 14/14(100%) | 0/14(0%) | Plus/Plus | |

Features:
**660216 bp at 5' side: type II inositol 3,4-bisphosphate 4-phosphatase93910 bp at 3' side: ubiquitin carboxyl-terminal hydrolase 38 isoform 2**

```
Query  274        AACAAGCTGGTGTT  287
                  ||||||||||||||
Sbjct  143989895  AACAAGCTGGTGTT  143989908
```

Range 36: 149481706 to 149481723

| Score | Expect | Identities | Gaps | Strand | Frame |
|---|---|---|---|---|---|
| 28.2 bits(14) | 256() | 17/18(94%) | 0/18(0%) | Plus/Plus | |

Features:
**145023 bp at 5' side: mineralocorticoid receptor isoform 1717321 bp at 3' side: uncharacterized protein LOC285423 isoform X3**

```
Query  281        TGGTGTTCTCTCCTTTAT  298
```

```
                      ||||| ||||||||||||
Sbjct  149481706  TGGTGATCTCTCCTTTAT  149481723
```

Range 37: 158312741 to 158312754

| Score | Expect | Identities | Gaps | Strand | Frame |
|---|---|---|---|---|---|
| 28.2 bits(14) | 256() | 14/14(100%) | 0/14(0%) | Plus/Plus | |

Features:
**51423 bp at 5' side: glutamate receptor 2 isoform 3712992 bp at 3' side: protein FAM198B isoform 2**

```
Query  281        TGGTGTTCTCTCCT  294
                  ||||||||||||||
Sbjct  158312741  TGGTGTTCTCTCCT  158312754
```

Range 38: 165186709 to 165186722

| Score | Expect | Identities | Gaps | Strand | Frame |
|---|---|---|---|---|---|
| 28.2 bits(14) | 256() | 14/14(100%) | 0/14(0%) | Plus/Plus | |

Features:
**91397 bp at 5' side: acidic leucine-rich nuclear phosphoprotein 32 family memb...588330 bp at 3' side: apelin receptor early endogenous ligand precursor**

```
Query  277        AAGCTGGTGTTCTC  290
                  ||||||||||||||
Sbjct  165186709  AAGCTGGTGTTCTC  165186722
```

Range 39: 177943846 to 177943859

| Score | Expect | Identities | Gaps | Strand | Frame |
|---|---|---|---|---|---|
| 28.2 bits(14) | 256() | 14/14(100%) | 0/14(0%) | Plus/Plus | |

Features:
**253863 bp at 5' side: vascular endothelial growth factor C preproprotein263403 bp at 3' side: endonuclease 8-like 3**

```
Query  285        GTTCTCTCCTTTAT  298
                  ||||||||||||||
Sbjct  177943846  GTTCTCTCCTTTAT  177943859
```

Range 40: 182655194 to 182655207

| Score | Expect | Identities | Gaps | Strand | Frame |
|---|---|---|---|---|---|
| 28.2 bits(14) | 256() | 14/14(100%) | 0/14(0%) | Plus/Plus | |

Features:
**4082033 bp at 5' side: uncharacterized protein LOC285500 isoform X2566502 bp at 3' side: teneurin-3**

```
Query  273        TAACAAGCTGGTGT  286
                  ||||||||||||||
Sbjct  182655194  TAACAAGCTGGTGT  182655207
```

Range 41: 6077291 to 6077308

| Score | Expect | Identities | Gaps | Strand | Frame |
|---|---|---|---|---|---|
| 28.2 bits(14) | 256() | 17/18(94%) | 0/18(0%) | Plus/Minus | |

Features:
**janus kinase and microtubule-interacting protein 1 isoform 1janus kinase and microtubule-interacting protein 1 isoform 2**

```
Query  279      GCTGGTGTTCTCTCCTTT  296
                |||||| |||||||||||
Sbjct  6077308  GCTGGTCTTCTCTCCTTT  6077291
```

Range 42: 8749499 to 8749512

| Score | Expect | Identities | Gaps | Strand | Frame |
|---|---|---|---|---|---|
| 28.2 bits(14) | 256() | 14/14(100%) | 0/14(0%) | Plus/Minus | |

Features:
**129830 bp at 5' side: carboxypeptidase Z isoform 3117853 bp at 3' side: homeobox protein HMX1**

```
Query  278      AGCTGGTGTTCTCT  291
                ||||||||||||||
Sbjct  8749512  AGCTGGTGTTCTCT  8749499
```

Range 43: 21597687 to 21597700

| Score | Expect | Identities | Gaps | Strand | Frame |
|---|---|---|---|---|---|
| 28.2 bits(14) | 256() | 14/14(100%) | 0/14(0%) | Plus/Minus | |

Features:
**Kv channel-interacting protein 4 isoform 2Kv channel-interacting protein 4 isoform 1**

```
Query  283       GTGTTCTCTCCTTT  296
                 ||||||||||||||
Sbjct  21597700  GTGTTCTCTCCTTT  21597687
```

Range 44: 22638495 to 22638508

| Score | Expect | Identities | Gaps | Strand | Frame |
|---|---|---|---|---|---|
| 28.2 bits(14) | 256() | 14/14(100%) | 0/14(0%) | Plus/Minus | |

Features:
**123070 bp at 5' side: probable G-protein coupled receptor 125 precursor54092 bp at 3' side: cytosolic beta-glucosidase isoform a**

```
Query  280       CTGGTGTTCTCTCC  293
                 ||||||||||||||
Sbjct  22638508  CTGGTGTTCTCTCC  22638495
```

Range 45: 34324050 to 34324063

| Score | Expect | Identities | Gaps | Strand | Frame |
|---|---|---|---|---|---|
| 28.2 bits(14) | 256() | 14/14(100%) | 0/14(0%) | Plus/Minus | |

Features:
**3596375 bp at 5' side: protocadherin-7 isoform b precursor1744580 bp at 3' side: arf-GAP with Rho-GAP domain, ANK repeat and PH domain-con...**

```
Query  273       TAACAAGCTGGTGT  286
                 ||||||||||||||
Sbjct  34324063  TAACAAGCTGGTGT  34324050
```

Range 46: 39168414 to 39168427

| Score | Expect | Identities | Gaps | Strand | Frame |
|---|---|---|---|---|---|
| 28.2 bits(14) | 256() | 14/14(100%) | 0/14(0%) | Plus/Minus | |

Features:
**46199 bp at 5' side: kelch-like protein 5 isoform 415287 bp at 3' side: WD repeat-containing protein 19**

```
Query  271       TGTAACAAGCTGGT  284
                 ||||||||||||||
Sbjct  39168427  TGTAACAAGCTGGT  39168414
```

Range 47: 44883678 to 44883691

| Score | Expect | Identities | Gaps | Strand | Frame |
|---|---|---|---|---|---|
| 28.2 bits(14) | 256() | 14/14(100%) | 0/14(0%) | Plus/Minus | |

Features:
**164250 bp at 5' side: glucosamine-6-phosphate isomerase 2 isoform 31158444 bp at 3' side: gamma-aminobutyric acid receptor subunit gamma-1 precursor**

```
Query  286       TTCTCTCCTTTATT  299
                 ||||||||||||||
Sbjct  44883691  TTCTCTCCTTTATT  44883678
```

Range 48: 56915305 to 56915318

| Score | Expect | Identities | Gaps | Strand | Frame |
|---|---|---|---|---|---|
| 28.2 bits(14) | 256() | 14/14(100%) | 0/14(0%) | Plus/Minus | |

Features:
**centrosomal protein of 135 kDa**

```
Query  281       TGGTGTTCTCTCCT  294
                 ||||||||||||||
Sbjct  56915318  TGGTGTTCTCTCCT  56915305
```

Range 49: 63443989 to 63444002

| Score | Expect | Identities | Gaps | Strand | Frame |
|---|---|---|---|---|---|
| 28.2 bits(14) | 256() | 14/14(100%) | 0/14(0%) | Plus/Minus | |

Features:
**471414 bp at 5' side: latrophilin-3 precursor1738348 bp at 3' side: trans-2,3-enoyl-CoA reductase-like**

```
Query  286       TTCTCTCCTTTATT  299
                 ||||||||||||||
Sbjct  63444002  TTCTCTCCTTTATT  63443989
```

Range 50: 69090162 to 69090175

| Score | Expect | Identities | Gaps | Strand | Frame |
|---|---|---|---|---|---|
| 28.2 bits(14) | 256() | 14/14(100%) | 0/14(0%) | Plus/Minus | |

Features:
**57696 bp at 5' side: transmembrane protease serine 11F40098 bp at 3' side: transmembrane protease serine 11B**

```
Query  283       GTGTTCTCTCCTTT  296
                 ||||||||||||||
Sbjct  69090175  GTGTTCTCTCCTTT  69090162
```

Range 51: 70352629 to 70352642

| Score | Expect | Identities | Gaps | Strand | Frame |
|---|---|---|---|---|---|
| 28.2 bits(14) | 256() | 14/14(100%) | 0/14(0%) | Plus/Minus | |

Features:
**235944 bp at 5' side: UDP-glucuronosyltransferase 2B11 precursor29557 bp at 3' side: UDP-glucuronosyltransferase 2B4 isoform 3**

```
Query  286       TTCTCTCCTTTATT  299
                 ||||||||||||||
Sbjct  70352642  TTCTCTCCTTTATT  70352629
```

Range 52: 100448163 to 100448176

| Score | Expect | Identities | Gaps | Strand | Frame |
|---|---|---|---|---|---|
| 28.2 bits(14) | 256() | 14/14(100%) | 0/14(0%) | Plus/Minus | |

Features:
**tRNA methyltransferase 10 homolog AtRNA methyltransferase 10 homolog A**

```
Query  286        TTCTCTCCTTTATT  299
                  ||||||||||||||
Sbjct  100448176  TTCTCTCCTTTATT  100448163
```

Range 53: 101483531 to 101483544

| Score | Expect | Identities | Gaps | Strand | Frame |
|---|---|---|---|---|---|
| 28.2 bits(14) | 256() | 14/14(100%) | 0/14(0%) | Plus/Minus | |

Features:
**67910 bp at 5' side: endomucin isoform 1 precursor439960 bp at 3' side: serine/threonine-protein phosphatase 2B catalytic subunit...**

```
Query  283        GTGTTCTCTCCTTT  296
                  ||||||||||||||
Sbjct  101483544  GTGTTCTCTCCTTT  101483531
```

Range 54: 111541240 to 111541253

| Score | Expect | Identities | Gaps | Strand | Frame |
|---|---|---|---|---|---|
| 28.2 bits(14) | 256() | 14/14(100%) | 0/14(0%) | Plus/Minus | |

Features:
**10183 bp at 5' side: pituitary homeobox 2 isoform a1183770 bp at 3' side: histone-lysine N-methyltransferase SETMAR-like**

```
Query  279        GCTGGTGTTCTCTC  292
                  ||||||||||||||
Sbjct  111541253  GCTGGTGTTCTCTC  111541240
```

Range 55: 112491607 to 112491620

| Score | Expect | Identities | Gaps | Strand | Frame |
|---|---|---|---|---|---|
| 28.2 bits(14) | 256() | 14/14(100%) | 0/14(0%) | Plus/Minus | |

Features:
**960550 bp at 5' side: pituitary homeobox 2 isoform a233403 bp at 3' side: histone-lysine N-methyltransferase SETMAR-like**

```
Query  286        TTCTCTCCTTTATT  299
                  ||||||||||||||
Sbjct  112491620  TTCTCTCCTTTATT  112491607
```

Range 56: 116354996 to 116355009

| Score | Expect | Identities | Gaps | Strand | Frame |
|---|---|---|---|---|---|
| 28.2 bits(14) | 256() | 14/14(100%) | 0/14(0%) | Plus/Minus | |

Features:
**380429 bp at 5' side: bifunctional heparan sulfate N-deacetylase/N-sulfotransfe...1626886 bp at 3' side: translocating chain-associated membrane protein 1-like 1**

```
Query  283        GTGTTCTCTCCTTT  296
                  ||||||||||||||
Sbjct  116355009  GTGTTCTCTCCTTT  116354996
```

Range 57: 135456831 to 135456844

| Score | Expect | Identities | Gaps | Strand | Frame |
|---|---|---|---|---|---|
| 28.2 bits(14) | 256() | 14/14(100%) | 0/14(0%) | Plus/Minus | |

Features:
**358030 bp at 5' side: polyadenylate-binding protein 4-like2961843 bp at 3' side: protocadherin-18 isoform 2 precursor**

```
Query  286        TTCTCTCCTTTATT  299
                  ||||||||||||||
Sbjct  135456844  TTCTCTCCTTTATT  135456831
```

Range 58: 135591419 to 135591436

| Score | Expect | Identities | Gaps | Strand | Frame |
|---|---|---|---|---|---|
| 28.2 bits(14) | 256() | 17/18(94%) | 0/18(0%) | Plus/Minus | |

Features:
**492618 bp at 5' side: polyadenylate-binding protein 4-like2827251 bp at 3' side: protocadherin-18 isoform 2 precursor**

```
Query  279        GCTGGTGTTCTCTCCTTT  296
                  ||||||||||| ||||||
Sbjct  135591436  GCTGGTGTTCTTTCCTTT  135591419
```

Range 59: 159741125 to 159741138

| Score | Expect | Identities | Gaps | Strand | Frame |
|---|---|---|---|---|---|

| | | | | |
|---|---|---|---|---|
| 28.2 bits(14) | 256() | 14/14(100%) | 0/14(0%) | Plus/Minus |

Features:
**folliculin-interacting protein 2**

```
Query  287        TCTCTCCTTTATTG  300
                  ||||||||||||||
Sbjct  159741138  TCTCTCCTTTATTG  159741125
```

Range 60: 180433279 to 180433292

| Score | Expect | Identities | Gaps | Strand | Frame |
|---|---|---|---|---|---|
| 28.2 bits(14) | 256() | 14/14(100%) | 0/14(0%) | Plus/Minus | |

Features:
**1860118 bp at 5' side: uncharacterized protein LOC285500 isoform X22788417 bp at 3' side: teneurin-3**

```
Query  286        TTCTCTCCTTTATT  299
                  ||||||||||||||
Sbjct  180433292  TTCTCTCCTTTATT  180433279
```

Range 61: 183759885 to 183759898

| Score | Expect | Identities | Gaps | Strand | Frame |
|---|---|---|---|---|---|
| 28.2 bits(14) | 256() | 14/14(100%) | 0/14(0%) | Plus/Minus | |

Features:
**61804 bp at 5' side: teneurin-327239 bp at 3' side: deoxycytidylate deaminase isoform a**

```
Query  286        TTCTCTCCTTTATT  299
                  ||||||||||||||
Sbjct  183759898  TTCTCTCCTTTATT  183759885
```

Range 62: 190701550 to 190701563

| Score | Expect | Identities | Gaps | Strand | Frame |
|---|---|---|---|---|---|
| 28.2 bits(14) | 256() | 14/14(100%) | 0/14(0%) | Plus/Minus | |

Features:
**1656569 bp at 5' side: probable E3 ubiquitin-protein ligase TRIML1144933 bp at 3' side: protein FRG1**

```
Query  282        GGTGTTCTCTCCTT  295
                  ||||||||||||||
Sbjct  190701563  GGTGTTCTCTCCTT  190701550
```

Homo sapiens chromosome 5, alternate assembly CHM1_1.1
Sequence ID: **ref|NC_018916.2|** Length: 180347728 Number of Matches: 74
Range 1: 123806586 to 123806606

| Score | Expect | Identities | Gaps | Strand | Frame |
|---|---|---|---|---|---|
| 34.2 bits(17) | 4.1() | 20/21(95%) | 0/21(0%) | Plus/Plus | |

Features:
**293260 bp at 5' side: zinc finger protein 6081322067 bp at 3' side: GRAM domain-containing protein 3 isoform 1**

```
Query  277        AAGCTGGTGTTCTCTCCTTTA  297
                  ||||||||||| |||||||||
Sbjct  123806586  AAGCTGGTGTTTTCTCCTTTA  123806606
```

Range 2: 5607063 to 5607079

| Score | Expect | Identities | Gaps | Strand | Frame |
|---|---|---|---|---|---|
| 34.2 bits(17) | 4.1() | 17/17(100%) | 0/17(0%) | Plus/Minus | |

Features:
**117620 bp at 5' side: little elongation complex subunit 1765604 bp at 3' side: mediator of RNA polymerase II transcription subunit 10**

```
Query  284      TGTTCTCTCCTTTATTG  300
```

```
                   |||||||||||||||||
Sbjct  5607079  TGTTCTCTCCTTTATTG  5607063
```

Range 3: 166996253 to 166996269

| Score | Expect | Identities | Gaps | Strand | Frame |
|---|---|---|---|---|---|
| 34.2 bits(17) | 4.1() | 17/17(100%) | 0/17(0%) | Plus/Minus | |

Features:
**teneurin-2**

```
Query  281        TGGTGTTCTCTCCTTTA  297
                  |||||||||||||||||
Sbjct  166996269  TGGTGTTCTCTCCTTTA  166996253
```

Range 4: 95557158 to 95557173

| Score | Expect | Identities | Gaps | Strand | Frame |
|---|---|---|---|---|---|
| 32.2 bits(16) | 16() | 16/16(100%) | 0/16(0%) | Plus/Minus | |

Features:
**endoplasmic reticulum aminopeptidase 1 isoform a precursorendoplasmic reticulum aminopeptidase 1 isoform b precursor**

```
Query  277       AAGCTGGTGTTCTCTC  292
                 ||||||||||||||||
Sbjct  95557173  AAGCTGGTGTTCTCTC  95557158
```

Range 5: 11468823 to 11468837

| Score | Expect | Identities | Gaps | Strand | Frame |
|---|---|---|---|---|---|
| 30.2 bits(15) | 65() | 15/15(100%) | 0/15(0%) | Plus/Plus | |

Features:
**catenin delta-2 isoform 1catenin delta-2 isoform 3**

```
Query  282       GGTGTTCTCTCCTTT  296
                 |||||||||||||||
Sbjct  11468823  GGTGTTCTCTCCTTT  11468837
```

Range 6: 69405951 to 69405965

| Score | Expect | Identities | Gaps | Strand | Frame |
|---|---|---|---|---|---|
| 30.2 bits(15) | 65() | 15/15(100%) | 0/15(0%) | Plus/Plus | |

Features:
**17706 bp at 5' side: small EDRK-rich factor 1 isoform 1241873 bp at 3' side: small EDRK-rich factor 1 isoform 2**

```
Query  286       TTCTCTCCTTTATTG  300
                 |||||||||||||||
Sbjct  69405951  TTCTCTCCTTTATTG  69405965
```

Range 7: 77304845 to 77304859

| Score | Expect | Identities | Gaps | Strand | Frame |
|---|---|---|---|---|---|
| 30.2 bits(15) | 65() | 15/15(100%) | 0/15(0%) | Plus/Plus | |

Features:
**65498 bp at 5' side: lipoma HMGIC fusion partner-like 2 protein204383 bp at 3' side: arylsulfatase B isoform 1 precursor**

```
Query  285       GTTCTCTCCTTTATT  299
                 |||||||||||||||
Sbjct  77304845  GTTCTCTCCTTTATT  77304859
```

Range 8: 90330409 to 90330423

| Score | Expect | Identities | Gaps | Strand | Frame |
|---|---|---|---|---|---|
| 30.2 bits(15) | 65() | 15/15(100%) | 0/15(0%) | Plus/Plus | |

Features:
**218786 bp at 5' side: arrestin domain-containing protein 32023006 bp at 3' side: COUP transcription factor 1**

```
Query  279       GCTGGTGTTCTCTCC  293
                 |||||||||||||||
Sbjct  90330409  GCTGGTGTTCTCTCC  90330423
```

Range 9: 100677513 to 100677527

| Score | Expect | Identities | Gaps | Strand | Frame |
|---|---|---|---|---|---|
| 30.2 bits(15) | 65() | 15/15(100%) | 0/15(0%) | Plus/Plus | |

Features:
**1006235 bp at 5' side: CMP-N-acetylneuraminate-poly-alpha-2,8-sialyltransferase ...327677 bp at 3' side: solute carrier organic anion transporter family member 4C1**

```
Query  284        TGTTCTCTCCTTTAT  298
                  |||||||||||||||
Sbjct  100677513  TGTTCTCTCCTTTAT  100677527
```

Range 10: 117316409 to 117316423

| Score | Expect | Identities | Gaps | Strand | Frame |
|---|---|---|---|---|---|
| 30.2 bits(15) | 65() | 15/15(100%) | 0/15(0%) | Plus/Plus | |

Features:
**2043086 bp at 5' side: semaphorin-6A isoform 1 precursor292938 bp at 3' side: DTW domain-containing protein 2**

```
Query  278        AGCTGGTGTTCTCTC  292
                  |||||||||||||||
Sbjct  117316409  AGCTGGTGTTCTCTC  117316423
```

Range 11: 121336612 to 121336626

| Score | Expect | Identities | Gaps | Strand | Frame |
|---|---|---|---|---|---|
| 30.2 bits(15) | 65() | 15/15(100%) | 0/15(0%) | Plus/Plus | |

Features:
**104724 bp at 5' side: synphilin-1 isoform 1A206862 bp at 3' side: sorting nexin-2 isoform 1**

```
Query  284        TGTTCTCTCCTTTAT  298
                  |||||||||||||||
Sbjct  121336612  TGTTCTCTCCTTTAT  121336626
```

Range 12: 122677387 to 122677405

| Score | Expect | Identities | Gaps | Strand | Frame |
|---|---|---|---|---|---|
| 30.2 bits(15) | 65() | 18/19(95%) | 0/19(0%) | Plus/Plus | |

Features:
**294679 bp at 5' side: casein kinase I isoform gamma-3 isoform 7728867 bp at 3' side: zinc finger protein 608**

```
Query  282        GGTGTTCTCTCCTTTATTG  300
                  |||||| ||||||||||||
Sbjct  122677387  GGTGTTTTCTCCTTTATTG  122677405
```

Range 13: 131964907 to 131964921

| Score | Expect | Identities | Gaps | Strand | Frame |
|---|---|---|---|---|---|
| 30.2 bits(15) | 65() | 15/15(100%) | 0/15(0%) | Plus/Plus | |

Features:
**91452 bp at 5' side: heat shock 70 kDa protein 43196 bp at 3' side: follistatin-related protein 4 precursor**

```
Query  285        GTTCTCTCCTTTATT  299
                  |||||||||||||||
Sbjct  131964907  GTTCTCTCCTTTATT  131964921
```

Range 14: 150661958 to 150661972

| Score | Expect | Identities | Gaps | Strand | Frame |
|---|---|---|---|---|---|
| 30.2 bits(15) | 65() | 15/15(100%) | 0/15(0%) | Plus/Plus | |

Features:
**glycine receptor subunit alpha-1 isoform 1 precursorglycine receptor subunit alpha-1 isoform 3**

```
Query  275        ACAAGCTGGTGTTCT  289
                  |||||||||||||||
Sbjct  150661958  ACAAGCTGGTGTTCT  150661972
```

Range 15: 157237463 to 157237477

| Score | Expect | Identities | Gaps | Strand | Frame |
|---|---|---|---|---|---|
| 30.2 bits(15) | 65() | 15/15(100%) | 0/15(0%) | Plus/Plus | |

Features:
**560340 bp at 5' side: clathrin interactor 1 isoform 3321411 bp at 3' side: transcription factor COE1 isoform 3**

```
Query  286        TTCTCTCCTTTATTG  300
                  |||||||||||||||
Sbjct  157237463  TTCTCTCCTTTATTG  157237477
```

Range 16: 5107918 to 5107932

| Score | Expect | Identities | Gaps | Strand | Frame |
|---|---|---|---|---|---|
| 30.2 bits(15) | 65() | 15/15(100%) | 0/15(0%) | Plus/Minus | |

Features:
**1506664 bp at 5' side: iroquois-class homeodomain protein IRX-132680 bp at 3' side: A disintegrin and metalloproteinase with thrombospondin m...**

```
Query  280      CTGGTGTTCTCTCCT  294
                |||||||||||||||
Sbjct  5107932  CTGGTGTTCTCTCCT  5107918
```

Range 17: 21998380 to 21998394

| Score | Expect | Identities | Gaps | Strand | Frame |
|---|---|---|---|---|---|
| 30.2 bits(15) | 65() | 15/15(100%) | 0/15(0%) | Plus/Minus | |

Features:
**cadherin-12 preproprotein**

```
Query  286       TTCTCTCCTTTATTG  300
                 |||||||||||||||
Sbjct  21998394  TTCTCTCCTTTATTG  21998380
```

Range 18: 39823727 to 39823741

| Score | Expect | Identities | Gaps | Strand | Frame |
|---|---|---|---|---|---|
| 30.2 bits(15) | 65() | 15/15(100%) | 0/15(0%) | Plus/Minus | |

Features:
**427404 bp at 5' side: disabled homolog 2 isoform 1859315 bp at 3' side: prostaglandin E2 receptor EP4 subtype**

```
Query  285       GTTCTCTCCTTTATT  299
                 |||||||||||||||
Sbjct  39823741  GTTCTCTCCTTTATT  39823727
```

Range 19: 82116792 to 82116806

| Score | Expect | Identities | Gaps | Strand | Frame |
|---|---|---|---|---|---|
| 30.2 bits(15) | 65() | 15/15(100%) | 0/15(0%) | Plus/Minus | |

Features:
**35014 bp at 5' side: DNA repair protein XRCC4 isoform 195230 bp at 3' side: versican core protein isoform 2 precursor**

```
Query  284       TGTTCTCTCCTTTAT  298
                 |||||||||||||||
```

```
Sbjct  82116806  TGTTCTCTCCTTTAT  82116792
```

Range 20: 83543348 to 83543362

| Score | Expect | Identities | Gaps | Strand | Frame |
|---|---|---|---|---|---|
| 30.2 bits(15) | 65() | 15/15(100%) | 0/15(0%) | Plus/Minus | |

Features:
**430182 bp at 5' side: EGF-like repeat and discoidin I-like domain-containing pr...1803444 bp at 3' side: cytochrome c oxidase subunit 7C, mitochondrial precursor**

```
Query  286       TTCTCTCCTTTATTG  300
                 |||||||||||||||
Sbjct  83543362  TTCTCTCCTTTATTG  83543348
```

Range 21: 99250573 to 99250587

| Score | Expect | Identities | Gaps | Strand | Frame |
|---|---|---|---|---|---|
| 30.2 bits(15) | 65() | 15/15(100%) | 0/15(0%) | Plus/Minus | |

Features:
**89696 bp at 5' side: putative POM121-like protein 1-like isoform X152655 bp at 3' side: membrane protein FAM174A precursor**

```
Query  286       TTCTCTCCTTTATTG  300
                 |||||||||||||||
Sbjct  99250587  TTCTCTCCTTTATTG  99250573
```

Range 22: 114570011 to 114570025

| Score | Expect | Identities | Gaps | Strand | Frame |
|---|---|---|---|---|---|
| 30.2 bits(15) | 65() | 15/15(100%) | 0/15(0%) | Plus/Minus | |

Features:
**175892 bp at 5' side: transmembrane emp24 domain-containing protein 7 precursor3716 bp at 3' side: cysteine dioxygenase type 1**

```
Query  286        TTCTCTCCTTTATTG  300
                  |||||||||||||||
Sbjct  114570025  TTCTCTCCTTTATTG  114570011
```

Range 23: 3680050 to 3680063

| Score | Expect | Identities | Gaps | Strand | Frame |
|---|---|---|---|---|---|
| 28.2 bits(14) | 256() | 14/14(100%) | 0/14(0%) | Plus/Plus | |

Features:
**78796 bp at 5' side: iroquois-class homeodomain protein IRX-11460549 bp at 3' side: A disintegrin and metalloproteinase with thrombospondin m...**

```
Query  286      TTCTCTCCTTTATT  299
                ||||||||||||||
Sbjct  3680050  TTCTCTCCTTTATT  3680063
```

Range 24: 6091313 to 6091326

| Score | Expect | Identities | Gaps | Strand | Frame |
|---|---|---|---|---|---|
| 28.2 bits(14) | 256() | 14/14(100%) | 0/14(0%) | Plus/Plus | |

Features:
**601870 bp at 5' side: little elongation complex subunit 1281357 bp at 3' side: mediator of RNA polymerase II transcription subunit 10**

```
Query  286      TTCTCTCCTTTATT  299
                ||||||||||||||
Sbjct  6091313  TTCTCTCCTTTATT  6091326
```

Range 25: 11492265 to 11492282

| Score | Expect | Identities | Gaps | Strand | Frame |
|---|---|---|---|---|---|

| Score | Expect | Identities | Gaps | Strand | Frame |
|---|---|---|---|---|---|
| 28.2 bits(14) | 256() | 17/18(94%) | 0/18(0%) | Plus/Plus | |

Features:
**catenin delta-2 isoform 1catenin delta-2 isoform 3**

```
Query  280       CTGGTGTTCTCTCCTTTA  297
                 ||||||||||| ||||||
Sbjct  11492265  CTGGTGTTCTCGCCTTTA  11492282
```

Range 26: 13278228 to 13278241

| Score | Expect | Identities | Gaps | Strand | Frame |
|---|---|---|---|---|---|
| 28.2 bits(14) | 256() | 14/14(100%) | 0/14(0%) | Plus/Plus | |

Features:
**1913539 bp at 5' side: catenin delta-2 isoform 4413723 bp at 3' side: dynein heavy chain 5, axonemal**

```
Query  286       TTCTCTCCTTTATT  299
                 ||||||||||||||
Sbjct  13278228  TTCTCTCCTTTATT  13278241
```

Range 27: 13996449 to 13996462

| Score | Expect | Identities | Gaps | Strand | Frame |
|---|---|---|---|---|---|
| 28.2 bits(14) | 256() | 14/14(100%) | 0/14(0%) | Plus/Plus | |

Features:
**52335 bp at 5' side: dynein heavy chain 5, axonemal147474 bp at 3' side: triple functional domain protein**

```
Query  281       TGGTGTTCTCTCCT  294
                 ||||||||||||||
Sbjct  13996449  TGGTGTTCTCTCCT  13996462
```

Range 28: 14638862 to 14638875

| Score | Expect | Identities | Gaps | Strand | Frame |
|---|---|---|---|---|---|
| 28.2 bits(14) | 256() | 14/14(100%) | 0/14(0%) | Plus/Plus | |

Features:
**28567 bp at 5' side: inactive ubiquitin thioesterase FAM105A25828 bp at 3' side: ubiquitin thioesterase otulin**

```
Query  283       GTGTTCTCTCCTTT  296
                 ||||||||||||||
Sbjct  14638862  GTGTTCTCTCCTTT  14638875
```

Range 29: 22195896 to 22195909

| Score | Expect | Identities | Gaps | Strand | Frame |
|---|---|---|---|---|---|
| 28.2 bits(14) | 256() | 14/14(100%) | 0/14(0%) | Plus/Plus | |

Features:
**117114 bp at 5' side: cadherin-12 preproprotein1313256 bp at 3' side: histone-lysine N-methyltransferase PRDM9**

```
Query  286       TTCTCTCCTTTATT  299
                 ||||||||||||||
Sbjct  22195896  TTCTCTCCTTTATT  22195909
```

Range 30: 39648101 to 39648118

| Score | Expect | Identities | Gaps | Strand | Frame |
|---|---|---|---|---|---|
| 28.2 bits(14) | 256() | 17/18(94%) | 0/18(0%) | Plus/Plus | |

Features:
**251778 bp at 5' side: disabled homolog 2 isoform 11034938 bp at 3' side: prostaglandin E2 receptor EP4 subtype**

```
Query  281       TGGTGTTCTCTCCTTTAT  298
                 ||||||| ||||||||||
Sbjct  39648101  TGGTGTTTTCTCCTTTAT  39648118
```

Range 31: 44910904 to 44910917

| Score | Expect | Identities | Gaps | Strand | Frame |
|---|---|---|---|---|---|
| 28.2 bits(14) | 256() | 14/14(100%) | 0/14(0%) | Plus/Plus | |

Features:
**94281 bp at 5' side: 28S ribosomal protein S30, mitochondrial350951 bp at 3' side: potassium/sodium hyperpolarization-activated cyclic nucle...**

```
Query  286       TTCTCTCCTTTATT  299
                 ||||||||||||||
Sbjct  44910904  TTCTCTCCTTTATT  44910917
```

Range 32: 55793108 to 55793121

| Score | Expect | Identities | Gaps | Strand | Frame |
|---|---|---|---|---|---|
| 28.2 bits(14) | 256() | 14/14(100%) | 0/14(0%) | Plus/Plus | |

Features:
**261387 bp at 5' side: ankyrin repeat domain-containing protein 5517410 bp at 3' side: uncharacterized protein LOC101928448**

```
Query  278       AGCTGGTGTTCTCT  291
                 ||||||||||||||
Sbjct  55793108  AGCTGGTGTTCTCT  55793121
```

Range 33: 58799978 to 58799991

| Score | Expect | Identities | Gaps | Strand | Frame |
|---|---|---|---|---|---|
| 28.2 bits(14) | 256() | 14/14(100%) | 0/14(0%) | Plus/Plus | |

Features:
**cAMP-specific 3',5'-cyclic phosphodiesterase 4D isoform P...cAMP-specific 3',5'-cyclic phosphodiesterase 4D isoform P...**

```
Query  283       GTGTTCTCTCCTTT  296
                 ||||||||||||||
Sbjct  58799978  GTGTTCTCTCCTTT  58799991
```

Range 34: 65778618 to 65778631

| Score | Expect | Identities | Gaps | Strand | Frame |
|---|---|---|---|---|---|
| 28.2 bits(14) | 256() | 14/14(100%) | 0/14(0%) | Plus/Plus | |

Features:
**303788 bp at 5' side: splicing regulatory glutamine/lysine-rich protein 1 isofo...113753 bp at 3' side: microtubule-associated serine/threonine-protein kinase 4 ...**

```
Query  285       GTTCTCTCCTTTAT  298
                 ||||||||||||||
Sbjct  65778618  GTTCTCTCCTTTAT  65778631
```

Range 35: 66359781 to 66359794

| Score | Expect | Identities | Gaps | Strand | Frame |
|---|---|---|---|---|---|
| 28.2 bits(14) | 256() | 14/14(100%) | 0/14(0%) | Plus/Plus | |

Features:
**microtubule-associated serine/threonine-protein kinase 4 ...microtubule-associated serine/threonine-protein kinase 4 ...**

```
Query  286       TTCTCTCCTTTATT  299
                 ||||||||||||||
Sbjct  66359781  TTCTCTCCTTTATT  66359794
```

Range 36: 68082964 to 68082977

| Score | Expect | Identities | Gaps | Strand | Frame |
|---|---|---|---|---|---|
| 28.2 bits(14) | 256() | 14/14(100%) | 0/14(0%) | Plus/Plus | |

Features:
**489437 bp at 5' side: phosphatidylinositol 3-kinase regulatory subunit alpha is...306790 bp at 3' side: zinc transporter 5 isoform 1**

```
Query  286       TTCTCTCCTTTATT  299
                 ||||||||||||||
Sbjct  68082964  TTCTCTCCTTTATT  68082977
```

Range 37: 70157426 to 70157439

| Score | Expect | Identities | Gaps | Strand | Frame |
|---|---|---|---|---|---|
| 28.2 bits(14) | 256() | 14/14(100%) | 0/14(0%) | Plus/Plus | |

Features:
**347725 bp at 5' side: general transcription factor IIH subunit 226983 bp at 3' side: transcription factor TFIIIB component B" homolog**

```
Query  286       TTCTCTCCTTTATT  299
                 ||||||||||||||
Sbjct  70157426  TTCTCTCCTTTATT  70157439
```

Range 38: 72234367 to 72234384

| Score | Expect | Identities | Gaps | Strand | Frame |
|---|---|---|---|---|---|
| 28.2 bits(14) | 256() | 17/18(94%) | 0/18(0%) | Plus/Plus | |

Features:
**578 bp at 5' side: transcription factor BTF3 isoform B46970 bp at 3' side: ankyrin repeat family A protein 2**

```
Query  282       GGTGTTCTCTCCTTTATT  299
                 |||||| |||||||||||
Sbjct  72234367  GGTGTTATCTCCTTTATT  72234384
```

Range 39: 73690442 to 73690455

| Score | Expect | Identities | Gaps | Strand | Frame |
|---|---|---|---|---|---|
| 28.2 bits(14) | 256() | 14/14(100%) | 0/14(0%) | Plus/Plus | |

Features:
**120083 bp at 5' side: soluble lamin-associated protein of 75 kDa66818 bp at 3' side: beta-1,3-galactosyl-O-glycosyl-glycoprotein beta-1,6-N-ac...**

```
Query  286       TTCTCTCCTTTATT  299
                 ||||||||||||||
Sbjct  73690442  TTCTCTCCTTTATT  73690455
```

Range 40: 88477804 to 88477817

| Score | Expect | Identities | Gaps | Strand | Frame |
|---|---|---|---|---|---|
| 28.2 bits(14) | 256() | 14/14(100%) | 0/14(0%) | Plus/Plus | |

Features:
**925513 bp at 5' side: myocyte-specific enhancer factor 2C isoform 1644743 bp at 3' side: centrin-3 isoform 1**

```
Query  283       GTGTTCTCTCCTTT  296
                 ||||||||||||||
Sbjct  88477804  GTGTTCTCTCCTTT  88477817
```

Range 41: 122158724 to 122158737

| Score | Expect | Identities | Gaps | Strand | Frame |
|---|---|---|---|---|---|
| 28.2 bits(14) | 256() | 14/14(100%) | 0/14(0%) | Plus/Plus | |

Features:
**centrosomal protein of 120 kDa isoform 2centrosomal protein of 120 kDa isoform 1**

```
Query  286        TTCTCTCCTTTATT  299
                  ||||||||||||||
Sbjct  122158724  TTCTCTCCTTTATT  122158737
```

Range 42: 124283903 to 124283916

| Score | Expect | Identities | Gaps | Strand | Frame |
|---|---|---|---|---|---|

| | | | | |
|---|---|---|---|---|
| 28.2 bits(14) | 256() | 14/14(100%) | 0/14(0%) | Plus/Plus |

Features:
**770577 bp at 5' side: zinc finger protein 608844757 bp at 3' side: GRAM domain-containing protein 3 isoform 1**

```
Query  286        TTCTCTCCTTTATT  299
                  ||||||||||||||
Sbjct  124283903  TTCTCTCCTTTATT  124283916
```

Range 43: 128255380 to 128255393

| Score | Expect | Identities | Gaps | Strand | Frame |
|---|---|---|---|---|---|
| 28.2 bits(14) | 256() | 14/14(100%) | 0/14(0%) | Plus/Plus | |

Features:
**A disintegrin and metalloproteinase with thrombospondin m...**

```
Query  286        TTCTCTCCTTTATT  299
                  ||||||||||||||
Sbjct  128255380  TTCTCTCCTTTATT  128255393
```

Range 44: 130085924 to 130085937

| Score | Expect | Identities | Gaps | Strand | Frame |
|---|---|---|---|---|---|
| 28.2 bits(14) | 256() | 14/14(100%) | 0/14(0%) | Plus/Plus | |

Features:
**117666 bp at 5' side: complex III assembly factor LYRM7 isoform 142160 bp at 3' side: CDC42 small effector protein 2**

```
Query  285        GTTCTCTCCTTTAT  298
                  ||||||||||||||
Sbjct  130085924  GTTCTCTCCTTTAT  130085937
```

Range 45: 136356715 to 136356728

| Score | Expect | Identities | Gaps | Strand | Frame |
|---|---|---|---|---|---|
| 28.2 bits(14) | 256() | 14/14(100%) | 0/14(0%) | Plus/Plus | |

Features:
**89778 bp at 5' side: testican-1 precursor31994 bp at 3' side: kelch-like protein 3 isoform 3**

```
Query  286        TTCTCTCCTTTATT  299
                  ||||||||||||||
Sbjct  136356715  TTCTCTCCTTTATT  136356728
```

Range 46: 141579367 to 141579380

| Score | Expect | Identities | Gaps | Strand | Frame |
|---|---|---|---|---|---|
| 28.2 bits(14) | 256() | 14/14(100%) | 0/14(0%) | Plus/Plus | |

Features:
**152012 bp at 5' side: fibroblast growth factor 1 isoform 2 precursor4613 bp at 3' side: rho GTPase-activating protein 26 isoform a**

```
Query  275        ACAAGCTGGTGTTC  288
                  ||||||||||||||
Sbjct  141579367  ACAAGCTGGTGTTC  141579380
```

Range 47: 146290606 to 146290619

| Score | Expect | Identities | Gaps | Strand | Frame |
|---|---|---|---|---|---|
| 28.2 bits(14) | 256() | 14/14(100%) | 0/14(0%) | Plus/Plus | |

Features:
**dihydropyrimidinase-related protein 3 isoform 1**

```
Query  287        TCTCTCCTTTATTG  300
                  ||||||||||||||
Sbjct  146290606  TCTCTCCTTTATTG  146290619
```

Range 48: 149882763 to 149882776

| Score | Expect | Identities | Gaps | Strand | Frame |
|---|---|---|---|---|---|
| 28.2 bits(14) | 256() | 14/14(100%) | 0/14(0%) | Plus/Plus | |

Features:
**5312 bp at 5' side: TNFAIP3-interacting protein 1 isoform 131028 bp at 3' side: annexin A6 isoform 1**

```
Query  284        TGTTCTCTCCTTTA  297
                  ||||||||||||||
Sbjct  149882763  TGTTCTCTCCTTTA  149882776
```

Range 49: 150935473 to 150935486

| Score | Expect | Identities | Gaps | Strand | Frame |
|---|---|---|---|---|---|
| 28.2 bits(14) | 256() | 14/14(100%) | 0/14(0%) | Plus/Plus | |

Features:
**198924 bp at 5' side: glycine receptor subunit alpha-1 isoform 2 precursor268813 bp at 3' side: neuromedin-U receptor 2**

```
Query  283        GTGTTCTCTCCTTT  296
                  ||||||||||||||
Sbjct  150935473  GTGTTCTCTCCTTT  150935486
```

Range 50: 154965264 to 154965277

| Score | Expect | Identities | Gaps | Strand | Frame |
|---|---|---|---|---|---|
| 28.2 bits(14) | 256() | 14/14(100%) | 0/14(0%) | Plus/Plus | |

Features:
**1135726 bp at 5' side: chromosome-associated kinesin KIF4B223731 bp at 3' side: delta-sarcoglycan isoform 1**

```
Query  284        TGTTCTCTCCTTTA  297
                  ||||||||||||||
Sbjct  154965264  TGTTCTCTCCTTTA  154965277
```

Range 51: 173845083 to 173845096

| Score | Expect | Identities | Gaps | Strand | Frame |
|---|---|---|---|---|---|
| 28.2 bits(14) | 256() | 14/14(100%) | 0/14(0%) | Plus/Plus | |

Features:
**256102 bp at 5' side: homeobox protein MSX-2456090 bp at 3' side: D(1A) dopamine receptor**

```
Query  283        GTGTTCTCTCCTTT  296
                  ||||||||||||||
Sbjct  173845083  GTGTTCTCTCCTTT  173845096
```

Range 52: 179940004 to 179940017

| Score | Expect | Identities | Gaps | Strand | Frame |
|---|---|---|---|---|---|
| 28.2 bits(14) | 256() | 14/14(100%) | 0/14(0%) | Plus/Plus | |

Features:
**20918 bp at 5' side: butyrophilin-like protein 9 precursor43565 bp at 3' side: olfactory receptor 2V1**

```
Query  282        GGTGTTCTCTCCTT  295
                  ||||||||||||||
Sbjct  179940004  GGTGTTCTCTCCTT  179940017
```

Range 53: 3329292 to 3329305

| Score | Expect | Identities | Gaps | Strand | Frame |
|---|---|---|---|---|---|
| 28.2 bits(14) | 256() | 14/14(100%) | 0/14(0%) | Plus/Minus | |

Features:
**368744 bp at 5' side: uncharacterized protein LOC105374620 isoform X3267016 bp at 3' side: iroquois-class homeodomain protein IRX-1**

```
Query  283      GTGTTCTCTCCTTT  296
```

```
                  ||||||||||||||
Sbjct  3329305  GTGTTCTCTCCTTT  3329292
```

Range 54: 9198468 to 9198489

| Score | Expect | Identities | Gaps | Strand | Frame |
|---|---|---|---|---|---|
| 28.2 bits(14) | 256() | 20/22(91%) | 0/22(0%) | Plus/Minus | |

Features:
**semaphorin-5A precursor**

```
Query  278      AGCTGGTGTTCTCTCCTTTATT  299
                ||||||||||| | ||||||||
Sbjct  9198489  AGCTGGTGTTCGCCCCTTTATT  9198468
```

Range 55: 10060919 to 10060932

| Score | Expect | Identities | Gaps | Strand | Frame |
|---|---|---|---|---|---|
| 28.2 bits(14) | 256() | 14/14(100%) | 0/14(0%) | Plus/Minus | |

Features:
**430816 bp at 5' side: taste receptor type 2 member 1166498 bp at 3' side: protein FAM173B isoform 2**

```
Query  286       TTCTCTCCTTTATT  299
                 ||||||||||||||
Sbjct  10060932  TTCTCTCCTTTATT  10060919
```

Range 56: 13376018 to 13376031

| Score | Expect | Identities | Gaps | Strand | Frame |
|---|---|---|---|---|---|
| 28.2 bits(14) | 256() | 14/14(100%) | 0/14(0%) | Plus/Minus | |

Features:
**2011329 bp at 5' side: catenin delta-2 isoform 4315933 bp at 3' side: dynein heavy chain 5, axonemal**

```
Query  285       GTTCTCTCCTTTAT  298
                 ||||||||||||||
Sbjct  13376031  GTTCTCTCCTTTAT  13376018
```

Range 57: 20292396 to 20292409

| Score | Expect | Identities | Gaps | Strand | Frame |
|---|---|---|---|---|---|
| 28.2 bits(14) | 256() | 14/14(100%) | 0/14(0%) | Plus/Minus | |

Features:
**453435 bp at 5' side: cadherin-18 isoform 3 preproprotein1186478 bp at 3' side: putative POM121-like protein 1-like isoform X1**

```
Query  286       TTCTCTCCTTTATT  299
                 ||||||||||||||
Sbjct  20292409  TTCTCTCCTTTATT  20292396
```

Range 58: 27286307 to 27286320

| Score | Expect | Identities | Gaps | Strand | Frame |
|---|---|---|---|---|---|
| 28.2 bits(14) | 256() | 14/14(100%) | 0/14(0%) | Plus/Minus | |

Features:
**297557 bp at 5' side: cadherin-9 preproprotein1522686 bp at 3' side: uncharacterized protein LOC105374700**

```
Query  285       GTTCTCTCCTTTAT  298
                 ||||||||||||||
Sbjct  27286320  GTTCTCTCCTTTAT  27286307
```

Range 59: 44636316 to 44636329

| Score | Expect | Identities | Gaps | Strand | Frame |
|---|---|---|---|---|---|
| 28.2 bits(14) | 256() | 14/14(100%) | 0/14(0%) | Plus/Minus | |

Features:
**246120 bp at 5' side: fibroblast growth factor 10 precursor174057 bp at 3' side: 28S ribosomal protein S30, mitochondrial**

```
Query  286       TTCTCTCCTTTATT  299
                 ||||||||||||||
Sbjct  44636329  TTCTCTCCTTTATT  44636316
```

Range 60: 45738033 to 45738046

| Score | Expect | Identities | Gaps | Strand | Frame |
|---|---|---|---|---|---|
| 28.2 bits(14) | 256() | 14/14(100%) | 0/14(0%) | Plus/Minus | |

Features:
**41987 bp at 5' side: potassium/sodium hyperpolarization-activated cyclic nucle...3959116 bp at 3' side: embigin precursor**

```
Query  286       TTCTCTCCTTTATT  299
                 ||||||||||||||
Sbjct  45738046  TTCTCTCCTTTATT  45738033
```

Range 61: 85586674 to 85586687

| Score | Expect | Identities | Gaps | Strand | Frame |
|---|---|---|---|---|---|
| 28.2 bits(14) | 256() | 14/14(100%) | 0/14(0%) | Plus/Minus | |

Features:
**238455 bp at 5' side: cytochrome c oxidase subunit 7C, mitochondrial precursor410644 bp at 3' side: ras GTPase-activating protein 1 isoform 1**

```
Query  287       TCTCTCCTTTATTG  300
                 ||||||||||||||
Sbjct  85586687  TCTCTCCTTTATTG  85586674
```

Range 62: 98321908 to 98321921

| Score | Expect | Identities | Gaps | Strand | Frame |
|---|---|---|---|---|---|
| 28.2 bits(14) | 256() | 14/14(100%) | 0/14(0%) | Plus/Minus | |

Features:
**28259 bp at 5' side: putative POM121-like protein 1-like isoform X1835116 bp at 3' side: putative POM121-like protein 1-like isoform X2**

```
Query  284       TGTTCTCTCCTTTA  297
                 ||||||||||||||
Sbjct  98321921  TGTTCTCTCCTTTA  98321908
```

Range 63: 108644707 to 108644720

| Score | Expect | Identities | Gaps | Strand | Frame |
|---|---|---|---|---|---|
| 28.2 bits(14) | 256() | 14/14(100%) | 0/14(0%) | Plus/Minus | |

Features:
**8684 bp at 5' side: alpha-mannosidase 2544882 bp at 3' side: transmembrane protein 232**

```
Query  280        CTGGTGTTCTCTCC  293
                  ||||||||||||||
Sbjct  108644720  CTGGTGTTCTCTCC  108644707
```

Range 64: 115773570 to 115773583

| Score | Expect | Identities | Gaps | Strand | Frame |
|---|---|---|---|---|---|
| 28.2 bits(14) | 256() | 14/14(100%) | 0/14(0%) | Plus/Minus | |

Features:
**500247 bp at 5' side: semaphorin-6A isoform 1 precursor1835778 bp at 3' side: DTW domain-containing protein 2**

```
Query  286        TTCTCTCCTTTATT  299
                  ||||||||||||||
Sbjct  115773583  TTCTCTCCTTTATT  115773570
```

Range 65: 122873794 to 122873807

| Score | Expect | Identities | Gaps | Strand | Frame |
|---|---|---|---|---|---|
| 28.2 bits(14) | 256() | 14/14(100%) | 0/14(0%) | Plus/Minus | |

Features:
**491086 bp at 5' side: casein kinase I isoform gamma-3 isoform 7532465 bp at 3' side: zinc finger protein 608**

```
Query  287        TCTCTCCTTTATTG  300
                  ||||||||||||||
Sbjct  122873807  TCTCTCCTTTATTG  122873794
```

Range 66: 123313304 to 123313317

| Score | Expect | Identities | Gaps | Strand | Frame |
|---|---|---|---|---|---|
| 28.2 bits(14) | 256() | 14/14(100%) | 0/14(0%) | Plus/Minus | |

Features:
**930596 bp at 5' side: casein kinase I isoform gamma-3 isoform 792955 bp at 3' side: zinc finger protein 608**

```
Query  280        CTGGTGTTCTCTCC  293
                  ||||||||||||||
Sbjct  123313317  CTGGTGTTCTCTCC  123313304
```

Range 67: 126166143 to 126166156

| Score | Expect | Identities | Gaps | Strand | Frame |
|---|---|---|---|---|---|
| 28.2 bits(14) | 256() | 14/14(100%) | 0/14(0%) | Plus/Minus | |

Features:
**multiple epidermal growth factor-like domains protein 10 ...multiple epidermal growth factor-like domains protein 10 ...**

```
Query  281        TGGTGTTCTCTCCT  294
                  ||||||||||||||
Sbjct  126166156  TGGTGTTCTCTCCT  126166143
```

Range 68: 129626558 to 129626571

| Score | Expect | Identities | Gaps | Strand | Frame |
|---|---|---|---|---|---|
| 28.2 bits(14) | 256() | 14/14(100%) | 0/14(0%) | Plus/Minus | |

Features:
**671777 bp at 5' side: chondroitin sulfate synthase 3301562 bp at 3' side: histidine triad nucleotide-binding protein 1**

```
Query  283        GTGTTCTCTCCTTT  296
                  ||||||||||||||
Sbjct  129626571  GTGTTCTCTCCTTT  129626558
```

Range 69: 136699241 to 136699254

| Score | Expect | Identities | Gaps | Strand | Frame |
|---|---|---|---|---|---|
| 28.2 bits(14) | 256() | 14/14(100%) | 0/14(0%) | Plus/Minus | |

Features:
**polycystic kidney disease 2-like 2 protein isoform 1polycystic kidney disease 2-like 2 protein isoform 4**

```
Query  286        TTCTCTCCTTTATT  299
                  ||||||||||||||
Sbjct  136699254  TTCTCTCCTTTATT  136699241
```

Range 70: 141507556 to 141507573

| Score | Expect | Identities | Gaps | Strand | Frame |
|---|---|---|---|---|---|
| 28.2 bits(14) | 256() | 17/18(94%) | 0/18(0%) | Plus/Minus | |

Features:
**80201 bp at 5' side: fibroblast growth factor 1 isoform 2 precursor76420 bp at 3' side: rho GTPase-activating protein 26 isoform a**

```
Query  278        AGCTGGTGTTCTCTCCTT  295
                  ||||||||||||| ||||
```

```
Sbjct  141507573  AGCTGGTGTTCTCACCTT  141507556
```

Range 71: 157209192 to 157209205

| Score | Expect | Identities | Gaps | Strand | Frame |
|---|---|---|---|---|---|
| 28.2 bits(14) | 256() | 14/14(100%) | 0/14(0%) | Plus/Minus | |

Features:
**532069 bp at 5' side: clathrin interactor 1 isoform 3349683 bp at 3' side: transcription factor COE1 isoform 3**

```
Query  278        AGCTGGTGTTCTCT  291
                  ||||||||||||||
Sbjct  157209205  AGCTGGTGTTCTCT  157209192
```

Range 72: 157773736 to 157773749

| Score | Expect | Identities | Gaps | Strand | Frame |
|---|---|---|---|---|---|
| 28.2 bits(14) | 256() | 14/14(100%) | 0/14(0%) | Plus/Minus | |

Features:
**transcription factor COE1 isoform 3transcription factor COE1 isoform 2**

```
Query  273        TAACAAGCTGGTGT  286
                  ||||||||||||||
Sbjct  157773749  TAACAAGCTGGTGT  157773736
```

Range 73: 170338575 to 170338588

| Score | Expect | Identities | Gaps | Strand | Frame |
|---|---|---|---|---|---|
| 28.2 bits(14) | 256() | 14/14(100%) | 0/14(0%) | Plus/Minus | |

Features:
**22237 bp at 5' side: fibroblast growth factor 18 precursor306770 bp at 3' side: small integral membrane protein 23**

```
Query  287        TCTCTCCTTTATTG  300
                  ||||||||||||||
Sbjct  170338588  TCTCTCCTTTATTG  170338575
```

Range 74: 170923389 to 170923406

| Score | Expect | Identities | Gaps | Strand | Frame |
|---|---|---|---|---|---|
| 28.2 bits(14) | 256() | 17/18(94%) | 0/18(0%) | Plus/Minus | |

Features:
**serine/threonine-protein kinase 10**

```
Query  276        CAAGCTGGTGTTCTCTCC  293
                  |||||| |||||||||||
Sbjct  170923406  CAAGCTTGTGTTCTCTCC  170923389
```

Homo sapiens chromosome 7, alternate assembly CHM1_1.1
Sequence ID: **ref|NC_018918.2|** Length: 159147065 Number of Matches: 51
Range 1: 1876161 to 1876177

| Score | Expect | Identities | Gaps | Strand | Frame |
|---|---|---|---|---|---|
| 34.2 bits(17) | 4.1() | 17/17(100%) | 0/17(0%) | Plus/Minus | |

Features:
**mitotic spindle assembly checkpoint protein MAD1 isoform amitotic spindle assembly checkpoint protein MAD1 isoform a**

```
Query  279      GCTGGTGTTCTCTCCTT  295
                |||||||||||||||||
Sbjct  1876177  GCTGGTGTTCTCTCCTT  1876161
```

Range 2: 457086 to 457104

| Score | Expect | Identities | Gaps | Strand | Frame |
|---|---|---|---|---|---|

| | | | | |
|---|---|---|---|---|
| 30.2 bits(15) | 65() | 18/19(95%) | 0/19(0%) | Plus/Plus |

Features:
**126759 bp at 5' side: forkhead box L1-like79851 bp at 3' side: platelet-derived growth factor subunit A isoform 2 prepro...**

```
Query  271     TGTAACAAGCTGGTGTTCT  289
               |||||||| ||||||||||
Sbjct  457086  TGTAACAACCTGGTGTTCT  457104
```

Range 3: 22318083 to 22318097

| Score | Expect | Identities | Gaps | Strand | Frame |
|---|---|---|---|---|---|
| 30.2 bits(15) | 65() | 15/15(100%) | 0/15(0%) | Plus/Plus | |

Features:
**rap guanine nucleotide exchange factor 5**

```
Query  285       GTTCTCTCCTTTATT  299
                 |||||||||||||||
Sbjct  22318083  GTTCTCTCCTTTATT  22318097
```

Range 4: 49461683 to 49461697

| Score | Expect | Identities | Gaps | Strand | Frame |
|---|---|---|---|---|---|
| 30.2 bits(15) | 65() | 15/15(100%) | 0/15(0%) | Plus/Plus | |

Features:
**569129 bp at 5' side: uncharacterized protein LOC105379727355627 bp at 3' side: brorin precursor**

```
Query  277       AAGCTGGTGTTCTCT  291
                 |||||||||||||||
Sbjct  49461683  AAGCTGGTGTTCTCT  49461697
```

Range 5: 79100673 to 79100687

| Score | Expect | Identities | Gaps | Strand | Frame |
|---|---|---|---|---|---|
| 30.2 bits(15) | 65() | 15/15(100%) | 0/15(0%) | Plus/Plus | |

Features:
**87838 bp at 5' side: membrane-associated guanylate kinase, WW and PDZ domain-c...593859 bp at 3' side: guanine nucleotide-binding protein G(i) subunit alpha-1 i...**

```
Query  286       TTCTCTCCTTTATTG  300
                 |||||||||||||||
Sbjct  79100673  TTCTCTCCTTTATTG  79100687
```

Range 6: 104043412 to 104043426

| Score | Expect | Identities | Gaps | Strand | Frame |
|---|---|---|---|---|---|
| 30.2 bits(15) | 65() | 15/15(100%) | 0/15(0%) | Plus/Plus | |

Features:
**lipoma HMGIC fusion partner-like 3 protein**

```
Query  281        TGGTGTTCTCTCCTT  295
                  |||||||||||||||
Sbjct  104043412  TGGTGTTCTCTCCTT  104043426
```

Range 7: 146107544 to 146107558

| Score | Expect | Identities | Gaps | Strand | Frame |
|---|---|---|---|---|---|
| 30.2 bits(15) | 65() | 15/15(100%) | 0/15(0%) | Plus/Plus | |

Features:
**contactin-associated protein-like 2 precursor**

```
Query  283        GTGTTCTCTCCTTTA  297
                  |||||||||||||||
Sbjct  146107544  GTGTTCTCTCCTTTA  146107558
```

Range 8: 86570722 to 86570736

| Score | Expect | Identities | Gaps | Strand | Frame |
|---|---|---|---|---|---|
| 30.2 bits(15) | 65() | 15/15(100%) | 0/15(0%) | Plus/Minus | |

Features:
**UPF0577 protein KIAA1324-like isoform 1 precursor**

```
Query  285       GTTCTCTCCTTTATT  299
                 |||||||||||||||
Sbjct  86570736  GTTCTCTCCTTTATT  86570722
```

Range 9: 93653025 to 93653039

| Score | Expect | Identities | Gaps | Strand | Frame |
|---|---|---|---|---|---|
| 30.2 bits(15) | 65() | 15/15(100%) | 0/15(0%) | Plus/Minus | |

Features:
**89264 bp at 5' side: BET1 homolog301404 bp at 3' side: collagen alpha-2(I) chain precursor**

```
Query  283       GTGTTCTCTCCTTTA  297
                 |||||||||||||||
Sbjct  93653039  GTGTTCTCTCCTTTA  93653025
```

Range 10: 16102717 to 16102734

| Score | Expect | Identities | Gaps | Strand | Frame |
|---|---|---|---|---|---|
| 28.2 bits(14) | 256() | 17/18(94%) | 0/18(0%) | Plus/Plus | |

Features:
**377095 bp at 5' side: homeobox protein MOX-228298 bp at 3' side: isoprenoid synthase domain-containing protein isoform b**

```
Query  278       AGCTGGTGTTCTCTCCTT  295
                 |||||||| |||||||||
Sbjct  16102717  AGCTGGTGCTCTCTCCTT  16102734
```

Range 11: 20432293 to 20432306

| Score | Expect | Identities | Gaps | Strand | Frame |
|---|---|---|---|---|---|
| 28.2 bits(14) | 256() | 14/14(100%) | 0/14(0%) | Plus/Plus | |

Features:
**integrin beta-8 precursor**

```
Query  275       ACAAGCTGGTGTTC  288
                 ||||||||||||||
Sbjct  20432293  ACAAGCTGGTGTTC  20432306
```

Range 12: 23663159 to 23663176

| Score | Expect | Identities | Gaps | Strand | Frame |
|---|---|---|---|---|---|
| 28.2 bits(14) | 256() | 17/18(94%) | 0/18(0%) | Plus/Plus | |

Features:
**coiled-coil domain-containing protein 126 precursor**

```
Query  281       TGGTGTTCTCTCCTTTAT  298
                 ||||| ||||||||||||
Sbjct  23663159  TGGTGATCTCTCCTTTAT  23663176
```

Range 13: 31881911 to 31881924

| Score | Expect | Identities | Gaps | Strand | Frame |
|---|---|---|---|---|---|
| 28.2 bits(14) | 256() | 14/14(100%) | 0/14(0%) | Plus/Plus | |

Features:
**calcium/calmodulin-dependent 3',5'-cyclic nucleotide phos...calcium/calmodulin-dependent 3',5'-cyclic nucleotide phos...**

```
Query  283       GTGTTCTCTCCTTT  296
                 ||||||||||||||
```

```
Sbjct  31881911  GTGTTCTCTCCTTT  31881924
```

Range 14: 34518917 to 34518930

| Score | Expect | Identities | Gaps | Strand | Frame |
|---|---|---|---|---|---|
| 28.2 bits(14) | 256() | 14/14(100%) | 0/14(0%) | Plus/Plus | |

Features:
**325775 bp at 5' side: BMP-binding endothelial regulator protein precursor179119 bp at 3' side: neuropeptide S receptor B short**

```
Query  287       TCTCTCCTTTATTG  300
                 ||||||||||||||
Sbjct  34518917  TCTCTCCTTTATTG  34518930
```

Range 15: 36292038 to 36292051

| Score | Expect | Identities | Gaps | Strand | Frame |
|---|---|---|---|---|---|
| 28.2 bits(14) | 256() | 14/14(100%) | 0/14(0%) | Plus/Plus | |

Features:
**endonuclease/exonuclease/phosphatase family domain-contai...**

```
Query  283       GTGTTCTCTCCTTT  296
                 ||||||||||||||
Sbjct  36292038  GTGTTCTCTCCTTT  36292051
```

Range 16: 42733743 to 42733760

| Score | Expect | Identities | Gaps | Strand | Frame |
|---|---|---|---|---|---|
| 28.2 bits(14) | 256() | 17/18(94%) | 0/18(0%) | Plus/Plus | |

Features:
**467284 bp at 5' side: transcriptional activator GLI3219036 bp at 3' side: UPF0415 protein C7orf25 isoform a**

```
Query  280       CTGGTGTTCTCTCCTTTA  297
                 ||||||| ||||||||||
Sbjct  42733743  CTGGTGTCCTCTCCTTTA  42733760
```

Range 17: 54624487 to 54624500

| Score | Expect | Identities | Gaps | Strand | Frame |
|---|---|---|---|---|---|
| 28.2 bits(14) | 256() | 14/14(100%) | 0/14(0%) | Plus/Plus | |

Features:
**V-set and transmembrane domain-containing protein 2A isof...V-set and transmembrane domain-containing protein 2A isof...**

```
Query  287       TCTCTCCTTTATTG  300
                 ||||||||||||||
Sbjct  54624487  TCTCTCCTTTATTG  54624500
```

Range 18: 69155083 to 69155096

| Score | Expect | Identities | Gaps | Strand | Frame |
|---|---|---|---|---|---|
| 28.2 bits(14) | 256() | 14/14(100%) | 0/14(0%) | Plus/Plus | |

Features:
**2358468 bp at 5' side: S-adenosyl-L-methionine-dependent tRNA 4-demethylwyosine ...4693 bp at 3' side: autism susceptibility gene 2 protein isoform 1**

```
Query  281       TGGTGTTCTCTCCT  294
                 ||||||||||||||
Sbjct  69155083  TGGTGTTCTCTCCT  69155096
```

Range 19: 69795035 to 69795048

| Score | Expect | Identities | Gaps | Strand | Frame |
|---|---|---|---|---|---|
| 28.2 bits(14) | 256() | 14/14(100%) | 0/14(0%) | Plus/Plus | |

Features:
**autism susceptibility gene 2 protein isoform 1autism susceptibility gene 2 protein isoform 2**

```
Query  281       TGGTGTTCTCTCCT  294
                 ||||||||||||||
Sbjct  69795035  TGGTGTTCTCTCCT  69795048
```

Range 20: 77418695 to 77418708

| Score | Expect | Identities | Gaps | Strand | Frame |
|---|---|---|---|---|---|
| 28.2 bits(14) | 256() | 14/14(100%) | 0/14(0%) | Plus/Plus | |

Features:
**putative homeodomain transcription factor 2 isoform 4putative homeodomain transcription factor 2 isoform 3**

```
Query  283       GTGTTCTCTCCTTT  296
                 ||||||||||||||
Sbjct  77418695  GTGTTCTCTCCTTT  77418708
```

Range 21: 81725741 to 81725754

| Score | Expect | Identities | Gaps | Strand | Frame |
|---|---|---|---|---|---|
| 28.2 bits(14) | 256() | 14/14(100%) | 0/14(0%) | Plus/Plus | |

Features:
**voltage-dependent calcium channel subunit alpha-2/delta-1...voltage-dependent calcium channel subunit alpha-2/delta-1...**

```
Query  272       GTAACAAGCTGGTG  285
                 ||||||||||||||
Sbjct  81725741  GTAACAAGCTGGTG  81725754
```

Range 22: 83755613 to 83755626

| Score | Expect | Identities | Gaps | Strand | Frame |
|---|---|---|---|---|---|
| 28.2 bits(14) | 256() | 14/14(100%) | 0/14(0%) | Plus/Plus | |

Features:
**1486 bp at 5' side: semaphorin-3A precursor803315 bp at 3' side: semaphorin-3D precursor**

```
Query  286       TTCTCTCCTTTATT  299
                 ||||||||||||||
Sbjct  83755613  TTCTCTCCTTTATT  83755626
```

Range 23: 88863793 to 88863806

| Score | Expect | Identities | Gaps | Strand | Frame |
|---|---|---|---|---|---|
| 28.2 bits(14) | 256() | 14/14(100%) | 0/14(0%) | Plus/Plus | |

Features:
**zinc finger protein 804B**

```
Query  287       TCTCTCCTTTATTG  300
                 ||||||||||||||
Sbjct  88863793  TCTCTCCTTTATTG  88863806
```

Range 24: 100998683 to 100998696

| Score | Expect | Identities | Gaps | Strand | Frame |
|---|---|---|---|---|---|
| 28.2 bits(14) | 256() | 14/14(100%) | 0/14(0%) | Plus/Plus | |

Features:
**collagen alpha-1(XXVI) chain isoform 1 precursorcollagen alpha-1(XXVI) chain isoform 2 precursor**

```
Query  284        TGTTCTCTCCTTTA  297
                  ||||||||||||||
Sbjct  100998683  TGTTCTCTCCTTTA  100998696
```

Range 25: 105762333 to 105762346

| Score | Expect | Identities | Gaps | Strand | Frame |
|---|---|---|---|---|---|
| 28.2 bits(14) | 256() | 14/14(100%) | 0/14(0%) | Plus/Plus | |

Features:
**76054 bp at 5' side: synaptophysin-like protein 1 isoform b62664 bp at 3' side: nicotinamide phosphoribosyltransferase precursor**

```
Query  285        GTTCTCTCCTTTAT  298
                  ||||||||||||||
Sbjct  105762333  GTTCTCTCCTTTAT  105762346
```

Range 26: 108920735 to 108920748

| Score | Expect | Identities | Gaps | Strand | Frame |
|---|---|---|---|---|---|
| 28.2 bits(14) | 256() | 14/14(100%) | 0/14(0%) | Plus/Plus | |

Features:
**462945 bp at 5' side: uncharacterized protein C7orf661316235 bp at 3' side: mitochondrial inner membrane protease subunit 2**

```
Query  286        TTCTCTCCTTTATT  299
                  ||||||||||||||
Sbjct  108920735  TTCTCTCCTTTATT  108920748
```

Range 27: 128186244 to 128186257

| Score | Expect | Identities | Gaps | Strand | Frame |
|---|---|---|---|---|---|
| 28.2 bits(14) | 256() | 14/14(100%) | 0/14(0%) | Plus/Plus | |

Features:
**110730 bp at 5' side: methyltransferase-like protein 2B59440 bp at 3' side: protein FAM71F2 isoform a**

```
Query  286        TTCTCTCCTTTATT  299
                  ||||||||||||||
Sbjct  128186244  TTCTCTCCTTTATT  128186257
```

Range 28: 130000112 to 130000125

| Score | Expect | Identities | Gaps | Strand | Frame |
|---|---|---|---|---|---|
| 28.2 bits(14) | 256() | 14/14(100%) | 0/14(0%) | Plus/Plus | |

Features:
**centrosomal protein of 41 kDa isoform 1centrosomal protein of 41 kDa isoform 3**

```
Query  287        TCTCTCCTTTATTG  300
                  ||||||||||||||
Sbjct  130000112  TCTCTCCTTTATTG  130000125
```

Range 29: 138365846 to 138365859

| Score | Expect | Identities | Gaps | Strand | Frame |
|---|---|---|---|---|---|
| 28.2 bits(14) | 256() | 14/14(100%) | 0/14(0%) | Plus/Plus | |

Features:
**V-type proton ATPase 116 kDa subunit a isoform 4V-type proton ATPase 116 kDa subunit a isoform 4**

```
Query  286        TTCTCTCCTTTATT  299
                  ||||||||||||||
Sbjct  138365846  TTCTCTCCTTTATT  138365859
```

Range 30: 143387739 to 143387752

| Score | Expect | Identities | Gaps | Strand | Frame |
|---|---|---|---|---|---|
| 28.2 bits(14) | 256() | 14/14(100%) | 0/14(0%) | Plus/Plus | |

Features:
**25961 bp at 5' side: protein FAM115C isoform B1741 bp at 3' side: cTAGE family member 6**

```
Query  283        GTGTTCTCTCCTTT  296
                  ||||||||||||||
```

```
Sbjct  143387739  GTGTTCTCTCCTTT  143387752
```

Range 31: 153342633 to 153342646

| Score | Expect | Identities | Gaps | Strand | Frame |
|---|---|---|---|---|---|
| 28.2 bits(14) | 256() | 14/14(100%) | 0/14(0%) | Plus/Plus | |

Features:
**492878 bp at 5' side: small ubiquitin-related modifier 2-like250595 bp at 3' side: dipeptidyl aminopeptidase-like protein 6 isoform 3**

```
Query  286        TTCTCTCCTTTATT  299
                  ||||||||||||||
Sbjct  153342633  TTCTCTCCTTTATT  153342646
```

Range 32: 154444404 to 154444417

| Score | Expect | Identities | Gaps | Strand | Frame |
|---|---|---|---|---|---|
| 28.2 bits(14) | 256() | 14/14(100%) | 0/14(0%) | Plus/Plus | |

Features:
**dipeptidyl aminopeptidase-like protein 6 isoform 3dipeptidyl aminopeptidase-like protein 6 isoform 1**

```
Query  278        AGCTGGTGTTCTCT  291
                  ||||||||||||||
Sbjct  154444404  AGCTGGTGTTCTCT  154444417
```

Range 33: 3564244 to 3564257

| Score | Expect | Identities | Gaps | Strand | Frame |
|---|---|---|---|---|---|
| 28.2 bits(14) | 256() | 14/14(100%) | 0/14(0%) | Plus/Minus | |

Features:
**protein sidekick-1 isoform 1**

```
Query  286      TTCTCTCCTTTATT  299
                ||||||||||||||
Sbjct  3564257  TTCTCTCCTTTATT  3564244
```

Range 34: 12017989 to 12018002

| Score | Expect | Identities | Gaps | Strand | Frame |
|---|---|---|---|---|---|
| 28.2 bits(14) | 256() | 14/14(100%) | 0/14(0%) | Plus/Minus | |

Features:
**145793 bp at 5' side: thrombospondin type-1 domain-containing protein 7A precursor237122 bp at 3' side: transmembrane protein 106B**

```
Query  286       TTCTCTCCTTTATT  299
                 ||||||||||||||
Sbjct  12018002  TTCTCTCCTTTATT  12017989
```

Range 35: 25855055 to 25855068

| Score | Expect | Identities | Gaps | Strand | Frame |
|---|---|---|---|---|---|
| 28.2 bits(14) | 256() | 14/14(100%) | 0/14(0%) | Plus/Minus | |

Features:
**145407 bp at 5' side: uncharacterized protein LOC646588330161 bp at 3' side: nuclear factor erythroid 2-related factor 3**

```
Query  274       AACAAGCTGGTGTT  287
                 ||||||||||||||
Sbjct  25855068  AACAAGCTGGTGTT  25855055
```

Range 36: 31687594 to 31687607

| Score | Expect | Identities | Gaps | Strand | Frame |
|---|---|---|---|---|---|
| 28.2 bits(14) | 256() | 14/14(100%) | 0/14(0%) | Plus/Minus | |

Features:
**coiled-coil domain-containing protein 129 isoform 3coiled-coil domain-containing protein 129 isoform 1**

```
Query  286       TTCTCTCCTTTATT  299
                 ||||||||||||||
Sbjct  31687607  TTCTCTCCTTTATT  31687594
```

Range 37: 42035917 to 42035930

| Score | Expect | Identities | Gaps | Strand | Frame |
|---|---|---|---|---|---|
| 28.2 bits(14) | 256() | 14/14(100%) | 0/14(0%) | Plus/Minus | |

Features:
**transcriptional activator GLI3**

```
Query  284       TGTTCTCTCCTTTA  297
                 ||||||||||||||
Sbjct  42035930  TGTTCTCTCCTTTA  42035917
```

Range 38: 56133918 to 56133931

| Score | Expect | Identities | Gaps | Strand | Frame |
|---|---|---|---|---|---|
| 28.2 bits(14) | 256() | 14/14(100%) | 0/14(0%) | Plus/Minus | |

Features:
**117 bp at 5' side: T-complex protein 1 subunit zeta isoform b1040 bp at 3' side: sulfatase-modifying factor 2 isoform e precursor**

```
Query  283       GTGTTCTCTCCTTT  296
                 ||||||||||||||
Sbjct  56133931  GTGTTCTCTCCTTT  56133918
```

Range 39: 64630077 to 64630090

| Score | Expect | Identities | Gaps | Strand | Frame |
|---|---|---|---|---|---|
| 28.2 bits(14) | 256() | 14/14(100%) | 0/14(0%) | Plus/Minus | |

Features:
**81583 bp at 5' side: endogenous retrovirus group 3 member 1 Env polyprotein pr...303826 bp at 3' side: zinc finger protein 92 isoform 2**

```
Query  283       GTGTTCTCTCCTTT  296
                 ||||||||||||||
Sbjct  64630090  GTGTTCTCTCCTTT  64630077
```

Range 40: 65323751 to 65323764

| Score | Expect | Identities | Gaps | Strand | Frame |
|---|---|---|---|---|---|
| 28.2 bits(14) | 256() | 14/14(100%) | 0/14(0%) | Plus/Minus | |

Features:
**209019 bp at 5' side: kinase suppressor of Ras 1-like isoform X2109793 bp at 3' side: vitamin K epoxide reductase complex subunit 1-like protei...**

```
Query  283       GTGTTCTCTCCTTT  296
                 ||||||||||||||
Sbjct  65323764  GTGTTCTCTCCTTT  65323751
```

Range 41: 90978066 to 90978083

| Score | Expect | Identities | Gaps | Strand | Frame |
|---|---|---|---|---|---|
| 28.2 bits(14) | 256() | 17/18(94%) | 0/18(0%) | Plus/Minus | |

Features:
**152035 bp at 5' side: frizzled-1454865 bp at 3' side: transcription termination factor 1, mitochondrial isoform...**

```
Query  279       GCTGGTGTTCTCTCCTTT  296
                 ||||||||||| ||||||
Sbjct  90978083  GCTGGTGTTCTTTCCTTT  90978066
```

Range 42: 91117319 to 91117332

| Score | Expect | Identities | Gaps | Strand | Frame |
|---|---|---|---|---|---|
| 28.2 bits(14) | 256() | 14/14(100%) | 0/14(0%) | Plus/Minus | |

Features:
**291288 bp at 5' side: frizzled-1315616 bp at 3' side: transcription termination factor 1, mitochondrial isoform...**

```
Query  281       TGGTGTTCTCTCCT  294
                 ||||||||||||||
Sbjct  91117332  TGGTGTTCTCTCCT  91117319
```

Range 43: 92332853 to 92332866

| Score | Expect | Identities | Gaps | Strand | Frame |
|---|---|---|---|---|---|
| 28.2 bits(14) | 256() | 14/14(100%) | 0/14(0%) | Plus/Minus | |

Features:
**cyclin-dependent kinase 6cyclin-dependent kinase 6**

```
Query  285       GTTCTCTCCTTTAT  298
                 ||||||||||||||
Sbjct  92332866  GTTCTCTCCTTTAT  92332853
```

Range 44: 103456094 to 103456111

| Score | Expect | Identities | Gaps | Strand | Frame |
|---|---|---|---|---|---|
| 28.2 bits(14) | 256() | 17/18(94%) | 0/18(0%) | Plus/Minus | |

Features:
**reelin isoform b precursorreelin isoform a precursor**

```
Query  281        TGGTGTTCTCTCCTTTAT  298
                  ||||| ||||||||||||
Sbjct  103456111  TGGTGATCTCTCCTTTAT  103456094
```

Range 45: 109465477 to 109465490

| Score | Expect | Identities | Gaps | Strand | Frame |
|---|---|---|---|---|---|
| 28.2 bits(14) | 256() | 14/14(100%) | 0/14(0%) | Plus/Minus | |

Features:
**1007687 bp at 5' side: uncharacterized protein C7orf66771493 bp at 3' side: mitochondrial inner membrane protease subunit 2**

```
Query  287        TCTCTCCTTTATTG  300
                  ||||||||||||||
Sbjct  109465490  TCTCTCCTTTATTG  109465477
```

Range 46: 109859522 to 109859535

| Score | Expect | Identities | Gaps | Strand | Frame |
|---|---|---|---|---|---|
| 28.2 bits(14) | 256() | 14/14(100%) | 0/14(0%) | Plus/Minus | |

Features:
**1401732 bp at 5' side: uncharacterized protein C7orf66377448 bp at 3' side: mitochondrial inner membrane protease subunit 2**

```
Query  284        TGTTCTCTCCTTTA  297
                  ||||||||||||||
Sbjct  109859535  TGTTCTCTCCTTTA  109859522
```

Range 47: 117625074 to 117625087

| Score | Expect | Identities | Gaps | Strand | Frame |
|---|---|---|---|---|---|
| 28.2 bits(14) | 256() | 14/14(100%) | 0/14(0%) | Plus/Minus | |

Features:
**178043 bp at 5' side: cortactin-binding protein 2132756 bp at 3' side: LSM8 homolog, U6 small nuclear RNA associated**

```
Query  286        TTCTCTCCTTTATT  299
                  ||||||||||||||
```

```
Sbjct  117625087  TTCTCTCCTTTATT  117625074
```

Range 48: 118281883 to 118281896

| Score | Expect | Identities | Gaps | Strand | Frame |
|---|---|---|---|---|---|
| 28.2 bits(14) | 256() | 14/14(100%) | 0/14(0%) | Plus/Minus | |

Features:
**468363 bp at 5' side: ankyrin repeat domain-containing protein 71566113 bp at 3' side: potassium voltage-gated channel subfamily D member 2 prec...**

```
Query  286        TTCTCTCCTTTATT  299
                  ||||||||||||||
Sbjct  118281896  TTCTCTCCTTTATT  118281883
```

Range 49: 123700464 to 123700477

| Score | Expect | Identities | Gaps | Strand | Frame |
|---|---|---|---|---|---|
| 28.2 bits(14) | 256() | 14/14(100%) | 0/14(0%) | Plus/Minus | |

Features:
**94082 bp at 5' side: transmembrane protein 229A619244 bp at 3' side: prosaposin receptor GPR37 precursor**

```
Query  281        TGGTGTTCTCTCCT  294
                  ||||||||||||||
Sbjct  123700477  TGGTGTTCTCTCCT  123700464
```

Range 50: 141196433 to 141196446

| Score | Expect | Identities | Gaps | Strand | Frame |
|---|---|---|---|---|---|
| 28.2 bits(14) | 256() | 14/14(100%) | 0/14(0%) | Plus/Minus | |

Features:
**acylglycerol kinase, mitochondrial precursor**

```
Query  278        AGCTGGTGTTCTCT  291
                  ||||||||||||||
Sbjct  141196446  AGCTGGTGTTCTCT  141196433
```

Range 51: 159096147 to 159096160

| Score | Expect | Identities | Gaps | Strand | Frame |
|---|---|---|---|---|---|
| 28.2 bits(14) | 256() | 14/14(100%) | 0/14(0%) | Plus/Minus | |

Features:
**150369 bp at 5' side: vasoactive intestinal polypeptide receptor 2 isoform 1 pr...**

```
Query  286        TTCTCTCCTTTATT  299
                  ||||||||||||||
Sbjct  159096160  TTCTCTCCTTTATT  159096147
```

Homo sapiens chromosome 8, alternate assembly CHM1_1.1
Sequence ID: **ref|NC_018919.2|** Length: 146399655 Number of Matches: 56
Range 1: 29518684 to 29518700

| Score | Expect | Identities | Gaps | Strand | Frame |
|---|---|---|---|---|---|
| 34.2 bits(17) | 4.1() | 17/17(100%) | 0/17(0%) | Plus/Minus | |

Features:
**113651 bp at 5' side: dual specificity protein phosphatase 4 isoform 2605044 bp at 3' side: store-operated calcium entry-associated regulatory factor...**

```
Query  277       AAGCTGGTGTTCTCTCC  293
                 |||||||||||||||||
Sbjct  29518700  AAGCTGGTGTTCTCTCC  29518684
```

Range 2: 120022443 to 120022458

| Score | Expect | Identities | Gaps | Strand | Frame |
|---|---|---|---|---|---|
| 32.2 bits(16) | 16() | 16/16(100%) | 0/16(0%) | Plus/Minus | |

Features:
**17368 bp at 5' side: tumor necrosis factor receptor superfamily member 11B pre...98079 bp at 3' side: collectin-10 precursor**

```
Query  284        TGTTCTCTCCTTTATT  299
                  ||||||||||||||||
Sbjct  120022458  TGTTCTCTCCTTTATT  120022443
```

Range 3: 22711044 to 22711058

| Score | Expect | Identities | Gaps | Strand | Frame |
|---|---|---|---|---|---|
| 30.2 bits(15) | 65() | 15/15(100%) | 0/15(0%) | Plus/Plus | |

Features:
**bridging integrator 3**

```
Query  282       GGTGTTCTCTCCTTT  296
                 |||||||||||||||
Sbjct  22711044  GGTGTTCTCTCCTTT  22711058
```

Range 4: 31046896 to 31046910

| Score | Expect | Identities | Gaps | Strand | Frame |
|---|---|---|---|---|---|
| 30.2 bits(15) | 65() | 15/15(100%) | 0/15(0%) | Plus/Plus | |

Features:
**138924 bp at 5' side: testis-expressed sequence 15 protein8368 bp at 3' side: purine-rich element-binding protein gamma isoform B**

```
Query  284       TGTTCTCTCCTTTAT  298
                 |||||||||||||||
Sbjct  31046896  TGTTCTCTCCTTTAT  31046910
```

Range 5: 38339361 to 38339375

| Score | Expect | Identities | Gaps | Strand | Frame |
|---|---|---|---|---|---|
| 30.2 bits(15) | 65() | 15/15(100%) | 0/15(0%) | Plus/Plus | |

Features:
**histone-lysine N-methyltransferase NSD3 isoform long**

```
Query  285       GTTCTCTCCTTTATT  299
                 |||||||||||||||
Sbjct  38339361  GTTCTCTCCTTTATT  38339375
```

Range 6: 132678980 to 132678994

| Score | Expect | Identities | Gaps | Strand | Frame |
|---|---|---|---|---|---|
| 30.2 bits(15) | 65() | 15/15(100%) | 0/15(0%) | Plus/Plus | |

Features:
**585364 bp at 5' side: adenylate cyclase type 8277967 bp at 3' side: protein EFR3 homolog A**

```
Query  284        TGTTCTCTCCTTTAT  298
                  |||||||||||||||
Sbjct  132678980  TGTTCTCTCCTTTAT  132678994
```

Range 7: 17577964 to 17577978

| Score | Expect | Identities | Gaps | Strand | Frame |
|---|---|---|---|---|---|
| 30.2 bits(15) | 65() | 15/15(100%) | 0/15(0%) | Plus/Minus | |

Features:
**105701 bp at 5' side: myotubularin-related protein 719837 bp at 3' side: cationic amino acid transporter 2 isoform 3**

```
Query  278       AGCTGGTGTTCTCTC  292
                 |||||||||||||||
```

```
Sbjct  17577978  AGCTGGTGTTCTCTC  17577964
```

Range 8: 81676046 to 81676064

| Score | Expect | Identities | Gaps | Strand | Frame |
|---|---|---|---|---|---|
| 30.2 bits(15) | 65() | 18/19(95%) | 0/19(0%) | Plus/Minus | |

Features:
**zinc finger protein 704**

```
Query  280       CTGGTGTTCTCTCCTTTAT  298
                 |||||||||| ||||||||
Sbjct  81676064  CTGGTGTTCTGTCCTTTAT  81676046
```

Range 9: 91414642 to 91414660

| Score | Expect | Identities | Gaps | Strand | Frame |
|---|---|---|---|---|---|
| 30.2 bits(15) | 65() | 18/19(95%) | 0/19(0%) | Plus/Minus | |

Features:
**278788 bp at 5' side: calbindin263824 bp at 3' side: transmembrane protein 64 isoform 2**

```
Query  273       TAACAAGCTGGTGTTCTCT  291
                 ||||||||| |||||||||
Sbjct  91414660  TAACAAGCTTGTGTTCTCT  91414642
```

Range 10: 122649963 to 122649977

| Score | Expect | Identities | Gaps | Strand | Frame |
|---|---|---|---|---|---|
| 30.2 bits(15) | 65() | 15/15(100%) | 0/15(0%) | Plus/Minus | |

Features:
**785502 bp at 5' side: beta-1-syntrophin16555 bp at 3' side: hyaluronan synthase 2**

```
Query  278        AGCTGGTGTTCTCTC  292
                  |||||||||||||||
Sbjct  122649977  AGCTGGTGTTCTCTC  122649963
```

Range 11: 1092825 to 1092838

| Score | Expect | Identities | Gaps | Strand | Frame |
|---|---|---|---|---|---|
| 28.2 bits(14) | 256() | 14/14(100%) | 0/14(0%) | Plus/Plus | |

Features:
**260956 bp at 5' side: uncharacterized protein LOC105377777 isoform X2403721 bp at 3' side: disks large-associated protein 2 isoform 1**

```
Query  285      GTTCTCTCCTTTAT  298
                ||||||||||||||
Sbjct  1092825  GTTCTCTCCTTTAT  1092838
```

Range 12: 10975434 to 10975451

| Score | Expect | Identities | Gaps | Strand | Frame |
|---|---|---|---|---|---|
| 28.2 bits(14) | 256() | 17/18(94%) | 0/18(0%) | Plus/Plus | |

Features:
**XK-related protein 6**

```
Query  277       AAGCTGGTGTTCTCTCCT  294
                 |||||||||| |||||||
Sbjct  10975434  AAGCTGGTGTCCTCTCCT  10975451
```

Range 13: 18351752 to 18351765

| Score | Expect | Identities | Gaps | Strand | Frame |
|---|---|---|---|---|---|
| 28.2 bits(14) | 256() | 14/14(100%) | 0/14(0%) | Plus/Plus | |

Features:
**69870 bp at 5' side: arylamine N-acetyltransferase 1 isoform a107214 bp at 3' side: arylamine N-acetyltransferase 2**

```
Query  286       TTCTCTCCTTTATT  299
                 ||||||||||||||
Sbjct  18351752  TTCTCTCCTTTATT  18351765
```

Range 14: 24771877 to 24771890

| Score | Expect | Identities | Gaps | Strand | Frame |
|---|---|---|---|---|---|
| 28.2 bits(14) | 256() | 14/14(100%) | 0/14(0%) | Plus/Plus | |

Features:
**203668 bp at 5' side: disintegrin and metalloproteinase domain-containing prote...201725 bp at 3' side: neurofilament medium polypeptide isoform 1**

```
Query  286       TTCTCTCCTTTATT  299
                 ||||||||||||||
Sbjct  24771877  TTCTCTCCTTTATT  24771890
```

Range 15: 24826605 to 24826618

| Score | Expect | Identities | Gaps | Strand | Frame |
|---|---|---|---|---|---|
| 28.2 bits(14) | 256() | 14/14(100%) | 0/14(0%) | Plus/Plus | |

Features:
**258396 bp at 5' side: disintegrin and metalloproteinase domain-containing prote...146997 bp at 3' side: neurofilament medium polypeptide isoform 1**

```
Query  286       TTCTCTCCTTTATT  299
                 ||||||||||||||
Sbjct  24826605  TTCTCTCCTTTATT  24826618
```

Range 16: 28210048 to 28210061

| Score | Expect | Identities | Gaps | Strand | Frame |
|---|---|---|---|---|---|
| 28.2 bits(14) | 256() | 14/14(100%) | 0/14(0%) | Plus/Plus | |

Features:
**elongator complex protein 3 isoform 1elongator complex protein 3 isoform 3**

```
Query  286       TTCTCTCCTTTATT  299
                 ||||||||||||||
Sbjct  28210048  TTCTCTCCTTTATT  28210061
```

Range 17: 31594505 to 31594518

| Score | Expect | Identities | Gaps | Strand | Frame |
|---|---|---|---|---|---|
| 28.2 bits(14) | 256() | 14/14(100%) | 0/14(0%) | Plus/Plus | |

Features:
**362427 bp at 5' side: Werner syndrome ATP-dependent helicase103849 bp at 3' side: pro-neuregulin-1, membrane-bound isoform isoform HRG-beta1d**

```
Query  286       TTCTCTCCTTTATT  299
                 ||||||||||||||
Sbjct  31594505  TTCTCTCCTTTATT  31594518
```

Range 18: 54981640 to 54981653

| Score | Expect | Identities | Gaps | Strand | Frame |
|---|---|---|---|---|---|
| 28.2 bits(14) | 256() | 14/14(100%) | 0/14(0%) | Plus/Plus | |

Features:
**transcription elongation factor A protein 1 isoform 2transcription elongation factor A protein 1 isoform 1**

```
Query  286       TTCTCTCCTTTATT  299
                 ||||||||||||||
Sbjct  54981640  TTCTCTCCTTTATT  54981653
```

Range 19: 57546866 to 57546879

| Score | Expect | Identities | Gaps | Strand | Frame |
|---|---|---|---|---|---|
| 28.2 bits(14) | 256() | 14/14(100%) | 0/14(0%) | Plus/Plus | |

Features:
**136477 bp at 5' side: proenkephalin-A preproprotein381229 bp at 3' side: inositol monophosphatase 3**

```
Query  278       AGCTGGTGTTCTCT  291
                 ||||||||||||||
Sbjct  57546866  AGCTGGTGTTCTCT  57546879
```

Range 20: 59059956 to 59059969

| Score | Expect | Identities | Gaps | Strand | Frame |
|---|---|---|---|---|---|
| 28.2 bits(14) | 256() | 14/14(100%) | 0/14(0%) | Plus/Plus | |

Features:
**1102056 bp at 5' side: inositol monophosphatase 351083 bp at 3' side: protein FAM110B**

```
Query  286       TTCTCTCCTTTATT  299
                 ||||||||||||||
Sbjct  59059956  TTCTCTCCTTTATT  59059969
```

Range 21: 59557373 to 59557386

| Score | Expect | Identities | Gaps | Strand | Frame |
|---|---|---|---|---|---|
| 28.2 bits(14) | 256() | 14/14(100%) | 0/14(0%) | Plus/Plus | |

Features:
**protein FAN isoform 1protein FAN isoform 2**

```
Query  287       TCTCTCCTTTATTG  300
                 ||||||||||||||
Sbjct  59557373  TCTCTCCTTTATTG  59557386
```

Range 22: 74514638 to 74514651

| Score | Expect | Identities | Gaps | Strand | Frame |
|---|---|---|---|---|---|
| 28.2 bits(14) | 256() | 14/14(100%) | 0/14(0%) | Plus/Plus | |

Features:
**double-stranded RNA-binding protein Staufen homolog 2 iso...double-stranded RNA-binding protein Staufen homolog 2 iso...**

```
Query  271       TGTAACAAGCTGGT  284
                 ||||||||||||||
Sbjct  74514638  TGTAACAAGCTGGT  74514651
```

Range 23: 85574744 to 85574757

| Score | Expect | Identities | Gaps | Strand | Frame |
|---|---|---|---|---|---|
| 28.2 bits(14) | 256() | 14/14(100%) | 0/14(0%) | Plus/Plus | |

Features:
**RNA-binding Raly-like protein isoform 1RNA-binding Raly-like protein isoform 2**

```
Query  284       TGTTCTCTCCTTTA  297
                 ||||||||||||||
Sbjct  85574744  TGTTCTCTCCTTTA  85574757
```

Range 24: 95875441 to 95875454

| Score | Expect | Identities | Gaps | Strand | Frame |
|---|---|---|---|---|---|
| 28.2 bits(14) | 256() | 14/14(100%) | 0/14(0%) | Plus/Plus | |

Features:
**32973 bp at 5' side: probable C-mannosyltransferase DPY19L4529 bp at 3' side: integrator complex subunit 8**

```
Query  277       AAGCTGGTGTTCTC  290
                 ||||||||||||||
```

```
Sbjct  95875441  AAGCTGGTGTTCTC  95875454
```

Range 25: 109711943 to 109711956

| Score | Expect | Identities | Gaps | Strand | Frame |
|---|---|---|---|---|---|
| 28.2 bits(14) | 256() | 14/14(100%) | 0/14(0%) | Plus/Plus | |

Features:
**172858 bp at 5' side: ER membrane protein complex subunit 2124366 bp at 3' side: transmembrane protein 74**

```
Query  277        AAGCTGGTGTTCTC  290
                  ||||||||||||||
Sbjct  109711943  AAGCTGGTGTTCTC  109711956
```

Range 26: 114211770 to 114211783

| Score | Expect | Identities | Gaps | Strand | Frame |
|---|---|---|---|---|---|
| 28.2 bits(14) | 256() | 14/14(100%) | 0/14(0%) | Plus/Plus | |

Features:
**CUB and sushi domain-containing protein 3 isoform 3CUB and sushi domain-containing protein 3 isoform 1**

```
Query  279        GCTGGTGTTCTCTC  292
                  ||||||||||||||
Sbjct  114211770  GCTGGTGTTCTCTC  114211783
```

Range 27: 135655842 to 135655855

| Score | Expect | Identities | Gaps | Strand | Frame |
|---|---|---|---|---|---|
| 28.2 bits(14) | 256() | 14/14(100%) | 0/14(0%) | Plus/Plus | |

Features:
**zinc finger protein ZFAT isoform 2zinc finger protein ZFAT isoform 2**

```
Query  275        ACAAGCTGGTGTTC  288
                  ||||||||||||||
Sbjct  135655842  ACAAGCTGGTGTTC  135655855
```

Range 28: 145188922 to 145188935

| Score | Expect | Identities | Gaps | Strand | Frame |
|---|---|---|---|---|---|
| 28.2 bits(14) | 256() | 14/14(100%) | 0/14(0%) | Plus/Plus | |

Features:
**7642 bp at 5' side: glycosylphosphatidylinositol anchor attachment 1 protein1303 bp at 3' side: cytochrome c1, heme protein, mitochondrial precursor**

```
Query  286        TTCTCTCCTTTATT  299
                  ||||||||||||||
Sbjct  145188922  TTCTCTCCTTTATT  145188935
```

Range 29: 1529873 to 1529886

| Score | Expect | Identities | Gaps | Strand | Frame |
|---|---|---|---|---|---|
| 28.2 bits(14) | 256() | 14/14(100%) | 0/14(0%) | Plus/Minus | |

Features:
**disks large-associated protein 2 isoform 1disks large-associated protein 2 isoform 2**

```
Query  278      AGCTGGTGTTCTCT  291
                ||||||||||||||
Sbjct  1529886  AGCTGGTGTTCTCT  1529873
```

Range 30: 2514166 to 2514179

| Score | Expect | Identities | Gaps | Strand | Frame |
|---|---|---|---|---|---|
| 28.2 bits(14) | 256() | 14/14(100%) | 0/14(0%) | Plus/Minus | |

Features:
**421311 bp at 5' side: myomesin-2370513 bp at 3' side: CUB and sushi domain-containing protein 1 precursor**

```
Query  271      TGTAACAAGCTGGT  284
                ||||||||||||||
Sbjct  2514179  TGTAACAAGCTGGT  2514166
```

Range 31: 2872470 to 2872483

| Score | Expect | Identities | Gaps | Strand | Frame |
|---|---|---|---|---|---|
| 28.2 bits(14) | 256() | 14/14(100%) | 0/14(0%) | Plus/Minus | |

Features:
**779615 bp at 5' side: myomesin-212209 bp at 3' side: CUB and sushi domain-containing protein 1 precursor**

```
Query  278      AGCTGGTGTTCTCT  291
                ||||||||||||||
Sbjct  2872483  AGCTGGTGTTCTCT  2872470
```

Range 32: 4300493 to 4300506

| Score | Expect | Identities | Gaps | Strand | Frame |
|---|---|---|---|---|---|
| 28.2 bits(14) | 256() | 14/14(100%) | 0/14(0%) | Plus/Minus | |

Features:
**CUB and sushi domain-containing protein 1 precursor**

```
Query  281      TGGTGTTCTCTCCT  294
                ||||||||||||||
Sbjct  4300506  TGGTGTTCTCTCCT  4300493
```

Range 33: 13157489 to 13157502

| Score | Expect | Identities | Gaps | Strand | Frame |
|---|---|---|---|---|---|
| 28.2 bits(14) | 256() | 14/14(100%) | 0/14(0%) | Plus/Minus | |

Features:
**rho GTPase-activating protein 7 isoform 1rho GTPase-activating protein 7 isoform 2**

```
Query  286       TTCTCTCCTTTATT  299
                 ||||||||||||||
Sbjct  13157502  TTCTCTCCTTTATT  13157489
```

Range 34: 13582991 to 13583004

| Score | Expect | Identities | Gaps | Strand | Frame |
|---|---|---|---|---|---|
| 28.2 bits(14) | 256() | 14/14(100%) | 0/14(0%) | Plus/Minus | |

Features:
**23688 bp at 5' side: rho GTPase-activating protein 7 isoform 343236 bp at 3' side: uncharacterized protein C8orf48**

```
Query  286       TTCTCTCCTTTATT  299
                 ||||||||||||||
Sbjct  13583004  TTCTCTCCTTTATT  13582991
```

Range 35: 15408156 to 15408169

| Score | Expect | Identities | Gaps | Strand | Frame |
|---|---|---|---|---|---|
| 28.2 bits(14) | 256() | 14/14(100%) | 0/14(0%) | Plus/Minus | |

Features:
**111574 bp at 5' side: zeta-sarcoglycan191233 bp at 3' side: tumor suppressor candidate 3 isoform b precursor**

```
Query  283       GTGTTCTCTCCTTT  296
                 ||||||||||||||
Sbjct  15408169  GTGTTCTCTCCTTT  15408156
```

Range 36: 21359387 to 21359400

| Score | Expect | Identities | Gaps | Strand | Frame |
|---|---|---|---|---|---|
| 28.2 bits(14) | 256() | 14/14(100%) | 0/14(0%) | Plus/Minus | |

Features:
**3227 bp at 5' side: uncharacterized protein LOC105379399393077 bp at 3' side: GDNF family receptor alpha-2 isoform c precursor**

```
Query  286       TTCTCTCCTTTATT  299
                 ||||||||||||||
Sbjct  21359400  TTCTCTCCTTTATT  21359387
```

Range 37: 36097240 to 36097253

| Score | Expect | Identities | Gaps | Strand | Frame |
|---|---|---|---|---|---|
| 28.2 bits(14) | 256() | 14/14(100%) | 0/14(0%) | Plus/Minus | |

Features:
**247581 bp at 5' side: netrin receptor UNC5D precursor746912 bp at 3' side: potassium channel subfamily U member 1**

```
Query  286       TTCTCTCCTTTATT  299
                 ||||||||||||||
Sbjct  36097253  TTCTCTCCTTTATT  36097240
```

Range 38: 40704526 to 40704539

| Score | Expect | Identities | Gaps | Strand | Frame |
|---|---|---|---|---|---|
| 28.2 bits(14) | 256() | 14/14(100%) | 0/14(0%) | Plus/Minus | |

Features:
**zinc finger matrin-type protein 4 isoform bzinc finger matrin-type protein 4 isoform a**

```
Query  286       TTCTCTCCTTTATT  299
                 ||||||||||||||
Sbjct  40704539  TTCTCTCCTTTATT  40704526
```

Range 39: 41705219 to 41705232

| Score | Expect | Identities | Gaps | Strand | Frame |
|---|---|---|---|---|---|
| 28.2 bits(14) | 256() | 14/14(100%) | 0/14(0%) | Plus/Minus | |

Features:
**ankyrin-1 isoform 9**

```
Query  279       GCTGGTGTTCTCTC  292
                 ||||||||||||||
Sbjct  41705232  GCTGGTGTTCTCTC  41705219
```

Range 40: 41836499 to 41836512

| Score | Expect | Identities | Gaps | Strand | Frame |
|---|---|---|---|---|---|
| 28.2 bits(14) | 256() | 14/14(100%) | 0/14(0%) | Plus/Minus | |

Features:
**265786 bp at 5' side: ankyrin-1 isoform 5 precursor1643 bp at 3' side: histone acetyltransferase KAT6A**

```
Query  286       TTCTCTCCTTTATT  299
                 ||||||||||||||
Sbjct  41836512  TTCTCTCCTTTATT  41836499
```

Range 41: 48489986 to 48490003

| Score | Expect | Identities | Gaps | Strand | Frame |
|---|---|---|---|---|---|
| 28.2 bits(14) | 256() | 17/18(94%) | 0/18(0%) | Plus/Minus | |

Features:
**DNA repair-scaffolding protein isoform 1DNA repair-scaffolding protein isoform 3**

```
Query  275       ACAAGCTGGTGTTCTCTC  292
                 |||| |||||||||||||
```

```
Sbjct  48490003  ACAACCTGGTGTTCTCTC  48489986
```

Range 42: 49838387 to 49838400

| Score | Expect | Identities | Gaps | Strand | Frame |
|---|---|---|---|---|---|
| 28.2 bits(14) | 256() | 14/14(100%) | 0/14(0%) | Plus/Minus | |

Features:
**138317 bp at 5' side: EF-hand calcium-binding domain-containing protein 1 isofo...45280 bp at 3' side: zinc finger protein SNAI2**

```
Query  286       TTCTCTCCTTTATT  299
                 ||||||||||||||
Sbjct  49838400  TTCTCTCCTTTATT  49838387
```

Range 43: 57586415 to 57586428

| Score | Expect | Identities | Gaps | Strand | Frame |
|---|---|---|---|---|---|
| 28.2 bits(14) | 256() | 14/14(100%) | 0/14(0%) | Plus/Minus | |

Features:
**176026 bp at 5' side: proenkephalin-A preproprotein341680 bp at 3' side: inositol monophosphatase 3**

```
Query  283       GTGTTCTCTCCTTT  296
                 ||||||||||||||
Sbjct  57586428  GTGTTCTCTCCTTT  57586415
```

Range 44: 58075778 to 58075791

| Score | Expect | Identities | Gaps | Strand | Frame |
|---|---|---|---|---|---|
| 28.2 bits(14) | 256() | 14/14(100%) | 0/14(0%) | Plus/Minus | |

Features:
**117878 bp at 5' side: inositol monophosphatase 31035261 bp at 3' side: protein FAM110B**

```
Query  279       GCTGGTGTTCTCTC  292
                 ||||||||||||||
Sbjct  58075791  GCTGGTGTTCTCTC  58075778
```

Range 45: 59025885 to 59025898

| Score | Expect | Identities | Gaps | Strand | Frame |
|---|---|---|---|---|---|
| 28.2 bits(14) | 256() | 14/14(100%) | 0/14(0%) | Plus/Minus | |

Features:
**1067985 bp at 5' side: inositol monophosphatase 385154 bp at 3' side: protein FAM110B**

```
Query  278       AGCTGGTGTTCTCT  291
                 ||||||||||||||
Sbjct  59025898  AGCTGGTGTTCTCT  59025885
```

Range 46: 66941977 to 66941990

| Score | Expect | Identities | Gaps | Strand | Frame |
|---|---|---|---|---|---|
| 28.2 bits(14) | 256() | 14/14(100%) | 0/14(0%) | Plus/Minus | |

Features:
**133629 bp at 5' side: high affinity cAMP-specific 3',5'-cyclic phosphodiesteras...76450 bp at 3' side: dnaJ homolog subfamily C member 5B**

```
Query  281       TGGTGTTCTCTCCT  294
                 ||||||||||||||
Sbjct  66941990  TGGTGTTCTCTCCT  66941977
```

Range 47: 72238412 to 72238425

| Score | Expect | Identities | Gaps | Strand | Frame |
|---|---|---|---|---|---|
| 28.2 bits(14) | 256() | 14/14(100%) | 0/14(0%) | Plus/Minus | |

Features:
**eyes absent homolog 1 isoform 3eyes absent homolog 1 isoform 1**

```
Query  286       TTCTCTCCTTTATT  299
                 ||||||||||||||
Sbjct  72238425  TTCTCTCCTTTATT  72238412
```

Range 48: 75466341 to 75466354

| Score | Expect | Identities | Gaps | Strand | Frame |
|---|---|---|---|---|---|
| 28.2 bits(14) | 256() | 14/14(100%) | 0/14(0%) | Plus/Minus | |

Features:
**137754 bp at 5' side: ganglioside-induced differentiation-associated protein 1 ...322749 bp at 3' side: peptidase inhibitor 15 preproprotein**

```
Query  276       CAAGCTGGTGTTCT  289
                 ||||||||||||||
Sbjct  75466354  CAAGCTGGTGTTCT  75466341
```

Range 49: 76729496 to 76729513

| Score | Expect | Identities | Gaps | Strand | Frame |
|---|---|---|---|---|---|
| 28.2 bits(14) | 256() | 17/18(94%) | 0/18(0%) | Plus/Minus | |

Features:
**201663 bp at 5' side: hepatocyte nuclear factor 4-gamma938419 bp at 3' side: zinc finger homeobox protein 4**

```
Query  281       TGGTGTTCTCTCCTTTAT  298
                 ||||| ||||||||||||
Sbjct  76729513  TGGTGATCTCTCCTTTAT  76729496
```

Range 50: 94375758 to 94375775

| Score | Expect | Identities | Gaps | Strand | Frame |
|---|---|---|---|---|---|
| 28.2 bits(14) | 256() | 17/18(94%) | 0/18(0%) | Plus/Minus | |

Features:
**156512 bp at 5' side: uncharacterized protein C8orf87377388 bp at 3' side: protein FAM92A1 isoform 2**

```
Query  283       GTGTTCTCTCCTTTATTG  300
                 |||||| |||||||||||
Sbjct  94375775  GTGTTCACTCCTTTATTG  94375758
```

Range 51: 100492627 to 100492640

| Score | Expect | Identities | Gaps | Strand | Frame |
|---|---|---|---|---|---|
| 28.2 bits(14) | 256() | 14/14(100%) | 0/14(0%) | Plus/Minus | |

Features:
**vacuolar protein sorting-associated protein 13B isoform 1vacuolar protein sorting-associated protein 13B isoform 5**

```
Query  286        TTCTCTCCTTTATT  299
                  ||||||||||||||
Sbjct  100492640  TTCTCTCCTTTATT  100492627
```

Range 52: 109753100 to 109753113

| Score | Expect | Identities | Gaps | Strand | Frame |
|---|---|---|---|---|---|
| 28.2 bits(14) | 256() | 14/14(100%) | 0/14(0%) | Plus/Minus | |

Features:
**214015 bp at 5' side: ER membrane protein complex subunit 283209 bp at 3' side: transmembrane protein 74**

```
Query  286        TTCTCTCCTTTATT  299
                  ||||||||||||||
Sbjct  109753113  TTCTCTCCTTTATT  109753100
```

Range 53: 112764549 to 112764562

| Score | Expect | Identities | Gaps | Strand | Frame |
|---|---|---|---|---|---|
| 28.2 bits(14) | 256() | 14/14(100%) | 0/14(0%) | Plus/Minus | |

Features:
**1737667 bp at 5' side: potassium voltage-gated channel subfamily V member 1512650 bp at 3' side: CUB and sushi domain-containing protein 3 isoform 3**

```
Query  286        TTCTCTCCTTTATT  299
                  ||||||||||||||
Sbjct  112764562  TTCTCTCCTTTATT  112764549
```

Range 54: 124578229 to 124578242

| Score | Expect | Identities | Gaps | Strand | Frame |
|---|---|---|---|---|---|
| 28.2 bits(14) | 256() | 14/14(100%) | 0/14(0%) | Plus/Minus | |

Features:
**F-box only protein 32 isoform 1F-box only protein 32 isoform 3**

```
Query  283        GTGTTCTCTCCTTT  296
                  ||||||||||||||
Sbjct  124578242  GTGTTCTCTCCTTT  124578229
```

Range 55: 135176113 to 135176126

| Score | Expect | Identities | Gaps | Strand | Frame |
|---|---|---|---|---|---|
| 28.2 bits(14) | 256() | 14/14(100%) | 0/14(0%) | Plus/Minus | |

Features:
**646325 bp at 5' side: CMP-N-acetylneuraminate-beta-galactosamide-alpha-2,3-sial...356716 bp at 3' side: zinc finger protein ZFAT isoform 2**

```
Query  283        GTGTTCTCTCCTTT  296
                  ||||||||||||||
Sbjct  135176126  GTGTTCTCTCCTTT  135176113
```

Range 56: 138719601 to 138719614

| Score | Expect | Identities | Gaps | Strand | Frame |
|---|---|---|---|---|---|
| 28.2 bits(14) | 256() | 14/14(100%) | 0/14(0%) | Plus/Minus | |

Features:
**2018524 bp at 5' side: KH domain-containing, RNA-binding, signal transduction-as...465456 bp at 3' side: protein FAM135B**

```
Query  286        TTCTCTCCTTTATT  299
                  ||||||||||||||
Sbjct  138719614  TTCTCTCCTTTATT  138719601
```

Homo sapiens chromosome 2, GRCh38.p2 Primary Assembly
Sequence ID: **ref|NC_000002.12|** Length: 242193529 Number of Matches: 74
Range 1: 194129972 to 194129988

| Score | Expect | Identities | Gaps | Strand | Frame |
|---|---|---|---|---|---|
| 34.2 bits(17) | 4.1() | 17/17(100%) | 0/17(0%) | Plus/Minus | |

Features:
**1935448 bp at 5' side: tomoregulin-2 isoform X21550055 bp at 3' side: zinc transporter ZIP10 isoform X1**

```
Query  273        TAACAAGCTGGTGTTCT  289
                  |||||||||||||||||
Sbjct  194129988  TAACAAGCTGGTGTTCT  194129972
```

Range 2: 5412256 to 5412271

| Score | Expect | Identities | Gaps | Strand | Frame |
|---|---|---|---|---|---|
| 32.2 bits(16) | 16() | 16/16(100%) | 0/16(0%) | Plus/Minus | |

Features:
**1303049 bp at 5' side: uncharacterized protein LOC105373397280451 bp at 3' side: transcription factor SOX-11**

```
Query  275      ACAAGCTGGTGTTCTC  290
                ||||||||||||||||
Sbjct  5412271  ACAAGCTGGTGTTCTC  5412256
```

Range 3: 19966083 to 19966098

| Score | Expect | Identities | Gaps | Strand | Frame |
|---|---|---|---|---|---|
| 32.2 bits(16) | 16() | 16/16(100%) | 0/16(0%) | Plus/Minus | |

Features:
**WD repeat-containing protein 35 isoform 1WD repeat-containing protein 35 isoform 2**

```
Query  279       GCTGGTGTTCTCTCCT  294
                 ||||||||||||||||
Sbjct  19966098  GCTGGTGTTCTCTCCT  19966083
```

Range 4: 17027032 to 17027046

| Score | Expect | Identities | Gaps | Strand | Frame |
|---|---|---|---|---|---|
| 30.2 bits(15) | 65() | 15/15(100%) | 0/15(0%) | Plus/Plus | |

Features:
**438913 bp at 5' side: protein FAM49A isoform X1483758 bp at 3' side: RAD51-associated protein 2 isoform X2**

```
Query  284       TGTTCTCTCCTTTAT  298
                 |||||||||||||||
Sbjct  17027032  TGTTCTCTCCTTTAT  17027046
```

Range 5: 48661130 to 48661144

| Score | Expect | Identities | Gaps | Strand | Frame |
|---|---|---|---|---|---|
| 30.2 bits(15) | 65() | 15/15(100%) | 0/15(0%) | Plus/Plus | |

Features:
**STON1-GTF2A1L protein isoform 2STON1-GTF2A1L protein isoform 1**

```
Query  284       TGTTCTCTCCTTTAT  298
                 |||||||||||||||
Sbjct  48661130  TGTTCTCTCCTTTAT  48661144
```

Range 6: 70460507 to 70460525

| Score | Expect | Identities | Gaps | Strand | Frame |
|---|---|---|---|---|---|
| 30.2 bits(15) | 65() | 18/19(95%) | 0/19(0%) | Plus/Plus | |

Features:
**protransforming growth factor alpha isoform 2 preproproteinprotransforming growth factor alpha isoform 1 preproprotein**

```
Query  278       AGCTGGTGTTCTCTCCTTT  296
                 ||||| |||||||||||||
Sbjct  70460507  AGCTGCTGTTCTCTCCTTT  70460525
```

Range 7: 222013416 to 222013430

| Score | Expect | Identities | Gaps | Strand | Frame |
|---|---|---|---|---|---|
| 30.2 bits(15) | 65() | 15/15(100%) | 0/15(0%) | Plus/Plus | |

Features:
**441168 bp at 5' side: ephrin type-A receptor 4 isoform X1187744 bp at 3' side: paired box protein Pax-3 isoform X1**

```
Query  280        CTGGTGTTCTCTCCT  294
                  |||||||||||||||
Sbjct  222013416  CTGGTGTTCTCTCCT  222013430
```

Range 8: 85263810 to 85263824

| Score | Expect | Identities | Gaps | Strand | Frame |
|---|---|---|---|---|---|
| 30.2 bits(15) | 65() | 15/15(100%) | 0/15(0%) | Plus/Minus | |

Features:
**transcription factor 7-like 1 isoform X1transcription factor 7-like 1**

```
Query  277       AAGCTGGTGTTCTCT  291
                 |||||||||||||||
Sbjct  85263824  AAGCTGGTGTTCTCT  85263810
```

Range 9: 127773179 to 127773193

| Score | Expect | Identities | Gaps | Strand | Frame |
|---|---|---|---|---|---|
| 30.2 bits(15) | 65() | 15/15(100%) | 0/15(0%) | Plus/Minus | |

Features:
**2198 bp at 5' side: pre-mRNA 3' end processing protein WDR33 isoform 274914 bp at 3' side: DNA-directed RNA polymerase II subunit RPB4**

```
Query  286        TTCTCTCCTTTATTG  300
                  |||||||||||||||
Sbjct  127773193  TTCTCTCCTTTATTG  127773179
```

Range 10: 140869619 to 140869633

| Score | Expect | Identities | Gaps | Strand | Frame |
|---|---|---|---|---|---|
| 30.2 bits(15) | 65() | 15/15(100%) | 0/15(0%) | Plus/Minus | |

Features:
**low-density lipoprotein receptor-related protein 1B precu...low-density lipoprotein receptor-related protein 1B isofo...**

```
Query  286        TTCTCTCCTTTATTG  300
                  |||||||||||||||
Sbjct  140869633  TTCTCTCCTTTATTG  140869619
```

Range 11: 150173346 to 150173360

| Score | Expect | Identities | Gaps | Strand | Frame |
|---|---|---|---|---|---|
| 30.2 bits(15) | 65() | 15/15(100%) | 0/15(0%) | Plus/Minus | |

Features:
**586249 bp at 5' side: methylmalonic aciduria and homocystinuria type D protein,...296627 bp at 3' side: rho-related GTP-binding protein RhoE precursor**

```
Query  286        TTCTCTCCTTTATTG  300
                  |||||||||||||||
Sbjct  150173360  TTCTCTCCTTTATTG  150173346
```

Range 12: 5354701 to 5354714

| Score | Expect | Identities | Gaps | Strand | Frame |
|---|---|---|---|---|---|
| 28.2 bits(14) | 256() | 14/14(100%) | 0/14(0%) | Plus/Plus | |

Features:
**1245494 bp at 5' side: uncharacterized protein LOC105373397338008 bp at 3' side: transcription factor SOX-11**

```
Query  286      TTCTCTCCTTTATT  299
                ||||||||||||||
Sbjct  5354701  TTCTCTCCTTTATT  5354714
```

Range 13: 22390786 to 22390799

| Score | Expect | Identities | Gaps | Strand | Frame |
|---|---|---|---|---|---|
| 28.2 bits(14) | 256() | 14/14(100%) | 0/14(0%) | Plus/Plus | |

Features:
**1247514 bp at 5' side: tudor domain-containing protein 15992028 bp at 3' side: transcription initiation factor TFIID subunit 4-like**

```
Query  283       GTGTTCTCTCCTTT  296
                 ||||||||||||||
Sbjct  22390786  GTGTTCTCTCCTTT  22390799
```

Range 14: 29574409 to 29574426

| Score | Expect | Identities | Gaps | Strand | Frame |
|---|---|---|---|---|---|
| 28.2 bits(14) | 256() | 17/18(94%) | 0/18(0%) | Plus/Plus | |

Features:
**ALK tyrosine kinase receptor precursor**

```
Query  279       GCTGGTGTTCTCTCCTTT  296
                 ||||||||||||| ||||
Sbjct  29574409  GCTGGTGTTCTCTGCTTT  29574426
```

Range 15: 40464960 to 40464973

| Score | Expect | Identities | Gaps | Strand | Frame |
|---|---|---|---|---|---|
| 28.2 bits(14) | 256() | 14/14(100%) | 0/14(0%) | Plus/Plus | |

Features:
**34680 bp at 5' side: sodium/calcium exchanger 1 isoform X111473630 bp at 3' side: uncharacterized protein C2orf91**

```
Query  282       GGTGTTCTCTCCTT  295
                 ||||||||||||||
Sbjct  40464960  GGTGTTCTCTCCTT  40464973
```

Range 16: 69929784 to 69929801

| Score | Expect | Identities | Gaps | Strand | Frame |
|---|---|---|---|---|---|
| 28.2 bits(14) | 256() | 17/18(94%) | 0/18(0%) | Plus/Plus | |

Features:
**max dimerization protein 1 isoform 1max dimerization protein 1 isoform 2**

```
Query  281       TGGTGTTCTCTCCTTTAT  298
                 ||||||| ||||||||||
Sbjct  69929784  TGGTGTTATCTCCTTTAT  69929801
```

Range 17: 77324157 to 77324170

| Score | Expect | Identities | Gaps | Strand | Frame |
|---|---|---|---|---|---|
| 28.2 bits(14) | 256() | 14/14(100%) | 0/14(0%) | Plus/Plus | |

Features:
**leucine-rich repeat transmembrane neuronal protein 4 isof...leucine-rich repeat transmembrane neuronal protein 4 isof...**

```
Query  280       CTGGTGTTCTCTCC  293
                 ||||||||||||||
Sbjct  77324157  CTGGTGTTCTCTCC  77324170
```

Range 18: 79568122 to 79568135

| Score | Expect | Identities | Gaps | Strand | Frame |
|---|---|---|---|---|---|
| 28.2 bits(14) | 256() | 14/14(100%) | 0/14(0%) | Plus/Plus | |

Features:
**catenin alpha-2 isoform 4**

```
Query  283       GTGTTCTCTCCTTT  296
                 ||||||||||||||
Sbjct  79568122  GTGTTCTCTCCTTT  79568135
```

Range 19: 109363199 to 109363212

| Score | Expect | Identities | Gaps | Strand | Frame |
|---|---|---|---|---|---|
| 28.2 bits(14) | 256() | 14/14(100%) | 0/14(0%) | Plus/Plus | |

Features:
**SH3 domain-containing RING finger protein 3 precursorSH3 domain-containing RING finger protein 3 isoform X1**

```
Query  286        TTCTCTCCTTTATT  299
                  ||||||||||||||
Sbjct  109363199  TTCTCTCCTTTATT  109363212
```

Range 20: 113405393 to 113405406

| Score | Expect | Identities | Gaps | Strand | Frame |
|---|---|---|---|---|---|
| 28.2 bits(14) | 256() | 14/14(100%) | 0/14(0%) | Plus/Plus | |

Features:
**126999 bp at 5' side: paired box protein Pax-8 isoform PAX8E990 bp at 3' side: Ig kappa chain V-I region Walker-like**

```
Query  285        GTTCTCTCCTTTAT  298
                  ||||||||||||||
Sbjct  113405393  GTTCTCTCCTTTAT  113405406
```

Range 21: 122001383 to 122001396

| Score | Expect | Identities | Gaps | Strand | Frame |
|---|---|---|---|---|---|
| 28.2 bits(14) | 256() | 14/14(100%) | 0/14(0%) | Plus/Plus | |

Features:
**236176 bp at 5' side: translin isoform 22024255 bp at 3' side: contactin-associated protein-like 5 isoform X1**

```
Query  279        GCTGGTGTTCTCTC  292
                  ||||||||||||||
Sbjct  122001383  GCTGGTGTTCTCTC  122001396
```

Range 22: 123878365 to 123878378

| Score | Expect | Identities | Gaps | Strand | Frame |
|---|---|---|---|---|---|
| 28.2 bits(14) | 256() | 14/14(100%) | 0/14(0%) | Plus/Plus | |

Features:
**2113158 bp at 5' side: translin isoform 2147273 bp at 3' side: contactin-associated protein-like 5 isoform X1**

```
Query  286        TTCTCTCCTTTATT  299
                  ||||||||||||||
Sbjct  123878365  TTCTCTCCTTTATT  123878378
```

Range 23: 125559221 to 125559234

| Score | Expect | Identities | Gaps | Strand | Frame |
|---|---|---|---|---|---|
| 28.2 bits(14) | 256() | 14/14(100%) | 0/14(0%) | Plus/Plus | |

Features:
**644933 bp at 5' side: contactin-associated protein-like 5 precursor977020 bp at 3' side: uncharacterized protein LOC105373602 isoform X1**

```
Query  284        TGTTCTCTCCTTTA  297
                  ||||||||||||||
Sbjct  125559221  TGTTCTCTCCTTTA  125559234
```

Range 24: 126274491 to 126274504

| Score | Expect | Identities | Gaps | Strand | Frame |
|---|---|---|---|---|---|
| 28.2 bits(14) | 256() | 14/14(100%) | 0/14(0%) | Plus/Plus | |

Features:
**1360203 bp at 5' side: contactin-associated protein-like 5 precursor261750 bp at 3' side: uncharacterized protein LOC105373602 isoform X1**

```
Query  282        GGTGTTCTCTCCTT  295
                  ||||||||||||||
Sbjct  126274491  GGTGTTCTCTCCTT  126274504
```

Range 25: 131131565 to 131131578

| Score | Expect | Identities | Gaps | Strand | Frame |
|---|---|---|---|---|---|
| 28.2 bits(14) | 256() | 14/14(100%) | 0/14(0%) | Plus/Plus | |

Features:
**pleckstrin homology domain-containing family B member 2 i...pleckstrin homology domain-containing family B member 2 i...**

```
Query  286        TTCTCTCCTTTATT  299
                  ||||||||||||||
Sbjct  131131565  TTCTCTCCTTTATT  131131578
```

Range 26: 137797488 to 137797501

| Score | Expect | Identities | Gaps | Strand | Frame |
|---|---|---|---|---|---|
| 28.2 bits(14) | 256() | 14/14(100%) | 0/14(0%) | Plus/Plus | |

Features:
**120883 bp at 5' side: thrombospondin type-1 domain-containing protein 7B166991 bp at 3' side: histamine N-methyltransferase isoform 1**

```
Query  283        GTGTTCTCTCCTTT  296
                  ||||||||||||||
Sbjct  137797488  GTGTTCTCTCCTTT  137797501
```

Range 27: 156230216 to 156230229

| Score | Expect | Identities | Gaps | Strand | Frame |
|---|---|---|---|---|---|
| 28.2 bits(14) | 256() | 14/14(100%) | 0/14(0%) | Plus/Plus | |

Features:
**1520389 bp at 5' side: G protein-activated inward rectifier potassium channel 1 ...95515 bp at 3' side: nuclear receptor subfamily 4 group A member 2 isoform X2**

```
Query  278        AGCTGGTGTTCTCT  291
                  ||||||||||||||
Sbjct  156230216  AGCTGGTGTTCTCT  156230229
```

Range 28: 158248303 to 158248316

| Score | Expect | Identities | Gaps | Strand | Frame |
|---|---|---|---|---|---|
| 28.2 bits(14) | 256() | 14/14(100%) | 0/14(0%) | Plus/Plus | |

Features:
**coiled-coil domain-containing protein 148 isoform 1coiled-coil domain-containing protein 148 isoform X2**

```
Query  286        TTCTCTCCTTTATT  299
                  ||||||||||||||
Sbjct  158248303  TTCTCTCCTTTATT  158248316
```

Range 29: 160470090 to 160470103

| Score | Expect | Identities | Gaps | Strand | Frame |
|---|---|---|---|---|---|
| 28.2 bits(14) | 256() | 14/14(100%) | 0/14(0%) | Plus/Plus | |

Features:
**RNA-binding motif, single-stranded-interacting protein 1 ...RNA-binding motif, single-stranded-interacting protein 1 ...**

```
Query  276        CAAGCTGGTGTTCT  289
                  ||||||||||||||
Sbjct  160470090  CAAGCTGGTGTTCT  160470103
```

Range 30: 160739520 to 160739533

| Score | Expect | Identities | Gaps | Strand | Frame |
|---|---|---|---|---|---|
| 28.2 bits(14) | 256() | 14/14(100%) | 0/14(0%) | Plus/Plus | |

Features:
**246157 bp at 5' side: RNA-binding motif, single-stranded-interacting protein 1 ...421261 bp at 3' side: TRAF family member-associated NF-kappa-B activator isofor...**

```
Query  286        TTCTCTCCTTTATT  299
                  ||||||||||||||
Sbjct  160739520  TTCTCTCCTTTATT  160739533
```

Range 31: 164936173 to 164936186

| Score | Expect | Identities | Gaps | Strand | Frame |
|---|---|---|---|---|---|
| 28.2 bits(14) | 256() | 14/14(100%) | 0/14(0%) | Plus/Plus | |

Features:
**putative sodium-coupled neutral amino acid transporter 11...putative sodium-coupled neutral amino acid transporter 11...**

```
Query  286        TTCTCTCCTTTATT  299
                  ||||||||||||||
Sbjct  164936173  TTCTCTCCTTTATT  164936186
```

Range 32: 182110032 to 182110045

| Score | Expect | Identities | Gaps | Strand | Frame |
|---|---|---|---|---|---|
| 28.2 bits(14) | 256() | 14/14(100%) | 0/14(0%) | Plus/Plus | |

Features:
**protein phosphatase 1 regulatory subunit 1C isoform 2protein phosphatase 1 regulatory subunit 1C isoform 2**

```
Query  276        CAAGCTGGTGTTCT  289
                  ||||||||||||||
Sbjct  182110032  CAAGCTGGTGTTCT  182110045
```

Range 33: 191352842 to 191352855

| Score | Expect | Identities | Gaps | Strand | Frame |
|---|---|---|---|---|---|
| 28.2 bits(14) | 256() | 14/14(100%) | 0/14(0%) | Plus/Plus | |

Features:
**unconventional myosin-Ib isoform 2unconventional myosin-Ib isoform X1**

```
Query  280        CTGGTGTTCTCTCC  293
                  ||||||||||||||
Sbjct  191352842  CTGGTGTTCTCTCC  191352855
```

Range 34: 200312585 to 200312598

| Score | Expect | Identities | Gaps | Strand | Frame |
|---|---|---|---|---|---|
| 28.2 bits(14) | 256() | 14/14(100%) | 0/14(0%) | Plus/Plus | |

Features:
**SPATS2-like protein isoform dSPATS2-like protein isoform X3**

```
Query  286        TTCTCTCCTTTATT  299
                  ||||||||||||||
Sbjct  200312585  TTCTCTCCTTTATT  200312598
```

Range 35: 211724479 to 211724492

| Score | Expect | Identities | Gaps | Strand | Frame |
|---|---|---|---|---|---|
| 28.2 bits(14) | 256() | 14/14(100%) | 0/14(0%) | Plus/Plus | |

Features:
**receptor tyrosine-protein kinase erbB-4 isoform JM-a/CVT-...receptor tyrosine-protein kinase erbB-4 isoform JM-a/CVT-...**

```
Query  277        AAGCTGGTGTTCTC  290
                  ||||||||||||||
Sbjct  211724479  AAGCTGGTGTTCTC  211724492
```

Range 36: 217044631 to 217044644

| Score | Expect | Identities | Gaps | Strand | Frame |
|---|---|---|---|---|---|
| 28.2 bits(14) | 256() | 14/14(100%) | 0/14(0%) | Plus/Plus | |

Features:
**184597 bp at 5' side: spermatid nuclear transition protein 1759815 bp at 3' side: tensin-1 isoform X13**

```
Query  286        TTCTCTCCTTTATT  299
                  ||||||||||||||
Sbjct  217044631  TTCTCTCCTTTATT  217044644
```

Range 37: 221398105 to 221398118

| Score | Expect | Identities | Gaps | Strand | Frame |
|---|---|---|---|---|---|
| 28.2 bits(14) | 256() | 14/14(100%) | 0/14(0%) | Plus/Plus | |

Features:
**1756377 bp at 5' side: anion exchange protein 3 isoform X527910 bp at 3' side: ephrin type-A receptor 4 isoform a precursor**

```
Query  287        TCTCTCCTTTATTG  300
                  ||||||||||||||
Sbjct  221398105  TCTCTCCTTTATTG  221398118
```

Range 38: 221665744 to 221665757

| Score | Expect | Identities | Gaps | Strand | Frame |
|---|---|---|---|---|---|
| 28.2 bits(14) | 256() | 14/14(100%) | 0/14(0%) | Plus/Plus | |

Features:
**93496 bp at 5' side: ephrin type-A receptor 4 isoform X1535417 bp at 3' side: paired box protein Pax-3 isoform X1**

```
Query  286        TTCTCTCCTTTATT  299
                  ||||||||||||||
Sbjct  221665744  TTCTCTCCTTTATT  221665757
```

Range 39: 224876858 to 224876871

| Score | Expect | Identities | Gaps | Strand | Frame |
|---|---|---|---|---|---|
| 28.2 bits(14) | 256() | 14/14(100%) | 0/14(0%) | Plus/Plus | |

Features:
**dedicator of cytokinesis protein 10 DOCK10.1dedicator of cytokinesis protein 10 isoform X4**

```
Query  286        TTCTCTCCTTTATT  299
                  ||||||||||||||
Sbjct  224876858  TTCTCTCCTTTATT  224876871
```

Range 40: 228961312 to 228961325

| Score | Expect | Identities | Gaps | Strand | Frame |
|---|---|---|---|---|---|
| 28.2 bits(14) | 256() | 14/14(100%) | 0/14(0%) | Plus/Plus | |

Features:
**779714 bp at 5' side: A-kinase anchor protein SPHKAP isoform 264307 bp at 3' side: PTB-containing, cubilin and LRP1-interacting protein isof...**

```
Query  286        TTCTCTCCTTTATT  299
                  ||||||||||||||
Sbjct  228961312  TTCTCTCCTTTATT  228961325
```

Range 41: 17505442 to 17505455

| Score | Expect | Identities | Gaps | Strand | Frame |
|---|---|---|---|---|---|
| 28.2 bits(14) | 256() | 14/14(100%) | 0/14(0%) | Plus/Minus | |

Features:
**917323 bp at 5' side: protein FAM49A isoform X15349 bp at 3' side: RAD51-associated protein 2 isoform X2**

```
Query  286       TTCTCTCCTTTATT  299
                 ||||||||||||||
Sbjct  17505455  TTCTCTCCTTTATT  17505442
```

Range 42: 21960168 to 21960181

| Score | Expect | Identities | Gaps | Strand | Frame |
|---|---|---|---|---|---|
| 28.2 bits(14) | 256() | 14/14(100%) | 0/14(0%) | Plus/Minus | |

Features:
**816896 bp at 5' side: tudor domain-containing protein 151422646 bp at 3' side: transcription initiation factor TFIID subunit 4-like**

```
Query  286       TTCTCTCCTTTATT  299
                 ||||||||||||||
Sbjct  21960181  TTCTCTCCTTTATT  21960168
```

Range 43: 35257623 to 35257636

| Score | Expect | Identities | Gaps | Strand | Frame |
|---|---|---|---|---|---|
| 28.2 bits(14) | 256() | 14/14(100%) | 0/14(0%) | Plus/Minus | |

Features:
**1658402 bp at 5' side: protein FAM98A isoform 11098657 bp at 3' side: cysteine-rich motor neuron 1 protein isoform X1**

```
Query  286       TTCTCTCCTTTATT  299
                 ||||||||||||||
Sbjct  35257636  TTCTCTCCTTTATT  35257623
```

Range 44: 58737947 to 58737960

| Score | Expect | Identities | Gaps | Strand | Frame |
|---|---|---|---|---|---|
| 28.2 bits(14) | 256() | 14/14(100%) | 0/14(0%) | Plus/Minus | |

Features:
**496707 bp at 5' side: E3 ubiquitin-protein ligase FANCL isoform X71714605 bp at 3' side: B-cell lymphoma/leukemia 11A isoform X4**

```
Query  283       GTGTTCTCTCCTTT  296
                 ||||||||||||||
Sbjct  58737960  GTGTTCTCTCCTTT  58737947
```

Range 45: 66242938 to 66242955

| Score | Expect | Identities | Gaps | Strand | Frame |
|---|---|---|---|---|---|
| 28.2 bits(14) | 256() | 17/18(94%) | 0/18(0%) | Plus/Minus | |

Features:
**808188 bp at 5' side: WAS/WASL-interacting protein family member 1-like isoform X1192902 bp at 3' side: homeobox protein Meis1 isoform X1**

```
Query  279       GCTGGTGTTCTCTCCTTT  296
                 ||||||||| ||||||||
Sbjct  66242955  GCTGGTGTTTTCTCCTTT  66242938
```

Range 46: 74969343 to 74969356

| Score | Expect | Identities | Gaps | Strand | Frame |
|---|---|---|---|---|---|
| 28.2 bits(14) | 256() | 14/14(100%) | 0/14(0%) | Plus/Minus | |

Features:
**DNA polymerase epsilon subunit 4**

```
Query  270       GTGTAACAAGCTGG  283
                 ||||||||||||||
Sbjct  74969356  GTGTAACAAGCTGG  74969343
```

Range 47: 80604706 to 80604719

| Score | Expect | Identities | Gaps | Strand | Frame |
|---|---|---|---|---|---|
| 28.2 bits(14) | 256() | 14/14(100%) | 0/14(0%) | Plus/Minus | |

Features:
**catenin alpha-2 isoform 4catenin alpha-2 isoform 3**

```
Query  286       TTCTCTCCTTTATT  299
                 ||||||||||||||
Sbjct  80604719  TTCTCTCCTTTATT  80604706
```

Range 48: 88923783 to 88923796

| Score | Expect | Identities | Gaps | Strand | Frame |
|---|---|---|---|---|---|
| 28.2 bits(14) | 256() | 14/14(100%) | 0/14(0%) | Plus/Minus | |

Features:
**8105 bp at 5' side: IGKV7-37855 bp at 3' side: IGKV2-4**

```
Query  286       TTCTCTCCTTTATT  299
                 ||||||||||||||
Sbjct  88923796  TTCTCTCCTTTATT  88923783
```

Range 49: 91750849 to 91750862

| Score | Expect | Identities | Gaps | Strand | Frame |
|---|---|---|---|---|---|
| 28.2 bits(14) | 256() | 14/14(100%) | 0/14(0%) | Plus/Minus | |

Features:
**237574 bp at 5' side: adhesive plaque matrix protein-like2837573 bp at 3' side: putative aquaporin-7-like protein 3 isoform X7**

```
Query  286       TTCTCTCCTTTATT  299
                 ||||||||||||||
Sbjct  91750862  TTCTCTCCTTTATT  91750849
```

Range 50: 122775392 to 122775409

| Score | Expect | Identities | Gaps | Strand | Frame |
|---|---|---|---|---|---|
| 28.2 bits(14) | 256() | 17/18(94%) | 0/18(0%) | Plus/Minus | |

Features:
**1010185 bp at 5' side: translin isoform 21250242 bp at 3' side: contactin-associated protein-like 5 isoform X1**

```
Query  281        TGGTGTTCTCTCCTTTAT  298
                  ||||| ||||||||||||
Sbjct  122775409  TGGTGATCTCTCCTTTAT  122775392
```

Range 51: 126335156 to 126335169

| Score | Expect | Identities | Gaps | Strand | Frame |
|---|---|---|---|---|---|
| 28.2 bits(14) | 256() | 14/14(100%) | 0/14(0%) | Plus/Minus | |

Features:
**1420868 bp at 5' side: contactin-associated protein-like 5 precursor201085 bp at 3' side: uncharacterized protein LOC105373602 isoform X1**

```
Query  276        CAAGCTGGTGTTCT  289
                  ||||||||||||||
Sbjct  126335169  CAAGCTGGTGTTCT  126335156
```

Range 52: 130868397 to 130868410

| Score | Expect | Identities | Gaps | Strand | Frame |
|---|---|---|---|---|---|
| 28.2 bits(14) | 256() | 14/14(100%) | 0/14(0%) | Plus/Minus | |

Features:
**rho guanine nucleotide exchange factor 4 isoform X1rho guanine nucleotide exchange factor 4 isoform X4**

```
Query  283        GTGTTCTCTCCTTT  296
                  ||||||||||||||
Sbjct  130868410  GTGTTCTCTCCTTT  130868397
```

Range 53: 132884731 to 132884744

| Score | Expect | Identities | Gaps | Strand | Frame |
|---|---|---|---|---|---|

| | | | | |
|---|---|---|---|---|
| 28.2 bits(14) | 256() | 14/14(100%) | 0/14(0%) | Plus/Minus |

Features:
**nck-associated protein 5 isoform X3nck-associated protein 5 isoform X1**

```
Query  285        GTTCTCTCCTTTAT  298
                  ||||||||||||||
Sbjct  132884744  GTTCTCTCCTTTAT  132884731
```

Range 54: 134314276 to 134314289

| Score | Expect | Identities | Gaps | Strand | Frame |
|---|---|---|---|---|---|
| 28.2 bits(14) | 256() | 14/14(100%) | 0/14(0%) | Plus/Minus | |

Features:
**alpha-1,6-mannosylglycoprotein 6-beta-N-acetylglucosaminy...alpha-1,6-mannosylglycoprotein 6-beta-N-acetylglucosaminy...**

```
Query  286        TTCTCTCCTTTATT  299
                  ||||||||||||||
Sbjct  134314289  TTCTCTCCTTTATT  134314276
```

Range 55: 139444509 to 139444522

| Score | Expect | Identities | Gaps | Strand | Frame |
|---|---|---|---|---|---|
| 28.2 bits(14) | 256() | 14/14(100%) | 0/14(0%) | Plus/Minus | |

Features:
**664268 bp at 5' side: neurexophilin-2 precursor788664 bp at 3' side: low-density lipoprotein receptor-related protein 1B precu...**

```
Query  286        TTCTCTCCTTTATT  299
                  ||||||||||||||
Sbjct  139444522  TTCTCTCCTTTATT  139444509
```

Range 56: 142636578 to 142636591

| Score | Expect | Identities | Gaps | Strand | Frame |
|---|---|---|---|---|---|
| 28.2 bits(14) | 256() | 14/14(100%) | 0/14(0%) | Plus/Minus | |

Features:
**505738 bp at 5' side: low-density lipoprotein receptor-related protein 1B isofo...248777 bp at 3' side: kynureninase isoform a**

```
Query  286        TTCTCTCCTTTATT  299
                  ||||||||||||||
Sbjct  142636591  TTCTCTCCTTTATT  142636578
```

Range 57: 145614127 to 145614140

| Score | Expect | Identities | Gaps | Strand | Frame |
|---|---|---|---|---|---|
| 28.2 bits(14) | 256() | 14/14(100%) | 0/14(0%) | Plus/Minus | |

Features:
**1176001 bp at 5' side: zinc finger E-box-binding homeobox 2 isoform X22231013 bp at 3' side: activin receptor type-2A isoform 1 precursor**

```
Query  283        GTGTTCTCTCCTTT  296
                  ||||||||||||||
Sbjct  145614140  GTGTTCTCTCCTTT  145614127
```

Range 58: 150402853 to 150402866

| Score | Expect | Identities | Gaps | Strand | Frame |
|---|---|---|---|---|---|
| 28.2 bits(14) | 256() | 14/14(100%) | 0/14(0%) | Plus/Minus | |

Features:
**815756 bp at 5' side: methylmalonic aciduria and homocystinuria type D protein,...67121 bp at 3' side: rho-related GTP-binding protein RhoE precursor**

```
Query  286        TTCTCTCCTTTATT  299
                  ||||||||||||||
Sbjct  150402866  TTCTCTCCTTTATT  150402853
```

Range 59: 150829585 to 150829598

| Score | Expect | Identities | Gaps | Strand | Frame |
|---|---|---|---|---|---|
| 28.2 bits(14) | 256() | 14/14(100%) | 0/14(0%) | Plus/Minus | |

Features:
**342168 bp at 5' side: rho-related GTP-binding protein RhoE precursor421308 bp at 3' side: RNA-binding protein 43**

```
Query  287        TCTCTCCTTTATTG  300
                  ||||||||||||||
Sbjct  150829598  TCTCTCCTTTATTG  150829585
```

Range 60: 151623928 to 151623941

| Score | Expect | Identities | Gaps | Strand | Frame |
|---|---|---|---|---|---|
| 28.2 bits(14) | 256() | 14/14(100%) | 0/14(0%) | Plus/Minus | |

Features:
**nebulin isoform X5nebulin isoform X22**

```
Query  283        GTGTTCTCTCCTTT  296
                  ||||||||||||||
Sbjct  151623941  GTGTTCTCTCCTTT  151623928
```

Range 61: 156189205 to 156189218

| Score | Expect | Identities | Gaps | Strand | Frame |
|---|---|---|---|---|---|
| 28.2 bits(14) | 256() | 14/14(100%) | 0/14(0%) | Plus/Minus | |

Features:
**1479378 bp at 5' side: G protein-activated inward rectifier potassium channel 1 ...136526 bp at 3' side: nuclear receptor subfamily 4 group A member 2 isoform X2**

```
Query  286        TTCTCTCCTTTATT  299
                  ||||||||||||||
Sbjct  156189218  TTCTCTCCTTTATT  156189205
```

Range 62: 157046508 to 157046521

| Score | Expect | Identities | Gaps | Strand | Frame |
|---|---|---|---|---|---|
| 28.2 bits(14) | 256() | 14/14(100%) | 0/14(0%) | Plus/Minus | |

Features:
**463590 bp at 5' side: glycerol-3-phosphate dehydrogenase, mitochondrial isoform X4211562 bp at 3' side: polypeptide N-acetylgalactosaminyltransferase 5**

```
Query  279        GCTGGTGTTCTCTC  292
                  ||||||||||||||
Sbjct  157046521  GCTGGTGTTCTCTC  157046508
```

Range 63: 174962382 to 174962399

| Score | Expect | Identities | Gaps | Strand | Frame |
|---|---|---|---|---|---|
| 28.2 bits(14) | 256() | 17/18(94%) | 0/18(0%) | Plus/Minus | |

Features:
**N-chimaerin isoform 2N-chimaerin isoform 1**

```
Query  274        AACAAGCTGGTGTTCTCT  291
                  ||||||||| ||||||||
Sbjct  174962399  AACAAGCTGCTGTTCTCT  174962382
```

Range 64: 177679026 to 177679039

| Score | Expect | Identities | Gaps | Strand | Frame |
|---|---|---|---|---|---|
| 28.2 bits(14) | 256() | 14/14(100%) | 0/14(0%) | Plus/Minus | |

Features:

**dual 3',5'-cyclic-AMP and -GMP phosphodiesterase 11A isof...dual 3',5'-cyclic-AMP and -GMP phosphodiesterase 11A isof...**

```
Query  286        TTCTCTCCTTTATT  299
                  ||||||||||||||
Sbjct  177679039  TTCTCTCCTTTATT  177679026
```

Range 65: 180902954 to 180902967

| Score | Expect | Identities | Gaps | Strand | Frame |
|---|---|---|---|---|---|
| 28.2 bits(14) | 256() | 14/14(100%) | 0/14(0%) | Plus/Minus | |

Features:
**909613 bp at 5' side: pre-mRNA-splicing factor CWC22 homolog isoform X179076 bp at 3' side: ubiquitin-conjugating enzyme E2 E3**

```
Query  286        TTCTCTCCTTTATT  299
                  ||||||||||||||
Sbjct  180902967  TTCTCTCCTTTATT  180902954
```

Range 66: 198610197 to 198610210

| Score | Expect | Identities | Gaps | Strand | Frame |
|---|---|---|---|---|---|
| 28.2 bits(14) | 256() | 14/14(100%) | 0/14(0%) | Plus/Minus | |

Features:
**463235 bp at 5' side: inactive phospholipase C-like protein 1 isoform X3662001 bp at 3' side: DNA-binding protein SATB2**

```
Query  286        TTCTCTCCTTTATT  299
                  ||||||||||||||
Sbjct  198610210  TTCTCTCCTTTATT  198610197
```

Range 67: 199313792 to 199313805

| Score | Expect | Identities | Gaps | Strand | Frame |
|---|---|---|---|---|---|
| 28.2 bits(14) | 256() | 14/14(100%) | 0/14(0%) | Plus/Minus | |

Features:
**DNA-binding protein SATB2DNA-binding protein SATB2 isoform X1**

```
Query  285        GTTCTCTCCTTTAT  298
                  ||||||||||||||
Sbjct  199313805  GTTCTCTCCTTTAT  199313792
```

Range 68: 211386018 to 211386031

| Score | Expect | Identities | Gaps | Strand | Frame |
|---|---|---|---|---|---|
| 28.2 bits(14) | 256() | 14/14(100%) | 0/14(0%) | Plus/Minus | |

Features:
**receptor tyrosine-protein kinase erbB-4 isoform JM-a/CVT-...receptor tyrosine-protein kinase erbB-4 isoform JM-a/CVT-...**

```
Query  284        TGTTCTCTCCTTTA  297
                  ||||||||||||||
Sbjct  211386031  TGTTCTCTCCTTTA  211386018
```

Range 69: 213962581 to 213962594

| Score | Expect | Identities | Gaps | Strand | Frame |
|---|---|---|---|---|---|
| 28.2 bits(14) | 256() | 14/14(100%) | 0/14(0%) | Plus/Minus | |

Features:
**sperm-associated antigen 16 protein isoform X12sperm-associated antigen 16 protein isoform X10**

```
Query  286        TTCTCTCCTTTATT  299
                  ||||||||||||||
Sbjct  213962594  TTCTCTCCTTTATT  213962581
```

Range 70: 214491694 to 214491707

| Score | Expect | Identities | Gaps | Strand | Frame |
|---|---|---|---|---|---|
| 28.2 bits(14) | 256() | 14/14(100%) | 0/14(0%) | Plus/Minus | |

Features:
**von Willebrand factor C domain-containing protein 2-like ...von Willebrand factor C domain-containing protein 2-like ...**

```
Query  283        GTGTTCTCTCCTTT  296
                  ||||||||||||||
Sbjct  214491707  GTGTTCTCTCCTTT  214491694
```

Range 71: 224899638 to 224899651

| Score | Expect | Identities | Gaps | Strand | Frame |
|---|---|---|---|---|---|
| 28.2 bits(14) | 256() | 14/14(100%) | 0/14(0%) | Plus/Minus | |

Features:
**dedicator of cytokinesis protein 10 DOCK10.1dedicator of cytokinesis protein 10 isoform X4**

```
Query  284        TGTTCTCTCCTTTA  297
                  ||||||||||||||
Sbjct  224899651  TGTTCTCTCCTTTA  224899638
```

Range 72: 228548581 to 228548594

| Score | Expect | Identities | Gaps | Strand | Frame |
|---|---|---|---|---|---|
| 28.2 bits(14) | 256() | 14/14(100%) | 0/14(0%) | Plus/Minus | |

Features:
**366983 bp at 5' side: A-kinase anchor protein SPHKAP isoform 2477038 bp at 3' side: PTB-containing, cubilin and LRP1-interacting protein isof...**

```
Query  283        GTGTTCTCTCCTTT  296
                  ||||||||||||||
Sbjct  228548594  GTGTTCTCTCCTTT  228548581
```

Range 73: 230289969 to 230289982

| Score | Expect | Identities | Gaps | Strand | Frame |
|---|---|---|---|---|---|
| 28.2 bits(14) | 256() | 14/14(100%) | 0/14(0%) | Plus/Minus | |

Features:
**nuclear body protein SP140 isoform 4nuclear body protein SP140 isoform X14**

```
Query  284        TGTTCTCTCCTTTA  297
                  ||||||||||||||
Sbjct  230289982  TGTTCTCTCCTTTA  230289969
```

Range 74: 231469990 to 231470003

| Score | Expect | Identities | Gaps | Strand | Frame |
|---|---|---|---|---|---|
| 28.2 bits(14) | 256() | 14/14(100%) | 0/14(0%) | Plus/Minus | |

Features:
**5637 bp at 5' side: nucleolin50379 bp at 3' side: neuromedin-U receptor 1 isoform X4**

```
Query  287        TCTCTCCTTTATTG  300
                  ||||||||||||||
Sbjct  231470003  TCTCTCCTTTATTG  231469990
```

Homo sapiens chromosome 4, GRCh38.p2 Primary Assembly
Sequence ID: **ref|NC_000004.12|** Length: 190214555 Number of Matches: 63
Range 1: 40780306 to 40780326

| Score | Expect | Identities | Gaps | Strand | Frame |
|---|---|---|---|---|---|
| 34.2 bits(17) | 4.1() | 20/21(95%) | 0/21(0%) | Plus/Minus | |

Features:

**putative methyltransferase NSUN7**

```
Query  280       CTGGTGTTCTCTCCTTTATTG  300
                 ||||| |||||||||||||||
Sbjct  40780326  CTGGTCTTCTCTCCTTTATTG  40780306
```

Range 2: 91721375 to 91721391

| Score | Expect | Identities | Gaps | Strand | Frame |
|---|---|---|---|---|---|
| 34.2 bits(17) | 4.1() | 17/17(100%) | 0/17(0%) | Plus/Minus | |

Features:
**782640 bp at 5' side: serine-rich coiled-coil domain-containing protein 1 isofo...583266 bp at 3' side: glutamate receptor ionotropic, delta-2 isoform 1 precursor**

```
Query  283       GTGTTCTCTCCTTTATT  299
                 |||||||||||||||||
Sbjct  91721391  GTGTTCTCTCCTTTATT  91721375
```

Range 3: 91529788 to 91529803

| Score | Expect | Identities | Gaps | Strand | Frame |
|---|---|---|---|---|---|
| 32.2 bits(16) | 16() | 16/16(100%) | 0/16(0%) | Plus/Plus | |

Features:
**serine-rich coiled-coil domain-containing protein 1 isofo...serine-rich coiled-coil domain-containing protein 1 isofo...**

```
Query  284       TGTTCTCTCCTTTATT  299
                 ||||||||||||||||
Sbjct  91529788  TGTTCTCTCCTTTATT  91529803
```

Range 4: 180849629 to 180849644

| Score | Expect | Identities | Gaps | Strand | Frame |
|---|---|---|---|---|---|
| 32.2 bits(16) | 16() | 16/16(100%) | 0/16(0%) | Plus/Plus | |

Features:
**3252036 bp at 5' side: uncharacterized protein LOC285500 isoform X31474377 bp at 3' side: teneurin-3**

```
Query  279        GCTGGTGTTCTCTCCT  294
                  ||||||||||||||||
Sbjct  180849629  GCTGGTGTTCTCTCCT  180849644
```

Range 5: 53685506 to 53685520

| Score | Expect | Identities | Gaps | Strand | Frame |
|---|---|---|---|---|---|
| 30.2 bits(15) | 65() | 15/15(100%) | 0/15(0%) | Plus/Plus | |

Features:
**109119 bp at 5' side: COMM domain-containing protein 5-like324575 bp at 3' side: cysteine-rich hydrophobic domain-containing protein 2 iso...**

```
Query  271       TGTAACAAGCTGGTG  285
                 |||||||||||||||
Sbjct  53685506  TGTAACAAGCTGGTG  53685520
```

Range 6: 150750232 to 150750246

| Score | Expect | Identities | Gaps | Strand | Frame |
|---|---|---|---|---|---|
| 30.2 bits(15) | 65() | 15/15(100%) | 0/15(0%) | Plus/Plus | |

Features:
**lipopolysaccharide-responsive and beige-like anchor prote...lipopolysaccharide-responsive and beige-like anchor prote...**

```
Query  284        TGTTCTCTCCTTTAT  298
                  |||||||||||||||
Sbjct  150750232  TGTTCTCTCCTTTAT  150750246
```

Range 7: 159162621 to 159162639

| Score | Expect | Identities | Gaps | Strand | Frame |
|---|---|---|---|---|---|
| 30.2 bits(15) | 65() | 18/19(95%) | 0/19(0%) | Plus/Plus | |

Features:
**rap guanine nucleotide exchange factor 2 isoform X6rap guanine nucleotide exchange factor 2 isoform X3**

```
Query  279        GCTGGTGTTCTCTCCTTTA  297
                  ||||||||||||||| ||||
Sbjct  159162621  GCTGGTGTTCTCTCATTTA  159162639
```

Range 8: 159937708 to 159937722

| Score | Expect | Identities | Gaps | Strand | Frame |
|---|---|---|---|---|---|
| 30.2 bits(15) | 65() | 15/15(100%) | 0/15(0%) | Plus/Plus | |

Features:
**579569 bp at 5' side: rap guanine nucleotide exchange factor 21448025 bp at 3' side: follistatin-related protein 5 isoform c precursor**

```
Query  283        GTGTTCTCTCCTTTA  297
                  |||||||||||||||
Sbjct  159937708  GTGTTCTCTCCTTTA  159937722
```

Range 9: 170217792 to 170217806

| Score | Expect | Identities | Gaps | Strand | Frame |
|---|---|---|---|---|---|
| 30.2 bits(15) | 65() | 15/15(100%) | 0/15(0%) | Plus/Plus | |

Features:
**111039 bp at 5' side: DNA dC->dU-editing enzyme APOBEC-3G-like1596775 bp at 3' side: polypeptide N-acetylgalactosaminyltransferase-like 6**

```
Query  273        TAACAAGCTGGTGTT  287
                  |||||||||||||||
Sbjct  170217792  TAACAAGCTGGTGTT  170217806
```

Range 10: 59259008 to 59259022

| Score | Expect | Identities | Gaps | Strand | Frame |
|---|---|---|---|---|---|
| 30.2 bits(15) | 65() | 15/15(100%) | 0/15(0%) | Plus/Minus | |

Features:
**2148657 bp at 5' side: insulin-like growth factor-binding protein 7 isoform 2 pr...2238272 bp at 3' side: latrophilin-3 isoform X6**

```
Query  285       GTTCTCTCCTTTATT  299
                 |||||||||||||||
Sbjct  59259022  GTTCTCTCCTTTATT  59259008
```

Range 11: 76550202 to 76550216

| Score | Expect | Identities | Gaps | Strand | Frame |
|---|---|---|---|---|---|
| 30.2 bits(15) | 65() | 15/15(100%) | 0/15(0%) | Plus/Minus | |

Features:
**protein Shroom3 isoform X3protein Shroom3 isoform X3**

```
Query  282       GGTGTTCTCTCCTTT  296
                 |||||||||||||||
Sbjct  76550216  GGTGTTCTCTCCTTT  76550202
```

Range 12: 130549844 to 130549858

| Score | Expect | Identities | Gaps | Strand | Frame |
|---|---|---|---|---|---|
| 30.2 bits(15) | 65() | 15/15(100%) | 0/15(0%) | Plus/Minus | |

Features:
**1439766 bp at 5' side: UPF0462 protein C4orf33 isoform X22600283 bp at 3' side: protocadherin-10 isoform X1**

```
Query  284        TGTTCTCTCCTTTAT  298
```

```
                    |||||||||||||||
Sbjct  130549858  TGTTCTCTCCTTTAT  130549844
```

Range 13: 173838989 to 173839003

| Score | Expect | Identities | Gaps | Strand | Frame |
|---|---|---|---|---|---|
| 30.2 bits(15) | 65() | 15/15(100%) | 0/15(0%) | Plus/Minus | |

Features:
**309700 bp at 5' side: heart- and neural crest derivatives-expressed protein 2398409 bp at 3' side: F-box only protein 8**

```
Query  273        TAACAAGCTGGTGTT  287
                  |||||||||||||||
Sbjct  173839003  TAACAAGCTGGTGTT  173838989
```

Range 14: 189191874 to 189191888

| Score | Expect | Identities | Gaps | Strand | Frame |
|---|---|---|---|---|---|
| 30.2 bits(15) | 65() | 15/15(100%) | 0/15(0%) | Plus/Minus | |

Features:
**1044502 bp at 5' side: probable E3 ubiquitin-protein ligase TRIML1 isoform X2749122 bp at 3' side: protein FRG1**

```
Query  280        CTGGTGTTCTCTCCT  294
                  |||||||||||||||
Sbjct  189191888  CTGGTGTTCTCTCCT  189191874
```

Range 15: 3195908 to 3195921

| Score | Expect | Identities | Gaps | Strand | Frame |
|---|---|---|---|---|---|
| 28.2 bits(14) | 256() | 14/14(100%) | 0/14(0%) | Plus/Plus | |

Features:
**huntingtin**

```
Query  279      GCTGGTGTTCTCTC  292
                ||||||||||||||
Sbjct  3195908  GCTGGTGTTCTCTC  3195921
```

Range 16: 3222569 to 3222582

| Score | Expect | Identities | Gaps | Strand | Frame |
|---|---|---|---|---|---|
| 28.2 bits(14) | 256() | 14/14(100%) | 0/14(0%) | Plus/Plus | |

Features:
**huntingtin**

```
Query  276      CAAGCTGGTGTTCT  289
                ||||||||||||||
Sbjct  3222569  CAAGCTGGTGTTCT  3222582
```

Range 17: 8398451 to 8398464

| Score | Expect | Identities | Gaps | Strand | Frame |
|---|---|---|---|---|---|
| 28.2 bits(14) | 256() | 14/14(100%) | 0/14(0%) | Plus/Plus | |

Features:
**peroxisomal acyl-coenzyme A oxidase 3 isoform X4peroxisomal acyl-coenzyme A oxidase 3 isoform X3**

```
Query  286      TTCTCTCCTTTATT  299
                ||||||||||||||
Sbjct  8398451  TTCTCTCCTTTATT  8398464
```

Range 18: 10767778 to 10767791

| Score | Expect | Identities | Gaps | Strand | Frame |
|---|---|---|---|---|---|
| 28.2 bits(14) | 256() | 14/14(100%) | 0/14(0%) | Plus/Plus | |

Features:
**168535 bp at 5' side: cytokine-dependent hematopoietic cell linker isoform X3631291 bp at 3' side: heparan sulfate glucosamine 3-O-sulfotransferase 1 precursor**

```
Query  285       GTTCTCTCCTTTAT  298
                 ||||||||||||||
Sbjct  10767778  GTTCTCTCCTTTAT  10767791
```

Range 19: 23298838 to 23298851

| Score | Expect | Identities | Gaps | Strand | Frame |
|---|---|---|---|---|---|
| 28.2 bits(14) | 256() | 14/14(100%) | 0/14(0%) | Plus/Plus | |

Features:
**783054 bp at 5' side: probable G-protein coupled receptor 125 isoform X3496971 bp at 3' side: peroxisome proliferator-activated receptor gamma coactiva...**

```
Query  281       TGGTGTTCTCTCCT  294
                 ||||||||||||||
Sbjct  23298838  TGGTGTTCTCTCCT  23298851
```

Range 20: 28459960 to 28459973

| Score | Expect | Identities | Gaps | Strand | Frame |
|---|---|---|---|---|---|
| 28.2 bits(14) | 256() | 14/14(100%) | 0/14(0%) | Plus/Plus | |

Features:
**1438923 bp at 5' side: stromal interaction molecule 2 isoform 3 precursor2261450 bp at 3' side: protocadherin-7 isoform c precursor**

```
Query  276       CAAGCTGGTGTTCT  289
                 ||||||||||||||
Sbjct  28459960  CAAGCTGGTGTTCT  28459973
```

Range 21: 55280092 to 55280105

| Score | Expect | Identities | Gaps | Strand | Frame |
|---|---|---|---|---|---|
| 28.2 bits(14) | 256() | 14/14(100%) | 0/14(0%) | Plus/Plus | |

Features:
**154799 bp at 5' side: vascular endothelial growth factor receptor 2 precursor66232 bp at 3' side: polyprenol reductase isoform X2**

```
Query  283       GTGTTCTCTCCTTT  296
                 ||||||||||||||
Sbjct  55280092  GTGTTCTCTCCTTT  55280105
```

Range 22: 60102953 to 60102970

| Score | Expect | Identities | Gaps | Strand | Frame |
|---|---|---|---|---|---|
| 28.2 bits(14) | 256() | 17/18(94%) | 0/18(0%) | Plus/Plus | |

Features:
**2992602 bp at 5' side: insulin-like growth factor-binding protein 7 isoform 2 pr...1394324 bp at 3' side: latrophilin-3 isoform X6**

```
Query  272       GTAACAAGCTGGTGTTCT  289
                 ||||||| ||||||||||
Sbjct  60102953  GTAACAATCTGGTGTTCT  60102970
```

Range 23: 66324693 to 66324706

| Score | Expect | Identities | Gaps | Strand | Frame |
|---|---|---|---|---|---|
| 28.2 bits(14) | 256() | 14/14(100%) | 0/14(0%) | Plus/Plus | |

Features:
**654951 bp at 5' side: ephrin type-A receptor 5 isoform X41147899 bp at 3' side: centromere protein C isoform X1**

```
Query  271       TGTAACAAGCTGGT  284
                 ||||||||||||||
Sbjct  66324693  TGTAACAAGCTGGT  66324706
```

Range 24: 68974948 to 68974961

| Score | Expect | Identities | Gaps | Strand | Frame |
|---|---|---|---|---|---|
| 28.2 bits(14) | 256() | 14/14(100%) | 0/14(0%) | Plus/Plus | |

Features:
**23188 bp at 5' side: UDP-glucuronosyltransferase 2A3 isoform X1121560 bp at 3' side: UDP-glucuronosyltransferase 2B7 precursor**

```
Query  286       TTCTCTCCTTTATT  299
                 ||||||||||||||
Sbjct  68974948  TTCTCTCCTTTATT  68974961
```

Range 25: 78178181 to 78178194

| Score | Expect | Identities | Gaps | Strand | Frame |
|---|---|---|---|---|---|
| 28.2 bits(14) | 256() | 14/14(100%) | 0/14(0%) | Plus/Plus | |

Features:
**extracellular matrix protein FRAS1 isoform 2 precursorextracellular matrix protein FRAS1 isoform X1**

```
Query  283       GTGTTCTCTCCTTT  296
                 ||||||||||||||
Sbjct  78178181  GTGTTCTCTCCTTT  78178194
```

Range 26: 85471750 to 85471763

| Score | Expect | Identities | Gaps | Strand | Frame |
|---|---|---|---|---|---|
| 28.2 bits(14) | 256() | 14/14(100%) | 0/14(0%) | Plus/Plus | |

Features:
**611159 bp at 5' side: WD repeat and FYVE domain-containing protein 3 isoform X898779 bp at 3' side: rho GTPase-activating protein 24 isoform 1**

```
Query  286       TTCTCTCCTTTATT  299
                 ||||||||||||||
Sbjct  85471750  TTCTCTCCTTTATT  85471763
```

Range 27: 93238356 to 93238369

| Score | Expect | Identities | Gaps | Strand | Frame |
|---|---|---|---|---|---|
| 28.2 bits(14) | 256() | 14/14(100%) | 0/14(0%) | Plus/Plus | |

Features:
**glutamate receptor ionotropic, delta-2 isoform 1 precursorglutamate receptor ionotropic, delta-2 isoform 2 precursor**

```
Query  284       TGTTCTCTCCTTTA  297
                 ||||||||||||||
Sbjct  93238356  TGTTCTCTCCTTTA  93238369
```

Range 28: 96483941 to 96483954

| Score | Expect | Identities | Gaps | Strand | Frame |
|---|---|---|---|---|---|
| 28.2 bits(14) | 256() | 14/14(100%) | 0/14(0%) | Plus/Plus | |

Features:
**642624 bp at 5' side: pyruvate dehydrogenase E1 component subunit alpha, testis...964797 bp at 3' side: sperm-tail PG-rich repeat-containing protein 2 isoform X1**

```
Query  278       AGCTGGTGTTCTCT  291
                 ||||||||||||||
Sbjct  96483941  AGCTGGTGTTCTCT  96483954
```

Range 29: 107878597 to 107878610

| Score | Expect | Identities | Gaps | Strand | Frame |
|---|---|---|---|---|---|
| 28.2 bits(14) | 256() | 14/14(100%) | 0/14(0%) | Plus/Plus | |

Features:
**177315 bp at 5' side: bifunctional 3'-phosphoadenosine 5'-phosphosulfate syntha...16944 bp at 3' side:**

**phosphatidylcholine:ceramide cholinephosphotransferase 2 ...**

```
Query  286        TTCTCTCCTTTATT  299
                  ||||||||||||||
Sbjct  107878597  TTCTCTCCTTTATT  107878610
```

Range 30: 109645138 to 109645151

| Score | Expect | Identities | Gaps | Strand | Frame |
|---|---|---|---|---|---|
| 28.2 bits(14) | 256() | 14/14(100%) | 0/14(0%) | Plus/Plus | |

Features:
**calcium uniporter regulatory subunit MCUb, mitochondrialcalcium uniporter regulatory subunit MCUb, mitochondrial ...**

```
Query  286        TTCTCTCCTTTATT  299
                  ||||||||||||||
Sbjct  109645138  TTCTCTCCTTTATT  109645151
```

Range 31: 112537460 to 112537473

| Score | Expect | Identities | Gaps | Strand | Frame |
|---|---|---|---|---|---|
| 28.2 bits(14) | 256() | 14/14(100%) | 0/14(0%) | Plus/Plus | |

Features:
**21985 bp at 5' side: neurogenin-22074 bp at 3' side: protein ZGRF1 isoform X1**

```
Query  286        TTCTCTCCTTTATT  299
                  ||||||||||||||
Sbjct  112537460  TTCTCTCCTTTATT  112537473
```

Range 32: 112732627 to 112732640

| Score | Expect | Identities | Gaps | Strand | Frame |
|---|---|---|---|---|---|
| 28.2 bits(14) | 256() | 14/14(100%) | 0/14(0%) | Plus/Plus | |

Features:
**75300 bp at 5' side: la-related protein 7 isoform 1171854 bp at 3' side: ankyrin-2 isoform X5**

```
Query  282        GGTGTTCTCTCCTT  295
                  ||||||||||||||
Sbjct  112732627  GGTGTTCTCTCCTT  112732640
```

Range 33: 115576936 to 115576949

| Score | Expect | Identities | Gaps | Strand | Frame |
|---|---|---|---|---|---|
| 28.2 bits(14) | 256() | 14/14(100%) | 0/14(0%) | Plus/Plus | |

Features:
**499900 bp at 5' side: bifunctional heparan sulfate N-deacetylase/N-sulfotransfe...1507335 bp at 3' side: translocating chain-associated membrane protein 1-like 1**

```
Query  286        TTCTCTCCTTTATT  299
                  ||||||||||||||
Sbjct  115576936  TTCTCTCCTTTATT  115576949
```

Range 34: 135614675 to 135614688

| Score | Expect | Identities | Gaps | Strand | Frame |
|---|---|---|---|---|---|
| 28.2 bits(14) | 256() | 14/14(100%) | 0/14(0%) | Plus/Plus | |

Features:
**1413482 bp at 5' side: polyadenylate-binding protein 4-like1906341 bp at 3' side: protocadherin-18 isoform 2 precursor**

```
Query  286        TTCTCTCCTTTATT  299
                  ||||||||||||||
Sbjct  135614675  TTCTCTCCTTTATT  135614688
```

Range 35: 143091521 to 143091534

| Score | Expect | Identities | Gaps | Strand | Frame |
|---|---|---|---|---|---|
| 28.2 bits(14) | 256() | 14/14(100%) | 0/14(0%) | Plus/Plus | |

Features:
**660262 bp at 5' side: type II inositol 3,4-bisphosphate 4-phosphatase isoform X1186493 bp at 3' side: uncharacterized protein LOC105377623**

```
Query  274        AACAAGCTGGTGTT  287
                  ||||||||||||||
Sbjct  143091521  AACAAGCTGGTGTT  143091534
```

Range 36: 148582335 to 148582352

| Score | Expect | Identities | Gaps | Strand | Frame |
|---|---|---|---|---|---|
| 28.2 bits(14) | 256() | 17/18(94%) | 0/18(0%) | Plus/Plus | |

Features:
**145475 bp at 5' side: mineralocorticoid receptor isoform X2719841 bp at 3' side: uncharacterized protein LOC285423 isoform X3**

```
Query  281        TGGTGTTCTCTCCTTTAT  298
                  ||||| ||||||||||||
Sbjct  148582335  TGGTGATCTCTCCTTTAT  148582352
```

Range 37: 157414497 to 157414510

| Score | Expect | Identities | Gaps | Strand | Frame |
|---|---|---|---|---|---|
| 28.2 bits(14) | 256() | 14/14(100%) | 0/14(0%) | Plus/Plus | |

Features:
**51453 bp at 5' side: glutamate receptor 2 isoform 3712897 bp at 3' side: protein FAM198B isoform 2**

```
Query  281        TGGTGTTCTCTCCT  294
                  ||||||||||||||
Sbjct  157414497  TGGTGTTCTCTCCT  157414510
```

Range 38: 163092750 to 163092763

| Score | Expect | Identities | Gaps | Strand | Frame |
|---|---|---|---|---|---|
| 28.2 bits(14) | 256() | 14/14(100%) | 0/14(0%) | Plus/Plus | |

Features:
**981354 bp at 5' side: follistatin-related protein 5 isoform a precursor34216 bp at 3' side: H/ACA ribonucleoprotein complex non-core subunit NAF1 iso...**

```
Query  284        TGTTCTCTCCTTTA  297
                  ||||||||||||||
Sbjct  163092750  TGTTCTCTCCTTTA  163092763
```

Range 39: 164289114 to 164289127

| Score | Expect | Identities | Gaps | Strand | Frame |
|---|---|---|---|---|---|
| 28.2 bits(14) | 256() | 14/14(100%) | 0/14(0%) | Plus/Plus | |

Features:
**91403 bp at 5' side: acidic leucine-rich nuclear phosphoprotein 32 family memb...588205 bp at 3' side: apelin receptor early endogenous ligand precursor**

```
Query  277        AAGCTGGTGTTCTC  290
                  ||||||||||||||
Sbjct  164289114  AAGCTGGTGTTCTC  164289127
```

Range 40: 177046566 to 177046579

| Score | Expect | Identities | Gaps | Strand | Frame |
|---|---|---|---|---|---|
| 28.2 bits(14) | 256() | 14/14(100%) | 0/14(0%) | Plus/Plus | |

Features:
**254255 bp at 5' side: vascular endothelial growth factor C preproprotein263375 bp at 3' side: endonuclease 8-like 3**

```
Query  285        GTTCTCTCCTTTAT  298
                  ||||||||||||||
Sbjct  177046566  GTTCTCTCCTTTAT  177046579
```

Range 41: 181757601 to 181757614

| Score | Expect | Identities | Gaps | Strand | Frame |
|---|---|---|---|---|---|
| 28.2 bits(14) | 256() | 14/14(100%) | 0/14(0%) | Plus/Plus | |

Features:
**4160008 bp at 5' side: uncharacterized protein LOC285500 isoform X3566407 bp at 3' side: teneurin-3**

```
Query  273        TAACAAGCTGGTGT  286
                  ||||||||||||||
Sbjct  181757601  TAACAAGCTGGTGT  181757614
```

Range 42: 6078104 to 6078121

| Score | Expect | Identities | Gaps | Strand | Frame |
|---|---|---|---|---|---|
| 28.2 bits(14) | 256() | 17/18(94%) | 0/18(0%) | Plus/Minus | |

Features:
**janus kinase and microtubule-interacting protein 1 isoform 1janus kinase and microtubule-interacting protein 1 isofor...**

```
Query  279      GCTGGTGTTCTCTCCTTT  296
                |||||| |||||||||||
Sbjct  6078121  GCTGGTCTTCTCTCCTTT  6078104
```

Range 43: 8749848 to 8749861

| Score | Expect | Identities | Gaps | Strand | Frame |
|---|---|---|---|---|---|
| 28.2 bits(14) | 256() | 14/14(100%) | 0/14(0%) | Plus/Minus | |

Features:
**130231 bp at 5' side: carboxypeptidase Z isoform 396285 bp at 3' side: homeobox protein HMX1 isoform X1**

```
Query  278      AGCTGGTGTTCTCT  291
                ||||||||||||||
Sbjct  8749861  AGCTGGTGTTCTCT  8749848
```

Range 44: 21598114 to 21598127

| Score | Expect | Identities | Gaps | Strand | Frame |
|---|---|---|---|---|---|
| 28.2 bits(14) | 256() | 14/14(100%) | 0/14(0%) | Plus/Minus | |

Features:
**Kv channel-interacting protein 4 isoform 2Kv channel-interacting protein 4 isoform 1**

```
Query  283       GTGTTCTCTCCTTT  296
                 ||||||||||||||
Sbjct  21598127  GTGTTCTCTCCTTT  21598114
```

Range 45: 22638874 to 22638887

| Score | Expect | Identities | Gaps | Strand | Frame |
|---|---|---|---|---|---|
| 28.2 bits(14) | 256() | 14/14(100%) | 0/14(0%) | Plus/Minus | |

Features:
**123090 bp at 5' side: probable G-protein coupled receptor 125 isoform X31156935 bp at 3' side: peroxisome proliferator-activated receptor gamma coactiva...**

```
Query  280       CTGGTGTTCTCTCC  293
                 ||||||||||||||
Sbjct  22638887  CTGGTGTTCTCTCC  22638874
```

Range 46: 34323282 to 34323295

| Score | Expect | Identities | Gaps | Strand | Frame |
|---|---|---|---|---|---|

| | | | | |
|---|---|---|---|---|
| 28.2 bits(14) | 256() | 14/14(100%) | 0/14(0%) | Plus/Minus |

Features:
**3180433 bp at 5' side: protocadherin-7 isoform X11744612 bp at 3' side: arf-GAP with Rho-GAP domain, ANK repeat and PH domain-con...**

```
Query  273       TAACAAGCTGGTGT  286
                 ||||||||||||||
Sbjct  34323295  TAACAAGCTGGTGT  34323282
```

Range 47: 39167257 to 39167270

| Score | Expect | Identities | Gaps | Strand | Frame |
|---|---|---|---|---|---|
| 28.2 bits(14) | 256() | 14/14(100%) | 0/14(0%) | Plus/Minus | |

Features:
**46191 bp at 5' side: kelch-like protein 5 isoform 415288 bp at 3' side: WD repeat-containing protein 19 isoform X1**

```
Query  271       TGTAACAAGCTGGT  284
                 ||||||||||||||
Sbjct  39167270  TGTAACAAGCTGGT  39167257
```

Range 48: 56013952 to 56013965

| Score | Expect | Identities | Gaps | Strand | Frame |
|---|---|---|---|---|---|
| 28.2 bits(14) | 256() | 14/14(100%) | 0/14(0%) | Plus/Minus | |

Features:
**centrosomal protein of 135 kDa isoform X1centrosomal protein of 135 kDa**

```
Query  281       TGGTGTTCTCTCCT  294
                 ||||||||||||||
Sbjct  56013965  TGGTGTTCTCTCCT  56013952
```

Range 49: 62542193 to 62542206

| Score | Expect | Identities | Gaps | Strand | Frame |
|---|---|---|---|---|---|
| 28.2 bits(14) | 256() | 14/14(100%) | 0/14(0%) | Plus/Minus | |

Features:
**472024 bp at 5' side: latrophilin-3 isoform X101734807 bp at 3' side: trans-2,3-enoyl-CoA reductase-like isoform X3**

```
Query  286       TTCTCTCCTTTATT  299
                 ||||||||||||||
Sbjct  62542206  TTCTCTCCTTTATT  62542193
```

Range 50: 68187859 to 68187872

| Score | Expect | Identities | Gaps | Strand | Frame |
|---|---|---|---|---|---|
| 28.2 bits(14) | 256() | 14/14(100%) | 0/14(0%) | Plus/Minus | |

Features:
**58039 bp at 5' side: transmembrane protease serine 11F isoform X140039 bp at 3' side: transmembrane protease serine 11B**

```
Query  283       GTGTTCTCTCCTTT  296
                 ||||||||||||||
Sbjct  68187872  GTGTTCTCTCCTTT  68187859
```

Range 51: 69451050 to 69451063

| Score | Expect | Identities | Gaps | Strand | Frame |
|---|---|---|---|---|---|
| 28.2 bits(14) | 256() | 14/14(100%) | 0/14(0%) | Plus/Minus | |

Features:
**156501 bp at 5' side: UDP-glucuronosyltransferase 2B28 isoform X129571 bp at 3' side: UDP-glucuronosyltransferase 2B4 isoform X1**

```
Query  286       TTCTCTCCTTTATT  299
                 ||||||||||||||
Sbjct  69451063  TTCTCTCCTTTATT  69451050
```

Range 52: 99550533 to 99550546

| Score | Expect | Identities | Gaps | Strand | Frame |
|---|---|---|---|---|---|
| 28.2 bits(14) | 256() | 14/14(100%) | 0/14(0%) | Plus/Minus | |

Features:
**tRNA methyltransferase 10 homolog AtRNA methyltransferase 10 homolog A**

```
Query  286       TTCTCTCCTTTATT  299
                 ||||||||||||||
Sbjct  99550546  TTCTCTCCTTTATT  99550533
```

Range 53: 100585972 to 100585985

| Score | Expect | Identities | Gaps | Strand | Frame |
|---|---|---|---|---|---|
| 28.2 bits(14) | 256() | 14/14(100%) | 0/14(0%) | Plus/Minus | |

Features:
**68058 bp at 5' side: endomucin isoform X1439880 bp at 3' side: serine/threonine-protein phosphatase 2B catalytic subunit...**

```
Query  283        GTGTTCTCTCCTTT  296
                  ||||||||||||||
Sbjct  100585985  GTGTTCTCTCCTTT  100585972
```

Range 54: 110643181 to 110643194

| Score | Expect | Identities | Gaps | Strand | Frame |
|---|---|---|---|---|---|
| 28.2 bits(14) | 256() | 14/14(100%) | 0/14(0%) | Plus/Minus | |

Features:
**10183 bp at 5' side: pituitary homeobox 2 isoform b1183565 bp at 3' side: histone-lysine N-methyltransferase SETMAR-like**

```
Query  279        GCTGGTGTTCTCTC  292
                  ||||||||||||||
Sbjct  110643194  GCTGGTGTTCTCTC  110643181
```

Range 55: 111593276 to 111593289

| Score | Expect | Identities | Gaps | Strand | Frame |
|---|---|---|---|---|---|
| 28.2 bits(14) | 256() | 14/14(100%) | 0/14(0%) | Plus/Minus | |

Features:
**960278 bp at 5' side: pituitary homeobox 2 isoform b233470 bp at 3' side: histone-lysine N-methyltransferase SETMAR-like**

```
Query  286        TTCTCTCCTTTATT  299
                  ||||||||||||||
Sbjct  111593289  TTCTCTCCTTTATT  111593276
```

Range 56: 115457367 to 115457380

| Score | Expect | Identities | Gaps | Strand | Frame |
|---|---|---|---|---|---|
| 28.2 bits(14) | 256() | 14/14(100%) | 0/14(0%) | Plus/Minus | |

Features:
**380331 bp at 5' side: bifunctional heparan sulfate N-deacetylase/N-sulfotransfe...1626904 bp at 3' side: translocating chain-associated membrane protein 1-like 1**

```
Query  283        GTGTTCTCTCCTTT  296
                  ||||||||||||||
Sbjct  115457380  GTGTTCTCTCCTTT  115457367
```

Range 57: 127803065 to 127803078

| Score | Expect | Identities | Gaps | Strand | Frame |
|---|---|---|---|---|---|
| 28.2 bits(14) | 256() | 14/14(100%) | 0/14(0%) | Plus/Minus | |

Features:
**heat shock 70 kDa protein 4Lheat shock 70 kDa protein 4L isoform X2**

```
Query  271        TGTAACAAGCTGGT  284
```

```
                          ||||||||||||||
Sbjct  127803078  TGTAACAAGCTGGT  127803065
```

Range 58: 134559244 to 134559257

| Score | Expect | Identities | Gaps | Strand | Frame |
|---|---|---|---|---|---|
| 28.2 bits(14) | 256() | 14/14(100%) | 0/14(0%) | Plus/Minus | |

Features:
**358051 bp at 5' side: polyadenylate-binding protein 4-like2961772 bp at 3' side: protocadherin-18 isoform 2 precursor**

```
Query  286        TTCTCTCCTTTATT  299
                  ||||||||||||||
Sbjct  134559257  TTCTCTCCTTTATT  134559244
```

Range 59: 134694757 to 134694774

| Score | Expect | Identities | Gaps | Strand | Frame |
|---|---|---|---|---|---|
| 28.2 bits(14) | 256() | 17/18(94%) | 0/18(0%) | Plus/Minus | |

Features:
**493564 bp at 5' side: polyadenylate-binding protein 4-like2826255 bp at 3' side: protocadherin-18 isoform 2 precursor**

```
Query  279        GCTGGTGTTCTCTCCTTT  296
                  ||||||||||| ||||||
Sbjct  134694774  GCTGGTGTTCTTTCCTTT  134694757
```

Range 60: 158843513 to 158843526

| Score | Expect | Identities | Gaps | Strand | Frame |
|---|---|---|---|---|---|
| 28.2 bits(14) | 256() | 14/14(100%) | 0/14(0%) | Plus/Minus | |

Features:
**folliculin-interacting protein 2 isoform X2folliculin-interacting protein 2**

```
Query  287        TCTCTCCTTTATTG  300
                  ||||||||||||||
Sbjct  158843526  TCTCTCCTTTATTG  158843513
```

Range 61: 179535664 to 179535677

| Score | Expect | Identities | Gaps | Strand | Frame |
|---|---|---|---|---|---|
| 28.2 bits(14) | 256() | 14/14(100%) | 0/14(0%) | Plus/Minus | |

Features:
**1938071 bp at 5' side: uncharacterized protein LOC285500 isoform X32788344 bp at 3' side: teneurin-3**

```
Query  286        TTCTCTCCTTTATT  299
                  ||||||||||||||
Sbjct  179535677  TTCTCTCCTTTATT  179535664
```

Range 62: 182864589 to 182864602

| Score | Expect | Identities | Gaps | Strand | Frame |
|---|---|---|---|---|---|
| 28.2 bits(14) | 256() | 14/14(100%) | 0/14(0%) | Plus/Minus | |

Features:
**64238 bp at 5' side: teneurin-3 isoform X126797 bp at 3' side: deoxycytidylate deaminase isoform X3**

```
Query  286        TTCTCTCCTTTATT  299
                  ||||||||||||||
Sbjct  182864602  TTCTCTCCTTTATT  182864589
```

Range 63: 189794178 to 189794191

| Score | Expect | Identities | Gaps | Strand | Frame |
|---|---|---|---|---|---|
| 28.2 bits(14) | 256() | 14/14(100%) | 0/14(0%) | Plus/Minus | |

Features:
**1646806 bp at 5' side: probable E3 ubiquitin-protein ligase TRIML1 isoform X2146819 bp at 3' side: protein FRG1**

```
Query  282        GGTGTTCTCTCCTT  295
                  ||||||||||||||
Sbjct  189794191  GGTGTTCTCTCCTT  189794178
```

Homo sapiens chromosome 5, GRCh38.p2 Primary Assembly

Sequence ID: **ref|NC_000005.10|** Length: 181538259 Number of Matches: 75

Range 1: 125038294 to 125038314

| Score | Expect | Identities | Gaps | Strand | Frame |
|---|---|---|---|---|---|
| 34.2 bits(17) | 4.1() | 20/21(95%) | 0/21(0%) | Plus/Plus | |

Features:
**293305 bp at 5' side: zinc finger protein 608 isoform X31322018 bp at 3' side: GRAM domain-containing protein 3 isoform X4**

```
Query  277        AAGCTGGTGTTCTCTCCTTTA  297
                  ||||||||||| |||||||||
Sbjct  125038294  AAGCTGGTGTTTTCTCCTTTA  125038314
```

Range 2: 5606549 to 5606565

| Score | Expect | Identities | Gaps | Strand | Frame |
|---|---|---|---|---|---|
| 34.2 bits(17) | 4.1() | 17/17(100%) | 0/17(0%) | Plus/Minus | |

Features:
**131873 bp at 5' side: little elongation complex subunit 1 isoform X1765938 bp at 3' side: mediator of RNA polymerase II transcription subunit 10**

```
Query  284      TGTTCTCTCCTTTATTG  300
                |||||||||||||||||
Sbjct  5606565  TGTTCTCTCCTTTATTG  5606549
```

Range 3: 168136544 to 168136560

| Score | Expect | Identities | Gaps | Strand | Frame |
|---|---|---|---|---|---|
| 34.2 bits(17) | 4.1() | 17/17(100%) | 0/17(0%) | Plus/Minus | |

Features:
**teneurin-2 isoform X1teneurin-2**

```
Query  281        TGGTGTTCTCTCCTTTA  297
                  |||||||||||||||||
Sbjct  168136560  TGGTGTTCTCTCCTTTA  168136544
```

Range 4: 96788742 to 96788757

| Score | Expect | Identities | Gaps | Strand | Frame |
|---|---|---|---|---|---|
| 32.2 bits(16) | 16() | 16/16(100%) | 0/16(0%) | Plus/Minus | |

Features:
**endoplasmic reticulum aminopeptidase 1 isoform a precursorendoplasmic reticulum aminopeptidase 1 isoform X2**

```
Query  277       AAGCTGGTGTTCTCTC  292
                 ||||||||||||||||
Sbjct  96788757  AAGCTGGTGTTCTCTC  96788742
```

Range 5: 11468838 to 11468852

| Score | Expect | Identities | Gaps | Strand | Frame |
|---|---|---|---|---|---|
| 30.2 bits(15) | 65() | 15/15(100%) | 0/15(0%) | Plus/Plus | |

Features:
**catenin delta-2 isoform X3catenin delta-2 isoform X2**

```
Query  282       GGTGTTCTCTCCTTT  296
                 |||||||||||||||
Sbjct  11468838  GGTGTTCTCTCCTTT  11468852
```

Range 6: 78575708 to 78575722

| Score | Expect | Identities | Gaps | Strand | Frame |
|---|---|---|---|---|---|
| 30.2 bits(15) | 65() | 15/15(100%) | 0/15(0%) | Plus/Plus | |

Features:
**65495 bp at 5' side: lipoma HMGIC fusion partner-like 2 protein204675 bp at 3' side: arylsulfatase B isoform X1**

```
Query  285       GTTCTCTCCTTTATT  299
                 |||||||||||||||
Sbjct  78575708  GTTCTCTCCTTTATT  78575722
```

Range 7: 91601888 to 91601902

| Score | Expect | Identities | Gaps | Strand | Frame |
|---|---|---|---|---|---|
| 30.2 bits(15) | 65() | 15/15(100%) | 0/15(0%) | Plus/Plus | |

Features:
**218796 bp at 5' side: arrestin domain-containing protein 31983122 bp at 3' side: COUP transcription factor 1**

```
Query  279       GCTGGTGTTCTCTCC  293
                 |||||||||||||||
Sbjct  91601888  GCTGGTGTTCTCTCC  91601902
```

Range 8: 101909103 to 101909117

| Score | Expect | Identities | Gaps | Strand | Frame |
|---|---|---|---|---|---|
| 30.2 bits(15) | 65() | 15/15(100%) | 0/15(0%) | Plus/Plus | |

Features:
**1006148 bp at 5' side: CMP-N-acetylneuraminate-poly-alpha-2,8-sialyltransferase ...327741 bp at 3' side: solute carrier organic anion transporter family member 4C...**

```
Query  284        TGTTCTCTCCTTTAT  298
                  |||||||||||||||
Sbjct  101909103  TGTTCTCTCCTTTAT  101909117
```

Range 9: 118548069 to 118548083

| Score | Expect | Identities | Gaps | Strand | Frame |
|---|---|---|---|---|---|
| 30.2 bits(15) | 65() | 15/15(100%) | 0/15(0%) | Plus/Plus | |

Features:
**2043125 bp at 5' side: semaphorin-6A isoform 1 precursor292834 bp at 3' side: DTW domain-containing protein 2**

```
Query  278        AGCTGGTGTTCTCTC  292
                  |||||||||||||||
Sbjct  118548069  AGCTGGTGTTCTCTC  118548083
```

Range 10: 122568332 to 122568346

| Score | Expect | Identities | Gaps | Strand | Frame |
|---|---|---|---|---|---|
| 30.2 bits(15) | 65() | 15/15(100%) | 0/15(0%) | Plus/Plus | |

Features:
**104757 bp at 5' side: synphilin-1 isoform X8206758 bp at 3' side: sorting nexin-2 isoform 1**

```
Query  284        TGTTCTCTCCTTTAT  298
                  |||||||||||||||
Sbjct  122568332  TGTTCTCTCCTTTAT  122568346
```

Range 11: 123908952 to 123908970

| Score | Expect | Identities | Gaps | Strand | Frame |
|---|---|---|---|---|---|
| 30.2 bits(15) | 65() | 18/19(95%) | 0/19(0%) | Plus/Plus | |

Features:
**294556 bp at 5' side: casein kinase I isoform gamma-3 isoform 7728930 bp at 3' side: zinc finger protein 608 isoform X7**

```
Query  282        GGTGTTCTCTCCTTTATTG  300
                  |||||| ||||||||||||
Sbjct  123908952  GGTGTTTTCTCCTTTATTG  123908970
```

Range 12: 133195879 to 133195893

| Score | Expect | Identities | Gaps | Strand | Frame |
|---|---|---|---|---|---|
| 30.2 bits(15) | 65() | 15/15(100%) | 0/15(0%) | Plus/Plus | |

Features:
**91443 bp at 5' side: heat shock 70 kDa protein 43202 bp at 3' side: follistatin-related protein 4 isoform X2**

```
Query  285        GTTCTCTCCTTTATT  299
                  |||||||||||||||
Sbjct  133195879  GTTCTCTCCTTTATT  133195893
```

Range 13: 151849956 to 151849970

| Score | Expect | Identities | Gaps | Strand | Frame |
|---|---|---|---|---|---|
| 30.2 bits(15) | 65() | 15/15(100%) | 0/15(0%) | Plus/Plus | |

Features:
**glycine receptor subunit alpha-1 isoform 2 precursorglycine receptor subunit alpha-1 isoform 1 precursor**

```
Query  275        ACAAGCTGGTGTTCT  289
                  |||||||||||||||
Sbjct  151849956  ACAAGCTGGTGTTCT  151849970
```

Range 14: 158377697 to 158377711

| Score | Expect | Identities | Gaps | Strand | Frame |
|---|---|---|---|---|---|
| 30.2 bits(15) | 65() | 15/15(100%) | 0/15(0%) | Plus/Plus | |

Features:
**518727 bp at 5' side: clathrin interactor 1 isoform X3321400 bp at 3' side: transcription factor COE1 isoform 2**

```
Query  286        TTCTCTCCTTTATTG  300
                  |||||||||||||||
Sbjct  158377697  TTCTCTCCTTTATTG  158377711
```

Range 15: 5107789 to 5107803

| Score | Expect | Identities | Gaps | Strand | Frame |
|---|---|---|---|---|---|
| 30.2 bits(15) | 65() | 15/15(100%) | 0/15(0%) | Plus/Minus | |

Features:
**1506749 bp at 5' side: iroquois-class homeodomain protein IRX-132665 bp at 3' side: A disintegrin and metalloproteinase with thrombospondin m...**

```
Query  280      CTGGTGTTCTCTCCT  294
                |||||||||||||||
Sbjct  5107803  CTGGTGTTCTCTCCT  5107789
```

Range 16: 21998274 to 21998288

| Score | Expect | Identities | Gaps | Strand | Frame |
|---|---|---|---|---|---|
| 30.2 bits(15) | 65() | 15/15(100%) | 0/15(0%) | Plus/Minus | |

Features:
**cadherin-12 preproproteincadherin-12 isoform X1**

```
Query  286       TTCTCTCCTTTATTG  300
                 |||||||||||||||
Sbjct  21998288  TTCTCTCCTTTATTG  21998274
```

Range 17: 39821399 to 39821413

| Score | Expect | Identities | Gaps | Strand | Frame |
|---|---|---|---|---|---|

| | | | | |
|---|---|---|---|---|
| 30.2 bits(15) | 65() | 15/15(100%) | 0/15(0%) | Plus/Minus |

Features:
**427079 bp at 5' side: disabled homolog 2 isoform 1859581 bp at 3' side: prostaglandin E2 receptor EP4 subtype isoform X3**

```
Query  285       GTTCTCTCCTTTATT  299
                 |||||||||||||||
Sbjct  39821413  GTTCTCTCCTTTATT  39821399
```

Range 18: 70008067 to 70008081

| Score | Expect | Identities | Gaps | Strand | Frame |
|---|---|---|---|---|---|
| 30.2 bits(15) | 65() | 15/15(100%) | 0/15(0%) | Plus/Minus | |

Features:
**415869 bp at 5' side: general transcription factor IIH subunit 2-like protein i...17371 bp at 3' side: small EDRK-rich factor 1 isoform 1**

```
Query  286       TTCTCTCCTTTATTG  300
                 |||||||||||||||
Sbjct  70008081  TTCTCTCCTTTATTG  70008067
```

Range 19: 70883145 to 70883159

| Score | Expect | Identities | Gaps | Strand | Frame |
|---|---|---|---|---|---|
| 30.2 bits(15) | 65() | 15/15(100%) | 0/15(0%) | Plus/Minus | |

Features:
**104333 bp at 5' side: LOW QUALITY PROTEIN: putative POM121-like protein 1-like ...17711 bp at 3' side: small EDRK-rich factor 1 isoform 1**

```
Query  286       TTCTCTCCTTTATTG  300
                 |||||||||||||||
Sbjct  70883159  TTCTCTCCTTTATTG  70883145
```

Range 20: 83388258 to 83388272

| Score | Expect | Identities | Gaps | Strand | Frame |
|---|---|---|---|---|---|
| 30.2 bits(15) | 65() | 15/15(100%) | 0/15(0%) | Plus/Minus | |

Features:
**35016 bp at 5' side: DNA repair protein XRCC4 isoform X495247 bp at 3' side: versican core protein isoform 1 precursor**

```
Query  284       TGTTCTCTCCTTTAT  298
                 |||||||||||||||
Sbjct  83388272  TGTTCTCTCCTTTAT  83388258
```

Range 21: 84814880 to 84814894

| Score | Expect | Identities | Gaps | Strand | Frame |
|---|---|---|---|---|---|
| 30.2 bits(15) | 65() | 15/15(100%) | 0/15(0%) | Plus/Minus | |

Features:
**430506 bp at 5' side: EGF-like repeat and discoidin I-like domain-containing pr...1803162 bp at 3' side: cytochrome c oxidase subunit 7C, mitochondrial precursor**

```
Query  286       TTCTCTCCTTTATTG  300
                 |||||||||||||||
Sbjct  84814894  TTCTCTCCTTTATTG  84814880
```

Range 22: 100482855 to 100482869

| Score | Expect | Identities | Gaps | Strand | Frame |
|---|---|---|---|---|---|
| 30.2 bits(15) | 65() | 15/15(100%) | 0/15(0%) | Plus/Minus | |

Features:
**89960 bp at 5' side: LOW QUALITY PROTEIN: putative POM121-like protein 1-like ...52662 bp at 3' side: membrane protein FAM174A precursor**

```
Query  286        TTCTCTCCTTTATTG  300
                  |||||||||||||||
Sbjct  100482869  TTCTCTCCTTTATTG  100482855
```

Range 23: 115802045 to 115802059

| Score | Expect | Identities | Gaps | Strand | Frame |
|---|---|---|---|---|---|
| 30.2 bits(15) | 65() | 15/15(100%) | 0/15(0%) | Plus/Minus | |

Features:
**176253 bp at 5' side: transmembrane emp24 domain-containing protein 7 precursor3374 bp at 3' side: cysteine dioxygenase type 1**

```
Query  286        TTCTCTCCTTTATTG  300
                  |||||||||||||||
Sbjct  115802059  TTCTCTCCTTTATTG  115802045
```

Range 24: 3680829 to 3680842

| Score | Expect | Identities | Gaps | Strand | Frame |
|---|---|---|---|---|---|
| 28.2 bits(14) | 256() | 14/14(100%) | 0/14(0%) | Plus/Plus | |

Features:
**79789 bp at 5' side: iroquois-class homeodomain protein IRX-11459626 bp at 3' side: A disintegrin and metalloproteinase with thrombospondin m...**

```
Query  286      TTCTCTCCTTTATT  299
                ||||||||||||||
Sbjct  3680829  TTCTCTCCTTTATT  3680842
```

Range 25: 6091043 to 6091056

| Score | Expect | Identities | Gaps | Strand | Frame |
|---|---|---|---|---|---|
| 28.2 bits(14) | 256() | 14/14(100%) | 0/14(0%) | Plus/Plus | |

Features:
**616367 bp at 5' side: little elongation complex subunit 1 isoform X1281447 bp at 3' side: mediator of RNA polymerase II transcription subunit 10**

```
Query  286      TTCTCTCCTTTATT  299
                ||||||||||||||
Sbjct  6091043  TTCTCTCCTTTATT  6091056
```

Range 26: 11492288 to 11492305

| Score | Expect | Identities | Gaps | Strand | Frame |
|---|---|---|---|---|---|
| 28.2 bits(14) | 256() | 17/18(94%) | 0/18(0%) | Plus/Plus | |

Features:
**catenin delta-2 isoform X3catenin delta-2 isoform X2**

```
Query  280       CTGGTGTTCTCTCCTTTA  297
                 ||||||||||| ||||||
Sbjct  11492288  CTGGTGTTCTCGCCTTTA  11492305
```

Range 27: 13278181 to 13278194

| Score | Expect | Identities | Gaps | Strand | Frame |
|---|---|---|---|---|---|
| 28.2 bits(14) | 256() | 14/14(100%) | 0/14(0%) | Plus/Plus | |

Features:
**1374328 bp at 5' side: catenin delta-2 isoform 1413790 bp at 3' side: dynein heavy chain 5, axonemal isoform X1**

```
Query  286       TTCTCTCCTTTATT  299
                 ||||||||||||||
Sbjct  13278181  TTCTCTCCTTTATT  13278194
```

Range 28: 13997135 to 13997148

| Score | Expect | Identities | Gaps | Strand | Frame |
|---|---|---|---|---|---|
| 28.2 bits(14) | 256() | 14/14(100%) | 0/14(0%) | Plus/Plus | |

Features:
**dynein heavy chain 5, axonemal isoform X1**

```
Query  281       TGGTGTTCTCTCCT  294
                 ||||||||||||||
Sbjct  13997135  TGGTGTTCTCTCCT  13997148
```

Range 29: 14639291 to 14639304

| Score | Expect | Identities | Gaps | Strand | Frame |
|---|---|---|---|---|---|
| 28.2 bits(14) | 256() | 14/14(100%) | 0/14(0%) | Plus/Plus | |

Features:
**28977 bp at 5' side: inactive ubiquitin thioesterase FAM105A isoform X125522 bp at 3' side: ubiquitin thioesterase otulin**

```
Query  283       GTGTTCTCTCCTTT  296
                 ||||||||||||||
Sbjct  14639291  GTGTTCTCTCCTTT  14639304
```

Range 30: 22195788 to 22195801

| Score | Expect | Identities | Gaps | Strand | Frame |
|---|---|---|---|---|---|
| 28.2 bits(14) | 256() | 14/14(100%) | 0/14(0%) | Plus/Plus | |

Features:
**117112 bp at 5' side: cadherin-12 isoform X11313233 bp at 3' side: histone-lysine N-methyltransferase PRDM9**

```
Query  286       TTCTCTCCTTTATT  299
                 ||||||||||||||
Sbjct  22195788  TTCTCTCCTTTATT  22195801
```

Range 31: 39645803 to 39645820

| Score | Expect | Identities | Gaps | Strand | Frame |
|---|---|---|---|---|---|
| 28.2 bits(14) | 256() | 17/18(94%) | 0/18(0%) | Plus/Plus | |

Features:
**251483 bp at 5' side: disabled homolog 2 isoform 11035174 bp at 3' side: prostaglandin E2 receptor EP4 subtype isoform X3**

```
Query  281       TGGTGTTCTCTCCTTTAT  298
                 ||||||| ||||||||||
Sbjct  39645803  TGGTGTTTTCTCCTTTAT  39645820
```

Range 32: 44909512 to 44909525

| Score | Expect | Identities | Gaps | Strand | Frame |
|---|---|---|---|---|---|
| 28.2 bits(14) | 256() | 14/14(100%) | 0/14(0%) | Plus/Plus | |

Features:
**94310 bp at 5' side: 28S ribosomal protein S30, mitochondrial352396 bp at 3' side: potassium/sodium hyperpolarization-activated cyclic nucle...**

```
Query  286       TTCTCTCCTTTATT  299
                 ||||||||||||||
Sbjct  44909512  TTCTCTCCTTTATT  44909525
```

Range 33: 56494278 to 56494291

| Score | Expect | Identities | Gaps | Strand | Frame |
|---|---|---|---|---|---|
| 28.2 bits(14) | 256() | 14/14(100%) | 0/14(0%) | Plus/Plus | |

Features:
**367236 bp at 5' side: ankyrin repeat domain-containing protein 55 isoform X217431 bp at 3' side: uncharacterized protein LOC101928448**

```
Query  278       AGCTGGTGTTCTCT  291
                 ||||||||||||||
Sbjct  56494278  AGCTGGTGTTCTCT  56494291
```

Range 34: 59504671 to 59504684

| Score | Expect | Identities | Gaps | Strand | Frame |
|---|---|---|---|---|---|
| 28.2 bits(14) | 256() | 14/14(100%) | 0/14(0%) | Plus/Plus | |

Features:
**cAMP-specific 3',5'-cyclic phosphodiesterase 4D isoform X6cAMP-specific 3',5'-cyclic phosphodiesterase 4D isoform X1**

```
Query  283       GTGTTCTCTCCTTT  296
                 ||||||||||||||
Sbjct  59504671  GTGTTCTCTCCTTT  59504684
```

Range 35: 66482623 to 66482636

| Score | Expect | Identities | Gaps | Strand | Frame |
|---|---|---|---|---|---|
| 28.2 bits(14) | 256() | 14/14(100%) | 0/14(0%) | Plus/Plus | |

Features:
**303755 bp at 5' side: splicing regulatory glutamine/lysine-rich protein 1 isofo...114020 bp at 3' side: microtubule-associated serine/threonine-protein kinase 4 ...**

```
Query  285       GTTCTCTCCTTTAT  298
                 ||||||||||||||
Sbjct  66482623  GTTCTCTCCTTTAT  66482636
```

Range 36: 67063537 to 67063550

| Score | Expect | Identities | Gaps | Strand | Frame |
|---|---|---|---|---|---|
| 28.2 bits(14) | 256() | 14/14(100%) | 0/14(0%) | Plus/Plus | |

Features:
**microtubule-associated serine/threonine-protein kinase 4 ...microtubule-associated serine/threonine-protein kinase 4 ...**

```
Query  286       TTCTCTCCTTTATT  299
                 ||||||||||||||
Sbjct  67063537  TTCTCTCCTTTATT  67063550
```

Range 37: 68787327 to 68787340

| Score | Expect | Identities | Gaps | Strand | Frame |
|---|---|---|---|---|---|
| 28.2 bits(14) | 256() | 14/14(100%) | 0/14(0%) | Plus/Plus | |

Features:
**489726 bp at 5' side: phosphatidylinositol 3-kinase regulatory subunit alpha is...306916 bp at 3' side: zinc transporter 5 isoform 1**

```
Query  286       TTCTCTCCTTTATT  299
                 ||||||||||||||
Sbjct  68787327  TTCTCTCCTTTATT  68787340
```

Range 38: 71428883 to 71428896

| Score | Expect | Identities | Gaps | Strand | Frame |
|---|---|---|---|---|---|
| 28.2 bits(14) | 256() | 14/14(100%) | 0/14(0%) | Plus/Plus | |

Features:
**366120 bp at 5' side: general transcription factor IIH subunit 226982 bp at 3' side: transcription factor TFIIIB component B'' homolog**

```
Query  286       TTCTCTCCTTTATT  299
                 ||||||||||||||
Sbjct  71428883  TTCTCTCCTTTATT  71428896
```

Range 39: 73505816 to 73505833

| Score | Expect | Identities | Gaps | Strand | Frame |
|---|---|---|---|---|---|
| 28.2 bits(14) | 256() | 17/18(94%) | 0/18(0%) | Plus/Plus | |

Features:
**578 bp at 5' side: transcription factor BTF3 isoform B46964 bp at 3' side: ankyrin repeat family A protein 2**

```
Query  282       GGTGTTCTCTCCTTTATT  299
                 |||||| |||||||||||
Sbjct  73505816  GGTGTTATCTCCTTTATT  73505833
```

Range 40: 74961859 to 74961872

| Score | Expect | Identities | Gaps | Strand | Frame |
|---|---|---|---|---|---|
| 28.2 bits(14) | 256() | 14/14(100%) | 0/14(0%) | Plus/Plus | |

Features:
**120183 bp at 5' side: soluble lamin-associated protein of 75 kDa isoform X166804 bp at 3' side: beta-1,3-galactosyl-O-glycosyl-glycoprotein beta-1,6-N-ac...**

```
Query  286       TTCTCTCCTTTATT  299
                 ||||||||||||||
Sbjct  74961859  TTCTCTCCTTTATT  74961872
```

Range 41: 89749267 to 89749280

| Score | Expect | Identities | Gaps | Strand | Frame |
|---|---|---|---|---|---|
| 28.2 bits(14) | 256() | 14/14(100%) | 0/14(0%) | Plus/Plus | |

Features:
**925479 bp at 5' side: myocyte-specific enhancer factor 2C isoform 3644784 bp at 3' side: centrin-3 isoform 3**

```
Query  283       GTGTTCTCTCCTTT  296
                 ||||||||||||||
Sbjct  89749267  GTGTTCTCTCCTTT  89749280
```

Range 42: 123390500 to 123390513

| Score | Expect | Identities | Gaps | Strand | Frame |
|---|---|---|---|---|---|
| 28.2 bits(14) | 256() | 14/14(100%) | 0/14(0%) | Plus/Plus | |

Features:
**centrosomal protein of 120 kDa isoform X2centrosomal protein of 120 kDa isoform 1**

```
Query  286        TTCTCTCCTTTATT  299
                  ||||||||||||||
Sbjct  123390500  TTCTCTCCTTTATT  123390513
```

Range 43: 125515626 to 125515639

| Score | Expect | Identities | Gaps | Strand | Frame |
|---|---|---|---|---|---|
| 28.2 bits(14) | 256() | 14/14(100%) | 0/14(0%) | Plus/Plus | |

Features:
**770637 bp at 5' side: zinc finger protein 608 isoform X3844693 bp at 3' side: GRAM domain-containing protein 3 isoform X4**

```
Query  286        TTCTCTCCTTTATT  299
                  ||||||||||||||
Sbjct  125515626  TTCTCTCCTTTATT  125515639
```

Range 44: 129486980 to 129486993

| Score | Expect | Identities | Gaps | Strand | Frame |
|---|---|---|---|---|---|
| 28.2 bits(14) | 256() | 14/14(100%) | 0/14(0%) | Plus/Plus | |

Features:
**A disintegrin and metalloproteinase with thrombospondin m...A disintegrin and metalloproteinase with thrombospondin m...**

```
Query  286        TTCTCTCCTTTATT  299
                  ||||||||||||||
Sbjct  129486980  TTCTCTCCTTTATT  129486993
```

Range 45: 131317282 to 131317295

| Score | Expect | Identities | Gaps | Strand | Frame |
|---|---|---|---|---|---|

| | | | | |
|---|---|---|---|---|
| 28.2 bits(14) | 256() | 14/14(100%) | 0/14(0%) | Plus/Plus |

Features:
**117722 bp at 5' side: complex III assembly factor LYRM7 isoform 242199 bp at 3' side: CDC42 small effector protein 2**

```
Query  285        GTTCTCTCCTTTAT  298
                  ||||||||||||||
Sbjct  131317282  GTTCTCTCCTTTAT  131317295
```

Range 46: 137590091 to 137590104

| Score | Expect | Identities | Gaps | Strand | Frame |
|---|---|---|---|---|---|
| 28.2 bits(14) | 256() | 14/14(100%) | 0/14(0%) | Plus/Plus | |

Features:
**91533 bp at 5' side: testican-1 precursor31994 bp at 3' side: kelch-like protein 3 isoform 1**

```
Query  286        TTCTCTCCTTTATT  299
                  ||||||||||||||
Sbjct  137590091  TTCTCTCCTTTATT  137590104
```

Range 47: 142766136 to 142766149

| Score | Expect | Identities | Gaps | Strand | Frame |
|---|---|---|---|---|---|
| 28.2 bits(14) | 256() | 14/14(100%) | 0/14(0%) | Plus/Plus | |

Features:
**152009 bp at 5' side: fibroblast growth factor 1 isoform 2 precursor4613 bp at 3' side: rho GTPase-activating protein 26 isoform a**

```
Query  275        ACAAGCTGGTGTTC  288
                  ||||||||||||||
Sbjct  142766136  ACAAGCTGGTGTTC  142766149
```

Range 48: 147478620 to 147478633

| Score | Expect | Identities | Gaps | Strand | Frame |
|---|---|---|---|---|---|
| 28.2 bits(14) | 256() | 14/14(100%) | 0/14(0%) | Plus/Plus | |

Features:
**dihydropyrimidinase-related protein 3 isoform 1dihydropyrimidinase-related protein 3 isoform X2**

```
Query  287        TCTCTCCTTTATTG  300
                  ||||||||||||||
Sbjct  147478620  TCTCTCCTTTATTG  147478633
```

Range 49: 151070407 to 151070420

| Score | Expect | Identities | Gaps | Strand | Frame |
|---|---|---|---|---|---|
| 28.2 bits(14) | 256() | 14/14(100%) | 0/14(0%) | Plus/Plus | |

Features:
**5312 bp at 5' side: TNFAIP3-interacting protein 1 isoform X131028 bp at 3' side: annexin A6 isoform 2**

```
Query  284        TGTTCTCTCCTTTA  297
                  ||||||||||||||
Sbjct  151070407  TGTTCTCTCCTTTA  151070420
```

Range 50: 152123503 to 152123516

| Score | Expect | Identities | Gaps | Strand | Frame |
|---|---|---|---|---|---|
| 28.2 bits(14) | 256() | 14/14(100%) | 0/14(0%) | Plus/Plus | |

Features:
**198954 bp at 5' side: glycine receptor subunit alpha-1 isoform X1268675 bp at 3' side: neuromedin-U receptor 2 isoform X1**

```
Query  283        GTGTTCTCTCCTTT  296
                  ||||||||||||||
Sbjct  152123503  GTGTTCTCTCCTTT  152123516
```

Range 51: 156105781 to 156105794

| Score | Expect | Identities | Gaps | Strand | Frame |
|---|---|---|---|---|---|
| 28.2 bits(14) | 256() | 14/14(100%) | 0/14(0%) | Plus/Plus | |

Features:
**1088217 bp at 5' side: chromosome-associated kinesin KIF4B223783 bp at 3' side: delta-sarcoglycan isoform 2**

```
Query  284        TGTTCTCTCCTTTA  297
                  ||||||||||||||
Sbjct  156105781  TGTTCTCTCCTTTA  156105794
```

Range 52: 174985008 to 174985021

| Score | Expect | Identities | Gaps | Strand | Frame |
|---|---|---|---|---|---|
| 28.2 bits(14) | 256() | 14/14(100%) | 0/14(0%) | Plus/Plus | |

Features:
**255425 bp at 5' side: homeobox protein MSX-2456738 bp at 3' side: D(1A) dopamine receptor**

```
Query  283        GTGTTCTCTCCTTT  296
                  ||||||||||||||
Sbjct  174985008  GTGTTCTCTCCTTT  174985021
```

Range 53: 181080601 to 181080614

| Score | Expect | Identities | Gaps | Strand | Frame |
|---|---|---|---|---|---|
| 28.2 bits(14) | 256() | 14/14(100%) | 0/14(0%) | Plus/Plus | |

Features:
**20739 bp at 5' side: butyrophilin-like protein 9 isoform X543743 bp at 3' side: olfactory receptor 2V1**

```
Query  282        GGTGTTCTCTCCTT  295
                  ||||||||||||||
Sbjct  181080601  GGTGTTCTCTCCTT  181080614
```

Range 54: 3329357 to 3329370

| Score | Expect | Identities | Gaps | Strand | Frame |
|---|---|---|---|---|---|
| 28.2 bits(14) | 256() | 14/14(100%) | 0/14(0%) | Plus/Minus | |

Features:
**403634 bp at 5' side: uncharacterized protein LOC105374620 isoform X6266736 bp at 3' side: iroquois-class homeodomain protein IRX-1**

```
Query  283      GTGTTCTCTCCTTT  296
                ||||||||||||||
Sbjct  3329370  GTGTTCTCTCCTTT  3329357
```

Range 55: 9198655 to 9198676

| Score | Expect | Identities | Gaps | Strand | Frame |
|---|---|---|---|---|---|
| 28.2 bits(14) | 256() | 20/22(91%) | 0/22(0%) | Plus/Minus | |

Features:
**semaphorin-5A isoform X2semaphorin-5A isoform X1**

```
Query  278      AGCTGGTGTTCTCTCCTTTATT  299
                ||||||||||| | ||||||||
Sbjct  9198676  AGCTGGTGTTCGCCCCTTTATT  9198655
```

Range 56: 10061397 to 10061410

| Score | Expect | Identities | Gaps | Strand | Frame |
|---|---|---|---|---|---|
| 28.2 bits(14) | 256() | 14/14(100%) | 0/14(0%) | Plus/Minus | |

Features:
**431365 bp at 5' side: taste receptor type 2 member 1166031 bp at 3' side: protein FAM173B isoform 2**

```
Query  286       TTCTCTCCTTTATT  299
```

```
                    ||||||||||||||
Sbjct  10061410  TTCTCTCCTTTATT  10061397
```

Range 57: 13376153 to 13376166

| Score | Expect | Identities | Gaps | Strand | Frame |
|---|---|---|---|---|---|
| 28.2 bits(14) | 256() | 14/14(100%) | 0/14(0%) | Plus/Minus | |

Features:
**1472300 bp at 5' side: catenin delta-2 isoform 1315818 bp at 3' side: dynein heavy chain 5, axonemal isoform X1**

```
Query  285       GTTCTCTCCTTTAT  298
                 ||||||||||||||
Sbjct  13376166  GTTCTCTCCTTTAT  13376153
```

Range 58: 20292360 to 20292373

| Score | Expect | Identities | Gaps | Strand | Frame |
|---|---|---|---|---|---|
| 28.2 bits(14) | 256() | 14/14(100%) | 0/14(0%) | Plus/Minus | |

Features:
**453374 bp at 5' side: cadherin-18 isoform X21188951 bp at 3' side: LOW QUALITY PROTEIN: putative POM121-like protein 1-like ...**

```
Query  286       TTCTCTCCTTTATT  299
                 ||||||||||||||
Sbjct  20292373  TTCTCTCCTTTATT  20292360
```

Range 59: 27285960 to 27285973

| Score | Expect | Identities | Gaps | Strand | Frame |
|---|---|---|---|---|---|
| 28.2 bits(14) | 256() | 14/14(100%) | 0/14(0%) | Plus/Minus | |

Features:
**297627 bp at 5' side: cadherin-9 isoform X11523075 bp at 3' side: uncharacterized protein LOC105374700**

```
Query  285       GTTCTCTCCTTTAT  298
                 ||||||||||||||
Sbjct  27285973  GTTCTCTCCTTTAT  27285960
```

Range 60: 44634805 to 44634818

| Score | Expect | Identities | Gaps | Strand | Frame |
|---|---|---|---|---|---|
| 28.2 bits(14) | 256() | 14/14(100%) | 0/14(0%) | Plus/Minus | |

Features:
**246123 bp at 5' side: fibroblast growth factor 10 isoform X1174145 bp at 3' side: 28S ribosomal protein S30, mitochondrial**

```
Query  286       TTCTCTCCTTTATT  299
                 ||||||||||||||
Sbjct  44634818  TTCTCTCCTTTATT  44634805
```

Range 61: 45738093 to 45738106

| Score | Expect | Identities | Gaps | Strand | Frame |
|---|---|---|---|---|---|
| 28.2 bits(14) | 256() | 14/14(100%) | 0/14(0%) | Plus/Minus | |

Features:
**42000 bp at 5' side: potassium/sodium hyperpolarization-activated cyclic nucle...4661167 bp at 3' side: embigin isoform X1**

```
Query  286       TTCTCTCCTTTATT  299
                 ||||||||||||||
Sbjct  45738106  TTCTCTCCTTTATT  45738093
```

Range 62: 86857649 to 86857662

| Score | Expect | Identities | Gaps | Strand | Frame |
|---|---|---|---|---|---|
| 28.2 bits(14) | 256() | 14/14(100%) | 0/14(0%) | Plus/Minus | |

Features:
**238180 bp at 5' side: cytochrome c oxidase subunit 7C, mitochondrial precursor410790 bp at 3' side: ras GTPase-activating protein 1 isoform 1**

```
Query  287       TCTCTCCTTTATTG  300
                 ||||||||||||||
Sbjct  86857662  TCTCTCCTTTATTG  86857649
```

Range 63: 99553617 to 99553630

| Score | Expect | Identities | Gaps | Strand | Frame |
|---|---|---|---|---|---|
| 28.2 bits(14) | 256() | 14/14(100%) | 0/14(0%) | Plus/Minus | |

Features:
**28256 bp at 5' side: putative POM121-like protein 1-like isoform X2835345 bp at 3' side: LOW QUALITY PROTEIN: putative POM121-like protein 1-like ...**

```
Query  284       TGTTCTCTCCTTTA  297
                 ||||||||||||||
Sbjct  99553630  TGTTCTCTCCTTTA  99553617
```

Range 64: 109875687 to 109875700

| Score | Expect | Identities | Gaps | Strand | Frame |
|---|---|---|---|---|---|
| 28.2 bits(14) | 256() | 14/14(100%) | 0/14(0%) | Plus/Minus | |

Features:
**8689 bp at 5' side: alpha-mannosidase 2 isoform X1538920 bp at 3' side: transmembrane protein 232 isoform X11**

```
Query  280        CTGGTGTTCTCTCC  293
                  ||||||||||||||
Sbjct  109875700  CTGGTGTTCTCTCC  109875687
```

Range 65: 117005240 to 117005253

| Score | Expect | Identities | Gaps | Strand | Frame |
|---|---|---|---|---|---|
| 28.2 bits(14) | 256() | 14/14(100%) | 0/14(0%) | Plus/Minus | |

Features:
**500296 bp at 5' side: semaphorin-6A isoform 1 precursor1835664 bp at 3' side: DTW domain-containing protein 2**

```
Query  286        TTCTCTCCTTTATT  299
                  ||||||||||||||
Sbjct  117005253  TTCTCTCCTTTATT  117005240
```

Range 66: 124105383 to 124105396

| Score | Expect | Identities | Gaps | Strand | Frame |
|---|---|---|---|---|---|
| 28.2 bits(14) | 256() | 14/14(100%) | 0/14(0%) | Plus/Minus | |

Features:
**490987 bp at 5' side: casein kinase I isoform gamma-3 isoform 7532504 bp at 3' side: zinc finger protein 608 isoform X7**

```
Query  287        TCTCTCCTTTATTG  300
                  ||||||||||||||
Sbjct  124105396  TCTCTCCTTTATTG  124105383
```

Range 67: 124544918 to 124544931

| Score | Expect | Identities | Gaps | Strand | Frame |
|---|---|---|---|---|---|
| 28.2 bits(14) | 256() | 14/14(100%) | 0/14(0%) | Plus/Minus | |

Features:
**930522 bp at 5' side: casein kinase I isoform gamma-3 isoform 792969 bp at 3' side: zinc finger protein 608 isoform X7**

```
Query  280        CTGGTGTTCTCTCC  293
                  ||||||||||||||
Sbjct  124544931  CTGGTGTTCTCTCC  124544918
```

Range 68: 127397865 to 127397878

| Score | Expect | Identities | Gaps | Strand | Frame |
|---|---|---|---|---|---|
| 28.2 bits(14) | 256() | 14/14(100%) | 0/14(0%) | Plus/Minus | |

Features:
**multiple epidermal growth factor-like domains protein 10 ...multiple epidermal growth factor-like domains protein 10 ...**

```
Query  281        TGGTGTTCTCTCCT  294
                  ||||||||||||||
Sbjct  127397878  TGGTGTTCTCTCCT  127397865
```

Range 69: 130857878 to 130857891

| Score | Expect | Identities | Gaps | Strand | Frame |
|---|---|---|---|---|---|
| 28.2 bits(14) | 256() | 14/14(100%) | 0/14(0%) | Plus/Minus | |

Features:
**672087 bp at 5' side: chondroitin sulfate synthase 3 isoform X2301556 bp at 3' side: histidine triad nucleotide-binding protein 1**

```
Query  283        GTGTTCTCTCCTTT  296
                  ||||||||||||||
Sbjct  130857891  GTGTTCTCTCCTTT  130857878
```

Range 70: 137930976 to 137930989

| Score | Expect | Identities | Gaps | Strand | Frame |
|---|---|---|---|---|---|
| 28.2 bits(14) | 256() | 14/14(100%) | 0/14(0%) | Plus/Minus | |

Features:
**polycystic kidney disease 2-like 2 protein isoform X5polycystic kidney disease 2-like 2 protein isoform X3**

```
Query  286        TTCTCTCCTTTATT  299
                  ||||||||||||||
Sbjct  137930989  TTCTCTCCTTTATT  137930976
```

Range 71: 142694321 to 142694338

| Score | Expect | Identities | Gaps | Strand | Frame |
|---|---|---|---|---|---|
| 28.2 bits(14) | 256() | 17/18(94%) | 0/18(0%) | Plus/Minus | |

Features:
**80194 bp at 5' side: fibroblast growth factor 1 isoform 2 precursor76424 bp at 3' side: rho GTPase-activating protein 26 isoform a**

```
Query  278        AGCTGGTGTTCTCTCCTT  295
                  ||||||||||||| ||||
Sbjct  142694338  AGCTGGTGTTCTCACCTT  142694321
```

Range 72: 158349419 to 158349432

| Score | Expect | Identities | Gaps | Strand | Frame |
|---|---|---|---|---|---|
| 28.2 bits(14) | 256() | 14/14(100%) | 0/14(0%) | Plus/Minus | |

Features:
**490449 bp at 5' side: clathrin interactor 1 isoform X3349679 bp at 3' side: transcription factor COE1 isoform 2**

```
Query  278        AGCTGGTGTTCTCT  291
                  ||||||||||||||
Sbjct  158349432  AGCTGGTGTTCTCT  158349419
```

Range 73: 158914139 to 158914152

| Score | Expect | Identities | Gaps | Strand | Frame |
|---|---|---|---|---|---|
| 28.2 bits(14) | 256() | 14/14(100%) | 0/14(0%) | Plus/Minus | |

Features:
**transcription factor COE1 isoform 2transcription factor COE1 isoform 1**

```
Query  273        TAACAAGCTGGTGT  286
```

```
                          ||||||||||||||
Sbjct  158914152  TAACAAGCTGGTGT  158914139
```

Range 74: 171479019 to 171479032

| Score | Expect | Identities | Gaps | Strand | Frame |
|---|---|---|---|---|---|
| 28.2 bits(14) | 256() | 14/14(100%) | 0/14(0%) | Plus/Minus | |

Features:
**22214 bp at 5' side: fibroblast growth factor 18 precursor294924 bp at 3' side: small integral membrane protein 23 isoform X2**

```
Query  287        TCTCTCCTTTATTG  300
                  ||||||||||||||
Sbjct  171479032  TCTCTCCTTTATTG  171479019
```

Range 75: 172063689 to 172063706

| Score | Expect | Identities | Gaps | Strand | Frame |
|---|---|---|---|---|---|
| 28.2 bits(14) | 256() | 17/18(94%) | 0/18(0%) | Plus/Minus | |

Features:
**serine/threonine-protein kinase 10 isoform X4serine/threonine-protein kinase 10 isoform X1**

```
Query  276        CAAGCTGGTGTTCTCTCC  293
                  |||||| |||||||||||
Sbjct  172063706  CAAGCTTGTGTTCTCTCC  172063689
```
